# Clonal Evolution of Acute Myeloid Leukemia Revealed by High-Throughput Single-Cell Genomics

**DOI:** 10.1101/2020.02.07.925743

**Authors:** Kiyomi Morita, Feng Wang, Katharina Jahn, Jack Kuipers, Yuanqing Yan, Jairo Matthews, Latasha Little, Curtis Gumbs, Shujuan Chen, Jianhua Zhang, Xingzhi Song, Erika Thompson, Keyur Patel, Carlos Bueso-Ramos, Courtney D DiNardo, Farhad Ravandi, Elias Jabbour, Michael Andreeff, Jorge Cortes, Marina Konopleva, Kapil Bhalla, Guillermo Garcia-Manero, Hagop Kantarjian, Niko Beerenwinkel, Nicholas Navin, P Andrew Futreal, Koichi Takahashi

**Affiliations:** Department of Leukemia, The University of Texas MD Anderson Cancer Center, Houston, Texas, USA; Department of Genomic Medicine, The University of Texas MD Anderson Cancer Center, Houston, Texas, USA; Department of Genetics, The University of Texas MD Anderson Cancer Center, Houston, Texas, USA; Department of Hematopathology, The University of Texas MD Anderson Cancer Center, Houston, Texas, USA; Department of Bioinformatics, The University of Texas MD Anderson Cancer Center, Houston, Texas, USA; Department of Biosystems Science and Engineering, Swiss Federal Institute of Technology in Zurich, Zurich, Switzerland; Department of Neurosurgery, The University of Texas Health Science Center at Houston, Houston, Texas, USA; Department of Hematology and Oncology, Graduate School of Medicine, The University of Tokyo, Tokyo, Japan

## Abstract

One of the pervasive features of cancer is the diversity of mutations found in malignant cells within the same tumor; a phenomenon called clonal diversity or intratumor heterogeneity. Clonal diversity allows tumors to adapt to the selective pressure of treatment and likely contributes to the development of treatment resistance and cancer recurrence. Thus, the ability to precisely delineate the clonal substructure of a tumor, including the evolutionary history of its development and the co-occurrence of its mutations, is necessary to understand and overcome treatment resistance. However, DNA sequencing of bulk tumor samples cannot accurately resolve complex clonal architectures. Here, we performed high-throughput single-cell DNA sequencing to quantitatively assess the clonal architecture of acute myeloid leukemia (AML). We sequenced a total of 556,951 cells from 77 patients with AML for 19 genes known to be recurrently mutated in AML. The data revealed clonal relationship among AML driver mutations and identified mutations that often co-occurred (e.g., *NPM1*/*FLT3-*ITD*, DNMT3A/NPM1, SRSF2*/*IDH2,* and *WT1/FLT3-*ITD) and those that were mutually exclusive (e.g., *NRAS*/*KRAS, FLT3-*D835/ITD, and *IDH1*/*IDH2*) at single-cell resolution. Reconstruction of the tumor phylogeny uncovered history of tumor development that is characterized by linear and branching clonal evolution patterns with latter involving functional convergence of separately evolved clones. Analysis of longitudinal samples revealed remodeling of clonal architecture in response to therapeutic pressure that is driven by clonal selection. Furthermore, in this AML cohort, higher clonal diversity (≥4 subclones) was associated with significantly worse overall survival. These data portray clonal relationship, architecture, and evolution of AML driver genes with unprecedented resolution, and illuminate the role of clonal diversity in therapeutic resistance, relapse and clinical outcome in AML.

## Main

A growing body of evidence supports the role of clonal diversity in therapeutic resistance, recurrence, and poor outcomes in cancer ^1^. Clonal diversity also reflects the history of the accumulation of somatic mutations within a tumor. Thus, a precise characterization of clonal diversity reveals not only the extent of a tumor’s clonal complexity but also the evolutionary history of the tumor’s development. Much of the work characterizing the clonal architecture of tumors has been done by computational inference using variant allele fraction (VAF) data from massively parallel DNA sequencing of bulk tumor samples ^2,3^. However, the ability to infer clonal heterogeneity and tumor phylogeny from bulk sequencing data is inherently limited, because bulk sequencing techniques cannot reliably infer mutation co-occurrences and hence often fail in reconstructing clonal substructure.

Single-cell DNA sequencing (scDNA-seq) can address some of these challenges ^4–8^. However, until recently, the available methods required laborious single-cell isolation protocols and suffered from low cell throughput, limited gene coverage, and technical artifacts from whole-genome amplification that hindered their ability to characterize clonal architecture with precision ^9^. Recent technological advances now allow rapid single-cell genotyping of targeted cancer-related genes in thousands of cells. We previously described the performance and feasibility of a new scDNA-seq platform (Tapestri®, Mission Bio, Inc.) in primary samples from 2 patients with acute myeloid leukemia (AML) ^10^. Here, using this method, we conducted scDNA-seq in 91 AML samples from 77 patients and uncovered the landscape of AML clonal architecture at single-cell resolution. Using the data, we reconstructed the mutational history of driver genes, some of which are therapeutic targets, and identified both linear and branching clonal evolution patterns in AML. Additionally, we studied dynamic changes of clonal architecture in response to therapies and analyzed the clinical implications of clonal diversity in AML.

### The landscape of driver mutations in AML at single-cell resolution

We analyzed bone marrow mononuclear cells (BMNC) from 77 AML patients, of which 64 (83%) were previously untreated, and 13 (17%) had relapsed or refractory disease (detailed clinical characteristics are summarized in Supplementary Table 1). The cohort was enriched with samples with normal diploid karyotype (*N* = 68, 88%) to avoid allelic imbalance affecting the genotype calling. The median bone marrow blast percentage was 44% (interquartile range [IQR]: 29%-67%). A median of 7,584 BMNC (IQR: 6,194-8,361) per sample were sequenced by the scDNA-seq platform (Fig. 1a). Across 40 amplicons targeting 19 known AML driver genes, scDNA-seq resulted in a median of 25× coverage per amplicon per cell (IQR: 12×-43×, Extended Data Fig. 1). The amplicons covering guanine-cytosine–rich sequences, such as *GATA2, SRSF2*, and parts of *RUNX1* and *TP53*, had lower coverage than others, such that relatively large numbers of cells had inconclusive genotype information for the mutations covered by these amplicons (Extended Data Fig. 2). The estimated median allele dropout (ADO) rate was 4.7% (IQR: 3.6%-5.7%) (Extended Data Fig. 3). The estimated lower limit of detection (LOD) of the platform was 0.1% of the cellular population based on the serial dilution assay of a cell line and also from mutation validation by droplet digital PCR (Supplementary Table 2 and Extended Data Fig. 4).

**Fig. 1.**
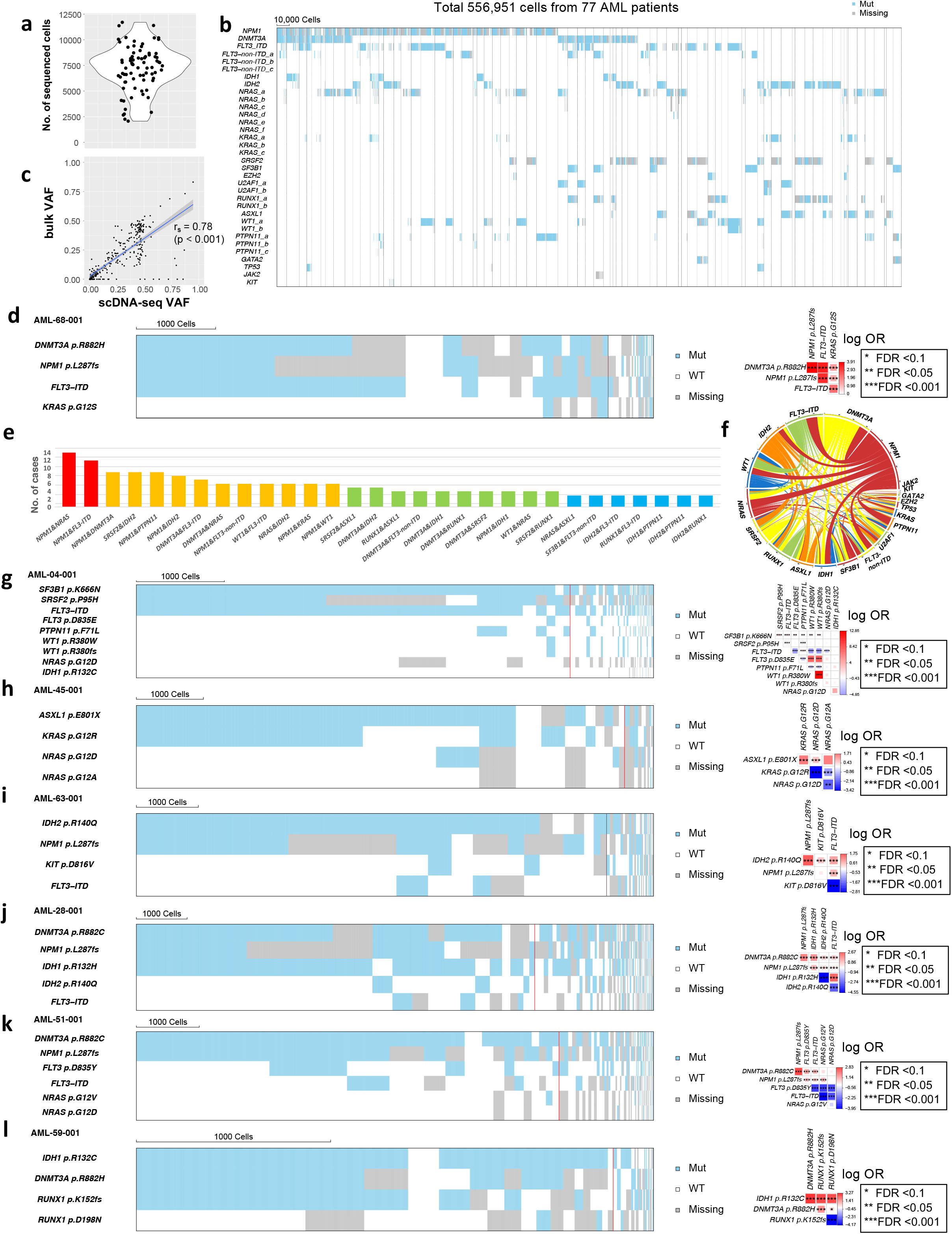
The Genetic landscape of AML based on single-cell DNA sequencing. **a,** Distribution of the number of total sequenced cells. Each point represents a sample from unique patients. **b**, Somatic mutations in 556,951 cells from 77 AML patients detected by single-cell DNA sequencing. Each column represents a cell, and cells from the same case are clustered together within the areas surrounded by the grey lines. Note that some cases are difficult to be segregated in print. Cells that were genotyped as being mutated or wild type for the indicated gene are colored in blue and white, respectively. Cells with missing genotypes are colored in grey. When one sample has multiple different mutations in the same gene, they were annotated differently (e.g., *NRAS*_a, *NRAS*_b). A total of 57,953 cells that were genotyped as wild type for all the variants screened are not shown. **c**, Correlation of the variant allele fraction (VAF) from bulk-sequencing and single cell DNA sequencing. The X-axis shows the VAF from the single-cell genotype data (scDNA-seq VAF). The Y-axis shows the VAF from the bulk next-generation sequencing (bulk VAF). Each dot represents a detected variant. The linear trendline was added to best fit the distribution of the dots. The shaded area around the trendline represents the 95% confidence intervals. **d**, Cellular-level co-occurrence of *DNMT3A*, *NPM1*, and *FLT3*-ITD mutations. Heat map (left) shows the genotype of each sequenced cell for each variant, with clustering based on the genotypes of driver mutations. Each column represents a cell at the indicated scale. Cells with mutations and wild-type cells are indicated in blue and white, respectively. Cells with missing genotypes are indicated in grey. The subclones located to the right of the red line comprised <1% of the total sequence cells, since such small subclones can represent false positive or negative genotypes as a result of ADO or multiplets. The figure on the right show the pairwise association of mutations. The color and size of each panel represent the degree of the logarithmic odds ratio (log OR). The bar on the right side is a key indicating the association of the colors with the log OR. Co-occurrence and mutual exclusivity are indicated by red and blue, respectively. The statistical significance of the associations based on the false discovery rate (FDR) is indicated by the asterisks (*FDR < 0.1, **FDR < 0.05, ***FDR < 0.001). **e**, Frequency of mutation combination showing statistically-significant cell-level co-occurrence (FDR < 0.001). x-axis represents the combination of mutations based on mutated genes, and y-axis shows the number of patients showing the significant cell-level co-occurrence of each mutation combination. Mutation combinations that were detected in 3 or more patients are plotted. Bars are colored based on the frequency (red if significantly co-occurred in >10 patients, orange if 6-10 patients, green if 4-5 patients, blue if 3 patients). **f,** Circos plot showing the patterns of mutation co-occurrence for all genes based on the single-cell genotype data. When 2 mutations co-occurred in the same cell, a ribbon connects the genes. The width of each ribbon is proportional to the frequency of mutational events. **g,** Cellular-level co-occurrence of *SF3B1* and *SRSF2* mutations. **h-l**, Cell-level mutual exclusivity patterns of somatic mutations in individual samples for 5 representative cases. (**h**) *KRAS* and *NRAS*, (**i**) *KIT* and *FLT3*-ITD, (**j**) *IDH1* and *IDH2*, (**k**) *FLT3-*non-ITD, *FLT3-*ITD, and *NRAS,* and (**l**) *RUNX1* p.K152fs and *RUNX1* p.D198N variants did not co-occur in the same cellular populations. Mut, mutated; WT, wild type; Missing, missing genotype.

In total, we sequenced 556,951 BMNC from 77 AML patient samples (Fig. 1b). The scDNA-seq approach detected 331 somatic mutations in 19 cancer genes, which included 238 (72%) single-nucleotide variants (SNV) and 93 (28%) small indels. Among those, 314 mutations (95%) were orthogonally validated: 274 (87%) by conventional bulk next-generation sequencing ^11^ (bulk-seq, median 407×), 29 (9%) by droplet digital PCR, and 11 (3%) by a quantitative PCR assay (all *FLT3*-internal tandem duplication [ITD], 4%). Therefore, the subsequent analyses used a final set of 314 validated mutations (Supplementary Table 3). Of note, among the shared genomic regions covered by the scDNA-seq and the bulk-seq platforms, all 274 (100%) mutations called by the bulk-seq were also detected by scDNA-seq. The VAF from bulk-seq (bulk VAF) and the VAF inferred from the scDNA-seq data (scDNA-seq VAF) had a good concordance (r_s_ = 0.78, p < 0.001) suggesting that the sequenced cells are a good representation of the total bulk samples. (Fig. 1c and Extended Data Fig. 5).

The most frequently detected mutations by scDNA-seq in the 77 patients were in *FLT3* (N = 37, 48%; 30 [39%] with ITD and 16 [21%] with non-ITD mutations), followed by *NRAS* (N = 35, 45%), *NPM1* (*N* = 32, 42%), *IDH2* (*N* = 23, 30%), *DNMT3A* (*N* = 20, 26%), *SRSF2* (*N* = 17, 22%), *RUNX1* (*N* = 14, 18%), *KRAS* (*N* = 14, 18%), *PTPN11* (*N* = 14, 18%), and *WT1* (N=14, 18%). scDNA-seq detected substantially more *FLT3* mutations (11 [79%] ITD and 3 [21%] non-ITD) than bulk-seq (Extended Data Fig. 6a). This is likely due to the capability of the scDNA-seq platform in detecting cryptic *FLT3* mutations in small cellular subpopulations (Extended Data Fig. 6b), which has been also reported previously using a different single-cell technology ^12^.

### Cellular-level co-occurrence and mutual exclusivity of AML driver mutations

Analysis of the cell-level co-occurrence and mutual exclusivity of AML driver mutations suggested cooperative mechanisms and functional redundancy among these mutations. For instance, sample number AML-68-001 carried *DNMT3A*, *NPM1*, and *FLT3*-ITD mutations, which are the most frequently co-occurring mutations in AML at the patient level ^13^. scDNA-seq unambiguously identified a cellular population carrying all 3 of these mutations (Fig. 1d). Analysis of statistically significant mutation co-occurrence (false discovery rate [FDR] < 0.001) identified frequently co-occurring mutations at the cell-level, which included *NPM1*/*NRAS*, *NPM1*/*FLT3*, *NPM1*/*DNMT3A*, *SRSF2/IDH2, NPM1*/*PTPN11*, *NPM1*/*IDH2, DNMT3A*/*FLT3*-ITD, *DNMT3A/NRAS, WT1*/*FLT3*-ITD, *NRAS/IDH2, NPM1*/*KRAS*, *NPM1/WT1,* and others (Fig. 1e-f, Extended Data Fig. 7a and 8, variant-level co-occurrence is summarized in Extended Data Fig. 7b). These cell-level co-occurrence data confirm the known cooperative relationship between the driver mutations that has been suggested by previous studies with bulk-seq^13,14^ or functional studies^15–21^, but also generates new hypotheses for the role of rare combinations in cellular oncogenesis. For instance, we detected the cell-level co-occurrence of *SF3B1* p.K666N and *SRSF2* p.P95H mutations in AML-04-001 (Fig. 1g). Mutations in RNA splicing genes are mutually exclusive in general and thought to have functional redundancy or synthetically lethal relationship ^22^; however, in rare instances, patients having both *SF3B1* and *SRSF2* mutations were reported ^23^. Our data confirm that the two mutations can co-occur at the cellular level, and suggests potential cooperativity between the two RNA splicing mutations.

In contrast, sample AML-45-001 carried *ASXL1*, *KRAS*, and 2 *NRA*S mutations. The *KRAS* and 2 *NRAS* mutations were mutually exclusive at the cellular level (Fig. 1h; FDR < 0.001). Mutually exclusive relationships were frequently identified among mutations that are in the same gene or part of the same molecular pathway (e.g., *KIT* p.D816V and *FLT3*-ITD; *IDH1* and *IDH2*; *FLT3*-p.D835Y and *FLT3*-ITD; *RUNX1* p.K152fs and *RUNX1* p.D198N) (Fig. 1i-l). These results support the widely explored hypothesis that co-occurrence of two or more functionally redundant oncogenic mutations does not provide a selective advantage to cancer cells and could be potentially synthetically lethal ^24,25^. Taken together, these data provide definitive evidence of mutation co-occurrence at cellular level and provide a landscape of clonal relationship among AML driver mutations (Extended Data Fig. 8 and 9).

### Zygosity of AML driver mutations

One of the unique aspects of scDNA-seq is its capability of calling mutations in individual cells with zygosity information. In fact, a previous single-cell study reported the cellular diversity in the zygosity of *NPM1* and *FLT3* mutations in AML^4^. However, the lack of validation method has made the interpretation of zygostiy difficult. In the current cohort, mutations in genes such as *FLT3*-ITD, *GATA2, JAK2, NPM1, RUNX1*, and *SRSF2* were frequently detected as homozygous (Extended Data Fig. 10). Because amplicons covering some of these mutations (*GATA2, RUNX1*, and *SRSF2*) had relatively low sequencing depth (Extended Data Fig. 1), it is possible that some homozygous calls were the result of low sequencing depth and ADO. To validate zygosity called by the scDNA-seq, we performed SNP arrays in selected samples with homozygous mutation calls. In AML-25-001, 97% of the cells had a homozygous *RUNX1* p.Q335X mutation determined by scDNA-seq data, and SNP array data detected a copy-neutral loss of heterozygosity (CN-LOH) on chromosome 21 involving *RUNX1* (Fig. 2a). Similarly, in AML-03-001, 66% of the cells had a homozygous *FLT3*-ITD mutation determined by scDNA-seq data, and the SNP array detected CN-LOH on chromosome 13 involving *FLT3* (Fig. 2b). These results indicate that the observation of homozygosity of the *RUNX1* and *FLT3-* ITD mutations in these cases was true and was a result of CN-LOH. In contrast, none of the samples with homozygous *SRSF2* (17% of the cells genotyped as homozygous in AML-57-001, Fig. 2c) or *NPM1* (13% of the cells genotyped as homozygous in AML-13-001, Fig. 2d) mutations had allelic imbalance involving the mutated loci. These results do not rule out the possibility that the SNP arrays missed the subclonal allelic imbalance. However, the cells that were genotyped as homozygous had significantly lower sequencing depth than did the cells that were genotyped as heterozygous (Fig. 2c-d and Extended Data Fig. 11), suggesting that the homozygous calls in these mutations may have resulted from low sequencing depth and ADO. For cases with validated homozygous calls, the zygosity of the mutations added further resolution to the interpretation of the clonal substructure of the samples (Fig. 2a-b and Extended Data Fig. 11).

**Fig. 2.**
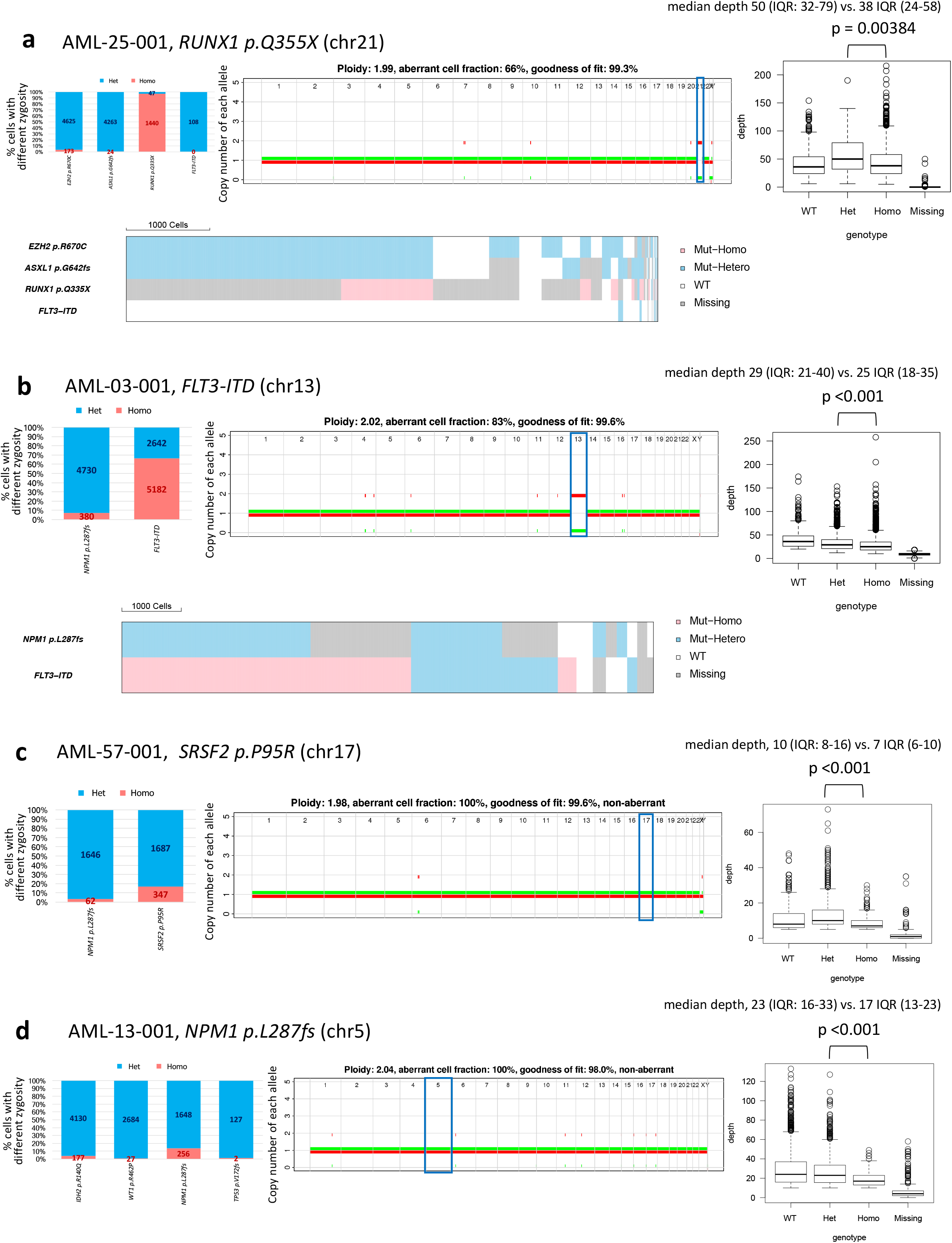
Homozygous variants involving copy-neutral loss of heterozygosity. **a-d,** Representative cases with highly homozygous variants analyzed by SNP array. The bar graphs on the left show the distribution of zygosity for each indicated variant. Cells that were genotyped as having heterozygous and homozygous mutations are shown in blue and red, respectively. The numbers on the bars represent the number of cells with each genotype. The figures in the middle show the distribution of the allele counts for the two alleles (green or red). The allele count is shown on the vertical axes, and the chromosomes are shown on the horizontal axes. The chromosomes on which the highly homozygous variants were located are highlighted with blue rectangles. Distributions of depth are shown in the figures on the right based on the genotype calling. Heat maps incorporating the zygosity information are also shown for cases with validated homozygosity. Copy-neutral loss of heterozygosity (CN-LOH) involving the homozygously called variant loci was detected by SNP array in cases with highly homozygous **(a)** *RUNX1* p.Q355X and **(b)** *FLT3*-ITD variants. Cases with homozygously called **(c)** *SRSF2* p.P95R and **(d)** *NPM1* p.L287fs variants did not have CN-LOH or any other copy-number alterations. WT, wild type; Het, heterozygous; Homo, homozygous; Missing, missing genotype; IQR, interquartile range.

### Reconstructing mutational histories in AML

To reconstruct mutational histories in AML, we used SCITE, a probabilistic model to infer phylogenetic trees from single-cell sequencing data that involves a flexible Markov-chain Monte Carlo (MCMC) algorithm ^26^ (Supplemental Methods). Reconstructed phylogenies revealed both linear and branching evolution patterns in AML (Extended Data Fig. 12). Patients showing branching evolution had a significantly higher number of mutations, compared with those with linear evolution (median number of mutations 5 [IQR: 4-6] vs. 3 [IQR: 2-4], p < 0.001, Fig. 3a). In cases with linear evolution pattern, ancestral mutations often involved *DNMT3A, IDH2,* and *SRSF2* mutations that have been detected as preleukemic clonal hematopoiesis ^27,28^ followed by sequential accrual of secondary mutations that frequently involved *NPM1, FLT3, RUNX1, NRAS*, *KRAS*, and *PTPN11.* (Fig. 3b-e and Extended Data Fig. 12) ^13,29^. In some cases with branching pattern, we observed the evolutionary history that is consistent with convergent evolution. For example, in AML-38-001, a putative founding mutation, *NPM1* p.L287fs, generated 2 independent branches with *IDH1* p.R132H or *IDH2* p.R140Q mutations. Each of these branches then separated into *PTPN11* p.D61H or *KRAS* p.G12A mutations and *FLT3*-ITD, *NRAS* p.G13R, *PTPN11* p.A72G, or *NRAS* p.G12A mutations, respectively. As a result, the sample carried 9 clones, each with a combination of similar, but separately evolved, molecular alterations (*NPM1-IDH-RAS/RTK* pathway alteration) and the same mutation order (Fig. 3f). Other cases exhibiting evidence of branching evolution are shown in Fig. 3g-i and Extended Data Fig. 12. These data with convergent evolution indicate the presence of selective pressure in a bone marrow ecosystem that favors AML clones having certain combination of molecular alterations with a fixed order.

**Fig. 3.**
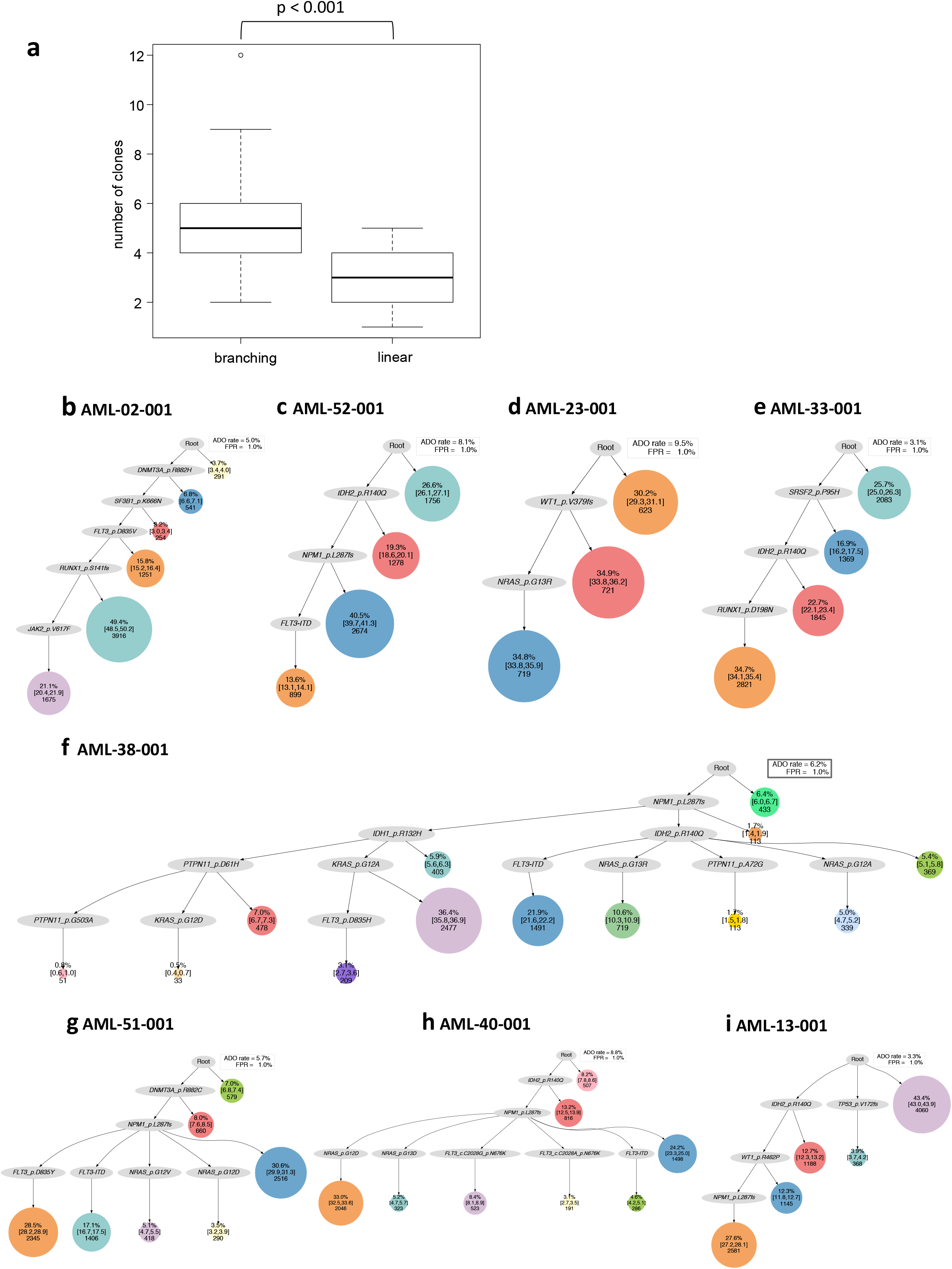
Inference of mutational history from single-cell genotype data using the SCITE algorithm. **a,** Distribution of the number of driver mutations based on the evolution patterns (branching or linear). The thick line within each box represents the median, and the top and bottom edges represent the 25th and 75th percentiles, respectively. The upper and lower whiskers represent the 75th percentile plus 1.5 times the interquartile range and the 25th percentile minus 1.5 times the interquartile range, respectively. **b-i,** Phylogenetic trees for representative cases illustrating distinct patterns of clonal evolution. The size of each circle is proportional to the clonal population. The numbers within each circle are the number of cells and the percentage of each clone among the total number of sequenced cells and the 95% credible intervals from the posterior sampling to illustrate the uncertainty in the subclone sizes. (**b)-(e)** show linear clonal evolution pattern, in which a subset of cells from the founder clone acquired additional mutations in a stepwise manner. The trunk clone exhibits a forked evolution pattern based on the status of additional mutations. (**f)-(i)** show branching clonal evolution pattern characterized by the parallel acquisition of multiple functionally redundant mutations in different cell populations. ADO, allele dropout; FPR, false positive rate.

### Clonal remodeling under therapeutic pressure

We then analyzed 24 longitudinal samples from 10 patients (8 with baseline and relapse pairs and 2 with multiple refractory timepoints) to study the evolution of clonal architecture in response to therapies (Fig. 4 and Extended Data Fig. 13). We observed clonal selection and adaptation under the therapeutic pressure that were associated with the patients’ clinical courses. For instance, AML-09 was a 74-year-old man with previously untreated AML with *NPM1* p.L287fs, *FLT3-*ITD, *FLT3* p.D835E, *FLT3* p.D835Y, and *KRAS* p.G13D mutations. The patient was treated with azacitidine and sorafenib and experienced morphological complete remission (i.e. leukemic blast less than 5% in marrow with normal recovery of blood counts). However, 5 months later, his AML relapsed. scDNA-seq of the baseline-relapse pair revealed that *NPM1*-*FLT3* p.D835Y clone, originally a subclone that constituted 1.7% of the diagnostic sample, was selected during the therapy, suggesting that clonal selection is the underlying mechanism of relapse in this case (Fig. 4a). This clonal selection is consistent with the known *in vitro* differential sensitivity of various *FLT3* mutations to sorafenib; indeed, the *FLT3* p.D835Y mutation was shown to be more resistant to sorafenib than were the D835E and ITD mutations ^30^.

**Fig. 4.**
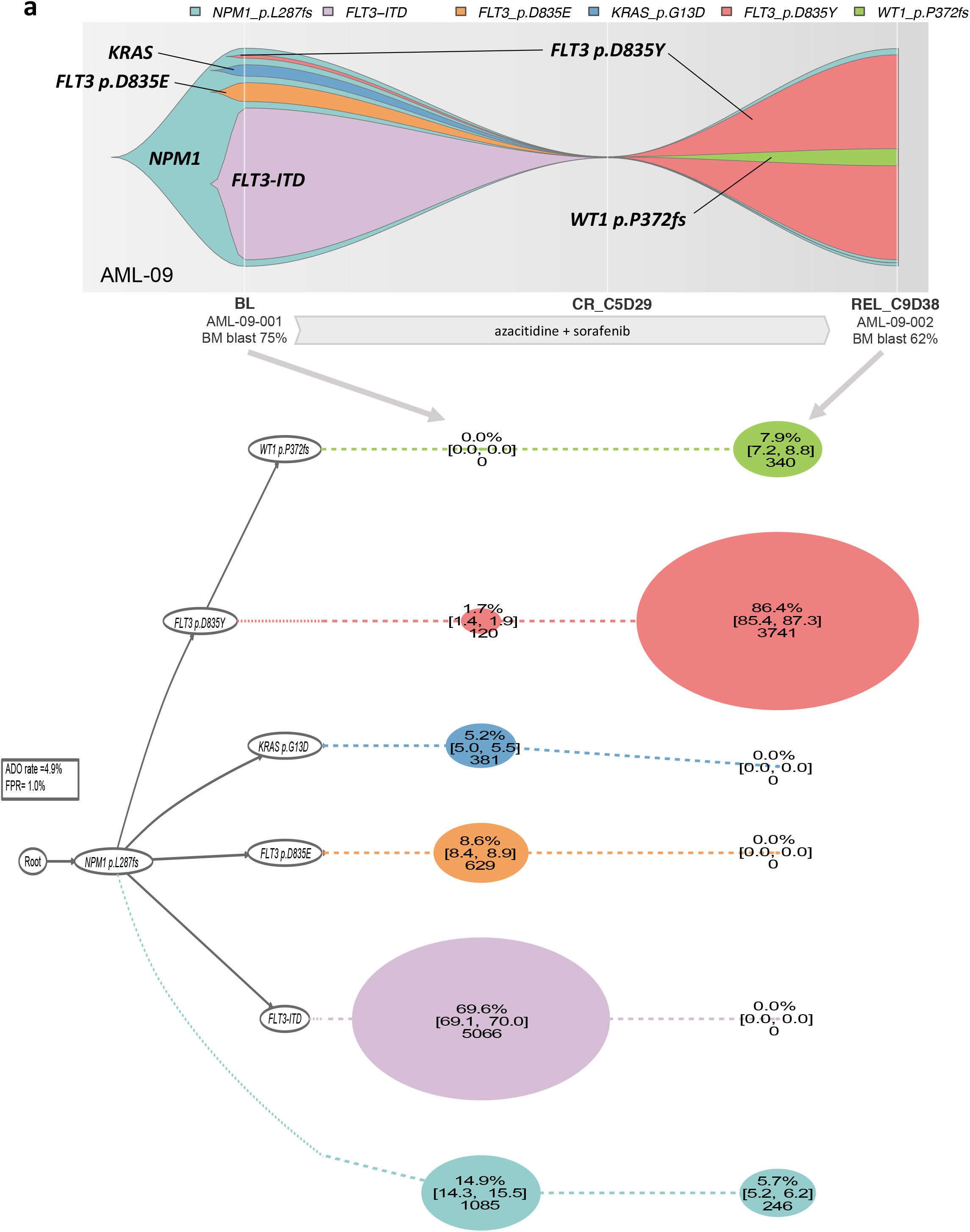
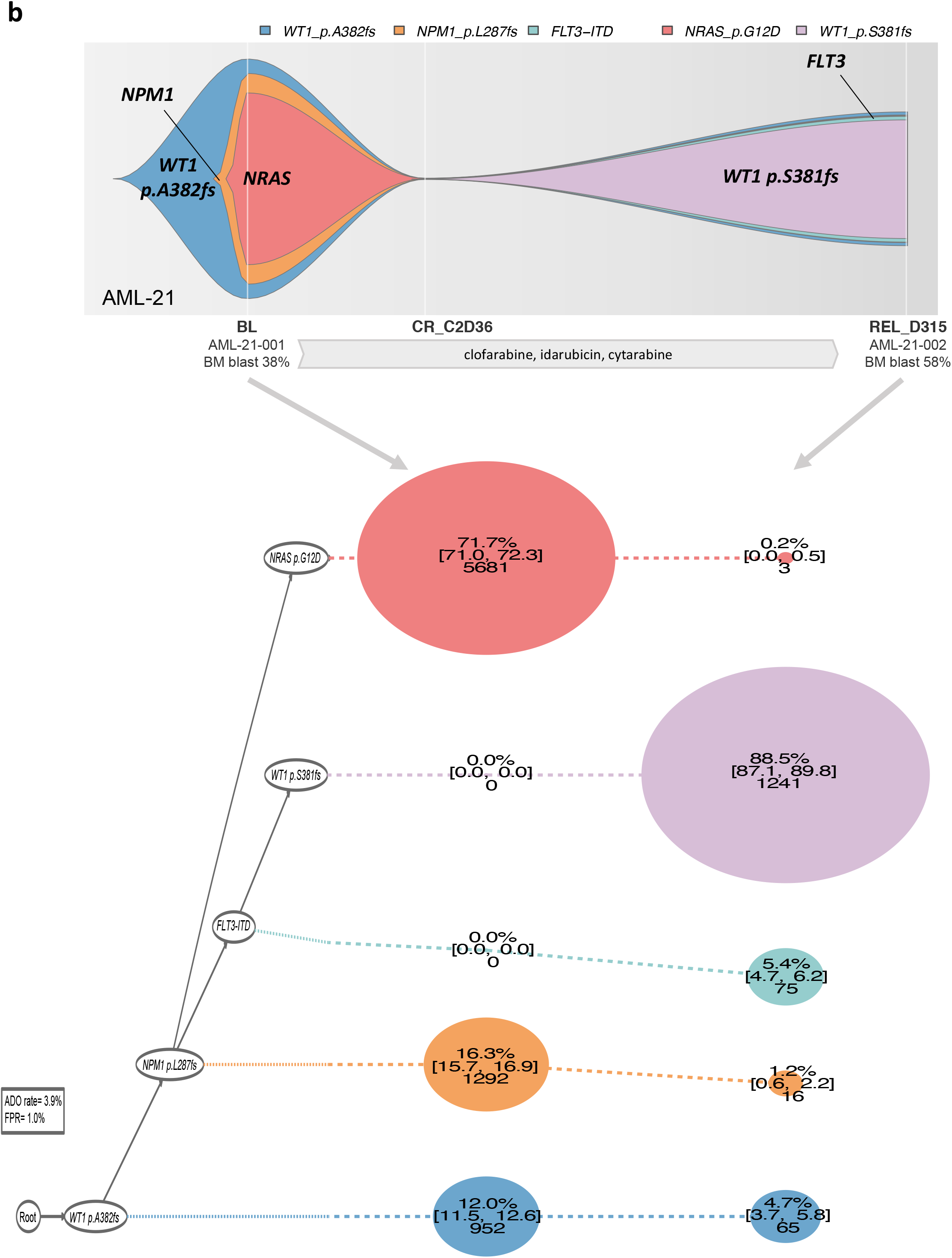
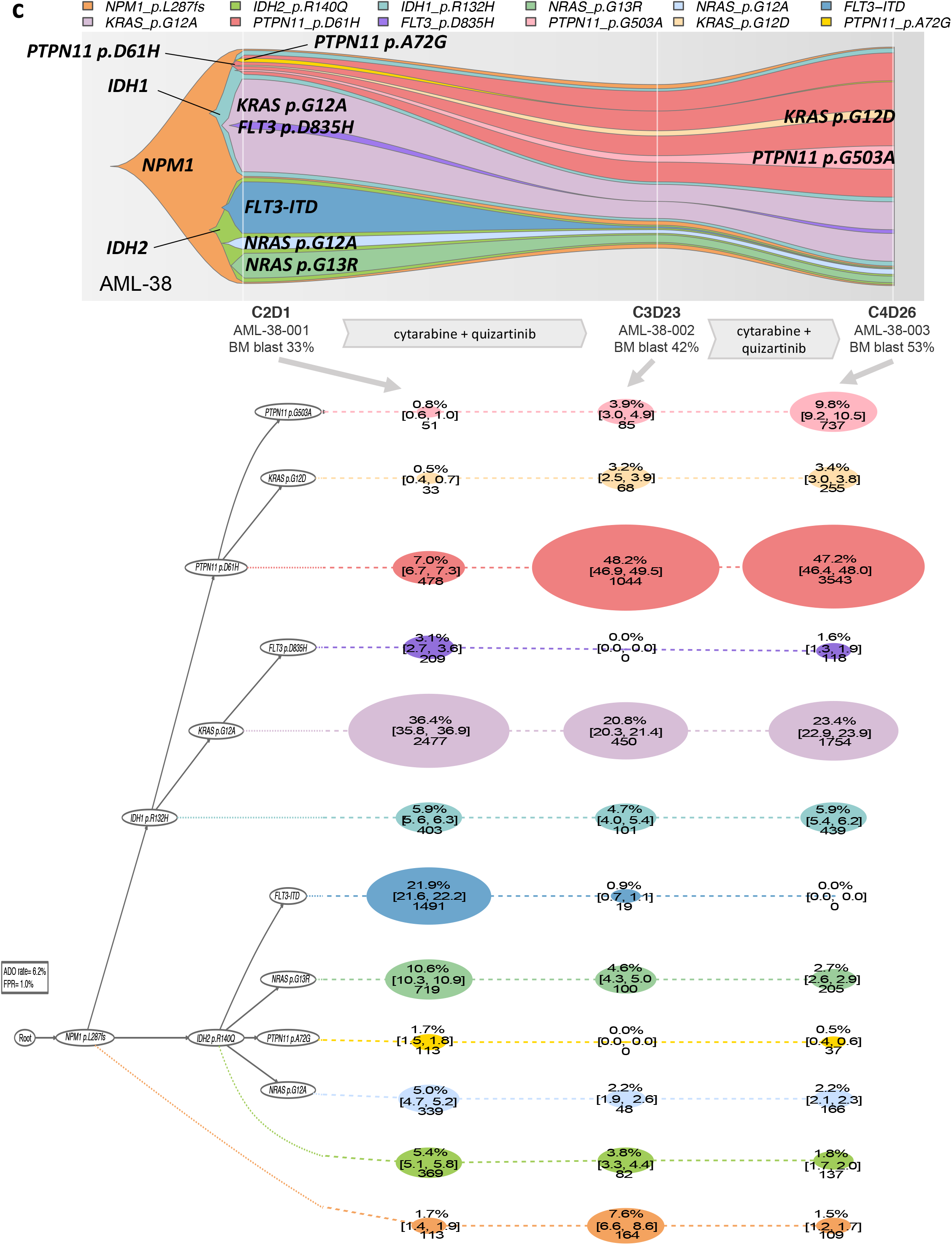
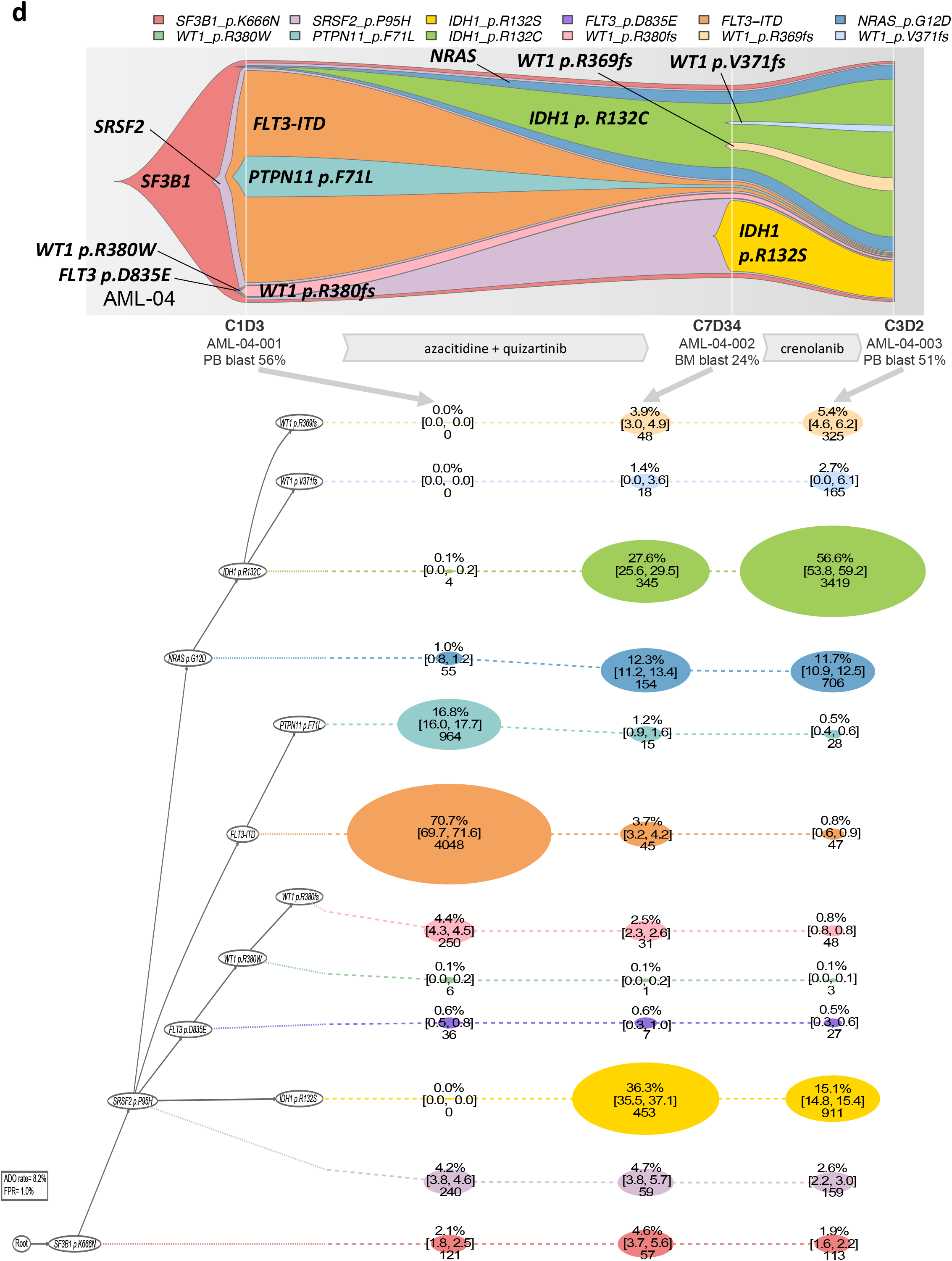
Various patterns of clonal evolution after therapy. The fish plot shows the inferred clonal evolution pattern based on the single-cell genotype data. The phylogenetic trees visualize the estimated order of mutation acquisition and the proportion of subclones with a different combination of mutations at each timepoint. **(a)** A 74-year-old man with newly diagnosed therapy-related acute myelomonocytic leukemia showing a clonal selection of *FLT3* p.D835Y-mutated clone, a small subclone at baseline. **(b)** A 56-year old woman with acute myelomonocytic leukemia showing the regression of *NRAS*-mutated clone replaced by *FLT3*-*WT1* p.S381fs-mutated clone. **(c)** A 58-year-old-man with refractory AML showing the dynamic change of subclonal architecture. The patient was started on cytarabine and quizartinib and initially responded, but had a recurrent disease, and was refractory to a total of 4 cycles of cytarabine and quizartinib therapy. *FLT3-*ITD mutated clone was cleared after the therapy, whereas the remaining subclones persisted or expanded. **(d)** A 76-year-old man with refractory secondary AML showing the adaptive behavior of subclones. Treatment with azacitidine plus quizartinib followed by crenolanib monotherapy substantially shrank *FLT3-*ITD–mutated clone, whereas the *NRAS-IDH1* p.R132C-mutated clone and *IDH1* p.R132S-mutated clone emerged in the context of *SF3B1* and *SRSF2* mutations and expanded. Full case description is available in the figure legends of Extended Data Fig. 13.

The second case AML-21 is a 56-year-old-woman with newly-diagnosed AML who was treated with induction chemotherapy with clofarabine, idarubicin, and cytarabine followed by 4 additional cycles of consolidation therapy. After approximately 7 months of remission, her disease relapsed. While both baseline and relapse samples shared ancestral *WT1* p.A382fs - *NPM1* mutations, the *NRAS* clone was replaced by the *FLT3*-ITD-*WT1* p.S381fs clone at relapse (the acquired *WT1* p.S381fs mutation was *in trans*, making biallelic *WT1* mutations at relapse, Extended Data Fig. 14). These *FLT3*-ITD and *WT1* p.S381fs mutations were undetectable at baseline. They were likely acquired *de novo* at relapse or sub-detectable at baseline and were selected during the therapy (Fig. 4b).

Finally, two treatment refractory AML cases showed highly adaptive clonal structure during therapy. Both AML-38 (Fig. 4c, the same case in Fig. 3f) and AML-04 (Fig. 4d) had AML with multiple branching clones. In both cases, treatment with a *FLT3* inhibitor-containing therapy decreased clones with *FLT3* mutations, however with a concurrent expansion or selection of other clones and development of new clones. High clonal diversity in both cases seems to have allowed these AMLs to flexibly re-configure their clonal composition during therapy, which likely contributed to the therapeutic resistance (Fig. 4c-d). These data from longitudinal cases illustrate the evolution of clonal architecture under the selective pressure of therapy and elucidate the role of clonal selection and adaptation in therapy resistance and relapse.

### Association between clonal diversity and clinical outcomes

We then analyzed the clinical implications of clonal diversity in AML that is uncovered by our single-cell sequencing. The median number of cell subclones per patient was 4 (IQR: 3-5). Using the median as a cut-off, we divided the patients into lower (<4 subclones) and higher clonal diversity (≥4 subclones) groups. Patients with higher clonal diversity were significantly older compared with those with lower clonal diversity (median age 63 vs. 56 years, p = 0.0283). Patients with higher clonal diversity were more likely to have a secondary or therapy-related disease, relapsed/refractory disease, and tended to harbor chromosomal abnormalities detected by karyotyping, although the differences were statistically not significant. Among the 64 previously-untreated cases, those with higher clonal diversity had a trend toward lower CR rate compared with those with lower clonal diversity (CR rate 78% vs. 97%, p = 0.0534, Fig. 5a). Also, AML patients with higher clonal diversity showed significantly worse overall survival (OS) compared with those with lower clonal diversity (2-year OS 25 vs. 59 months, p = 0.0469; Fig. 5b).

**Fig. 5.**
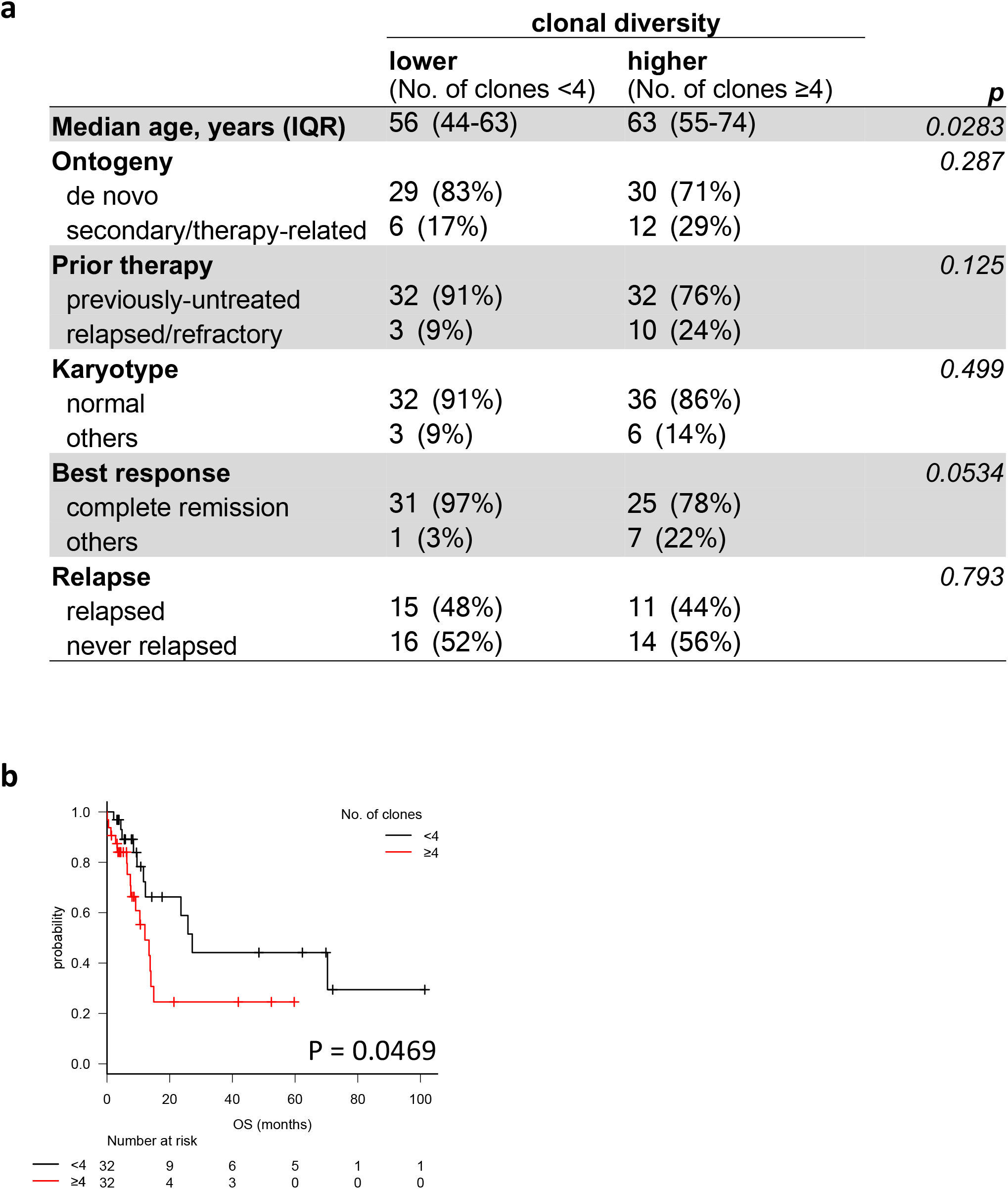
Clonal diversity and its association with clinical outcome. **a,** Association between clonal diversity and clinical characteristics. **b,** Overall survival (OS) for previously-untreated patients (N=64) according to clonal diversity based on single-cell DNA sequencing data.

## Discussion

Using a novel high-throughput scDNA-seq platform, we determined the clonal architecture of AML at single-cell resolution and described the clonal relationships among AML driver mutations. Cell-level co-occurrence and mutual exclusivity data obtained from this study provide the rationale for future studies investigating the cooperative mechanism, functional redundancy, and synthetic lethality among oncogenic mutations. Reconstruction of mutational history based on the single-cell data not only revealed inter-tumor diversity in the evolutionary history of AML but also provided evidence for linear and branching evolution patterns in AML with some cases exhibiting convergent evolution. In cases with convergent evolution, we observed clones that evolved separately but with a similar coalescence of molecular alterations, which is similar to the observations in other studies utilizing multi-region/site sequencing or single-cell analysis for different tumors^6,31–34^. These observations indicate for an underlying genetic instability and evolutionary adaptation of AML clones to selective pressure in tissue ecosystem. Cancer therapies, particularly molecularly targeted therapies, provide additional selective pressure to AML clones, which facilitates selection of resistant clones and acquisition of new mutations or clones, leading to recurrence or treatment resistance.

This work represents the largest cohort of AML patients yet examined at single-cell resolution, moves a growing body of data^4,6^ forward into a deeper understanding of the fundamental clonal architectures of AML. The depth of both patient numbers and cells sequenced allowed a robust analysis of the clonal relationship and phylogeny in this study despite the technical challenges associated with single-cell sequencing, such as but not limited to, ADO, multiplets, coverage inconsistency, and false positives. Moreover, a large sample size allowed the description of inter-tumor diversity in the patterns of clonal evolution, and offered some indication that clonal diversity affects prognosis in AML. Here, we interrogated 19 known leukemia driver genes that have given rise to a remarkable level of clonal complexity in AML. It is noteworthy that this is still an underestimation of the true extent of clonal diversity. Future studies with even more cells, broader coverage of the genome, and integration with single-cell transcriptomic and epigenomic states, that is becoming a reality with the recent technological advancement ^35 36^, will further elucidate the clonal diversity and evolutionary trajectories of AML, which may contribute to the development of predictive biomarkers or therapies targeting clonal diversity.

## METHODS

### Patients and samples

We included in the analysis 91 samples (89 bone marrow mononuclear cells and 2 peripheral blood mononuclear cells) from 75 patients with AML and 2 patients with high-risk myelodysplastic syndrome who had at least one somatic mutation covered by the targeted panel for scDNA-seq. In order to avoid allelic imbalance, samples with a normal karyotype were prioritized for analysis. For samples that exhibited cytogenetic abnormalities, we confirmed that the chromosomal position of the examined somatic mutations did not overlap with the regions with a structural variation. Of the 77 patients, 67 patients were analyzed for the single-timepoint sample including pre-treatment (N=59), relapse (N=5), and random timepoint with refractory disease (N=3). For the remaining 10 patients, we analyzed the longitudinal samples obtained at pre-treatment and relapse (N=6), pre-treatment, during treatment, and relapse (N=2), and 3 random refractory timepoints (N=2). All the patients provided written informed consent for sample banking and analysis. The study was approved by the MD Anderson institutional review board and was in accordance with the Declaration of Helsinki.

### Variant detection by single-cell DNA sequencing

We used a novel microfluidic approach with molecular barcode technology to amplify the DNA from individual cells as previously described ^10^. Briefly, cryopreserved bone marrow mononuclear cells were thawed, and cells were quantified using a Countess Automated Cell Counter (Thermo Fisher). The cells were resuspended in cell buffer and diluted to a concentration of 2,000,000 to 4,000,000 cells/mL. Next, 100 μL of cell suspension was loaded onto a microfluidics cartridge and cells were encapsulated on the Tapestri instrument followed by the cell lysis and protease digestion on a thermal cycler within the individual droplet. The cell lysate was then barcoded such that each cell had a unique label. The barcoded samples were then thermocycled using 50 primer pairs specific to a panel of 19 mutated genes covering known AML-related hotspot loci and 10 commonly heterozygous SNP loci for ADO determination (Supplementary Table 4).

The pooled library was sequenced on an Illumina MiSeq system with 150- or 250-base pair (bp) paired-end multiplexed runs. Detailed methods are provided in the Supplemental Methods. Briefly, fastq files generated by the MiSeq instrument were processed using the Tapestri Analysis Pipeline for adapter trimming, sequence alignment, barcode correction, cell finding, and variant calling. Loom files that were generated by the pipeline via GATK-based haplotype calling were then processed using in-house filtering criteria. We included cells for downstream analysis that met the following criteria for genotyping: total read count (depth, DP) ≥ 10× and alternative allele count ≥ 3 (scVAF ≥ 15% if 20× ≤ DP ≤ 99×; scVAF ≥ 10% if DP ≥ 100×). Cells that did not satisfy these criteria were considered to have missing genotypes.

The ADO rate was calculated on the basis of common SNP information using 10 amplicons designed to cover 10 highly polymorphic loci in the Tapestri Single-Cell DNA AML Panel.

### Mutation detection by bulk sequencing

As an orthogonal validation, all samples were concurrently sequenced by conventional bulk next-generation sequencing (NGS) using target-capture deep sequencing (*N* = 66, median coverage: 432×, IQR: 283×-610×) or whole-exome sequencing (*N* = 11, median coverage: 146×, IQR: 86×-158×). Target-capture NGS was performed using a SureSelect (Agilent Technologies) custom panel of 295 genes that are recurrently mutated in hematological malignancies (Supplementary Table 5). Detailed methods were previously described ^11^. Briefly, genomic DNA was extracted using an Autopure extractor (QIAGEN/Gentra) and was fragmented and bait-captured in solution according to the manufacturer’s protocols. Captured DNA libraries were then sequenced using a HiSeq 2000 sequencer (Illumina) with 76-bp paired-end reads. Whole-exome sequencing was performed using SureSelect V4 exome probes (Agilent Technologies) and a HiSeq 2000 sequencer (Illumina) with 76-bp paired-end reads. Modified Mutect and Pindel algorithms were used for mutation calling as described previously ^11^.

### Comparison of genotype results from scDNA-seq and bulk sequencing

To determine how the models of clonal architecture obtained using the 2 sequencing methods differed, we compared the VAF from bulk sequencing (bulk VAF) and the VAF from single-cell genotype data (scDNA-seq VAF). scDNA-seq VAF was calculated as follows based on the sequencing reads from the pooled single cells: (number of the single-cell sequencing reads with alternate allele) / (number of total single-cell sequencing reads).

### Inference of mutational histories

We used the SCITE (Single Cell Inference of Tumor Evolution) software to infer phylogenetic trees of the driver mutations from scDNA-seq data as previously described ^26^. SCITE is an MCMC-based Bayesian inference scheme that can be used to find a mutation tree (a partial temporal order of mutations) that best fits the observed single-cell genotypes. The concentration on the mutation tree (as opposed to a cell lineage tree) makes the use of SCITE very efficient for use with our data that is characterized by few mutational events and many cells.

SCITE operates with 2 parameters, one for the false positive rate (FPR) and one for the false negative rate, which can be either set to predefined values or inferred in the MCMC model along with the tree structure. We used a global estimate of the sequencing error rate as the FPR (1%) and dataset-specific estimates of the dropout rate (ADO provided by the platform) as the false negative rate (FNR). In cases where no dropout rate was estimated, we let SCITE learn the value from the data by giving it the average value of the estimates across all patients as a prior estimate. We ran SCITE separately for each patient, providing the table of mutation calls as the input (encoding 0 for wild-type, 1 for mutation, and 3 for missing data point). To obtain a robust model, we ran SCITE with 4 different combinations of parameters: 1) use all cells including missing genotype information with 1% FPR and SCITE inferred FNR, 2) use all cells including missing genotype information with 1% FPR and platform provided FNR, 3) use only cells with full genotype information with 1% FPR and SCITE inferred FNR, and 4) use only cells with full genotype information with 1% FPR and platform provided FNR. When provided with an incomplete genotype for a cell, SCITE is still able to use the partial genotyping information in the tree inference and assigns cells into subclones based on the available information.

The inference procedure underlying SCITE is fully Bayesian, which allowed us to quantify uncertainty in the inferred clonal architectures by sampling trees from the model’s posterior distribution. We summarized the sampled trees by reporting 95% credible intervals for the inferred subclones.

The tree structure (branching vs. linear) were mostly consistent among the 4 models (47 of 76 [62%] cases showing consistent tree structure, Extended data. Fig. 12). Phylogeny figures that are shown in Fig. 3 are based on model 2 (all cells, 1% FPR, and platform provided FNR). For longitudinal samples, we combined the scDNA-seq data from all time points from the same patient and ran SCITE for the pooled data, and reconstructed the tumor phylogeny. To obtain time point-specific estimates of subclone sizes, we performed the cell to subclone assignment in the posterior sampling separately for each time point. As in some cases not all mutations were observed at all time points, we adjusted the assignment probabilities such that a cell cannot be placed below any mutation unobserved at the cell’s sampling time. This leads to subclones with a temporary prevalence of 0%. This does not necessarily mean that the subclone was non-existent/extinct at that time, but simply reflects the lack of evidence for its existence based on the cells sampled at the respective time point. The number of subclones was defined as the number of distinct cellular populations carrying at least one mutations based on model 2.

### SNP array

Genomic DNA from 28 samples in which scDNA-seq data showed at least 5% of homozygously mutated clones were analyzed by Illumina Omni2.5-8 SNP array. The raw data retrieved from an Illumina Omni2.5-8 SNP array was processed using GenomeStudio 2.0. The raw log R ratio and B allele frequency were used for ASCAT (allele-specific copy number analysis of tumors) algorithm ^37^ to identify copy-number alterations.

### Droplet digital PCR

We performed droplet digital PCR (ddPCR) using QX200™ Droplet Digital ™ System (Bio-Rad Laboratories) to confirm the variants that were detected by scDNA-seq but were not detected by bulk NGS. ddPCR™ Supermix for Probes (No dUTP) was used with 50ng of genomic DNA as a template for ddPCR assay in a 96-well plate according to the manufacture’s protocol. 7ng of synthesized mutant DNA (designed through Bio-Rad Laboratories and ordered through Integrated DNA Technologies) in a background of 130ng of normal human genomic DNA (Promega) was used as a positive control. 50ng of normal human genomic DNA (Promega) was used as a negative control. Water was used instead of DNAs for no-template control reactions. Each reaction was tested in duplicate. Variant-specific primers/probes (ddPCR™ Mutation Detection Assays, FAM/HEX for mutant/wildtype) were designed and ordered through Bio-Rad Laboratories and are summarized in Supplementary Table 6. Data was analyzed using Quanta-Soft Analysis Pro software v1.0.596 (Bio-Rad Laboratories).

### Statistical analysis

Categorical variables were compared using Chi-squared or Fisher’s exact tests. Continuous variables were analyzed by Student’s *t*-tests or Mann-Whitney *U* test depending on the satisfaction of the statistical testing assumptions. Spearman’s rank correlation coefficient (r_s_) was used to assess the relationships between two continuous variables that did not follow a normal distribution. To evaluate cell-level co-occurrence and mutual exclusivity, a contingency table was constructed to compute the log2-transformed odds ratios. Fisher’s exact test was used to evaluate the statistical significance of associations. The Benjamini-Hochberg method was used to adjust for multiple testing ^38^. In order to assess the prognostic relevance of clonal heterogeneity, we collected survival information for previously-untreated 64 AML patients. Overall survival was calculated from the date of pretreatment sample collection to the date of death from any cause, and censored on the date of last follow-up if alive. Those who underwent stem cell transplantation was censored on the date of transplantation. Kaplan-Meier plots were used to visualize survival distributions. Differences in survival between groups were analyzed using log-rank tests. We considered *P* value of less than 0.05 to be statistically significant. R (ver. 3.4.3) and EZR ^39^ software packages were used for statistical analysis.

## Code availability

Publicly available codes were used with a citation for data analysis. In-house codes that were used for single-cell sequencing data variant calling are available from the corresponding author on reasonable request.

## Data availability

Deidentified clinical and genetic data is available in supplementary information.

## Acknowledgments

This study was supported in part by the Cancer Prevention and Research Institute of Texas (grant R120501 to PAF), the Welch Foundation (grant G-0040 to PAF), the University of Texas System STARS Award (grant PS100149 to PAF), Physician Scientist Program at MD Anderson (to KT), Lyda Hill Foundation (to PAF), the Charif Souki Cancer Research Fund (to HK), the MD Anderson Cancer Center Leukemia SPORE grant (NIH P50 CA100632) (to HK), the MD Anderson Cancer Center Support Grant (NIH/NCI P30 CA016672), Research Fellowships of the Japan Society for the Promotion of Science for Young Scientists (to KM), and generous philanthropic contributions to MD Anderson’s Moon Shot Program (to PAF, KT, GGM, and HK). We thank Amy Ninetto at Department of Scientific Publications at MD Anderson for providing scientific editing of the manuscript. We also thank Charles Silver, Dennis Eastburn, Robert Durruthy-Durruthy, Matt Cato, Hannah Viernes, Anup Parikh, Sombeet Sahu, Kelly Kaihara, and all others members of Mission Bio Inc. for the technical support.

## Author contributions

KM performed the experiments, analyzed the data, and wrote the initial draft of the manuscript. KT designed the study and wrote the manuscript. KJ, JK, and NB performed the phylogenetic analysis. FW, JZ, and XS performed the bioinformatic analysis. YY performed the statistical analysis. JM collected samples. LL, CG, SC, and ET performed sequencing. KP and CBR performed pathologic analyses. CD, FR, EJ, MA, JC, MK, KB, GGM, and HK collected samples and treated patients. NN and PAF critically reviewed the manuscript. PAF and KT provided leadership and managed the study team. All authors read and approved the manuscript.

