## Extended Data for "Clonal Evolution of Acute Myeloid Leukemia Revealed by High-Throughput Single-Cell Genomics"

#### Extended Data Fig. 1

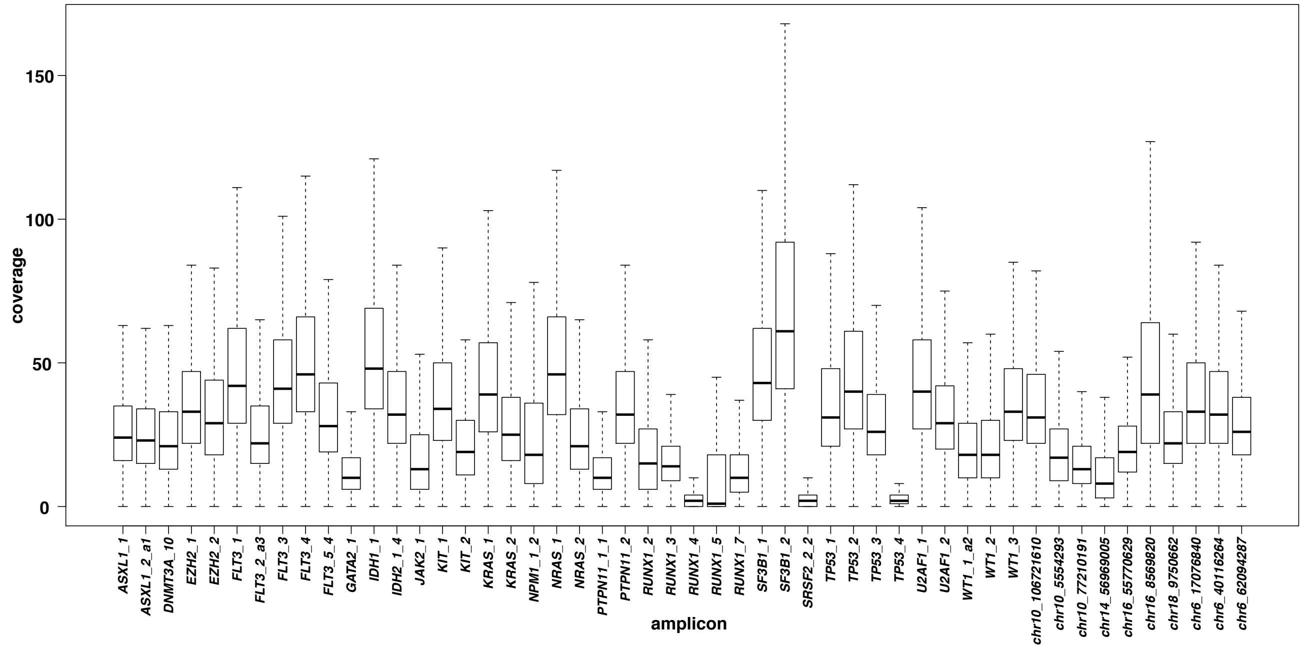


**Extended Data Fig. 1. Distribution of coverage per cell per amplicon in all sequenced cells from 77 patients.** The amplicons are shown on the X axis. The first 40 amplicons cover the hotspots of 19 AML-associated genes. The remaining 10 amplicons cover commonly heterozygous SNP loci. The Y axis represents the coverage for each sequenced cell. The thick line within each box represents the median, and the top and bottom edges of the box represent the 25th and 75th percentiles, respectively. The upper and lower whiskers represent the 75th percentile plus 1.5 times the interquartile range and the 25th percentile minus 1.5 times the interquartile range, respectively. Data points that fell outside of the upper and lower whiskers were considered outliers and are not shown.

**Extended Data Fig. 2**

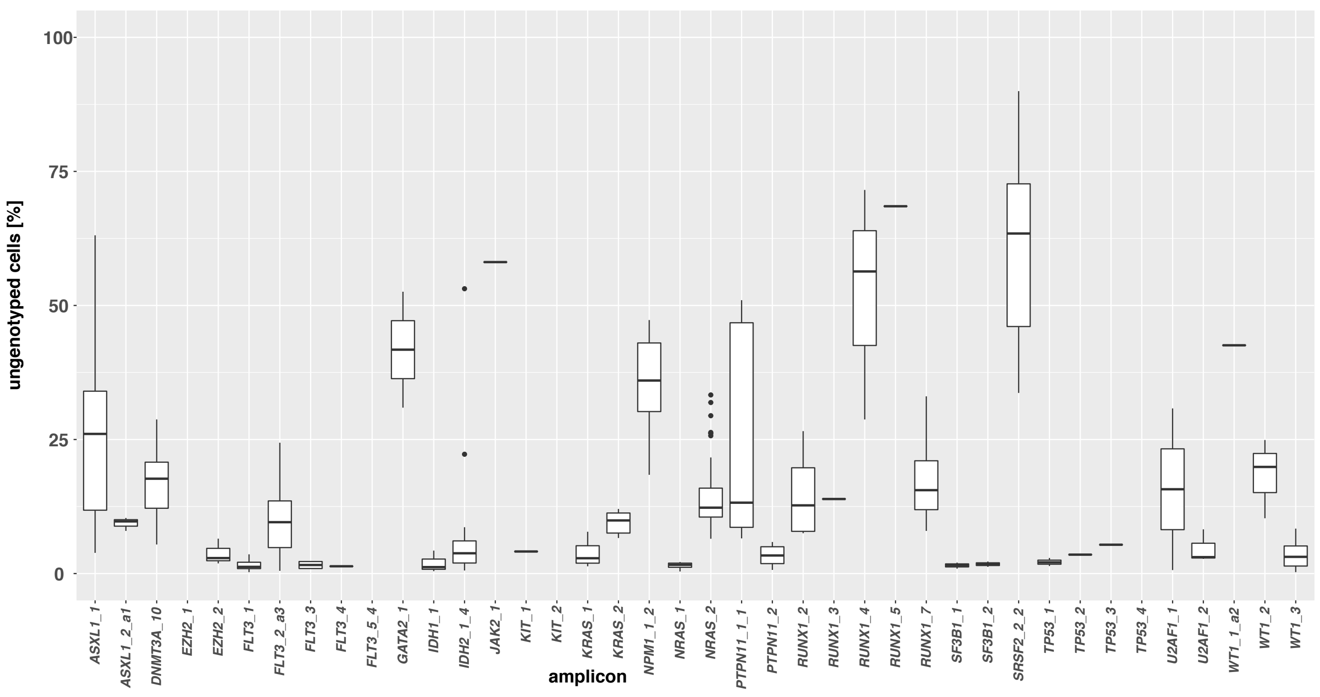


**Extended Data Fig. 2. Distribution of the percentage of ungenotyped cells for each variant based on the amplicon.** The 40 amplicons covering the hotspots of 19 AML-associated genes are shown on the X axis. The Y axis represents the percentage of ungenotyped cells for each variant. The percentage of ungenotyped cells were calculated for each variant from each sample as follows: (number of genotyped cells [wildtype, heterozygous, or homozygous]) / (number of total sequenced cells) × 100.

**Extended Data Fig. 3**

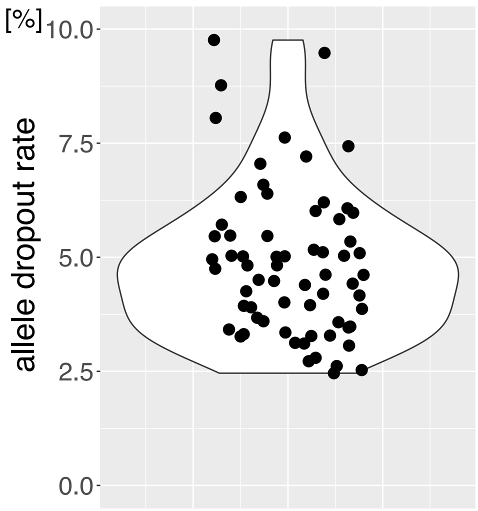


**Extended Data Fig. 3. Violin plot with jittered points showing the distribution of the allele dropout (ADO) rate.** The Y axis shows the ADO rate from unique patients. Each point represents a sample. ADO was not obtainable for 11 of the 77 patients.

**Extended Data Fig. 4**

*KRAS* exon2:c.G35A:p.G12D
(mutated in 67 of 7053 [0.9%] cells sequenced)
bulk NGS VAF :undetectable

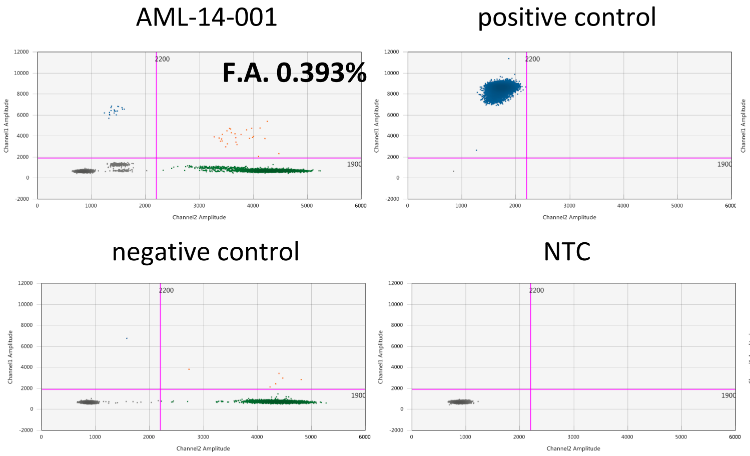


*IDH2* exon4:c.G419A:p.R140Q
(mutated in 56 of 9864 [0.6%] cells sequenced)
bulk NGS VAF: undetectable

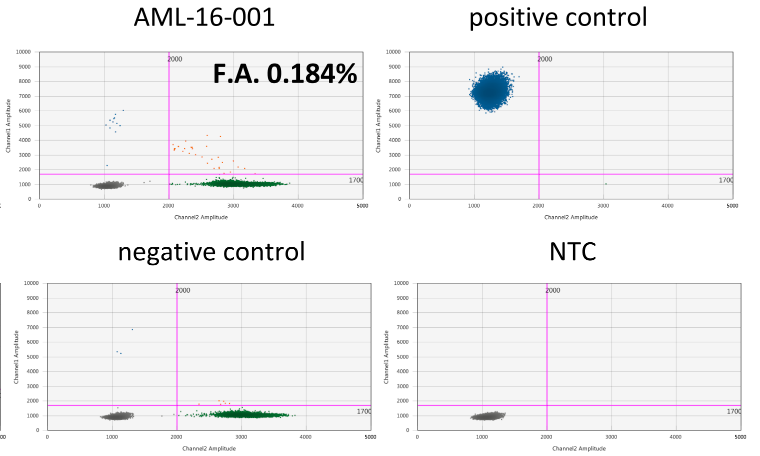


*NRAS* exon2:c.G35A:p.G12D
(mutated in 42 of 8219 [0.5%] cells sequenced)
bulk NGS VAF: undetectable

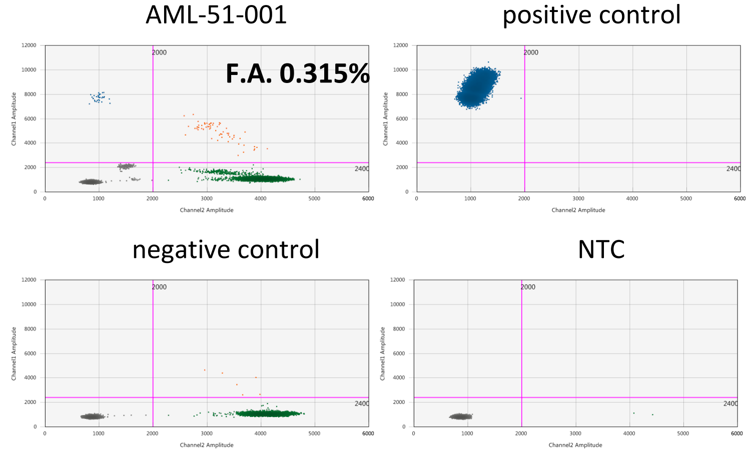


*NRAS* exon2:c.G38T:p.G13V
(mutated in 10 of 2252 [0.4%] cells sequenced)
bulk NGS VAF: undetectable

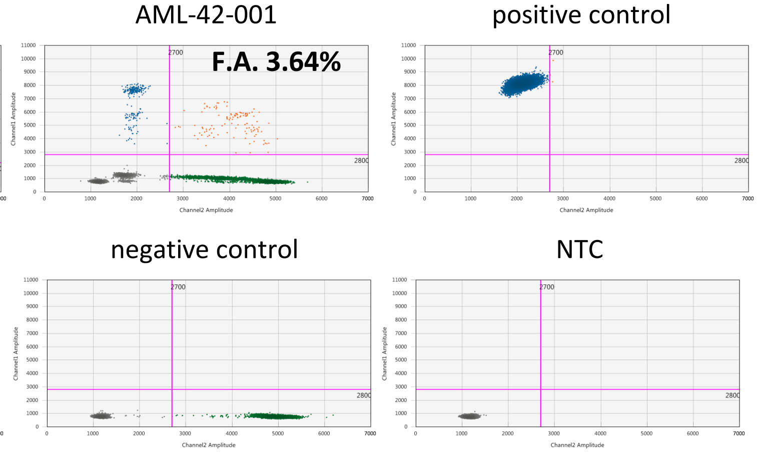


*NRAS* exon2:c.G38A:p.G13D
(mutated in 32 of 11398 [0.3%] cells sequenced)
bulk NGS VAF: undetectable

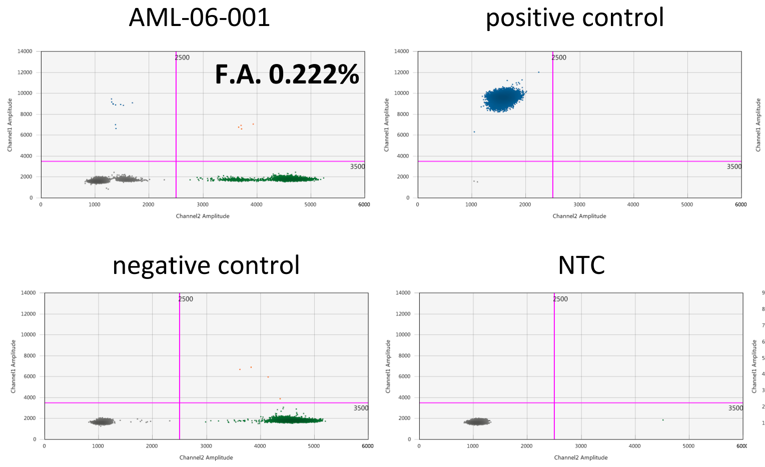


*NRAS* exon2:c.G35A:p.G12D
(mutated in 6 of 4359 [0.1%] cells sequenced)
bulk NGS VAF: undetectable

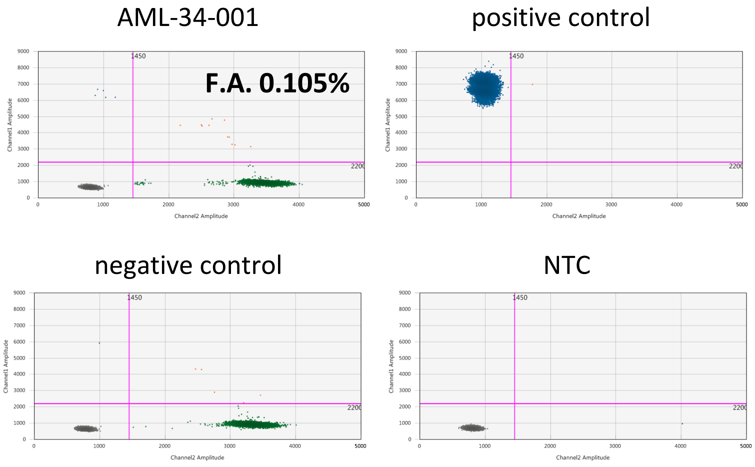

**Extended Data Fig. 4. Validation of single-cell DNA sequencing-specific variants by droplet digital PCR (ddPCR).** Representative cases of single-cell DNA sequencing-specific variants that were undetectable by bulk next generation sequencing but were validated by ddPCR. 2D plots for ddPCR results are shown for the indicated sample (upper left), a positive control using synthesized DNA (upper right), a negative control using wild-type human DNA (lower left), and a no-template control without DNA (lower right). Within each 2D plot, the blue cluster in the upper left quadrant represents droplets with mutated DNA only, and the orange cluster in the upper right quadrant represents droplets with both mutated and wild-type DNA. The green cluster in the lower right quadrant represents droplets with wild-type DNA only, and the grey cluster in the lower left quadrant represents droplets without DNA from the targeted locus. Fractional abundance (F.A.) was calculated as follows: (number of droplets with mutated DNA) / (number of droplets with mutated DNA + number of droplets with wild-type DNA) × 100. The smallest mutations that were validated by ddPCR assay was mutated in 0.1% of the total sequenced cells. Therefore, along with the cell line data shown in Supplemental Table 2, the limit of detection of the platform was estimated as 0.1%.

NGS, next-generation sequencing; VAF, variant allele fraction; NTC, no-template control.

**Extended Data Fig. 5**

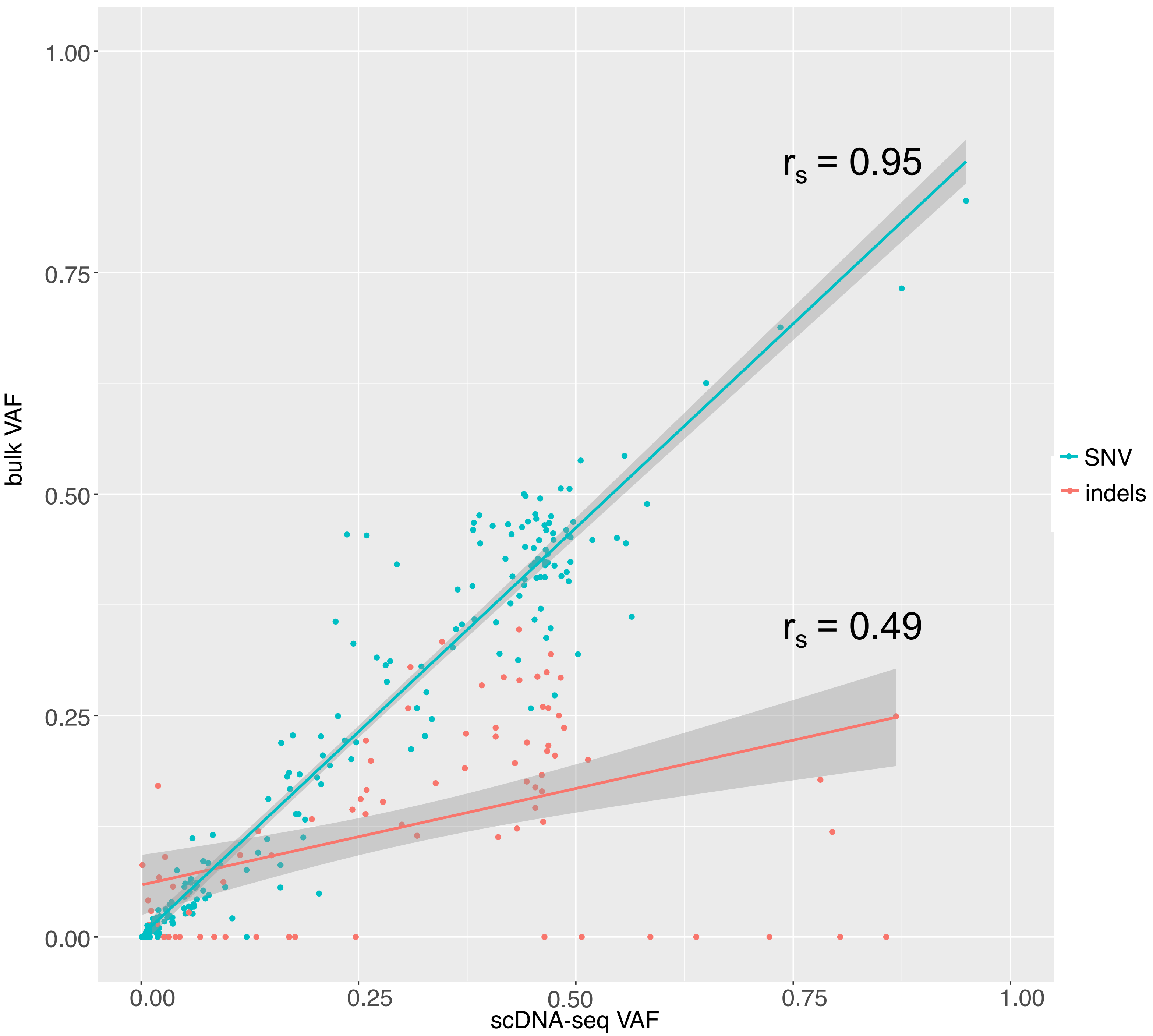


**Extended Data Fig. 5. Scatter plot showing the correlation of the bulk-sequencing VAF and the VAF inferred from the single cell genotype data based on the types of mutations.** The X-axis shows the VAF from the single-cell genotype data (scDNA-seq VAF). The Y-axis shows the VAF from the bulk next-generation sequencing (bulk VAF). Green dots represent single-nucleotide variants (SNV), and red dots represent insertion/deletion variants (indels). The linear trendlines were added to best fit the distribution of the dots. The shaded areas around the trendlines represent the 95% confidence intervals. scDNA-seq VAF and bulk VAF matched well for SNV ($r_s = 0.95$, $p < 0.001$), whereas the concordance was weaker for indels ($r_s = 0.49$, $p < 0.001$).

**Extended Data Fig. 6**

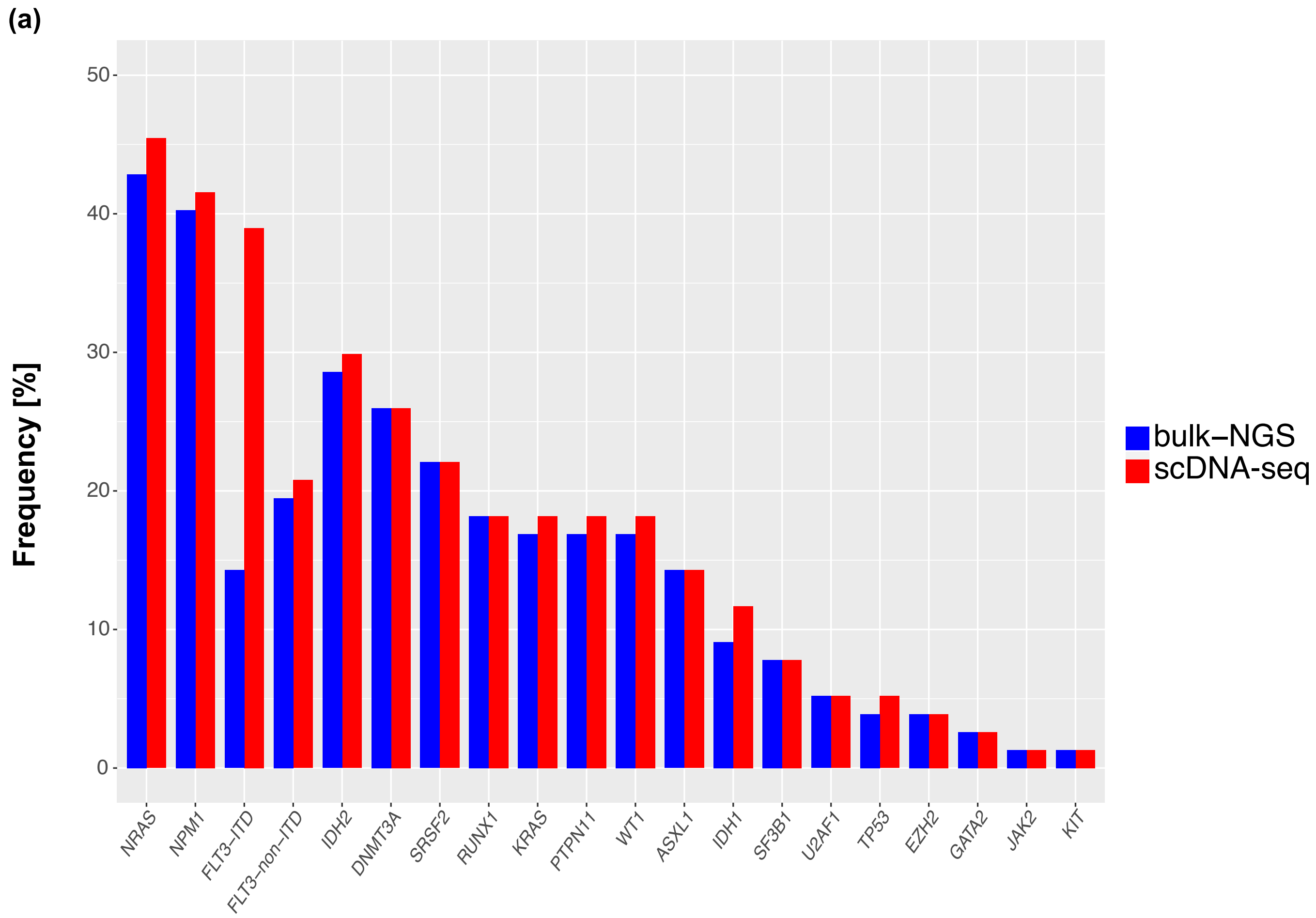


**Extended Data Fig. 6. (a) Frequency of driver mutations in 77 patients based on the data from bulk next generation sequencing (bulk-NGS) and single-cell DNA sequencing (scDNA-seq).** The X axis represents the gene. The Y axis shows the frequency of patients harboring at least one mutation within each gene. Blue bars represent the frequency based on the bulk-NGS data that was used for orthogonal validation, and red bars represent the frequency based on scDNA-seq data.

**Extended Data Fig. 6**

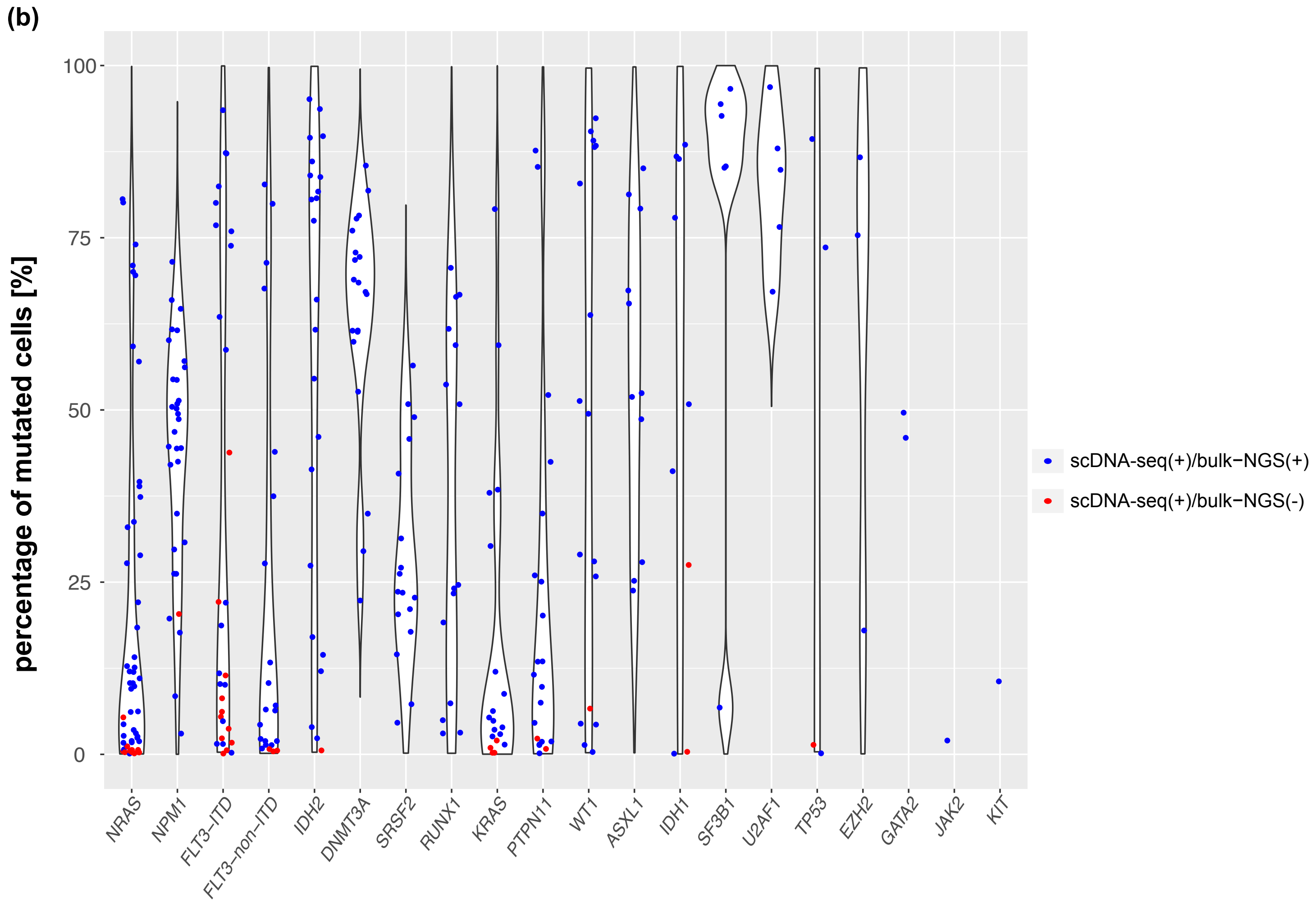


**Extended Data Fig. 6. (b) Distribution of the percentage of mutated cells for each driver mutation detected by the scDNA-seq.** Jittered plots represent variants. The X axis represents the gene. The Y axis shows the percentage of mutated cells for each variant detected by scDNA-seq. Blue plots represent the variants that were detected by both scDNA-seq and bulk-NGS. Red plots represent the variants that were detected by scDNA-seq but were undetectable by bulk-NGS. Percentage of mutated cells were calculated as follows: (number of cells that were single-cell genotyped as heterozygously- or homozygously-mutated) / (number of total sequenced cells) × 100.

**Extended Data Fig. 7**

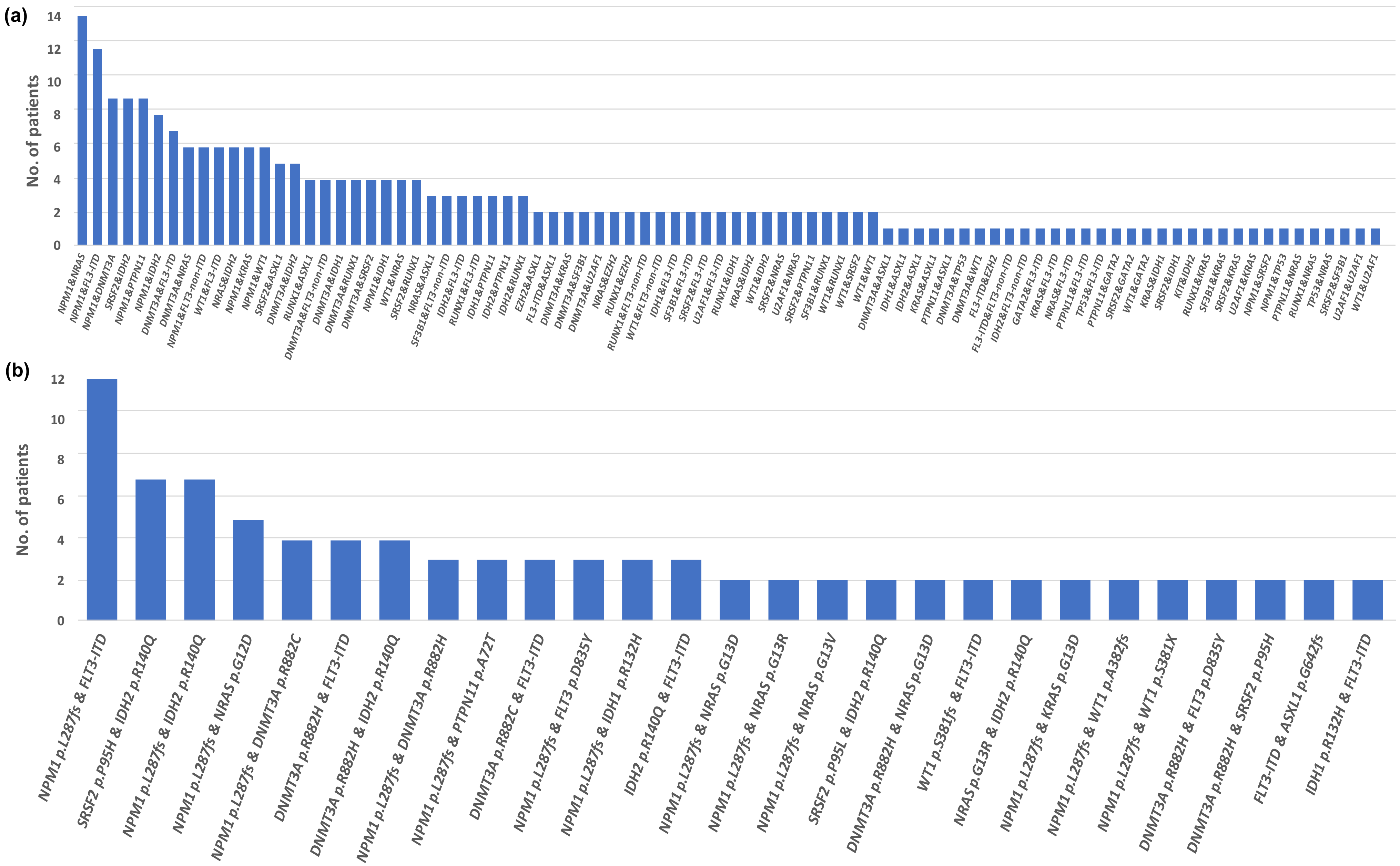


**Extended data Fig. 7. Distribution of variant combination showing statistically-significant cell-level co-occurrence (FDR < 0.001) based on the (a) mutated genes or (b) variants.** X axis represents combinations of mutated genes or variants, and y axis shows the number of patients showing the significant cell-level co-occurrence of each mutation combination. The variant combinations that were detected in only one patient are not plotted in Extended Data Fig. 7b.

**Extended Data Fig. 8**

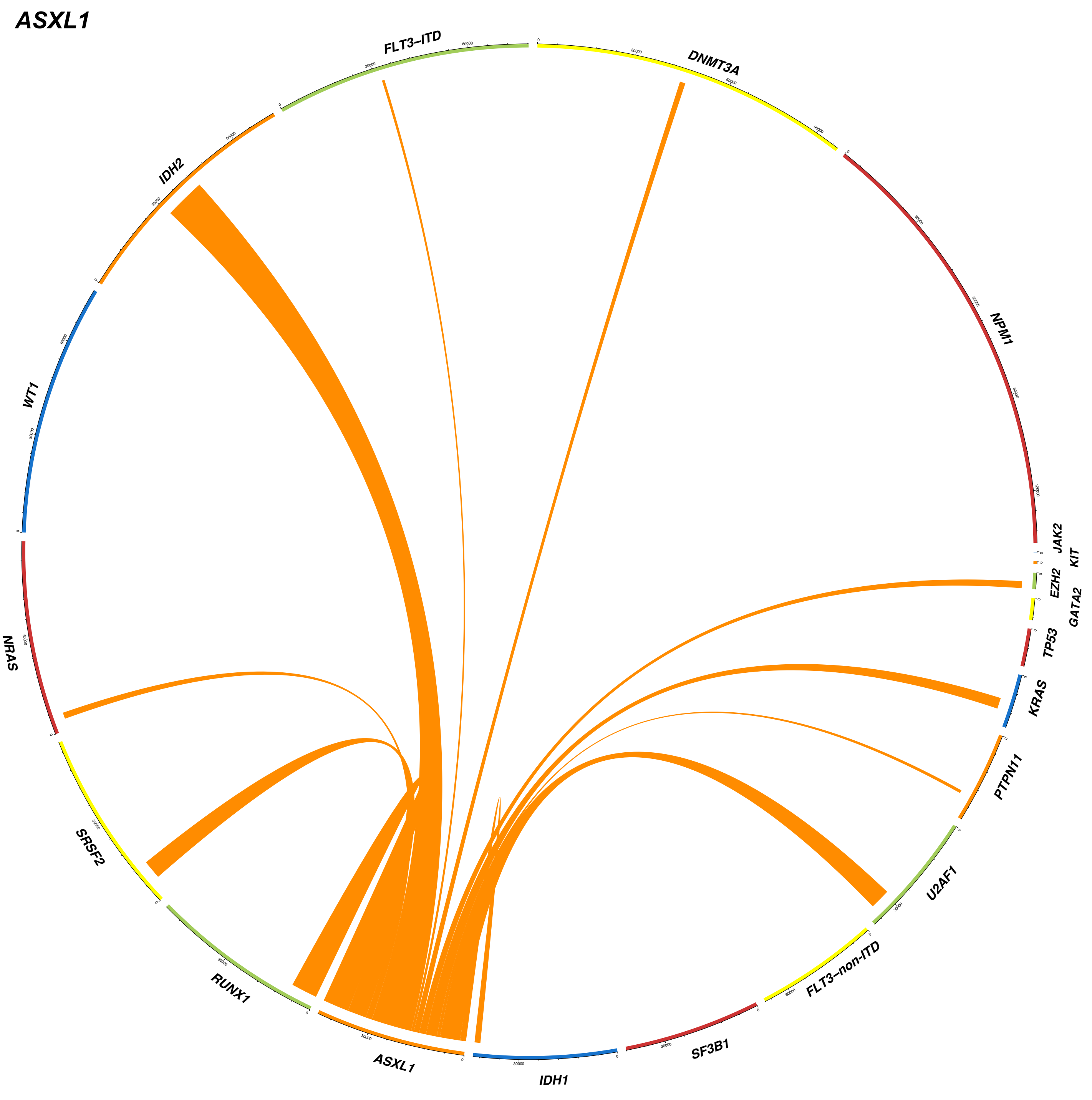

DNMT3A

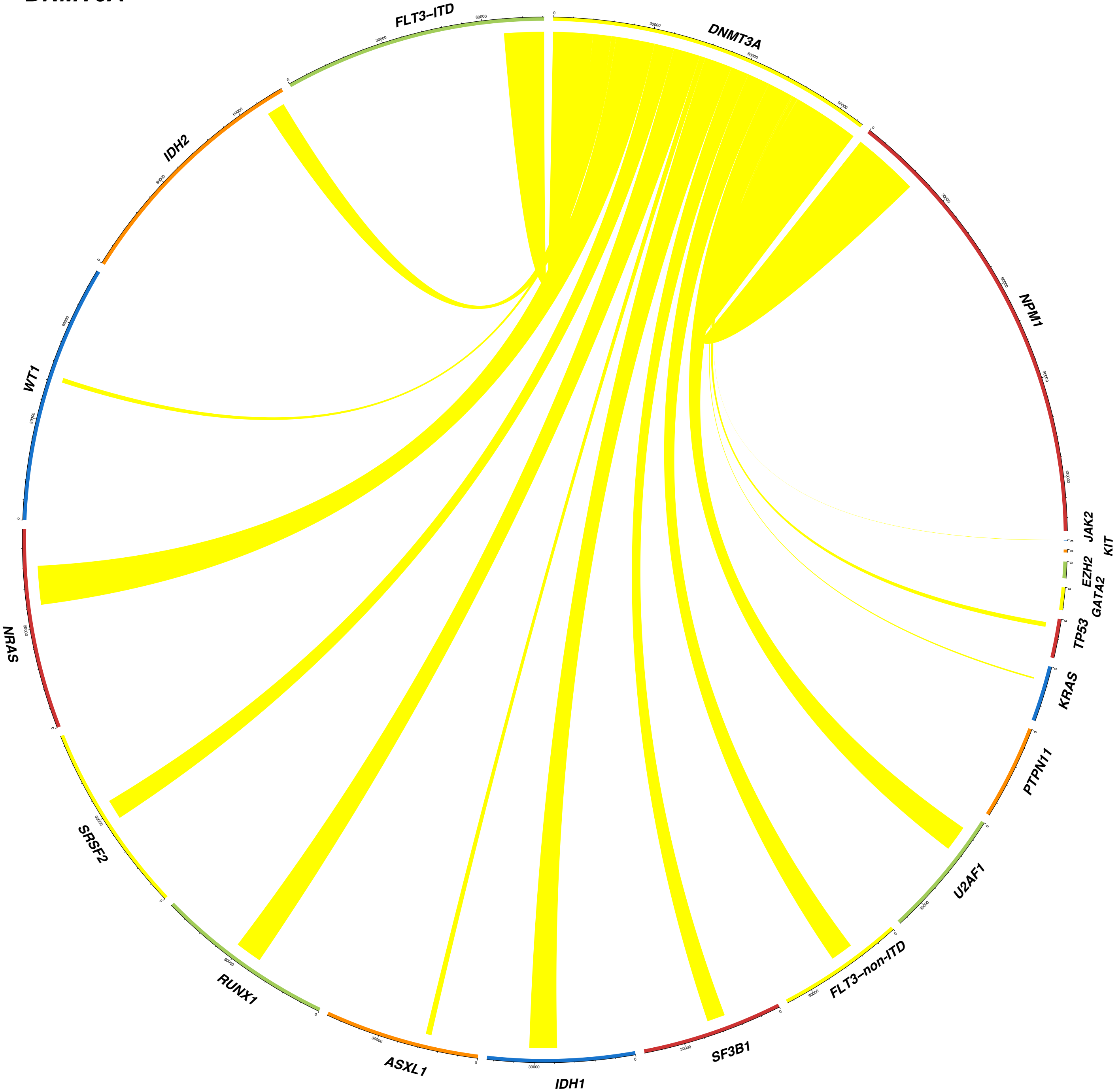
DNMT3A
FLT3–ITD
IDH2
WT1
NRAS
SRSF2
RUNX1
ASXL1
IDH1
SF3B1
FLT3–non-ITD
U2AF1
PTPN11
KRAS
TP53
GATA2
EZH2
KIT
JAK2
NPM1

#### *EZH2*

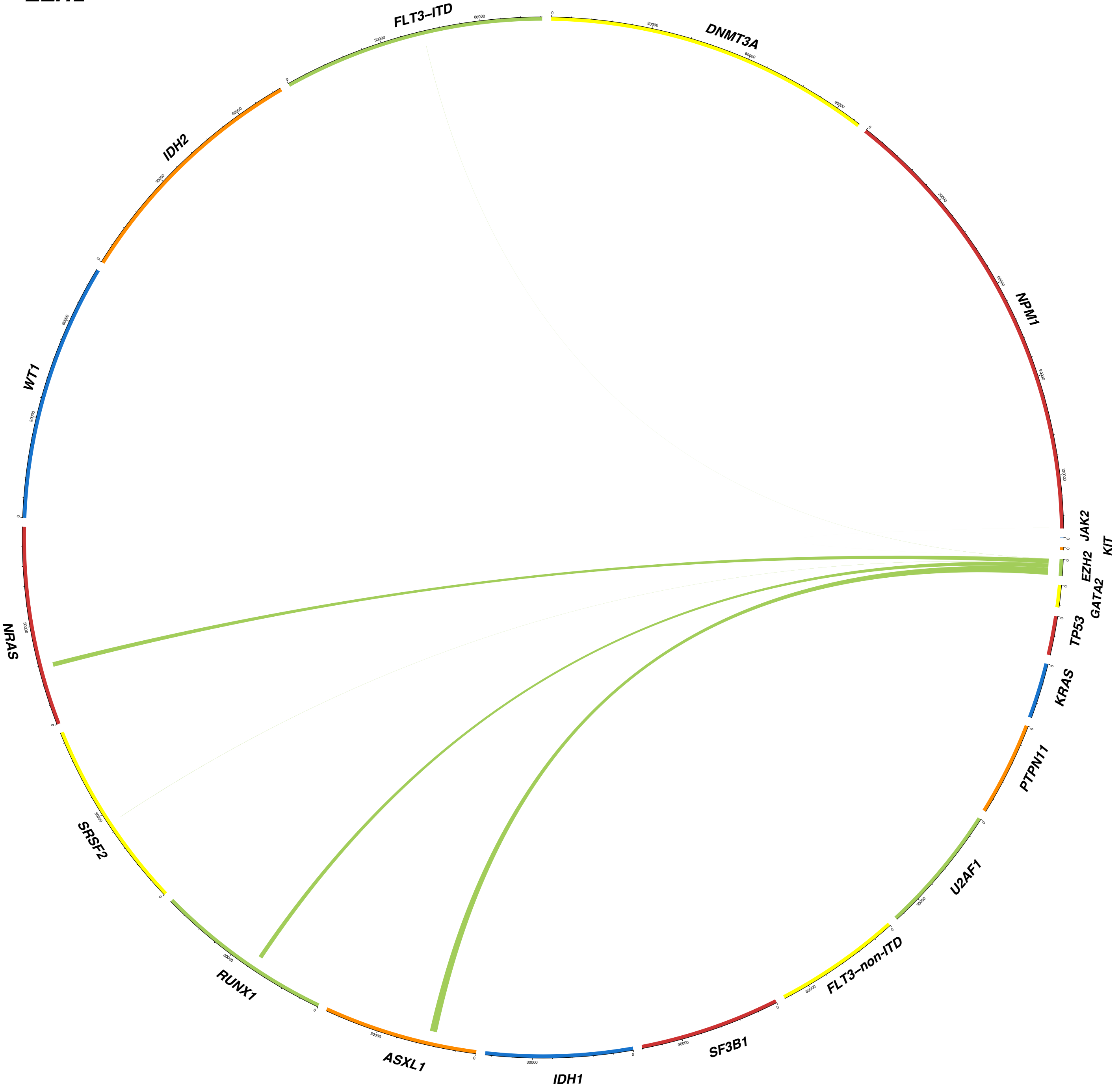
FLT3–ITD
DNMT3A
NPM1
JAK2
KIT
EZH2
GATA2
TP53
KRAS
PTPN11
U2AF1
FLT3–non-ITD
SF3B1
IDH1
ASXL1
RUNX1
SRSF2
NRAS
WT1
IDH2

*FLT3-ITD*

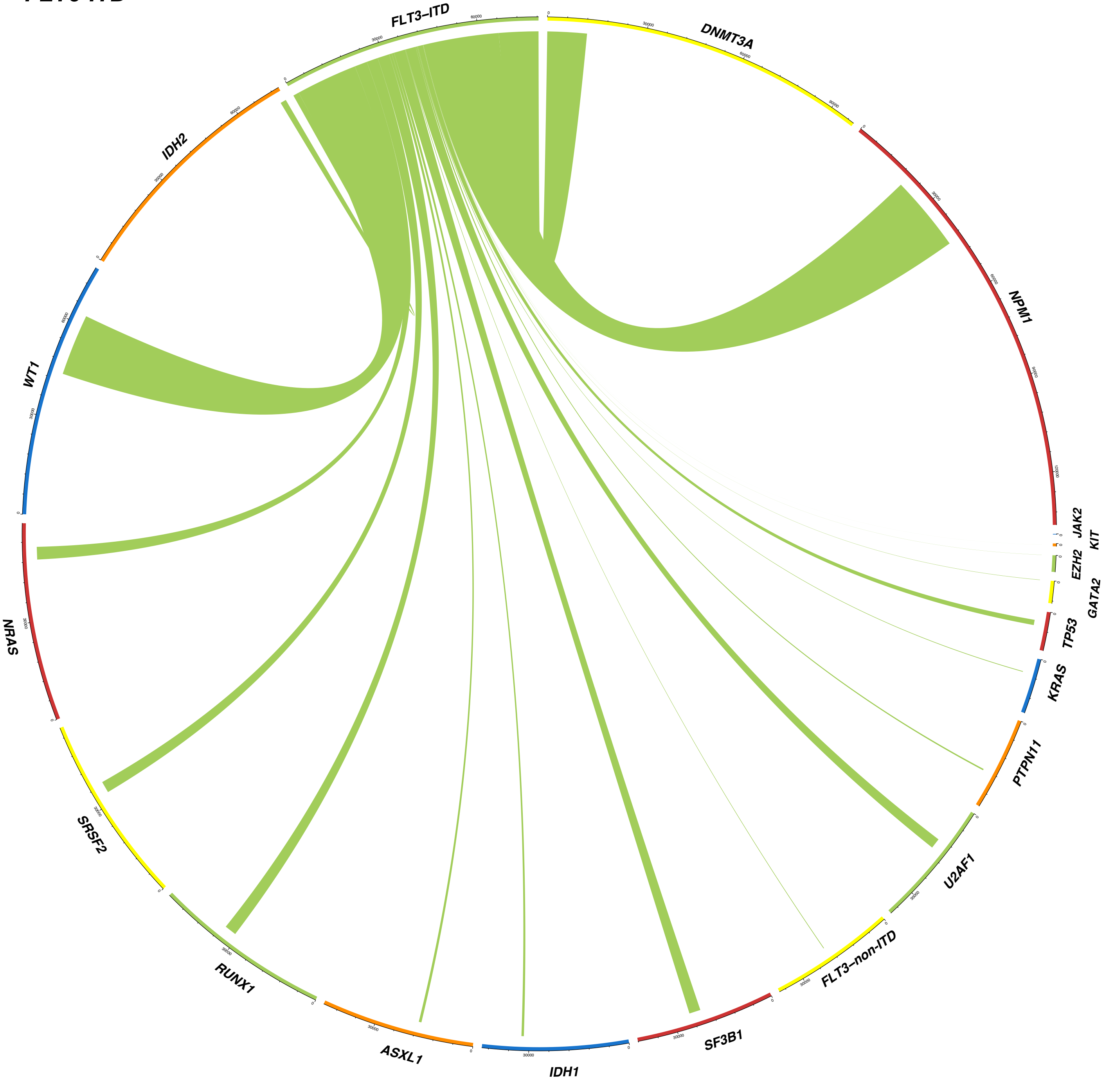

FLT3–ITD
DNMT3A
NPM1
JAK2
KIT
EZH2
GATA2
TP53
KRAS
PTPN11
U2AF1
FLT3–non-ITD
SF3B1
IDH1
ASXL1
RUNX1
SRSF2
NRAS
WT1
IDH2

#### *FLT3-non-ITD*

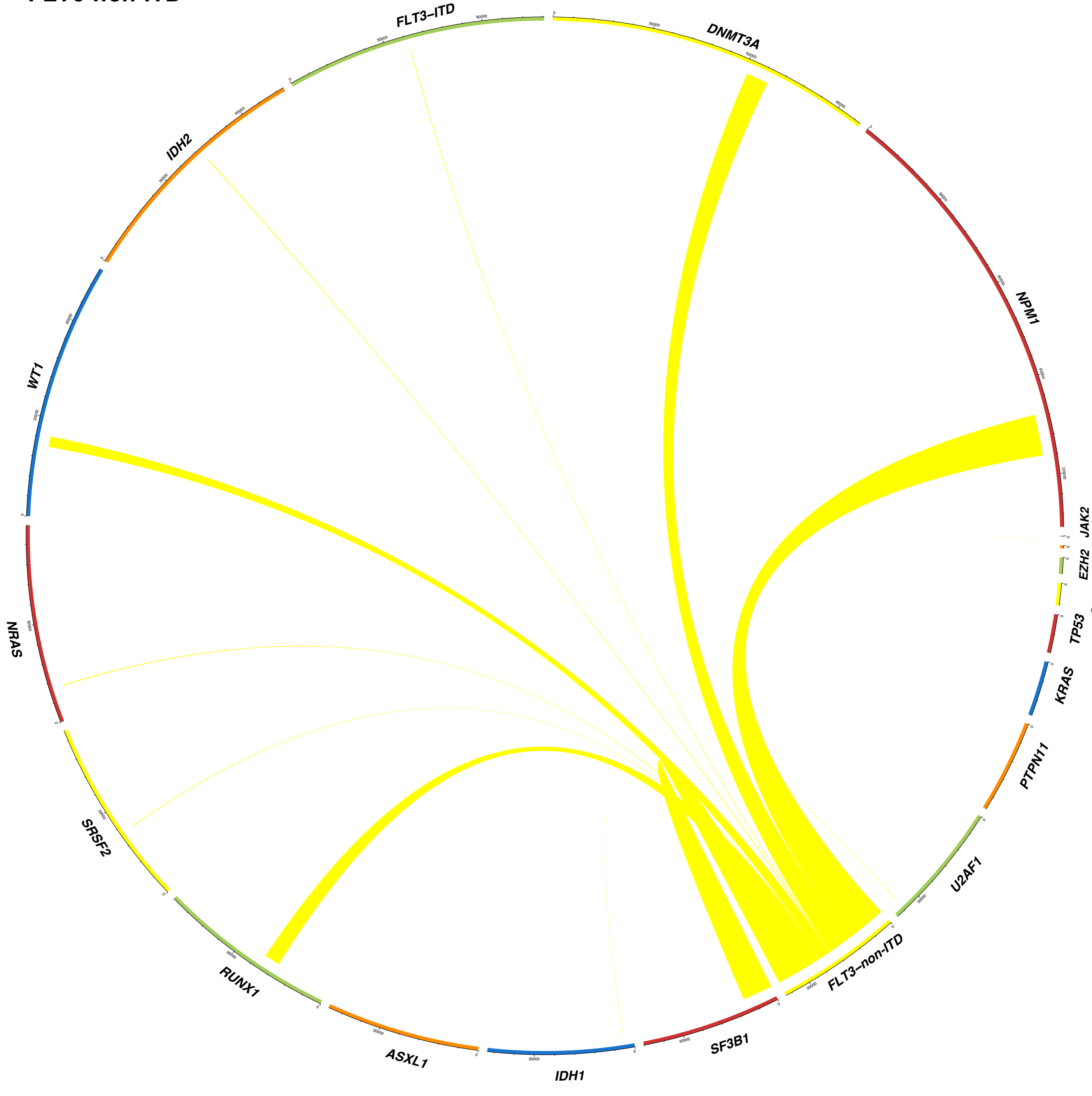

FLT3–ITD
DNMT3A
NPM1
JAK2
KIT
EZH2
GATA2
TP53
KRAS
PTPN11
U2AF1
FLT3–non-ITD
SF3B1
IDH1
ASXL1
RUNX1
SRSF2
NRAS
WT1
IDH2

#### GATA2

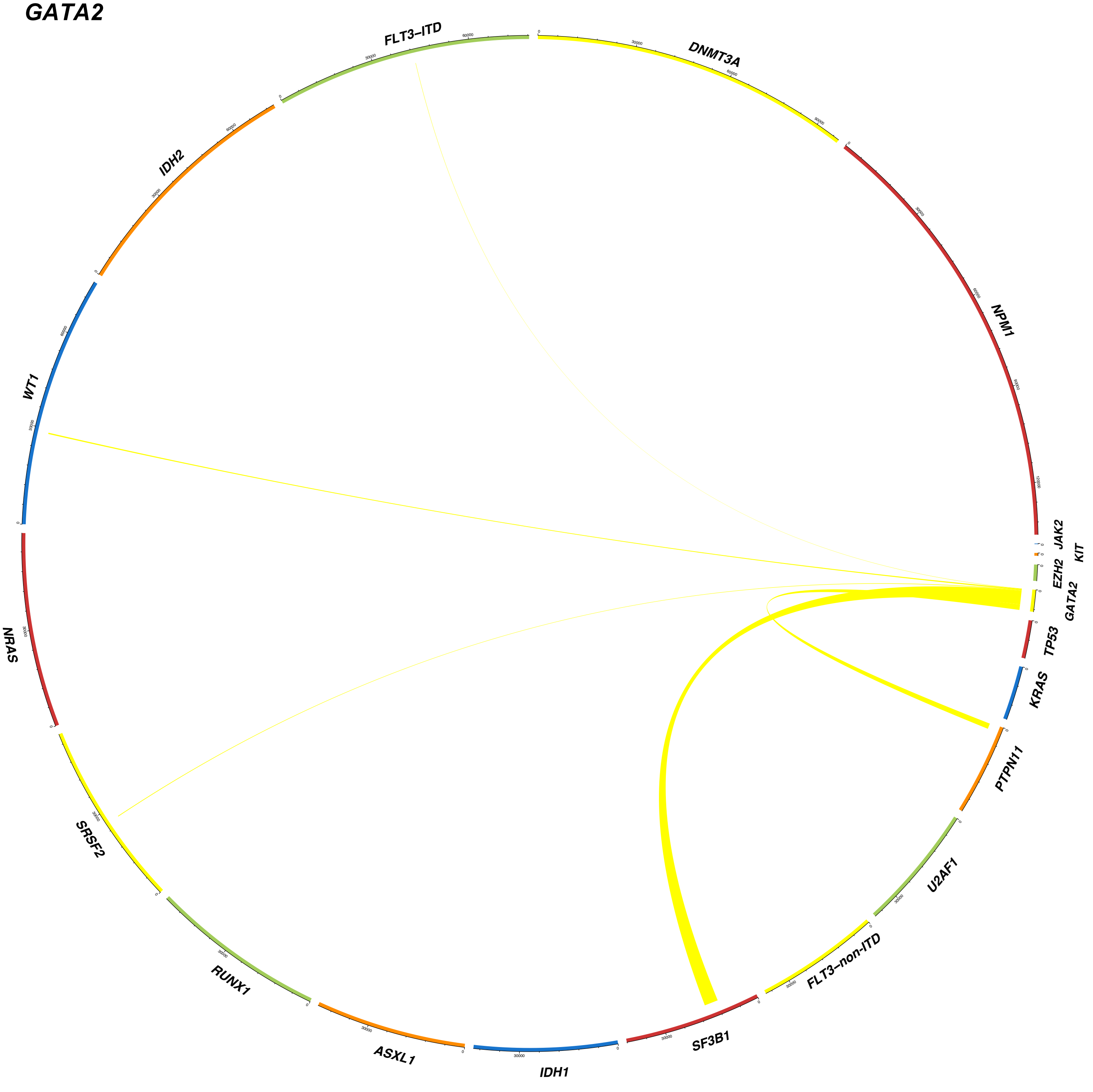
FLT3–ITD
DNMT3A
NPM1
JAK2
KIT
EZH2
GATA2
TP53
KRAS
PTPN11
U2AF1
FLT3–non-ITD
SF3B1
IDH1
ASXL1
RUNX1
SRSF2
NRAS
WT1
IDH2

#### IDH1

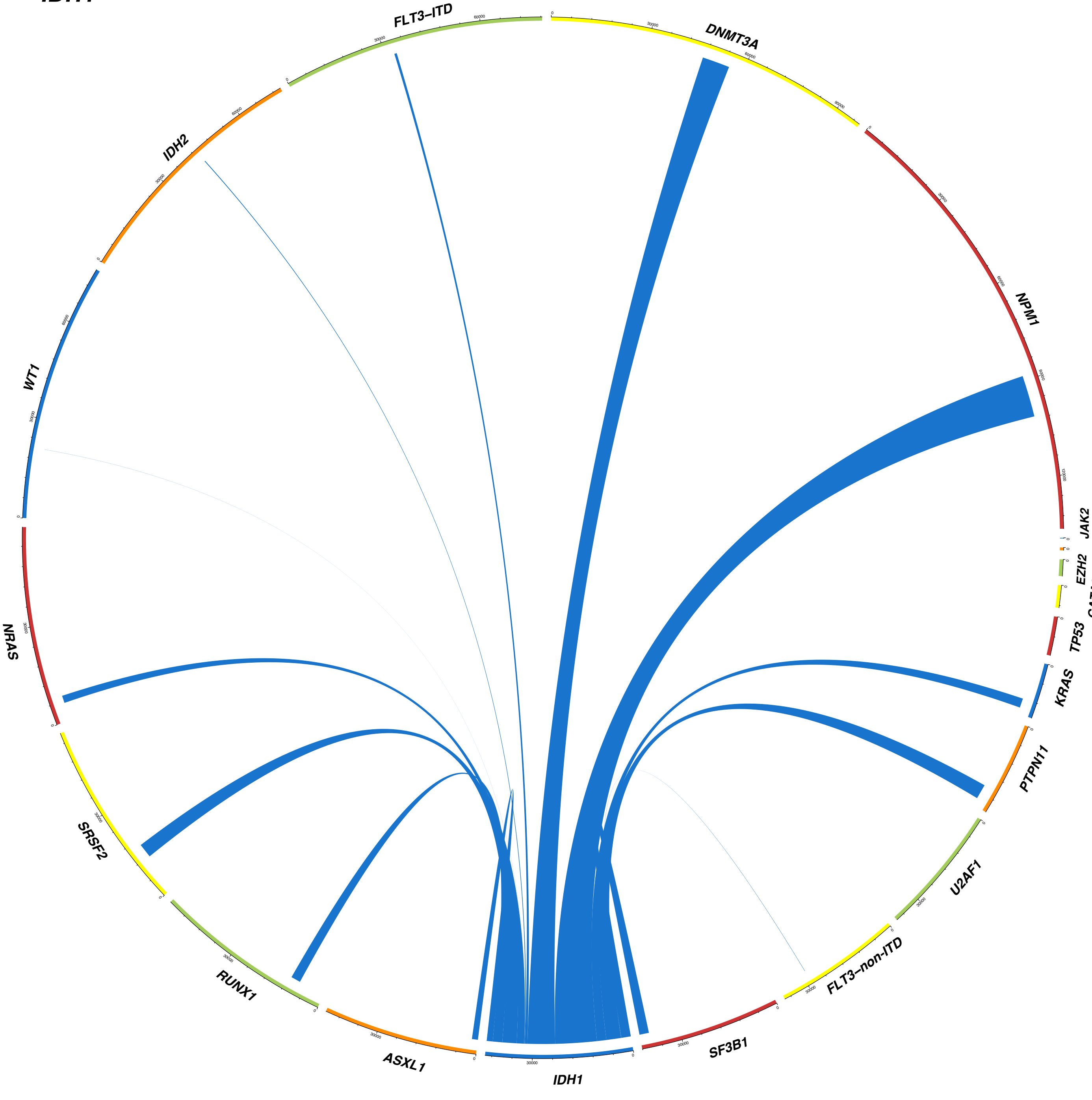

DNMT3A
NPM1
JAK2
KIT
EZH2
GATA2
TP53
KRAS
PTPN11
U2AF1
FLT3–non-ITD
SF3B1
IDH1
ASXL1
RUNX1
SRSF2
NRAS
WT1
IDH2
FLT3–ITD

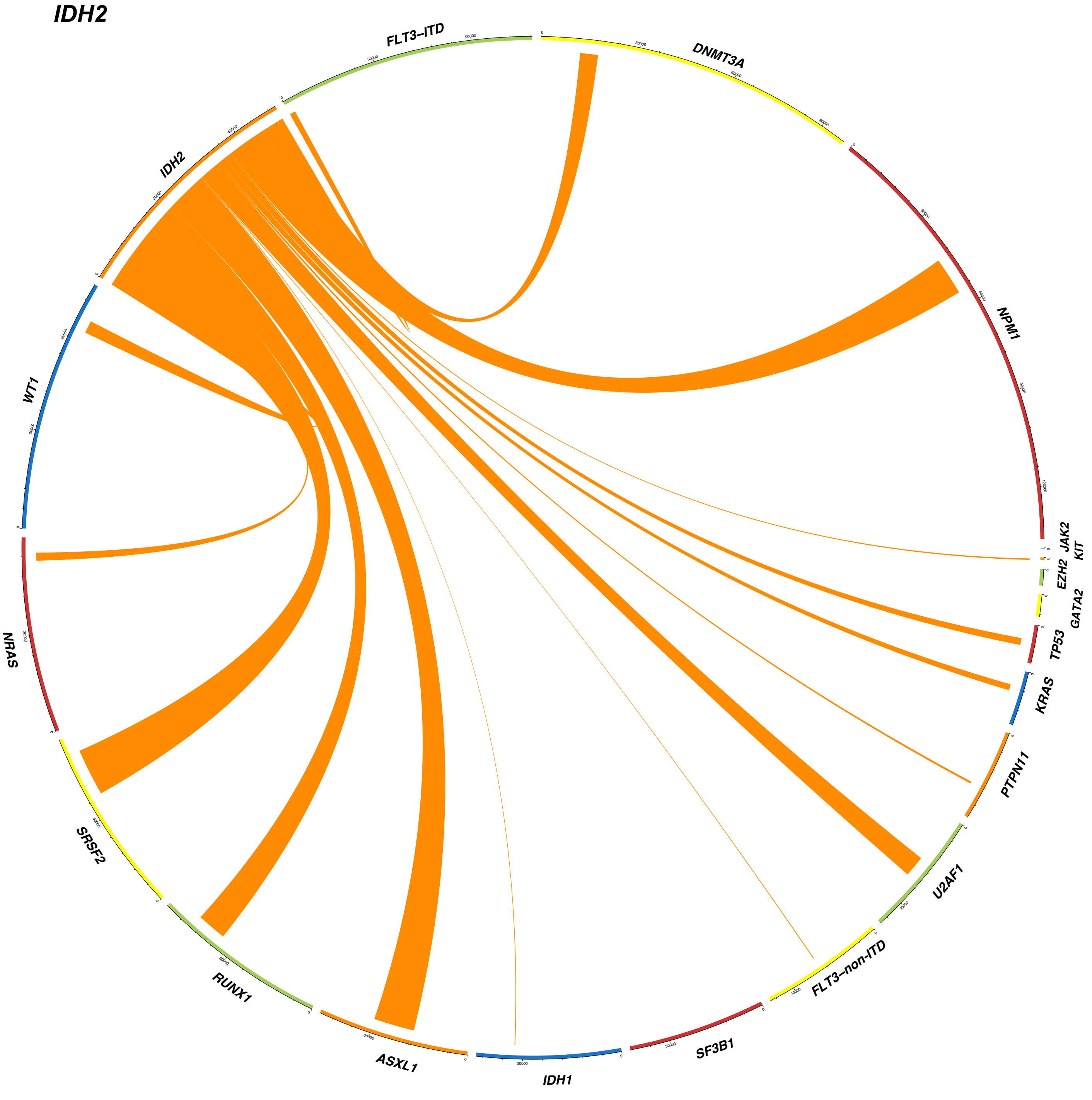
IDH2
FLT3–ITD
DNMT3A
NPM1
JAK2
KIT
EZH2
GATA2
TP53
KRAS
PTPN11
U2AF1
FLT3–non-ITD
SF3B1
IDH1
ASXL1
RUNX1
SRSF2
NRAS
WT1
IDH2

#### JAK2

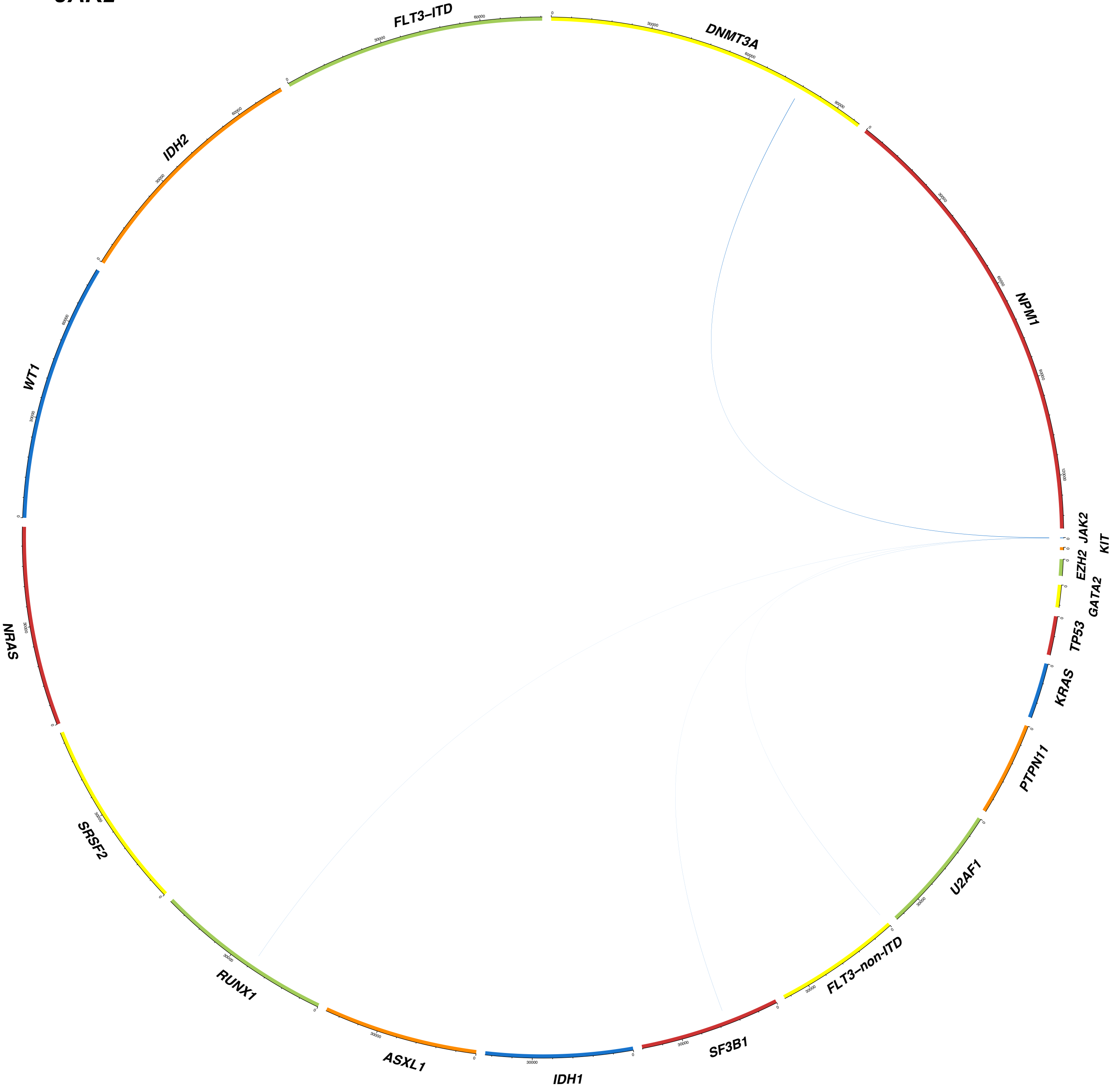

DNMT3A
NPM1
JAK2
KIT
EZH2
GATA2
TP53
KRAS
PTPN11
U2AF1
FLT3–non-ITD
SF3B1
IDH1
ASXL1
RUNX1
SRSF2
NRAS
WT1
IDH2
FLT3–ITD

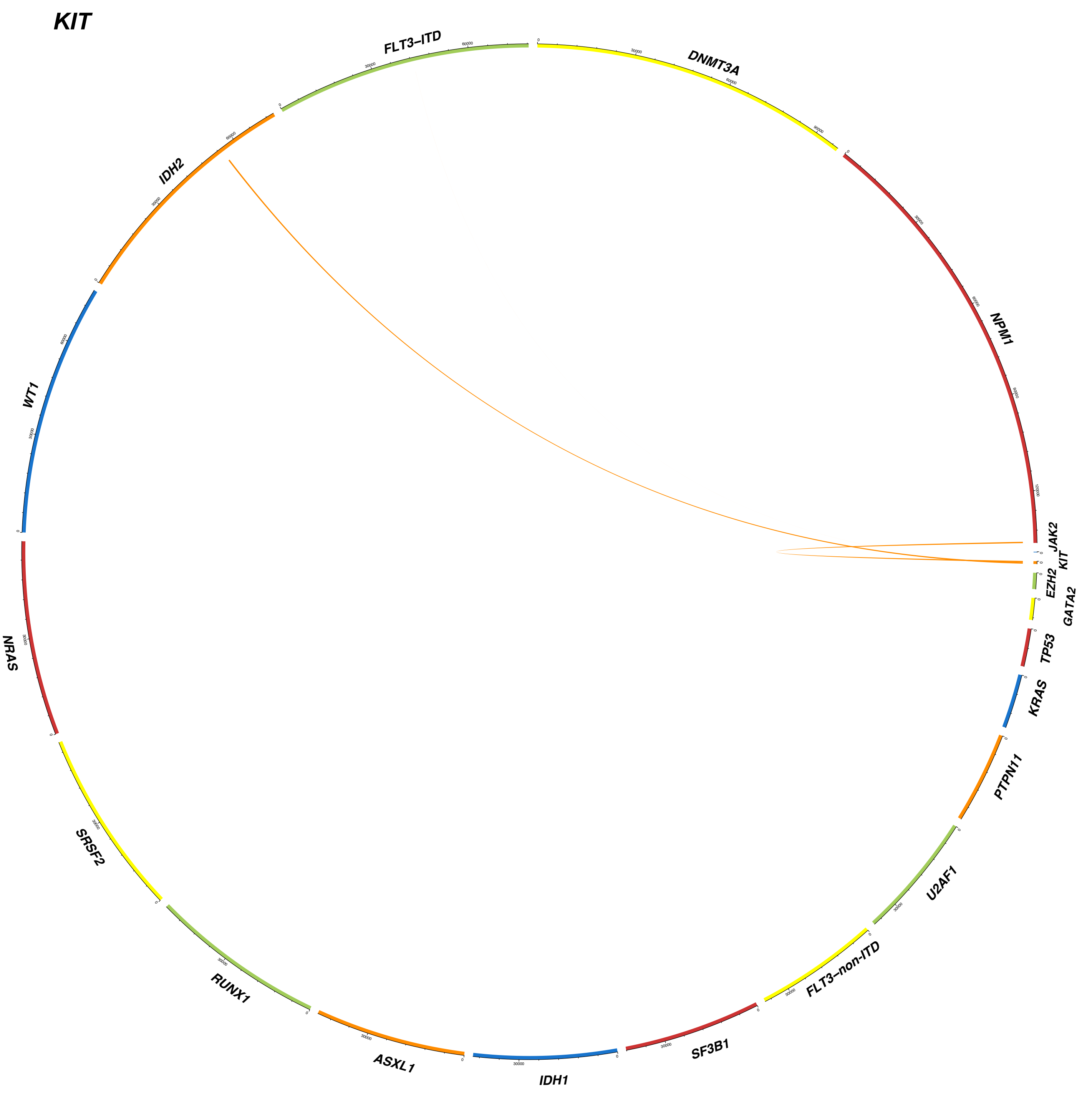
KIT
FLT3–ITD
DNMT3A
NPM1
JAK2
KIT
EZH2
GATA2
TP53
KRAS
PTPN11
U2AF1
FLT3–non-ITD
SF3B1
IDH1
ASXL1
RUNX1
SRSF2
NRAS
WT1
IDH2

#### *KRAS*

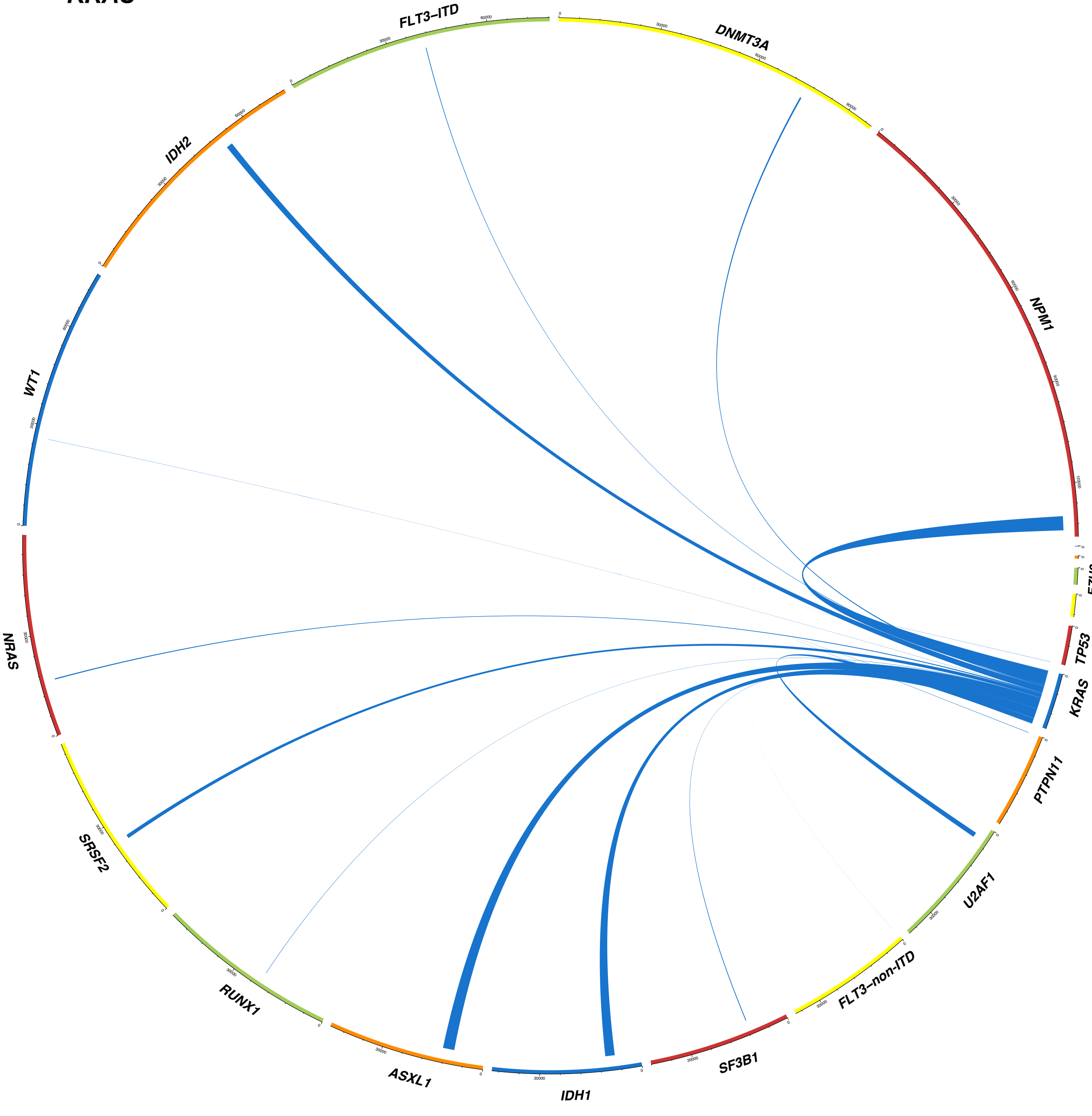

FLT3–ITD
DNMT3A
NPM1
JAK2
KIT
EZH2
GATA2
TP53
KRAS
PTPN11
U2AF1
FLT3–non-ITD
SF3B1
IDH1
ASXL1
RUNX1
SRSF2
NRAS
WT1
IDH2

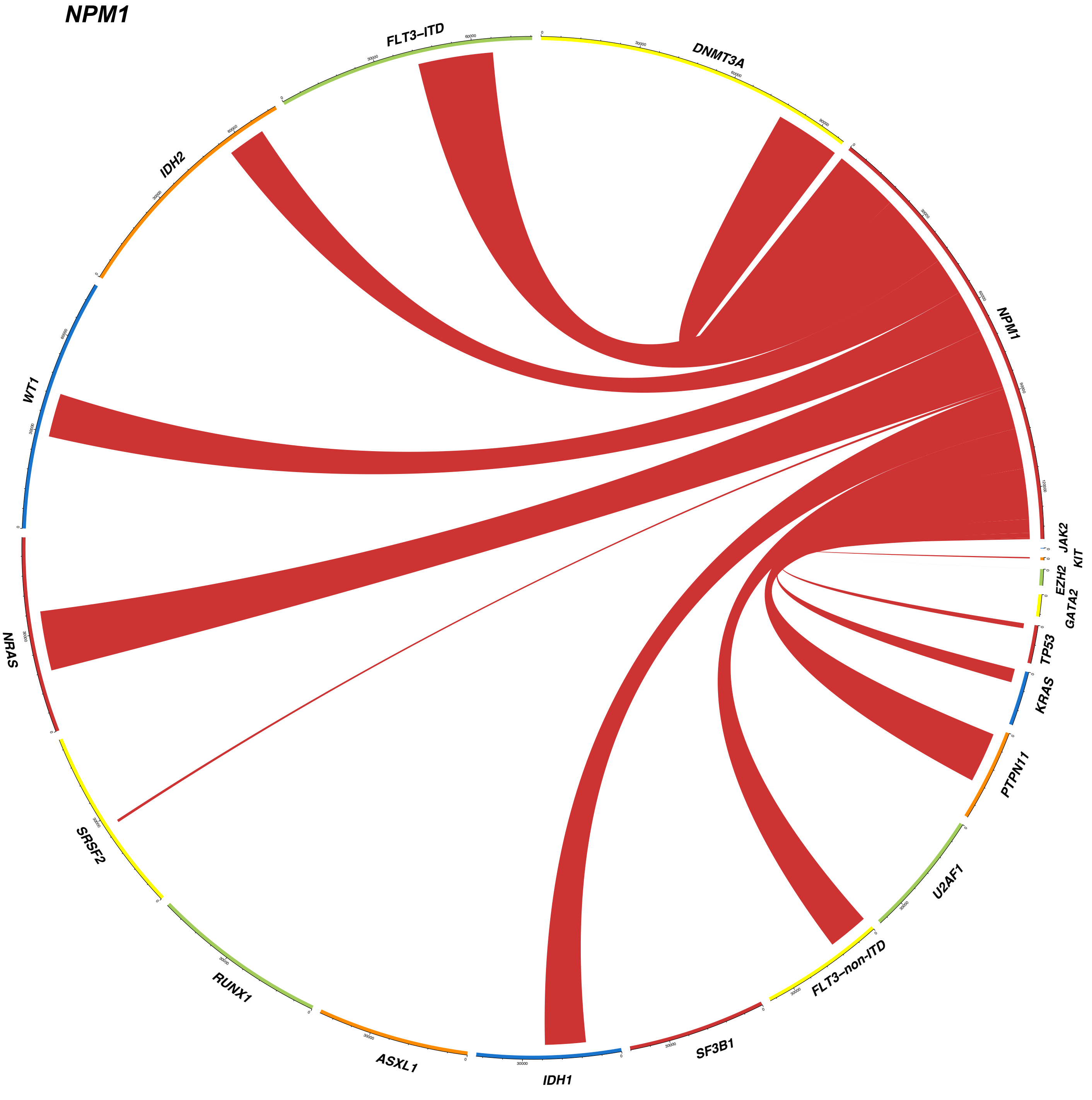
NPM1
FLT3–ITD
DNMT3A
IDH2
WT1
NPM1
JAK2
KIT
EZH2
GATA2
TP53
KRAS
PTPN11
U2AF1
FLT3–non-ITD
SF3B1
IDH1
ASXL1
RUNX1
SRSF2
NRAS

**NRAS**

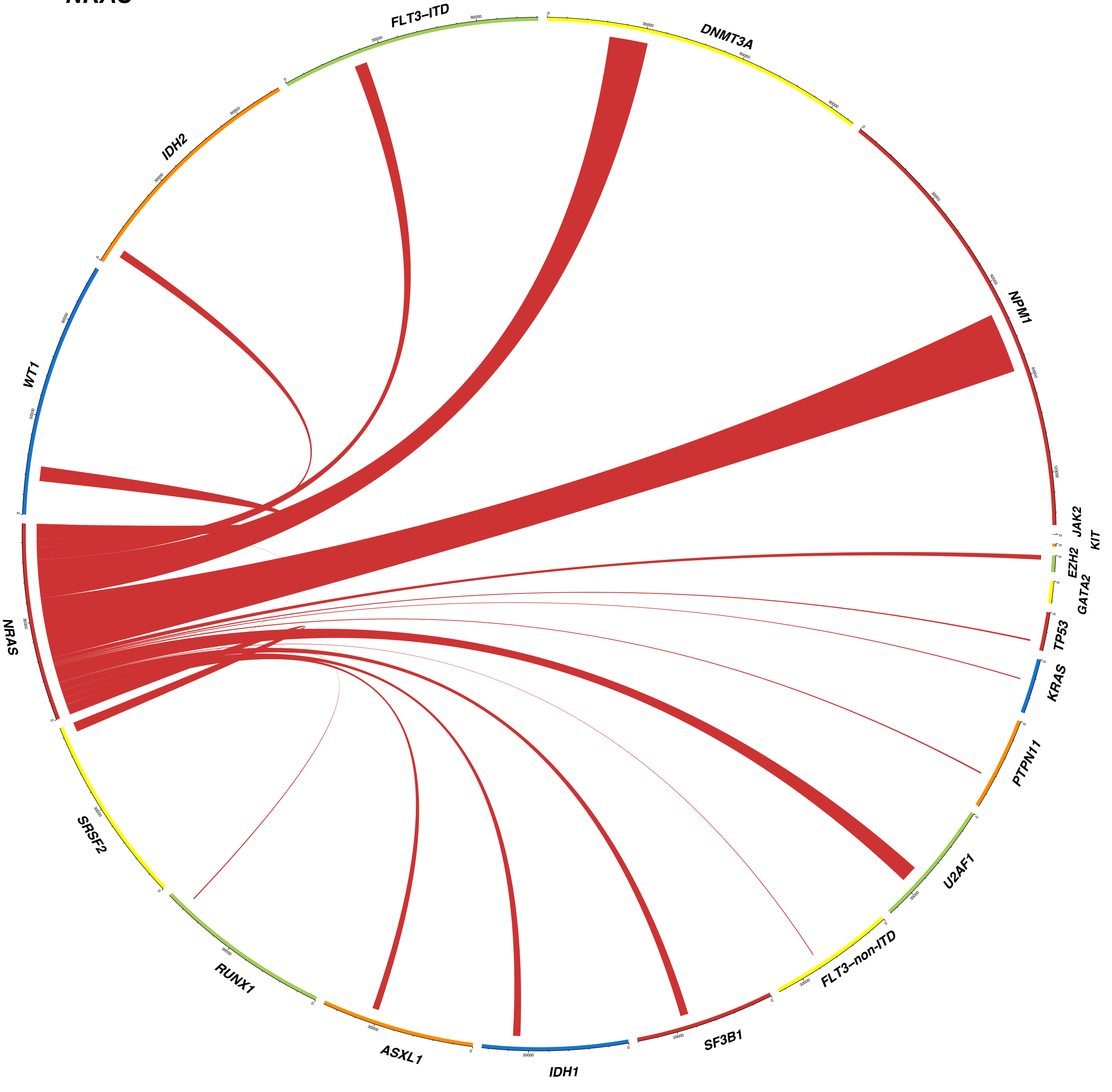
FLT3–ITD
DNMT3A
IDH2
NPM1
WT1
JAK2
KIT
EZH2
GATA2
NRAS
TP53
KRAS
PTPN11
SRSF2
U2AF1
RUNX1
FLT3–non-ITD
ASXL1
IDH1
SF3B1

PTPN11

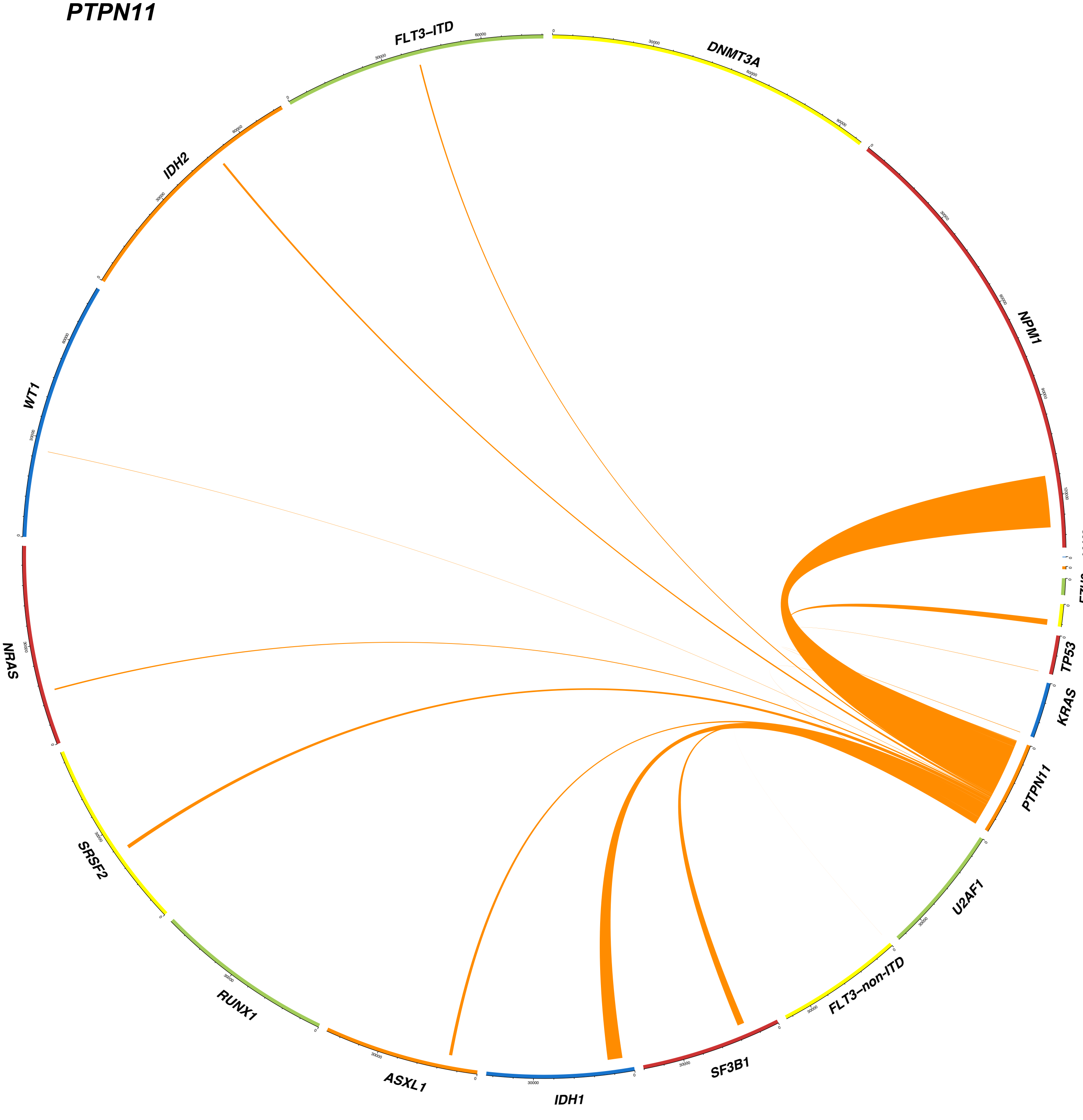
FLT3–ITD
DNMT3A
NPM1
JAK2
KIT
EZH2
GATA2
TP53
KRAS
PTPN11
U2AF1
FLT3–non-ITD
SF3B1
IDH1
ASXL1
RUNX1
SRSF2
NRAS
WT1
IDH2

### RUNX1

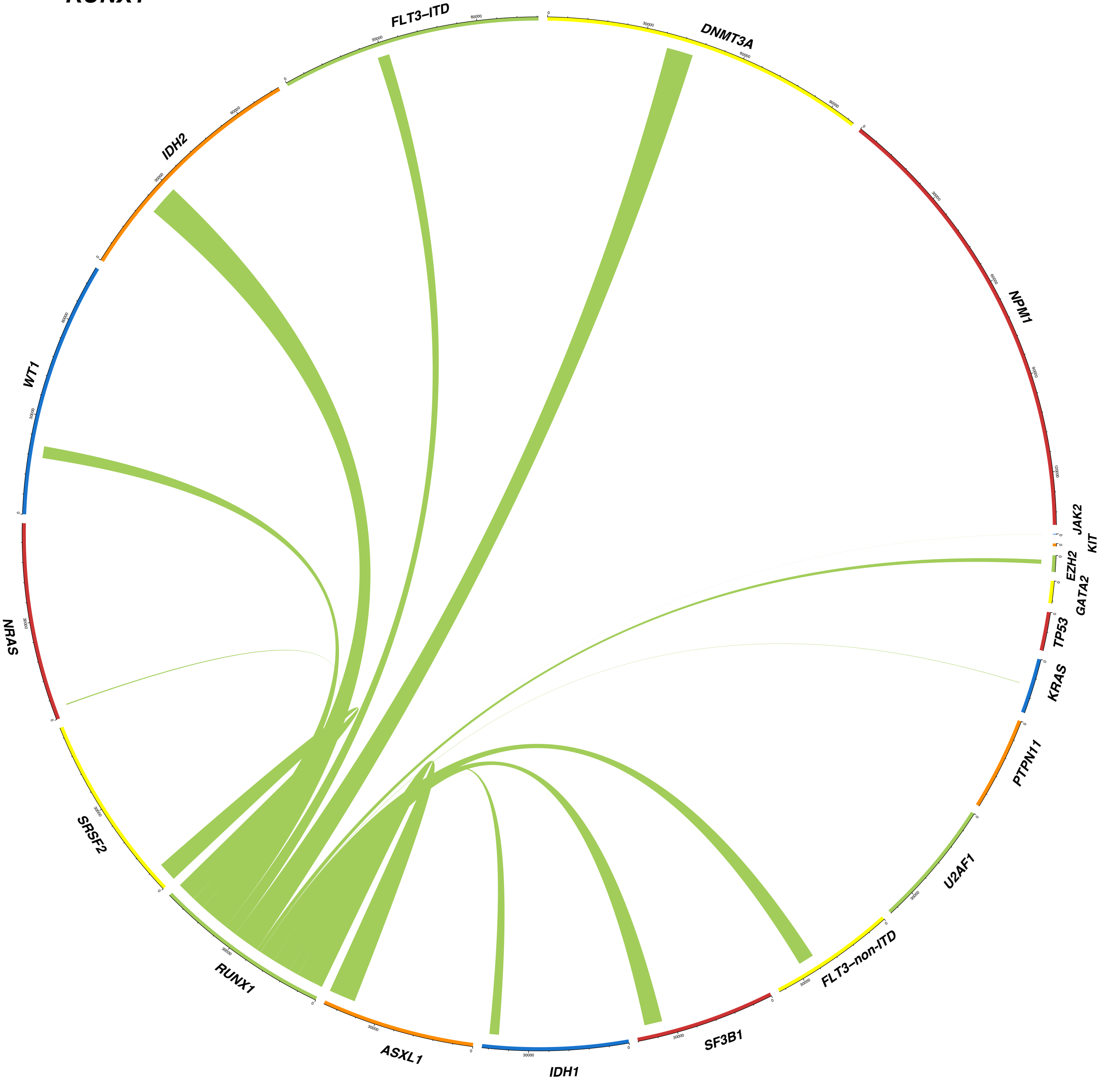

FLT3–ITD
DNMT3A
NPM1
JAK2
KIT
EZH2
GATA2
TP53
KRAS
PTPN11
U2AF1
FLT3–non-ITD
SF3B1
IDH1
ASXL1
RUNX1
SRSF2
NRAS
WT1
IDH2

## *SF3B1*

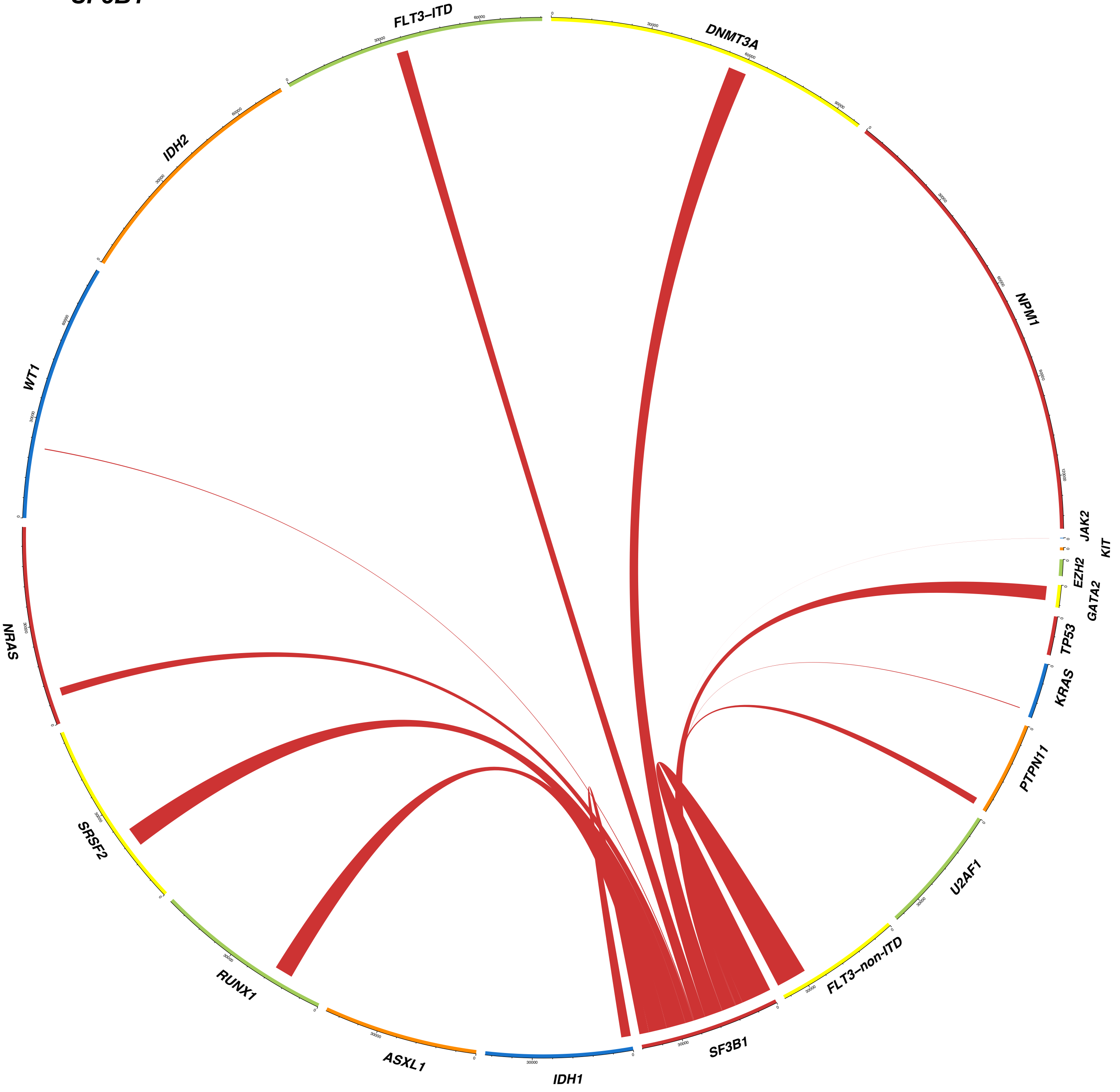
FLT3–ITD
DNMT3A
NPM1
JAK2
KIT
EZH2
GATA2
TP53
KRAS
PTPN11
U2AF1
FLT3–non-ITD
SF3B1
IDH1
ASXL1
RUNX1
SRSF2
NRAS
WT1
IDH2

#### SRSF2

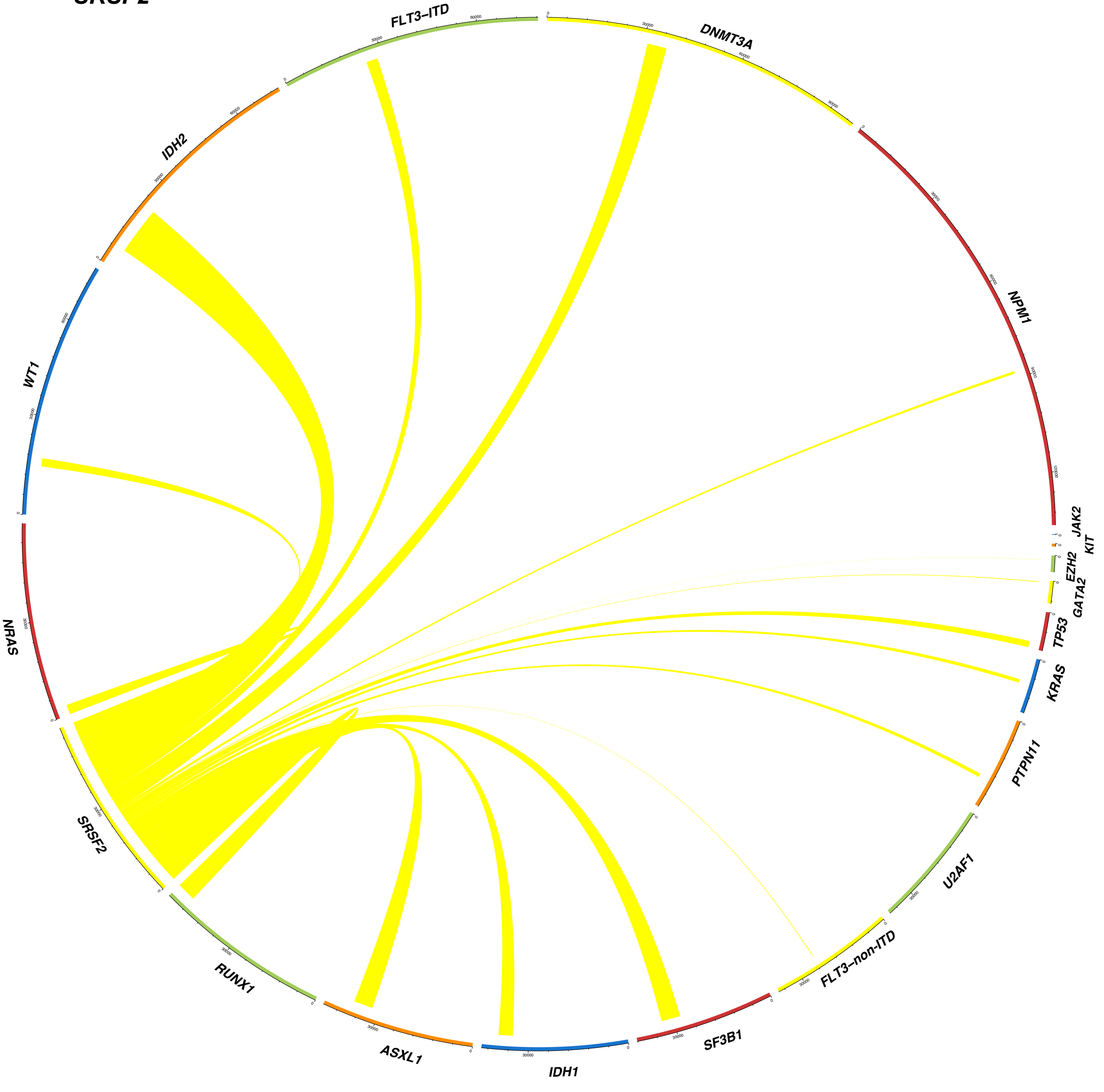
FLT3–ITD
DNMT3A
NPM1
JAK2
KIT
EZH2
GATA2
TP53
KRAS
PTPN11
U2AF1
FLT3–non-ITD
SF3B1
IDH1
ASXL1
RUNX1
SRSF2
NRAS
WT1
IDH2

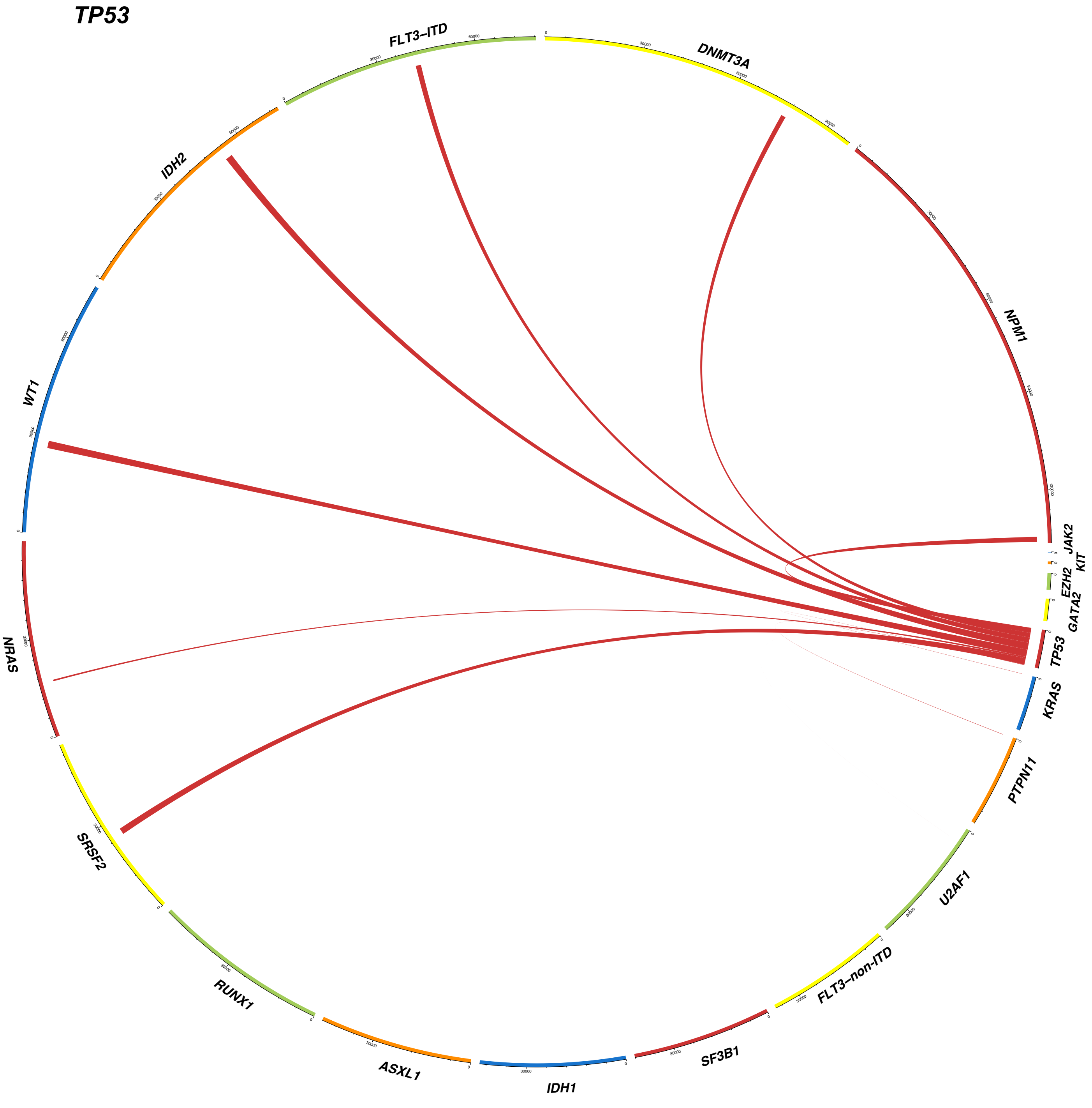
TP53
FLT3–ITD
DNMT3A
NPM1
JAK2
KIT
EZH2
GATA2
TP53
KRAS
PTPN11
U2AF1
FLT3–non-ITD
SF3B1
IDH1
ASXL1
RUNX1
SRSF2
NRAS
WT1
IDH2

## U2AF1

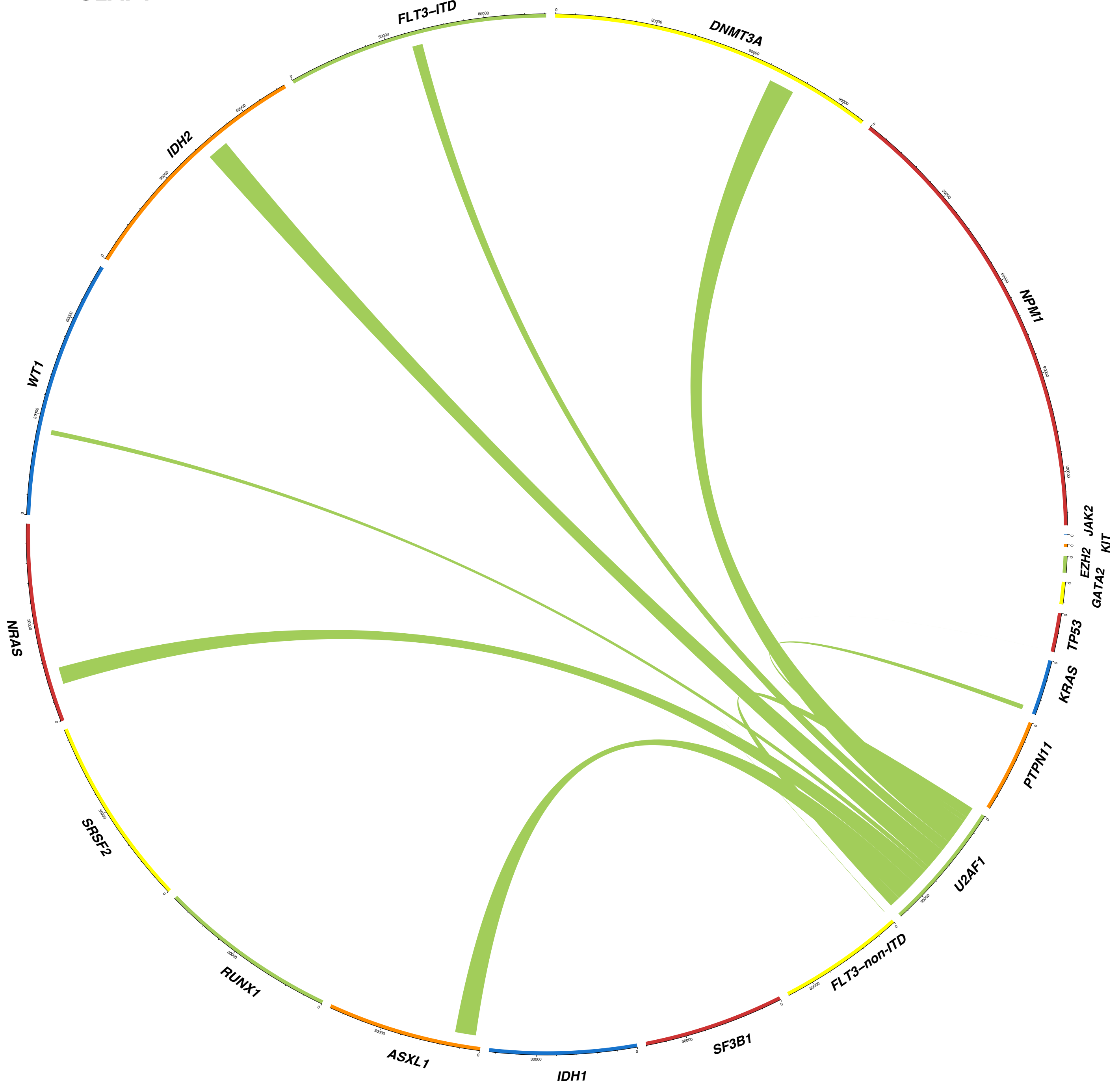

FLT3–ITD
DNMT3A
NPM1
JAK2
KIT
EZH2
GATA2
TP53
KRAS
PTPN11
U2AF1
FLT3–non-ITD
SF3B1
IDH1
ASXL1
RUNX1
SRSF2
NRAS
WT1
IDH2

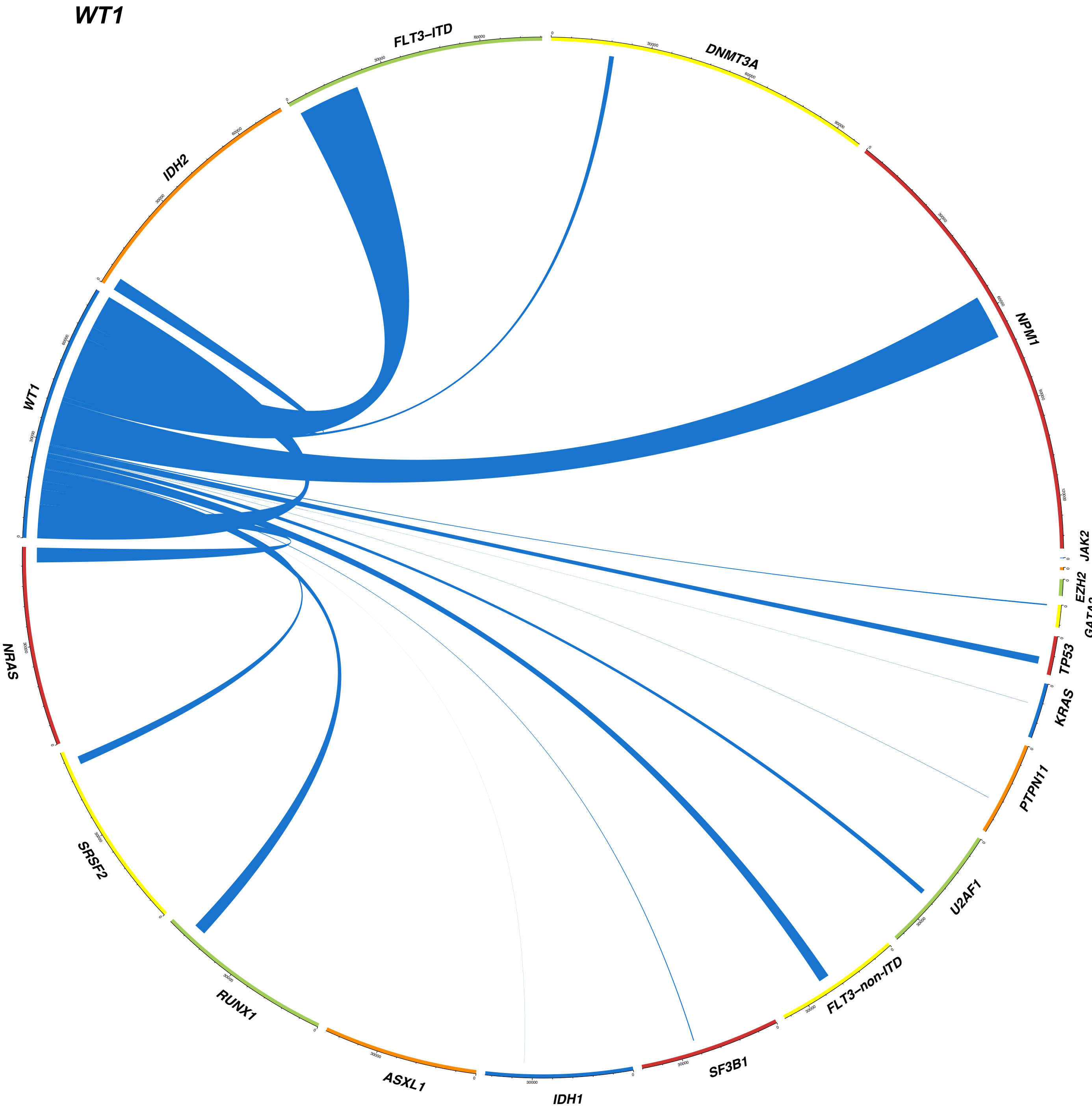
WT1
FLT3–ITD
DNMT3A
NPM1
JAK2
KIT
EZH2
GATA2
TP53
KRAS
PTPN11
U2AF1
FLT3–non-ITD
SF3B1
IDH1
ASXL1
RUNX1
SRSF2
NRAS
IDH2

**Extended Data Fig. 8. Circos plots showing patterns of mutation co-occurrence at single-cell resolution.** Patterns of mutation co-occurrence and mutual exclusivity based on the single cell genotype data are shown for each gene. When 2 mutations co-occurred in the same cell, a ribbon connects the genes. The width of each ribbon is proportional to the frequency of mutational events. *ASXL1* mutations frequently co-occurred with *IDH2, SRSF2, RUNX1,* and *U2AF1* mutations within the same cells, suggesting that these mutations have cooperative functions. Similarly, *DNMT3A* or *NPM1* mutations frequently co-occurred with mutations in almost all of the genes that were included in the single-cell DNA sequencing panel. *FLT3*-ITD frequently co-occurred with *NPM1, DNMT3A,* and *WT1* mutations and rarely co-occurred with *FLT3*-non-ITD*,* suggesting that *FLT3*-ITD and *FLT3*-non-ITD mutations are functionally redundant and therefore tend not to co-occur in the same cells*. IDH1* mutations frequently co-occurred with *NPM1* or *DNMT3A* mutations*,* but rarely with *IDH2* mutations. *IDH2* mutations frequently co-occurred with *NPM1, SRSF2, RUNX1*, and *ASXL1* mutations but rarely co-occurred with *IDH1* mutations, suggesting that *IDH1* and *IDH2* mutations share the same biological consequences and are not likely to co-occur in the same cellular populations. *NRAS, KRAS*, and *PTPN11* mutations rarely co-occur in the same cell, suggesting the functional redundancy of these mutations.

### Extended Data Fig. 9

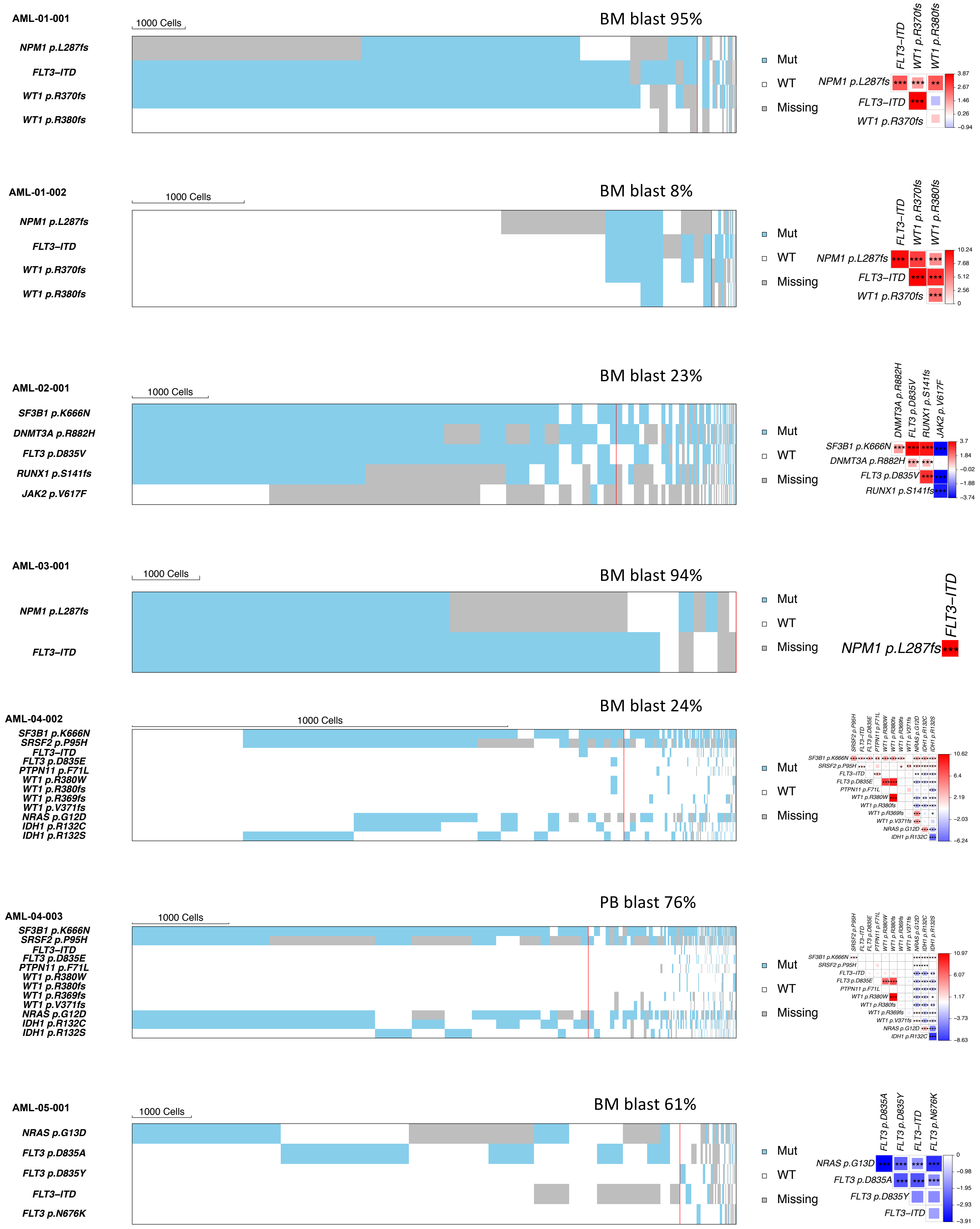

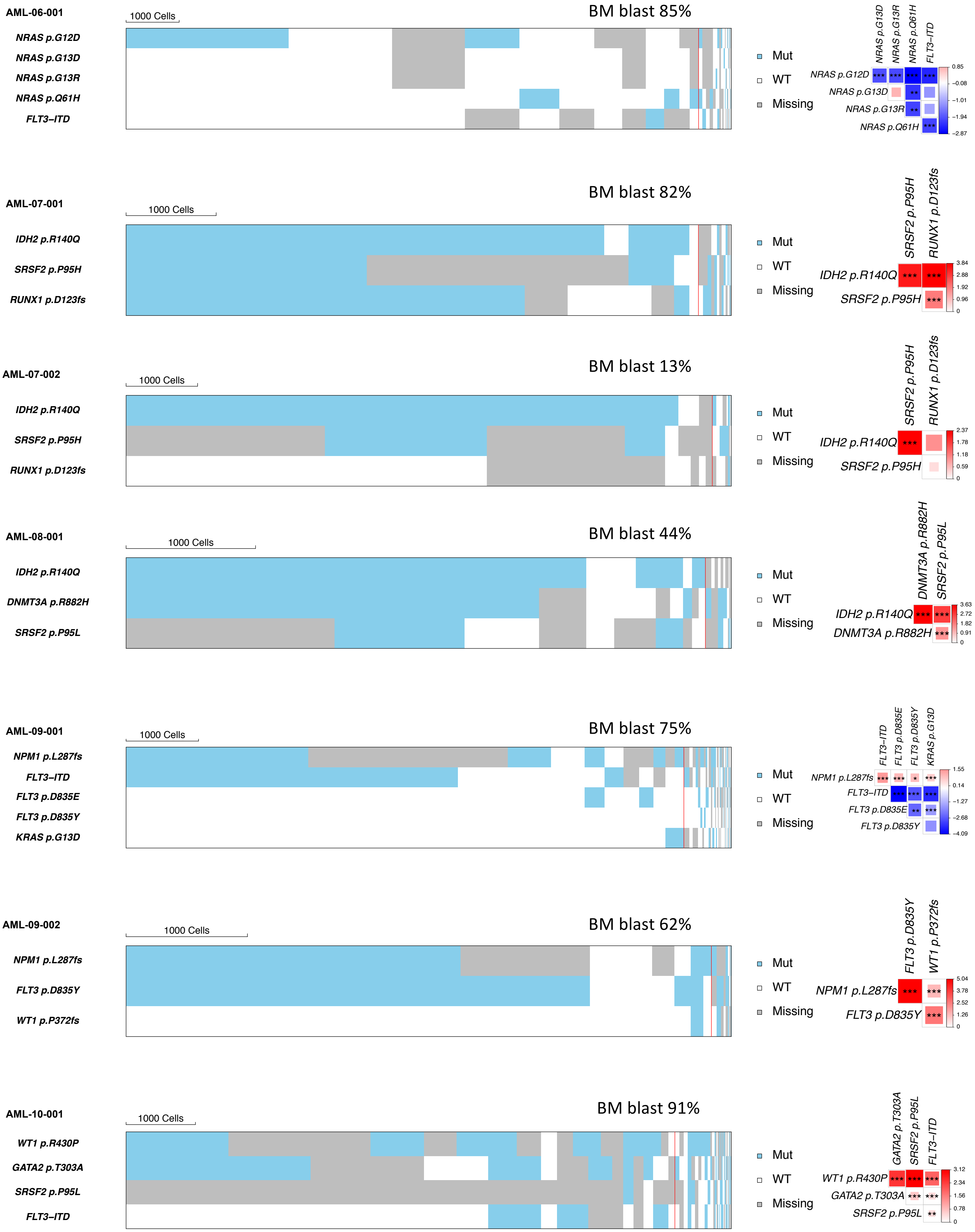

AML-06-001
1000 Cells
BM blast 85%
NRAS p.G12D
NRAS p.G13D
NRAS p.G13R
NRAS p.Q61H
FLT3–ITD
Mut
WT
Missing
AML-07-001
1000 Cells
BM blast 82%
IDH2 p.R140Q
SRSF2 p.P95H
RUNX1 p.D123fs
Mut
WT
Missing
AML-07-002
1000 Cells
BM blast 13%
IDH2 p.R140Q
SRSF2 p.P95H
RUNX1 p.D123fs
Mut
WT
Missing
AML-08-001
1000 Cells
BM blast 44%
IDH2 p.R140Q
DNMT3A p.R882H
SRSF2 p.P95L
Mut
WT
Missing
AML-09-001
1000 Cells
BM blast 75%
NPM1 p.L287fs
FLT3–ITD
FLT3 p.D835E
FLT3 p.D835Y
KRAS p.G13D
Mut
WT
Missing
AML-09-002
1000 Cells
BM blast 62%
NPM1 p.L287fs
FLT3 p.D835Y
WT1 p.P372fs
Mut
WT
Missing
AML-10-001
1000 Cells
BM blast 91%
WT1 p.R430P
GATA2 p.T303A
SRSF2 p.P95L
FLT3–ITD
Mut
WT
Missing

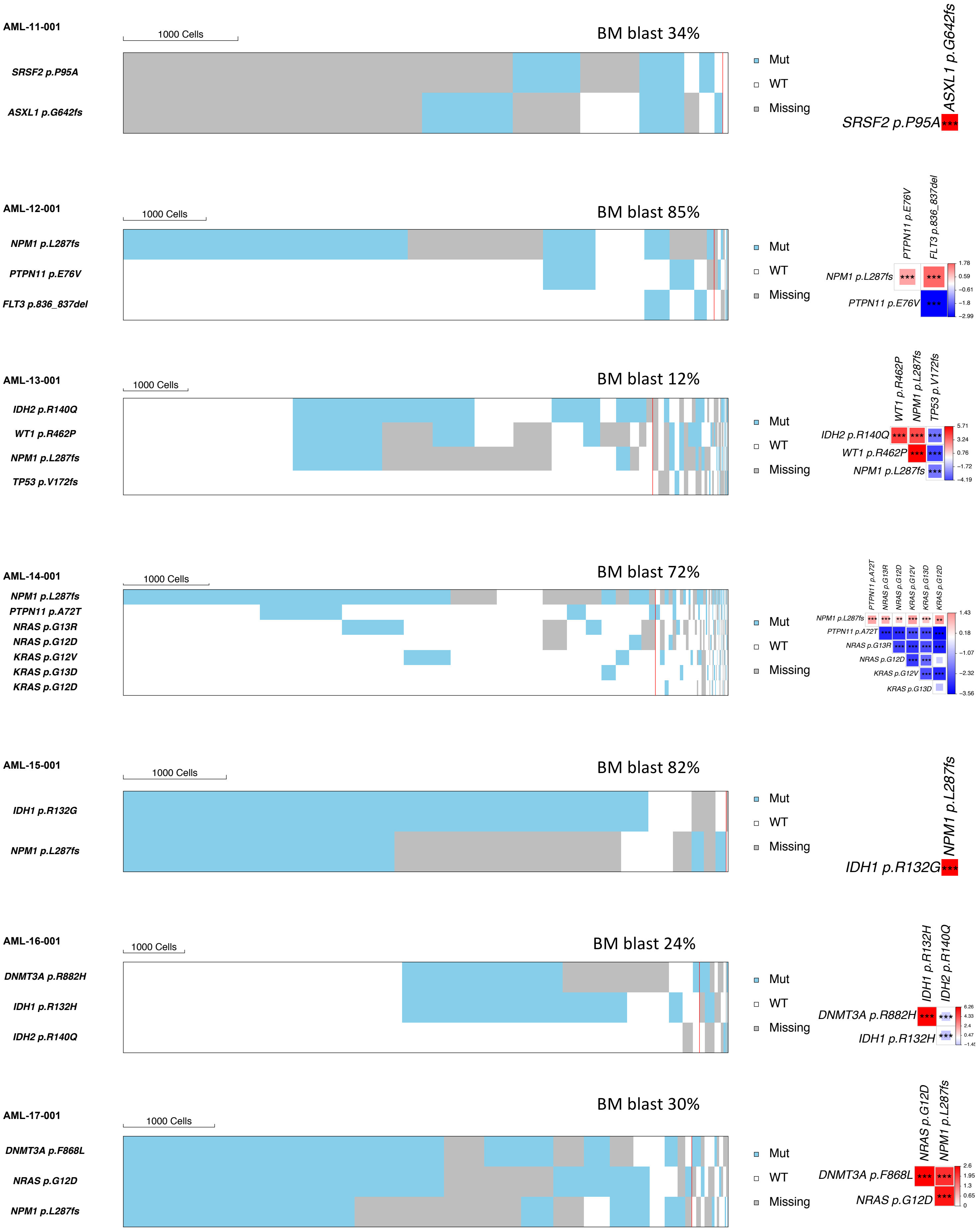

AML-11-001
1000 Cells
BM blast 34%
SRSF2 p.P95A
ASXL1 p.G642fs
Mut
WT
Missing
ASXL1 p.G642fs
SRSF2 p.P95A
***
AML-12-001
1000 Cells
BM blast 85%
NPM1 p.L287fs
PTPN11 p.E76V
FLT3 p.836_837del
Mut
WT
Missing
PTPN11 p.E76V
FLT3 p.836_837del
NPM1 p.L287fs
PTPN11 p.E76V
1.78
0.59
−0.61
−1.8
−2.99
AML-13-001
1000 Cells
BM blast 12%
IDH2 p.R140Q
WT1 p.R462P
NPM1 p.L287fs
TP53 p.V172fs
Mut
WT
Missing
WT1 p.R462P
NPM1 p.L287fs
TP53 p.V172fs
IDH2 p.R140Q
WT1 p.R462P
NPM1 p.L287fs
5.71
3.24
0.76
−1.72
−4.19
AML-14-001
1000 Cells
BM blast 72%
NPM1 p.L287fs
PTPN11 p.A72T
NRAS p.G13R
NRAS p.G12D
KRAS p.G12V
KRAS p.G13D
KRAS p.G12D
Mut
WT
Missing
PTPN11 p.A72T
NRAS p.G13R
NRAS p.G12D
KRAS p.G12V
KRAS p.G13D
KRAS p.G12D
NPM1 p.L287fs
PTPN11 p.A72T
NRAS p.G13R
NRAS p.G12D
KRAS p.G12V
KRAS p.G13D
1.43
0.18
−1.07
−2.32
−3.56
AML-15-001
1000 Cells
BM blast 82%
IDH1 p.R132G
NPM1 p.L287fs
Mut
WT
Missing
NPM1 p.L287fs
IDH1 p.R132G
AML-16-001
1000 Cells
BM blast 24%
DNMT3A p.R882H
IDH1 p.R132H
IDH2 p.R140Q
Mut
WT
Missing
IDH1 p.R132H
IDH2 p.R140Q
DNMT3A p.R882H
IDH1 p.R132H
6.26
4.33
2.4
0.47
AML-17-001
1000 Cells
BM blast 30%
DNMT3A p.F868L
NRAS p.G12D
NPM1 p.L287fs
Mut
WT
Missing
NRAS p.G12D
NPM1 p.L287fs
DNMT3A p.F868L
NRAS p.G12D
2.6
1.95
1.3
0.65
0

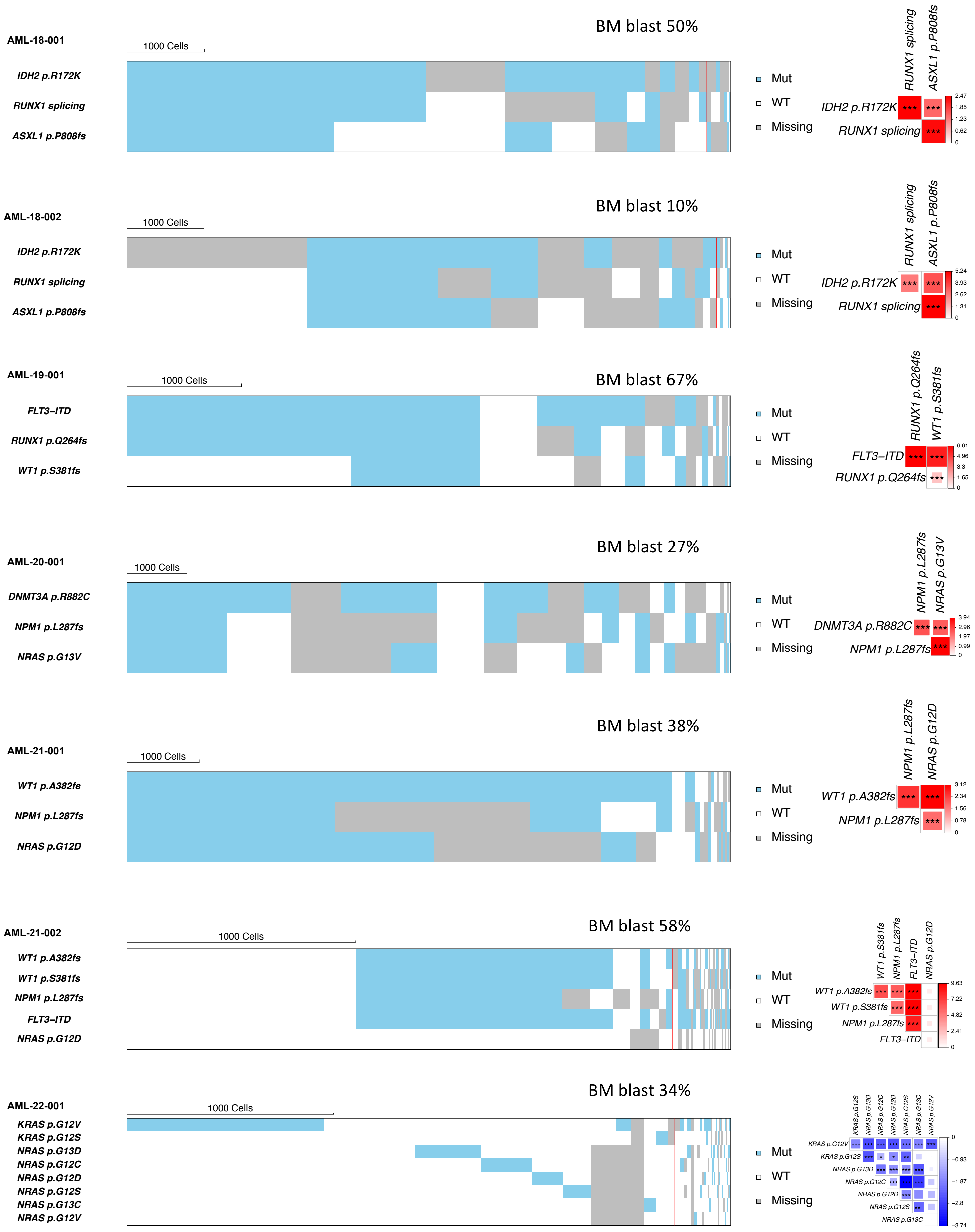

AML-18-001
BM blast 50%
1000 Cells
IDH2 p.R172K
RUNX1 splicing
ASXL1 p.P808fs
Mut
WT
Missing
AML-18-002
BM blast 10%
1000 Cells
IDH2 p.R172K
RUNX1 splicing
ASXL1 p.P808fs
AML-19-001
BM blast 67%
1000 Cells
FLT3–ITD
RUNX1 p.Q264fs
WT1 p.S381fs
AML-20-001
BM blast 27%
1000 Cells
DNMT3A p.R882C
NPM1 p.L287fs
NRAS p.G13V
AML-21-001
BM blast 38%
1000 Cells
WT1 p.A382fs
NPM1 p.L287fs
NRAS p.G12D
AML-21-002
BM blast 58%
1000 Cells
WT1 p.A382fs
WT1 p.S381fs
NPM1 p.L287fs
FLT3–ITD
NRAS p.G12D
AML-22-001
BM blast 34%
1000 Cells
KRAS p.G12V
KRAS p.G12S
NRAS p.G13D
NRAS p.G12C
NRAS p.G12D
NRAS p.G12S
NRAS p.G13C
NRAS p.G12V

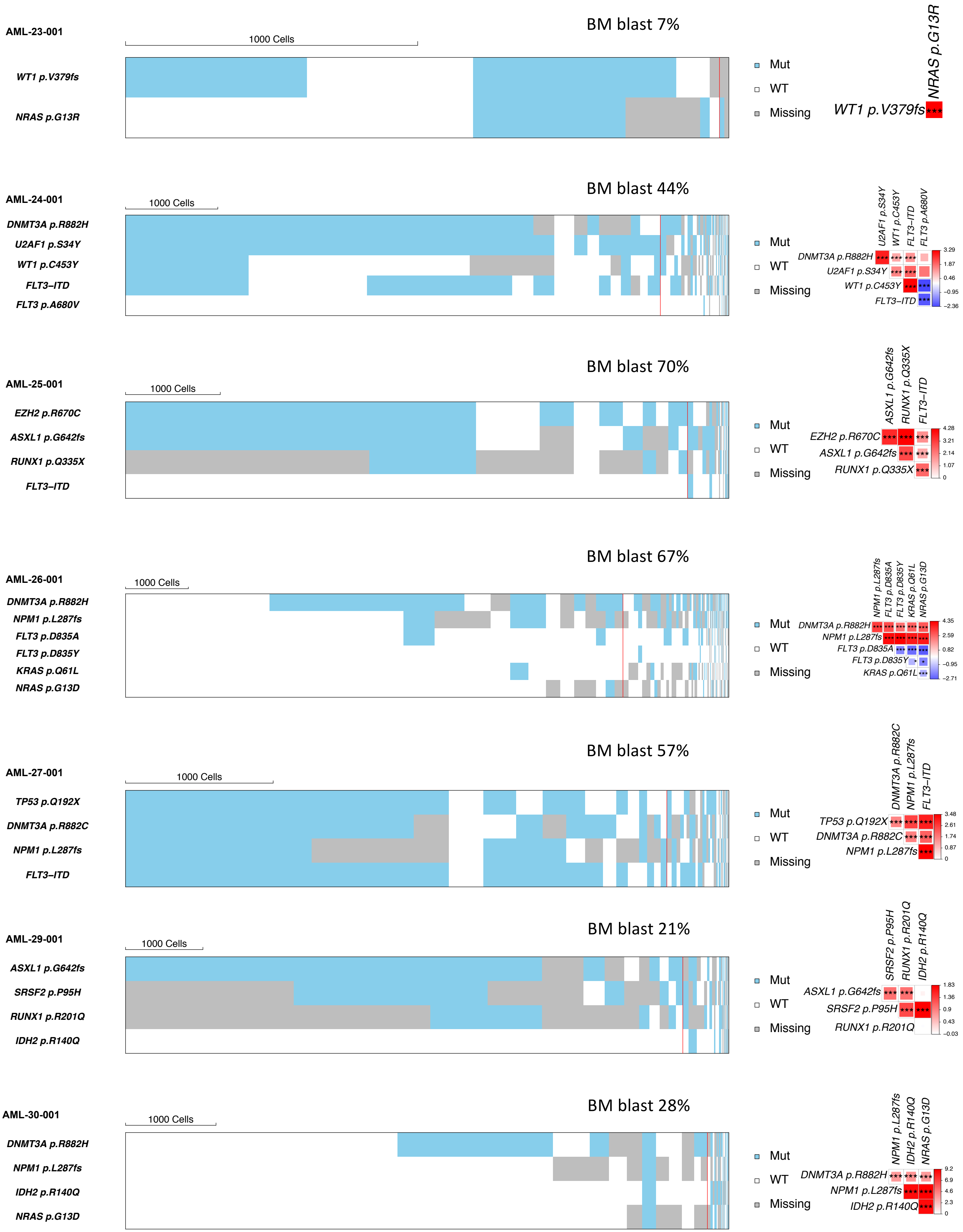

AML-23-001
BM blast 7%
1000 Cells
WT1 p.V379fs
NRAS p.G13R
Mut
WT
Missing
NRAS p.G13R
WT1 p.V379fs ***
AML-24-001
BM blast 44%
1000 Cells
DNMT3A p.R882H
U2AF1 p.S34Y
WT1 p.C453Y
FLT3–ITD
FLT3 p.A680V
Mut
WT
Missing
3.29
1.87
0.46
−0.95
−2.36
AML-25-001
BM blast 70%
1000 Cells
EZH2 p.R670C
ASXL1 p.G642fs
RUNX1 p.Q335X
FLT3–ITD
Mut
WT
Missing
4.28
3.21
2.14
1.07
0
AML-26-001
BM blast 67%
1000 Cells
DNMT3A p.R882H
NPM1 p.L287fs
FLT3 p.D835A
FLT3 p.D835Y
KRAS p.Q61L
NRAS p.G13D
Mut
WT
Missing
4.35
2.59
0.82
−0.95
−2.71
AML-27-001
BM blast 57%
1000 Cells
TP53 p.Q192X
DNMT3A p.R882C
NPM1 p.L287fs
FLT3–ITD
Mut
WT
Missing
3.48
2.61
1.74
0.87
0
AML-29-001
BM blast 21%
1000 Cells
ASXL1 p.G642fs
SRSF2 p.P95H
RUNX1 p.R201Q
IDH2 p.R140Q
Mut
WT
Missing
1.83
1.36
0.9
0.43
−0.03
AML-30-001
BM blast 28%
1000 Cells
DNMT3A p.R882H
NPM1 p.L287fs
IDH2 p.R140Q
NRAS p.G13D
Mut
WT
Missing
9.2
6.9
4.6
2.3
0

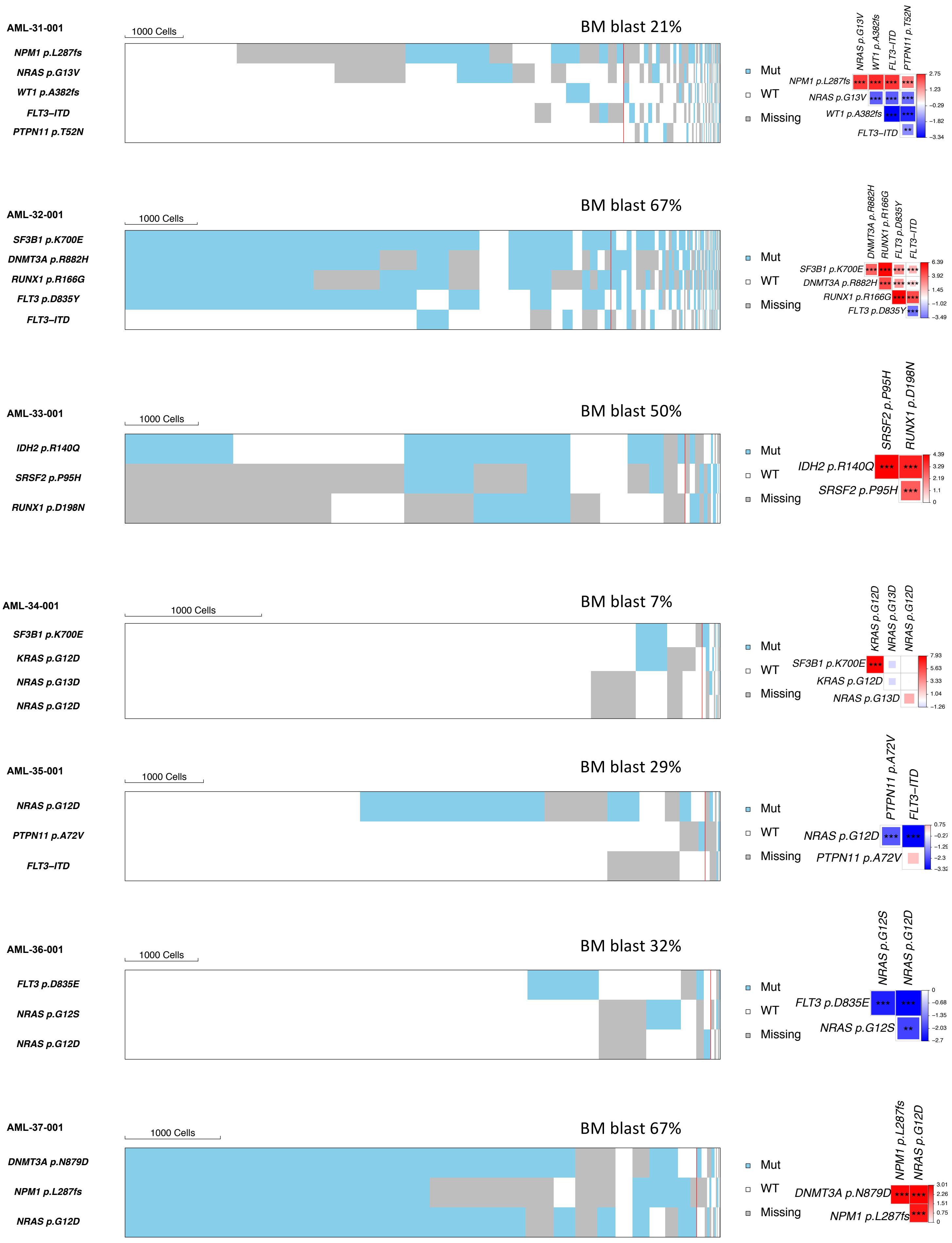

AML-31-001
1000 Cells
BM blast 21%
NPM1 p.L287fs
NRAS p.G13V
WT1 p.A382fs
FLT3–ITD
PTPN11 p.T52N
Mut
WT
Missing
AML-32-001
1000 Cells
BM blast 67%
SF3B1 p.K700E
DNMT3A p.R882H
RUNX1 p.R166G
FLT3 p.D835Y
FLT3–ITD
Mut
WT
Missing
AML-33-001
1000 Cells
BM blast 50%
IDH2 p.R140Q
SRSF2 p.P95H
RUNX1 p.D198N
Mut
WT
Missing
AML-34-001
1000 Cells
BM blast 7%
SF3B1 p.K700E
KRAS p.G12D
NRAS p.G13D
NRAS p.G12D
Mut
WT
Missing
AML-35-001
1000 Cells
BM blast 29%
NRAS p.G12D
PTPN11 p.A72V
FLT3–ITD
Mut
WT
Missing
AML-36-001
1000 Cells
BM blast 32%
FLT3 p.D835E
NRAS p.G12S
NRAS p.G12D
Mut
WT
Missing
AML-37-001
1000 Cells
BM blast 67%
DNMT3A p.N879D
NPM1 p.L287fs
NRAS p.G12D
Mut
WT
Missing

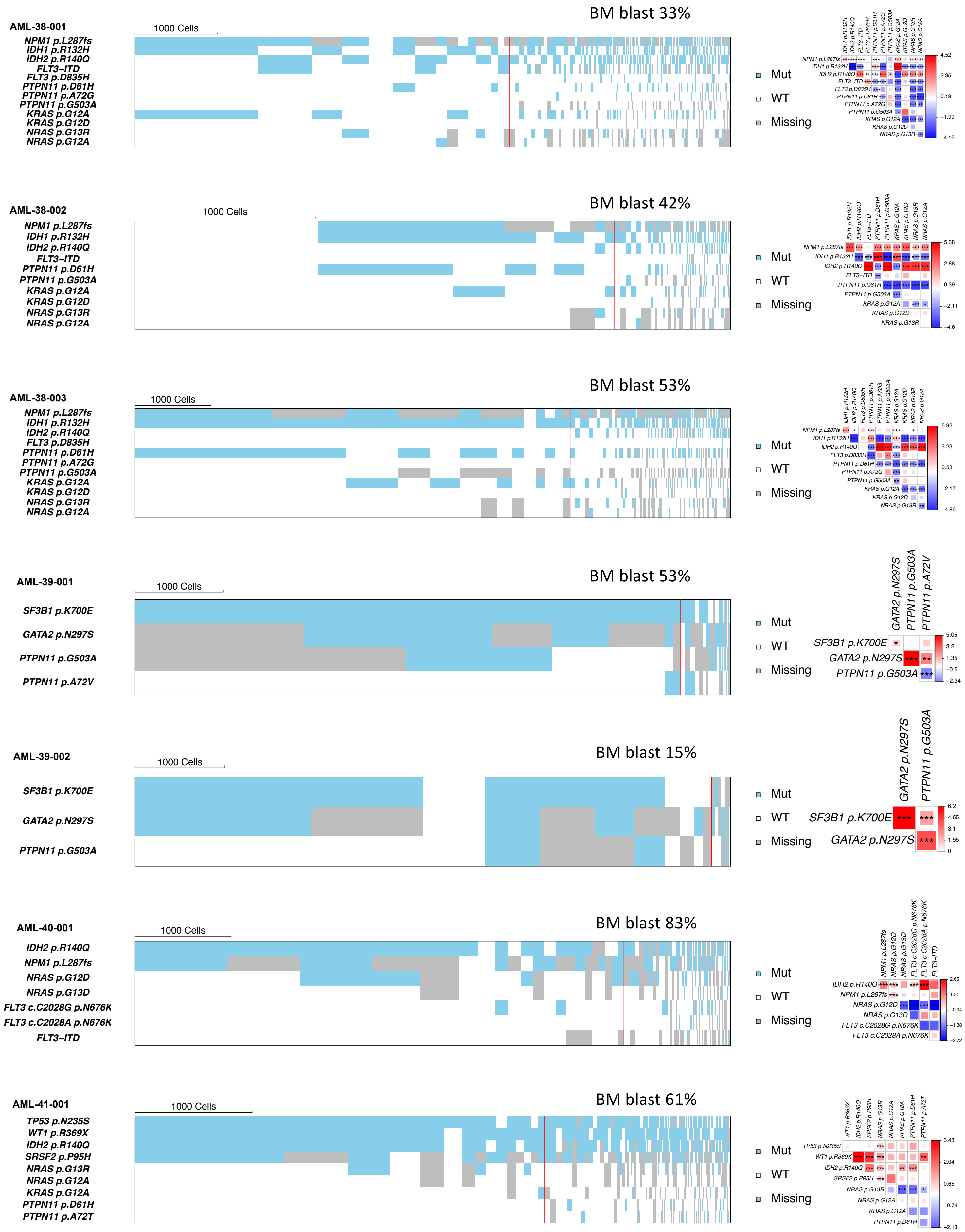
AML-38-001
1000 Cells
BM blast 33%
NPM1 p.L287fs
IDH1 p.R132H
IDH2 p.R140Q
FLT3–ITD
FLT3 p.D835H
PTPN11 p.D61H
PTPN11 p.A72G
PTPN11 p.G503A
KRAS p.G12A
KRAS p.G12D
NRAS p.G13R
NRAS p.G12A
Mut
WT
Missing
AML-38-002
1000 Cells
BM blast 42%
NPM1 p.L287fs
IDH1 p.R132H
IDH2 p.R140Q
FLT3–ITD
PTPN11 p.D61H
PTPN11 p.G503A
KRAS p.G12A
KRAS p.G12D
NRAS p.G13R
NRAS p.G12A
Mut
WT
Missing
AML-38-003
1000 Cells
BM blast 53%
NPM1 p.L287fs
IDH1 p.R132H
IDH2 p.R140Q
FLT3 p.D835H
PTPN11 p.D61H
PTPN11 p.A72G
PTPN11 p.G503A
KRAS p.G12A
KRAS p.G12D
NRAS p.G13R
NRAS p.G12A
Mut
WT
Missing
AML-39-001
1000 Cells
BM blast 53%
SF3B1 p.K700E
GATA2 p.N297S
PTPN11 p.G503A
PTPN11 p.A72V
Mut
WT
Missing
AML-39-002
1000 Cells
BM blast 15%
SF3B1 p.K700E
GATA2 p.N297S
PTPN11 p.G503A
Mut
WT
Missing
AML-40-001
1000 Cells
BM blast 83%
IDH2 p.R140Q
NPM1 p.L287fs
NRAS p.G12D
NRAS p.G13D
FLT3 c.C2028G p.N676K
FLT3 c.C2028A p.N676K
FLT3–ITD
Mut
WT
Missing
AML-41-001
1000 Cells
BM blast 61%
TP53 p.N235S
WT1 p.R369X
IDH2 p.R140Q
SRSF2 p.P95H
NRAS p.G13R
NRAS p.G12A
KRAS p.G12A
PTPN11 p.D61H
PTPN11 p.A72T
Mut
WT
Missing

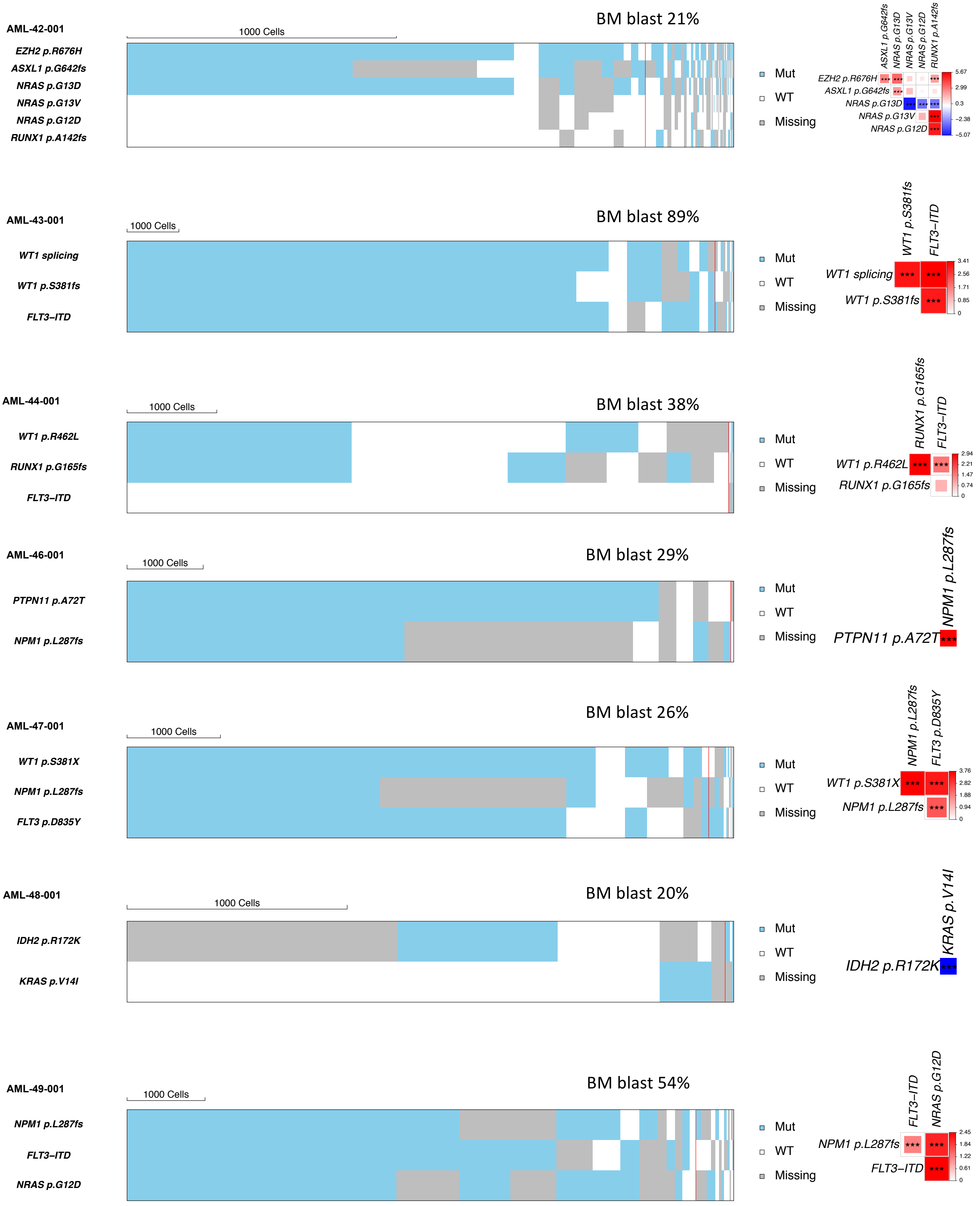
AML-42-001
1000 Cells
BM blast 21%
EZH2 p.R676H
ASXL1 p.G642fs
NRAS p.G13D
NRAS p.G13V
NRAS p.G12D
RUNX1 p.A142fs
Mut
WT
Missing
AML-43-001
1000 Cells
BM blast 89%
WT1 splicing
WT1 p.S381fs
FLT3–ITD
AML-44-001
1000 Cells
BM blast 38%
WT1 p.R462L
RUNX1 p.G165fs
FLT3–ITD
AML-46-001
1000 Cells
BM blast 29%
PTPN11 p.A72T
NPM1 p.L287fs
AML-47-001
1000 Cells
BM blast 26%
WT1 p.S381X
NPM1 p.L287fs
FLT3 p.D835Y
AML-48-001
1000 Cells
BM blast 20%
IDH2 p.R172K
KRAS p.V14I
AML-49-001
1000 Cells
BM blast 54%
NPM1 p.L287fs
FLT3–ITD
NRAS p.G12D

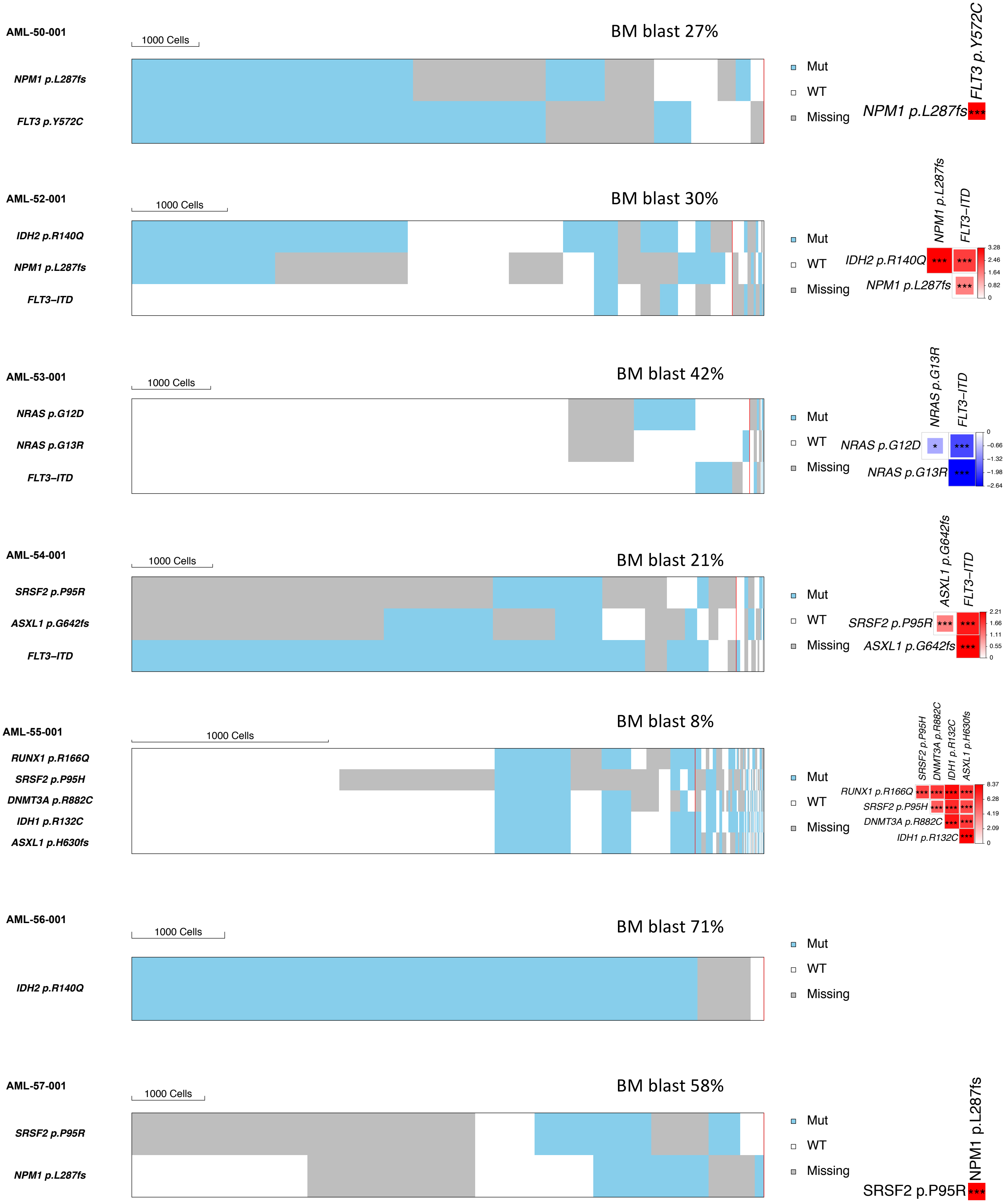
AML-50-001
1000 Cells
BM blast 27%
NPM1 p.L287fs
FLT3 p.Y572C
Mut
WT
Missing
FLT3 p.Y572C
NPM1 p.L287fs
***
AML-52-001
1000 Cells
BM blast 30%
IDH2 p.R140Q
NPM1 p.L287fs
FLT3–ITD
Mut
WT
Missing
NPM1 p.L287fs
FLT3–ITD
IDH2 p.R140Q
NPM1 p.L287fs
3.28
2.46
1.64
0.82
0
AML-53-001
1000 Cells
BM blast 42%
NRAS p.G12D
NRAS p.G13R
FLT3–ITD
Mut
WT
Missing
NRAS p.G13R
FLT3–ITD
NRAS p.G12D
NRAS p.G13R
0
−0.66
−1.32
−1.98
−2.64
AML-54-001
1000 Cells
BM blast 21%
SRSF2 p.P95R
ASXL1 p.G642fs
FLT3–ITD
Mut
WT
Missing
ASXL1 p.G642fs
FLT3–ITD
SRSF2 p.P95R
ASXL1 p.G642fs
2.21
1.66
1.11
0.55
0
AML-55-001
1000 Cells
BM blast 8%
RUNX1 p.R166Q
SRSF2 p.P95H
DNMT3A p.R882C
IDH1 p.R132C
ASXL1 p.H630fs
Mut
WT
Missing
SRSF2 p.P95H
DNMT3A p.R882C
IDH1 p.R132C
ASXL1 p.H630fs
RUNX1 p.R166Q
SRSF2 p.P95H
DNMT3A p.R882C
IDH1 p.R132C
8.37
6.28
4.19
2.09
0
AML-56-001
1000 Cells
BM blast 71%
IDH2 p.R140Q
Mut
WT
Missing
AML-57-001
1000 Cells
BM blast 58%
SRSF2 p.P95R
NPM1 p.L287fs
Mut
WT
Missing
NPM1 p.L287fs
SRSF2 p.P95R
***

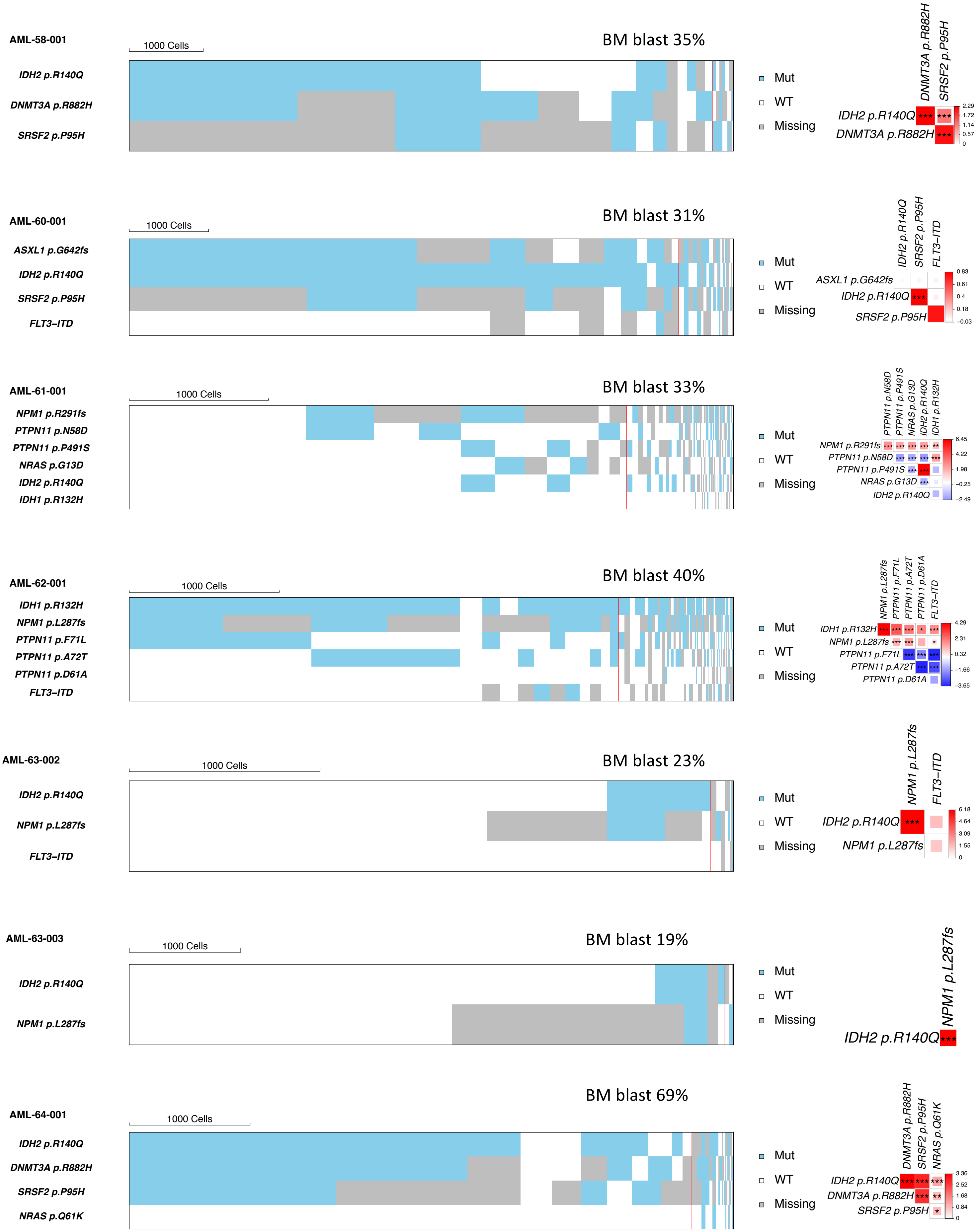
AML-58-001
1000 Cells
BM blast 35%
IDH2 p.R140Q
DNMT3A p.R882H
SRSF2 p.P95H
Mut
WT
Missing
AML-60-001
1000 Cells
BM blast 31%
ASXL1 p.G642fs
IDH2 p.R140Q
SRSF2 p.P95H
FLT3–ITD
AML-61-001
1000 Cells
BM blast 33%
NPM1 p.R291fs
PTPN11 p.N58D
PTPN11 p.P491S
NRAS p.G13D
IDH2 p.R140Q
IDH1 p.R132H
AML-62-001
1000 Cells
BM blast 40%
IDH1 p.R132H
NPM1 p.L287fs
PTPN11 p.F71L
PTPN11 p.A72T
PTPN11 p.D61A
FLT3–ITD
AML-63-002
1000 Cells
BM blast 23%
IDH2 p.R140Q
NPM1 p.L287fs
FLT3–ITD
AML-63-003
1000 Cells
BM blast 19%
IDH2 p.R140Q
NPM1 p.L287fs
AML-64-001
1000 Cells
BM blast 69%
IDH2 p.R140Q
DNMT3A p.R882H
SRSF2 p.P95H
NRAS p.Q61K

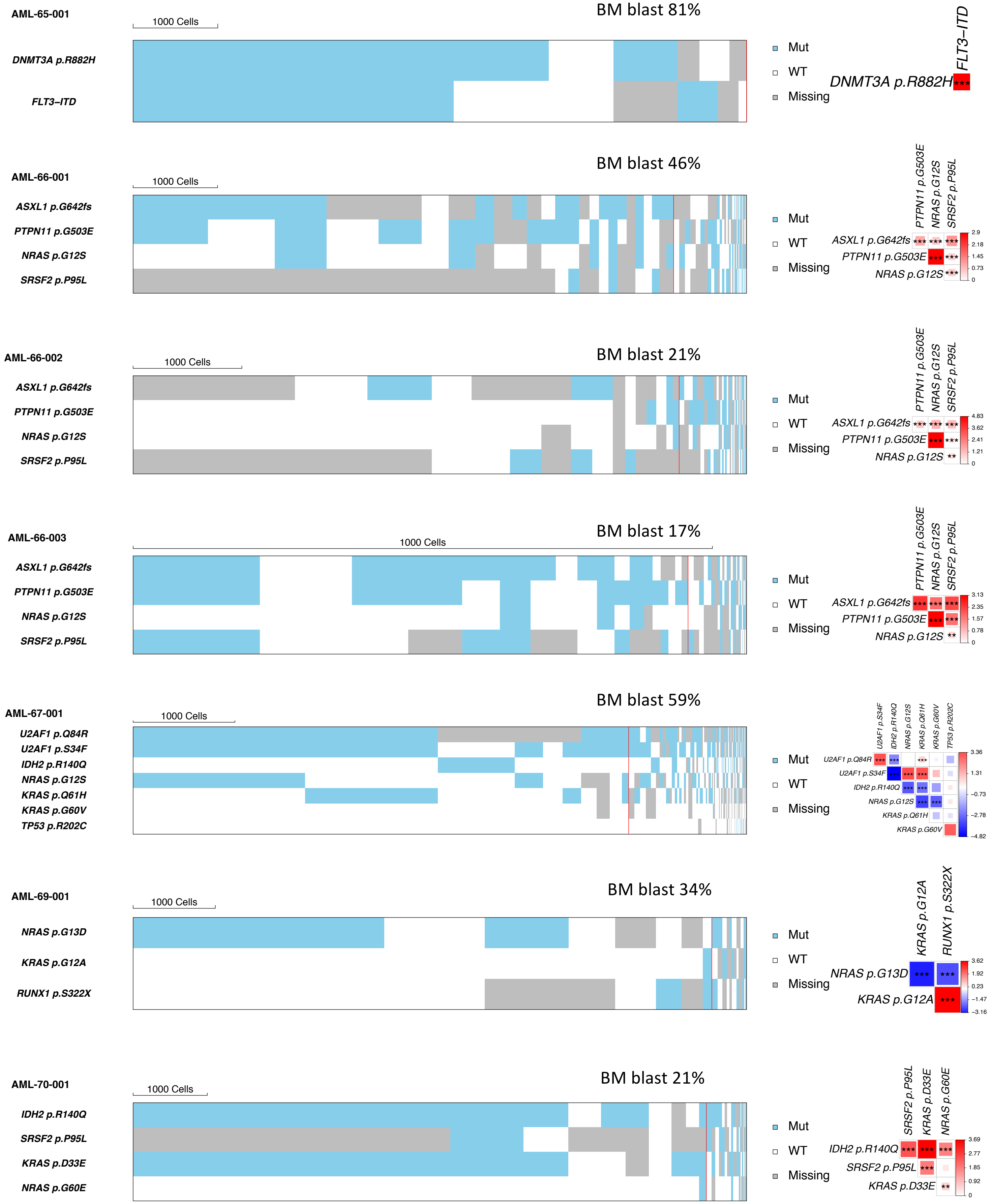
AML-65-001
BM blast 81%
1000 Cells
DNMT3A p.R882H
FLT3–ITD
Mut
WT
Missing
AML-66-001
BM blast 46%
ASXL1 p.G642fs
PTPN11 p.G503E
NRAS p.G12S
SRSF2 p.P95L
AML-66-002
BM blast 21%
AML-66-003
BM blast 17%
AML-67-001
BM blast 59%
U2AF1 p.Q84R
U2AF1 p.S34F
IDH2 p.R140Q
KRAS p.Q61H
KRAS p.G60V
TP53 p.R202C
AML-69-001
BM blast 34%
NRAS p.G13D
KRAS p.G12A
RUNX1 p.S322X
AML-70-001
BM blast 21%
KRAS p.D33E
NRAS p.G60E

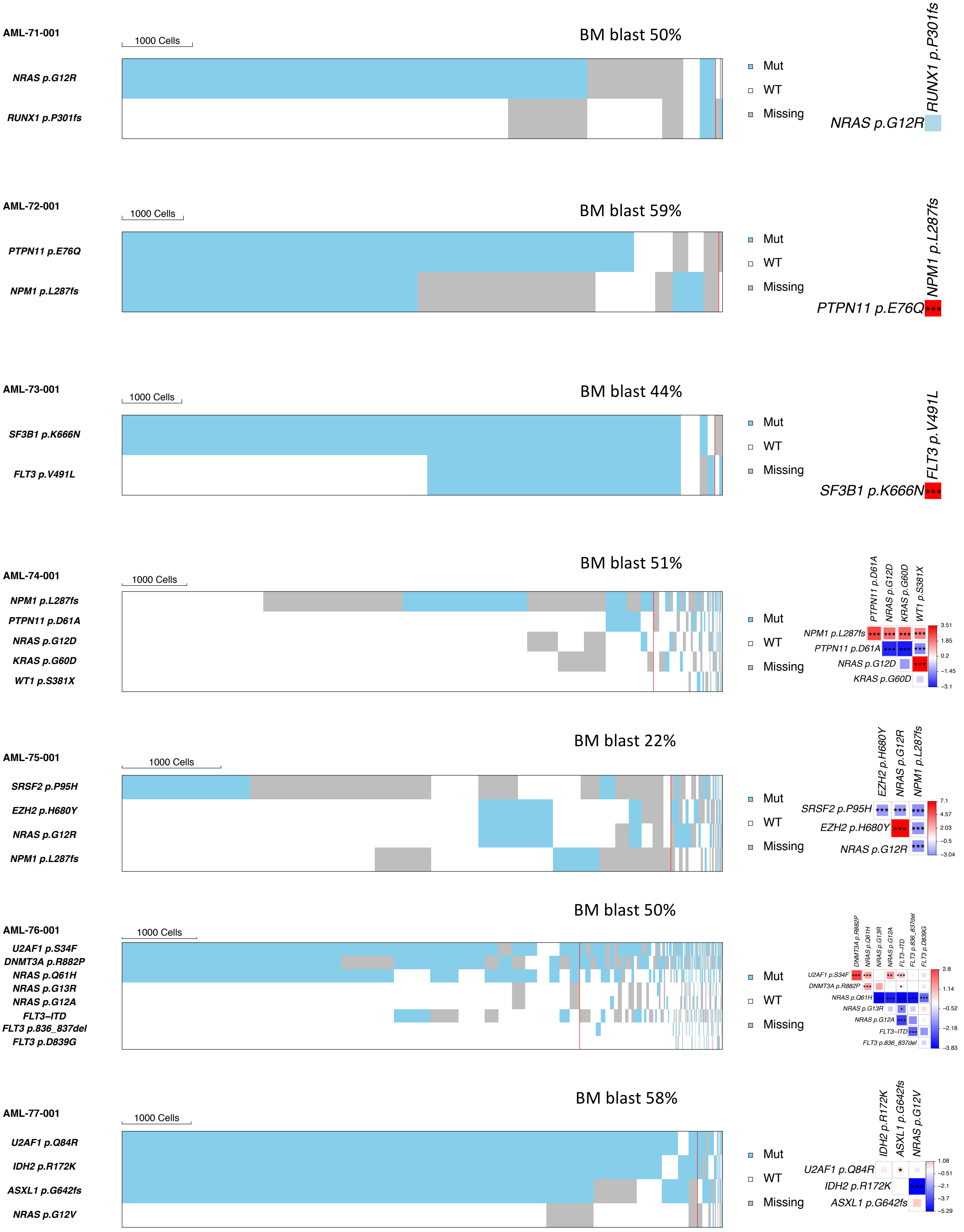
AML-71-001
1000 Cells
BM blast 50%
NRAS p.G12R
RUNX1 p.P301fs
Mut
WT
Missing
RUNX1 p.P301fs
NRAS p.G12R
AML-72-001
1000 Cells
BM blast 59%
PTPN11 p.E76Q
NPM1 p.L287fs
Mut
WT
Missing
NPM1 p.L287fs
PTPN11 p.E76Q
***
AML-73-001
1000 Cells
BM blast 44%
SF3B1 p.K666N
FLT3 p.V491L
Mut
WT
Missing
FLT3 p.V491L
SF3B1 p.K666N
***
AML-74-001
1000 Cells
BM blast 51%
NPM1 p.L287fs
PTPN11 p.D61A
NRAS p.G12D
KRAS p.G60D
WT1 p.S381X
Mut
WT
Missing
PTPN11 p.D61A
NRAS p.G12D
KRAS p.G60D
WT1 p.S381X
NPM1 p.L287fs
PTPN11 p.D61A
NRAS p.G12D
KRAS p.G60D
3.51
1.85
0.2
−1.45
−3.1
AML-75-001
1000 Cells
BM blast 22%
SRSF2 p.P95H
EZH2 p.H680Y
NRAS p.G12R
NPM1 p.L287fs
Mut
WT
Missing
EZH2 p.H680Y
NRAS p.G12R
NPM1 p.L287fs
SRSF2 p.P95H
EZH2 p.H680Y
NRAS p.G12R
7.1
4.57
2.03
−0.5
−3.04
AML-76-001
1000 Cells
BM blast 50%
U2AF1 p.S34F
DNMT3A p.R882P
NRAS p.Q61H
NRAS p.G13R
NRAS p.G12A
FLT3–ITD
FLT3 p.836_837del
FLT3 p.D839G
Mut
WT
Missing
DNMT3A p.R882P
NRAS p.Q61H
NRAS p.G13R
NRAS p.G12A
FLT3–ITD
FLT3 p.836_837del
FLT3 p.D839G
U2AF1 p.S34F
DNMT3A p.R882P
NRAS p.Q61H
NRAS p.G13R
NRAS p.G12A
FLT3–ITD
FLT3 p.836_837del
2.8
1.14
−0.52
−2.18
−3.83
AML-77-001
1000 Cells
BM blast 58%
U2AF1 p.Q84R
IDH2 p.R172K
ASXL1 p.G642fs
NRAS p.G12V
Mut
WT
Missing
IDH2 p.R172K
ASXL1 p.G642fs
NRAS p.G12V
U2AF1 p.Q84R
IDH2 p.R172K
ASXL1 p.G642fs
1.08
−0.51
−2.1
−3.7
−5.29

**Extended data Fig. 9. Genetic landscape of AML based on the single-cell genotype data.** Heat maps (left) show the genotype of each sequenced cell for each variant, with clustering based on the genotypes of driver mutations. Each column represents a cell at the indicated scale. Cells with mutations and wild-type cells are indicated in blue and white, respectively. Cells with missing genotypes are indicated in grey. The subclones located to the right of the red line comprised <1% of the total sequence cells, since such small subclones can represent false positive or negative genotypes as a result of ADO or multiplets. The figures on the right show the pairwise association of mutations. The color and size of each panel represent the degree of the logarithmic odds ratio (log OR). The bar on the right side is a key indicating the association of the colors with the log OR. Co-occurrence and mutual exclusivity are indicated by red and blue, respectively. The statistical significance of the associations based on the false discovery rate (FDR) is indicated by the asterisks (*FDR < 0.1, **FDR < 0.05, ***FDR < 0.001).

**AML-01-001**

*NPM1, FLT3*-ITD, and *WT1* p.R370fs mutations co-occurred in the same cellular population.

**AML-01-002**

*NPM1, FLT3*-ITD, and the 2 *WT1* mutations co-occurred in the same cellular population.

**AML-02-001**

*SF3B1, DNMT3A, FLT3*-non-ITD*,* and *RUNX1* mutations co-occurred in the same cellular population. A large proportion of cells were ungenotyped for the *JAK2* mutation.

**AML-03-001**

*NPM1* and *FLT3*-ITD mutations co-occurred in the same cellular population.

**AML-04-002**

*SF3B1, SRSF2,* and one of the *IDH1* mutations co-occurred in the same cellular population. *FLT3*-ITD, and *FLT3*-non-ITD, and *NRAS* mutations were mutually exclusive at the cellular level. The 2 *IDH1* mutations (p.R132C and p.R132S) were mutually exclusive at the cellular level.

**AML-04-003**

*SF3B1, SRSF2, NRAS,* and *IDH1* p.R132C mutations co-occurred in the same cellular population. An *IDH1* p.R132S mutation co-occurred with *SF3B1* and *SRSF2* mutations, but was mutually exclusive with an *NRAS* mutation.

**AML-05-001**

*NRAS, FLT3*-ITD, and 3 *FLT3*-non-ITD mutations were mutually exclusive at the cellular level.

**AML-06-001**

*FLT3*-ITD and 4 *NRAS* mutations were mutually exclusive at the cellular level.

**AML-07-001**

*IDH2, SRSF2,* and *RUNX1* mutations co-occurred in the same cellular population.

**AML-07-002**

*IDH2* and *SRSF2* mutations co-occurred in the same cellular population. The number of *RUNX1*-mutated cells was too small to describe the pattern of co-occurrence/mutual exclusivity.

**AML-08-001**

*IDH2, DNMT3A,* and *SRSF2* mutations co-occurred in the same cellular population.

**AML-09-001**

An *NPM1* mutation co-occurred with *FLT3*-ITD, *FLT3*-non-ITD, or *KRAS* mutations in the same cellular population, whereas *FLT3*-ITD, 2 *FLT3*-non-ITD, and *KRAS* mutations were mutually exclusive with each other at the cellular level.

**AML-09-002**

*NPM1*, *FLT3*-non-ITD, and *WT1* mutations co-occurred in the same cellular population, although the number of *WT1*-mutated cells was relatively small.

**AML-10-001**

*WT1, GATA2,* and *FLT3*-ITD mutations co-occurred in the same cellular population. A large proportion of cells were ungenotyped for the *SRSF2* mutation.

**AML-11-001**

*SRSF2* and *ASXL1* mutations co-occurred in the same cellular population.

**AML-12-001**

An *NPM1* mutation co-occurred with *PTPN11* or *FLT3* mutations in the same cellular population, whereas *PTPN11* and *FLT3* mutations were mutually exclusive at the cellular level.

**AML-13-001**

*IDH2, WT1*, and *NPM1* mutations co-occurred in the same cellular population. The number of *TP53*-mutated cells was relatively small and *TP53* mutation showed the cellular-level mutual exclusivity with all other mutations.

**AML-14-001**

An *NPM1* mutation co-occurred with *PTPN11, NRAS*, or *KRAS* mutations in the same cellular population, whereas *PTPN11,* 2 *NRAS*, and 3 *KRAS* mutations were mutually exclusive at the cellular level.

**AML-15-001**

*IDH2* and *NPM1* mutations co-occurred in the same cellular population.

**AML-16-001**

*DNMT3A* and *IDH1* mutations co-occurred in the same cellular population. The number of *IDH2*-mutated cells was small and *IDH2* mutation showed the cellular-level mutual exclusivity with *IDH1* and *DNMT3A* mutations.

**AML-17-001**

*DNMT3A, NRAS,* and *NPM1* mutations co-occurred in the same cellular population.

**AML-18-001**

*IDH2, RUNX1,* and *ASXL1* mutations co-occurred in the same cellular population.

**AML-18-002**

*IDH2, RUNX1*, and *ASXL1* mutations co-occurred in the same cellular population.

**AML-19-001**

*FLT3*-ITD*, RUNX1*, and *WT1* mutations co-occurred in the same cellular population.

**AML-20-001**

*DNMT3A, NPM1*, and *NRAS* mutations co-occurred in the same cellular population.

**AML-21-001**

*WT1, NPM1,* and *NRAS* mutations co-occurred in the same cellular population.

**AML-21-002**

*NPM1, FLT3*-ITD, and 2 *WT1* mutations co-occurred in the same cellular population. The number of *NRAS*-mutated cells was too small to describe the pattern of co-occurrence/mutual exclusivity.

**AML-22-001**

The 2 *KRAS* and 6 *NRAS* mutations were mutually exclusive at the cellular level.

**AML-23-001**

*WT1* and *NRAS* mutations co-occurred in the same cellular population.

**AML-24-001**

*DNMT3A, U2AF1, WT1*, and *FLT3*-ITD mutations co-occurred in the same cellular population. The number of *FLT3*-non-ITD-mutated cells was too small to describe the pattern of co-occurrence/mutual exclusivity.

**AML-25-001**

*EZH2, ASXL1*, and *RUNX1* mutations co-occurred in the same cellular population. The number of *FLT3*-ITD-mutated cells was small, and *FLT3*-ITD co-occurred with all other mutations in the same cells.

**AML-26-001**

*DNMT3A* and *NPM1* mutations co-occurred in the same cellular population with either *FLT3* p.D835A, *FLT3* p.D835Y, *KRAS,* or *NRAS* mutations. *FLT3* p.D835A, *FLT3* p.D835Y, *KRAS*, and *NRAS* mutations were mutually exclusive at the cellular level.

**AML-27-001**

*TP53, DNMT3A, NPM1,* and *FLT3*-ITD mutations co-occurred in the same cellular population.

**AML-29-001**

*ASXL1, SRSF2,* and *RUNX1* mutations co-occurred in the same cellular population. A large proportion of cells were ungenotyped for *SRSF2* and *RUNX1* mutations.

**AML-30-001**

*DNMT3A, NPM1, IDH2,* and *NRAS* mutations co-occurred in the same cellular population.

**AML-31-001**

An *NPM1* mutation co-occurred in the same cellular population with either *NRAS, WT1, FLT3*-ITD, or *PTPN11* mutations, whereas *NRAS, WT1, FLT3*-ITD, and *PTPN11* mutations were mutually exclusive at the cellular level.

**AML-32-001**

*SF3B1, DNMT3A, RUNX1* and one of the *FLT3* mutations co-occurred in the same cellular population, whereas *FLT3*-non-ITD and *FLT3*-ITD mutations were mutually exclusive at the cellular level.

**AML-33-001**

*IDH2, SRSF2,* and *RUNX1* mutations co-occurred in the same cellular population.

**AML-34-001**

The majority of cells had no mutations, which is in accordance with the low bone marrow blast percentage (7%). A small subclone showing the cellular-level co-occurrence of *SF3B1* and *KRAS* mutations was detected. The number of cells that were mutated with *NRAS* p.G13D or *NRAS* p.G12D was too small to describe the pattern of co-occurrence/mutual exclusivity.

**AML-35-001**

*NRAS* and *PTPN11* mutations were mutually exclusive at the cellular level. The number of *FLT3*-ITD-mutated cells was too small to describe the pattern of co-occurrence/mutual exclusivity.

**AML-36-001**

*FLT3*-non-ITD*, NRAS* p.G12S, and *NRAS* p.G12D mutations were mutually exclusive at the cellular level.

**AML-37-001**

*DNMT3A*, *NPM1*, and *NRAS* mutations co-occurred in the same cellular population.

**AML-38-001**

*IDH1* and *IDH2* mutations were mutually exclusive at the cellular level. *FLT3*-ITD*, PTPN11*, 2 *KRAS*, and 2 *NRAS* mutations were mutually exclusive at the cellular level.

**AML-38-002**

*PTPN11*, *KRAS*, and *NRAS* mutations were mutually exclusive at the cellular level.

**AML-38-003**

*PTPN11, KRAS,* and *NRAS* mutations were mutually exclusive at the cellular level.

**AML-39-001**

*SF3B1, GATA2,* and one of the *PTPN11* mutations co-occurred in the same cellular population, whereas the 2 *PTPN11* mutations (p.G503A and p.A72V) were mutually exclusive at the cellular level.

**AML-39-002**

*SF3B1, GATA2,* and *PTPN11* mutations co-occurred in the same cellular population.

**AML-40-001**

*IDH2, NPM1,* and *NRAS* p.G12D mutations co-occurred in the same cellular population. *NRAS* p.G12D and *FLT3* mutations were mutually exclusive at the cellular level. The number of cells that were mutated with *NRAS* p.G13D or *FLT3*-ITD was too small to describe the pattern of co-occurrence/mutual exclusivity.

**AML-41-001**

*TP53, WT1, IDH2*, and *SRSF2* mutations co-occurred in the same cellular population. Although the number of mutated cells was small, *NRAS, KRAS*, and *PTPN11* mutations tended to be mutually exclusive at the cellular level.

**AML-42-001**

*EZH2, ASXL1,* and *NRAS* p.G13D mutations co-occurred in the same cellular population. Although the number of mutated cells was small for *NRAS* p.G13V and *NRAS* p.G12D mutations, the 3 *NRAS* mutations (p.G13D, p.G13V, and p.G12D) tended to be mutually exclusive at the cellular level.

**AML-43-001**

*FLT3*-ITD and 2 *WT1* mutations co-occurred in the same cellular population.

**AML-44-001**

*WT1* and *RUNX1* mutations co-occurred in the same cellular population. The number of *FLT3*-ITD-mutated cells was too small to describe the pattern of co-occurrence/mutual exclusivity.

**AML-46-001**

*PTPN11* and *NPM1* mutations co-occurred in the same cellular population.

**AML-47-001**

*WT1, NPM1,* and *FLT3*-non-ITD mutations co-occurred in the same cellular population.

**AML-48-001**

*IDH2* and *KRAS* mutations were mutually exclusive at the cellular level.

**AML-49-001**

*NPM1, FLT3*-ITD, and *NRAS* mutations co-occurred in the same cellular population.

**AML-50-001**

*NPM1* and *FLT3*-non-ITD mutations co-occurred in the same cellular population.

**AML-52-001**

*IDH2, NPM1*, and *FLT3*-ITD mutations co-occurred in the same cellular population.

**AML-53-001**

*FLT3*-ITD and 2 *NRAS* mutations were mutually exclusive at the cellular level.

**AML-54-001**

*SRSF2, ASXL1*, and *FLT3*-ITD mutations co-occurred in the same cellular population. Note that the number of ungenotyped cells was relatively large for *SRSF2* and *ASXL1* mutations.

**AML-55-001**

*RUNX1, SRSF2, DNMT3A, IDH1*, and *ASXL1* mutations co-occurred in the same cellular population.

**AML-56-001**

Only a mutation in *IDH2* was detected in this sample.

**AML-57-001**

*SRSF2* and *NPM1* mutations co-occurred in the same cellular population.

**AML-58-001**

*IDH2, DNMT3A* and *SRSF2* mutations co-occurred in the same cellular population.

**AML-60-001**

*ASXL1, IDH2* and *SRSF2* mutations co-occurred in the same cellular population. The number of *FLT3*-ITD-mutated cells was too small to describe the pattern of co-occurrence/mutual exclusivity.

**AML-61-001**

An *NPM1* mutation co-occurred with one of the *PTPN11* or *NRAS* mutations in the same cellular population. The 2 *PTPN11* mutations (p.N58D and p.P491S) and *NRAS* mutations were mutually exclusive at the cellular level.

**AML-62-001**

*IDH1* and *NPM1* mutations co-occurred in the same cellular population with either one of the *PTPN11* mutations or with an *FLT3*-ITD mutation. The 3 *PTPN11* mutations (p.F71L, p.A72T, and p.D61A) and *FLT3*-ITD mutation were mutually exclusive at the cellular level.

**AML-63-002**

An *IDH2* mutation co-occurred with an *NPM1* mutation in the same cellular population. The number of *FLT3*-ITD-mutated cells was too small to describe the pattern of co-occurrence/mutual exclusivity.

**AML-63-003**

An *IDH2* mutation co-occurred with an *NPM1* mutation in the same cellular population.

**AML-64-001**

*IDH2, DNMT3A, SRSF2*, and *NRAS* mutations co-occurred in the same cellular population.

**AML-65-001**

A *DNMT3A* mutation co-occurred with an *FLT3*-ITD mutation in the same cellular population.

**AML-66-001**

*ASXL1, PTPN11, NRAS,* and *SRSF2* mutations co-occurred in the same cellular population. A large proportion of cells were ungenotyped for *SRSF2* mutation.

**AML-66-002**

*ASXL1, PTPN11, NRAS,* and *SRSF2* mutations co-occurred in the same cellular population. A large proportion of cells were ungenotyped for *SRSF2* mutation.

**AML-66-003**

*ASXL1, PTPN11, NRAS*, and *SRSF2* mutations co-occurred in the same cellular population.

**AML-67-001**

Two *U2AF1* mutations co-occurred in the same cellular population with either *NRAS* or *KRAS* mutations. *NRAS* and 2 *KRAS* mutations were mutually exclusive at the cellular level. *IDH2* mutation tended to be mutually exclusive with other mutations. The number of *TP53*-mutated cells was too small to describe the pattern of co-occurrence/mutual exclusivity.

**AML-69-001**

*NRAS* and *KRAS* mutations were mutually exclusive at the cellular level. *KRAS* and *RUNX1* mutations co-occurred in the same cellular population.

**AML-70-001**

*IDH2, SRSF2,* and *KRAS* mutations co-occurred in the same cellular population. *KRAS* and *NRAS* mutations co-occurred in a small number of cells.

**AML-71-001**

Although the number of cells with *RUNX1* mutation was small, a small subclone showing the cellular-level co-occurrence of *RUNX1* and *NRAS* mutations was detected.

**AML-72-001**

*PTPN11* and *NPM1* mutations co-occurred in the same cellular population.

**AML-73-001**

*SF3B1* and *FLT3*-non-ITD mutations co-occurred in the same cellular population.

**AML-74-001**

An *NPM1* mutation co-occurred with either *PTPN11, NRAS*, *KRAS*, or *WT1* mutations in the same cellular population, whereas *PTPN11, NRAS*, *KRAS*, and *WT1* mutations tended to be mutually exclusive at the cellular level.

**AML-75-001**

*EZH2* and *NRAS* mutations co-occurred in the same cellular population. A large proportion of cells were ungenotyped for *SRSF2* mutation.

**AML-76-001**

*U2AF1* and *DNMT3A* mutations co-occurred in the same cellular population with either *NRAS* p.Q61H or *FLT3*-ITD mutations. *FLT3*-ITD and 2 *NRAS* mutations (p.Q61H and p.G13R) were mutually exclusive at the cellular level. The number of cells that were mutated with *FLT3* p.836_837del or *FLT3* p.D839G was too small to describe the pattern of co-occurrence/mutual exclusivity.

**AML-77-001**

*U2AF1, IDH2*, and *ASXL1* mutations co-occurred in the same cellular population.

**Extended Data Fig. 10**

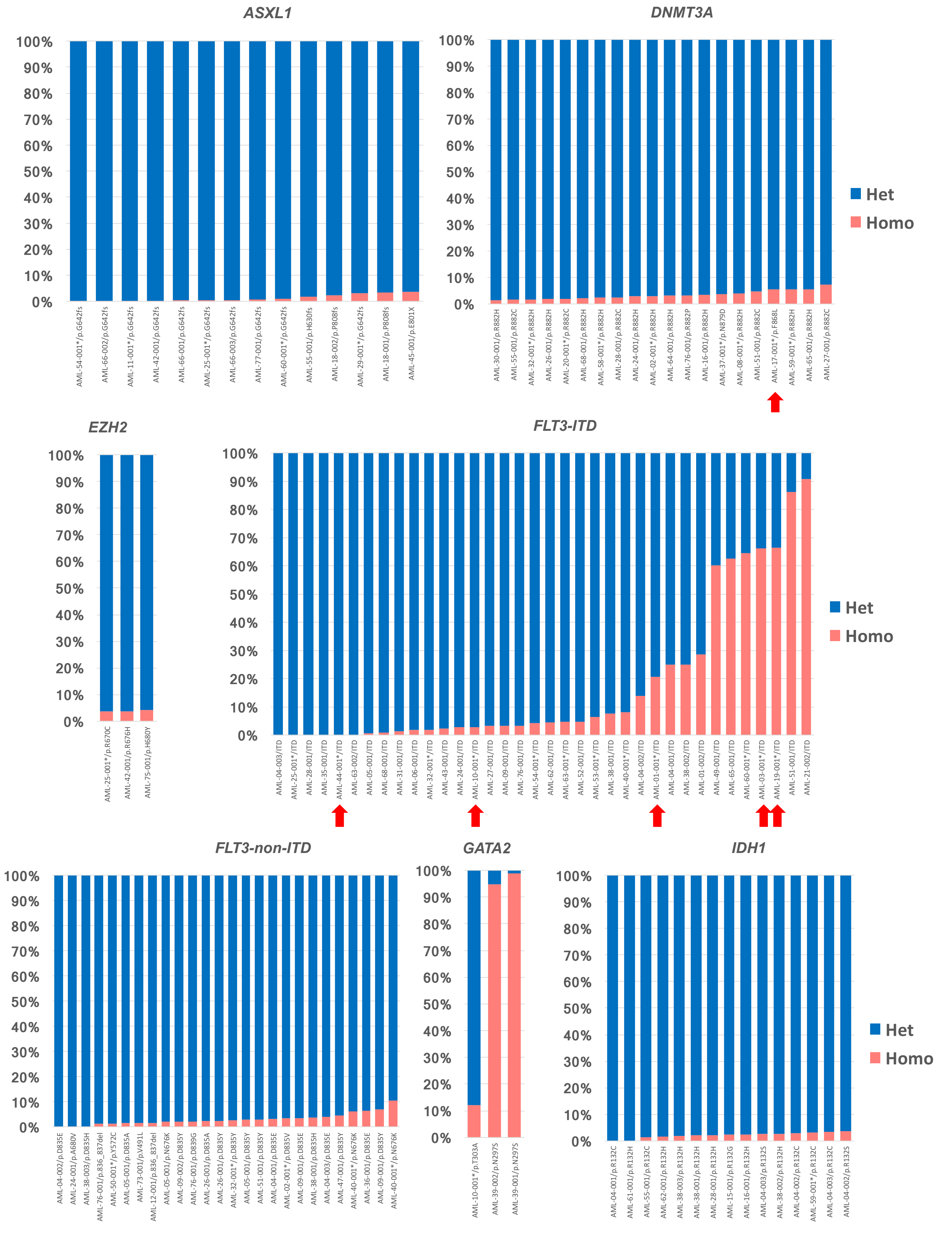

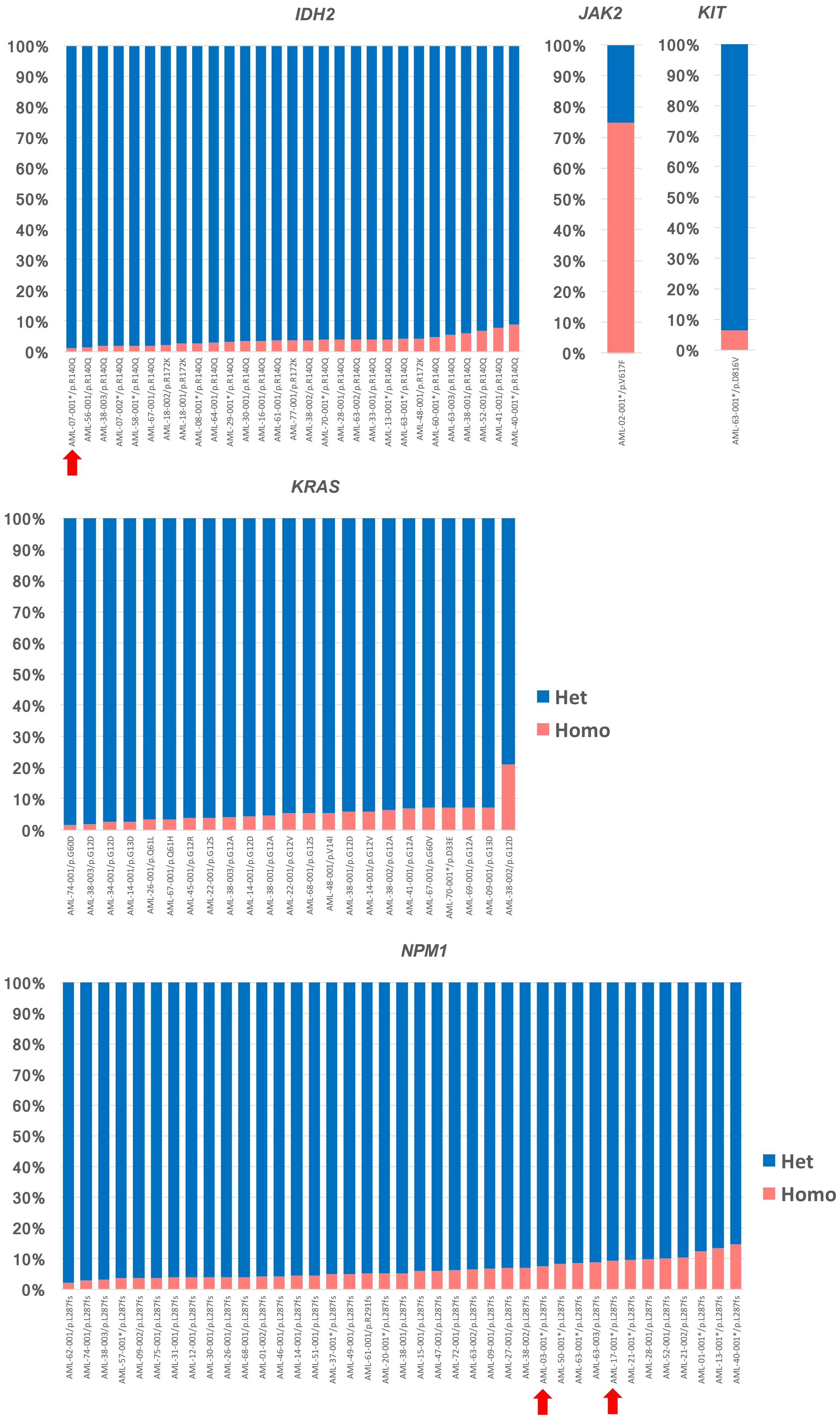
IDH2
JAK2
KIT
100%
90%
80%
70%
60%
50%
40%
30%
20%
10%
0%
AML-07-001*/p.R140Q
AML-56-001/p.R140Q
AML-38-003/p.R140Q
AML-07-002*/p.R140Q
AML-58-001*/p.R140Q
AML-67-001/p.R140Q
AML-18-002/p.R172K
AML-18-001/p.R172K
AML-08-001*/p.R140Q
AML-64-001/p.R140Q
AML-29-001*/p.R140Q
AML-30-001/p.R140Q
AML-16-001/p.R140Q
AML-61-001/p.R140Q
AML-77-001/p.R172K
AML-38-002/p.R140Q
AML-70-001*/p.R140Q
AML-28-001/p.R140Q
AML-63-002/p.R140Q
AML-33-001/p.R140Q
AML-13-001*/p.R140Q
AML-63-001*/p.R140Q
AML-48-001/p.R172K
AML-60-001*/p.R140Q
AML-63-003/p.R140Q
AML-38-001/p.R140Q
AML-52-001/p.R140Q
AML-41-001/p.R140Q
AML-40-001*/p.R140Q
AML-02-001*/p.V617F
AML-63-001*/p.D816V
KRAS
AML-74-001/p.G60D
AML-38-003/p.G12D
AML-34-001/p.G12D
AML-14-001/p.G13D
AML-26-001/p.Q61L
AML-67-001/p.Q61H
AML-45-001/p.G12R
AML-22-001/p.G12S
AML-38-003/p.G12A
AML-14-001/p.G12D
AML-38-001/p.G12A
AML-22-001/p.G12V
AML-68-001/p.G12S
AML-48-001/p.V14I
AML-38-001/p.G12D
AML-14-001/p.G12V
AML-38-002/p.G12A
AML-41-001/p.G12A
AML-67-001/p.G60V
AML-70-001*/p.D33E
AML-69-001/p.G12A
AML-09-001/p.G13D
AML-38-002/p.G12D
Het
Homo
NPM1
AML-62-001/p.L287fs
AML-74-001/p.L287fs
AML-38-003/p.L287fs
AML-57-001*/p.L287fs
AML-09-002/p.L287fs
AML-75-001/p.L287fs
AML-31-001/p.L287fs
AML-12-001/p.L287fs
AML-30-001/p.L287fs
AML-26-001/p.L287fs
AML-68-001/p.L287fs
AML-01-002/p.L287fs
AML-46-001/p.L287fs
AML-14-001/p.L287fs
AML-51-001/p.L287fs
AML-37-001*/p.L287fs
AML-49-001/p.L287fs
AML-61-001/p.R291fs
AML-20-001*/p.L287fs
AML-38-001/p.L287fs
AML-15-001/p.L287fs
AML-47-001/p.L287fs
AML-72-001/p.L287fs
AML-63-002/p.L287fs
AML-09-001/p.L287fs
AML-27-001/p.L287fs
AML-38-002/p.L287fs
AML-03-001*/p.L287fs
AML-50-001*/p.L287fs
AML-63-001*/p.L287fs
AML-63-003/p.L287fs
AML-17-001*/p.L287fs
AML-21-001*/p.L287fs
AML-28-001/p.L287fs
AML-52-001/p.L287fs
AML-21-002/p.L287fs
AML-01-001*/p.L287fs
AML-13-001*/p.L287fs
AML-40-001*/p.L287fs
Het
Homo

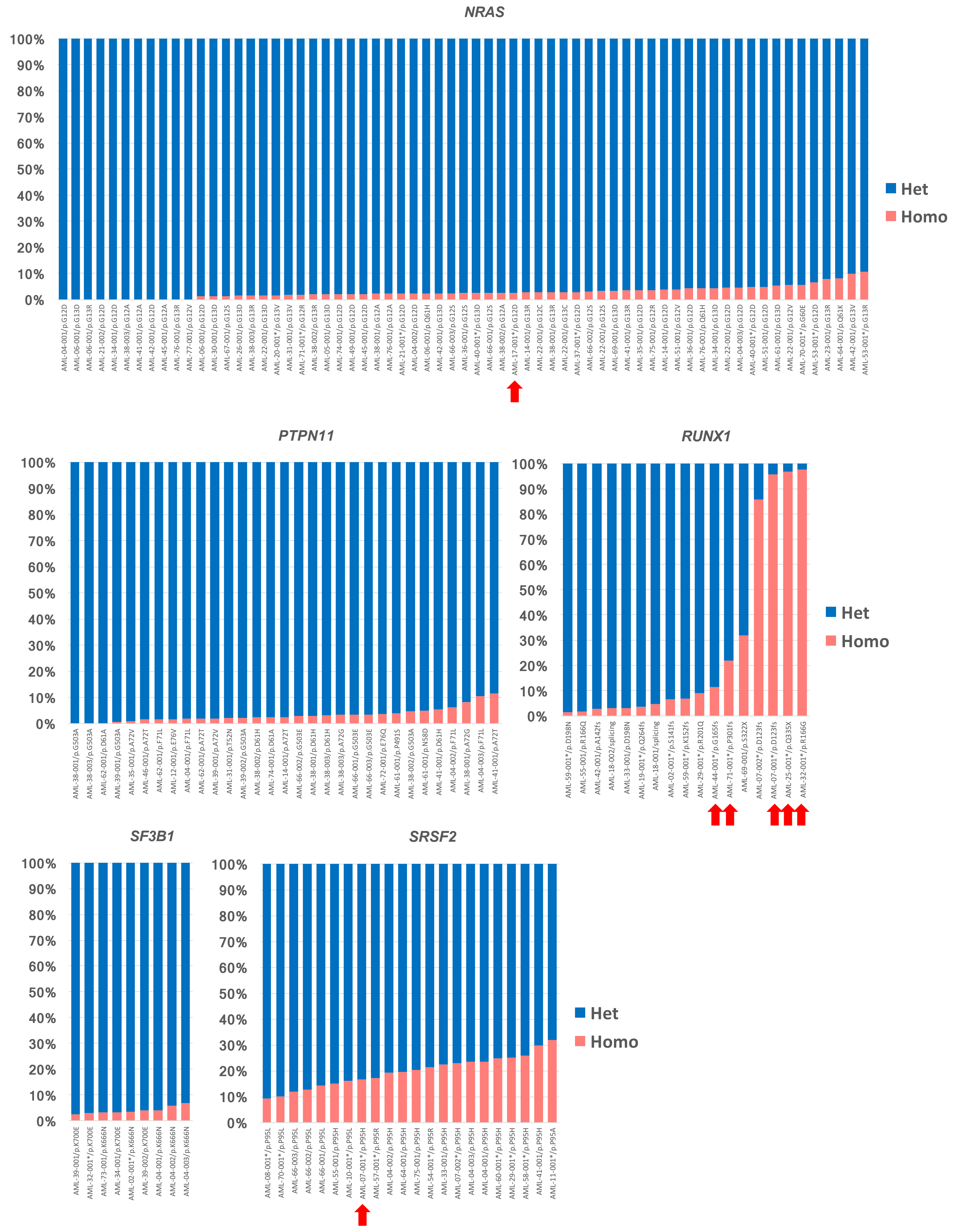
NRAS
100%
90%
80%
70%
60%
50%
40%
30%
20%
10%
0%
Het
Homo
AML-04-001/p.G12D
AML-06-001/p.G13D
AML-06-001/p.G13R
AML-21-002/p.G12D
AML-34-001/p.G12D
AML-38-003/p.G12A
AML-41-001/p.G12A
AML-42-001/p.G12D
AML-45-001/p.G12A
AML-76-001/p.G13R
AML-77-001/p.G12V
AML-06-001/p.G12D
AML-30-001/p.G13D
AML-67-001/p.G12S
AML-26-001/p.G13D
AML-38-003/p.G13R
AML-22-001/p.G13D
AML-20-001*/p.G13V
AML-31-001/p.G13V
AML-71-001*/p.G12R
AML-38-002/p.G13R
AML-05-001/p.G13D
AML-74-001/p.G12D
AML-49-001/p.G12D
AML-45-001/p.G12D
AML-38-001/p.G12A
AML-76-001/p.G12A
AML-21-001*/p.G12D
AML-04-002/p.G12D
AML-06-001/p.Q61H
AML-42-001/p.G13D
AML-66-003/p.G12S
AML-36-001/p.G12S
AML-40-001*/p.G13D
AML-66-001/p.G12S
AML-38-002/p.G12A
AML-17-001*/p.G12D
AML-14-001/p.G13R
AML-22-001/p.G12C
AML-38-001/p.G13R
AML-22-001*/p.G13C
AML-37-001*/p.G12D
AML-66-002/p.G12S
AML-22-001/p.G12S
AML-69-001/p.G13D
AML-41-001/p.G13R
AML-35-001/p.G12D
AML-75-001/p.G12R
AML-14-001/p.G12D
AML-51-001/p.G12V
AML-36-001/p.G12D
AML-76-001/p.Q61H
AML-34-001/p.G13D
AML-22-001/p.G13D
AML-04-003/p.G12D
AML-40-001*/p.G12D
AML-51-001/p.G12D
AML-61-001/p.G13D
AML-22-001/p.G12V
AML-70-001*/p.G60E
AML-53-001*/p.G12D
AML-23-001/p.G13R
AML-64-001/p.Q61K
AML-42-001/p.G13V
AML-53-001*/p.G13R
PTPN11
100%
90%
80%
70%
60%
50%
40%
30%
20%
10%
0%
AML-38-001/p.G503A
AML-38-003/p.G503A
AML-62-001/p.D61A
AML-39-001/p.G503A
AML-35-001/p.A72V
AML-46-001/p.A72T
AML-62-001/p.F71L
AML-12-001/p.E76V
AML-04-001/p.F71L
AML-62-001/p.A72T
AML-39-001/p.A72V
AML-31-001/p.T52N
AML-39-002/p.G503A
AML-38-002/p.D61H
AML-74-001/p.D61A
AML-14-001/p.A72T
AML-66-002/p.G503E
AML-38-001/p.D61H
AML-38-003/p.D61H
AML-38-003/p.A72G
AML-66-001/p.G503E
AML-66-003/p.G503E
AML-72-001/p.E76Q
AML-61-001/p.P491S
AML-38-002/p.G503A
AML-61-001/p.N58D
AML-41-001/p.D61H
AML-04-002/p.F71L
AML-38-001/p.A72G
AML-04-003/p.F71L
AML-41-001/p.A72T
RUNX1
100%
90%
80%
70%
60%
50%
40%
30%
20%
10%
0%
Het
Homo
AML-59-001*/p.D198N
AML-55-001/p.R166Q
AML-42-001/p.A142fs
AML-18-002/splicing
AML-33-001/p.D198N
AML-19-001*/p.Q264fs
AML-18-001/splicing
AML-02-001*/p.S141fs
AML-59-001*/p.K152fs
AML-29-001*/p.R201Q
AML-44-001*/p.G165fs
AML-71-001*/p.P301fs
AML-69-001/p.S322X
AML-07-002*/p.D123fs
AML-07-001*/p.D123fs
AML-25-001*/p.Q335X
AML-32-001*/p.R166G
SF3B1
100%
90%
80%
70%
60%
50%
40%
30%
20%
10%
0%
AML-39-001/p.K700E
AML-32-001*/p.K700E
AML-73-001/p.K666N
AML-34-001/p.K700E
AML-02-001*/p.K666N
AML-39-002/p.K700E
AML-04-001/p.K666N
AML-04-002/p.K666N
AML-04-003/p.K666N
SRSF2
100%
90%
80%
70%
60%
50%
40%
30%
20%
10%
0%
Het
Homo
AML-08-001*/p.P95L
AML-70-001*/p.P95L
AML-66-003/p.P95L
AML-66-002/p.P95L
AML-66-001/p.P95L
AML-55-001/p.P95H
AML-10-001*/p.P95L
AML-07-001*/p.P95H
AML-57-001*/p.P95R
AML-04-002/p.P95H
AML-64-001/p.P95H
AML-75-001/p.P95H
AML-54-001*/p.P95R
AML-33-001/p.P95H
AML-07-002*/p.P95H
AML-04-003/p.P95H
AML-04-001/p.P95H
AML-60-001*/p.P95H
AML-29-001*/p.P95H
AML-58-001*/p.P95H
AML-41-001/p.P95H
AML-11-001*/p.P95A

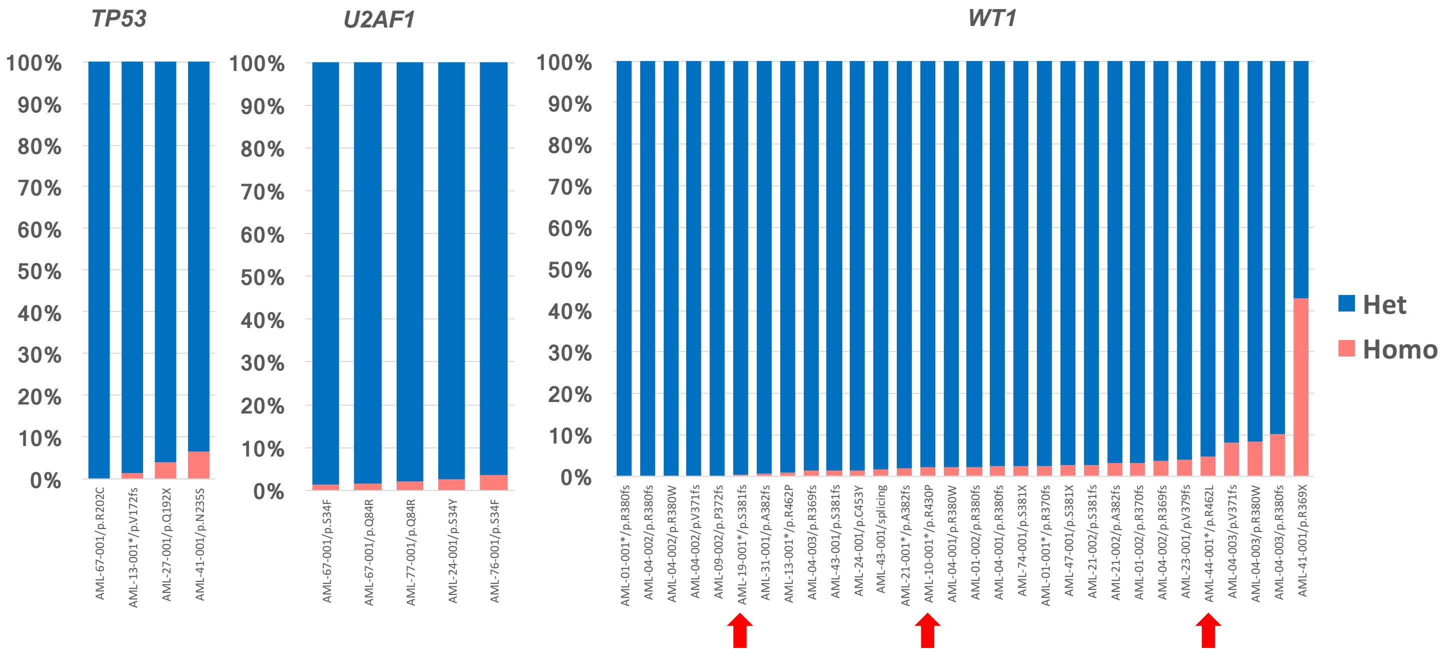


**Extended Data Fig. 10. Zygosity distributions for genotyped cells.** Bar charts show the proportion of mutated cells with different zygosity among the total number of genotyped cells. Cells genotyped as having heterozygous and homozygous mutations are shown in blue and red, respectively. *FLT3*-ITD*, GATA2, NPM1, RUNX1*, and *SRSF2* variants were frequently genotyped as homozygous. The X axis shows the sample ID, and the Y axis shows the percentage of cells with each genotype. The bars are sorted in ascending order based on the percentage of homozygously genotyped cells for each gene. Samples analyzed by SNP array are indicated with asterisks in X axis. The variants with SNP-array-confirmed copy number alterations are indicated with red arrows. Het, heterozygous; Homo, homozygous.

**Extended Data Fig. 11**

AML-71-001

Karyotype: 46,XX[20]

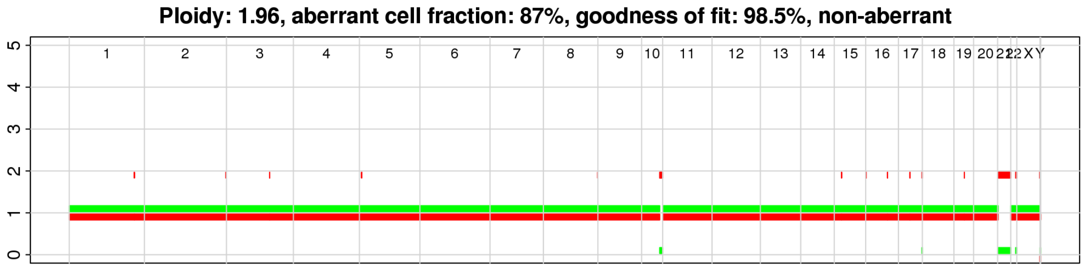


| chr | Gene | AA change | WT | Het | Homo | Missing | Homo [%] | allele count |
|---|---|---|---|---|---|---|---|---|
| 1 | *NRAS* | p.G12R | 309 | 6703 | 127 | 1386 | 2 | 1,1 |
| 21 | *RUNX1* | p.P301fs | 6779 | 202 | 57 | 1487 | 22 | 2,0 |

#### Heatmap ignoring zygosity

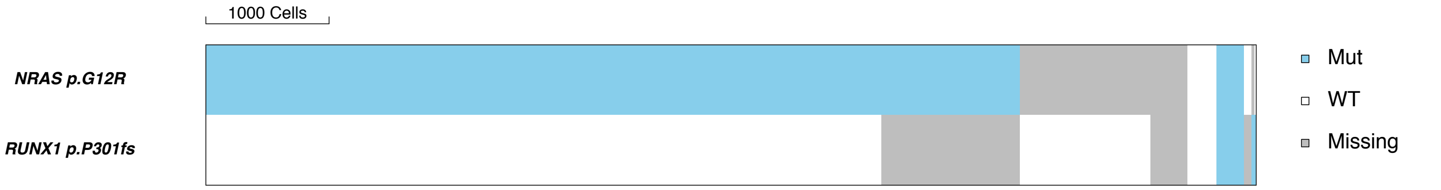


#### Heatmap considering zygosity if SNP array-confirmed

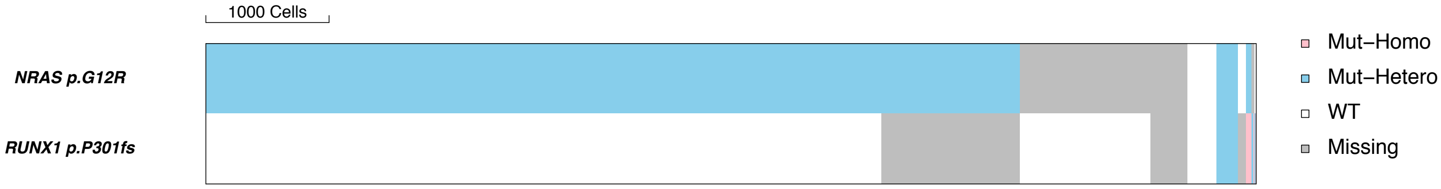


#### Distribution of depth for *RUNX1* p.P301fs based on genotype

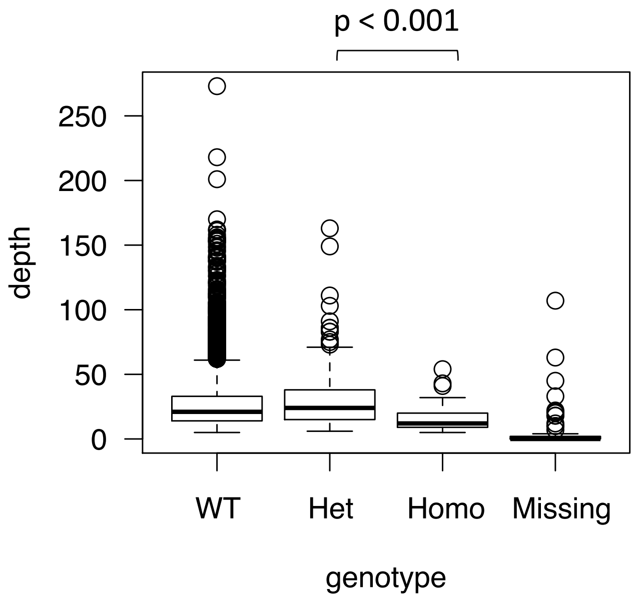

AML-32-001

Karyotype: 47,XY,+13[1];46,XY[19]

Ploidy: 1.93, aberrant cell fraction: 56%, goodness of fit: 86.9%

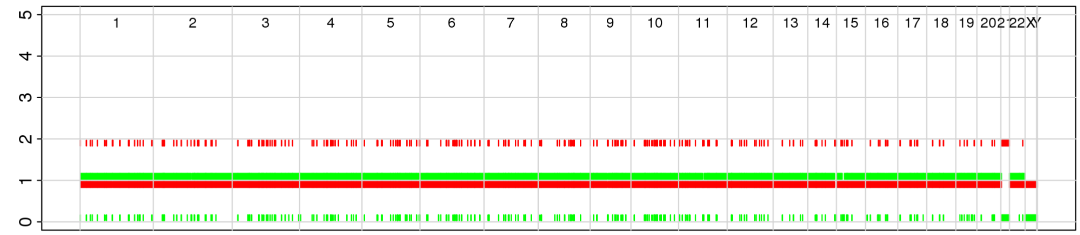


| chr | Gene | AA change | WT | Het | Homo | Missing | Homo [%] | allele count |
|---|---|---|---|---|---|---|---|---|
| 2 | *SF3B1* | p.K700E | 1038 | 6830 | 205 | 168 | 3 | 1,1 |
| 2 | *DNMT3A* | p.R882H | 902 | 5551 | 93 | 1695 | 2 | 1,1 |
| 21 | *RUNX1* | p.R166G | 676 | 125 | 4966 | 2474 | 98 | 2,0 |
| 13 | *FLT3* | p.D835Y | 2500 | 5422 | 151 | 168 | 3 | 1,1 |
| 13 | *FLT3* | ITD | 6345 | 878 | 17 | 1001 | 2 | 1,1 |

### Heatmap ignoring zygosity

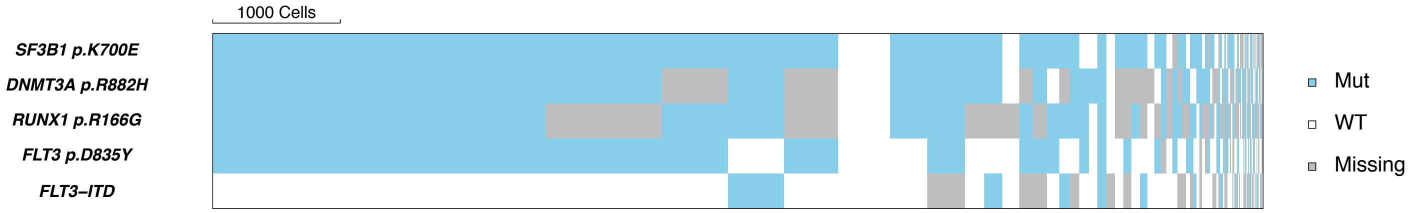


### Heatmap considering zygosity if SNP array-confirmed

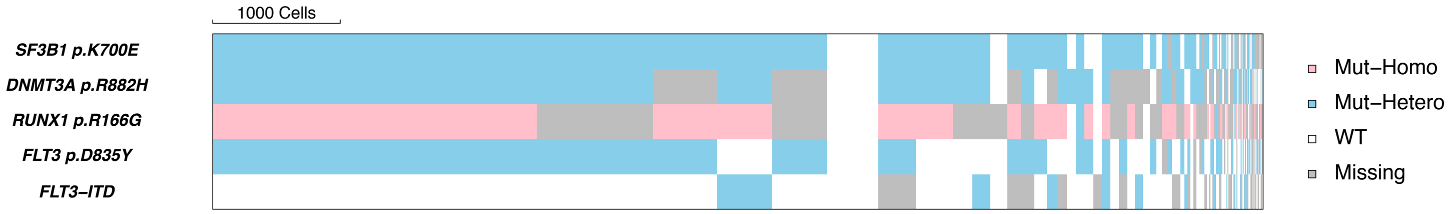


Distribution of depth for *RUNX1* p.R166G based on genotype

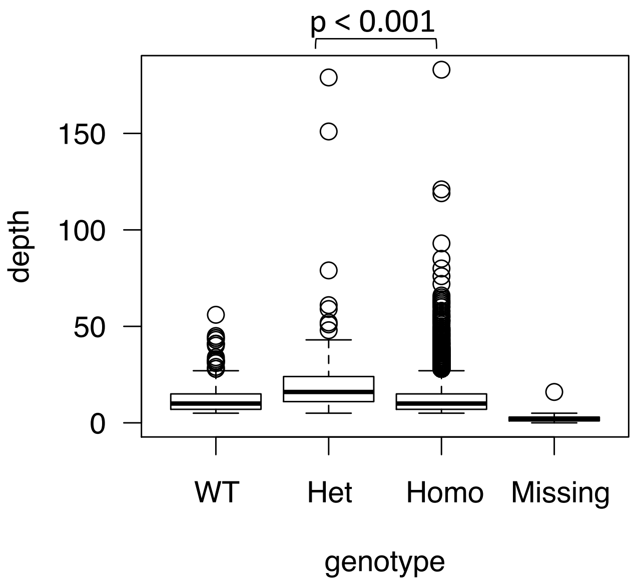

AML-19-001 46,XY[20]

Ploidy: 3.35, aberrant cell fraction: 90%, goodness of fit: 85.0%

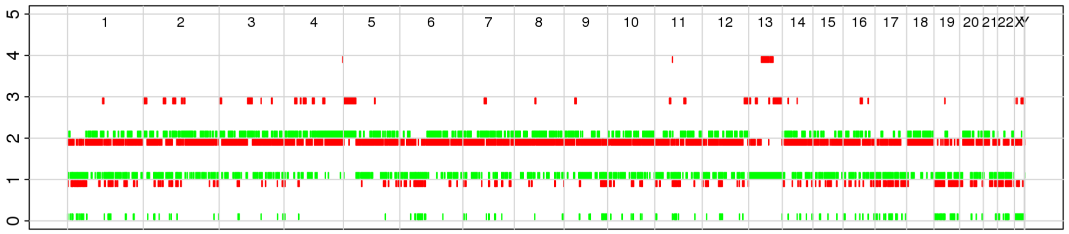


| chr | Gene | AA change | WT | Het | Homo | Missing | Homo [%] | allele count |
|---|---|---|---|---|---|---|---|---|
| 13 | *FLT3* | ITD | 575 | 1420 | 2815 | 456 | 66 | 2,1 |
| 21 | *RUNX1* | p.Q264fs | 1020 | 3389 | 125 | 732 | 4 | 1,1 |
| 11 | *WT1* | p.S381fs | 3350 | 1470 | 5 | 441 | 0 | 2,1 |

### Heatmap ignoring zygosity

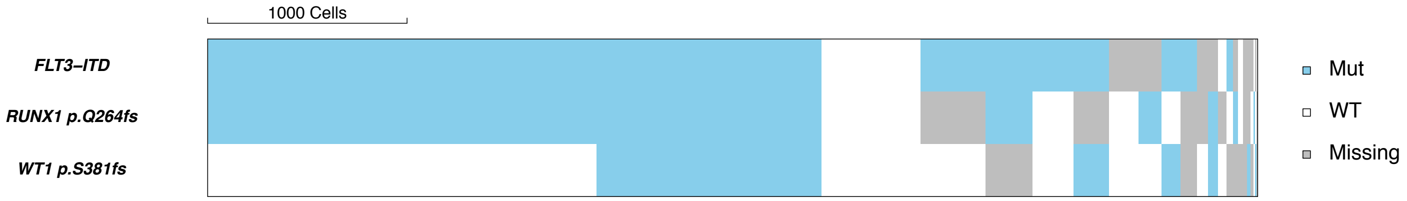


### Heatmap considering zygosity if SNP array-confirmed

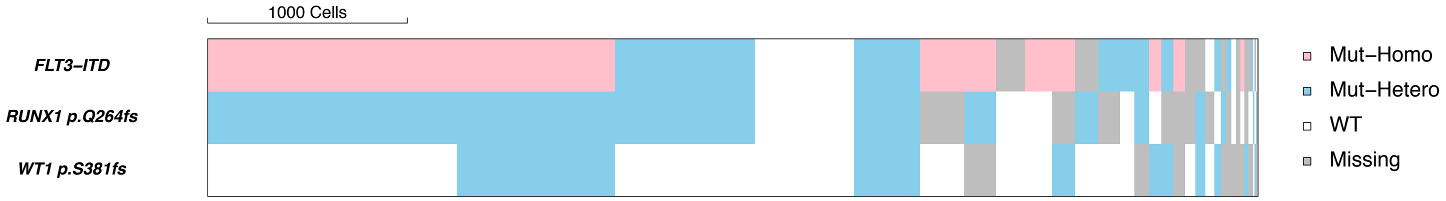


#### Distribution of depth for *FLT3-ITD* and *WT1* p.S381fs based on genotype

*FLT3-ITD*

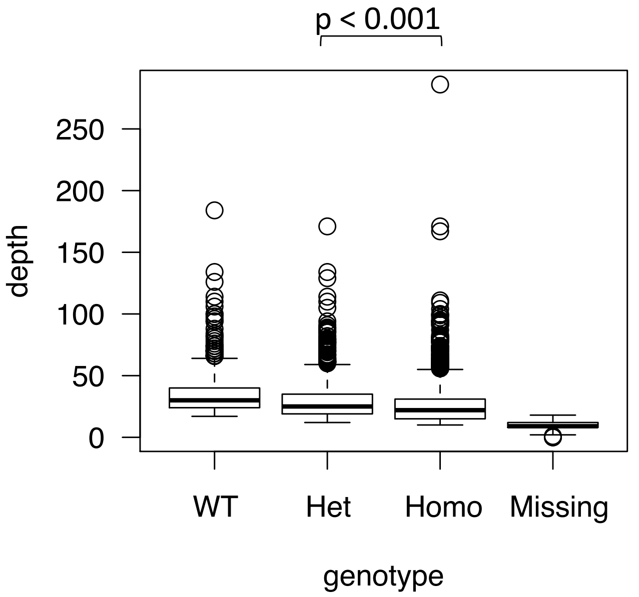


*WT1* p.S381fs

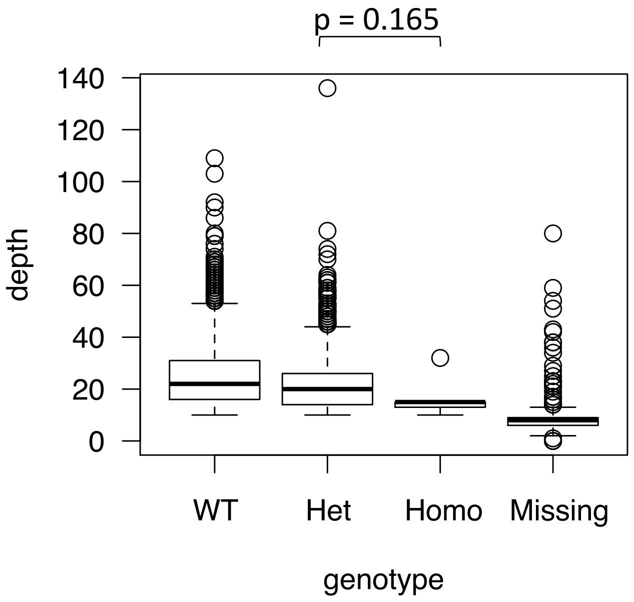

AML-44-001

Karyotype: 47,XY,+mar[1]; 46,XY[19]

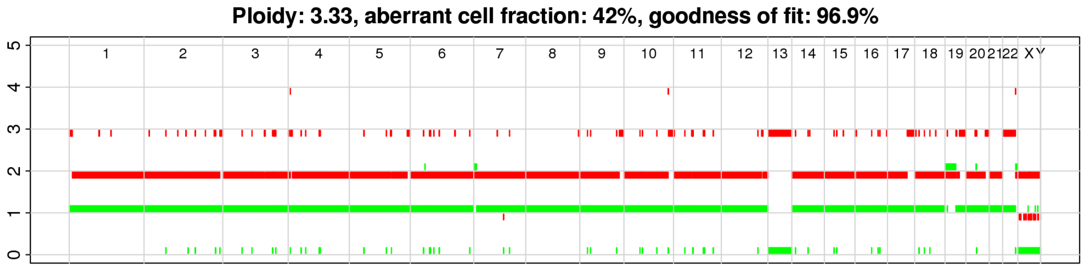


| chr | Gene | AA change | WT | Het | Homo | Missing | Homo [%] | allele count |
|---|---|---|---|---|---|---|---|---|
| 11 | *WT1* | p.R462L | 2720 | 3183 | 159 | 697 | 5 | 2,1 |
| 21 | *RUNX1* | p.G165fs | 2272 | 3038 | 398 | 1051 | 12 | 2,1 |
| 13 | *FLT3* | ITD | 6702 | 23 | 0 | 34 | 0 | 3,0 |

### Heatmap ignoring zygosity

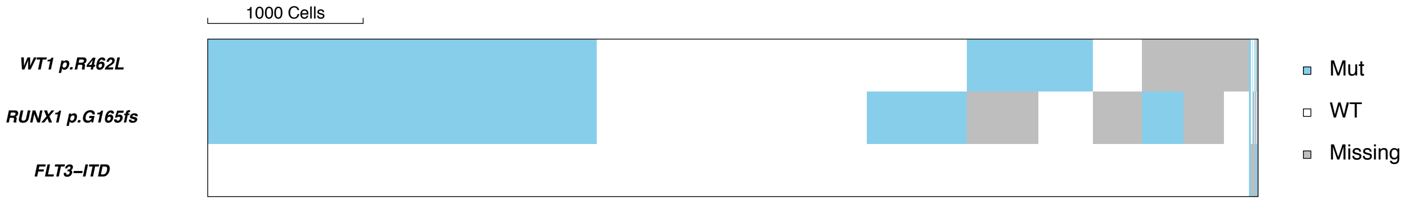


### Heatmap considering zygosity if SNP array-confirmed

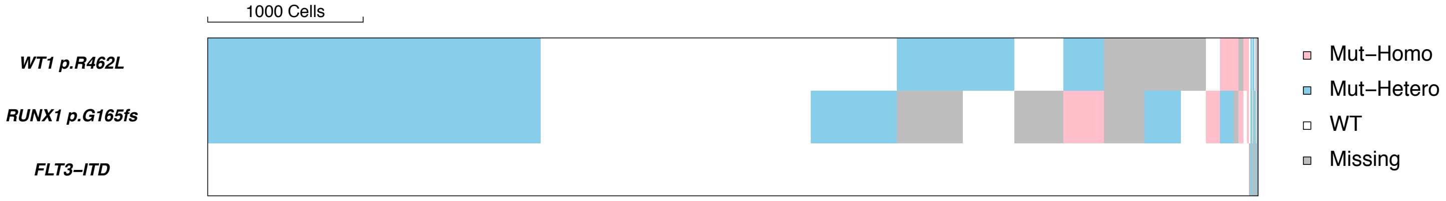


#### Distribution of depth for *RUNX1* p.G165fs based on genotype

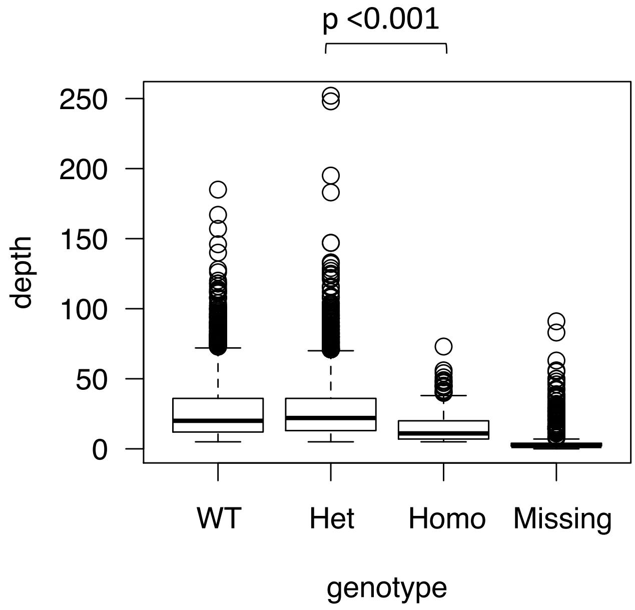

AML-01-001

karyotype: 45XY,add(6)(q27),-20[1]; 46XY[19]

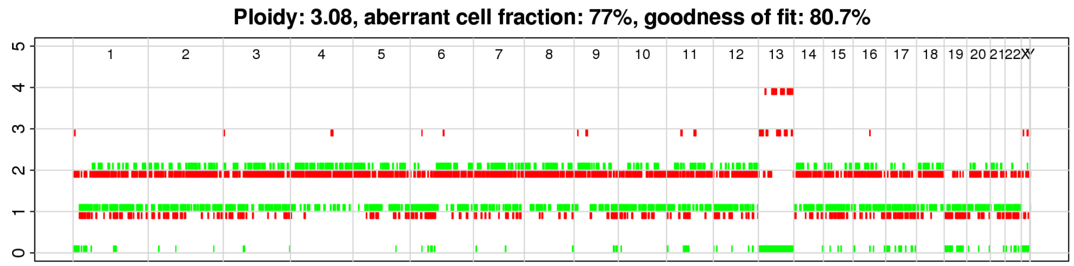


| chr | Gene | AA change | WT | Het | Homo | Missing | Homo [%] | allele count |
|---|---|---|---|---|---|---|---|---|
| 5 | *NPM1* | p.L287fs | 1285 | 4226 | 606 | 5254 | 13 | 1,1 |
| 13 | *FLT3* | ITD | 239 | 8444 | 2190 | 498 | 21 | 3,0 |
| 11 | *WT1* | p.R370fs | 568 | 9799 | 248 | 756 | 2 | 1,1 |
| 11 | *WT1* | p.R380fs | 10779 | 39 | 0 | 553 | 0 | 1,1 |

#### Heatmap ignoring zygosity

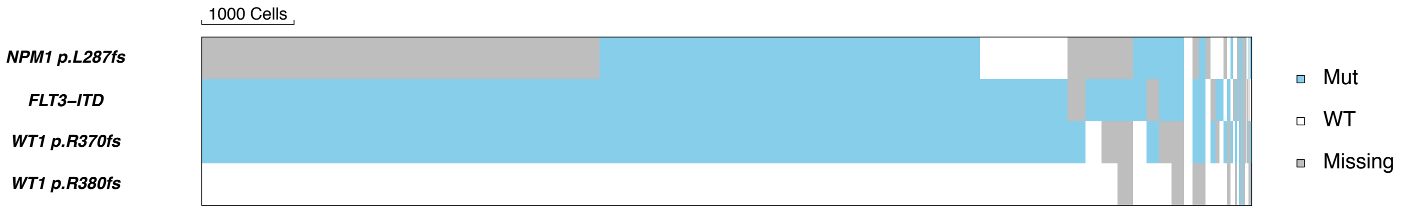


#### Heatmap considering zygosity if SNP array-confirmed

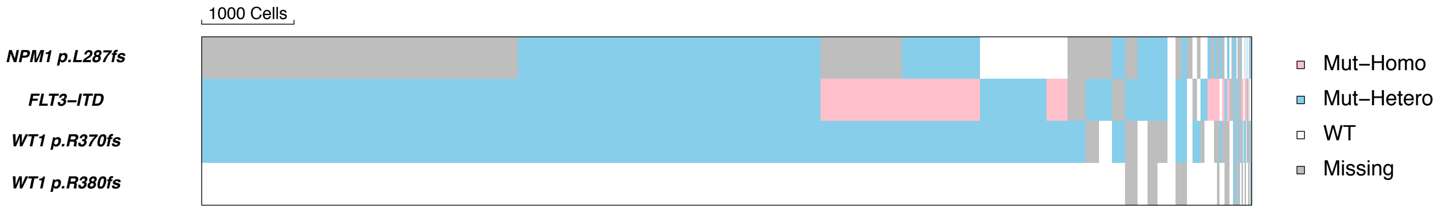


#### Distribution of depth for *NPM1* p.L287fs and *FLT3-ITD* based on genotype

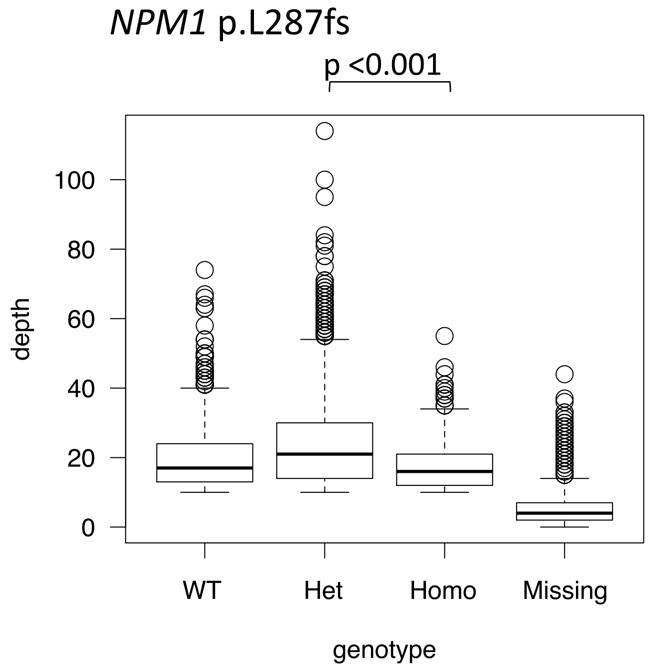


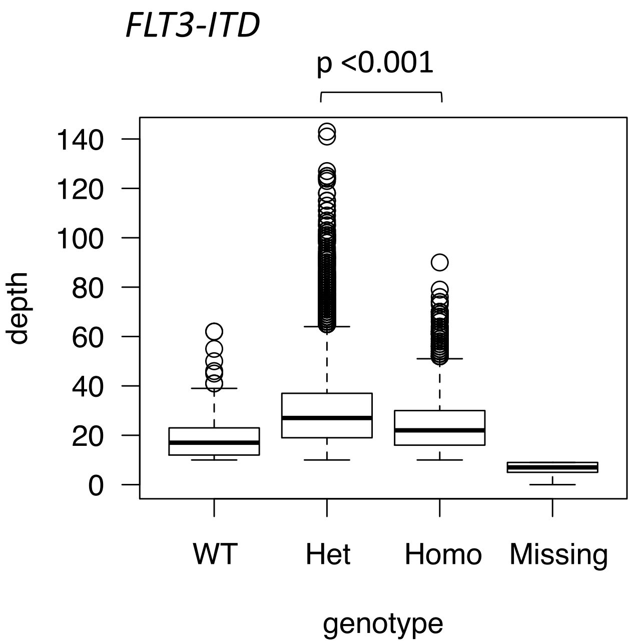

AML-17-001

Karyotype: 46,XY[20]

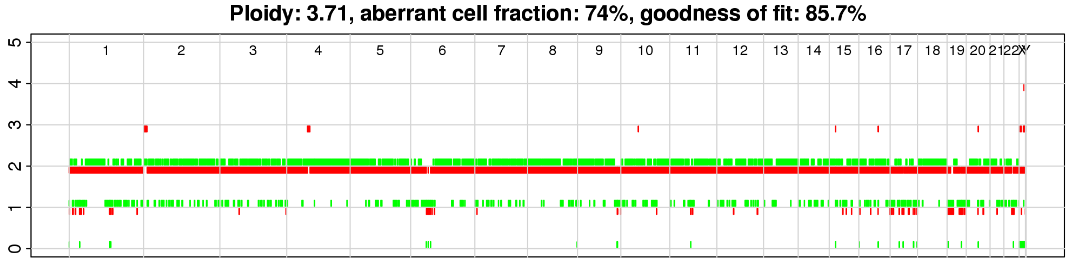


| chr | Gene | AA change | WT | Het | Homo | Missing | Homo [%] | allele count |
|---|---|---|---|---|---|---|---|---|
| 2 | *DNMT3A* | p.F868L | 571 | 4564 | 262 | 1227 | 5 | 2,1 |
| 1 | *NRAS* | p.G12D | 489 | 4576 | 125 | 1434 | 3 | 2,1 |
| 5 | *NPM1* | p.L287fs | 641 | 3265 | 336 | 2382 | 9 | 2,2 |

#### Heatmap ignoring zygosity

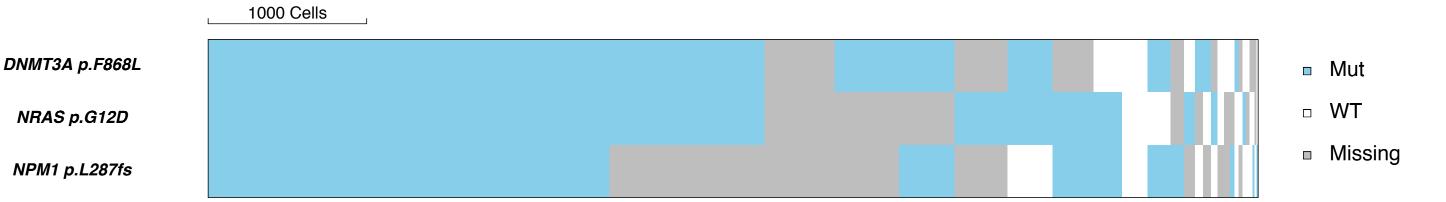


#### Heatmap considering zygosity if SNP array-confirmed

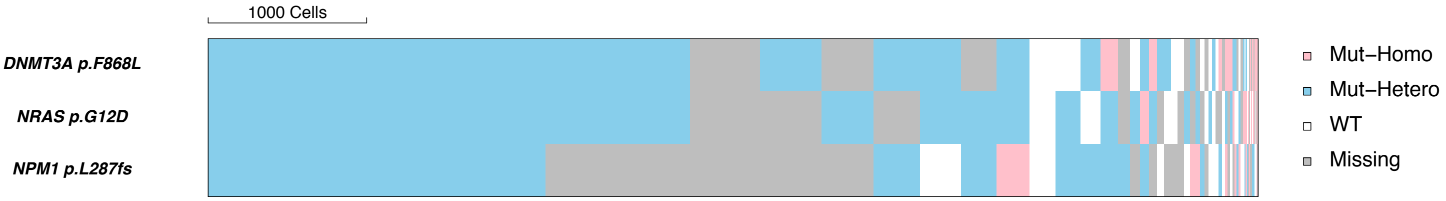

AML-07-001

Karyotype: 46,XY[19]

Ploidy: 5.04, aberrant cell fraction: 64%, goodness of fit: 84.2%

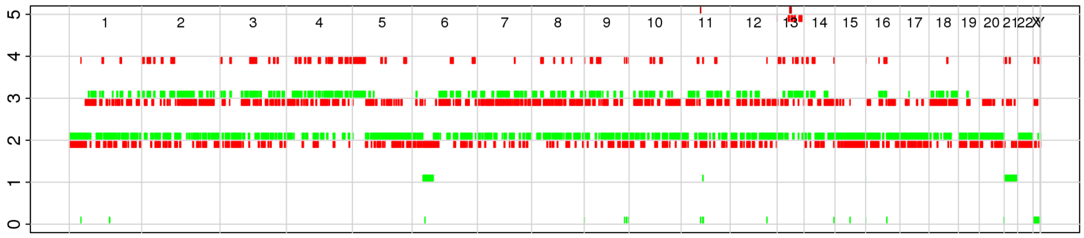

| chr | Gene | AA change | WT | Het | Homo | Missing | Homo [%] | allele count |
|---|---|---|---|---|---|---|---|---|
| 15 | *IDH2* | p.R140Q | 487 | 5952 | 77 | 202 | 1 | 2,2 |
| 17 | *SRSF2* | p.P95H | 334 | 2744 | 545 | 3095 | 17 | 2,2 |
| 21 | *RUNX1* | p.D123fs | 1152 | 199 | 4546 | 821 | 96 | 2,1 |

#### Heatmap ignoring zygosity

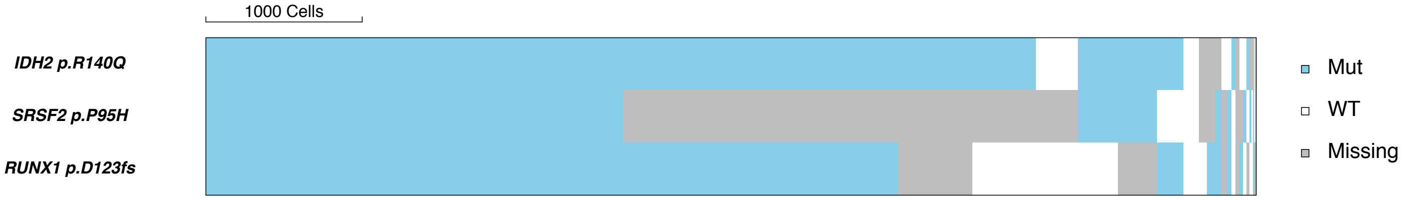


#### Heatmap considering zygosity if SNP array-confirmed

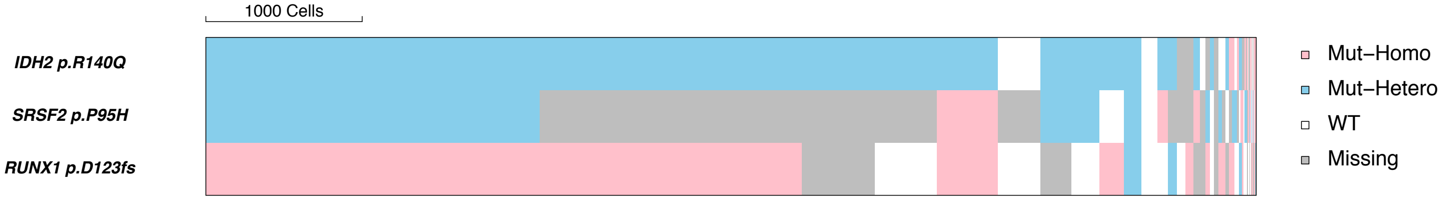


#### Distribution of depth for *SRSF2* p.P95H and *RUNX1* p.D123fs based on genotype

*SRSF2* p.P95H

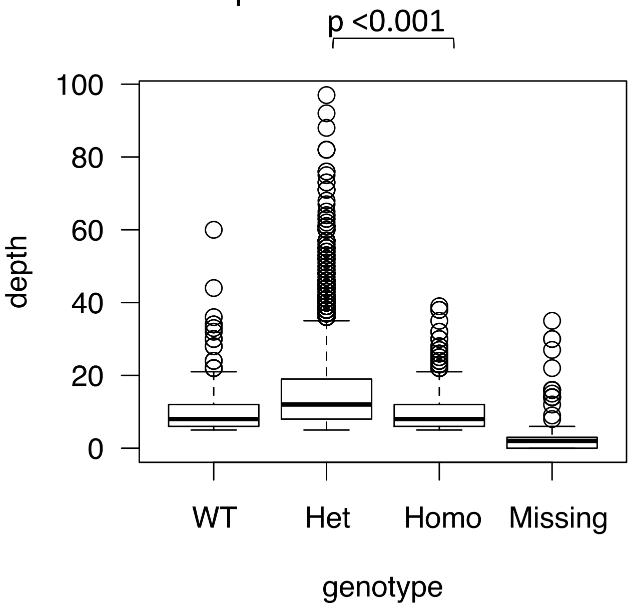


*RUNX1* p.D123fs

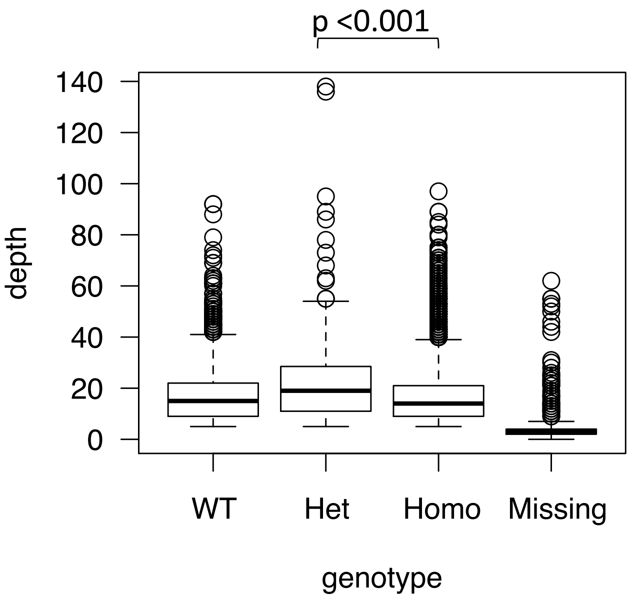

AML-40-001

Karyotype: 46,XY[20]

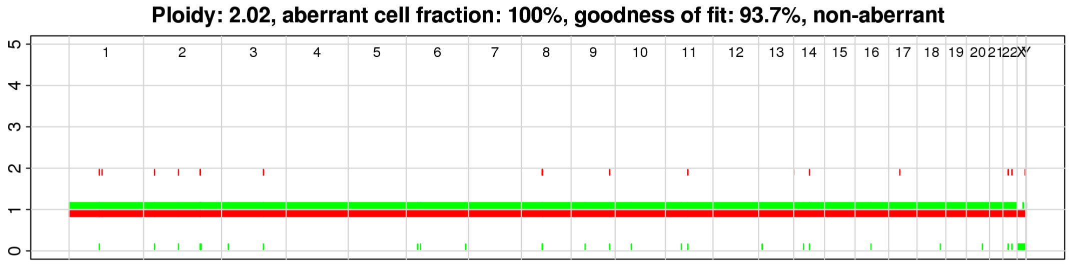


| chr | Gene | AA change | WT | Het | Homo | Missing | Homo [%] | allele count |
|---|---|---|---|---|---|---|---|---|
| 15 | *IDH2* | p.R140Q | 785 | 4560 | 441 | 408 | 9 | 1,1 |
| 5 | *NPM1* | p.L287fs | 736 | 2344 | 406 | 2708 | 15 | 1,1 |
| 1 | *NRAS* | p.G12D | 3343 | 1705 | 85 | 1061 | 5 | 1,1 |
| 1 | *NRAS* | p.G13D | 5225 | 38 | 1 | 930 | 3 | 1,1 |
| 13 | *FLT3* | c.C2028G p.N676K | 5661 | 370 | 24 | 139 | 6 | 1,1 |
| 13 | *FLT3* | c.C2028A p.N676K | 6008 | 42 | 5 | 139 | 11 | 1,1 |
| 13 | *FLT3* | ITD | 5436 | 34 | 3 | 721 | 8 | 1,1 |

Distribution of depth for *NPM1* p.L287fs based on genotype

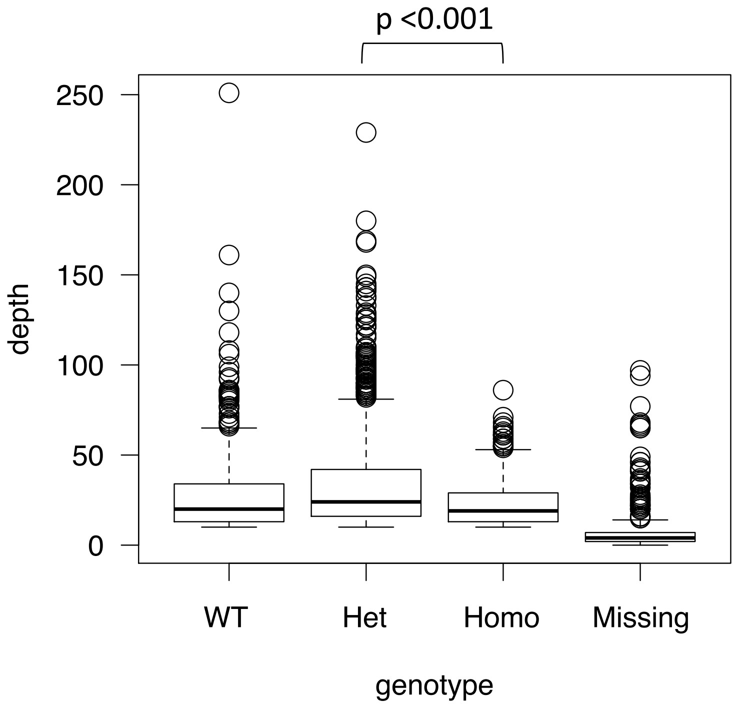

AML-21-001

Karyotype: 46,XX[20]

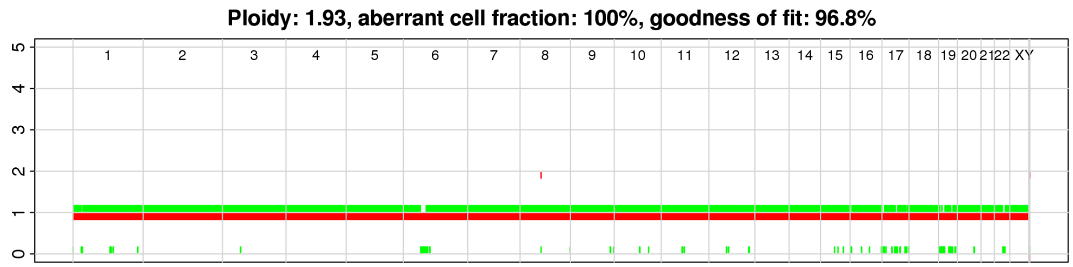


| chr | Gene | AA change | WT | Het | Homo | Missing | Homo [%] | allele count |
|---|---|---|---|---|---|---|---|---|
| 11 | *WT1* | p.A382fs | 484 | 7567 | 154 | 156 | 2 | 1,1 |
| 5 | *NPM1* | p.L287fs | 1093 | 3844 | 411 | 3013 | 10 | 1,1 |
| 1 | *NRAS* | p.G12D | 624 | 4841 | 111 | 2785 | 2 | 1,1 |

Distribution of depth for *NPM1* p.L287fs based on genotype

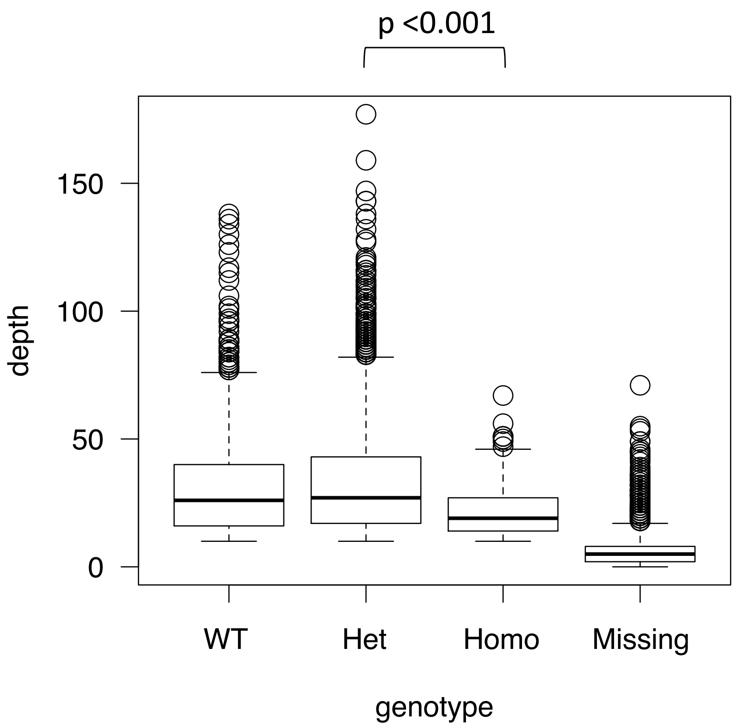

AML-58-001 Karyotype: 46,XY[20]

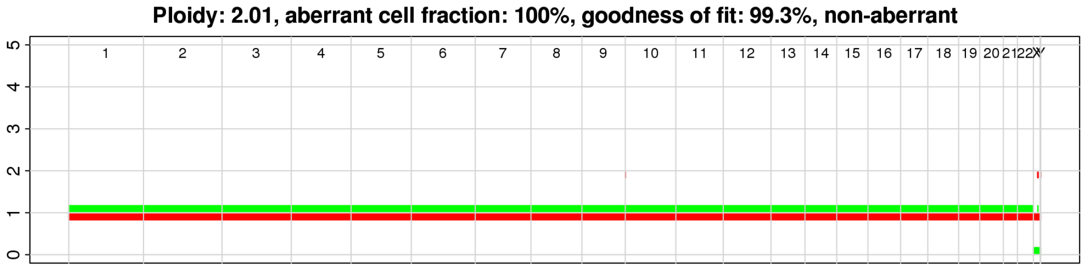


| chr | Gene | AA change | WT | Het | Homo | Missing | Homo [%] | allele count |
|---|---|---|---|---|---|---|---|---|
| 15 | *IDH2* | p.R140Q | 2421 | 5293 | 101 | 355 | 2 | 1,1 |
| 2 | *DNMT3A* | p.R882H | 797 | 4912 | 113 | 2348 | 2 | 1,1 |
| 17 | *SRSF2* | p.P95H | 541 | 1377 | 482 | 5770 | 26 | 1,1 |

Distribution of depth for *SRSF2* p.P95H based on genotype

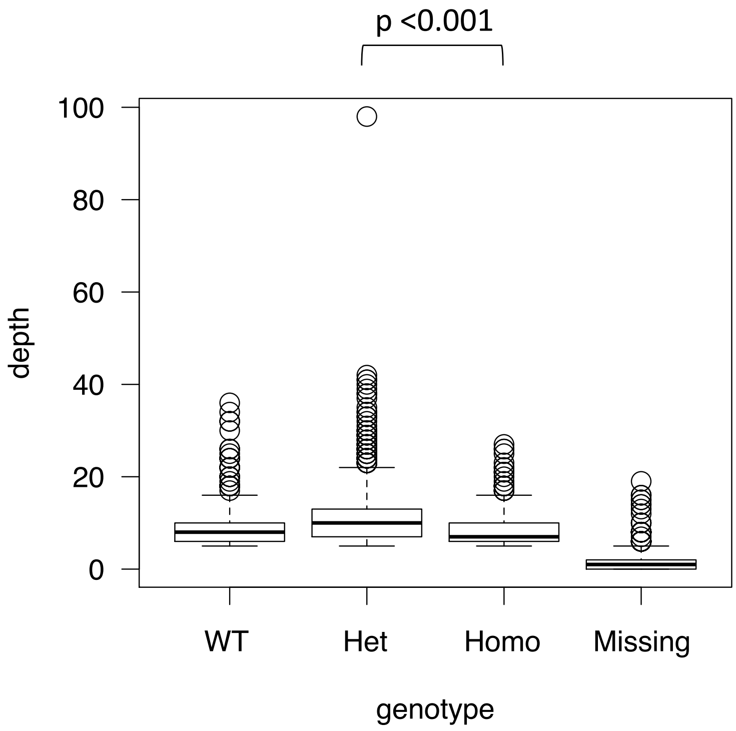

AML-29-001

Karyotype: 46,XY[20]

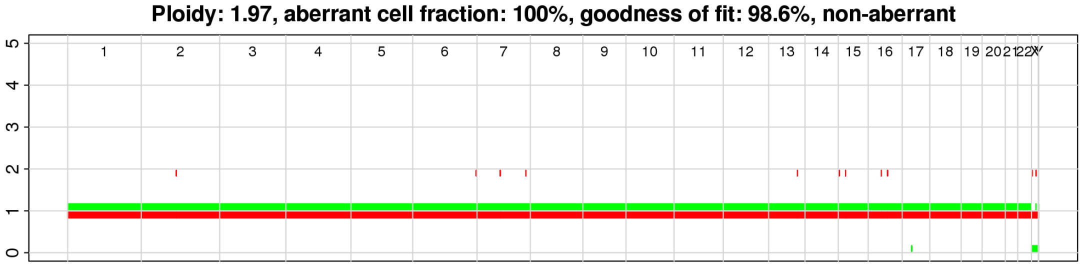


| chr | Gene | AA change | WT | Het | Homo | Missing | Homo [%] | allele count |
|---|---|---|---|---|---|---|---|---|
| 20 | *ASXL1* | p.G642fs | 477 | 5983 | 193 | 1142 | 3 | 1,1 |
| 17 | *SRSF2* | p.P95H | 538 | 2379 | 798 | 4080 | 25 | 1,1 |
| 21 | *RUNX1* | p.R201Q | 340 | 1708 | 170 | 5577 | 9 | 1,1 |
| 15 | *IDH2* | p.R140Q | 7569 | 176 | 6 | 44 | 3 | 1,1 |

Distribution of depth for *SRSF2* p.P95H based on genotype

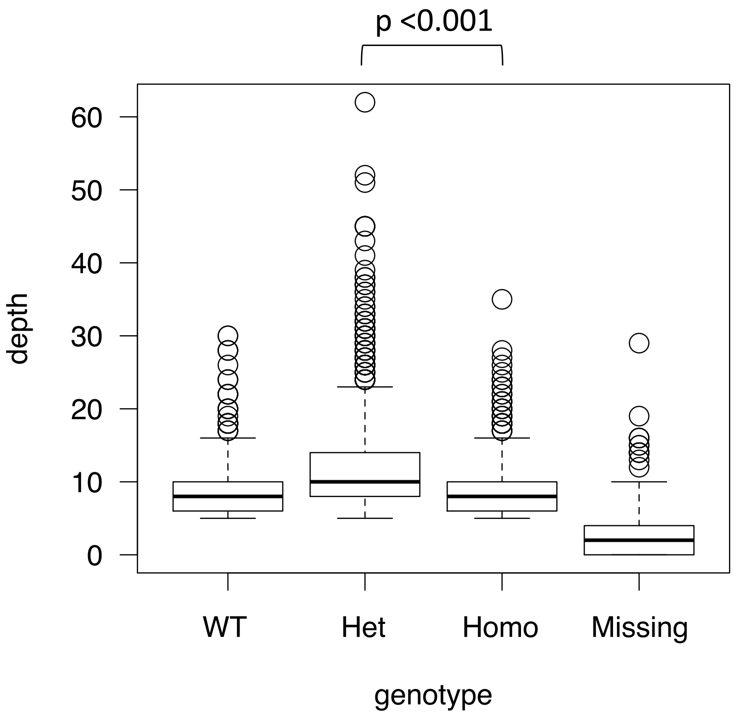

AML-54-001 Karyotype: 46,XY[20]

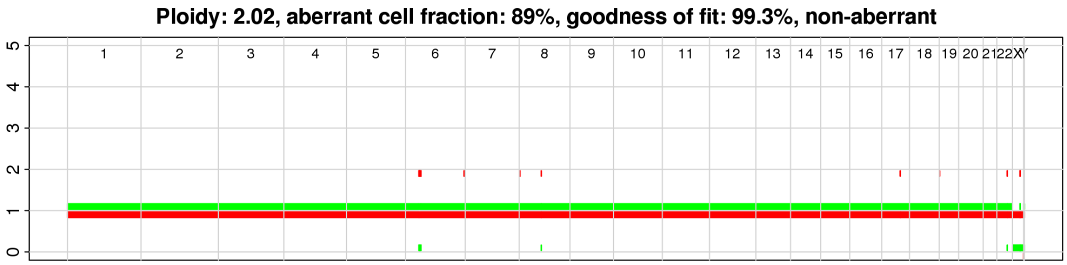


| chr | Gene | AA change | WT | Het | Homo | Missing | Homo [%] | allele count |
|---|---|---|---|---|---|---|---|---|
| 17 | *SRSF2* | p.P95R | 546 | 1250 | 339 | 5681 | 21 | 1,1 |
| 20 | *ASXL1* | p.G642fs | 1032 | 2178 | 3 | 4603 | 0 | 1,1 |
| 13 | *FLT3* | ITD | 387 | 6628 | 286 | 515 | 4 | 1,1 |

Distribution of depth for *SRSF2* p.P95R based on genotype

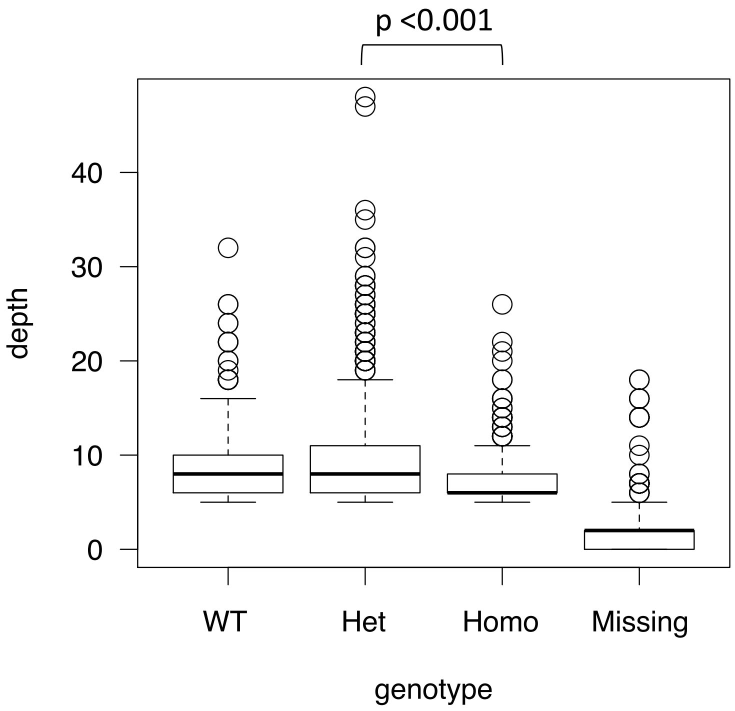

AML-70-001

Karyotype: 48,XY,+13,+14[1]/46,XY[19]

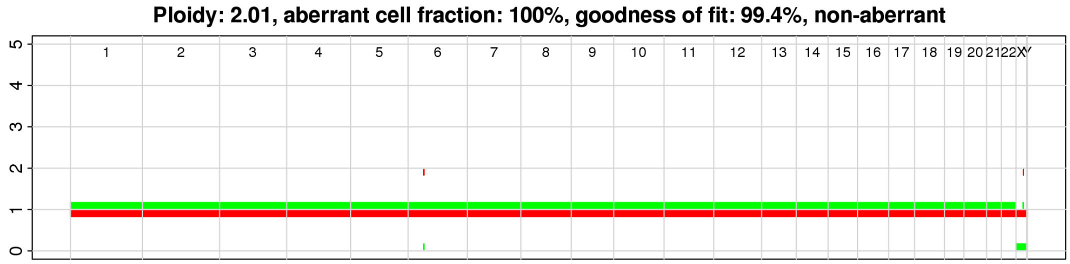


| chr | Gene | AA change | WT | Het | Homo | Missing | Homo [%] | allele count |
|---|---|---|---|---|---|---|---|---|
| 15 | *IDH2* | p.R140Q | 1033 | 6660 | 270 | 305 | 4 | 1,1 |
| 17 | *SRSF2* | p.P95L | 1053 | 1079 | 121 | 6015 | 10 | 1,1 |
| 12 | *KRAS* | p.D33E | 1269 | 6070 | 475 | 454 | 7 | 1,1 |
| 1 | *NRAS* | p.G60E | 8011 | 152 | 9 | 96 | 6 | 1,1 |

Distribution of depth for *SRSF2* p.P95L based on genotype

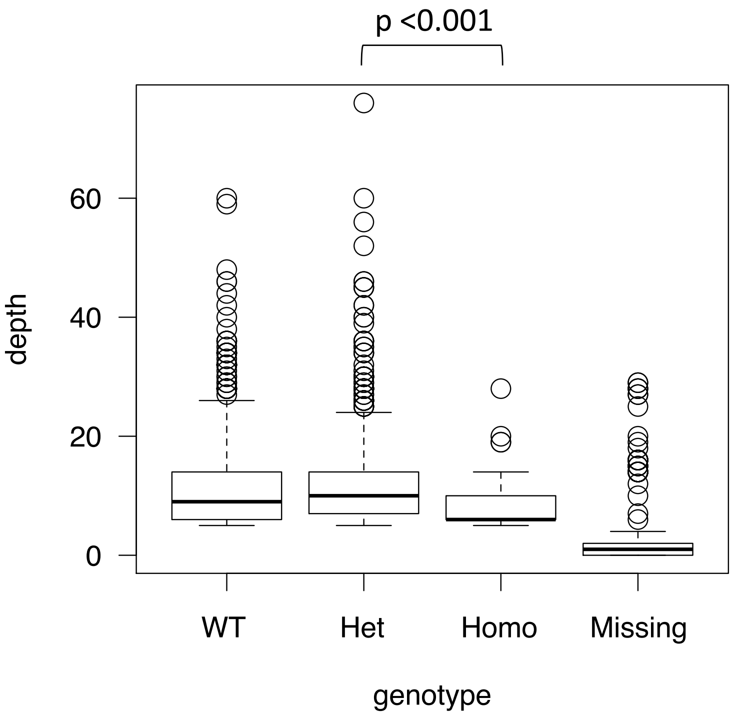

AML-11-001

Karyotype: 46,XY[20]

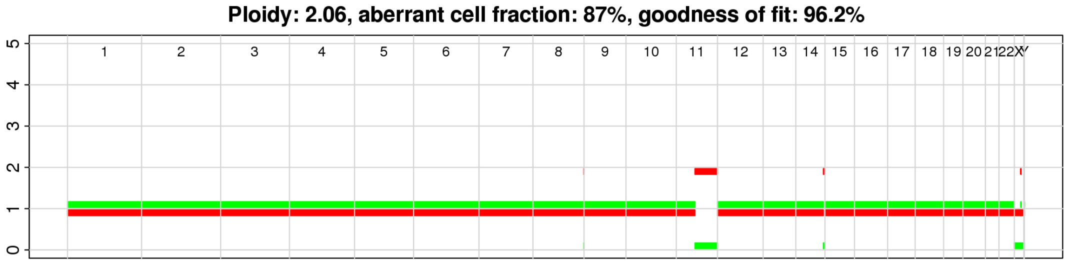


| chr | Gene | AA change | WT | Het | Homo | Missing | Homo [%] | allele count |
|---|---|---|---|---|---|---|---|---|
| 17 | *SRSF2* | p.P95A | 251 | 759 | 353 | 3911 | 32 | 1,1 |
| 20 | *ASXL1* | p.G642fs | 693 | 1252 | 2 | 3327 | 0 | 1,1 |

Distribution of depth for *SRSF2* p.P95A based on genotype

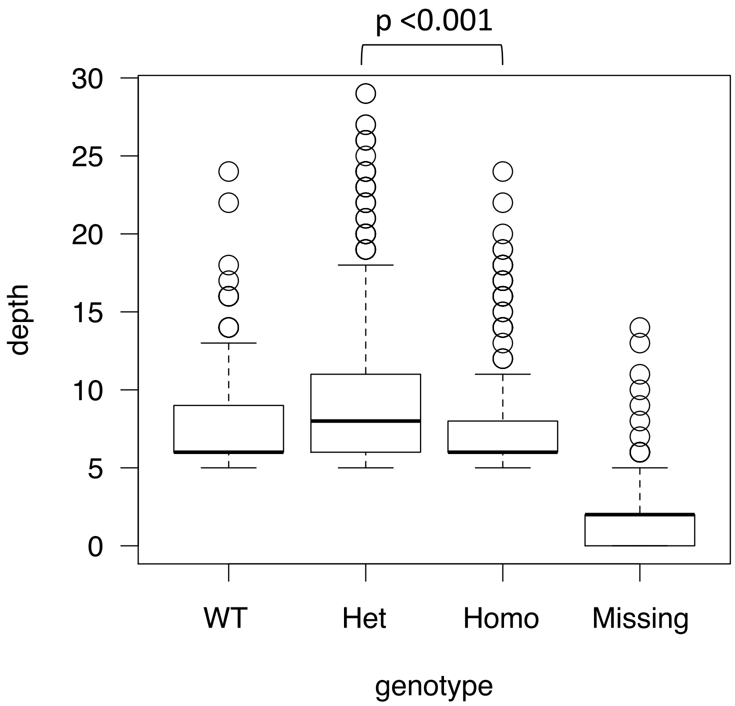

AML-60-001 Karyotype: 46,XY[20]

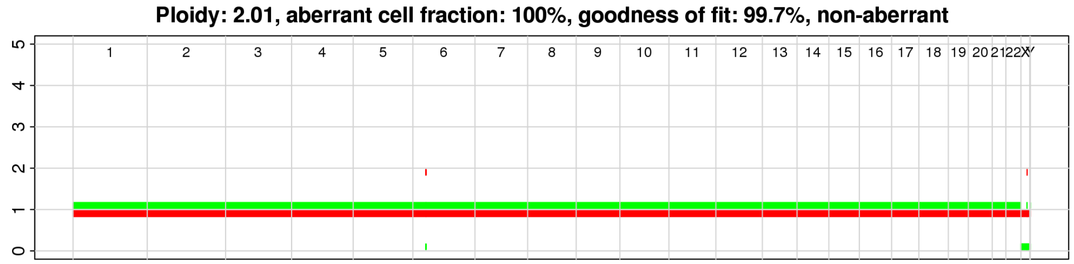


| chr | Gene | AA change | WT | Het | Homo | Missing | Homo [%] | allele count |
|---|---|---|---|---|---|---|---|---|
| 20 | *ASXL1* | p.G642fs | 648 | 4929 | 55 | 1983 | 1 | 1,1 |
| 15 | *IDH2* | p.R140Q | 362 | 6798 | 336 | 119 | 5 | 1,1 |
| 17 | *SRSF2* | p.P95H | 397 | 1797 | 590 | 4831 | 25 | 1,1 |
| 13 | *FLT3* | ITD | 6027 | 97 | 176 | 1315 | 64 | 1,1 |

Distribution of depth for *FLT3-ITD* and *SRSF2* p.P95R based on genotype

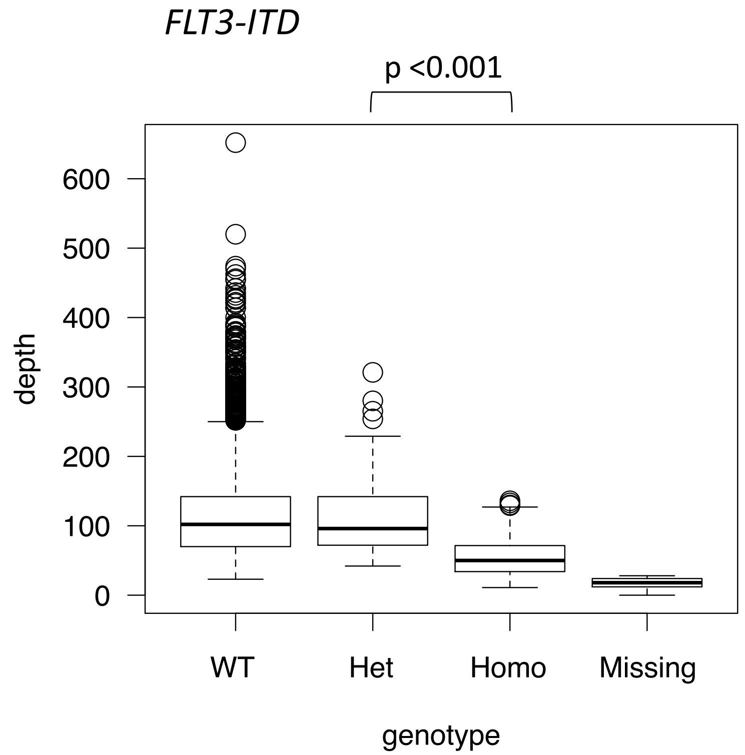


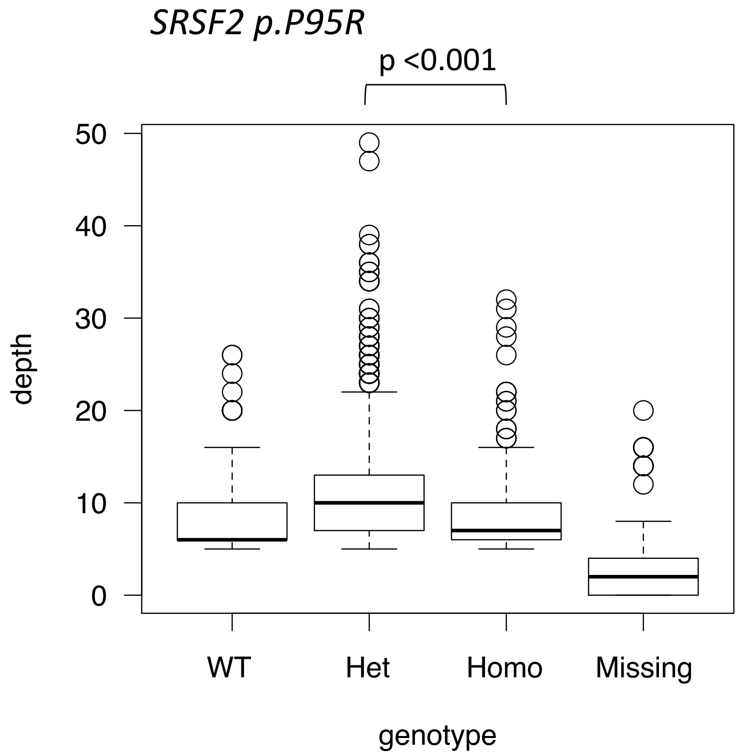

AML-53-001

Karyotype: 46,XX[20]

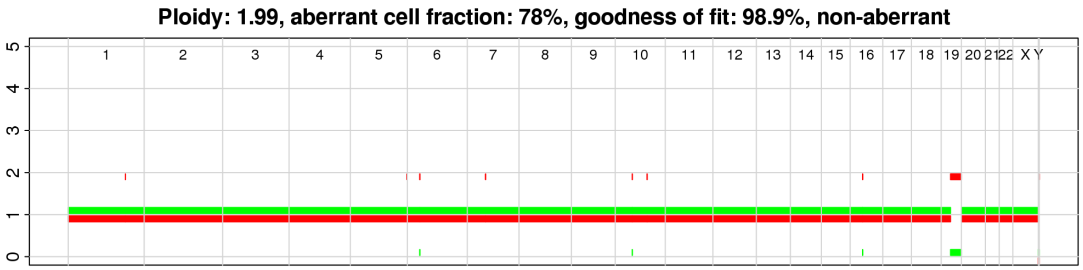


| chr | Gene | AA change | WT | Het | Homo | Missing | Homo [%] | allele count |
|---|---|---|---|---|---|---|---|---|
| 1 | *NRAS* | p.G12D | 6233 | 774 | 54 | 952 | 7 | 1,1 |
| 1 | *NRAS* | p.G13R | 7008 | 83 | 10 | 912 | 11 | 1,1 |
| 13 | *FLT3* | ITD | 7313 | 482 | 33 | 185 | 6 | 1,1 |

Distribution of depth for *NRAS* p.G13R based on genotype

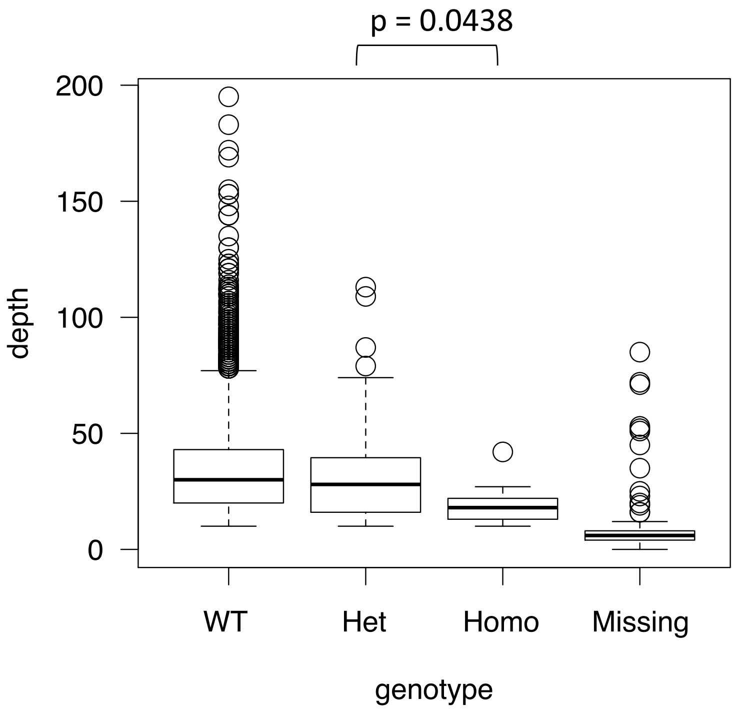

### AML-07-002

Karyotype: 91,XXY,-Y,-5,+13,-18,-21,+2mar[1], 46,XY[19] (tested 3 weeks after sample collection date)

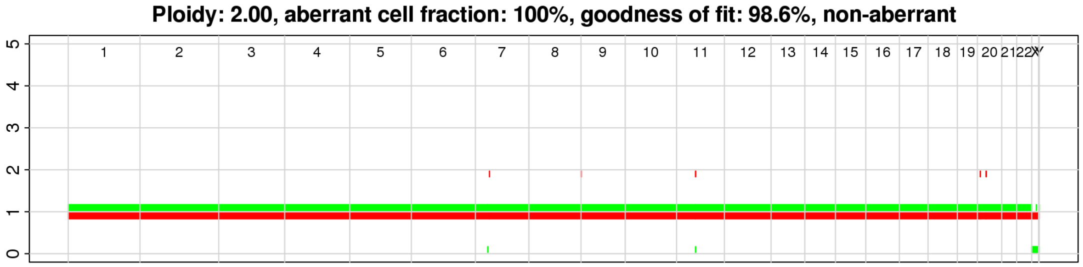


| chr | Gene | AA change | WT | Het | Homo | Missing | Homo [%] | allele count |
|---|---|---|---|---|---|---|---|---|
| 15 | *IDH2* | p.R140Q | 401 | 7664 | 145 | 252 | 2 | 1,1 |
| 17 | *SRSF2* | p.P95H | 303 | 2276 | 681 | 5202 | 23 | 1,1 |
| 21 | *RUNX1* | p.D123fs | 5624 | 4 | 24 | 2810 | 86 | 1,1 |

Distribution of depth for *RUNX1* p.D123fs and *SRSF2* p.P95H based on genotype

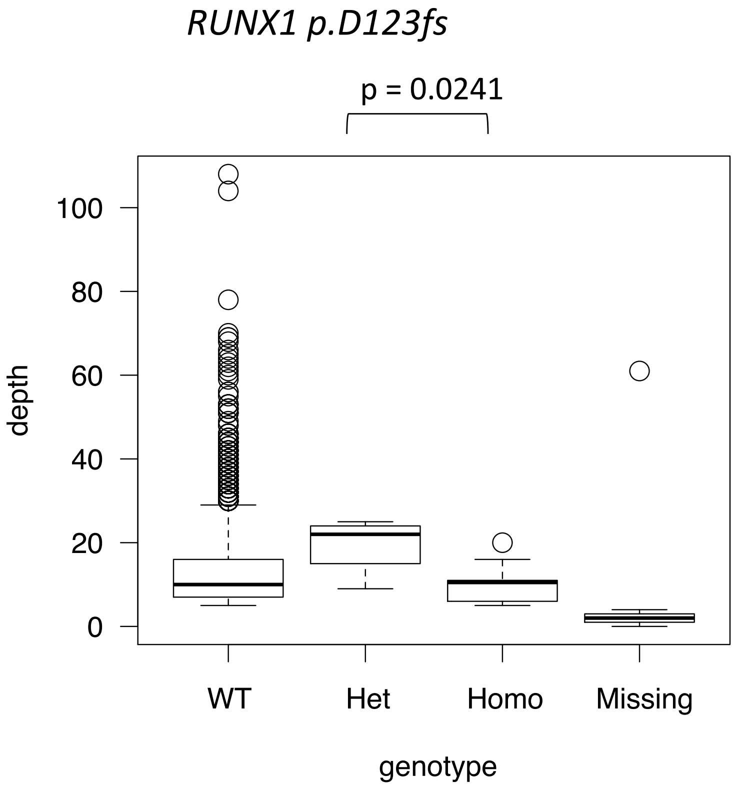


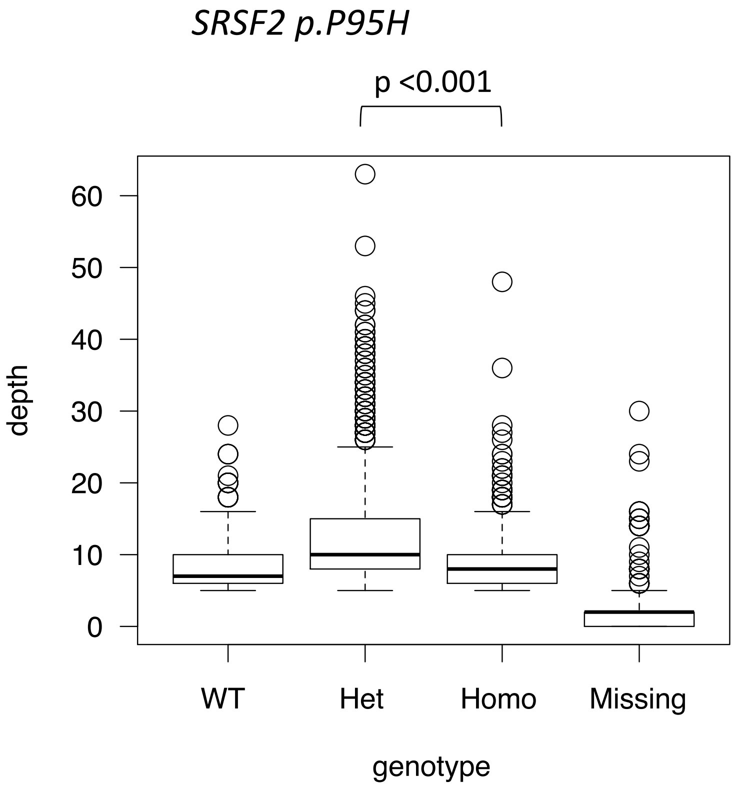

AML-02-001

Karyotype: 46,XY[20]

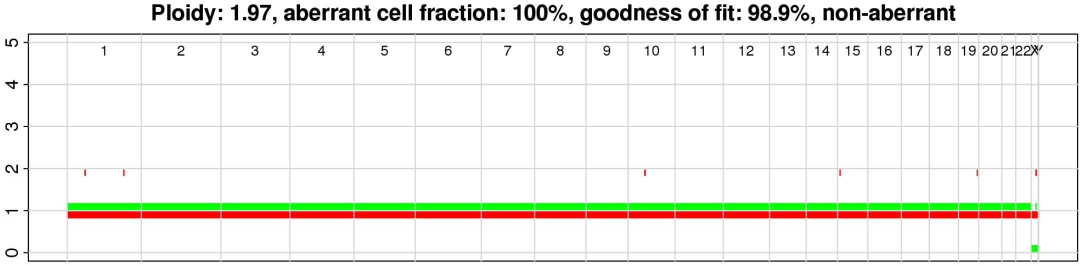


| chr | Gene | AA change | WT | Het | Homo | Missing | Homo [%] | allele count |
|---|---|---|---|---|---|---|---|---|
| 2 | *SF3B1* | p.K666N | 1038 | 6517 | 237 | 139 | 4 | 1,1 |
| 2 | *DNMT3A* | p.R882H | 468 | 5861 | 169 | 1433 | 3 | 1,1 |
| 13 | *FLT3* | p.D835V | 1186 | 6335 | 227 | 183 | 3 | 1,1 |
| 21 | *RUNX1* | p.S141fs | 1052 | 3971 | 287 | 2621 | 7 | 1,1 |
| 9 | *JAK2* | p.V617F | 3165 | 40 | 119 | 4607 | 75 | 1,1 |

Distribution of depth for *JAK2* p.V617F based on genotype

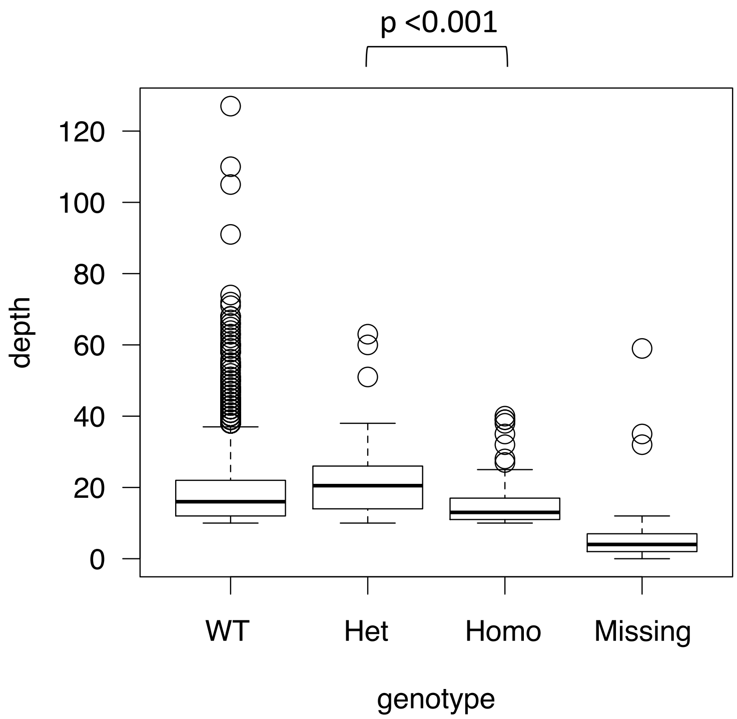

AML-08-001

Karyotype: 46,XY[20]

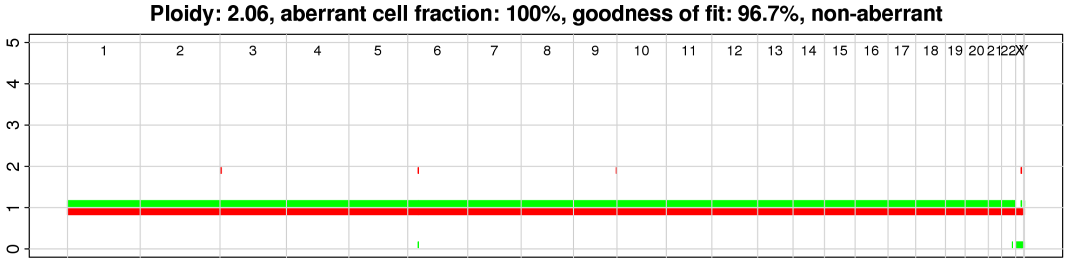


| chr | Gene | AA change | WT | Het | Homo | Missing | Homo [%] | allele count |
|---|---|---|---|---|---|---|---|---|
| 15 | *IDH2* | p.R140Q | 524 | 3912 | 112 | 127 | 3 | 1,1 |
| 2 | *DNMT3A* | p.R882H | 706 | 3246 | 130 | 593 | 4 | 1,1 |
| 17 | *SRSF2* | p.P95L | 936 | 1149 | 118 | 2472 | 9 | 1,1 |

AML-37-001

Karyotype: 46,XX[20]

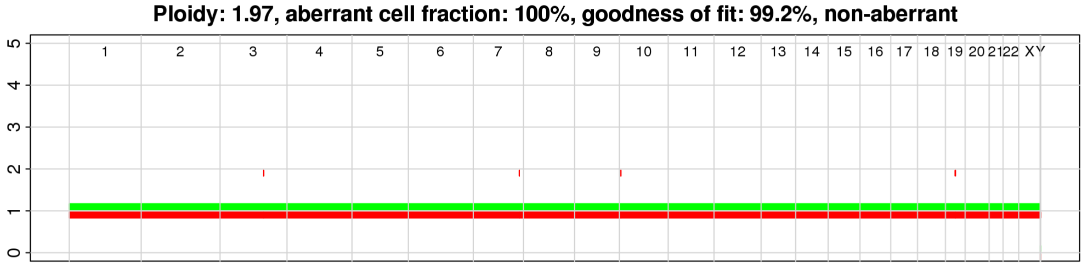


| chr | Gene | AA change | WT | Het | Homo | Missing | Homo [%] | allele count |
|---|---|---|---|---|---|---|---|---|
| 2 | *DNMT3A* | p.N879D | 469 | 4924 | 186 | 665 | 4 | 1,1 |
| 5 | *NPM1* | p.L287fs | 498 | 3657 | 187 | 1902 | 5 | 1,1 |
| 1 | *NRAS* | p.G12D | 474 | 4887 | 146 | 737 | 3 | 1,1 |

AML-20-001

Karyotype: 46,XY,del(1)(p21)[1], 46,XY[19]

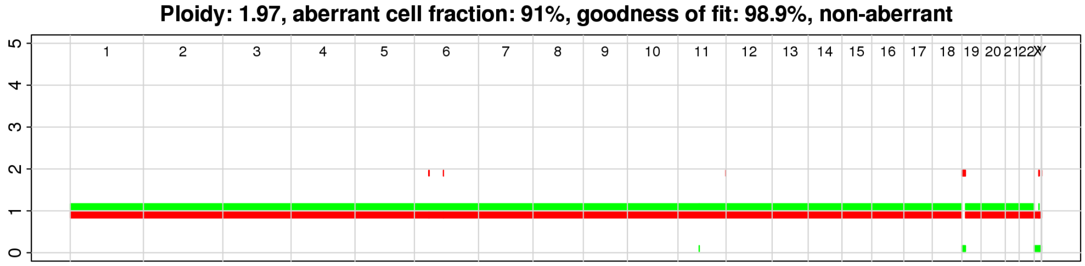


| chr | Gene | AA change | WT | Het | Homo | Missing | Homo [%] | allele count |
|---|---|---|---|---|---|---|---|---|
| 2 | *DNMT3A* | p.R882C | 1516 | 6048 | 117 | 2369 | 2 | 1,1 |
| 5 | *NPM1* | p.L287fs | 2967 | 2932 | 161 | 3990 | 5 | 1,1 |
| 1 | *NRAS* | p.G13V | 3530 | 3260 | 54 | 3206 | 2 | 1,1 |

AML-50-001

Karyotype: 46,XX[20]

Ploidy: 2.01, aberrant cell fraction: 100%, goodness of fit: 99.2%, non-aberrant

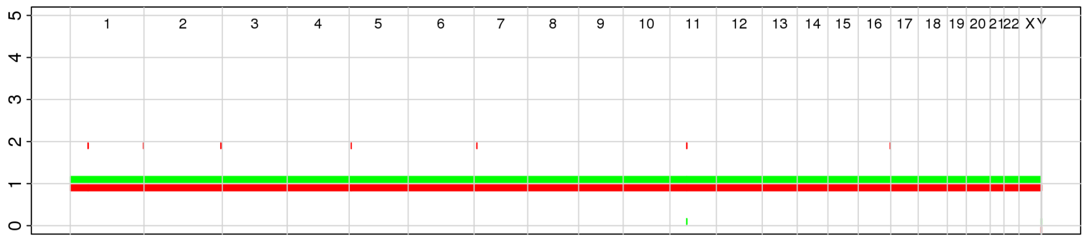

| chr | Gene | AA change | WT | Het | Homo | Missing | Homo [%] | allele count |
|---|---|---|---|---|---|---|---|---|
| 5 | *NPM1* | p.L287fs | 1141 | 4843 | 443 | 2980 | 8 | 1,1 |
| 13 | *FLT3* | p.Y572C | 887 | 6615 | 98 | 1807 | 1 | 1,1 |

AML-63-001

Karyotype: 46,XX[20]

Ploidy: 1.93, aberrant cell fraction: 52%, goodness of fit: 82.2%, non-aberrant

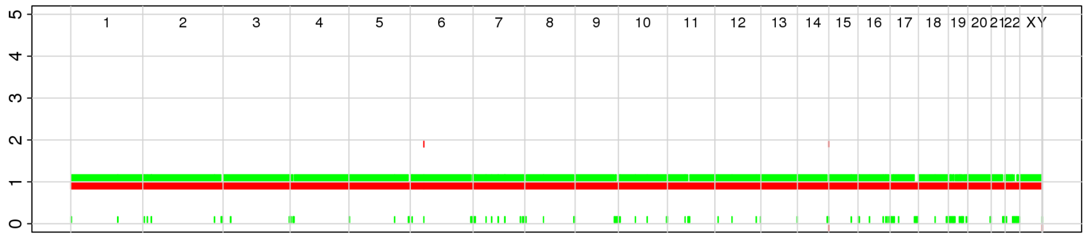

| chr | Gene | AA change | WT | Het | Homo | Missing | Homo [%] | allele count |
|---|---|---|---|---|---|---|---|---|
| 15 | *IDH2* | p.R140Q | 663 | 6709 | 306 | 669 | 4 | 1,1 |
| 5 | *NPM1* | p.L287fs | 734 | 3921 | 365 | 3327 | 9 | 1,1 |
| 4 | *KIT* | p.D816V | 7120 | 827 | 57 | 343 | 6 | 1,1 |
| 13 | *FLT3* | p.S585fs | 6484 | 914 | 44 | 905 | 5 | 1,1 |

AML-59-001

Karyotype: 47,XX,+11[3]/47,idem,del(7)(q22q34)[16]/46,idem,-7[1]

Ploidy: 2.08, aberrant cell fraction: 62%, goodness of fit: 95.6%

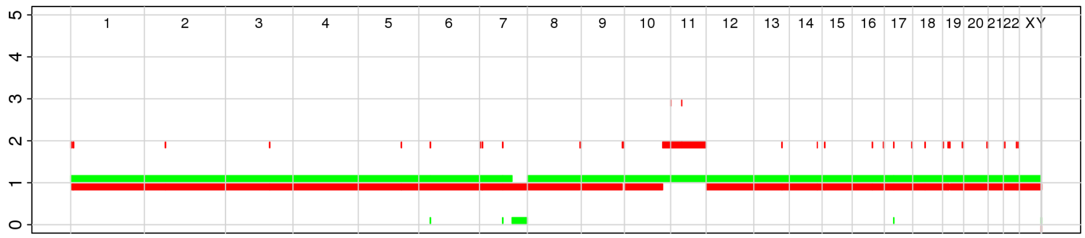

| chr | Gene | AA change | WT | Het | Homo | Missing | Homo [%] | allele count |
|---|---|---|---|---|---|---|---|---|
| 2 | *IDH1* | p.R132C | 278 | 2281 | 75 | 28 | 3 | 1,1 |
| 2 | *DNMT3A* | p.R882H | 488 | 1549 | 89 | 536 | 5 | 1,1 |
| 21 | *RUNX1* | p.K152fs | 584 | 1645 | 123 | 310 | 7 | 1,1 |
| 21 | *RUNX1* | p.D198N | 1765 | 130 | 2 | 765 | 2 | 1,1 |

**Extended Data Fig. 11. Analysis of copy-number alterations by SNP array in selected samples.** Copy-number alteration (CNA) data was retrieved from an Illumina Omni2.5-8 SNP array using the ASCAT algorithm. The karyotype as determined by G-banding is shown alongside the sample ID. In the top figure, the green and red lines each represent an allele. The vertical axis shows the allele count, and the horizontal axis shows the chromosomes. The table below the top figure summarizes the number of cells with each genotype. The percentage of homozygously mutated cells (Homo [%]) was calculated as follows: (the number of homozygously called cells) / (the total number of mutated cells [including heterozygously called and homozygously called cells]) × 100. Highly homozygous variants (Homo [%] >10) and variants involving CNA are highlighted in red. Heat maps were updated to incorporate the zygosity information when homozygously called variants were located within the loci involving CNA. For highly homozygous variants, box plots show the distribution of depth for each cell grouped by the genotype. The thick line within each box represents the median, and the top and bottom edges represent the 25th and 75th percentiles, respectively. The upper and lower whiskers represent the 75th percentile plus 1.5 times the interquartile range and the 25th percentile minus 1.5 times the interquartile range, respectively. The dots represent the actual depth data.

**AML-71-001, AML-32-001**

The SNP array detected copy-neutral loss of heterozygosity (CN-LOH) of the mutant loci of highly homozygous variants, suggesting that loss of wild-type allele and amplification of mutant allele had resulted in the homozygous call.

**AML-19-001, AML-44-001, AML-01-001, AML-17-001, AML-07-001**

The SNP array detected LOH likely with copy-number gain of the mutant allele. These data should be interpreted with caution because the SNP array data exhibit some noise.

**AML-40-001, AML-21-001, AML-58-001, AML-29-001, AML-54-001, AML-70-001, AML-11-001, AML-60-001, AML-53-001, AML-07-002, AML-02-001, AML-08-001, AML-37-001, AML-20-001, AML-50-001, AML-63-001, AML-59-001**

The SNP array did not detect LOH of the mutant loci where the highly homozygous variants were located. The depth for homozygously genotyped cells was significantly lower than for heterozygously genotyped cells, suggesting that these homozygous calls might have been affected by insufficient sequencing coverage or allele dropout.

AA change, amino acid change; WT, wild type; Het, heterozygous; Homo, homozygous; Missing, missing genotype; Homo [%], percentage of homozygously genotyped cells among total mutated cells.

### Extended Data Fig. 12

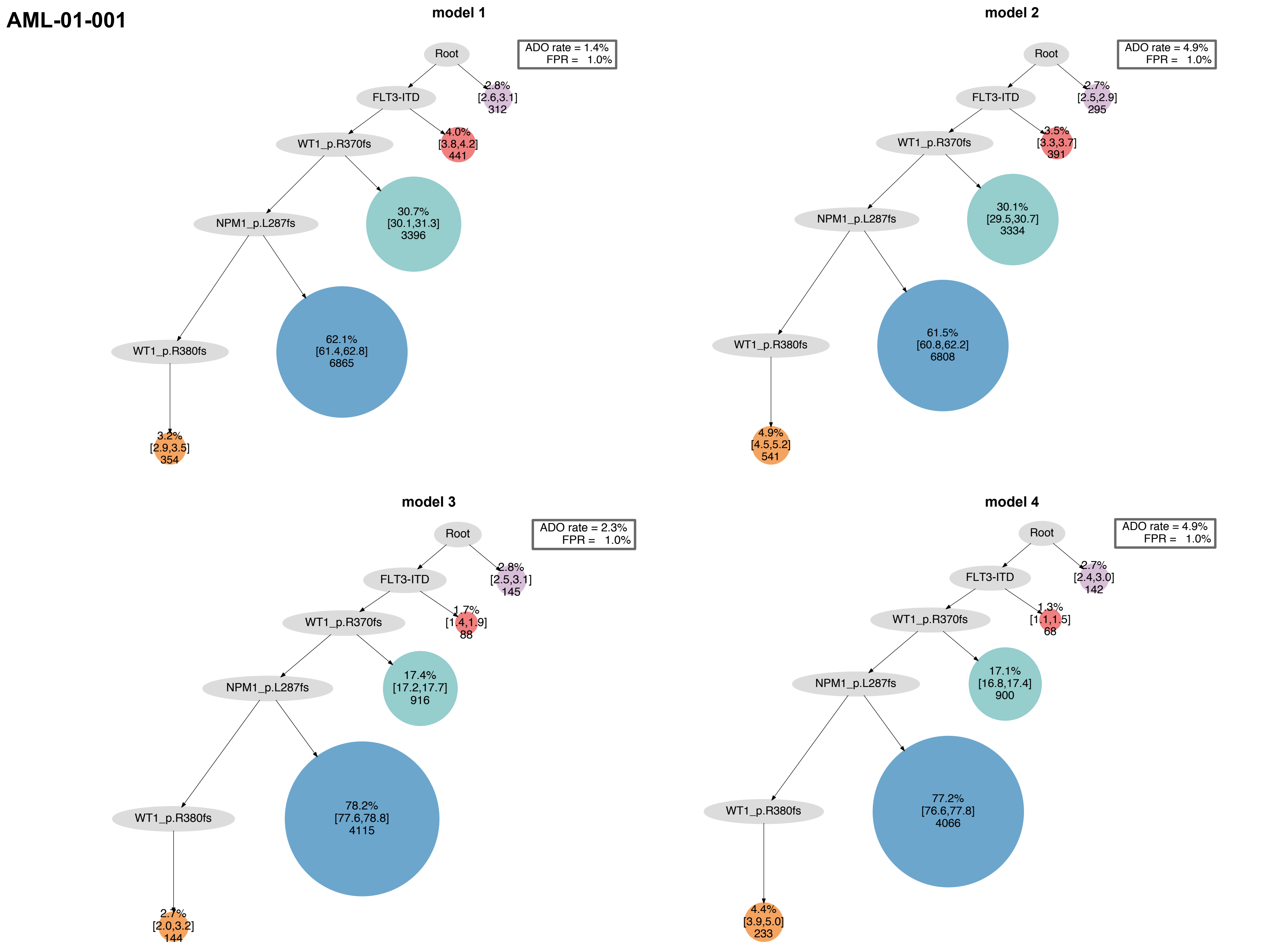

#### AML-02-001

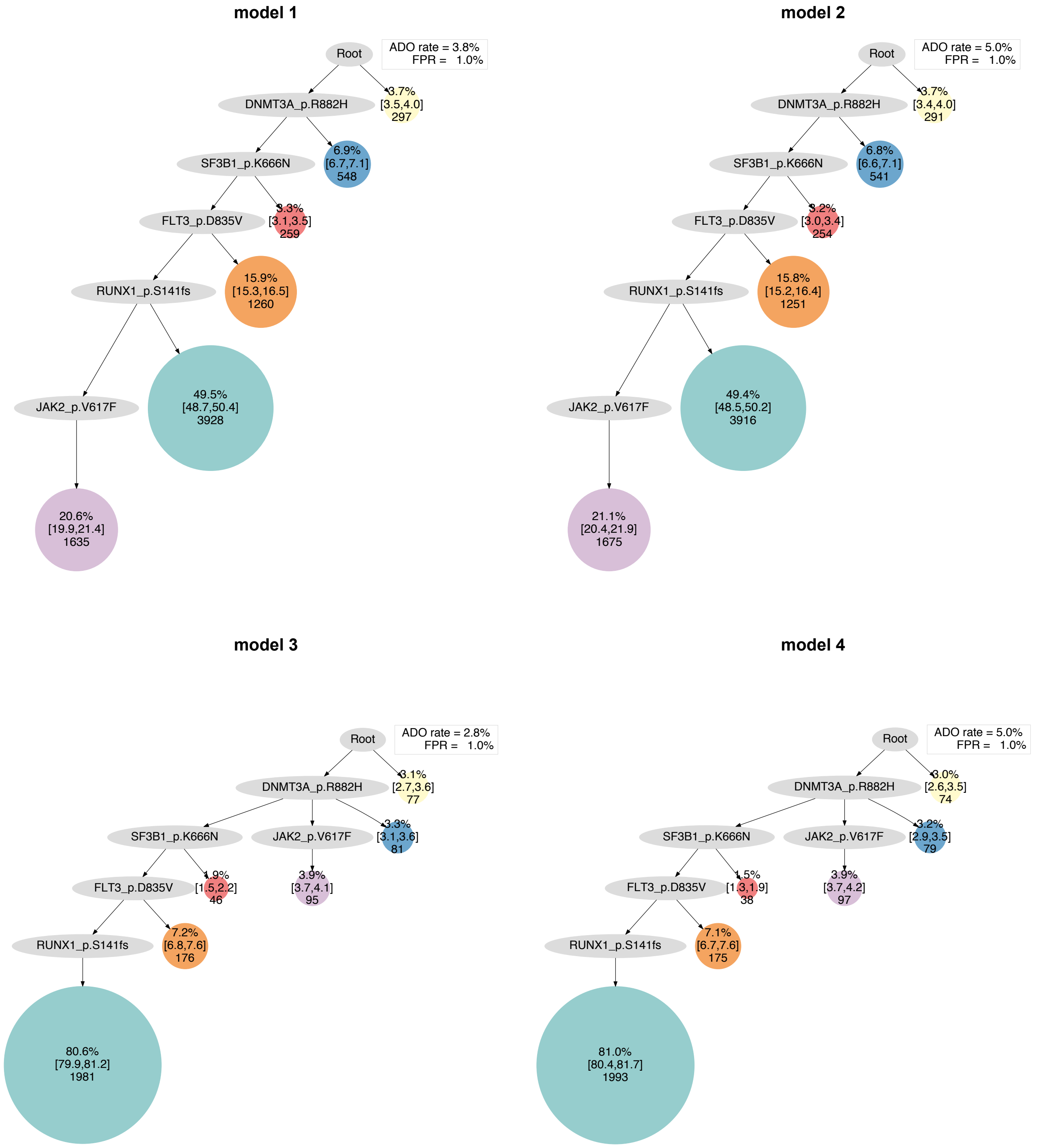

model 1
Root
ADO rate = 3.8%
FPR = 1.0%
DNMT3A_p.R882H
3.7%
[3.5,4.0]
297
SF3B1_p.K666N
6.9%
[6.7,7.1]
548
FLT3_p.D835V
3.3%
[3.1,3.5]
259
RUNX1_p.S141fs
15.9%
[15.3,16.5]
1260
JAK2_p.V617F
49.5%
[48.7,50.4]
3928
20.6%
[19.9,21.4]
1635
model 2
Root
ADO rate = 5.0%
FPR = 1.0%
DNMT3A_p.R882H
3.7%
[3.4,4.0]
291
SF3B1_p.K666N
6.8%
[6.6,7.1]
541
FLT3_p.D835V
3.2%
[3.0,3.4]
254
RUNX1_p.S141fs
15.8%
[15.2,16.4]
1251
JAK2_p.V617F
49.4%
[48.5,50.2]
3916
21.1%
[20.4,21.9]
1675
model 3
Root
ADO rate = 2.8%
FPR = 1.0%
DNMT3A_p.R882H
3.1%
[2.7,3.6]
77
SF3B1_p.K666N
JAK2_p.V617F
3.3%
[3.1,3.6]
81
FLT3_p.D835V
1.9%
[1.5,2.2]
46
3.9%
[3.7,4.1]
95
RUNX1_p.S141fs
7.2%
[6.8,7.6]
176
80.6%
[79.9,81.2]
1981
model 4
Root
ADO rate = 5.0%
FPR = 1.0%
DNMT3A_p.R882H
3.0%
[2.6,3.5]
74
SF3B1_p.K666N
JAK2_p.V617F
3.2%
[2.9,3.5]
79
FLT3_p.D835V
1.5%
[1.3,1.9]
38
3.9%
[3.7,4.2]
97
RUNX1_p.S141fs
7.1%
[6.7,7.6]
175
81.0%
[80.4,81.7]
1993

### AML-03-001

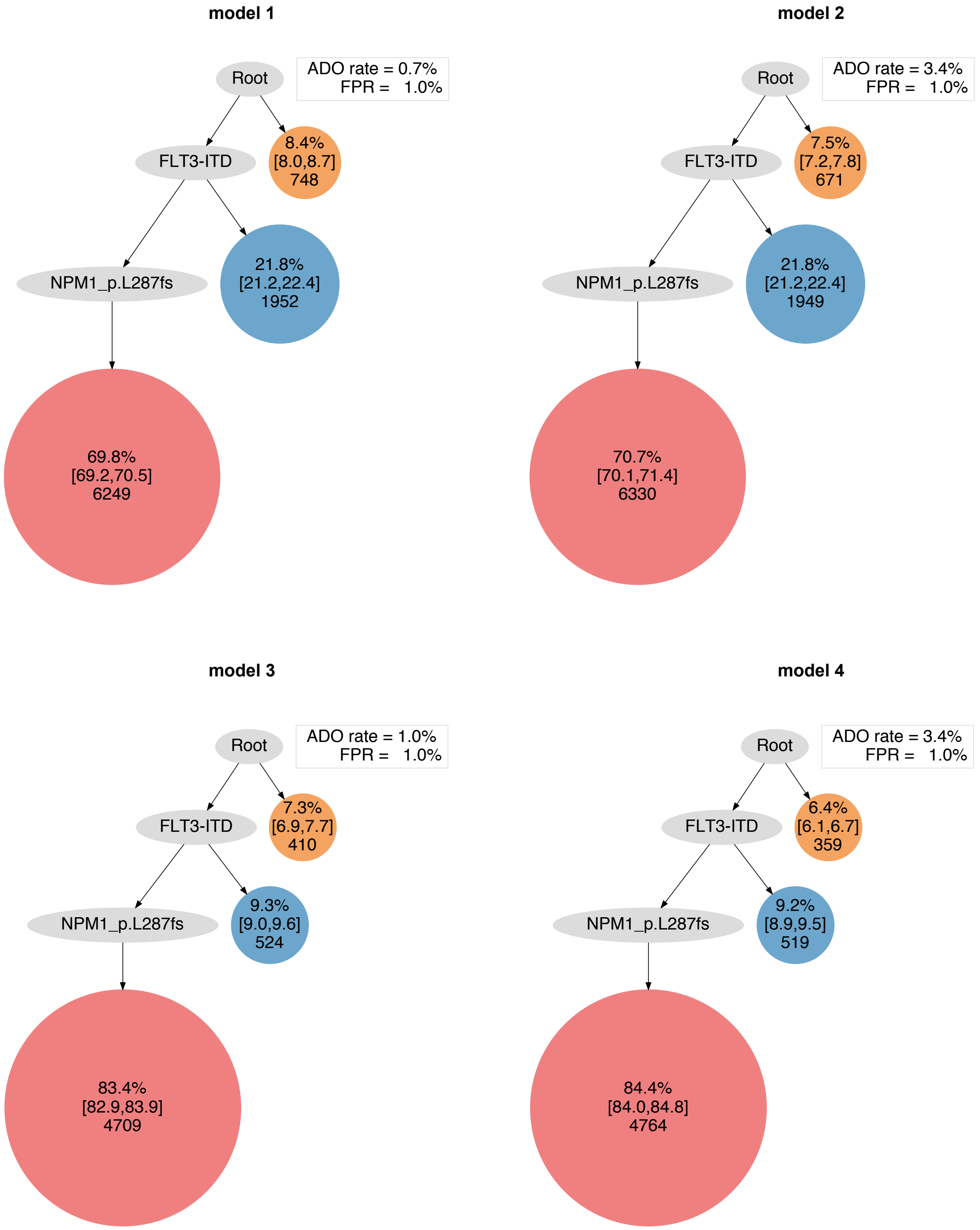

model 1
Root
ADO rate = 0.7%
FPR = 1.0%
FLT3-ITD
8.4%
[8.0,8.7]
748
NPM1_p.L287fs
21.8%
[21.2,22.4]
1952
69.8%
[69.2,70.5]
6249
model 2
Root
ADO rate = 3.4%
FPR = 1.0%
FLT3-ITD
7.5%
[7.2,7.8]
671
NPM1_p.L287fs
21.8%
[21.2,22.4]
1949
70.7%
[70.1,71.4]
6330
model 3
Root
ADO rate = 1.0%
FPR = 1.0%
FLT3-ITD
7.3%
[6.9,7.7]
410
NPM1_p.L287fs
9.3%
[9.0,9.6]
524
83.4%
[82.9,83.9]
4709
model 4
Root
ADO rate = 3.4%
FPR = 1.0%
FLT3-ITD
6.4%
[6.1,6.7]
359
NPM1_p.L287fs
9.2%
[8.9,9.5]
519
84.4%
[84.0,84.8]
4764

### AML-04-001

#### model 1

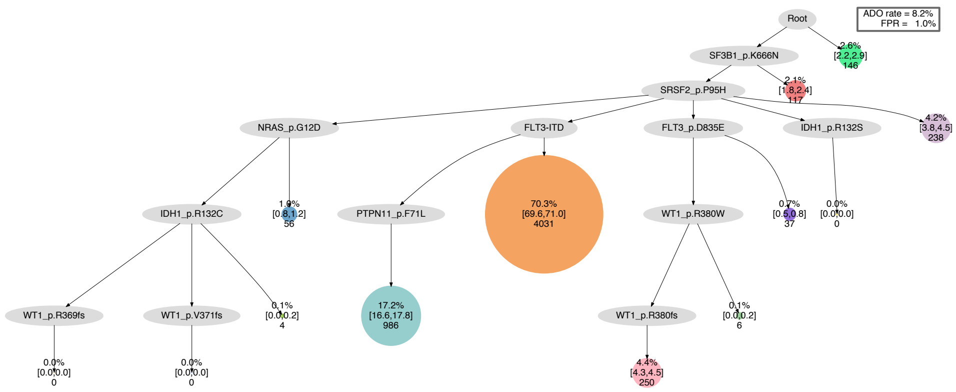


#### model 2

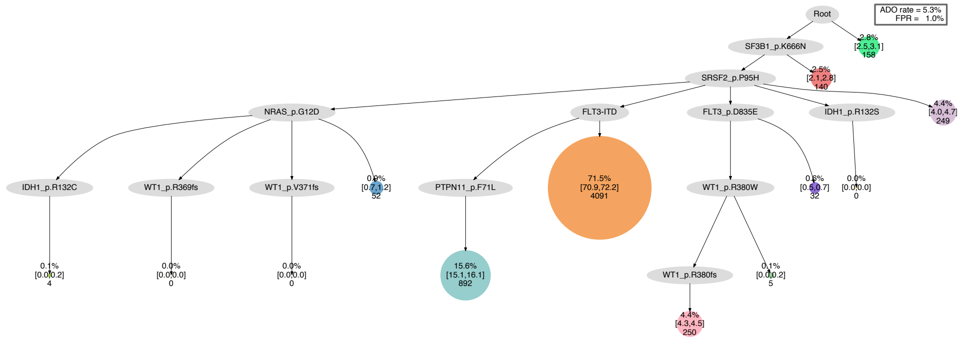


#### model 3

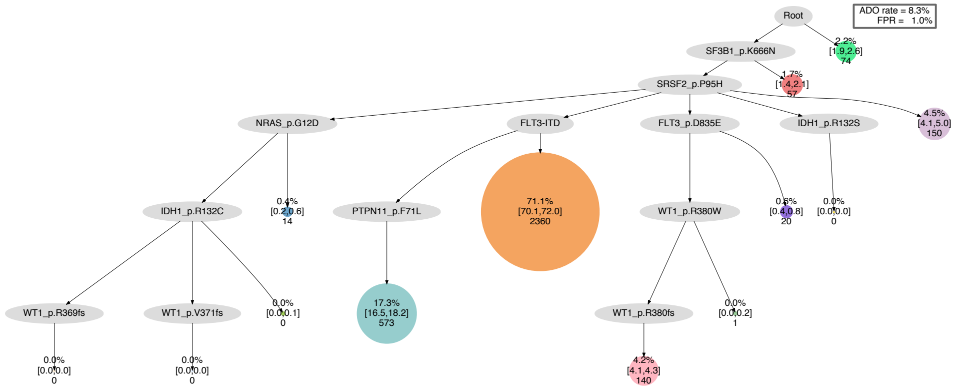


#### model 4

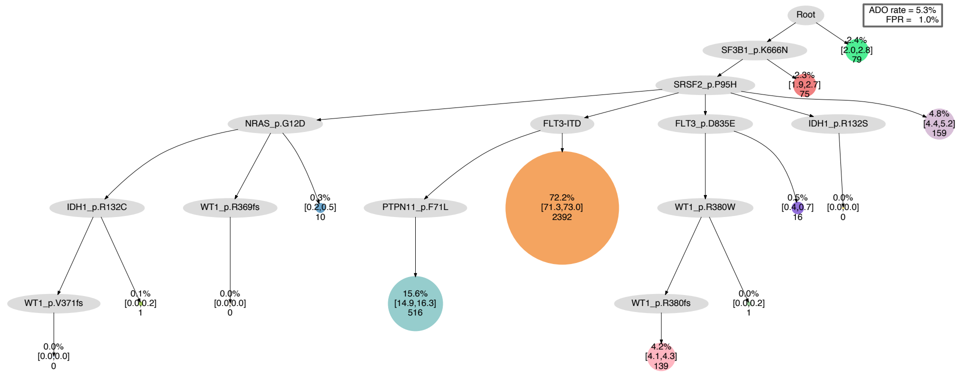

### AML-05-001

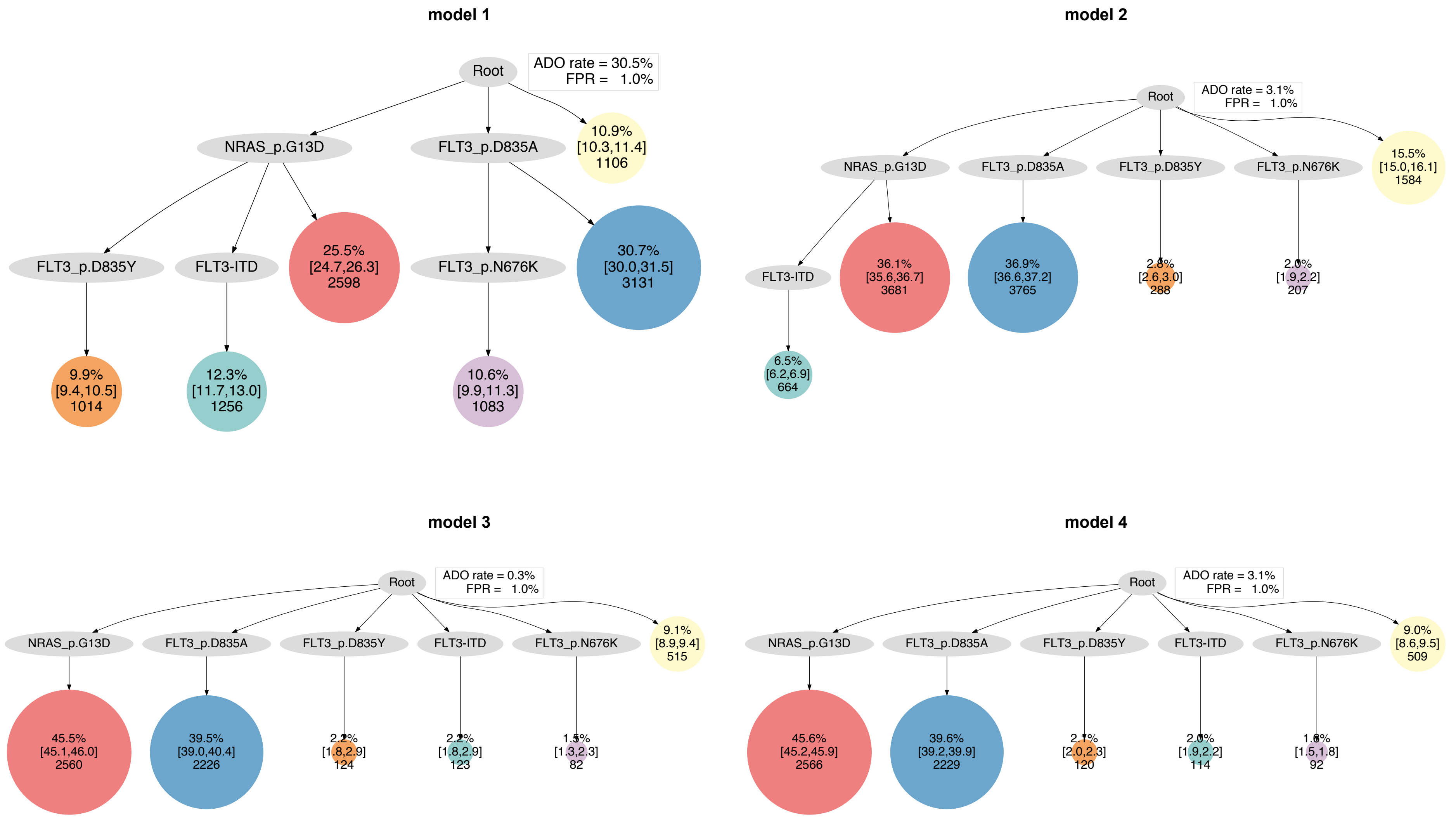

model 1
Root
ADO rate = 30.5%
FPR = 1.0%
NRAS_p.G13D
FLT3_p.D835A
10.9%
[10.3,11.4]
1106
FLT3_p.D835Y
FLT3-ITD
25.5%
[24.7,26.3]
2598
FLT3_p.N676K
30.7%
[30.0,31.5]
3131
9.9%
[9.4,10.5]
1014
12.3%
[11.7,13.0]
1256
10.6%
[9.9,11.3]
1083
model 2
Root
ADO rate = 3.1%
FPR = 1.0%
NRAS_p.G13D
FLT3_p.D835A
FLT3_p.D835Y
FLT3_p.N676K
15.5%
[15.0,16.1]
1584
FLT3-ITD
36.1%
[35.6,36.7]
3681
36.9%
[36.6,37.2]
3765
2.8%
[2.6,3.0]
288
2.0%
[1.9,2.2]
207
6.5%
[6.2,6.9]
664
model 3
Root
ADO rate = 0.3%
FPR = 1.0%
NRAS_p.G13D
FLT3_p.D835A
FLT3_p.D835Y
FLT3-ITD
FLT3_p.N676K
9.1%
[8.9,9.4]
515
45.5%
[45.1,46.0]
2560
39.5%
[39.0,40.4]
2226
2.2%
[1.8,2.9]
124
2.2%
[1.8,2.9]
123
1.5%
[1.3,2.3]
82
model 4
Root
ADO rate = 3.1%
FPR = 1.0%
NRAS_p.G13D
FLT3_p.D835A
FLT3_p.D835Y
FLT3-ITD
FLT3_p.N676K
9.0%
[8.6,9.5]
509
45.6%
[45.2,45.9]
2566
39.6%
[39.2,39.9]
2229
2.1%
[2.0,2.3]
120
2.0%
[1.9,2.2]
114
1.6%
[1.5,1.8]
92

### AML-06-001

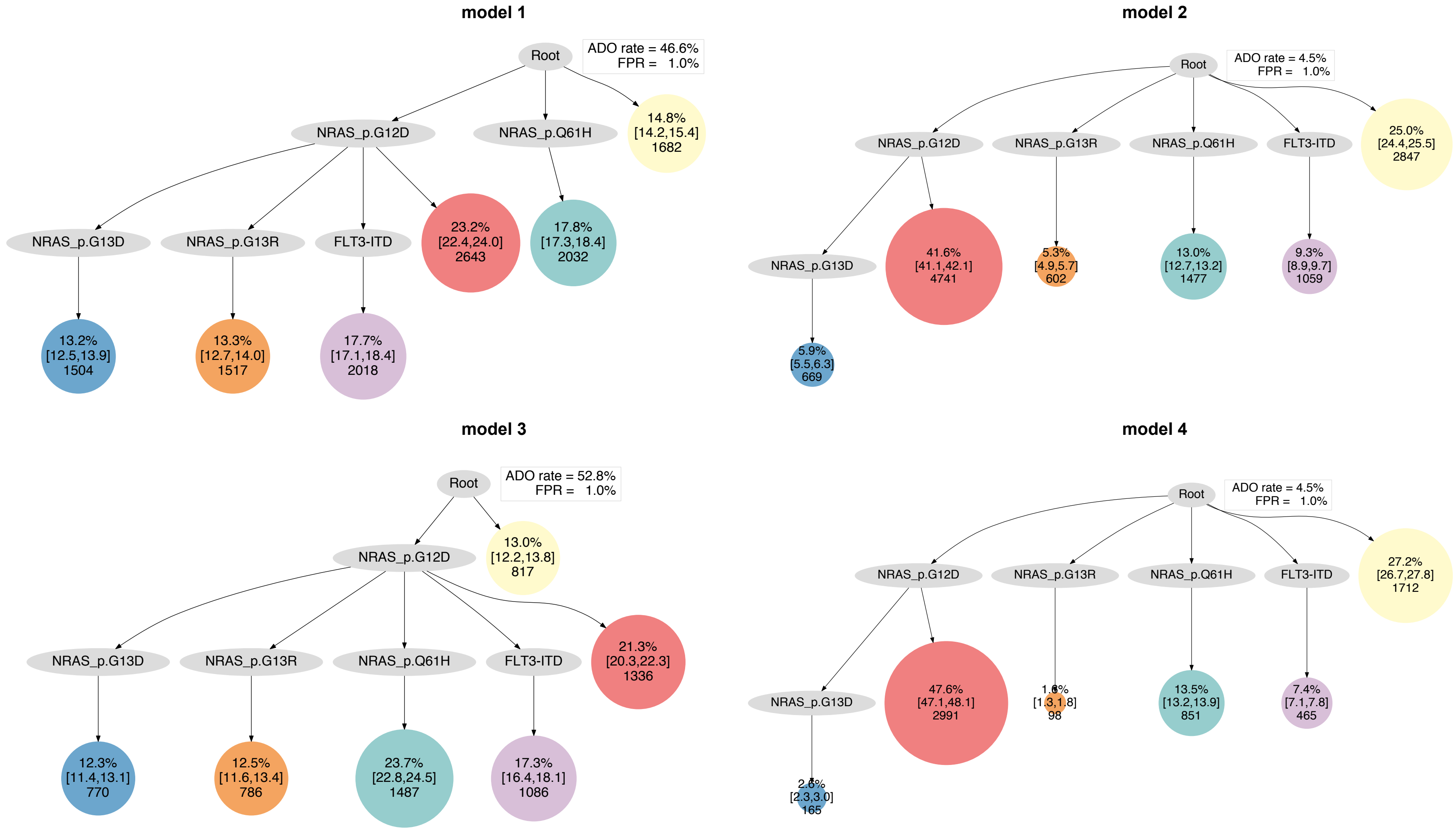

model 1
Root
ADO rate = 46.6%
FPR = 1.0%
NRAS_p.G12D
NRAS_p.Q61H
14.8%
[14.2,15.4]
1682
NRAS_p.G13D
NRAS_p.G13R
FLT3-ITD
23.2%
[22.4,24.0]
2643
17.8%
[17.3,18.4]
2032
13.2%
[12.5,13.9]
1504
13.3%
[12.7,14.0]
1517
17.7%
[17.1,18.4]
2018
model 2
Root
ADO rate = 4.5%
FPR = 1.0%
NRAS_p.G12D
NRAS_p.G13R
NRAS_p.Q61H
FLT3-ITD
25.0%
[24.4,25.5]
2847
NRAS_p.G13D
41.6%
[41.1,42.1]
4741
5.3%
[4.9,5.7]
602
13.0%
[12.7,13.2]
1477
9.3%
[8.9,9.7]
1059
5.9%
[5.5,6.3]
669
model 3
Root
ADO rate = 52.8%
FPR = 1.0%
NRAS_p.G12D
13.0%
[12.2,13.8]
817
NRAS_p.G13D
NRAS_p.G13R
NRAS_p.Q61H
FLT3-ITD
21.3%
[20.3,22.3]
1336
12.3%
[11.4,13.1]
770
12.5%
[11.6,13.4]
786
23.7%
[22.8,24.5]
1487
17.3%
[16.4,18.1]
1086
model 4
Root
ADO rate = 4.5%
FPR = 1.0%
NRAS_p.G12D
NRAS_p.G13R
NRAS_p.Q61H
FLT3-ITD
27.2%
[26.7,27.8]
1712
NRAS_p.G13D
47.6%
[47.1,48.1]
2991
[1.3,1.8]
98
13.5%
[13.2,13.9]
851
7.4%
[7.1,7.8]
465
[2.3,3.0]
165

#### AML-07-001

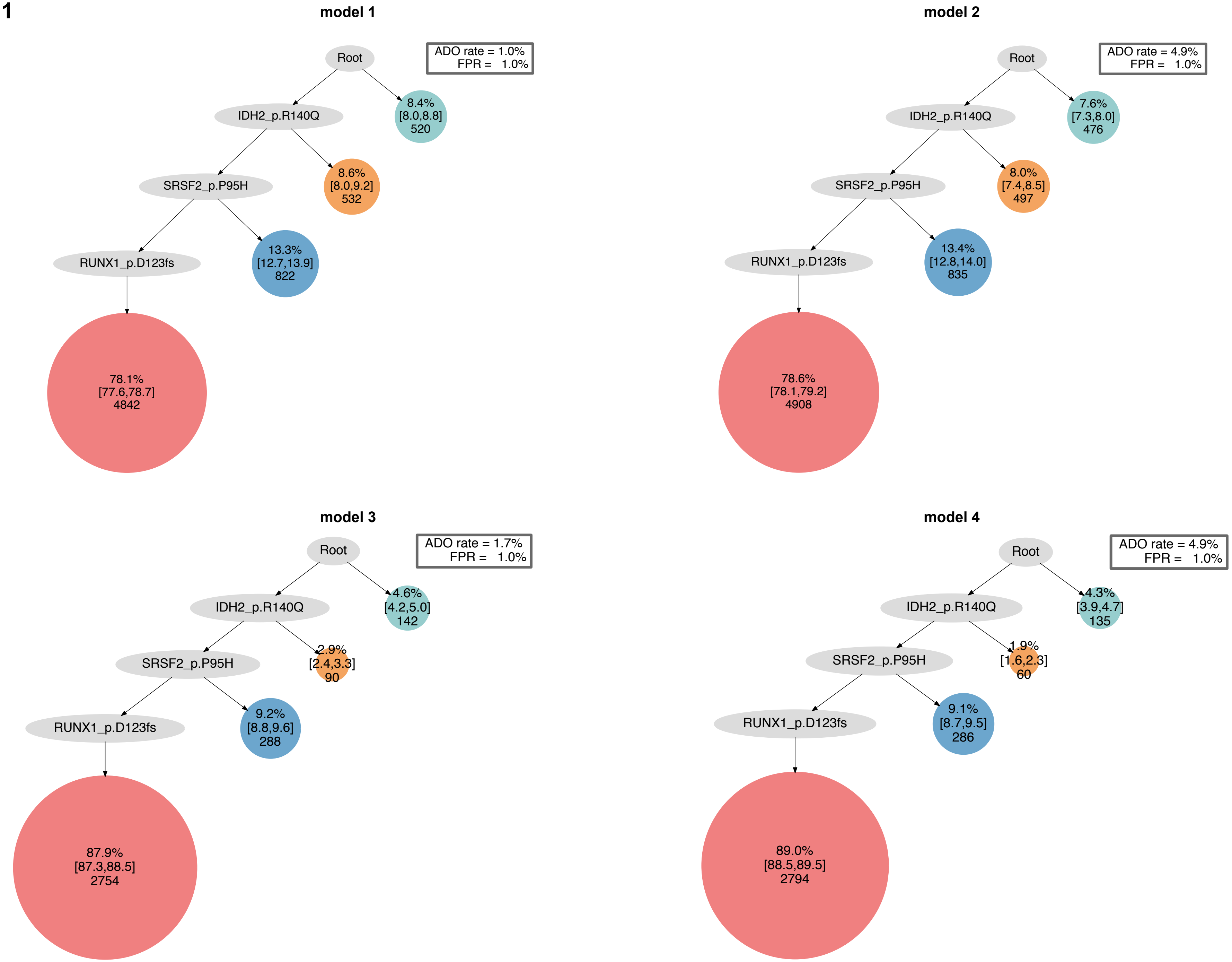

model 1
ADO rate = 1.0%
FPR = 1.0%
Root
IDH2_p.R140Q
SRSF2_p.P95H
RUNX1_p.D123fs
8.4%
[8.0,8.8]
520
8.6%
[8.0,9.2]
532
13.3%
[12.7,13.9]
822
78.1%
[77.6,78.7]
4842
model 2
ADO rate = 4.9%
FPR = 1.0%
Root
IDH2_p.R140Q
SRSF2_p.P95H
RUNX1_p.D123fs
7.6%
[7.3,8.0]
476
8.0%
[7.4,8.5]
497
13.4%
[12.8,14.0]
835
78.6%
[78.1,79.2]
4908
model 3
ADO rate = 1.7%
FPR = 1.0%
Root
IDH2_p.R140Q
SRSF2_p.P95H
RUNX1_p.D123fs
4.6%
[4.2,5.0]
142
2.9%
[2.4,3.3]
90
9.2%
[8.8,9.6]
288
87.9%
[87.3,88.5]
2754
model 4
ADO rate = 4.9%
FPR = 1.0%
Root
IDH2_p.R140Q
SRSF2_p.P95H
RUNX1_p.D123fs
4.3%
[3.9,4.7]
135
1.9%
[1.6,2.3]
60
9.1%
[8.7,9.5]
286
89.0%
[88.5,89.5]
2794

### AML-08-001

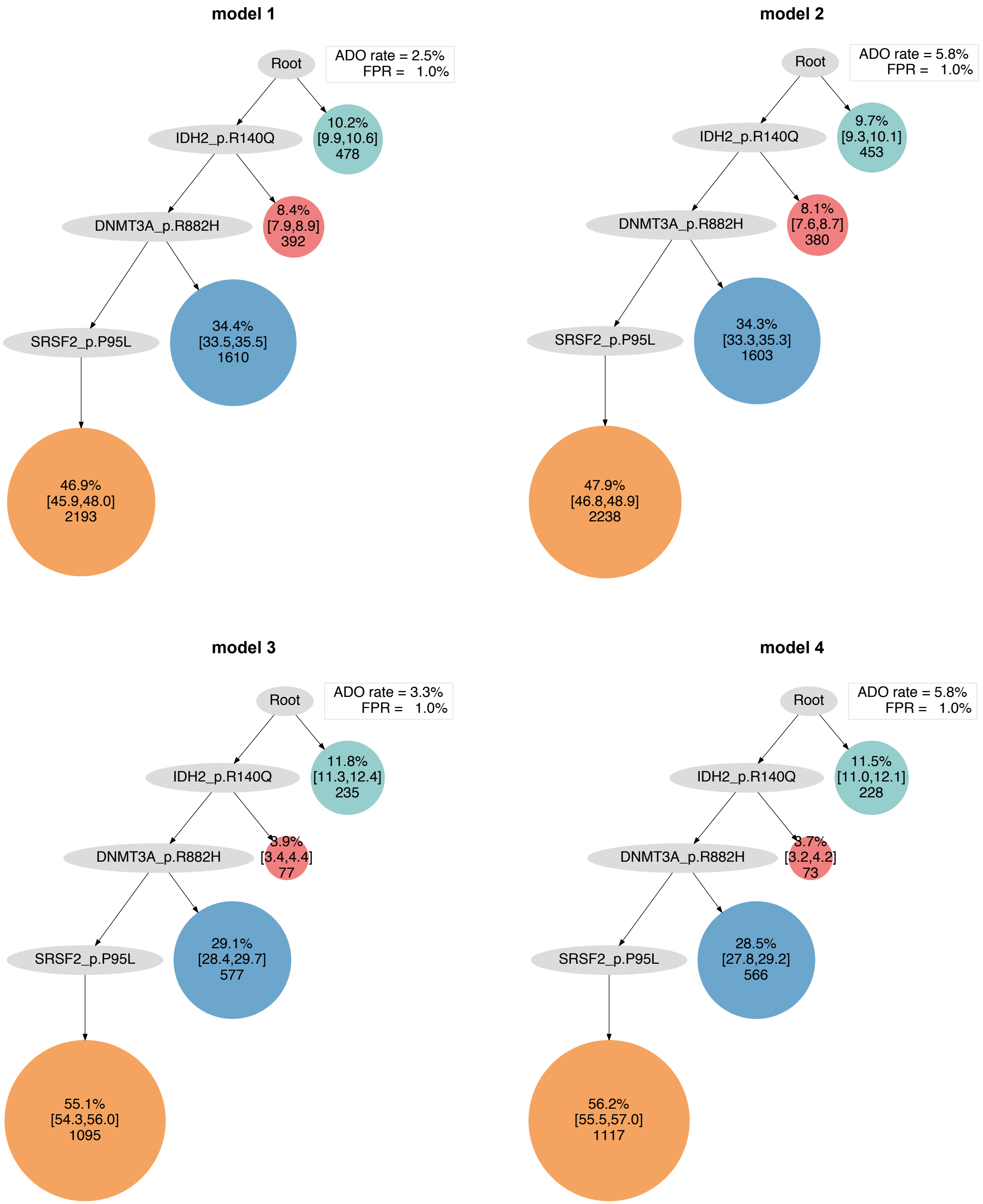
model 1
Root
ADO rate = 2.5%
FPR = 1.0%
IDH2_p.R140Q
10.2%
[9.9,10.6]
478
DNMT3A_p.R882H
8.4%
[7.9,8.9]
392
SRSF2_p.P95L
34.4%
[33.5,35.5]
1610
46.9%
[45.9,48.0]
2193
model 2
Root
ADO rate = 5.8%
FPR = 1.0%
IDH2_p.R140Q
9.7%
[9.3,10.1]
453
DNMT3A_p.R882H
8.1%
[7.6,8.7]
380
SRSF2_p.P95L
34.3%
[33.3,35.3]
1603
47.9%
[46.8,48.9]
2238
model 3
Root
ADO rate = 3.3%
FPR = 1.0%
IDH2_p.R140Q
11.8%
[11.3,12.4]
235
DNMT3A_p.R882H
3.9%
[3.4,4.4]
77
SRSF2_p.P95L
29.1%
[28.4,29.7]
577
55.1%
[54.3,56.0]
1095
model 4
Root
ADO rate = 5.8%
FPR = 1.0%
IDH2_p.R140Q
11.5%
[11.0,12.1]
228
DNMT3A_p.R882H
3.7%
[3.2,4.2]
73
SRSF2_p.P95L
28.5%
[27.8,29.2]
566
56.2%
[55.5,57.0]
1117

### AML-09-001

#### model 1

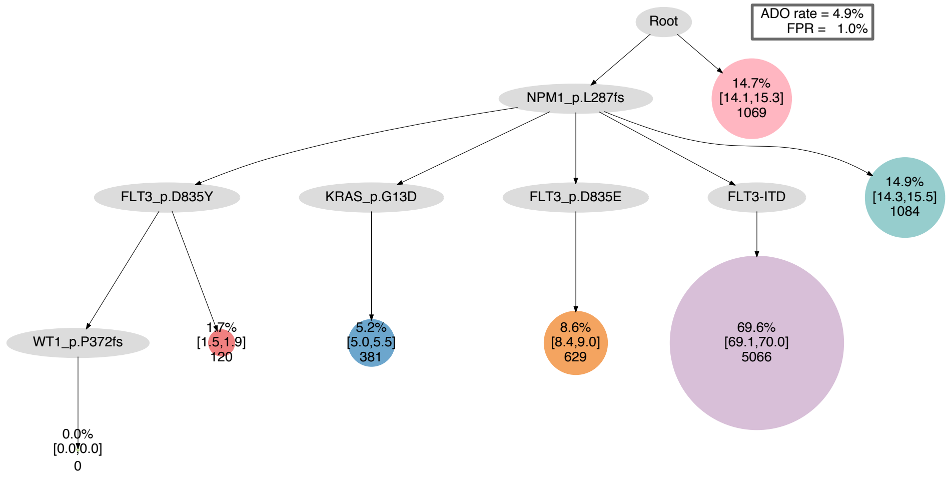


#### model 2

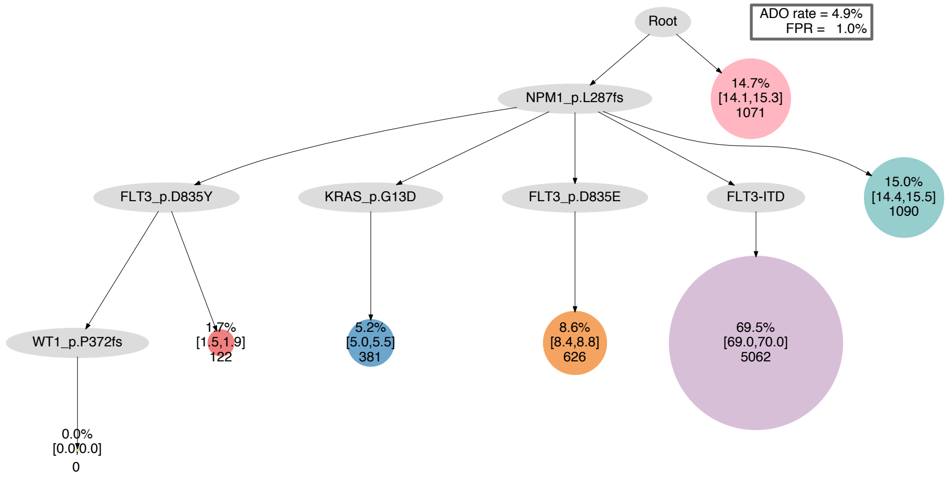


#### model 3

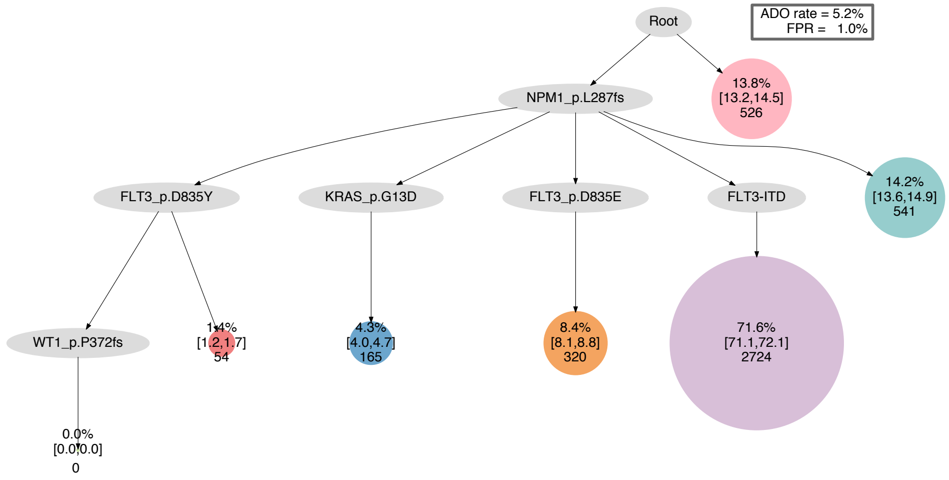


#### model 4

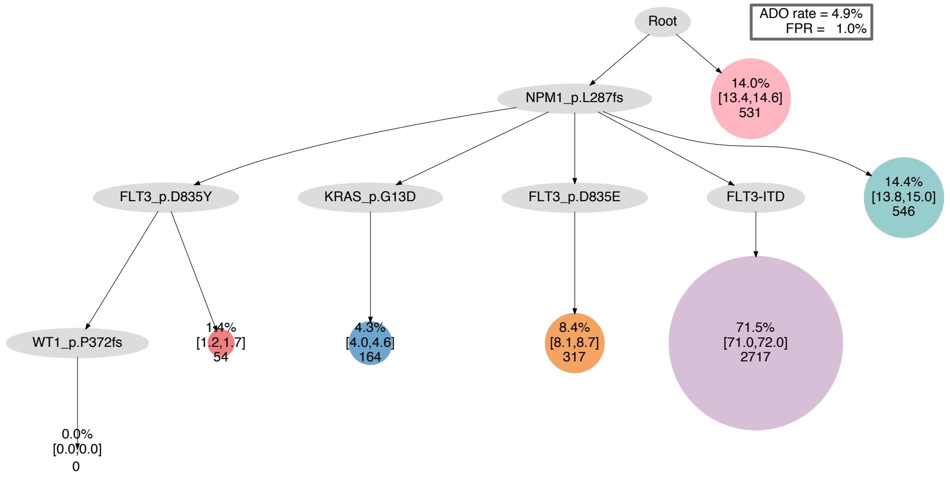

### AML-10-001

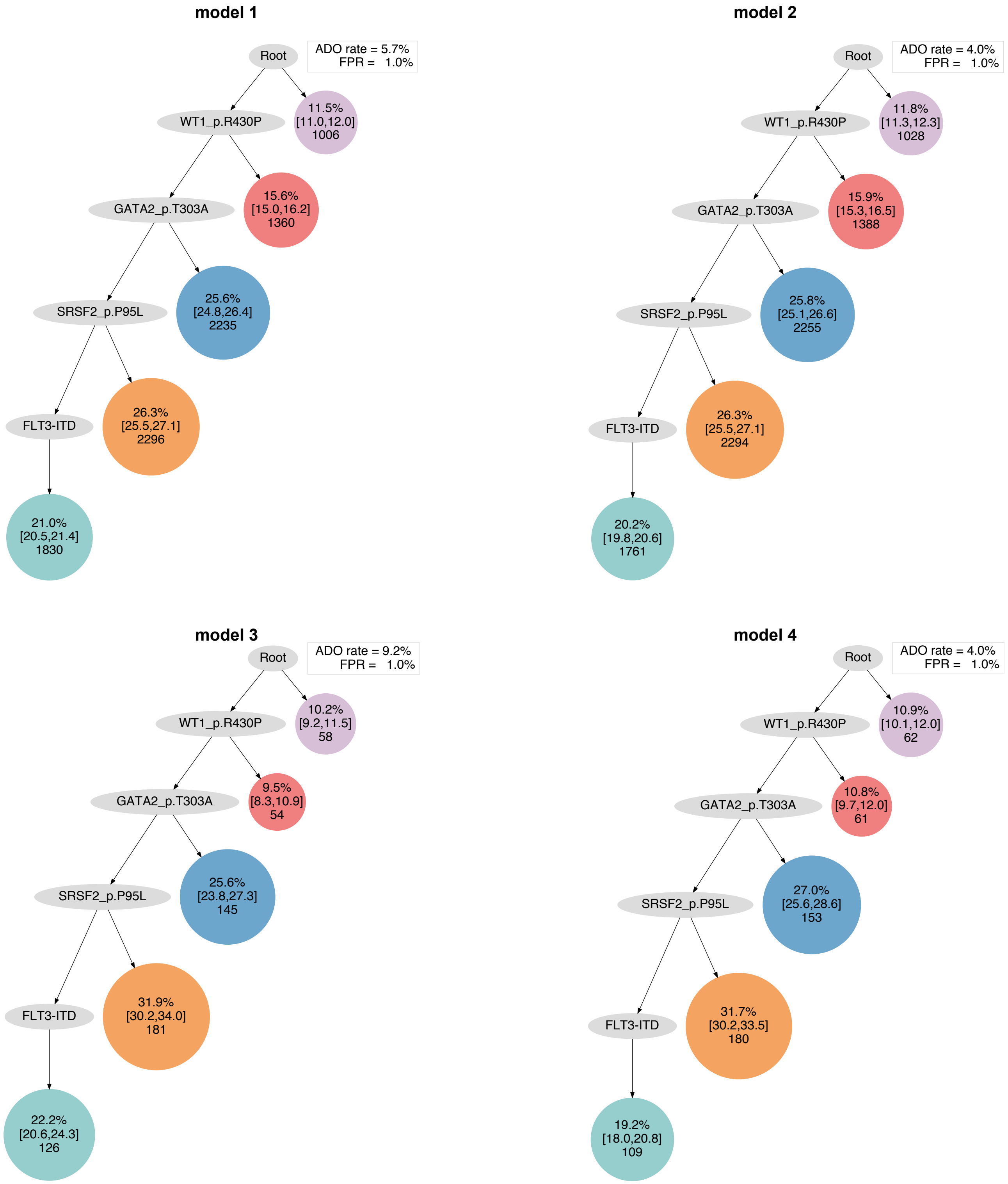

model 1
ADO rate = 5.7%
FPR = 1.0%
Root
11.5%
[11.0,12.0]
1006
WT1_p.R430P
15.6%
[15.0,16.2]
1360
GATA2_p.T303A
25.6%
[24.8,26.4]
2235
SRSF2_p.P95L
26.3%
[25.5,27.1]
2296
FLT3-ITD
21.0%
[20.5,21.4]
1830
model 2
ADO rate = 4.0%
FPR = 1.0%
Root
11.8%
[11.3,12.3]
1028
WT1_p.R430P
15.9%
[15.3,16.5]
1388
GATA2_p.T303A
25.8%
[25.1,26.6]
2255
SRSF2_p.P95L
26.3%
[25.5,27.1]
2294
FLT3-ITD
20.2%
[19.8,20.6]
1761
model 3
ADO rate = 9.2%
FPR = 1.0%
Root
10.2%
[9.2,11.5]
58
WT1_p.R430P
9.5%
[8.3,10.9]
54
GATA2_p.T303A
25.6%
[23.8,27.3]
145
SRSF2_p.P95L
31.9%
[30.2,34.0]
181
FLT3-ITD
22.2%
[20.6,24.3]
126
model 4
ADO rate = 4.0%
FPR = 1.0%
Root
10.9%
[10.1,12.0]
62
WT1_p.R430P
10.8%
[9.7,12.0]
61
GATA2_p.T303A
27.0%
[25.6,28.6]
153
SRSF2_p.P95L
31.7%
[30.2,33.5]
180
FLT3-ITD
19.2%
[18.0,20.8]
109

**AML-11-001**

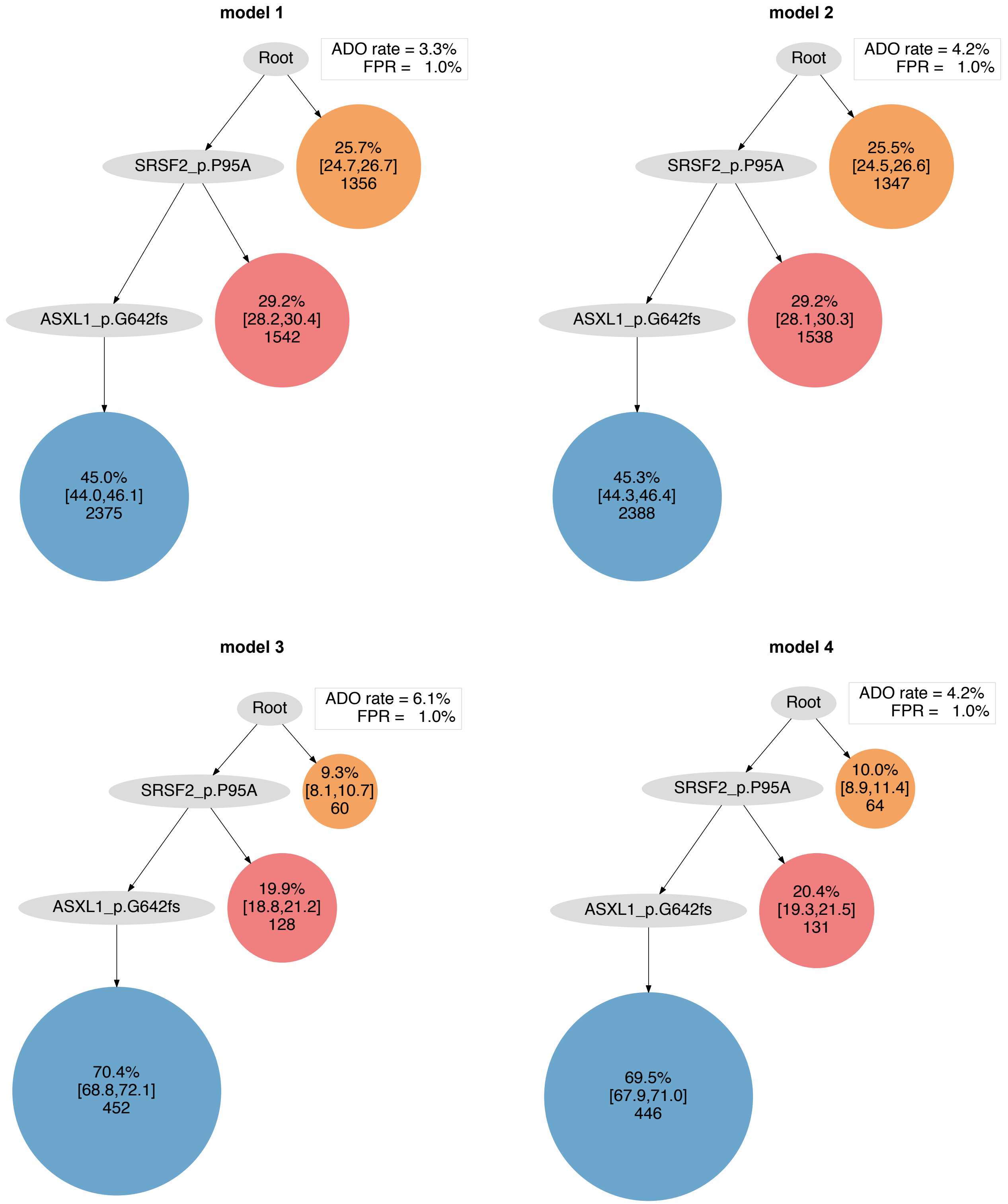
model 1
Root
ADO rate = 3.3%
FPR = 1.0%
SRSF2_p.P95A
25.7%
[24.7,26.7]
1356
ASXL1_p.G642fs
29.2%
[28.2,30.4]
1542
45.0%
[44.0,46.1]
2375
model 2
Root
ADO rate = 4.2%
FPR = 1.0%
SRSF2_p.P95A
25.5%
[24.5,26.6]
1347
ASXL1_p.G642fs
29.2%
[28.1,30.3]
1538
45.3%
[44.3,46.4]
2388
model 3
Root
ADO rate = 6.1%
FPR = 1.0%
SRSF2_p.P95A
9.3%
[8.1,10.7]
60
ASXL1_p.G642fs
19.9%
[18.8,21.2]
128
70.4%
[68.8,72.1]
452
model 4
Root
ADO rate = 4.2%
FPR = 1.0%
SRSF2_p.P95A
10.0%
[8.9,11.4]
64
ASXL1_p.G642fs
20.4%
[19.3,21.5]
131
69.5%
[67.9,71.0]
446

### AML-12-001

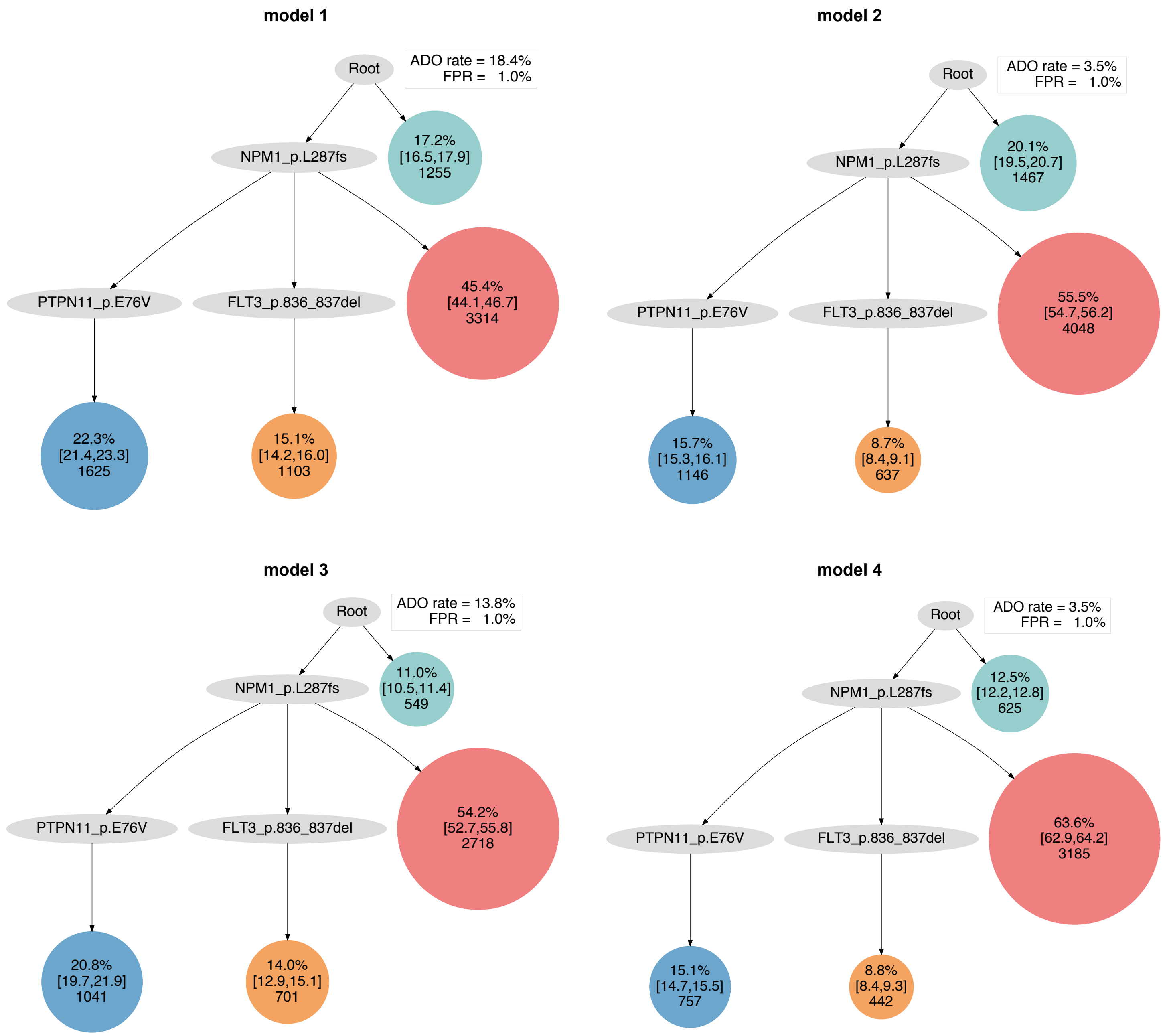
model 1
Root
ADO rate = 18.4%
FPR = 1.0%
NPM1_p.L287fs
17.2%
[16.5,17.9]
1255
PTPN11_p.E76V
FLT3_p.836_837del
45.4%
[44.1,46.7]
3314
22.3%
[21.4,23.3]
1625
15.1%
[14.2,16.0]
1103
model 2
Root
ADO rate = 3.5%
FPR = 1.0%
NPM1_p.L287fs
20.1%
[19.5,20.7]
1467
PTPN11_p.E76V
FLT3_p.836_837del
55.5%
[54.7,56.2]
4048
15.7%
[15.3,16.1]
1146
8.7%
[8.4,9.1]
637
model 3
Root
ADO rate = 13.8%
FPR = 1.0%
NPM1_p.L287fs
11.0%
[10.5,11.4]
549
PTPN11_p.E76V
FLT3_p.836_837del
54.2%
[52.7,55.8]
2718
20.8%
[19.7,21.9]
1041
14.0%
[12.9,15.1]
701
model 4
Root
ADO rate = 3.5%
FPR = 1.0%
NPM1_p.L287fs
12.5%
[12.2,12.8]
625
PTPN11_p.E76V
FLT3_p.836_837del
63.6%
[62.9,64.2]
3185
15.1%
[14.7,15.5]
757
8.8%
[8.4,9.3]
442

### AML-13-001

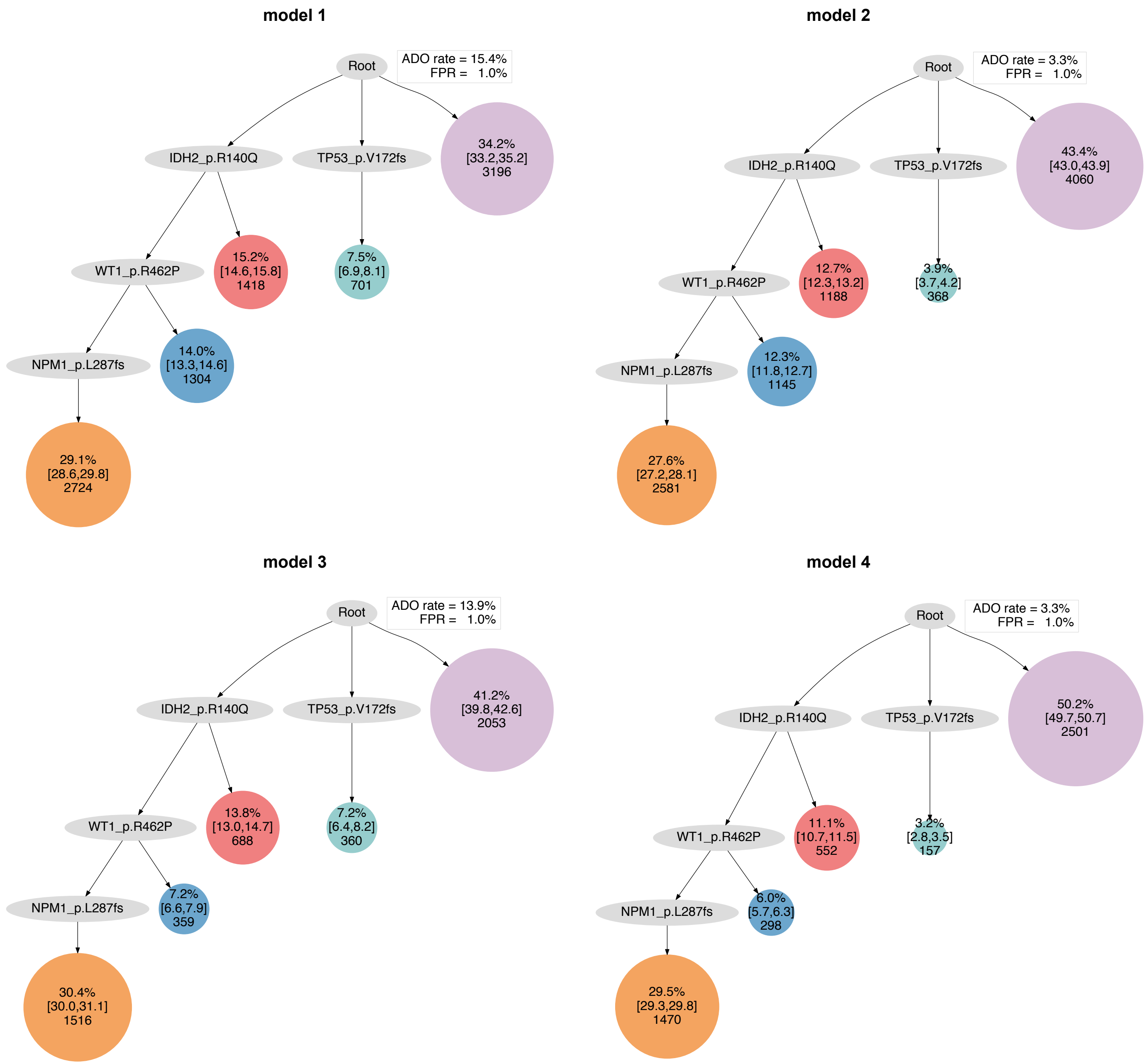

model 1
Root
ADO rate = 15.4%
FPR = 1.0%
IDH2_p.R140Q
TP53_p.V172fs
34.2%
[33.2,35.2]
3196
WT1_p.R462P
15.2%
[14.6,15.8]
1418
7.5%
[6.9,8.1]
701
NPM1_p.L287fs
14.0%
[13.3,14.6]
1304
29.1%
[28.6,29.8]
2724
model 2
Root
ADO rate = 3.3%
FPR = 1.0%
IDH2_p.R140Q
TP53_p.V172fs
43.4%
[43.0,43.9]
4060
WT1_p.R462P
12.7%
[12.3,13.2]
1188
3.9%
[3.7,4.2]
368
NPM1_p.L287fs
12.3%
[11.8,12.7]
1145
27.6%
[27.2,28.1]
2581
model 3
Root
ADO rate = 13.9%
FPR = 1.0%
IDH2_p.R140Q
TP53_p.V172fs
41.2%
[39.8,42.6]
2053
WT1_p.R462P
13.8%
[13.0,14.7]
688
7.2%
[6.4,8.2]
360
NPM1_p.L287fs
7.2%
[6.6,7.9]
359
30.4%
[30.0,31.1]
1516
model 4
Root
ADO rate = 3.3%
FPR = 1.0%
IDH2_p.R140Q
TP53_p.V172fs
50.2%
[49.7,50.7]
2501
WT1_p.R462P
11.1%
[10.7,11.5]
552
3.2%
[2.8,3.5]
157
NPM1_p.L287fs
6.0%
[5.7,6.3]
298
29.5%
[29.3,29.8]
1470

### AML-14-001

#### model 1

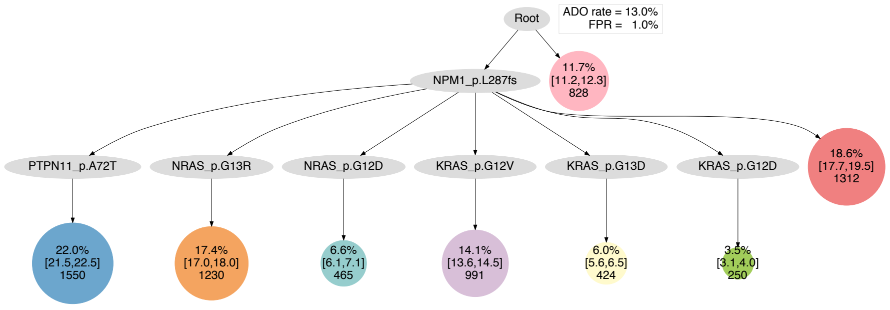


#### model 2

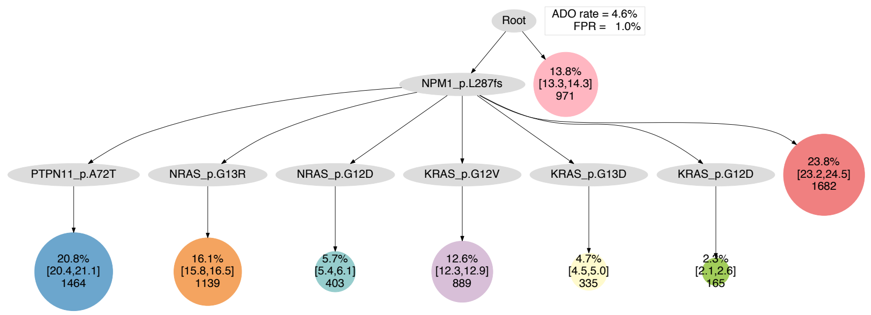


#### model 3

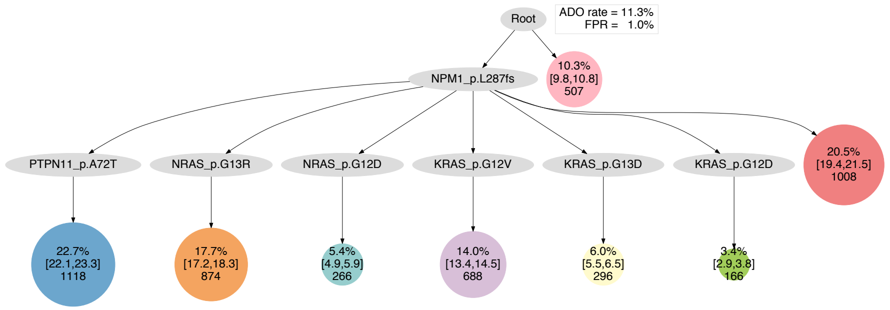


#### model 4

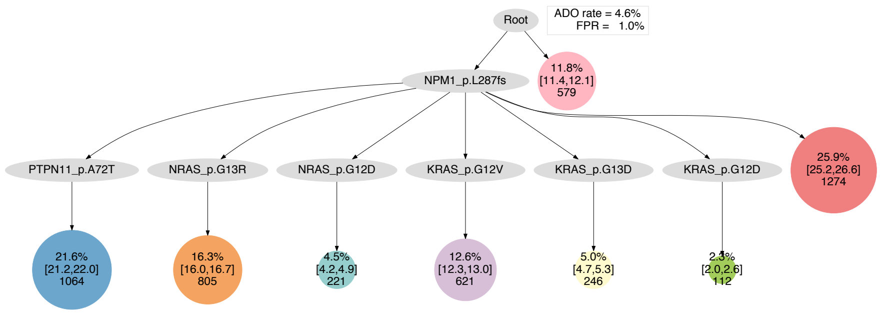

**AML-15-001**

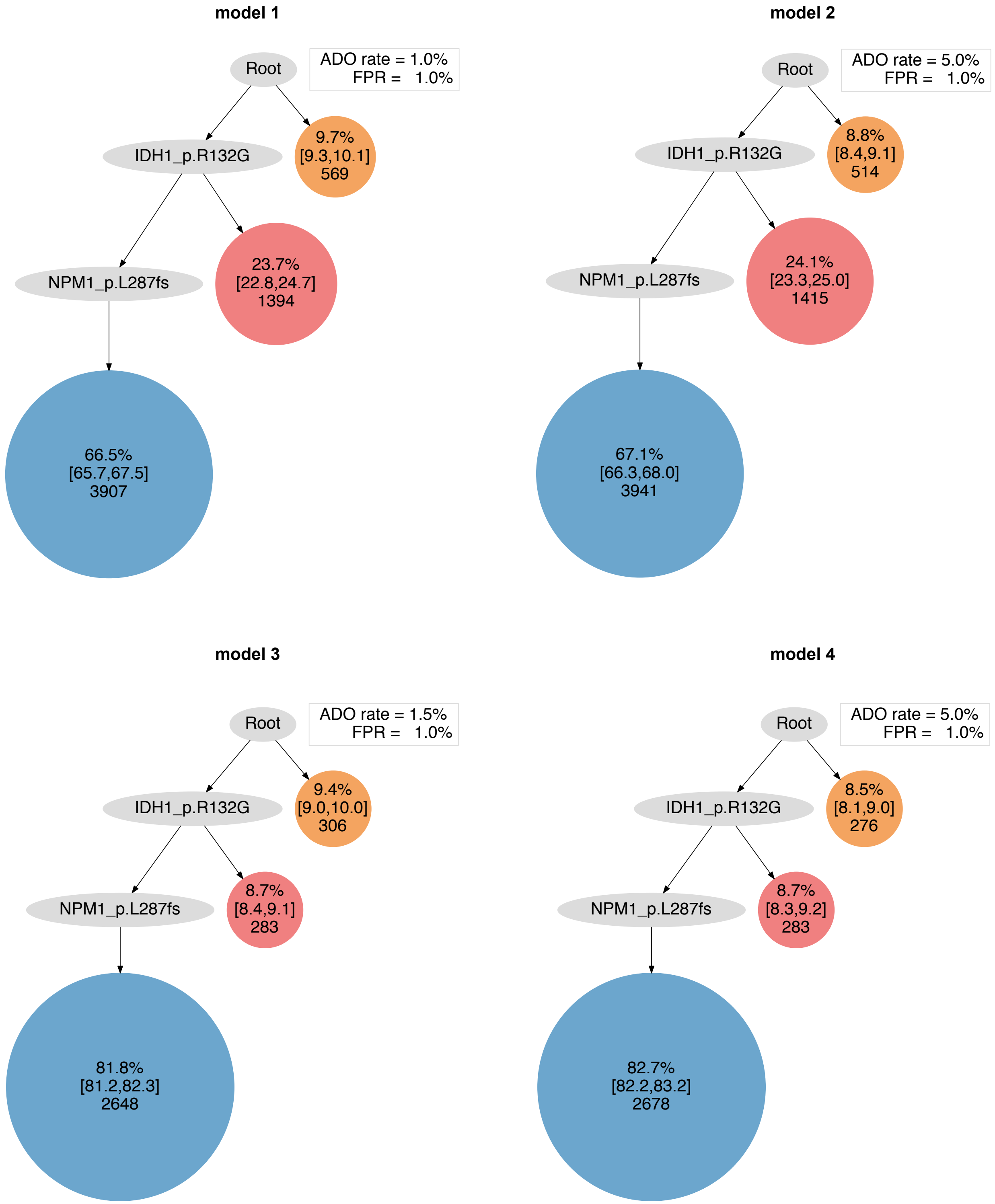
model 1
Root
ADO rate = 1.0%
FPR = 1.0%
IDH1_p.R132G
9.7%
[9.3,10.1]
569
NPM1_p.L287fs
23.7%
[22.8,24.7]
1394
66.5%
[65.7,67.5]
3907
model 2
Root
ADO rate = 5.0%
FPR = 1.0%
IDH1_p.R132G
8.8%
[8.4,9.1]
514
NPM1_p.L287fs
24.1%
[23.3,25.0]
1415
67.1%
[66.3,68.0]
3941
model 3
Root
ADO rate = 1.5%
FPR = 1.0%
IDH1_p.R132G
9.4%
[9.0,10.0]
306
NPM1_p.L287fs
8.7%
[8.4,9.1]
283
81.8%
[81.2,82.3]
2648
model 4
Root
ADO rate = 5.0%
FPR = 1.0%
IDH1_p.R132G
8.5%
[8.1,9.0]
276
NPM1_p.L287fs
8.7%
[8.3,9.2]
283
82.7%
[82.2,83.2]
2678

### AML-16-001

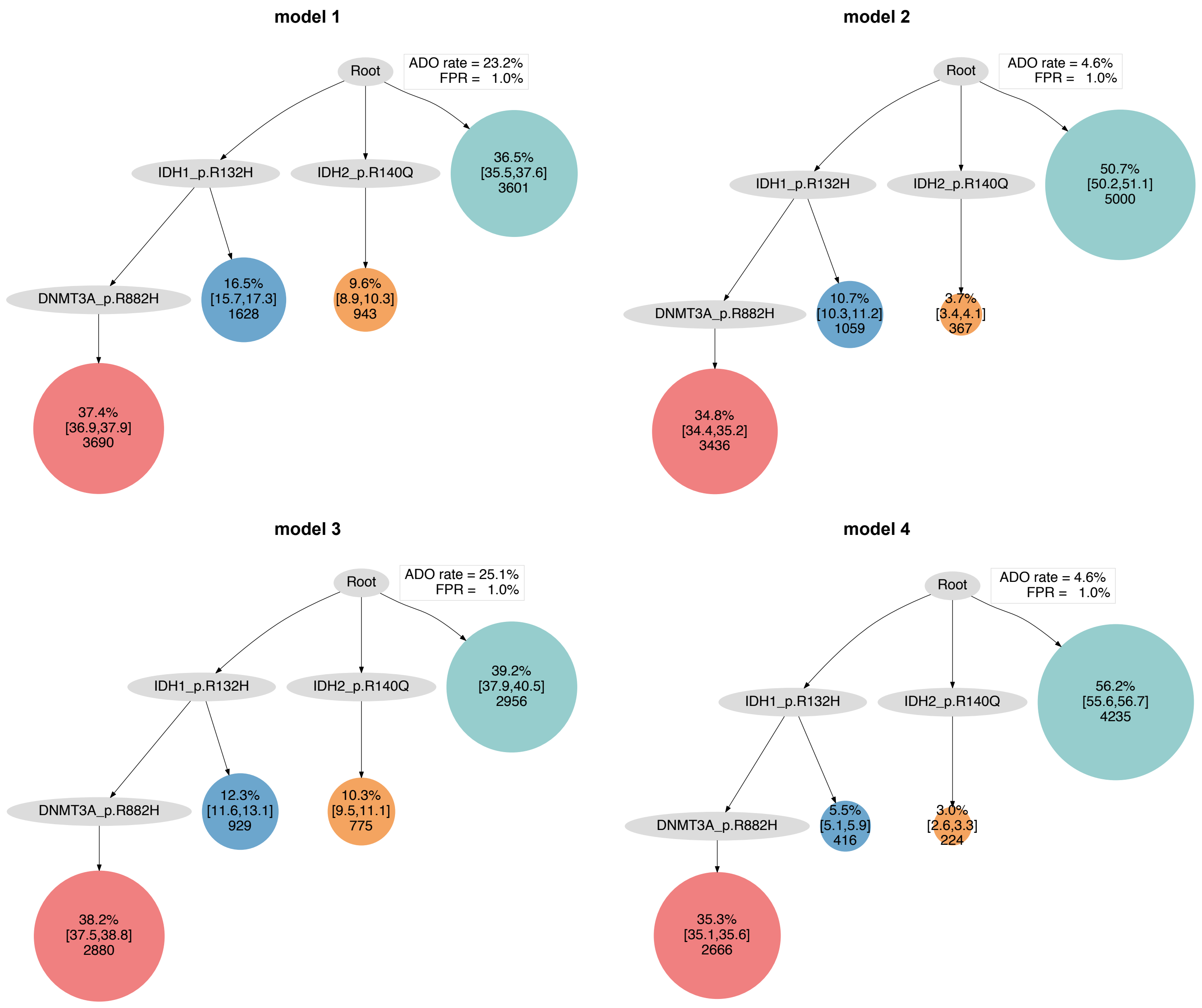
model 1
Root
ADO rate = 23.2%
FPR = 1.0%
IDH1_p.R132H
IDH2_p.R140Q
36.5%
[35.5,37.6]
3601
DNMT3A_p.R882H
16.5%
[15.7,17.3]
1628
9.6%
[8.9,10.3]
943
37.4%
[36.9,37.9]
3690
model 2
Root
ADO rate = 4.6%
FPR = 1.0%
IDH1_p.R132H
IDH2_p.R140Q
50.7%
[50.2,51.1]
5000
DNMT3A_p.R882H
10.7%
[10.3,11.2]
1059
3.7%
[3.4,4.1]
367
34.8%
[34.4,35.2]
3436
model 3
Root
ADO rate = 25.1%
FPR = 1.0%
IDH1_p.R132H
IDH2_p.R140Q
39.2%
[37.9,40.5]
2956
DNMT3A_p.R882H
12.3%
[11.6,13.1]
929
10.3%
[9.5,11.1]
775
38.2%
[37.5,38.8]
2880
model 4
Root
ADO rate = 4.6%
FPR = 1.0%
IDH1_p.R132H
IDH2_p.R140Q
56.2%
[55.6,56.7]
4235
DNMT3A_p.R882H
5.5%
[5.1,5.9]
416
3.0%
[2.6,3.3]
224
35.3%
[35.1,35.6]
2666

#### AML-17-001

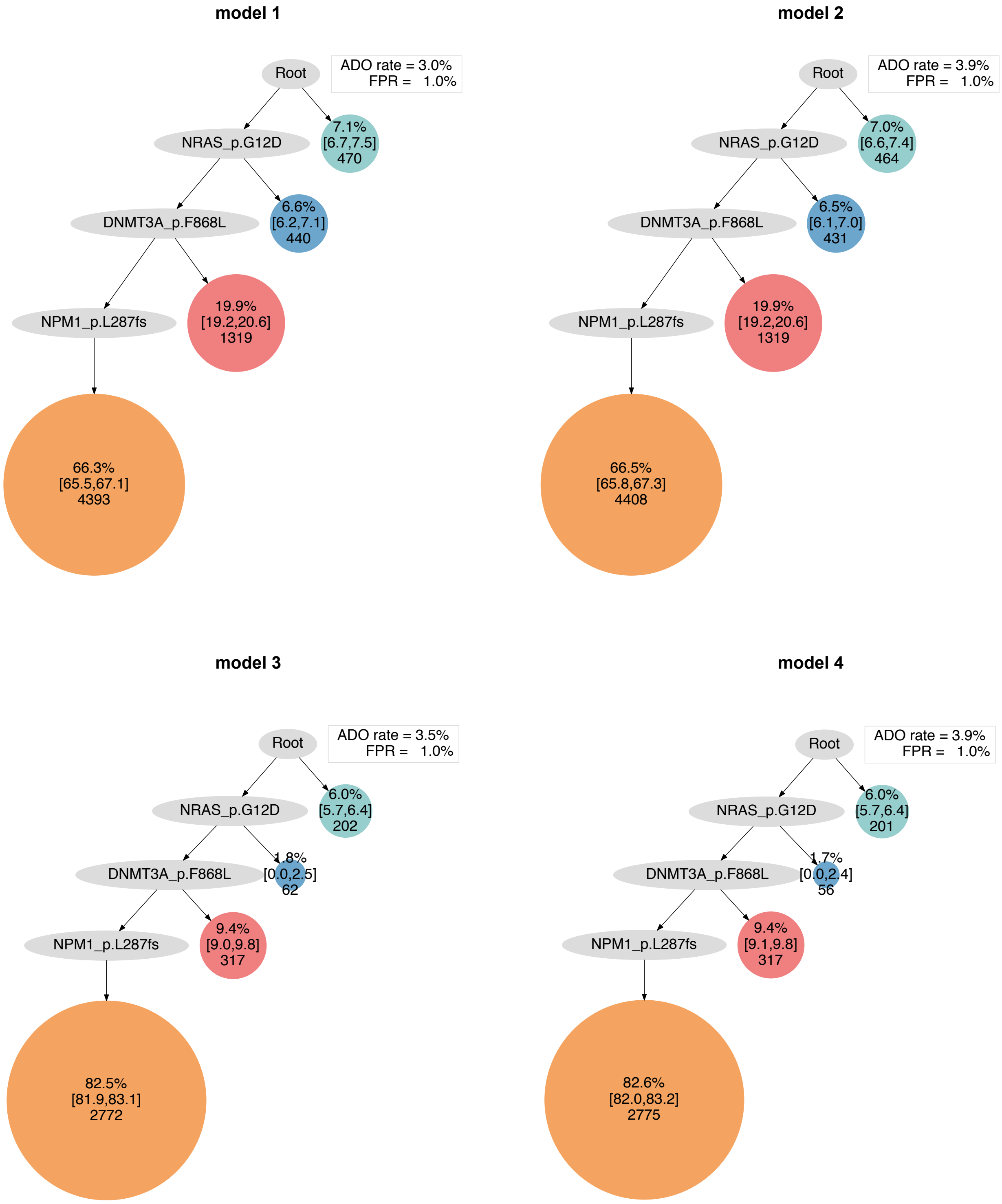

model 1
Root
ADO rate = 3.0%
FPR = 1.0%
NRAS_p.G12D
7.1%
[6.7,7.5]
470
DNMT3A_p.F868L
6.6%
[6.2,7.1]
440
NPM1_p.L287fs
19.9%
[19.2,20.6]
1319
66.3%
[65.5,67.1]
4393
model 2
Root
ADO rate = 3.9%
FPR = 1.0%
NRAS_p.G12D
7.0%
[6.6,7.4]
464
DNMT3A_p.F868L
6.5%
[6.1,7.0]
431
NPM1_p.L287fs
19.9%
[19.2,20.6]
1319
66.5%
[65.8,67.3]
4408
model 3
Root
ADO rate = 3.5%
FPR = 1.0%
NRAS_p.G12D
6.0%
[5.7,6.4]
202
DNMT3A_p.F868L
1.8%
[0.0,2.5]
62
NPM1_p.L287fs
9.4%
[9.0,9.8]
317
82.5%
[81.9,83.1]
2772
model 4
Root
ADO rate = 3.9%
FPR = 1.0%
NRAS_p.G12D
6.0%
[5.7,6.4]
201
DNMT3A_p.F868L
1.7%
[0.0,2.4]
56
NPM1_p.L287fs
9.4%
[9.1,9.8]
317
82.6%
[82.0,83.2]
2775

#### AML-18-001

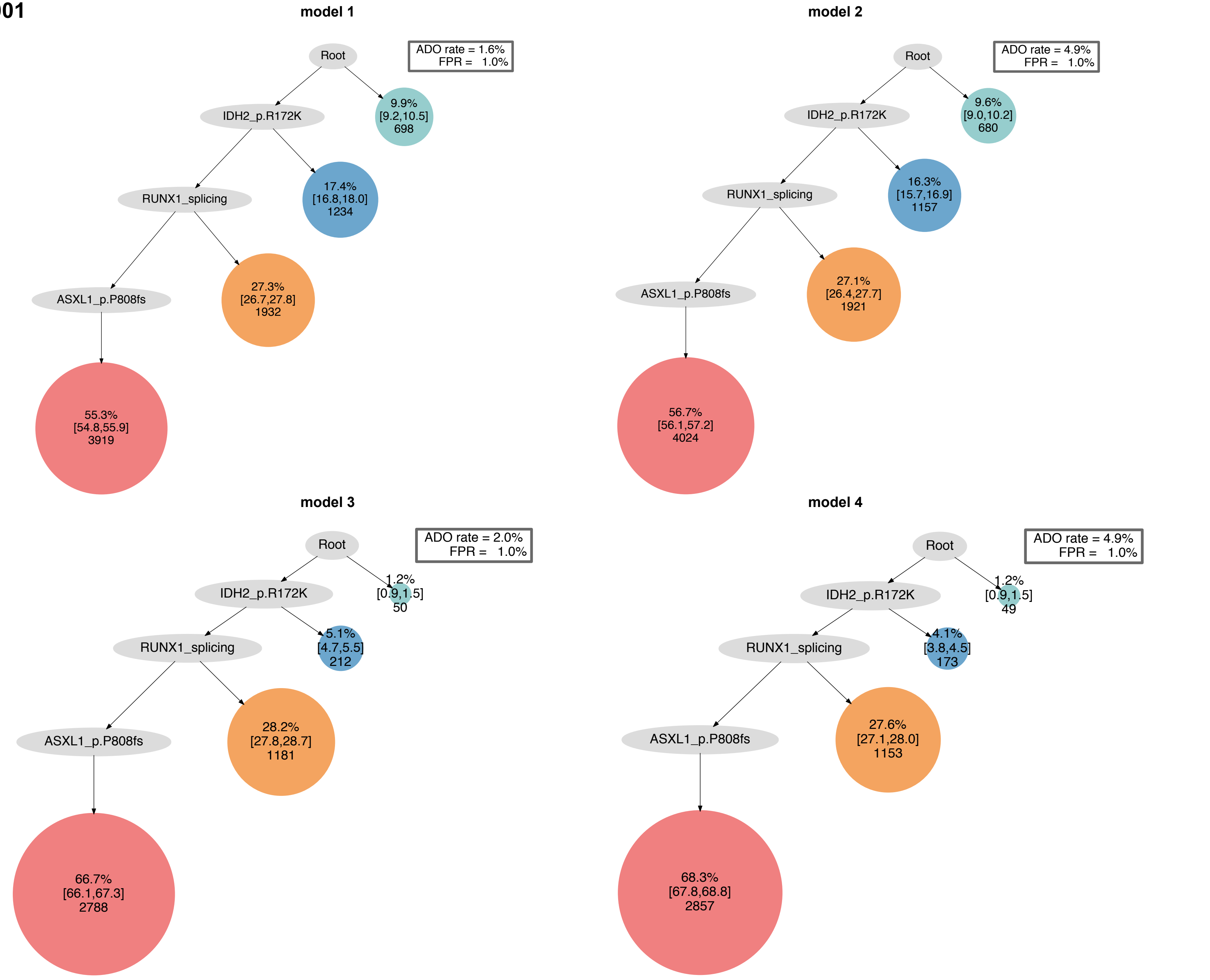

model 1
Root
ADO rate = 1.6%
FPR = 1.0%
IDH2_p.R172K
9.9%
[9.2,10.5]
698
RUNX1_splicing
17.4%
[16.8,18.0]
1234
ASXL1_p.P808fs
27.3%
[26.7,27.8]
1932
55.3%
[54.8,55.9]
3919
model 2
Root
ADO rate = 4.9%
FPR = 1.0%
IDH2_p.R172K
9.6%
[9.0,10.2]
680
RUNX1_splicing
16.3%
[15.7,16.9]
1157
ASXL1_p.P808fs
27.1%
[26.4,27.7]
1921
56.7%
[56.1,57.2]
4024
model 3
Root
ADO rate = 2.0%
FPR = 1.0%
1.2%
[0.9,1.5]
50
IDH2_p.R172K
5.1%
[4.7,5.5]
212
RUNX1_splicing
ASXL1_p.P808fs
28.2%
[27.8,28.7]
1181
66.7%
[66.1,67.3]
2788
model 4
Root
ADO rate = 4.9%
FPR = 1.0%
1.2%
[0.9,1.5]
49
IDH2_p.R172K
4.1%
[3.8,4.5]
173
RUNX1_splicing
ASXL1_p.P808fs
27.6%
[27.1,28.0]
1153
68.3%
[67.8,68.8]
2857

#### AML-19-001

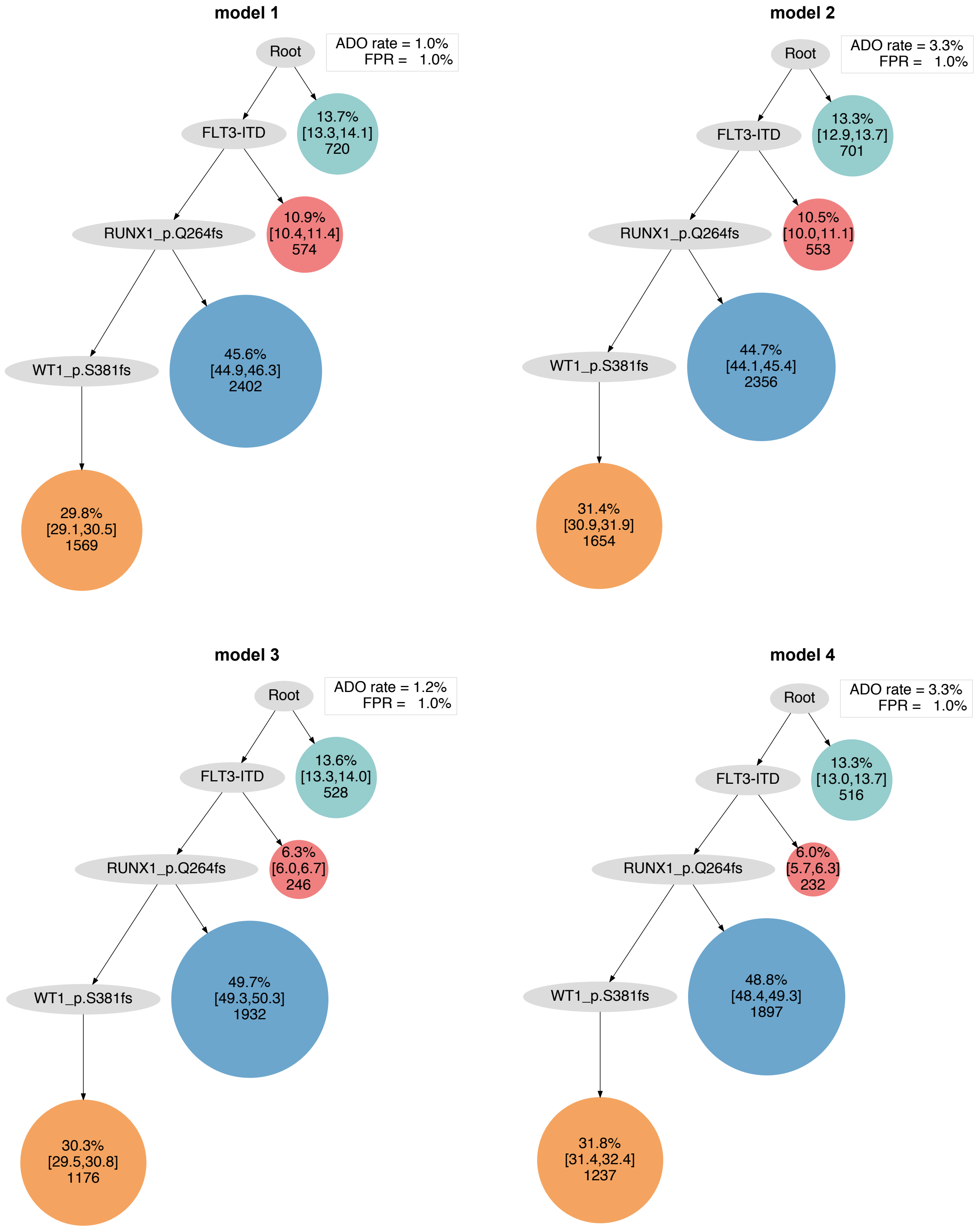

model 1
ADO rate = 1.0%
FPR = 1.0%
Root
FLT3-ITD
13.7%
[13.3,14.1]
720
RUNX1_p.Q264fs
10.9%
[10.4,11.4]
574
WT1_p.S381fs
45.6%
[44.9,46.3]
2402
29.8%
[29.1,30.5]
1569
model 2
ADO rate = 3.3%
FPR = 1.0%
Root
FLT3-ITD
13.3%
[12.9,13.7]
701
RUNX1_p.Q264fs
10.5%
[10.0,11.1]
553
WT1_p.S381fs
44.7%
[44.1,45.4]
2356
31.4%
[30.9,31.9]
1654
model 3
ADO rate = 1.2%
FPR = 1.0%
Root
FLT3-ITD
13.6%
[13.3,14.0]
528
RUNX1_p.Q264fs
6.3%
[6.0,6.7]
246
WT1_p.S381fs
49.7%
[49.3,50.3]
1932
30.3%
[29.5,30.8]
1176
model 4
ADO rate = 3.3%
FPR = 1.0%
Root
FLT3-ITD
13.3%
[13.0,13.7]
516
RUNX1_p.Q264fs
6.0%
[5.7,6.3]
232
WT1_p.S381fs
48.8%
[48.4,49.3]
1897
31.8%
[31.4,32.4]
1237

### AML-20-001

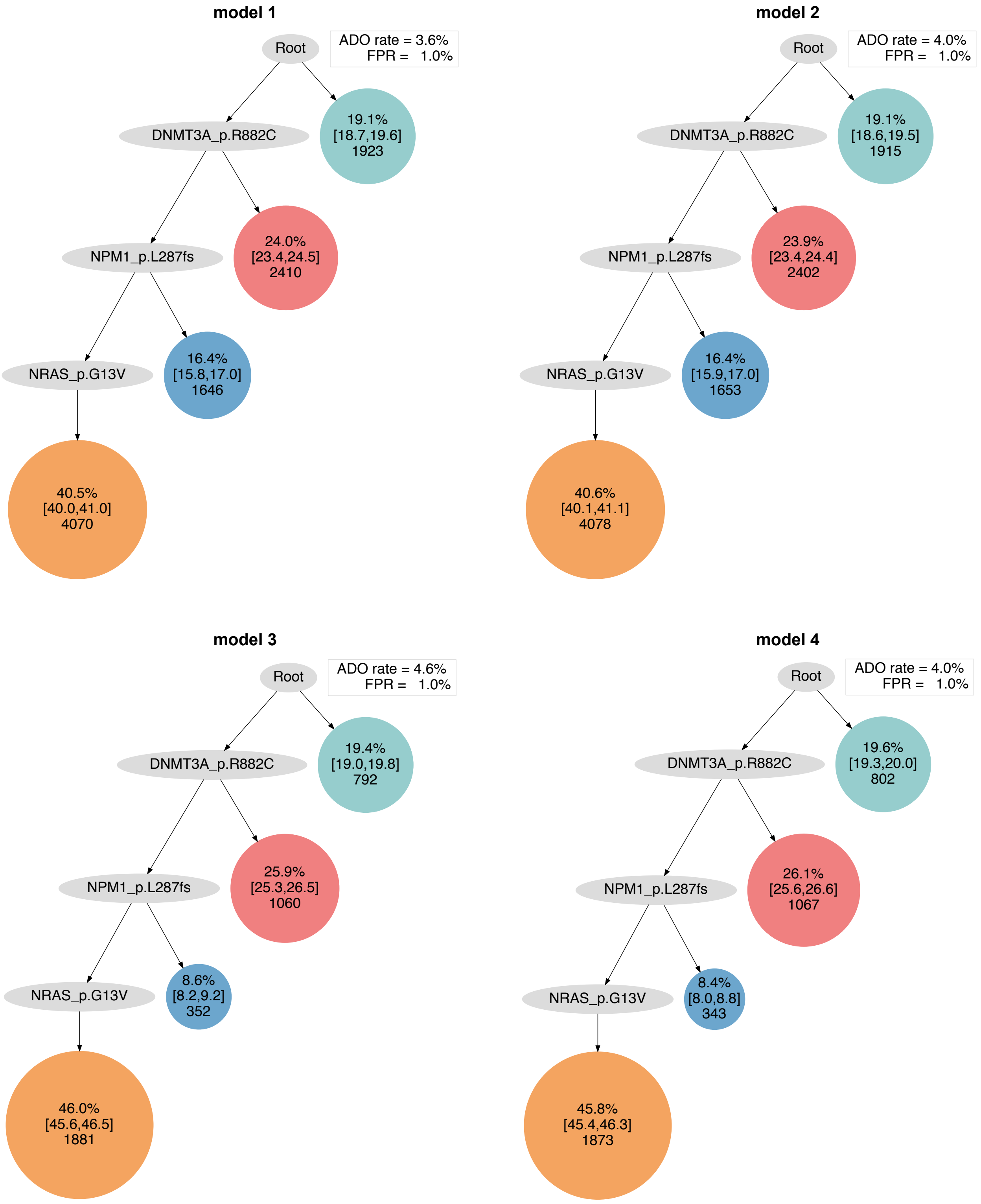
model 1
Root
ADO rate = 3.6%
FPR = 1.0%
DNMT3A_p.R882C
19.1%
[18.7,19.6]
1923
NPM1_p.L287fs
24.0%
[23.4,24.5]
2410
NRAS_p.G13V
16.4%
[15.8,17.0]
1646
40.5%
[40.0,41.0]
4070
model 2
Root
ADO rate = 4.0%
FPR = 1.0%
DNMT3A_p.R882C
19.1%
[18.6,19.5]
1915
NPM1_p.L287fs
23.9%
[23.4,24.4]
2402
NRAS_p.G13V
16.4%
[15.9,17.0]
1653
40.6%
[40.1,41.1]
4078
model 3
Root
ADO rate = 4.6%
FPR = 1.0%
DNMT3A_p.R882C
19.4%
[19.0,19.8]
792
NPM1_p.L287fs
25.9%
[25.3,26.5]
1060
NRAS_p.G13V
8.6%
[8.2,9.2]
352
46.0%
[45.6,46.5]
1881
model 4
Root
ADO rate = 4.0%
FPR = 1.0%
DNMT3A_p.R882C
19.6%
[19.3,20.0]
802
NPM1_p.L287fs
26.1%
[25.6,26.6]
1067
NRAS_p.G13V
8.4%
[8.0,8.8]
343
45.8%
[45.4,46.3]
1873

### AML-21-001

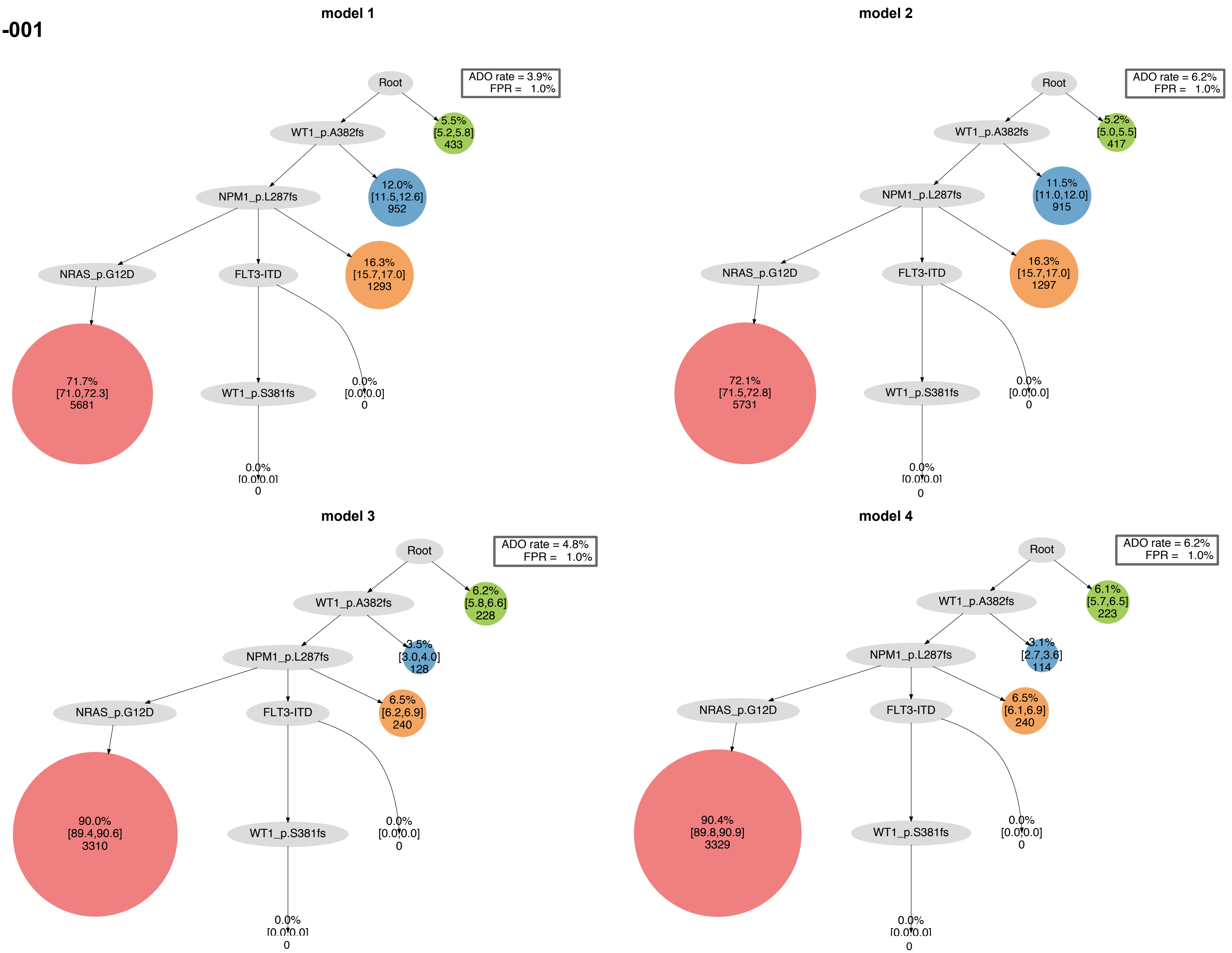

model 1
Root
ADO rate = 3.9%
FPR = 1.0%
WT1_p.A382fs
5.5%
[5.2,5.8]
433
NPM1_p.L287fs
12.0%
[11.5,12.6]
952
NRAS_p.G12D
FLT3-ITD
16.3%
[15.7,17.0]
1293
71.7%
[71.0,72.3]
5681
WT1_p.S381fs
0.0%
[0.0,0.0]
0
0.0%
[0.0,0.0]
0
model 2
Root
ADO rate = 6.2%
FPR = 1.0%
WT1_p.A382fs
5.2%
[5.0,5.5]
417
NPM1_p.L287fs
11.5%
[11.0,12.0]
915
NRAS_p.G12D
FLT3-ITD
16.3%
[15.7,17.0]
1297
72.1%
[71.5,72.8]
5731
WT1_p.S381fs
0.0%
[0.0,0.0]
0
0.0%
[0.0,0.0]
0
model 3
Root
ADO rate = 4.8%
FPR = 1.0%
WT1_p.A382fs
6.2%
[5.8,6.6]
228
NPM1_p.L287fs
3.5%
[3.0,4.0]
128
NRAS_p.G12D
FLT3-ITD
6.5%
[6.2,6.9]
240
90.0%
[89.4,90.6]
3310
WT1_p.S381fs
0.0%
[0.0,0.0]
0
0.0%
[0.0,0.0]
0
model 4
Root
ADO rate = 6.2%
FPR = 1.0%
WT1_p.A382fs
6.1%
[5.7,6.5]
223
NPM1_p.L287fs
3.1%
[2.7,3.6]
114
NRAS_p.G12D
FLT3-ITD
6.5%
[6.1,6.9]
240
90.4%
[89.8,90.9]
3329
WT1_p.S381fs
0.0%
[0.0,0.0]
0
0.0%
[0.0,0.0]
0

### AML-22-001

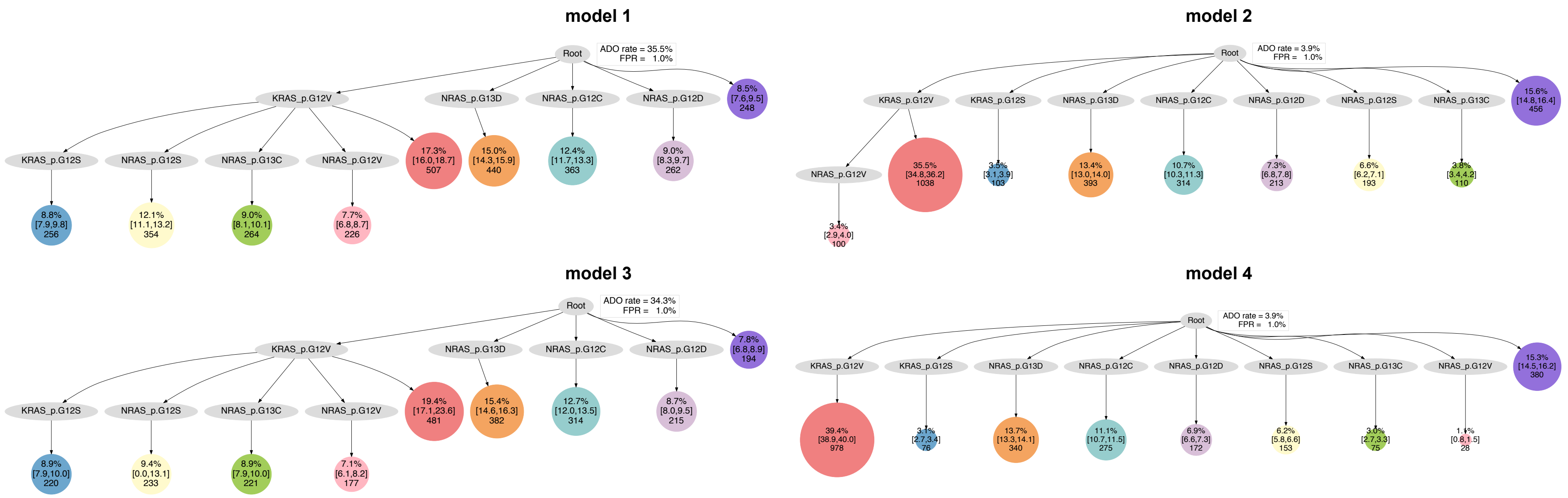
model 1
Root
ADO rate = 35.5%
FPR = 1.0%
KRAS_p.G12V
NRAS_p.G13D
NRAS_p.G12C
NRAS_p.G12D
8.5%
[7.6,9.5]
248
KRAS_p.G12S
NRAS_p.G12S
NRAS_p.G13C
NRAS_p.G12V
17.3%
[16.0,18.7]
507
15.0%
[14.3,15.9]
440
12.4%
[11.7,13.3]
363
9.0%
[8.3,9.7]
262
8.8%
[7.9,9.8]
256
12.1%
[11.1,13.2]
354
9.0%
[8.1,10.1]
264
7.7%
[6.8,8.7]
226
model 2
Root
ADO rate = 3.9%
FPR = 1.0%
KRAS_p.G12V
KRAS_p.G12S
NRAS_p.G13D
NRAS_p.G12C
NRAS_p.G12D
NRAS_p.G12S
NRAS_p.G13C
15.6%
[14.8,16.4]
456
NRAS_p.G12V
35.5%
[34.8,36.2]
1038
3.5%
[3.1,3.9]
103
13.4%
[13.0,14.0]
393
10.7%
[10.3,11.3]
314
7.3%
[6.8,7.8]
213
6.6%
[6.2,7.1]
193
3.8%
[3.4,4.2]
110
3.4%
[2.9,4.0]
100
model 3
Root
ADO rate = 34.3%
FPR = 1.0%
KRAS_p.G12V
NRAS_p.G13D
NRAS_p.G12C
NRAS_p.G12D
7.8%
[6.8,8.9]
194
KRAS_p.G12S
NRAS_p.G12S
NRAS_p.G13C
NRAS_p.G12V
19.4%
[17.1,23.6]
481
15.4%
[14.6,16.3]
382
12.7%
[12.0,13.5]
314
8.7%
[8.0,9.5]
215
8.9%
[7.9,10.0]
220
9.4%
[0.0,13.1]
233
8.9%
[7.9,10.0]
221
7.1%
[6.1,8.2]
177
model 4
Root
ADO rate = 3.9%
FPR = 1.0%
KRAS_p.G12V
KRAS_p.G12S
NRAS_p.G13D
NRAS_p.G12C
NRAS_p.G12D
NRAS_p.G12S
NRAS_p.G13C
NRAS_p.G12V
15.3%
[14.5,16.2]
380
39.4%
[38.9,40.0]
978
3.1%
[2.7,3.4]
76
13.7%
[13.3,14.1]
340
11.1%
[10.7,11.5]
275
6.9%
[6.6,7.3]
172
6.2%
[5.8,6.6]
153
3.0%
[2.7,3.3]
75
1.4%
[0.8,1.5]
28

**AML-23-001**

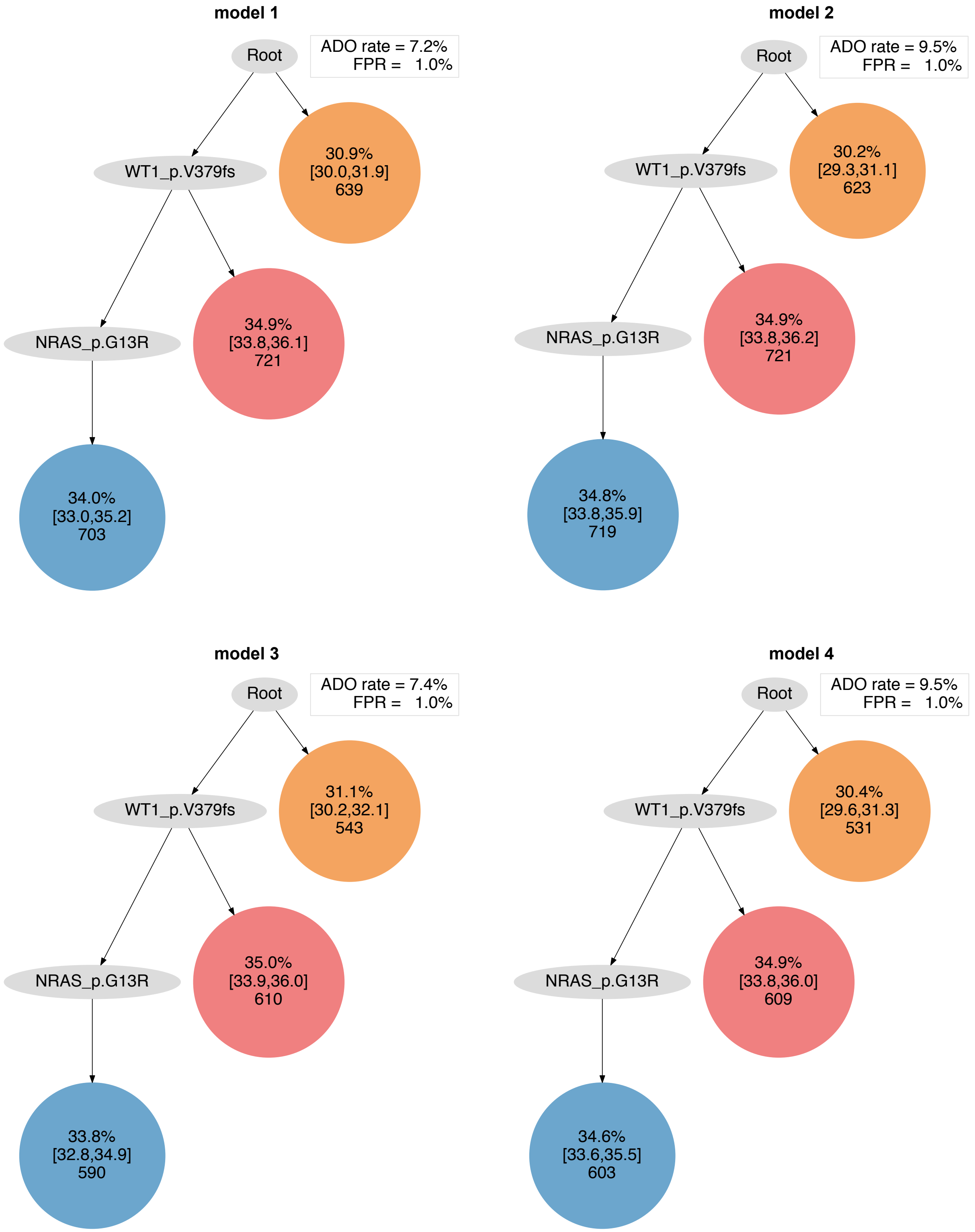
model 1
Root
ADO rate = 7.2%
FPR = 1.0%
WT1_p.V379fs
30.9%
[30.0,31.9]
639
NRAS_p.G13R
34.9%
[33.8,36.1]
721
34.0%
[33.0,35.2]
703
model 2
Root
ADO rate = 9.5%
FPR = 1.0%
WT1_p.V379fs
30.2%
[29.3,31.1]
623
NRAS_p.G13R
34.9%
[33.8,36.2]
721
34.8%
[33.8,35.9]
719
model 3
Root
ADO rate = 7.4%
FPR = 1.0%
WT1_p.V379fs
31.1%
[30.2,32.1]
543
NRAS_p.G13R
35.0%
[33.9,36.0]
610
33.8%
[32.8,34.9]
590
model 4
Root
ADO rate = 9.5%
FPR = 1.0%
WT1_p.V379fs
30.4%
[29.6,31.3]
531
NRAS_p.G13R
34.9%
[33.8,36.0]
609
34.6%
[33.6,35.5]
603

### AML-24-001

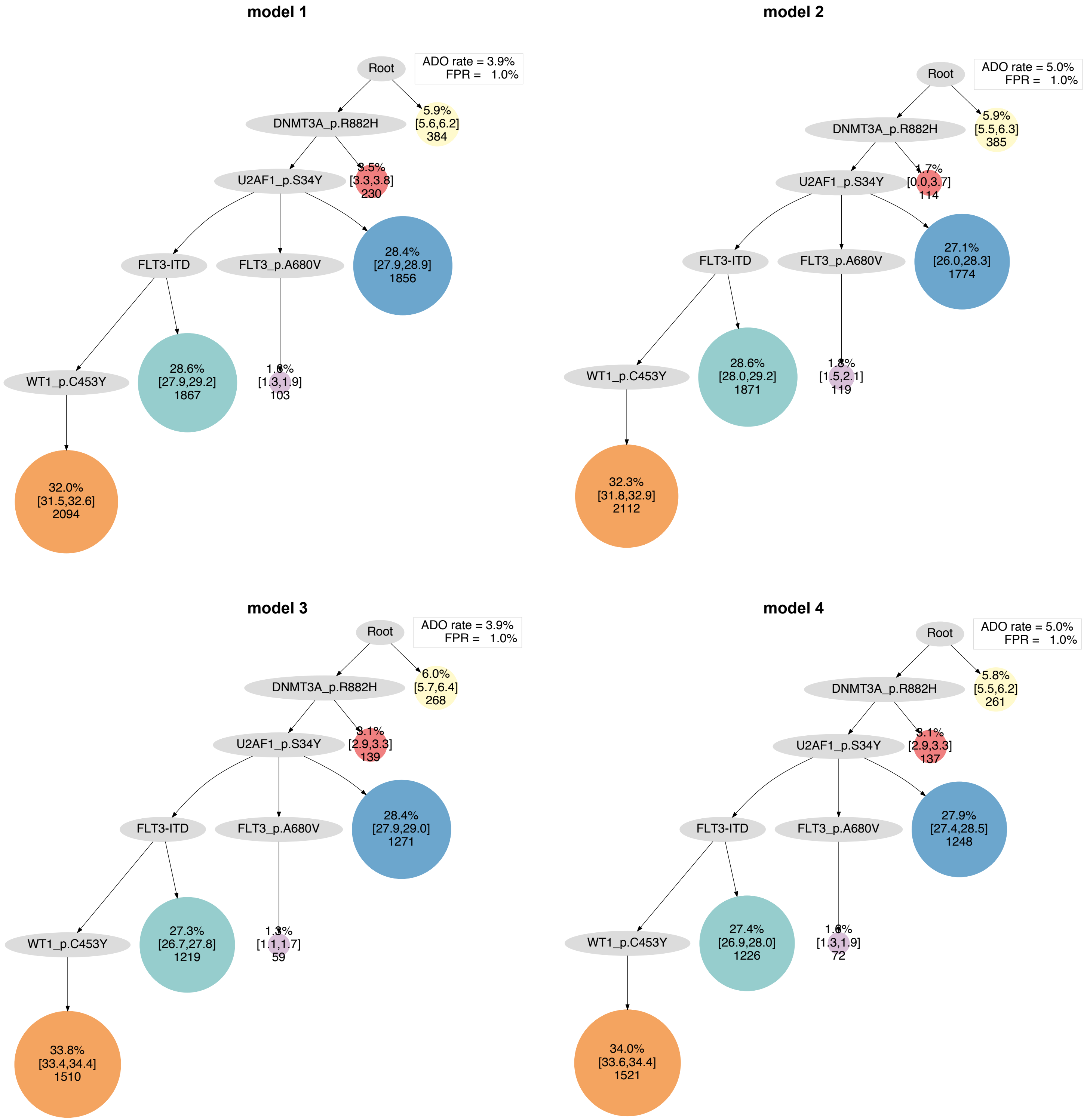

model 1
Root
ADO rate = 3.9%
FPR = 1.0%
DNMT3A_p.R882H
5.9%
[5.6,6.2]
384
U2AF1_p.S34Y
[3.3,3.8]
230
FLT3-ITD
FLT3_p.A680V
28.4%
[27.9,28.9]
1856
WT1_p.C453Y
28.6%
[27.9,29.2]
1867
[1.3,1.9]
103
32.0%
[31.5,32.6]
2094
model 2
Root
ADO rate = 5.0%
FPR = 1.0%
DNMT3A_p.R882H
5.9%
[5.5,6.3]
385
U2AF1_p.S34Y
1.7%
[0.0,3.7]
114
FLT3-ITD
FLT3_p.A680V
27.1%
[26.0,28.3]
1774
WT1_p.C453Y
28.6%
[28.0,29.2]
1871
[1.5,2.1]
119
32.3%
[31.8,32.9]
2112
model 3
Root
ADO rate = 3.9%
FPR = 1.0%
DNMT3A_p.R882H
6.0%
[5.7,6.4]
268
U2AF1_p.S34Y
[2.9,3.3]
139
FLT3-ITD
FLT3_p.A680V
28.4%
[27.9,29.0]
1271
WT1_p.C453Y
27.3%
[26.7,27.8]
1219
[1.1,1.7]
59
33.8%
[33.4,34.4]
1510
model 4
Root
ADO rate = 5.0%
FPR = 1.0%
DNMT3A_p.R882H
5.8%
[5.5,6.2]
261
U2AF1_p.S34Y
[2.9,3.3]
137
FLT3-ITD
FLT3_p.A680V
27.9%
[27.4,28.5]
1248
WT1_p.C453Y
27.4%
[26.9,28.0]
1226
[1.3,1.9]
72
34.0%
[33.6,34.4]
1521

**AML-25-001**

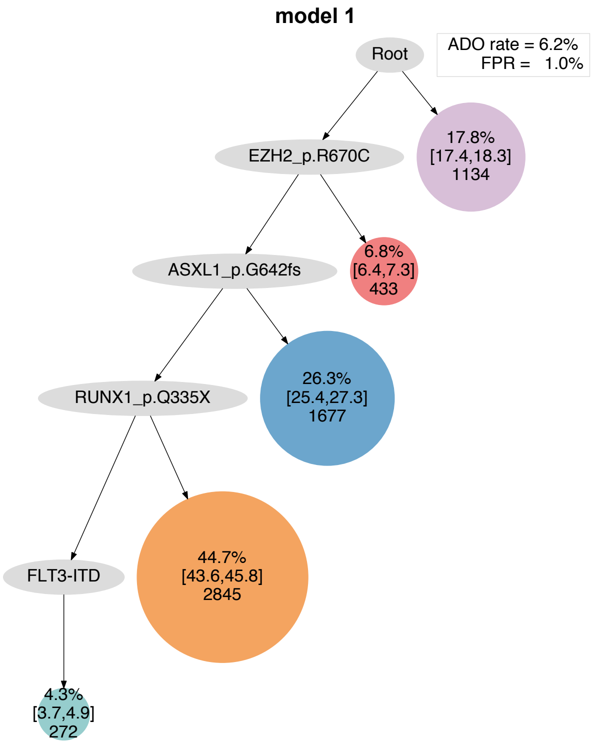
model 1
Root
ADO rate = 6.2%
FPR = 1.0%
EZH2_p.R670C
17.8%
[17.4,18.3]
1134
ASXL1_p.G642fs
6.8%
[6.4,7.3]
433
RUNX1_p.Q335X
26.3%
[25.4,27.3]
1677
FLT3-ITD
44.7%
[43.6,45.8]
2845
4.3%
[3.7,4.9]
272

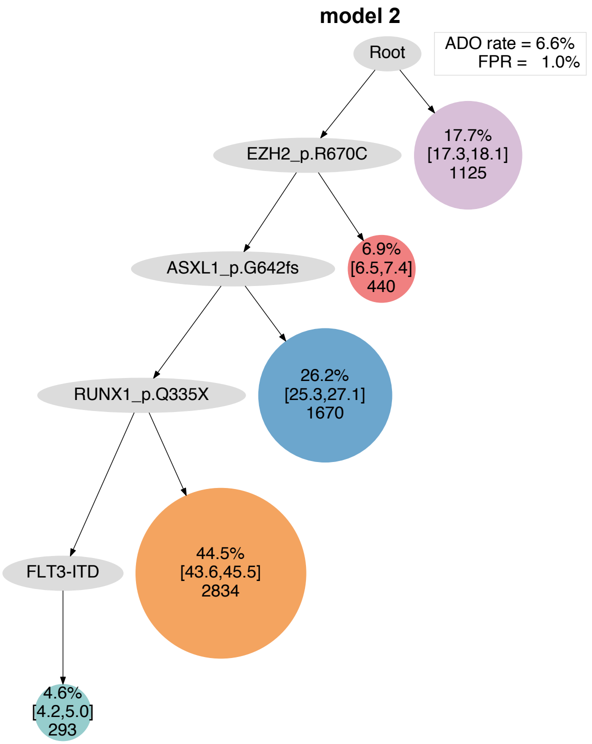
model 2
Root
ADO rate = 6.6%
FPR = 1.0%
EZH2_p.R670C
17.7%
[17.3,18.1]
1125
ASXL1_p.G642fs
6.9%
[6.5,7.4]
440
RUNX1_p.Q335X
26.2%
[25.3,27.1]
1670
FLT3-ITD
44.5%
[43.6,45.5]
2834
4.6%
[4.2,5.0]
293

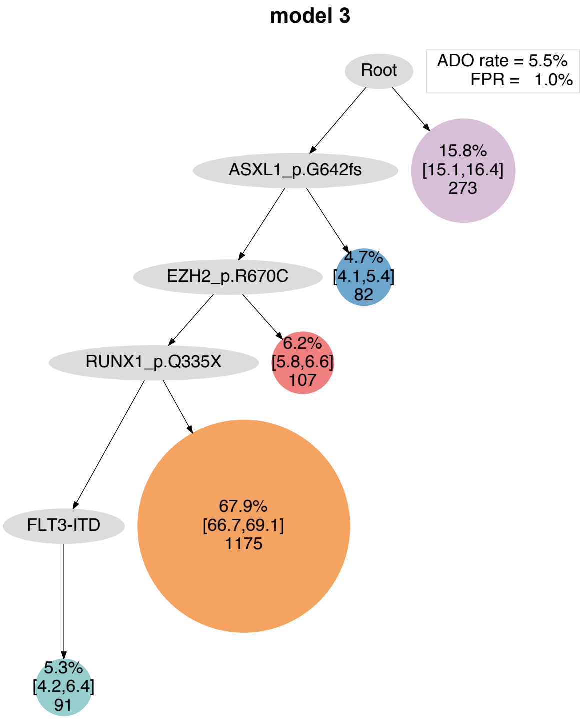
model 3
Root
ADO rate = 5.5%
FPR = 1.0%
ASXL1_p.G642fs
15.8%
[15.1,16.4]
273
EZH2_p.R670C
4.7%
[4.1,5.4]
82
RUNX1_p.Q335X
6.2%
[5.8,6.6]
107
FLT3-ITD
67.9%
[66.7,69.1]
1175
5.3%
[4.2,6.4]
91

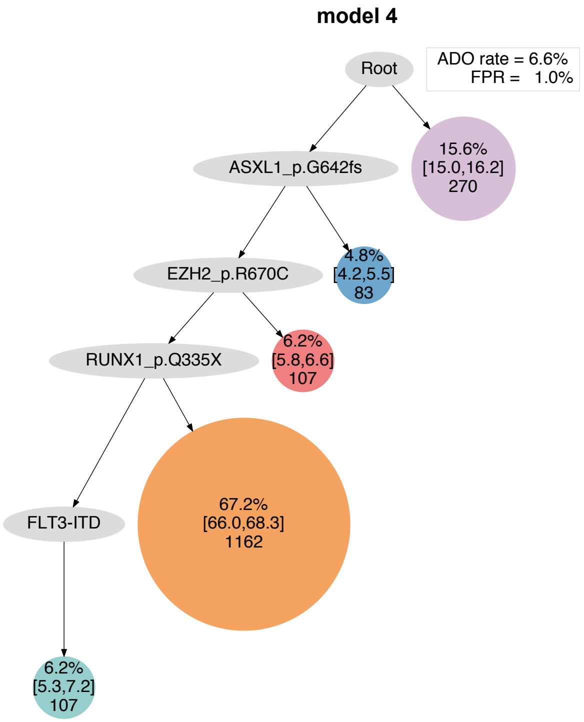
model 4
Root
ADO rate = 6.6%
FPR = 1.0%
ASXL1_p.G642fs
15.6%
[15.0,16.2]
270
EZH2_p.R670C
4.8%
[4.2,5.5]
83
RUNX1_p.Q335X
6.2%
[5.8,6.6]
107
FLT3-ITD
67.2%
[66.0,68.3]
1162
6.2%
[5.3,7.2]
107

### AML-26-001

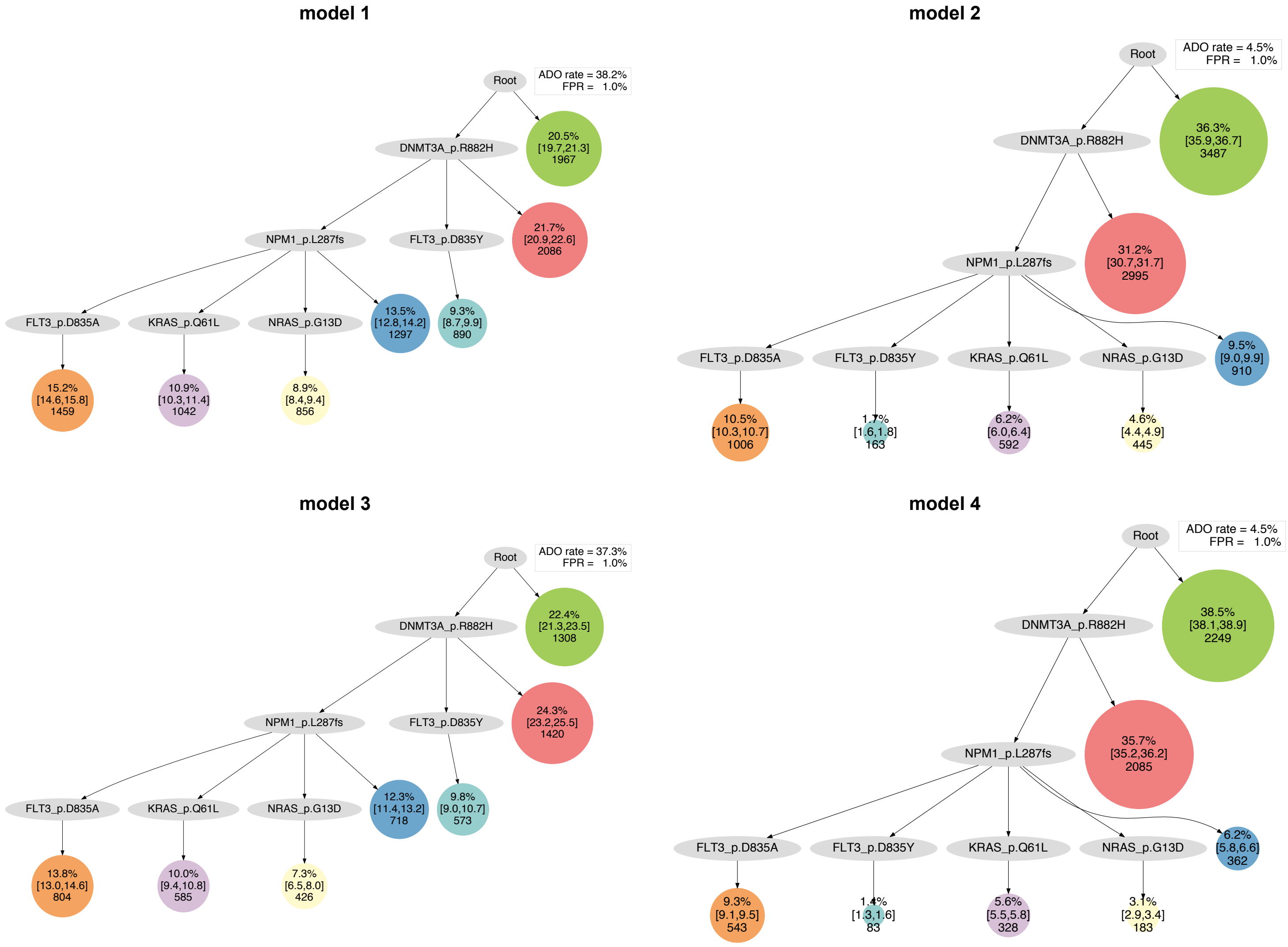

model 1
Root
ADO rate = 38.2%
FPR = 1.0%
DNMT3A_p.R882H
20.5%
[19.7,21.3]
1967
NPM1_p.L287fs
FLT3_p.D835Y
21.7%
[20.9,22.6]
2086
FLT3_p.D835A
KRAS_p.Q61L
NRAS_p.G13D
13.5%
[12.8,14.2]
1297
9.3%
[8.7,9.9]
890
15.2%
[14.6,15.8]
1459
10.9%
[10.3,11.4]
1042
8.9%
[8.4,9.4]
856
model 2
Root
ADO rate = 4.5%
FPR = 1.0%
DNMT3A_p.R882H
36.3%
[35.9,36.7]
3487
31.2%
[30.7,31.7]
2995
NPM1_p.L287fs
FLT3_p.D835A
FLT3_p.D835Y
KRAS_p.Q61L
NRAS_p.G13D
9.5%
[9.0,9.9]
910
10.5%
[10.3,10.7]
1006
1.7%
[1.6,1.8]
163
6.2%
[6.0,6.4]
592
4.6%
[4.4,4.9]
445
model 3
Root
ADO rate = 37.3%
FPR = 1.0%
DNMT3A_p.R882H
22.4%
[21.3,23.5]
1308
NPM1_p.L287fs
FLT3_p.D835Y
24.3%
[23.2,25.5]
1420
FLT3_p.D835A
KRAS_p.Q61L
NRAS_p.G13D
12.3%
[11.4,13.2]
718
9.8%
[9.0,10.7]
573
13.8%
[13.0,14.6]
804
10.0%
[9.4,10.8]
585
7.3%
[6.5,8.0]
426
model 4
Root
ADO rate = 4.5%
FPR = 1.0%
DNMT3A_p.R882H
38.5%
[38.1,38.9]
2249
35.7%
[35.2,36.2]
2085
NPM1_p.L287fs
FLT3_p.D835A
FLT3_p.D835Y
KRAS_p.Q61L
NRAS_p.G13D
6.2%
[5.8,6.6]
362
9.3%
[9.1,9.5]
543
1.4%
[1.3,1.6]
83
5.6%
[5.5,5.8]
328
3.1%
[2.9,3.4]
183

#### AML-27-001

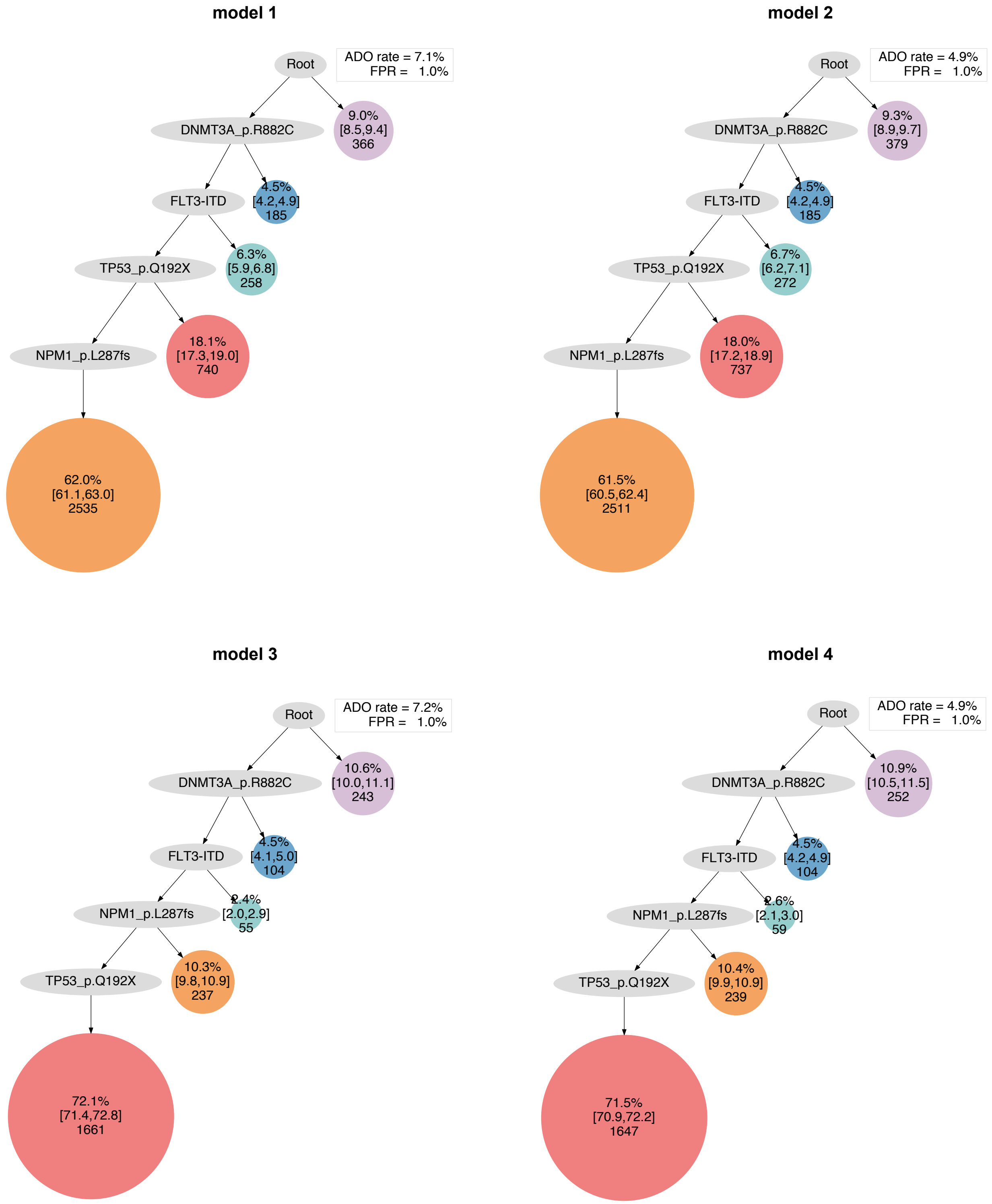

model 1
Root
ADO rate = 7.1%
FPR = 1.0%
DNMT3A_p.R882C
9.0%
[8.5,9.4]
366
FLT3-ITD
4.5%
[4.2,4.9]
185
TP53_p.Q192X
6.3%
[5.9,6.8]
258
NPM1_p.L287fs
18.1%
[17.3,19.0]
740
62.0%
[61.1,63.0]
2535
model 2
Root
ADO rate = 4.9%
FPR = 1.0%
DNMT3A_p.R882C
9.3%
[8.9,9.7]
379
FLT3-ITD
4.5%
[4.2,4.9]
185
TP53_p.Q192X
6.7%
[6.2,7.1]
272
NPM1_p.L287fs
18.0%
[17.2,18.9]
737
61.5%
[60.5,62.4]
2511
model 3
Root
ADO rate = 7.2%
FPR = 1.0%
DNMT3A_p.R882C
10.6%
[10.0,11.1]
243
FLT3-ITD
4.5%
[4.1,5.0]
104
NPM1_p.L287fs
2.4%
[2.0,2.9]
55
TP53_p.Q192X
10.3%
[9.8,10.9]
237
72.1%
[71.4,72.8]
1661
model 4
Root
ADO rate = 4.9%
FPR = 1.0%
DNMT3A_p.R882C
10.9%
[10.5,11.5]
252
FLT3-ITD
4.5%
[4.2,4.9]
104
NPM1_p.L287fs
2.6%
[2.1,3.0]
59
TP53_p.Q192X
10.4%
[9.9,10.9]
239
71.5%
[70.9,72.2]
1647

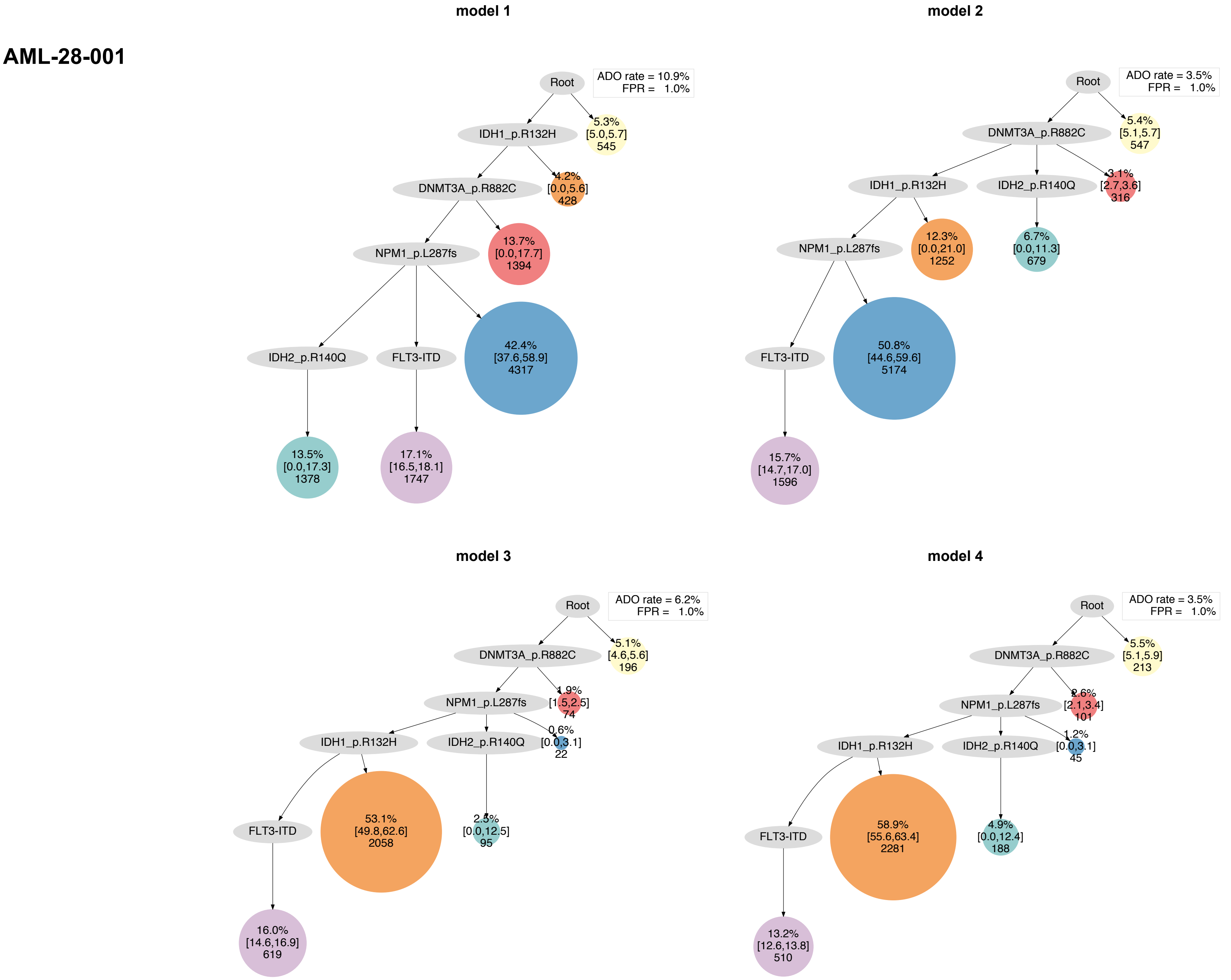
AML-28-001
model 1
Root
ADO rate = 10.9%
FPR = 1.0%
IDH1_p.R132H
5.3%
[5.0,5.7]
545
DNMT3A_p.R882C
4.2%
[0.0,5.6]
428
NPM1_p.L287fs
13.7%
[0.0,17.7]
1394
IDH2_p.R140Q
FLT3-ITD
42.4%
[37.6,58.9]
4317
13.5%
[0.0,17.3]
1378
17.1%
[16.5,18.1]
1747
model 2
Root
ADO rate = 3.5%
FPR = 1.0%
DNMT3A_p.R882C
5.4%
[5.1,5.7]
547
IDH1_p.R132H
IDH2_p.R140Q
3.1%
[2.7,3.6]
316
NPM1_p.L287fs
12.3%
[0.0,21.0]
1252
6.7%
[0.0,11.3]
679
FLT3-ITD
50.8%
[44.6,59.6]
5174
15.7%
[14.7,17.0]
1596
model 3
Root
ADO rate = 6.2%
FPR = 1.0%
DNMT3A_p.R882C
5.1%
[4.6,5.6]
196
NPM1_p.L287fs
1.9%
[1.5,2.5]
74
IDH1_p.R132H
IDH2_p.R140Q
0.6%
[0.0,3.1]
22
FLT3-ITD
53.1%
[49.8,62.6]
2058
2.5%
[0.0,12.5]
95
16.0%
[14.6,16.9]
619
model 4
Root
ADO rate = 3.5%
FPR = 1.0%
DNMT3A_p.R882C
5.5%
[5.1,5.9]
213
NPM1_p.L287fs
2.6%
[2.1,3.4]
101
IDH1_p.R132H
IDH2_p.R140Q
1.2%
[0.0,3.1]
45
FLT3-ITD
58.9%
[55.6,63.4]
2281
4.9%
[0.0,12.4]
188
13.2%
[12.6,13.8]
510

**AML-29-001**

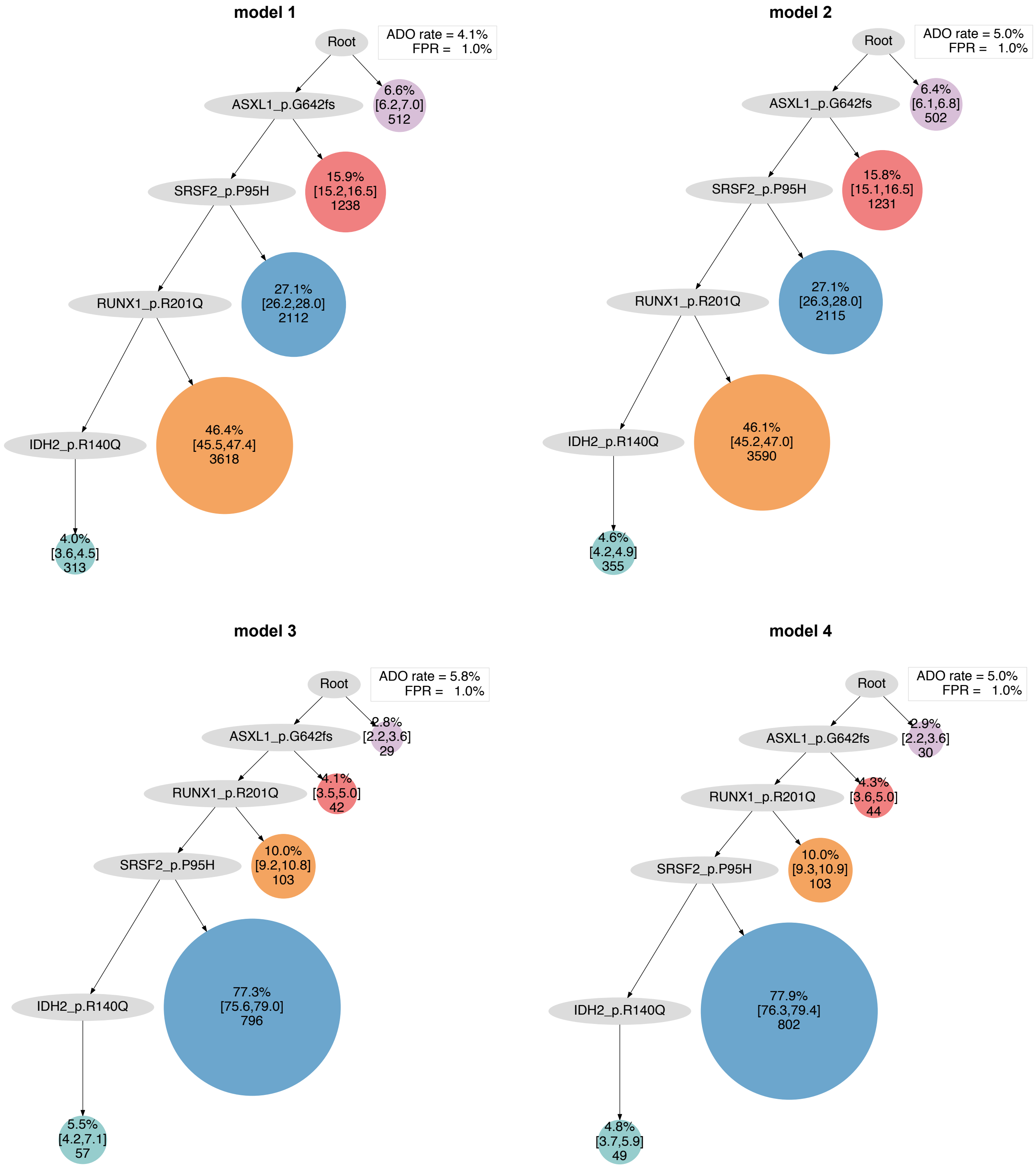
model 1
Root
ADO rate = 4.1%
FPR = 1.0%
ASXL1_p.G642fs
6.6%
[6.2,7.0]
512
SRSF2_p.P95H
15.9%
[15.2,16.5]
1238
RUNX1_p.R201Q
27.1%
[26.2,28.0]
2112
IDH2_p.R140Q
46.4%
[45.5,47.4]
3618
4.0%
[3.6,4.5]
313
model 2
Root
ADO rate = 5.0%
FPR = 1.0%
ASXL1_p.G642fs
6.4%
[6.1,6.8]
502
SRSF2_p.P95H
15.8%
[15.1,16.5]
1231
RUNX1_p.R201Q
27.1%
[26.3,28.0]
2115
IDH2_p.R140Q
46.1%
[45.2,47.0]
3590
4.6%
[4.2,4.9]
355
model 3
Root
ADO rate = 5.8%
FPR = 1.0%
ASXL1_p.G642fs
2.8%
[2.2,3.6]
29
RUNX1_p.R201Q
4.1%
[3.5,5.0]
42
SRSF2_p.P95H
10.0%
[9.2,10.8]
103
IDH2_p.R140Q
77.3%
[75.6,79.0]
796
5.5%
[4.2,7.1]
57
model 4
Root
ADO rate = 5.0%
FPR = 1.0%
ASXL1_p.G642fs
2.9%
[2.2,3.6]
30
RUNX1_p.R201Q
4.3%
[3.6,5.0]
44
SRSF2_p.P95H
10.0%
[9.3,10.9]
103
IDH2_p.R140Q
77.9%
[76.3,79.4]
802
4.8%
[3.7,5.9]
49

### AML-30-001

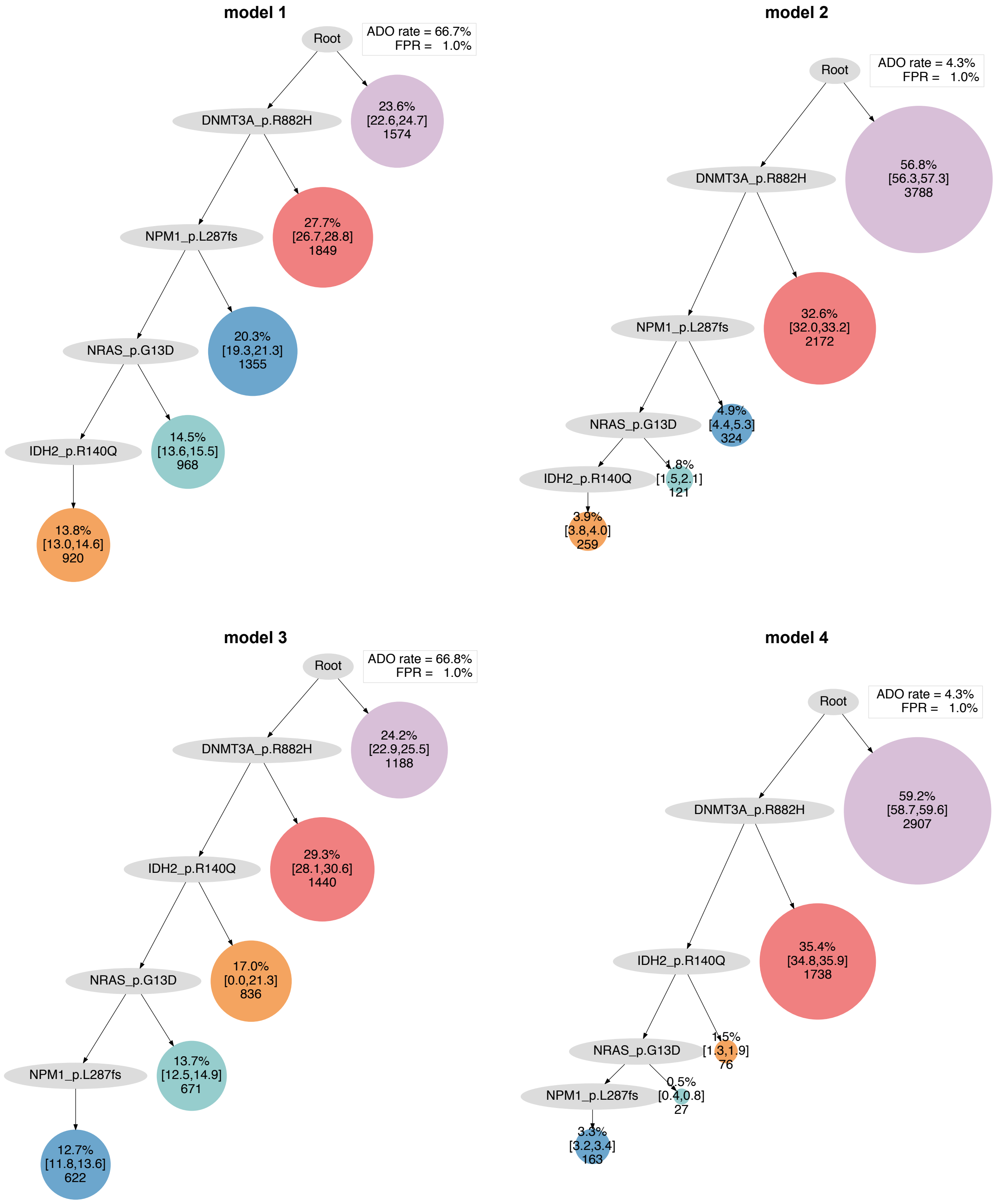

model 1
Root
ADO rate = 66.7%
FPR = 1.0%
DNMT3A_p.R882H
23.6%
[22.6,24.7]
1574
NPM1_p.L287fs
27.7%
[26.7,28.8]
1849
NRAS_p.G13D
20.3%
[19.3,21.3]
1355
IDH2_p.R140Q
14.5%
[13.6,15.5]
968
13.8%
[13.0,14.6]
920
model 2
Root
ADO rate = 4.3%
FPR = 1.0%
DNMT3A_p.R882H
56.8%
[56.3,57.3]
3788
NPM1_p.L287fs
32.6%
[32.0,33.2]
2172
NRAS_p.G13D
4.9%
[4.4,5.3]
324
IDH2_p.R140Q
1.8%
[1.5,2.1]
121
3.9%
[3.8,4.0]
259
model 3
Root
ADO rate = 66.8%
FPR = 1.0%
DNMT3A_p.R882H
24.2%
[22.9,25.5]
1188
IDH2_p.R140Q
29.3%
[28.1,30.6]
1440
NRAS_p.G13D
17.0%
[0.0,21.3]
836
NPM1_p.L287fs
13.7%
[12.5,14.9]
671
12.7%
[11.8,13.6]
622
model 4
Root
ADO rate = 4.3%
FPR = 1.0%
DNMT3A_p.R882H
59.2%
[58.7,59.6]
2907
IDH2_p.R140Q
35.4%
[34.8,35.9]
1738
NRAS_p.G13D
1.5%
[1.3,1.9]
76
NPM1_p.L287fs
0.5%
[0.4,0.8]
27
3.3%
[3.2,3.4]
163

### AML-31-001

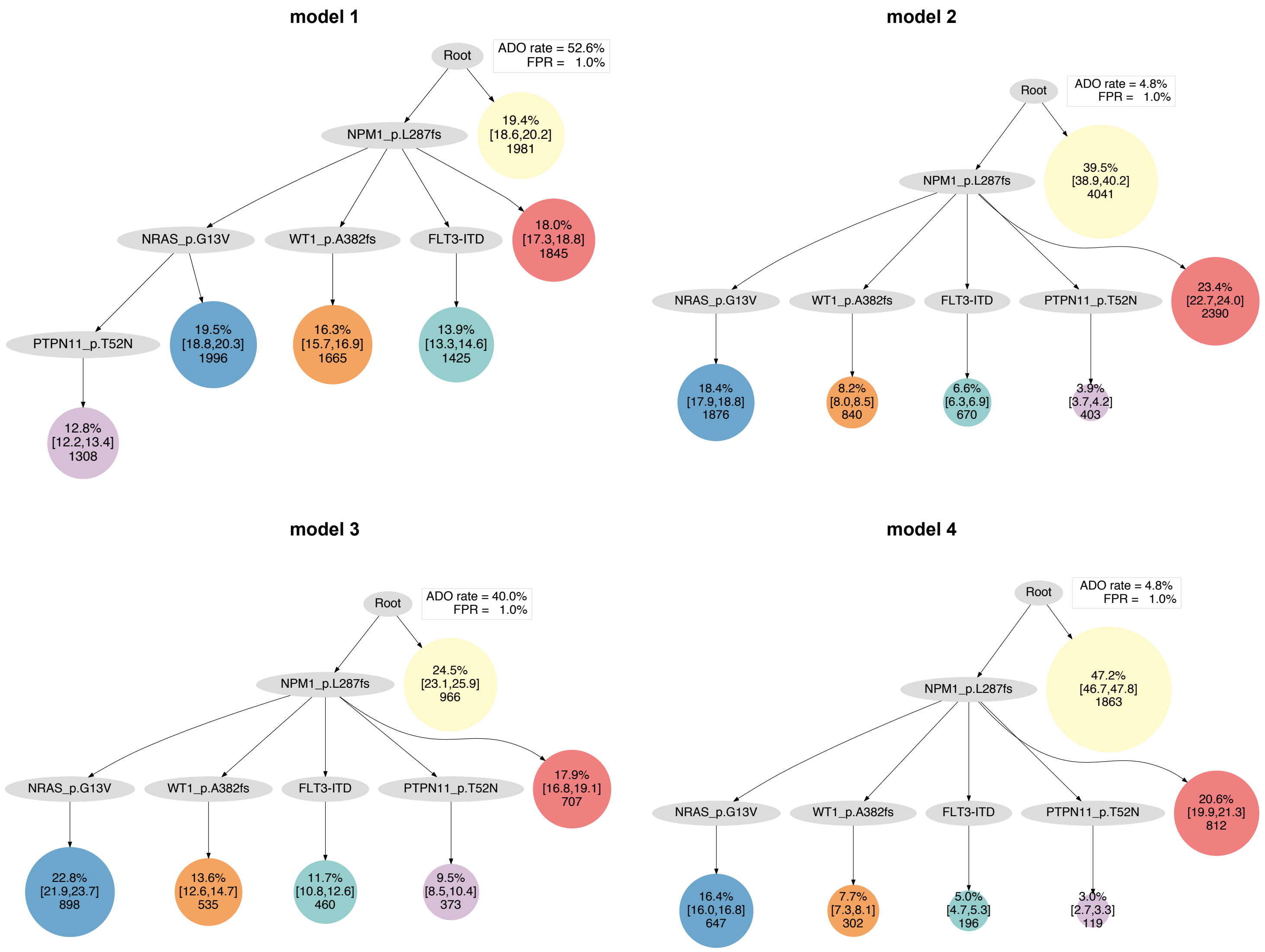

model 1
Root
ADO rate = 52.6%
FPR = 1.0%
NPM1_p.L287fs
19.4% [18.6,20.2] 1981
NRAS_p.G13V
WT1_p.A382fs
FLT3-ITD
18.0% [17.3,18.8] 1845
PTPN11_p.T52N
19.5% [18.8,20.3] 1996
16.3% [15.7,16.9] 1665
13.9% [13.3,14.6] 1425
12.8% [12.2,13.4] 1308
model 2
Root
ADO rate = 4.8%
FPR = 1.0%
NPM1_p.L287fs
39.5% [38.9,40.2] 4041
NRAS_p.G13V
WT1_p.A382fs
FLT3-ITD
PTPN11_p.T52N
23.4% [22.7,24.0] 2390
18.4% [17.9,18.8] 1876
8.2% [8.0,8.5] 840
6.6% [6.3,6.9] 670
3.9% [3.7,4.2] 403
model 3
Root
ADO rate = 40.0%
FPR = 1.0%
NPM1_p.L287fs
24.5% [23.1,25.9] 966
NRAS_p.G13V
WT1_p.A382fs
FLT3-ITD
PTPN11_p.T52N
17.9% [16.8,19.1] 707
22.8% [21.9,23.7] 898
13.6% [12.6,14.7] 535
11.7% [10.8,12.6] 460
9.5% [8.5,10.4] 373
model 4
Root
ADO rate = 4.8%
FPR = 1.0%
NPM1_p.L287fs
47.2% [46.7,47.8] 1863
NRAS_p.G13V
WT1_p.A382fs
FLT3-ITD
PTPN11_p.T52N
20.6% [19.9,21.3] 812
16.4% [16.0,16.8] 647
7.7% [7.3,8.1] 302
5.0% [4.7,5.3] 196
3.0% [2.7,3.3] 119

### AML-32-001

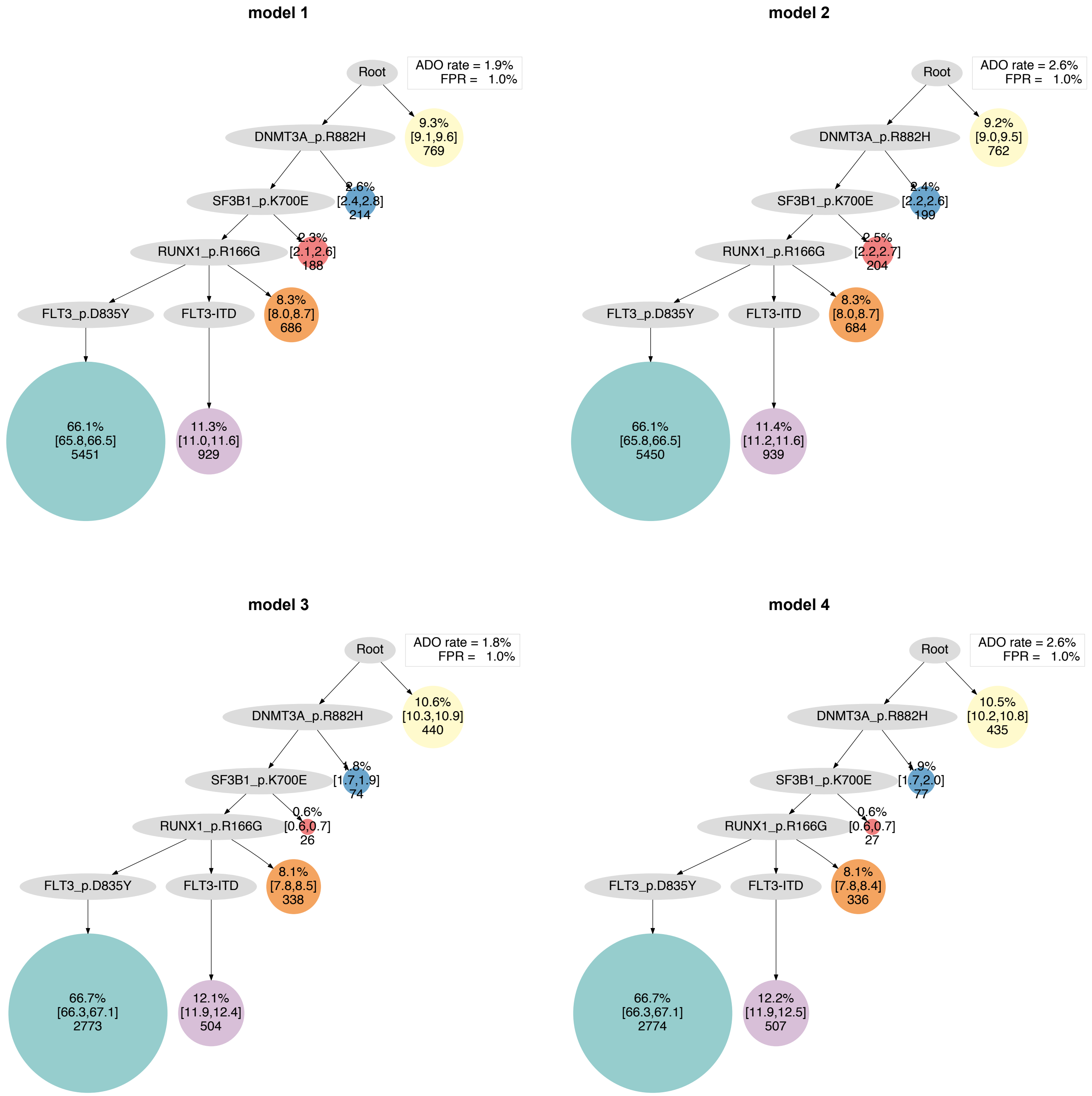

model 1
Root
ADO rate = 1.9%
FPR = 1.0%
DNMT3A_p.R882H
9.3% [9.1,9.6] 769
SF3B1_p.K700E
2.6% [2.4,2.8] 214
RUNX1_p.R166G
2.3% [2.1,2.6] 188
FLT3_p.D835Y
FLT3-ITD
8.3% [8.0,8.7] 686
66.1% [65.8,66.5] 5451
11.3% [11.0,11.6] 929
model 2
Root
ADO rate = 2.6%
FPR = 1.0%
DNMT3A_p.R882H
9.2% [9.0,9.5] 762
SF3B1_p.K700E
2.4% [2.2,2.6] 199
RUNX1_p.R166G
2.5% [2.2,2.7] 204
FLT3_p.D835Y
FLT3-ITD
8.3% [8.0,8.7] 684
66.1% [65.8,66.5] 5450
11.4% [11.2,11.6] 939
model 3
Root
ADO rate = 1.8%
FPR = 1.0%
DNMT3A_p.R882H
10.6% [10.3,10.9] 440
SF3B1_p.K700E
1.8% [1.7,1.9] 74
RUNX1_p.R166G
0.6% [0.6,0.7] 26
FLT3_p.D835Y
FLT3-ITD
8.1% [7.8,8.5] 338
66.7% [66.3,67.1] 2773
12.1% [11.9,12.4] 504
model 4
Root
ADO rate = 2.6%
FPR = 1.0%
DNMT3A_p.R882H
10.5% [10.2,10.8] 435
SF3B1_p.K700E
1.9% [1.7,2.0] 77
RUNX1_p.R166G
0.6% [0.6,0.7] 27
FLT3_p.D835Y
FLT3-ITD
8.1% [7.8,8.4] 336
66.7% [66.3,67.1] 2774
12.2% [11.9,12.5] 507

#### AML-33-001

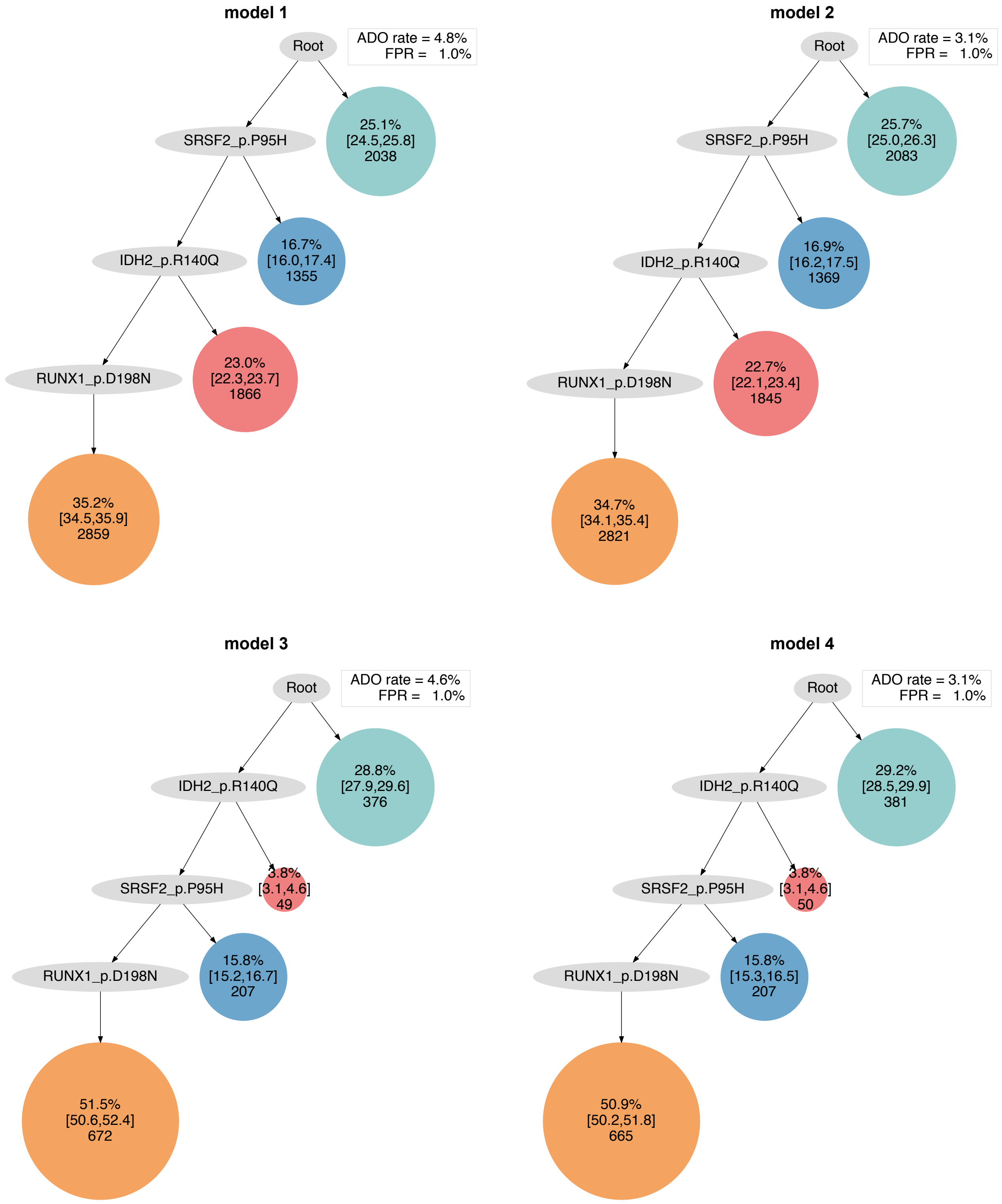
model 1
Root
ADO rate = 4.8%
FPR = 1.0%
SRSF2_p.P95H
25.1%
[24.5,25.8]
2038
IDH2_p.R140Q
16.7%
[16.0,17.4]
1355
RUNX1_p.D198N
23.0%
[22.3,23.7]
1866
35.2%
[34.5,35.9]
2859
model 2
Root
ADO rate = 3.1%
FPR = 1.0%
SRSF2_p.P95H
25.7%
[25.0,26.3]
2083
IDH2_p.R140Q
16.9%
[16.2,17.5]
1369
RUNX1_p.D198N
22.7%
[22.1,23.4]
1845
34.7%
[34.1,35.4]
2821
model 3
Root
ADO rate = 4.6%
FPR = 1.0%
IDH2_p.R140Q
28.8%
[27.9,29.6]
376
SRSF2_p.P95H
3.8%
[3.1,4.6]
49
RUNX1_p.D198N
15.8%
[15.2,16.7]
207
51.5%
[50.6,52.4]
672
model 4
Root
ADO rate = 3.1%
FPR = 1.0%
IDH2_p.R140Q
29.2%
[28.5,29.9]
381
SRSF2_p.P95H
3.8%
[3.1,4.6]
50
RUNX1_p.D198N
15.8%
[15.3,16.5]
207
50.9%
[50.2,51.8]
665

### AML-34-001

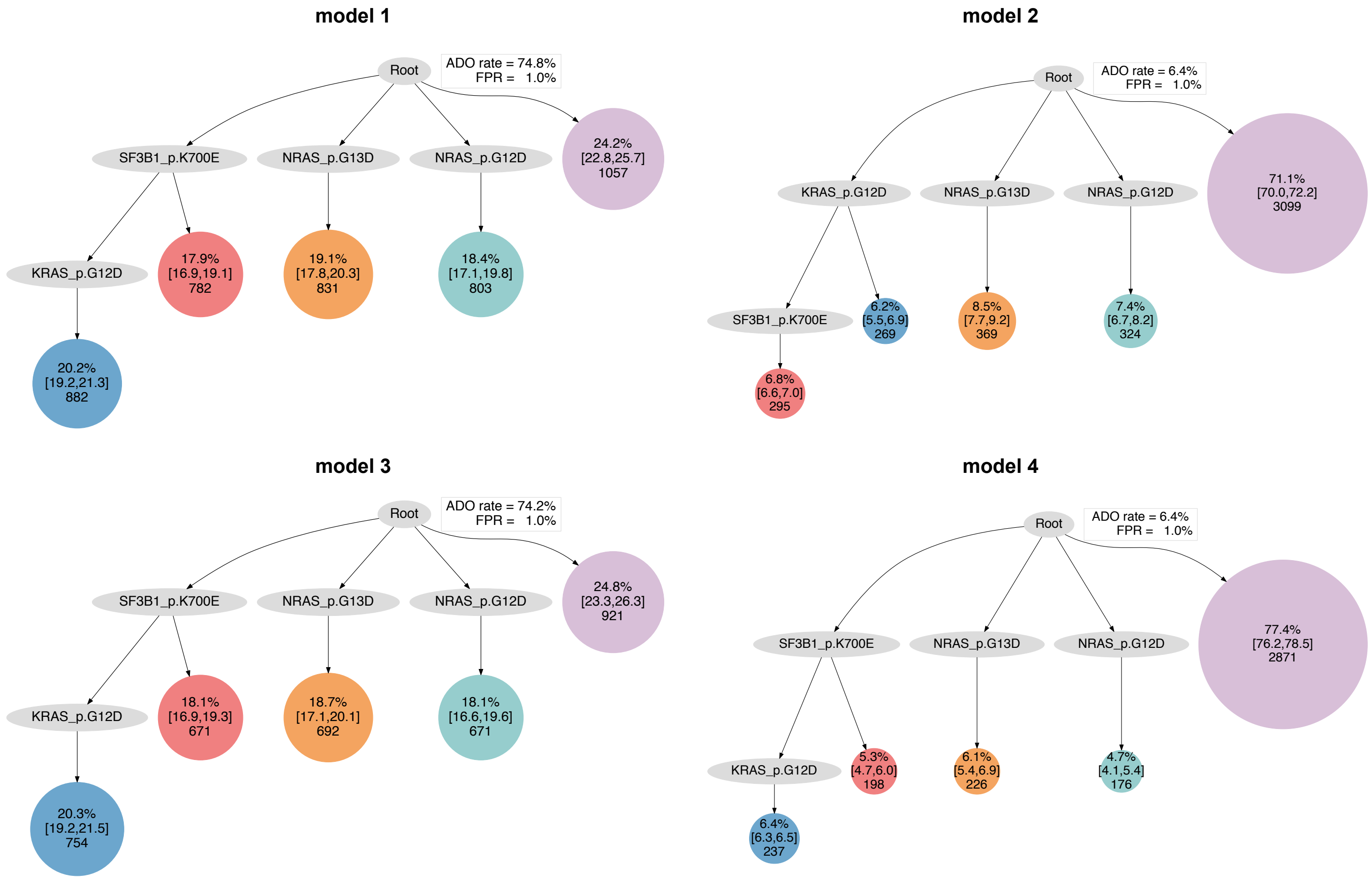

model 1
Root
ADO rate = 74.8%
FPR = 1.0%
SF3B1_p.K700E
NRAS_p.G13D
NRAS_p.G12D
24.2% [22.8,25.7] 1057
KRAS_p.G12D
17.9% [16.9,19.1] 782
19.1% [17.8,20.3] 831
18.4% [17.1,19.8] 803
20.2% [19.2,21.3] 882
model 2
Root
ADO rate = 6.4%
FPR = 1.0%
KRAS_p.G12D
NRAS_p.G13D
NRAS_p.G12D
71.1% [70.0,72.2] 3099
SF3B1_p.K700E
6.2% [5.5,6.9] 269
8.5% [7.7,9.2] 369
7.4% [6.7,8.2] 324
6.8% [6.6,7.0] 295
model 3
Root
ADO rate = 74.2%
FPR = 1.0%
SF3B1_p.K700E
NRAS_p.G13D
NRAS_p.G12D
24.8% [23.3,26.3] 921
KRAS_p.G12D
18.1% [16.9,19.3] 671
18.7% [17.1,20.1] 692
18.1% [16.6,19.6] 671
20.3% [19.2,21.5] 754
model 4
Root
ADO rate = 6.4%
FPR = 1.0%
SF3B1_p.K700E
NRAS_p.G13D
NRAS_p.G12D
77.4% [76.2,78.5] 2871
KRAS_p.G12D
5.3% [4.7,6.0] 198
6.1% [5.4,6.9] 226
4.7% [4.1,5.4] 176
6.4% [6.3,6.5] 237

**AML-35-001**

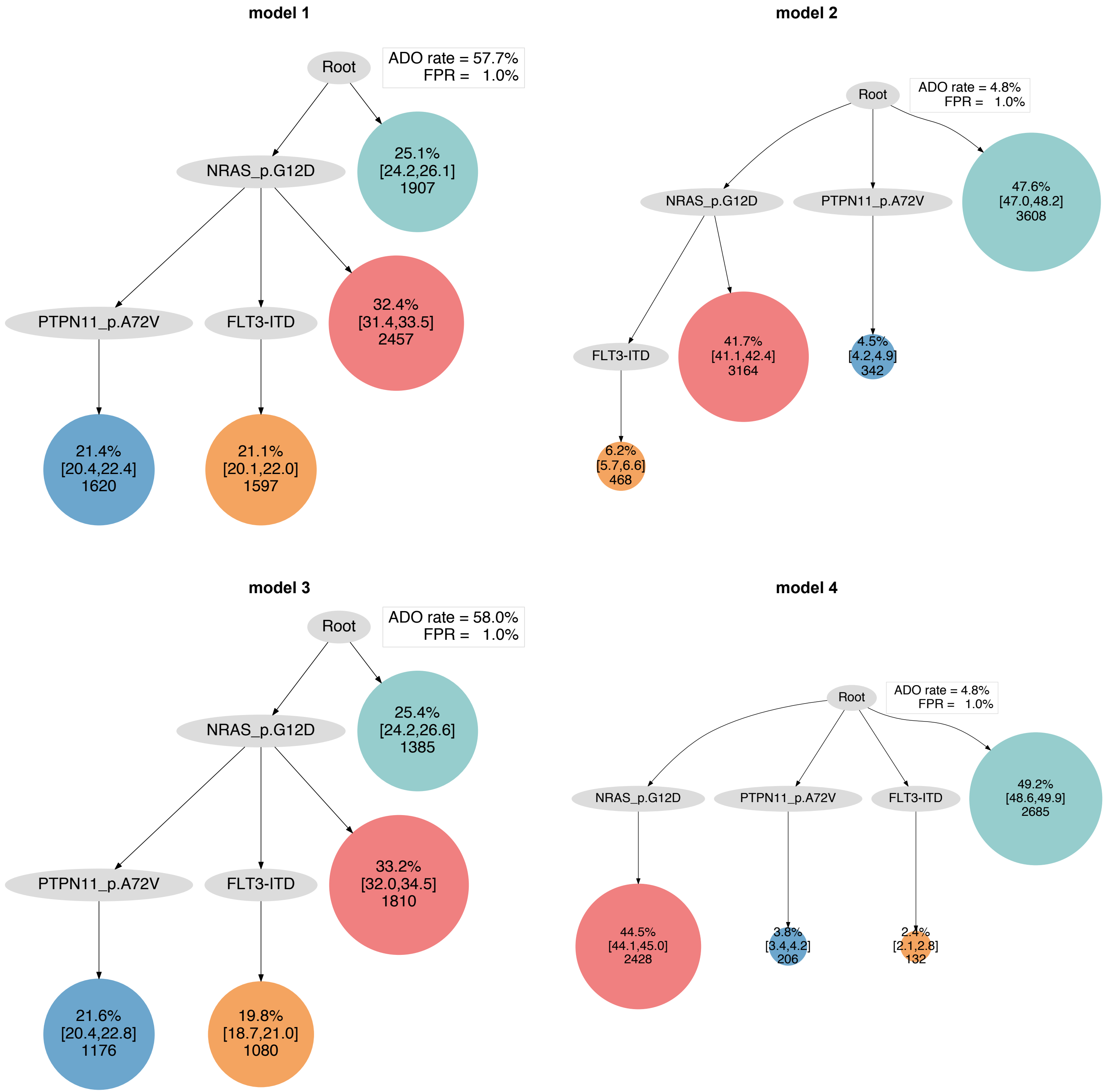
model 1
Root
ADO rate = 57.7%
FPR = 1.0%
NRAS_p.G12D
25.1%
[24.2,26.1]
1907
PTPN11_p.A72V
FLT3-ITD
32.4%
[31.4,33.5]
2457
21.4%
[20.4,22.4]
1620
21.1%
[20.1,22.0]
1597
model 2
Root
ADO rate = 4.8%
FPR = 1.0%
NRAS_p.G12D
PTPN11_p.A72V
47.6%
[47.0,48.2]
3608
FLT3-ITD
41.7%
[41.1,42.4]
3164
4.5%
[4.2,4.9]
342
6.2%
[5.7,6.6]
468
model 3
Root
ADO rate = 58.0%
FPR = 1.0%
NRAS_p.G12D
25.4%
[24.2,26.6]
1385
PTPN11_p.A72V
FLT3-ITD
33.2%
[32.0,34.5]
1810
21.6%
[20.4,22.8]
1176
19.8%
[18.7,21.0]
1080
model 4
Root
ADO rate = 4.8%
FPR = 1.0%
NRAS_p.G12D
PTPN11_p.A72V
FLT3-ITD
49.2%
[48.6,49.9]
2685
44.5%
[44.1,45.0]
2428
3.8%
[3.4,4.2]
206
2.4%
[2.1,2.8]
132

### AML-36-001

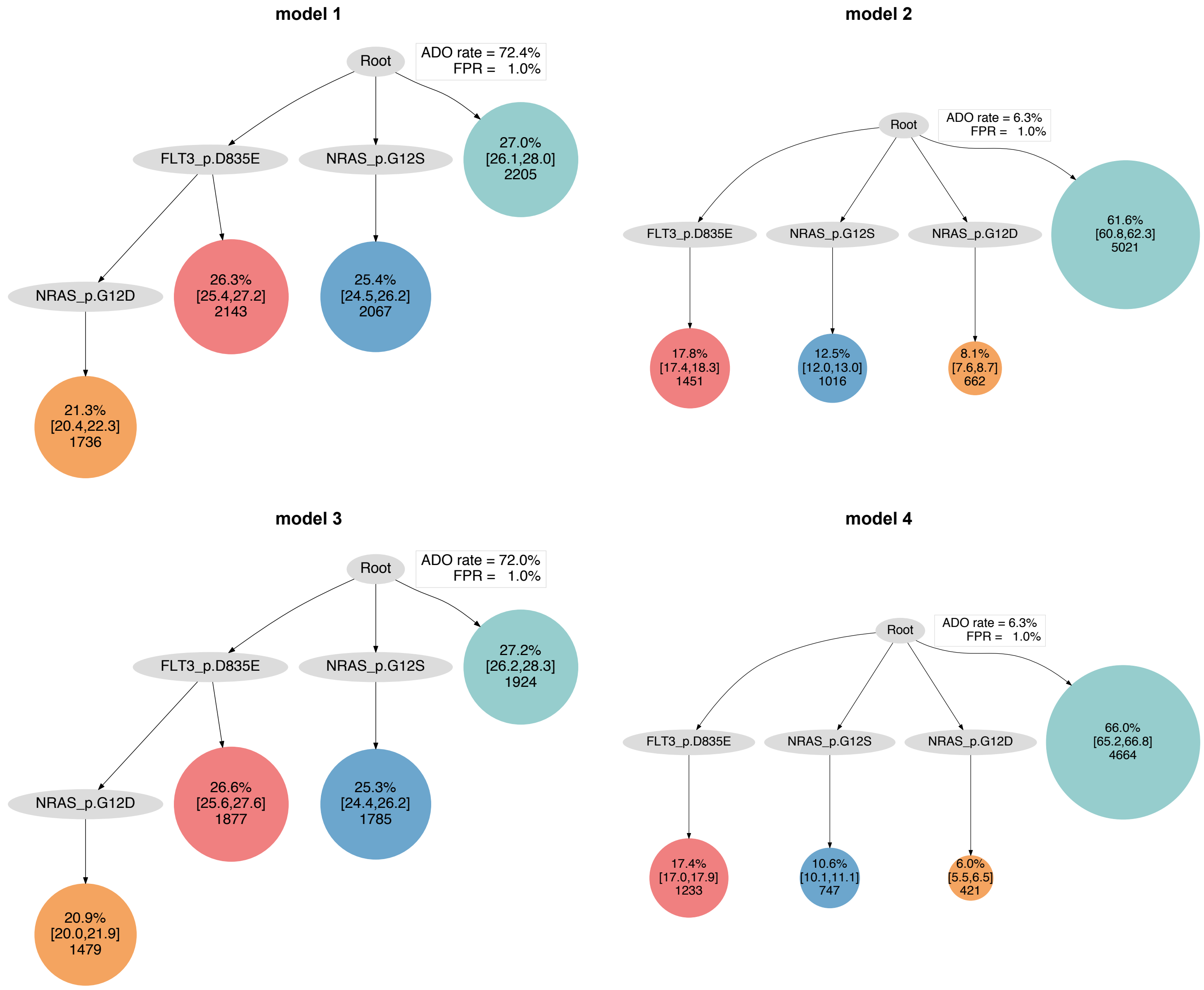

model 1
Root
ADO rate = 72.4%
FPR = 1.0%
FLT3_p.D835E
NRAS_p.G12S
27.0%
[26.1,28.0]
2205
NRAS_p.G12D
26.3%
[25.4,27.2]
2143
25.4%
[24.5,26.2]
2067
21.3%
[20.4,22.3]
1736
model 2
Root
ADO rate = 6.3%
FPR = 1.0%
FLT3_p.D835E
NRAS_p.G12S
NRAS_p.G12D
61.6%
[60.8,62.3]
5021
17.8%
[17.4,18.3]
1451
12.5%
[12.0,13.0]
1016
8.1%
[7.6,8.7]
662
model 3
Root
ADO rate = 72.0%
FPR = 1.0%
FLT3_p.D835E
NRAS_p.G12S
27.2%
[26.2,28.3]
1924
NRAS_p.G12D
26.6%
[25.6,27.6]
1877
25.3%
[24.4,26.2]
1785
20.9%
[20.0,21.9]
1479
model 4
Root
ADO rate = 6.3%
FPR = 1.0%
FLT3_p.D835E
NRAS_p.G12S
NRAS_p.G12D
66.0%
[65.2,66.8]
4664
17.4%
[17.0,17.9]
1233
10.6%
[10.1,11.1]
747
6.0%
[5.5,6.5]
421

### AML-37-001

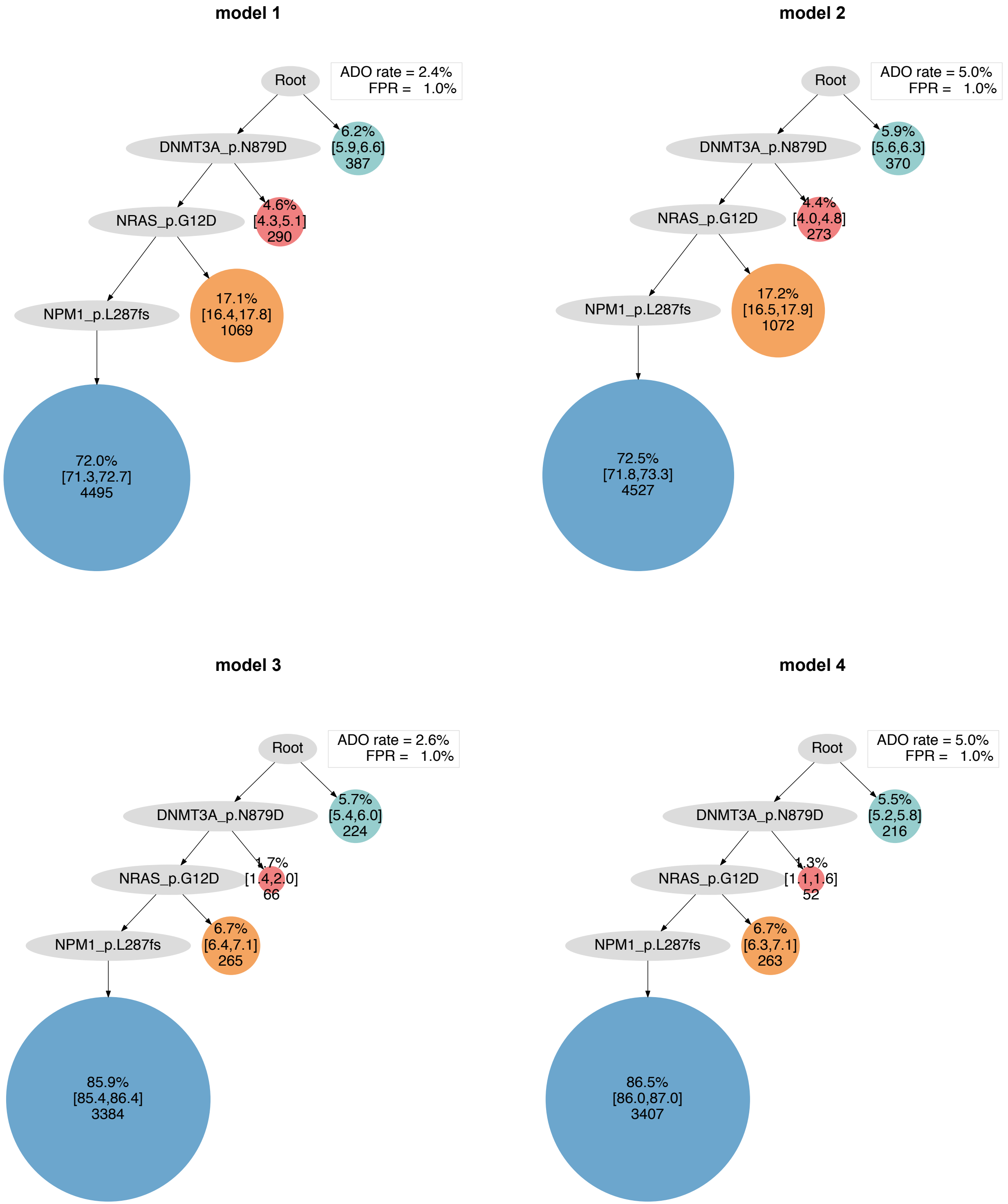

model 1
Root
ADO rate = 2.4%
FPR = 1.0%
DNMT3A_p.N879D
6.2% [5.9,6.6] 387
NRAS_p.G12D
4.6% [4.3,5.1] 290
NPM1_p.L287fs
17.1% [16.4,17.8] 1069
72.0% [71.3,72.7] 4495
model 2
Root
ADO rate = 5.0%
FPR = 1.0%
DNMT3A_p.N879D
5.9% [5.6,6.3] 370
NRAS_p.G12D
4.4% [4.0,4.8] 273
NPM1_p.L287fs
17.2% [16.5,17.9] 1072
72.5% [71.8,73.3] 4527
model 3
Root
ADO rate = 2.6%
FPR = 1.0%
DNMT3A_p.N879D
5.7% [5.4,6.0] 224
NRAS_p.G12D
1.7% [1.4,2.0] 66
NPM1_p.L287fs
6.7% [6.4,7.1] 265
85.9% [85.4,86.4] 3384
model 4
Root
ADO rate = 5.0%
FPR = 1.0%
DNMT3A_p.N879D
5.5% [5.2,5.8] 216
NRAS_p.G12D
1.3% [1.1,1.6] 52
NPM1_p.L287fs
6.7% [6.3,7.1] 263
86.5% [86.0,87.0] 3407

### AML-38-001

#### model 1

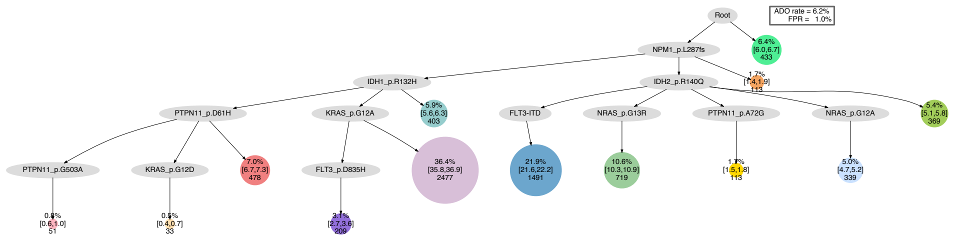


#### model 2

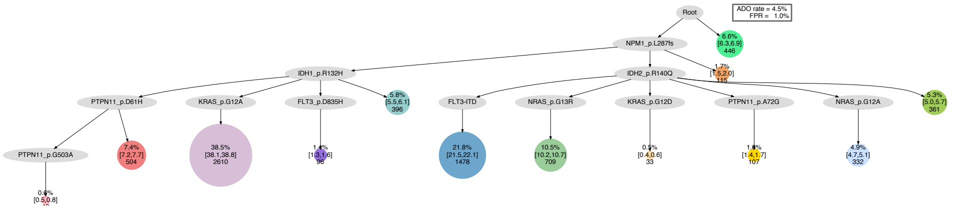


#### model 3

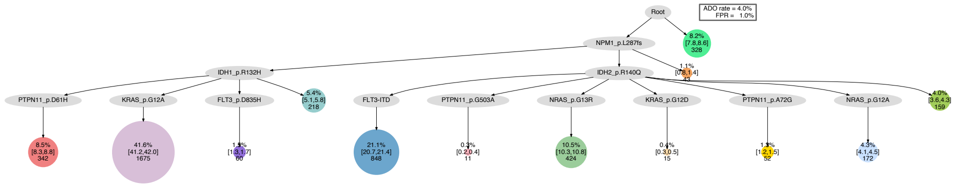


#### model 4

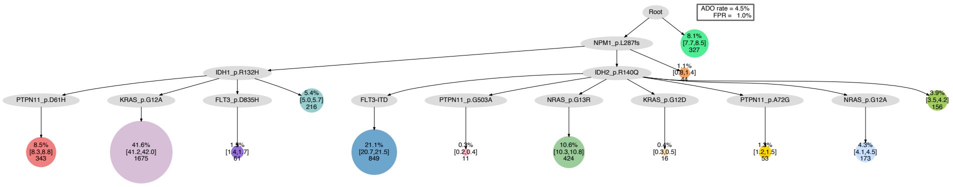

### AML-39-001

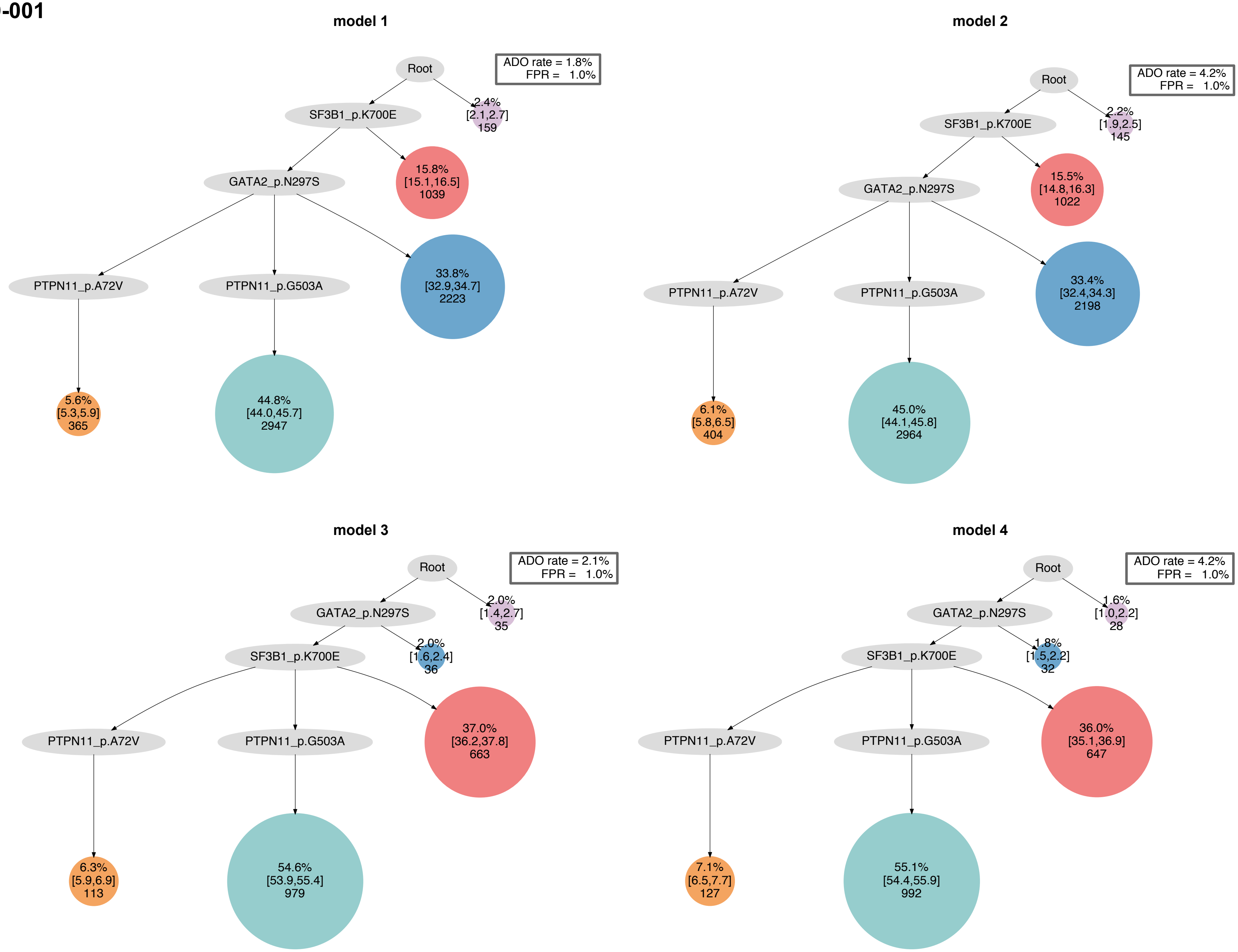

model 1
Root
ADO rate = 1.8%
FPR = 1.0%
SF3B1_p.K700E
2.4%
[2.1,2.7]
159
GATA2_p.N297S
15.8%
[15.1,16.5]
1039
PTPN11_p.A72V
PTPN11_p.G503A
33.8%
[32.9,34.7]
2223
5.6%
[5.3,5.9]
365
44.8%
[44.0,45.7]
2947
model 2
Root
ADO rate = 4.2%
FPR = 1.0%
SF3B1_p.K700E
2.2%
[1.9,2.5]
145
GATA2_p.N297S
15.5%
[14.8,16.3]
1022
PTPN11_p.A72V
PTPN11_p.G503A
33.4%
[32.4,34.3]
2198
6.1%
[5.8,6.5]
404
45.0%
[44.1,45.8]
2964
model 3
Root
ADO rate = 2.1%
FPR = 1.0%
GATA2_p.N297S
2.0%
[1.4,2.7]
35
SF3B1_p.K700E
2.0%
[1.6,2.4]
36
PTPN11_p.A72V
PTPN11_p.G503A
37.0%
[36.2,37.8]
663
6.3%
[5.9,6.9]
113
54.6%
[53.9,55.4]
979
model 4
Root
ADO rate = 4.2%
FPR = 1.0%
GATA2_p.N297S
1.6%
[1.0,2.2]
28
SF3B1_p.K700E
1.8%
[1.5,2.2]
32
PTPN11_p.A72V
PTPN11_p.G503A
36.0%
[35.1,36.9]
647
7.1%
[6.5,7.7]
127
55.1%
[54.4,55.9]
992

### AML-40-001

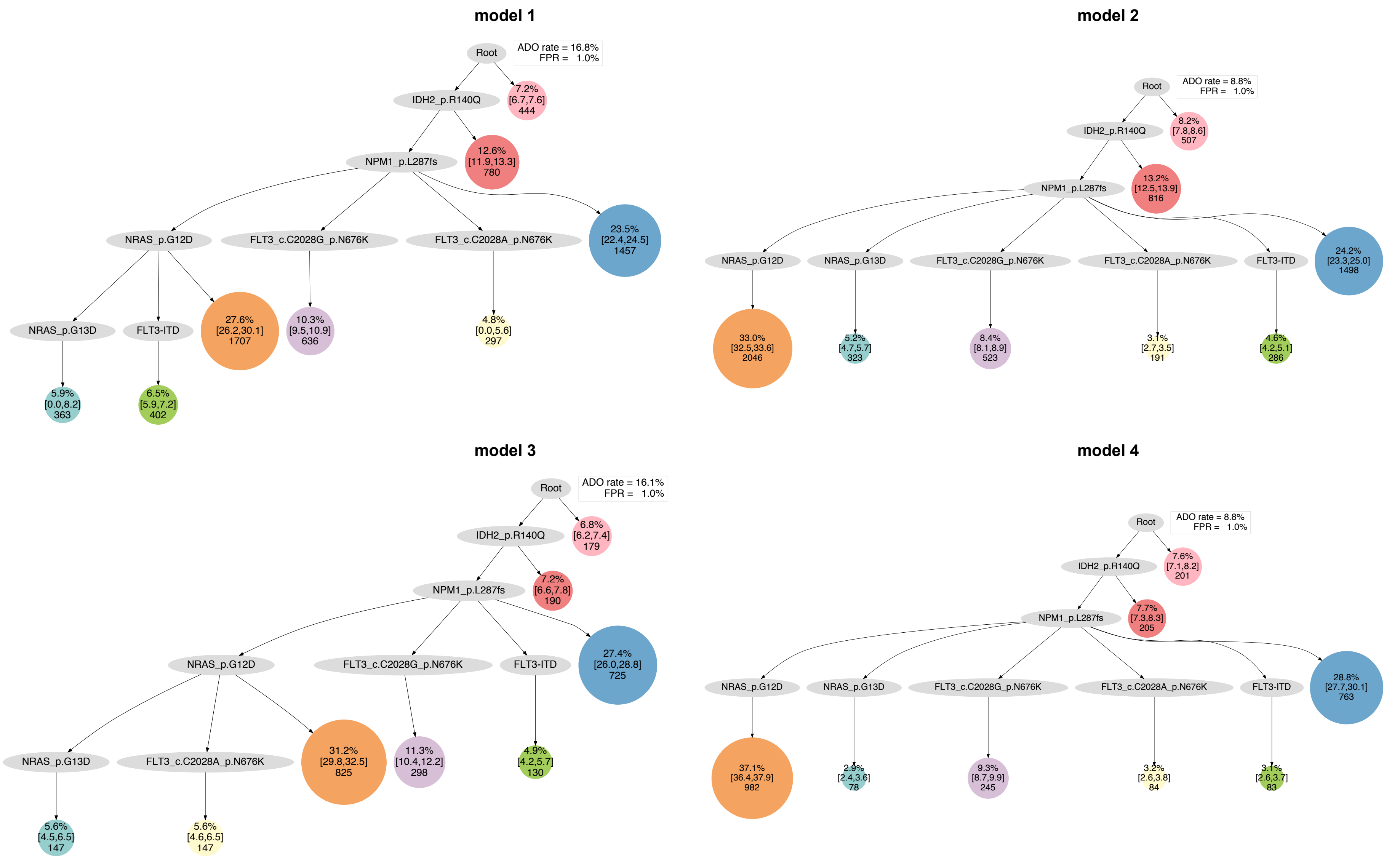

model 1
Root
ADO rate = 16.8%
FPR = 1.0%
IDH2_p.R140Q
7.2% [6.7,7.6] 444
NPM1_p.L287fs
12.6% [11.9,13.3] 780
NRAS_p.G12D
FLT3_c.C2028G_p.N676K
FLT3_c.C2028A_p.N676K
23.5% [22.4,24.5] 1457
NRAS_p.G13D
FLT3-ITD
27.6% [26.2,30.1] 1707
10.3% [9.5,10.9] 636
4.8% [0.0,5.6] 297
5.9% [0.0,8.2] 363
6.5% [5.9,7.2] 402
model 2
Root
ADO rate = 8.8%
FPR = 1.0%
IDH2_p.R140Q
8.2% [7.8,8.6] 507
NPM1_p.L287fs
13.2% [12.5,13.9] 816
NRAS_p.G12D
NRAS_p.G13D
FLT3_c.C2028G_p.N676K
FLT3_c.C2028A_p.N676K
FLT3-ITD
24.2% [23.3,25.0] 1498
33.0% [32.5,33.6] 2046
5.2% [4.7,5.7] 323
8.4% [8.1,8.9] 523
3.1% [2.7,3.5] 191
4.6% [4.2,5.1] 286
model 3
Root
ADO rate = 16.1%
FPR = 1.0%
IDH2_p.R140Q
6.8% [6.2,7.4] 179
NPM1_p.L287fs
7.2% [6.6,7.8] 190
NRAS_p.G12D
FLT3_c.C2028G_p.N676K
FLT3-ITD
27.4% [26.0,28.8] 725
NRAS_p.G13D
FLT3_c.C2028A_p.N676K
31.2% [29.8,32.5] 825
11.3% [10.4,12.2] 298
4.9% [4.2,5.7] 130
5.6% [4.5,6.5] 147
5.6% [4.6,6.5] 147
model 4
Root
ADO rate = 8.8%
FPR = 1.0%
IDH2_p.R140Q
7.6% [7.1,8.2] 201
NPM1_p.L287fs
7.7% [7.3,8.3] 205
NRAS_p.G12D
NRAS_p.G13D
FLT3_c.C2028G_p.N676K
FLT3_c.C2028A_p.N676K
FLT3-ITD
28.8% [27.7,30.1] 763
37.1% [36.4,37.9] 982
2.9% [2.4,3.6] 78
9.3% [8.7,9.9] 245
3.2% [2.6,3.8] 84
3.1% [2.6,3.7] 83

### AML-41-001

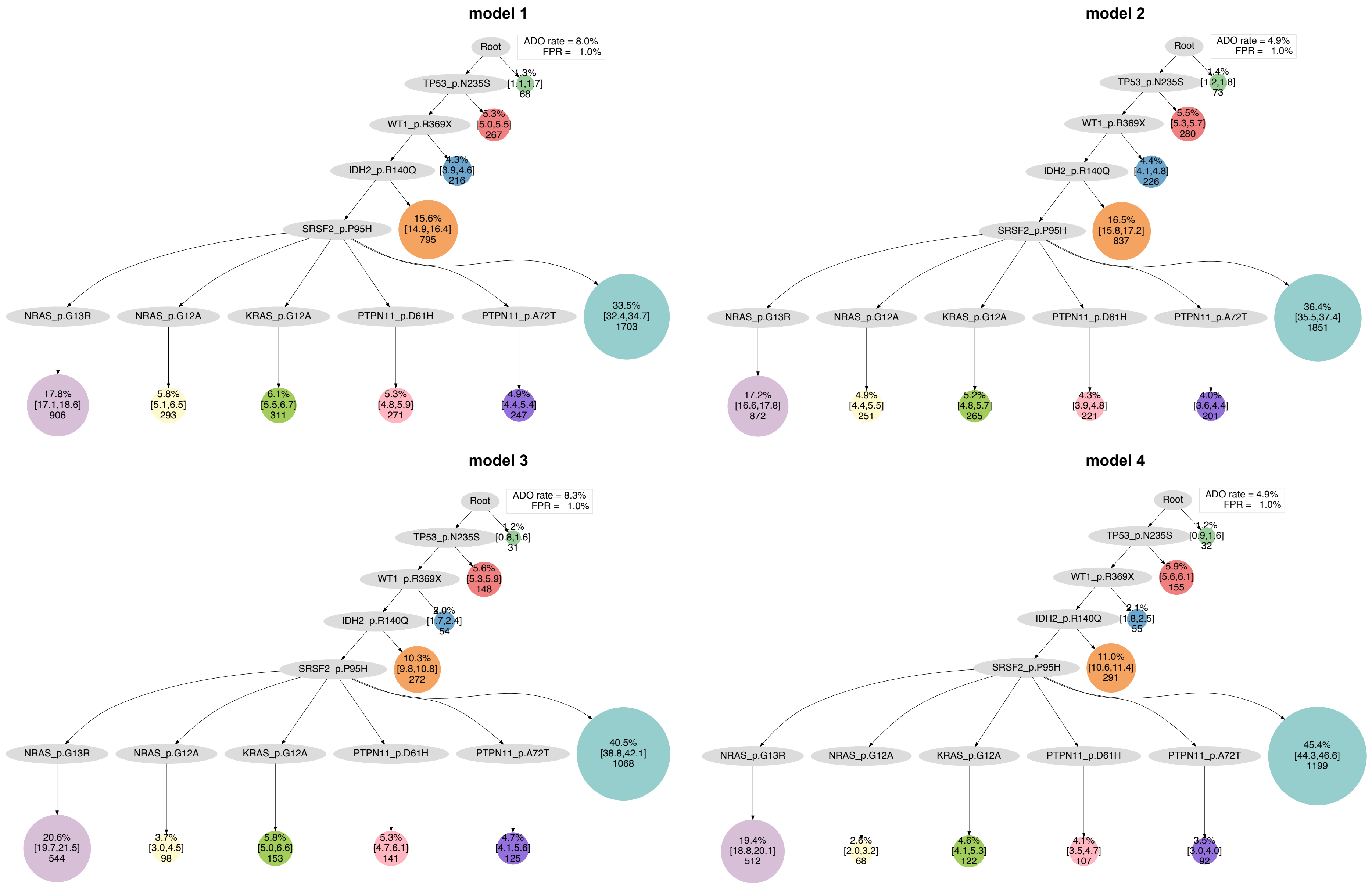

model 1
Root
ADO rate = 8.0%
FPR = 1.0%
TP53_p.N235S
1.3% [1.1,1.7] 68
WT1_p.R369X
5.3% [5.0,5.5] 267
IDH2_p.R140Q
4.3% [3.9,4.6] 216
SRSF2_p.P95H
15.6% [14.9,16.4] 795
NRAS_p.G13R
NRAS_p.G12A
KRAS_p.G12A
PTPN11_p.D61H
PTPN11_p.A72T
33.5% [32.4,34.7] 1703
17.8% [17.1,18.6] 906
5.8% [5.1,6.5] 293
6.1% [5.5,6.7] 311
5.3% [4.8,5.9] 271
4.9% [4.4,5.4] 247
model 2
Root
ADO rate = 4.9%
FPR = 1.0%
TP53_p.N235S
1.4% [1.2,1.8] 73
WT1_p.R369X
5.5% [5.3,5.7] 280
IDH2_p.R140Q
4.4% [4.1,4.8] 226
SRSF2_p.P95H
16.5% [15.8,17.2] 837
NRAS_p.G13R
NRAS_p.G12A
KRAS_p.G12A
PTPN11_p.D61H
PTPN11_p.A72T
36.4% [35.5,37.4] 1851
17.2% [16.6,17.8] 872
4.9% [4.4,5.5] 251
5.2% [4.8,5.7] 265
4.3% [3.9,4.8] 221
4.0% [3.6,4.4] 201
model 3
Root
ADO rate = 8.3%
FPR = 1.0%
TP53_p.N235S
1.2% [0.8,1.6] 31
WT1_p.R369X
5.6% [5.3,5.9] 148
IDH2_p.R140Q
2.0% [1.7,2.4] 54
SRSF2_p.P95H
10.3% [9.8,10.8] 272
NRAS_p.G13R
NRAS_p.G12A
KRAS_p.G12A
PTPN11_p.D61H
PTPN11_p.A72T
40.5% [38.8,42.1] 1068
20.6% [19.7,21.5] 544
3.7% [3.0,4.5] 98
5.8% [5.0,6.6] 153
5.3% [4.7,6.1] 141
4.7% [4.1,5.6] 125
model 4
Root
ADO rate = 4.9%
FPR = 1.0%
TP53_p.N235S
1.2% [0.9,1.6] 32
WT1_p.R369X
5.9% [5.6,6.1] 155
IDH2_p.R140Q
2.1% [1.8,2.5] 55
SRSF2_p.P95H
11.0% [10.6,11.4] 291
NRAS_p.G13R
NRAS_p.G12A
KRAS_p.G12A
PTPN11_p.D61H
PTPN11_p.A72T
45.4% [44.3,46.6] 1199
19.4% [18.8,20.1] 512
2.6% [2.0,3.2] 68
4.6% [4.1,5.3] 122
4.1% [3.5,4.7] 107
3.5% [3.0,4.0] 92

### AML-42-001

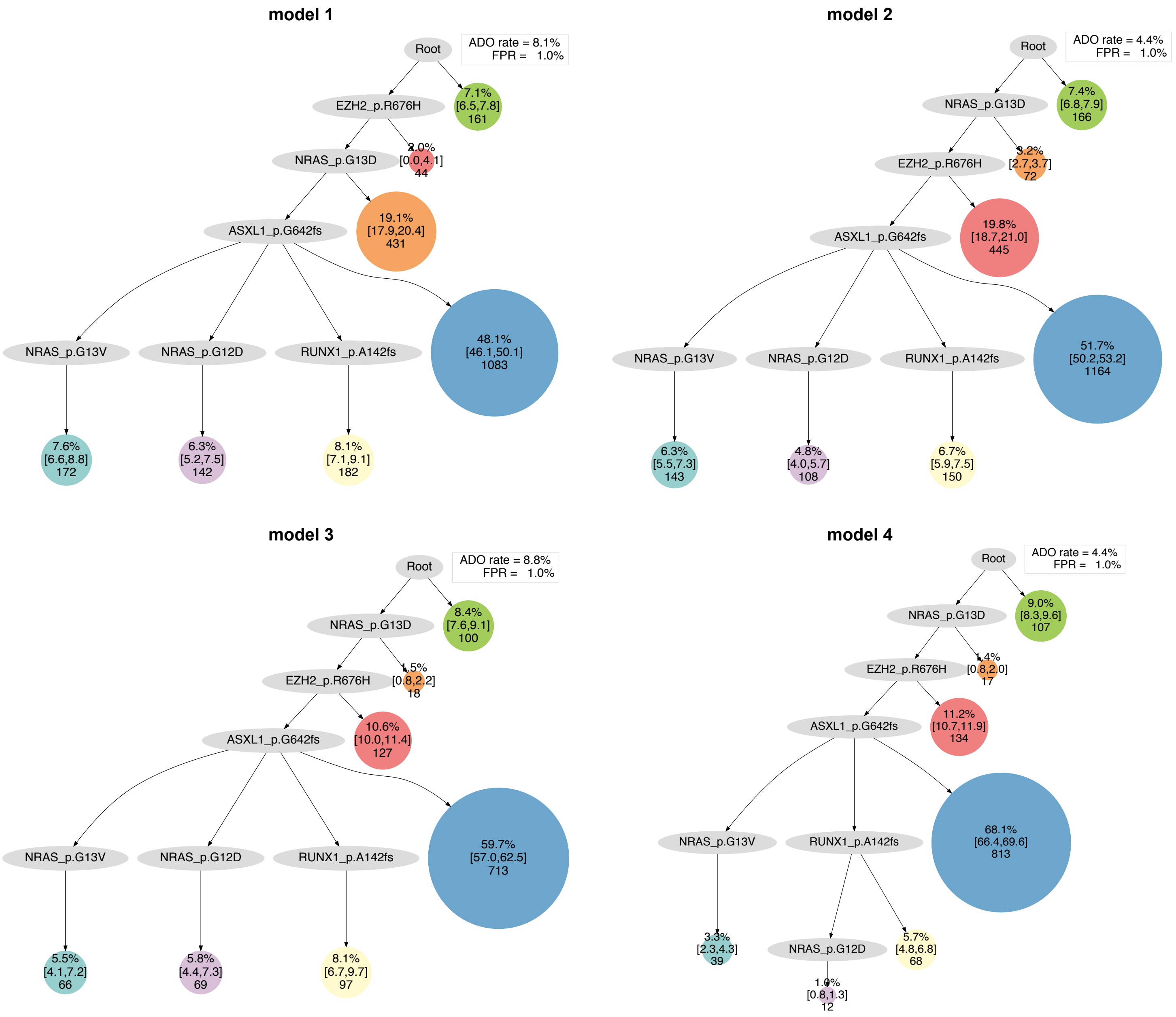

model 1
Root
ADO rate = 8.1%
FPR = 1.0%
EZH2_p.R676H
7.1%
[6.5,7.8]
161
NRAS_p.G13D
2.0%
[0.0,4.1]
44
ASXL1_p.G642fs
19.1%
[17.9,20.4]
431
NRAS_p.G13V
NRAS_p.G12D
RUNX1_p.A142fs
48.1%
[46.1,50.1]
1083
7.6%
[6.6,8.8]
172
6.3%
[5.2,7.5]
142
8.1%
[7.1,9.1]
182
model 2
Root
ADO rate = 4.4%
FPR = 1.0%
NRAS_p.G13D
7.4%
[6.8,7.9]
166
EZH2_p.R676H
3.2%
[2.7,3.7]
72
ASXL1_p.G642fs
19.8%
[18.7,21.0]
445
NRAS_p.G13V
NRAS_p.G12D
RUNX1_p.A142fs
51.7%
[50.2,53.2]
1164
6.3%
[5.5,7.3]
143
4.8%
[4.0,5.7]
108
6.7%
[5.9,7.5]
150
model 3
Root
ADO rate = 8.8%
FPR = 1.0%
NRAS_p.G13D
8.4%
[7.6,9.1]
100
EZH2_p.R676H
1.5%
[0.8,2.2]
18
ASXL1_p.G642fs
10.6%
[10.0,11.4]
127
NRAS_p.G13V
NRAS_p.G12D
RUNX1_p.A142fs
59.7%
[57.0,62.5]
713
5.5%
[4.1,7.2]
66
5.8%
[4.4,7.3]
69
8.1%
[6.7,9.7]
97
model 4
Root
ADO rate = 4.4%
FPR = 1.0%
NRAS_p.G13D
9.0%
[8.3,9.6]
107
EZH2_p.R676H
1.4%
[0.8,2.0]
17
ASXL1_p.G642fs
11.2%
[10.7,11.9]
134
NRAS_p.G13V
RUNX1_p.A142fs
68.1%
[66.4,69.6]
813
3.3%
[2.3,4.3]
39
NRAS_p.G12D
5.7%
[4.8,6.8]
68
1.0%
[0.8,1.3]
12

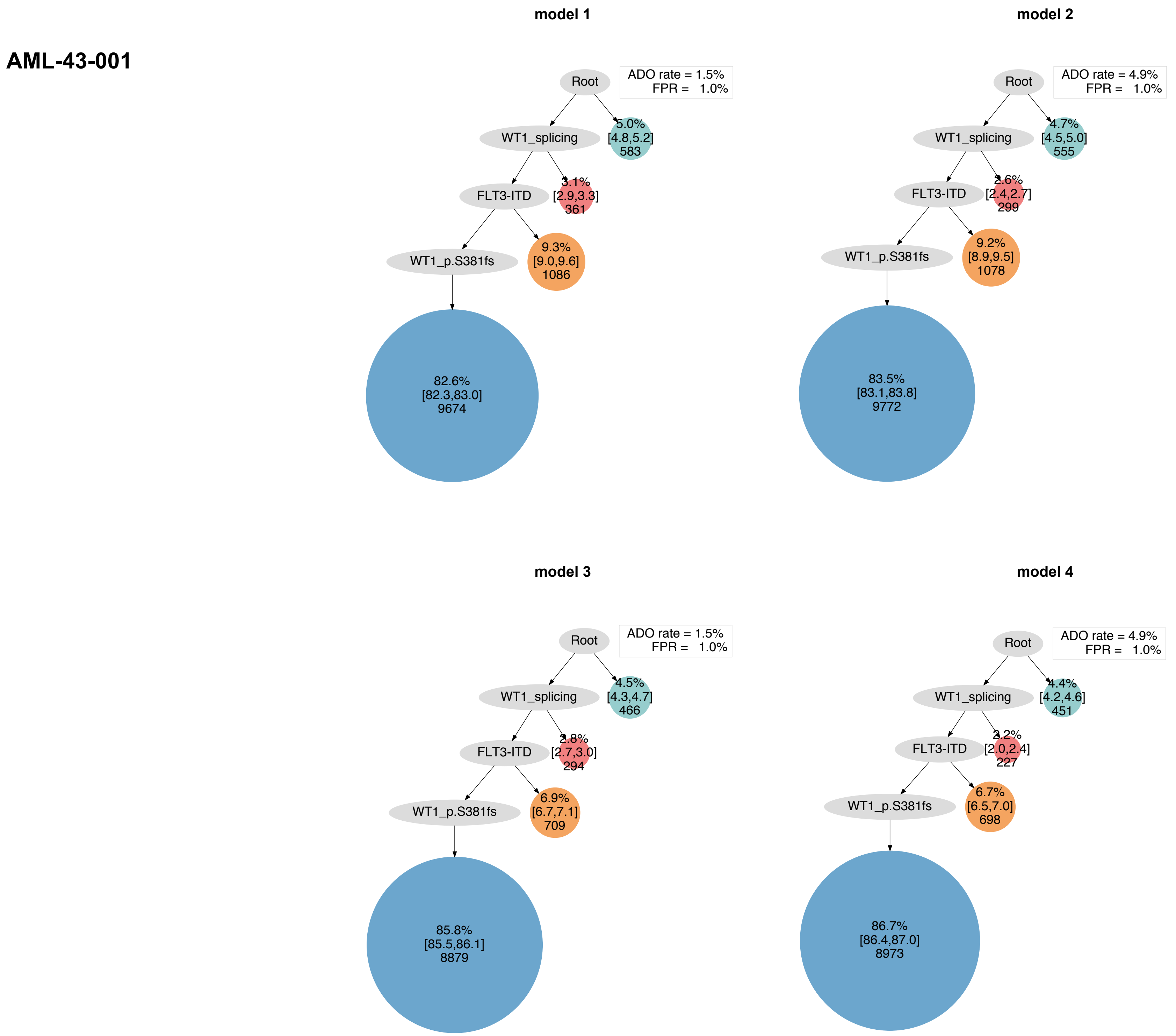
AML-43-001
model 1
Root
ADO rate = 1.5%
FPR = 1.0%
WT1_splicing
5.0%
[4.8,5.2]
583
FLT3-ITD
3.1%
[2.9,3.3]
361
WT1_p.S381fs
9.3%
[9.0,9.6]
1086
82.6%
[82.3,83.0]
9674
model 2
Root
ADO rate = 4.9%
FPR = 1.0%
WT1_splicing
4.7%
[4.5,5.0]
555
FLT3-ITD
2.6%
[2.4,2.7]
299
WT1_p.S381fs
9.2%
[8.9,9.5]
1078
83.5%
[83.1,83.8]
9772
model 3
Root
ADO rate = 1.5%
FPR = 1.0%
WT1_splicing
4.5%
[4.3,4.7]
466
FLT3-ITD
2.8%
[2.7,3.0]
294
WT1_p.S381fs
6.9%
[6.7,7.1]
709
85.8%
[85.5,86.1]
8879
model 4
Root
ADO rate = 4.9%
FPR = 1.0%
WT1_splicing
4.4%
[4.2,4.6]
451
FLT3-ITD
2.2%
[2.0,2.4]
227
WT1_p.S381fs
6.7%
[6.5,7.0]
698
86.7%
[86.4,87.0]
8973

### AML-44-001

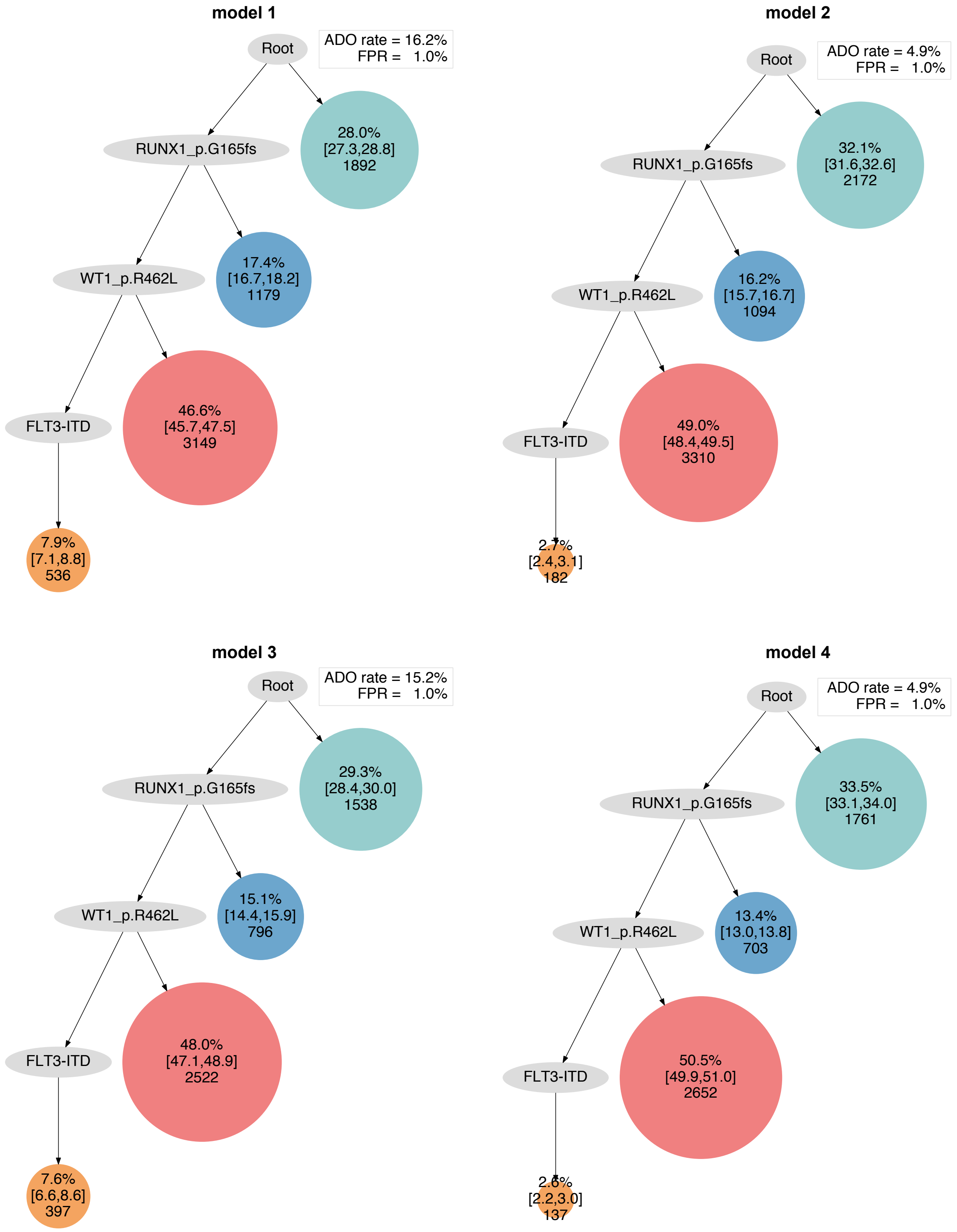

model 1
Root
ADO rate = 16.2%
FPR = 1.0%
28.0% [27.3,28.8] 1892
RUNX1_p.G165fs
17.4% [16.7,18.2] 1179
WT1_p.R462L
46.6% [45.7,47.5] 3149
FLT3-ITD
7.9% [7.1,8.8] 536
model 2
Root
ADO rate = 4.9%
FPR = 1.0%
32.1% [31.6,32.6] 2172
RUNX1_p.G165fs
16.2% [15.7,16.7] 1094
WT1_p.R462L
49.0% [48.4,49.5] 3310
FLT3-ITD
2.7% [2.4,3.1] 182
model 3
Root
ADO rate = 15.2%
FPR = 1.0%
29.3% [28.4,30.0] 1538
RUNX1_p.G165fs
15.1% [14.4,15.9] 796
WT1_p.R462L
48.0% [47.1,48.9] 2522
FLT3-ITD
7.6% [6.6,8.6] 397
model 4
Root
ADO rate = 4.9%
FPR = 1.0%
33.5% [33.1,34.0] 1761
RUNX1_p.G165fs
13.4% [13.0,13.8] 703
WT1_p.R462L
50.5% [49.9,51.0] 2652
FLT3-ITD
2.6% [2.2,3.0] 137

### AML-45-001

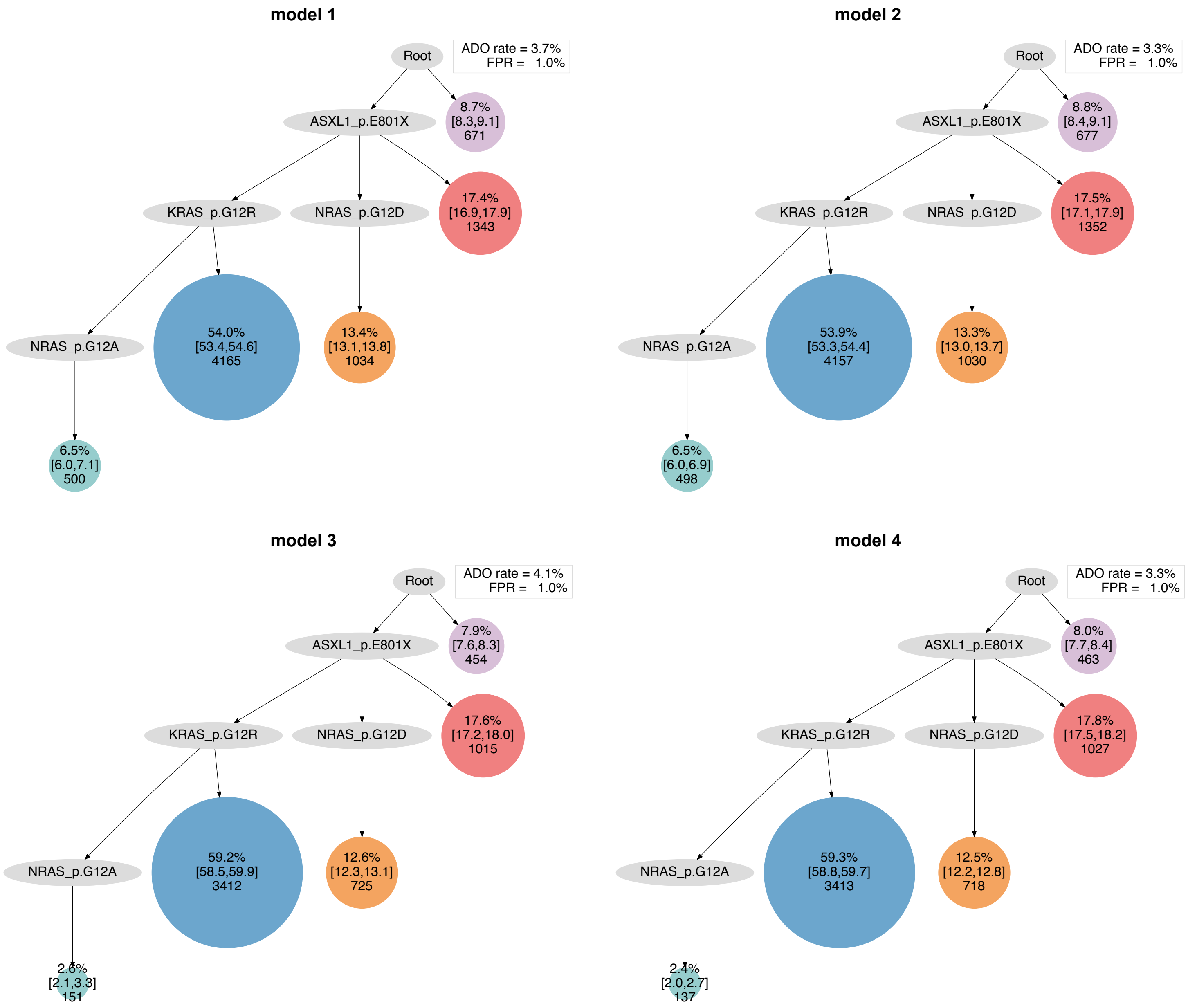
model 1
Root
ADO rate = 3.7%
FPR = 1.0%
ASXL1_p.E801X
8.7%
[8.3,9.1]
671
KRAS_p.G12R
NRAS_p.G12D
17.4%
[16.9,17.9]
1343
NRAS_p.G12A
54.0%
[53.4,54.6]
4165
13.4%
[13.1,13.8]
1034
6.5%
[6.0,7.1]
500
model 2
Root
ADO rate = 3.3%
FPR = 1.0%
ASXL1_p.E801X
8.8%
[8.4,9.1]
677
KRAS_p.G12R
NRAS_p.G12D
17.5%
[17.1,17.9]
1352
NRAS_p.G12A
53.9%
[53.3,54.4]
4157
13.3%
[13.0,13.7]
1030
6.5%
[6.0,6.9]
498
model 3
Root
ADO rate = 4.1%
FPR = 1.0%
ASXL1_p.E801X
7.9%
[7.6,8.3]
454
KRAS_p.G12R
NRAS_p.G12D
17.6%
[17.2,18.0]
1015
NRAS_p.G12A
59.2%
[58.5,59.9]
3412
12.6%
[12.3,13.1]
725
2.6%
[2.1,3.3]
151
model 4
Root
ADO rate = 3.3%
FPR = 1.0%
ASXL1_p.E801X
8.0%
[7.7,8.4]
463
KRAS_p.G12R
NRAS_p.G12D
17.8%
[17.5,18.2]
1027
NRAS_p.G12A
59.3%
[58.8,59.7]
3413
12.5%
[12.2,12.8]
718
2.4%
[2.0,2.7]
137

**AML-46-001**

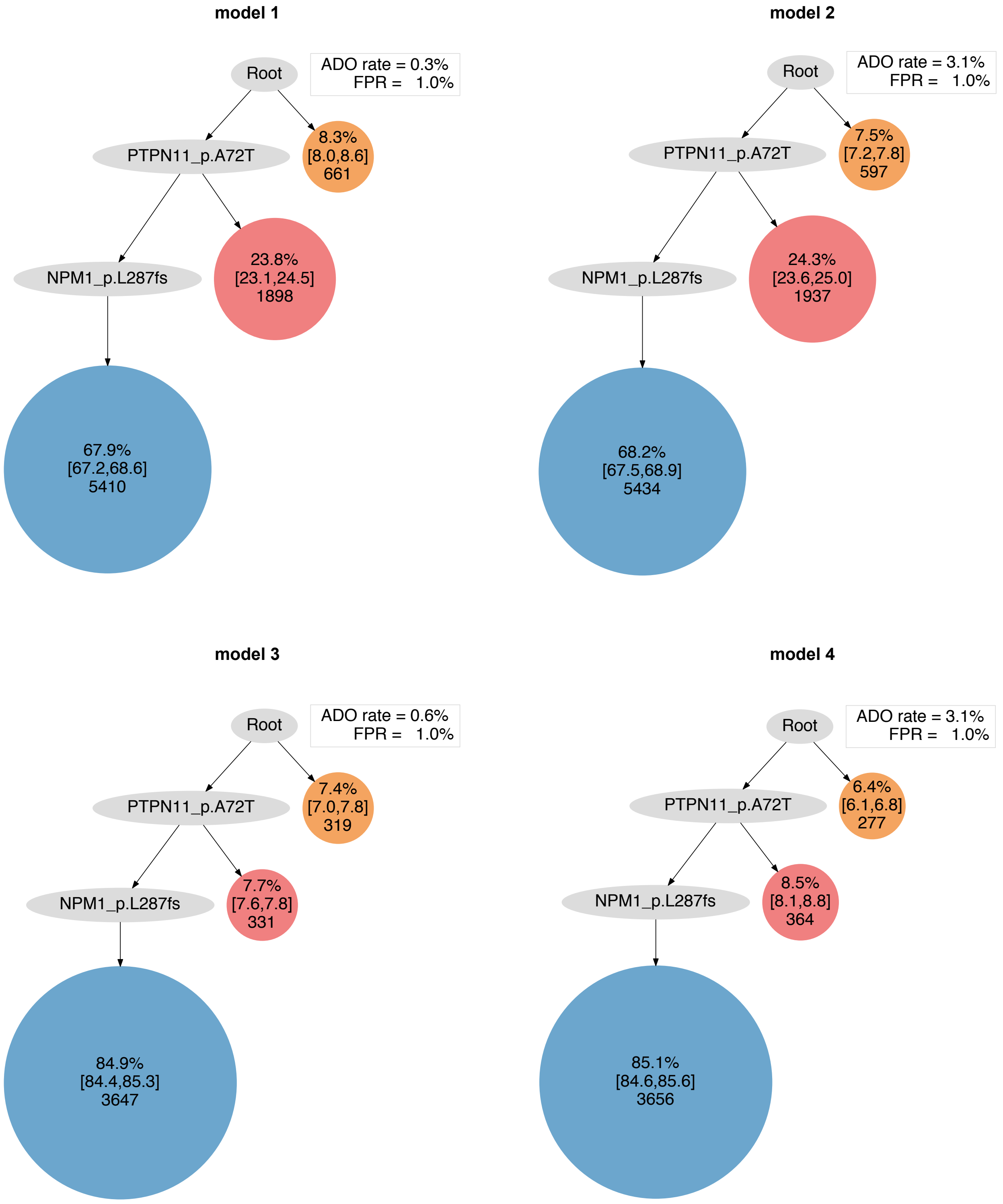
model 1
Root
ADO rate = 0.3%
FPR = 1.0%
PTPN11_p.A72T
8.3%
[8.0,8.6]
661
NPM1_p.L287fs
23.8%
[23.1,24.5]
1898
67.9%
[67.2,68.6]
5410
model 2
Root
ADO rate = 3.1%
FPR = 1.0%
PTPN11_p.A72T
7.5%
[7.2,7.8]
597
NPM1_p.L287fs
24.3%
[23.6,25.0]
1937
68.2%
[67.5,68.9]
5434
model 3
Root
ADO rate = 0.6%
FPR = 1.0%
PTPN11_p.A72T
7.4%
[7.0,7.8]
319
NPM1_p.L287fs
7.7%
[7.6,7.8]
331
84.9%
[84.4,85.3]
3647
model 4
Root
ADO rate = 3.1%
FPR = 1.0%
PTPN11_p.A72T
6.4%
[6.1,6.8]
277
NPM1_p.L287fs
8.5%
[8.1,8.8]
364
85.1%
[84.6,85.6]
3656

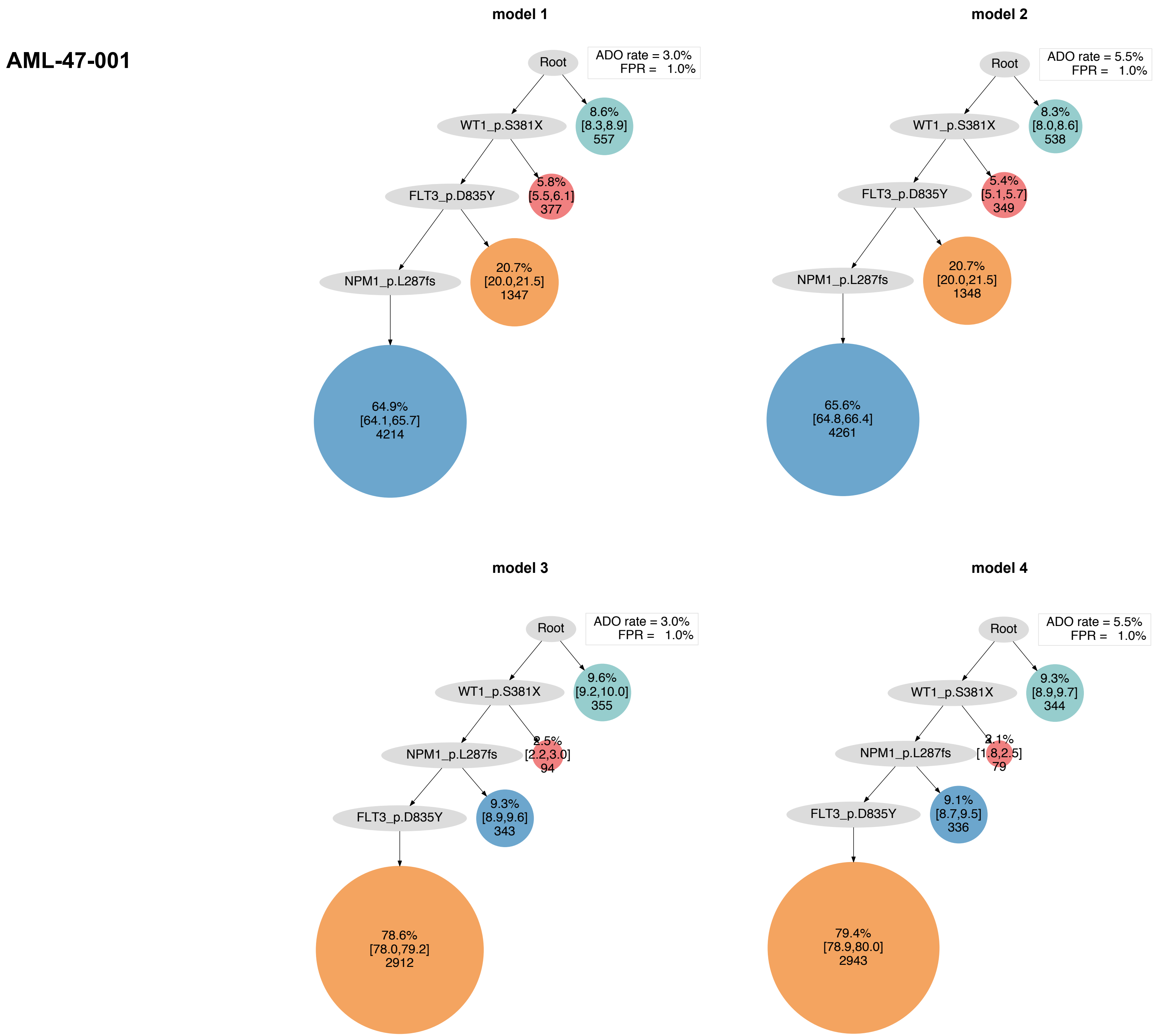

AML-47-001
model 1
Root
ADO rate = 3.0%
FPR = 1.0%
WT1_p.S381X
8.6%
[8.3,8.9]
557
FLT3_p.D835Y
5.8%
[5.5,6.1]
377
NPM1_p.L287fs
20.7%
[20.0,21.5]
1347
64.9%
[64.1,65.7]
4214
model 2
Root
ADO rate = 5.5%
FPR = 1.0%
WT1_p.S381X
8.3%
[8.0,8.6]
538
FLT3_p.D835Y
5.4%
[5.1,5.7]
349
NPM1_p.L287fs
20.7%
[20.0,21.5]
1348
65.6%
[64.8,66.4]
4261
model 3
Root
ADO rate = 3.0%
FPR = 1.0%
WT1_p.S381X
9.6%
[9.2,10.0]
355
NPM1_p.L287fs
2.5%
[2.2,3.0]
94
FLT3_p.D835Y
9.3%
[8.9,9.6]
343
78.6%
[78.0,79.2]
2912
model 4
Root
ADO rate = 5.5%
FPR = 1.0%
WT1_p.S381X
9.3%
[8.9,9.7]
344
NPM1_p.L287fs
2.1%
[1.8,2.5]
79
FLT3_p.D835Y
9.1%
[8.7,9.5]
336
79.4%
[78.9,80.0]
2943

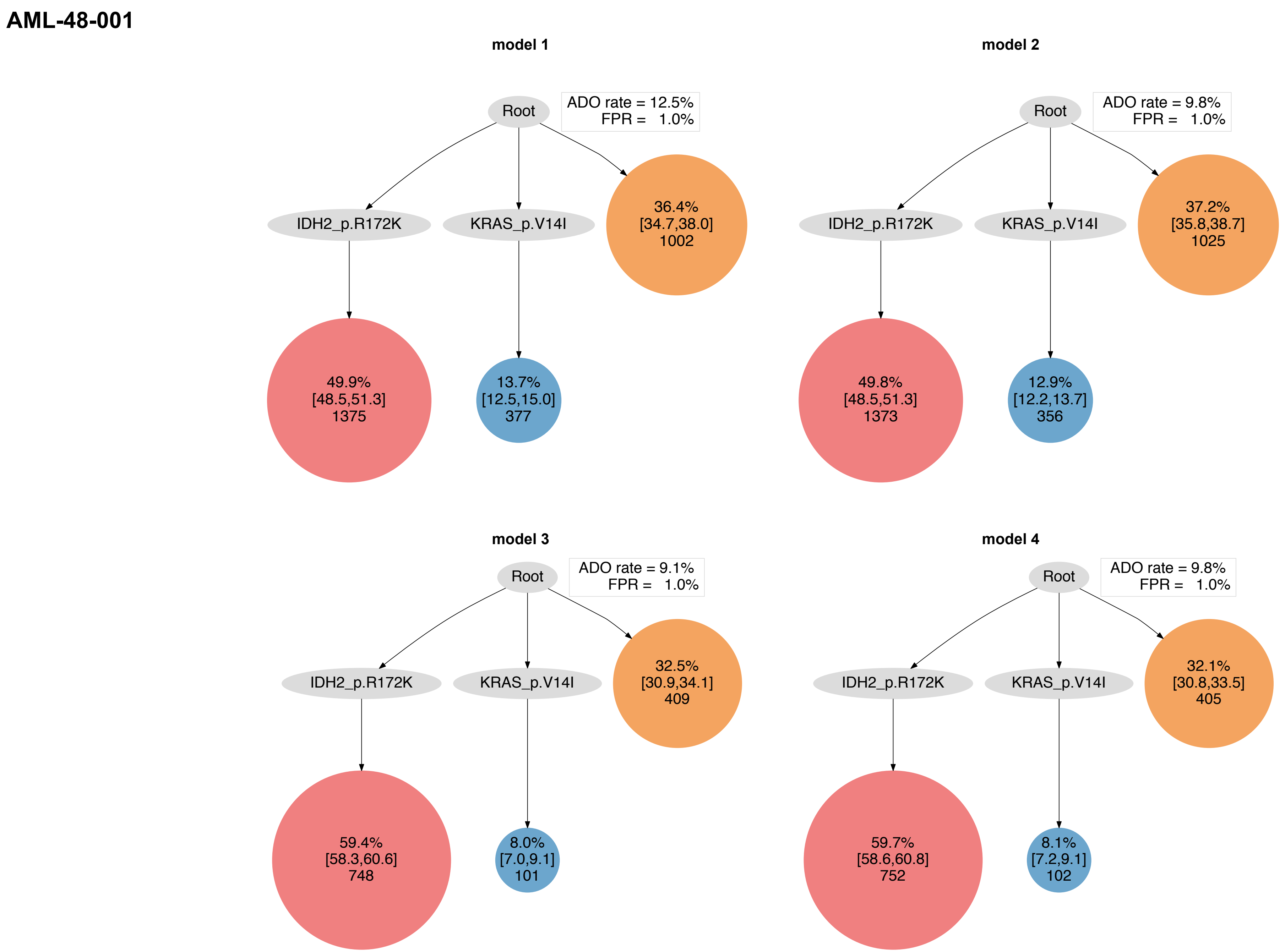
AML-48-001
model 1
Root
ADO rate = 12.5%
FPR = 1.0%
IDH2_p.R172K
KRAS_p.V14I
36.4%
[34.7,38.0]
1002
49.9%
[48.5,51.3]
1375
13.7%
[12.5,15.0]
377
model 2
Root
ADO rate = 9.8%
FPR = 1.0%
IDH2_p.R172K
KRAS_p.V14I
37.2%
[35.8,38.7]
1025
49.8%
[48.5,51.3]
1373
12.9%
[12.2,13.7]
356
model 3
Root
ADO rate = 9.1%
FPR = 1.0%
IDH2_p.R172K
KRAS_p.V14I
32.5%
[30.9,34.1]
409
59.4%
[58.3,60.6]
748
8.0%
[7.0,9.1]
101
model 4
Root
ADO rate = 9.8%
FPR = 1.0%
IDH2_p.R172K
KRAS_p.V14I
32.1%
[30.8,33.5]
405
59.7%
[58.6,60.8]
752
8.1%
[7.2,9.1]
102

### AML-49-001

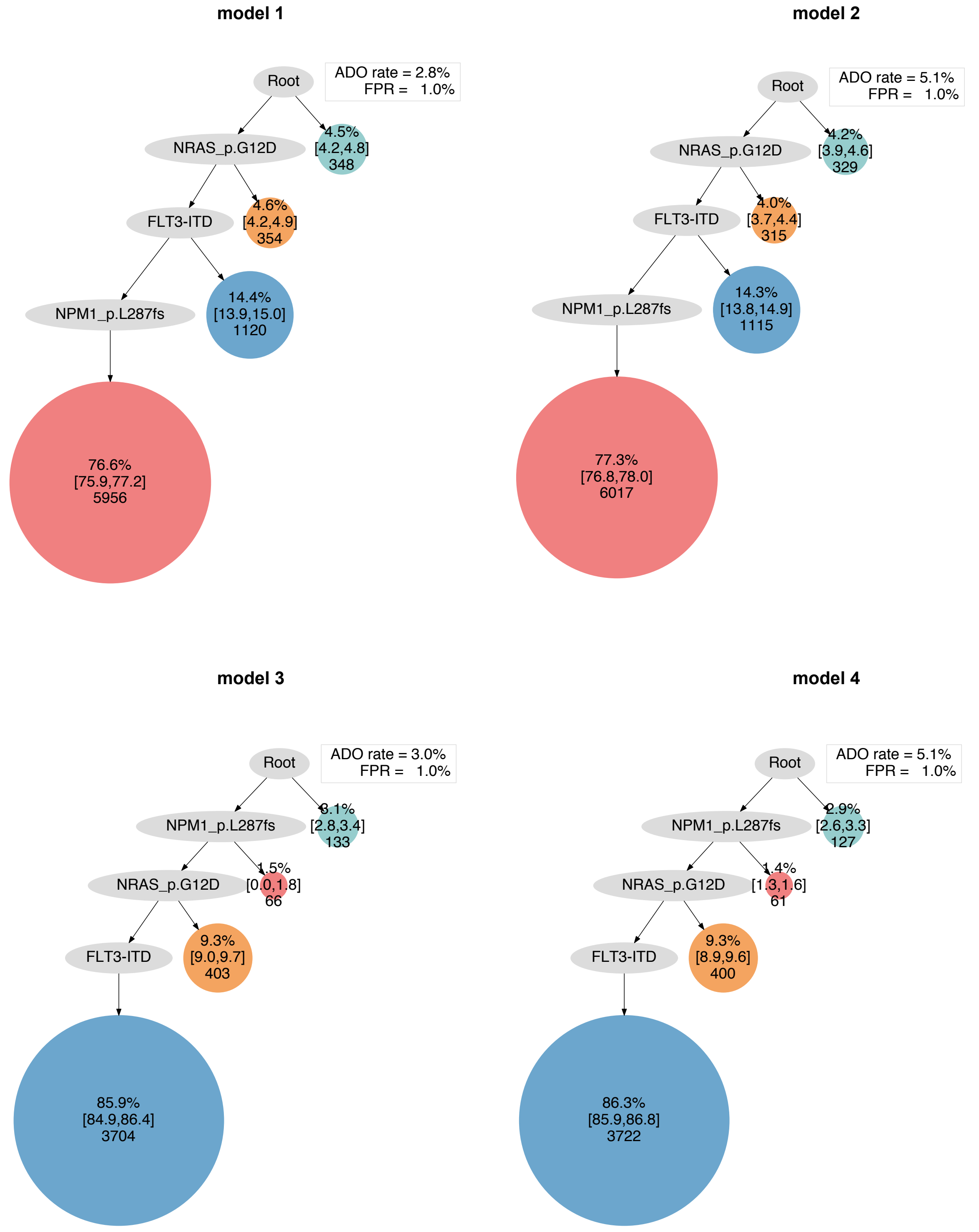

model 1
Root
ADO rate = 2.8%
FPR = 1.0%
NRAS_p.G12D
4.5%
[4.2,4.8]
348
FLT3-ITD
4.6%
[4.2,4.9]
354
NPM1_p.L287fs
14.4%
[13.9,15.0]
1120
76.6%
[75.9,77.2]
5956
model 2
Root
ADO rate = 5.1%
FPR = 1.0%
NRAS_p.G12D
4.2%
[3.9,4.6]
329
FLT3-ITD
4.0%
[3.7,4.4]
315
NPM1_p.L287fs
14.3%
[13.8,14.9]
1115
77.3%
[76.8,78.0]
6017
model 3
Root
ADO rate = 3.0%
FPR = 1.0%
NPM1_p.L287fs
3.1%
[2.8,3.4]
133
NRAS_p.G12D
1.5%
[0.0,1.8]
66
FLT3-ITD
9.3%
[9.0,9.7]
403
85.9%
[84.9,86.4]
3704
model 4
Root
ADO rate = 5.1%
FPR = 1.0%
NPM1_p.L287fs
2.9%
[2.6,3.3]
127
NRAS_p.G12D
1.4%
[1.3,1.6]
61
FLT3-ITD
9.3%
[8.9,9.6]
400
86.3%
[85.9,86.8]
3722

**AML-50-001**

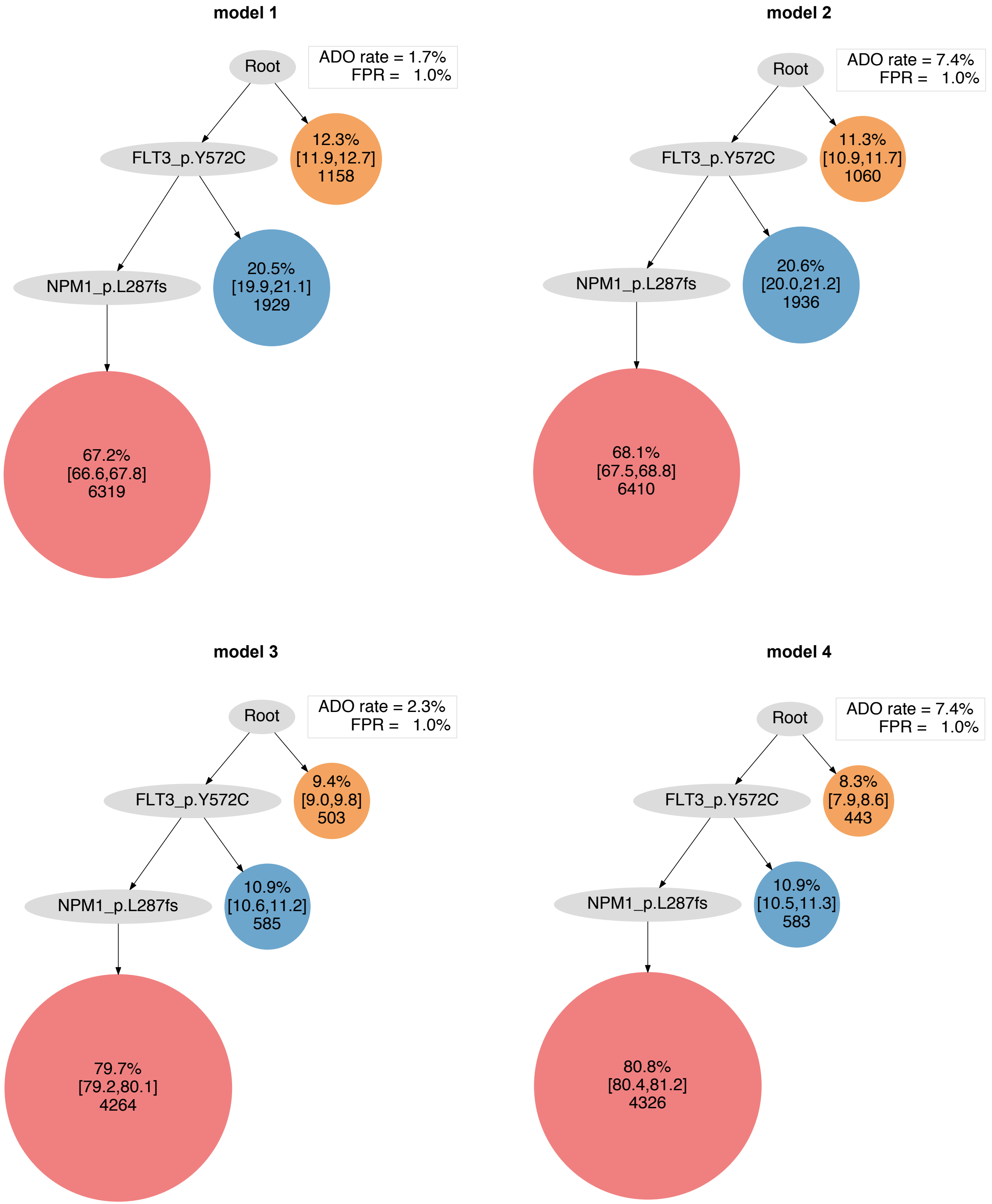
model 1
Root
ADO rate = 1.7%
FPR = 1.0%
FLT3_p.Y572C
12.3%
[11.9,12.7]
1158
NPM1_p.L287fs
20.5%
[19.9,21.1]
1929
67.2%
[66.6,67.8]
6319
model 2
Root
ADO rate = 7.4%
FPR = 1.0%
FLT3_p.Y572C
11.3%
[10.9,11.7]
1060
NPM1_p.L287fs
20.6%
[20.0,21.2]
1936
68.1%
[67.5,68.8]
6410
model 3
Root
ADO rate = 2.3%
FPR = 1.0%
FLT3_p.Y572C
9.4%
[9.0,9.8]
503
NPM1_p.L287fs
10.9%
[10.6,11.2]
585
79.7%
[79.2,80.1]
4264
model 4
Root
ADO rate = 7.4%
FPR = 1.0%
FLT3_p.Y572C
8.3%
[7.9,8.6]
443
NPM1_p.L287fs
10.9%
[10.5,11.3]
583
80.8%
[80.4,81.2]
4326

### AML-51-001

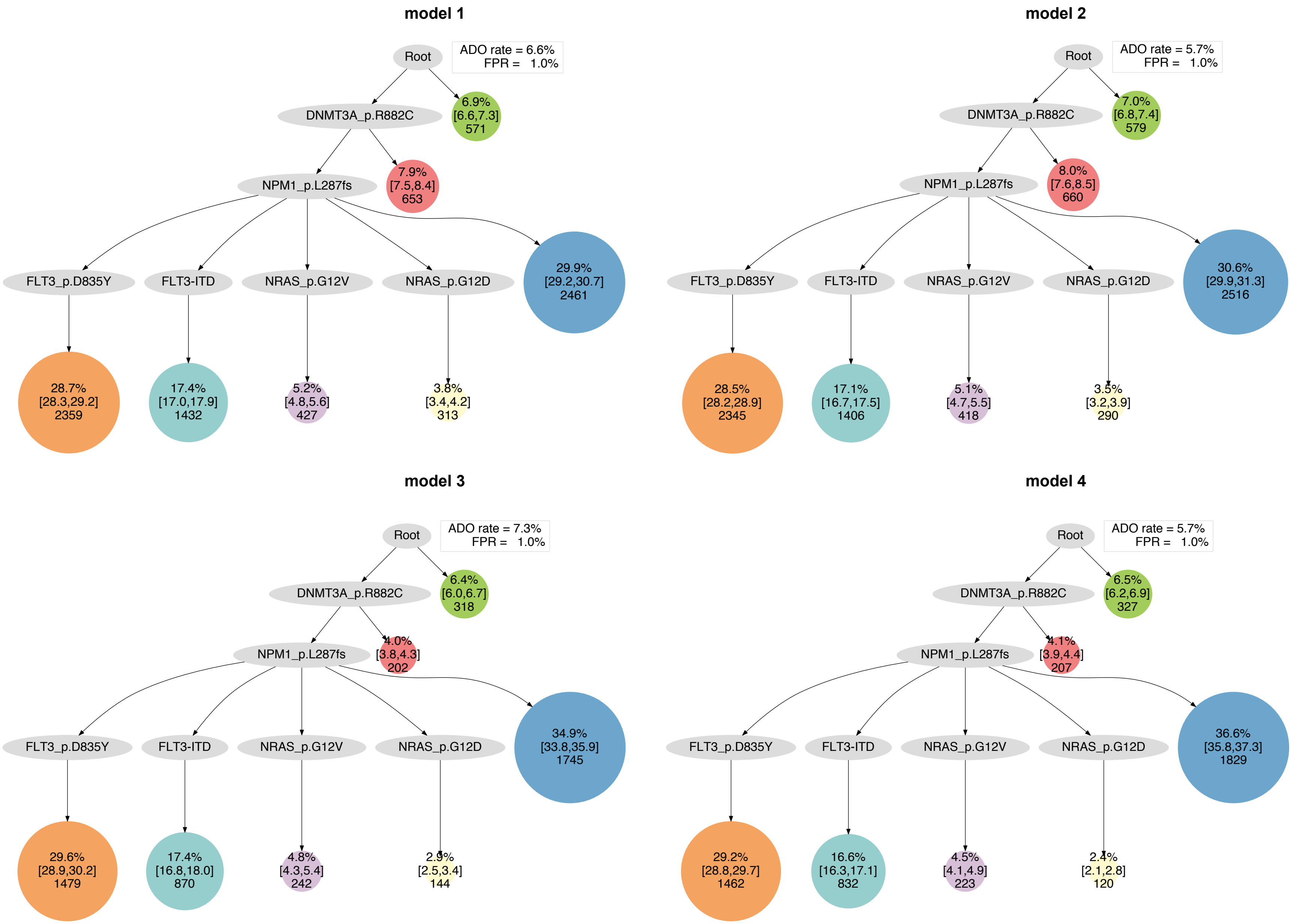

model 1
Root
ADO rate = 6.6%
FPR = 1.0%
DNMT3A_p.R882C
6.9%
[6.6,7.3]
571
NPM1_p.L287fs
7.9%
[7.5,8.4]
653
FLT3_p.D835Y
FLT3-ITD
NRAS_p.G12V
NRAS_p.G12D
29.9%
[29.2,30.7]
2461
28.7%
[28.3,29.2]
2359
17.4%
[17.0,17.9]
1432
5.2%
[4.8,5.6]
427
3.8%
[3.4,4.2]
313
model 2
Root
ADO rate = 5.7%
FPR = 1.0%
DNMT3A_p.R882C
7.0%
[6.8,7.4]
579
NPM1_p.L287fs
8.0%
[7.6,8.5]
660
FLT3_p.D835Y
FLT3-ITD
NRAS_p.G12V
NRAS_p.G12D
30.6%
[29.9,31.3]
2516
28.5%
[28.2,28.9]
2345
17.1%
[16.7,17.5]
1406
5.1%
[4.7,5.5]
418
3.5%
[3.2,3.9]
290
model 3
Root
ADO rate = 7.3%
FPR = 1.0%
DNMT3A_p.R882C
6.4%
[6.0,6.7]
318
NPM1_p.L287fs
4.0%
[3.8,4.3]
202
FLT3_p.D835Y
FLT3-ITD
NRAS_p.G12V
NRAS_p.G12D
34.9%
[33.8,35.9]
1745
29.6%
[28.9,30.2]
1479
17.4%
[16.8,18.0]
870
4.8%
[4.3,5.4]
242
2.9%
[2.5,3.4]
144
model 4
Root
ADO rate = 5.7%
FPR = 1.0%
DNMT3A_p.R882C
6.5%
[6.2,6.9]
327
NPM1_p.L287fs
4.1%
[3.9,4.4]
207
FLT3_p.D835Y
FLT3-ITD
NRAS_p.G12V
NRAS_p.G12D
36.6%
[35.8,37.3]
1829
29.2%
[28.8,29.7]
1462
16.6%
[16.3,17.1]
832
4.5%
[4.1,4.9]
223
2.4%
[2.1,2.8]
120

#### AML-52-001

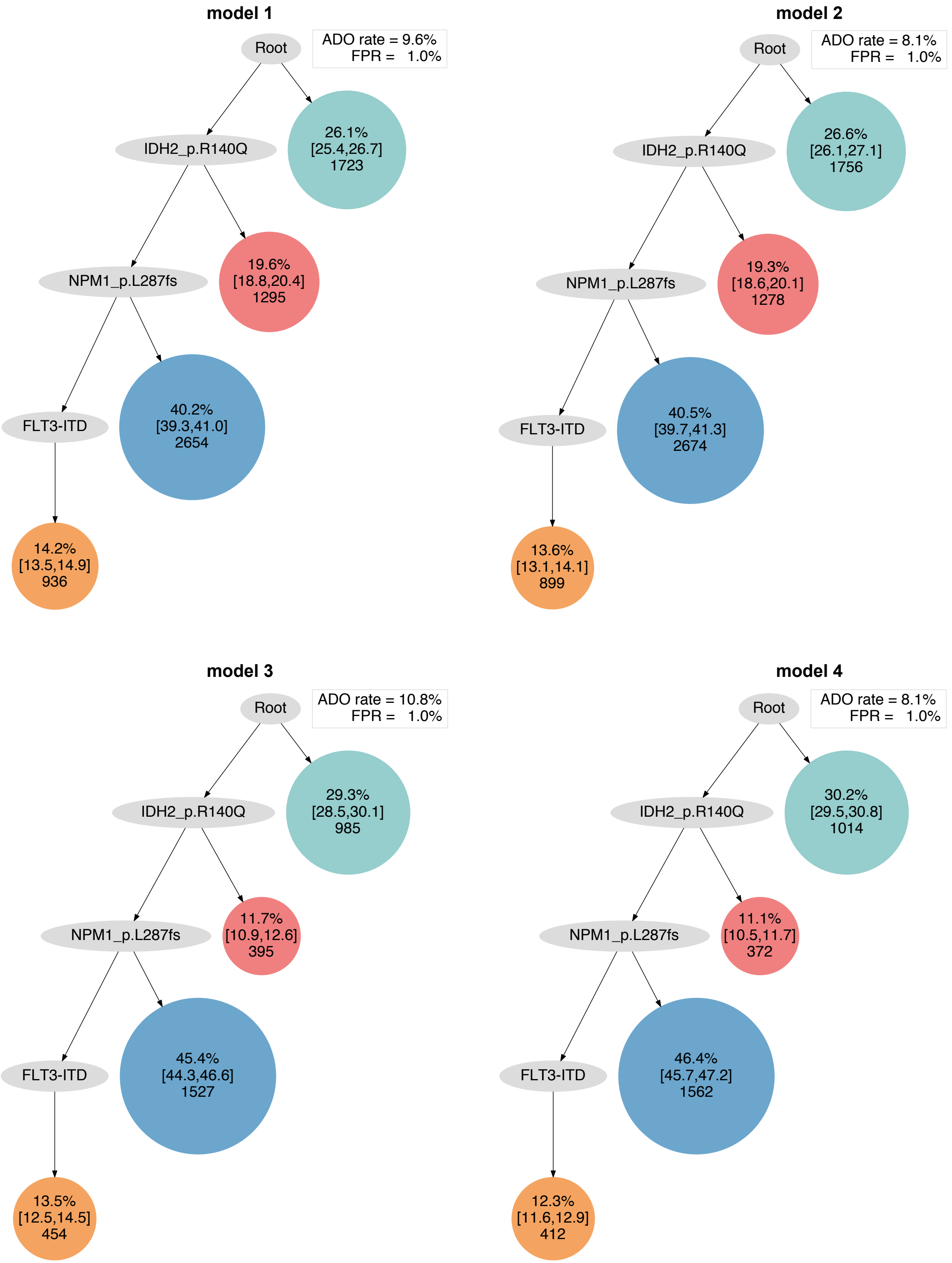
model 1
Root
ADO rate = 9.6%
FPR = 1.0%
IDH2_p.R140Q
26.1%
[25.4,26.7]
1723
NPM1_p.L287fs
19.6%
[18.8,20.4]
1295
FLT3-ITD
40.2%
[39.3,41.0]
2654
14.2%
[13.5,14.9]
936
model 2
Root
ADO rate = 8.1%
FPR = 1.0%
IDH2_p.R140Q
26.6%
[26.1,27.1]
1756
NPM1_p.L287fs
19.3%
[18.6,20.1]
1278
FLT3-ITD
40.5%
[39.7,41.3]
2674
13.6%
[13.1,14.1]
899
model 3
Root
ADO rate = 10.8%
FPR = 1.0%
IDH2_p.R140Q
29.3%
[28.5,30.1]
985
NPM1_p.L287fs
11.7%
[10.9,12.6]
395
FLT3-ITD
45.4%
[44.3,46.6]
1527
13.5%
[12.5,14.5]
454
model 4
Root
ADO rate = 8.1%
FPR = 1.0%
IDH2_p.R140Q
30.2%
[29.5,30.8]
1014
NPM1_p.L287fs
11.1%
[10.5,11.7]
372
FLT3-ITD
46.4%
[45.7,47.2]
1562
12.3%
[11.6,12.9]
412

#### AML-53-001

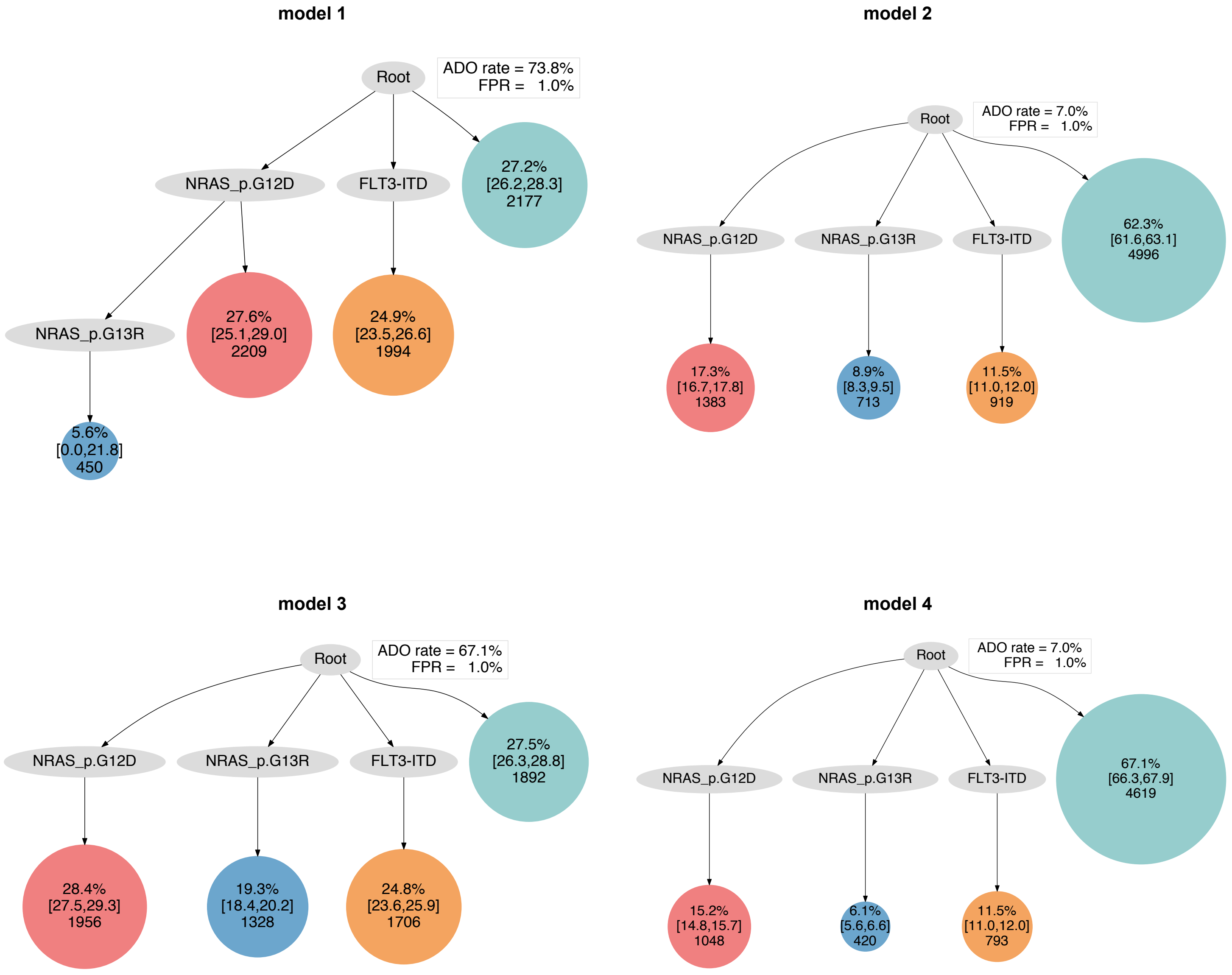

model 1
Root
ADO rate = 73.8%
FPR = 1.0%
NRAS_p.G12D
FLT3-ITD
27.2%
[26.2,28.3]
2177
NRAS_p.G13R
27.6%
[25.1,29.0]
2209
24.9%
[23.5,26.6]
1994
5.6%
[0.0,21.8]
450
model 2
Root
ADO rate = 7.0%
FPR = 1.0%
NRAS_p.G12D
NRAS_p.G13R
FLT3-ITD
62.3%
[61.6,63.1]
4996
17.3%
[16.7,17.8]
1383
8.9%
[8.3,9.5]
713
11.5%
[11.0,12.0]
919
model 3
Root
ADO rate = 67.1%
FPR = 1.0%
NRAS_p.G12D
NRAS_p.G13R
FLT3-ITD
27.5%
[26.3,28.8]
1892
28.4%
[27.5,29.3]
1956
19.3%
[18.4,20.2]
1328
24.8%
[23.6,25.9]
1706
model 4
Root
ADO rate = 7.0%
FPR = 1.0%
NRAS_p.G12D
NRAS_p.G13R
FLT3-ITD
67.1%
[66.3,67.9]
4619
15.2%
[14.8,15.7]
1048
6.1%
[5.6,6.6]
420
11.5%
[11.0,12.0]
793

### AML-54-001

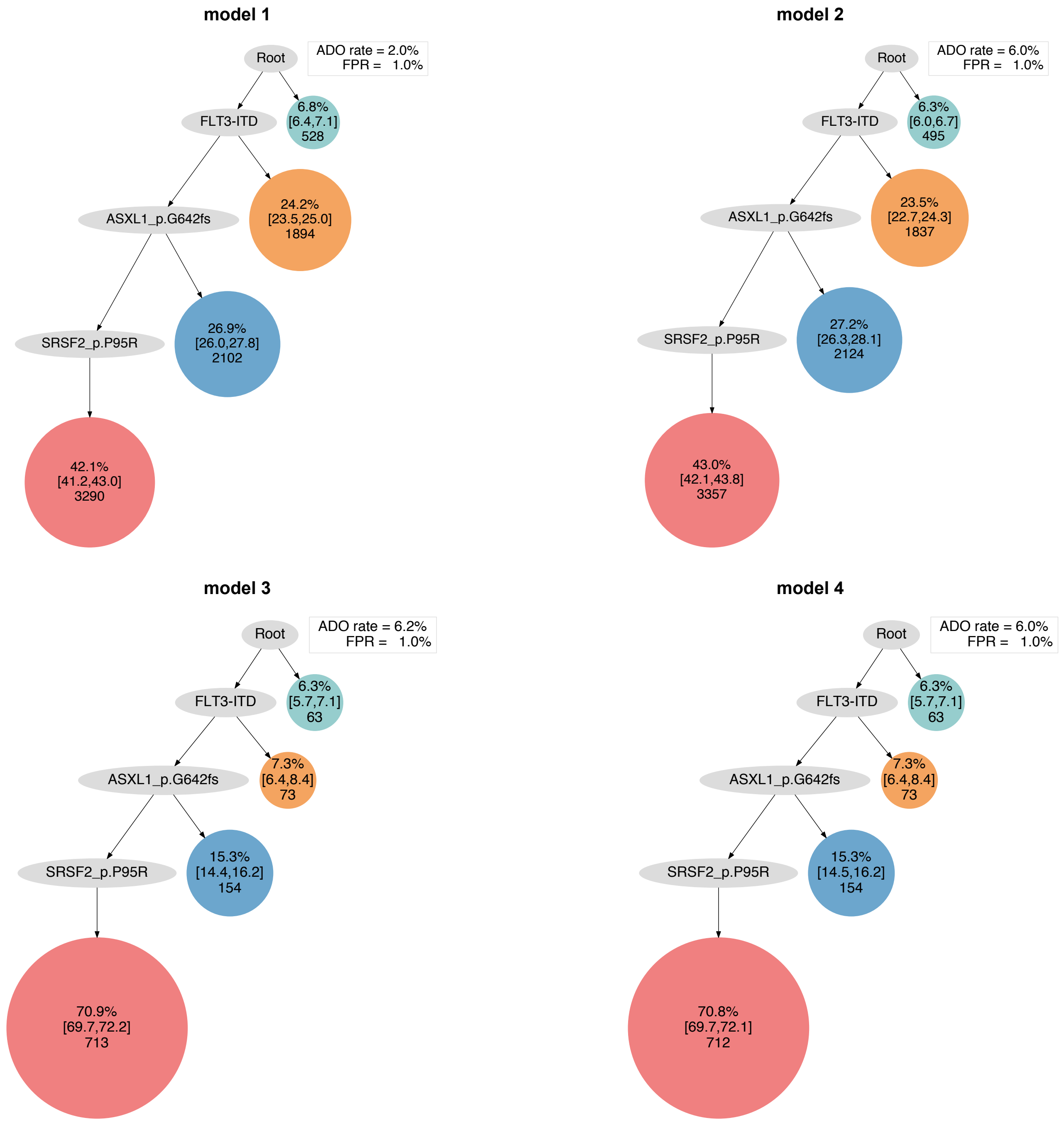
model 1
Root
ADO rate = 2.0%
FPR = 1.0%
FLT3-ITD
6.8%
[6.4,7.1]
528
ASXL1_p.G642fs
24.2%
[23.5,25.0]
1894
SRSF2_p.P95R
26.9%
[26.0,27.8]
2102
42.1%
[41.2,43.0]
3290
model 2
Root
ADO rate = 6.0%
FPR = 1.0%
FLT3-ITD
6.3%
[6.0,6.7]
495
ASXL1_p.G642fs
23.5%
[22.7,24.3]
1837
SRSF2_p.P95R
27.2%
[26.3,28.1]
2124
43.0%
[42.1,43.8]
3357
model 3
Root
ADO rate = 6.2%
FPR = 1.0%
FLT3-ITD
6.3%
[5.7,7.1]
63
ASXL1_p.G642fs
7.3%
[6.4,8.4]
73
SRSF2_p.P95R
15.3%
[14.4,16.2]
154
70.9%
[69.7,72.2]
713
model 4
Root
ADO rate = 6.0%
FPR = 1.0%
FLT3-ITD
6.3%
[5.7,7.1]
63
ASXL1_p.G642fs
7.3%
[6.4,8.4]
73
SRSF2_p.P95R
15.3%
[14.5,16.2]
154
70.8%
[69.7,72.1]
712

### AML-55-001

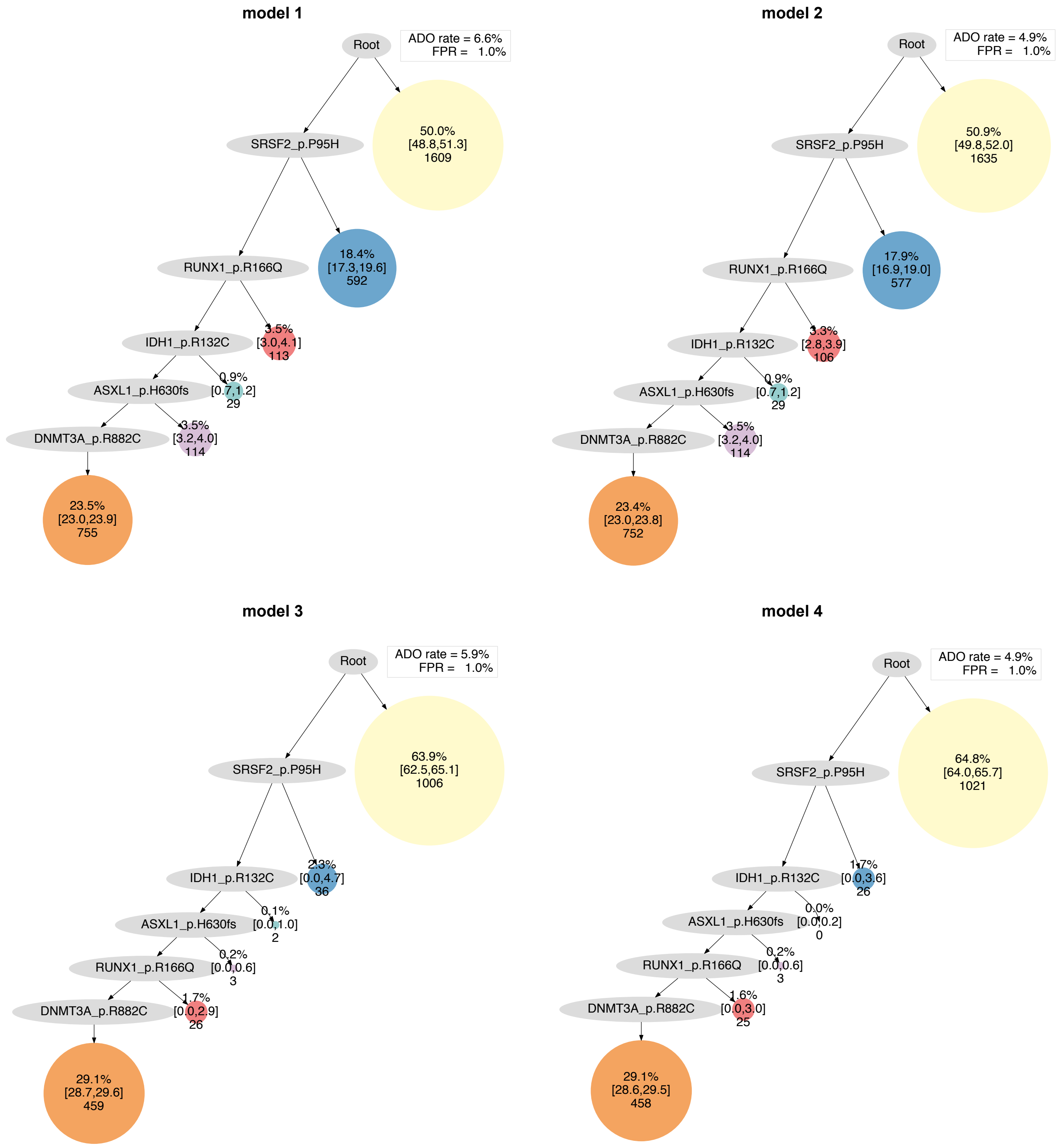
model 1
Root
ADO rate = 6.6%
FPR = 1.0%
SRSF2_p.P95H
50.0%
[48.8,51.3]
1609
RUNX1_p.R166Q
18.4%
[17.3,19.6]
592
IDH1_p.R132C
3.5%
[3.0,4.1]
113
ASXL1_p.H630fs
0.9%
[0.7,1.2]
29
DNMT3A_p.R882C
3.5%
[3.2,4.0]
114
23.5%
[23.0,23.9]
755
model 2
Root
ADO rate = 4.9%
FPR = 1.0%
SRSF2_p.P95H
50.9%
[49.8,52.0]
1635
RUNX1_p.R166Q
17.9%
[16.9,19.0]
577
IDH1_p.R132C
3.3%
[2.8,3.9]
106
ASXL1_p.H630fs
0.9%
[0.7,1.2]
29
DNMT3A_p.R882C
3.5%
[3.2,4.0]
114
23.4%
[23.0,23.8]
752
model 3
Root
ADO rate = 5.9%
FPR = 1.0%
SRSF2_p.P95H
63.9%
[62.5,65.1]
1006
IDH1_p.R132C
2.3%
[0.0,4.7]
36
ASXL1_p.H630fs
0.1%
[0.0,1.0]
2
RUNX1_p.R166Q
0.2%
[0.0,0.6]
3
DNMT3A_p.R882C
1.7%
[0.0,2.9]
26
29.1%
[28.7,29.6]
459
model 4
Root
ADO rate = 4.9%
FPR = 1.0%
SRSF2_p.P95H
64.8%
[64.0,65.7]
1021
IDH1_p.R132C
1.7%
[0.0,3.6]
26
ASXL1_p.H630fs
0.0%
[0.0,0.2]
0
RUNX1_p.R166Q
0.2%
[0.0,0.6]
3
DNMT3A_p.R882C
1.6%
[0.0,3.0]
25
29.1%
[28.6,29.5]
458

AML-57-001

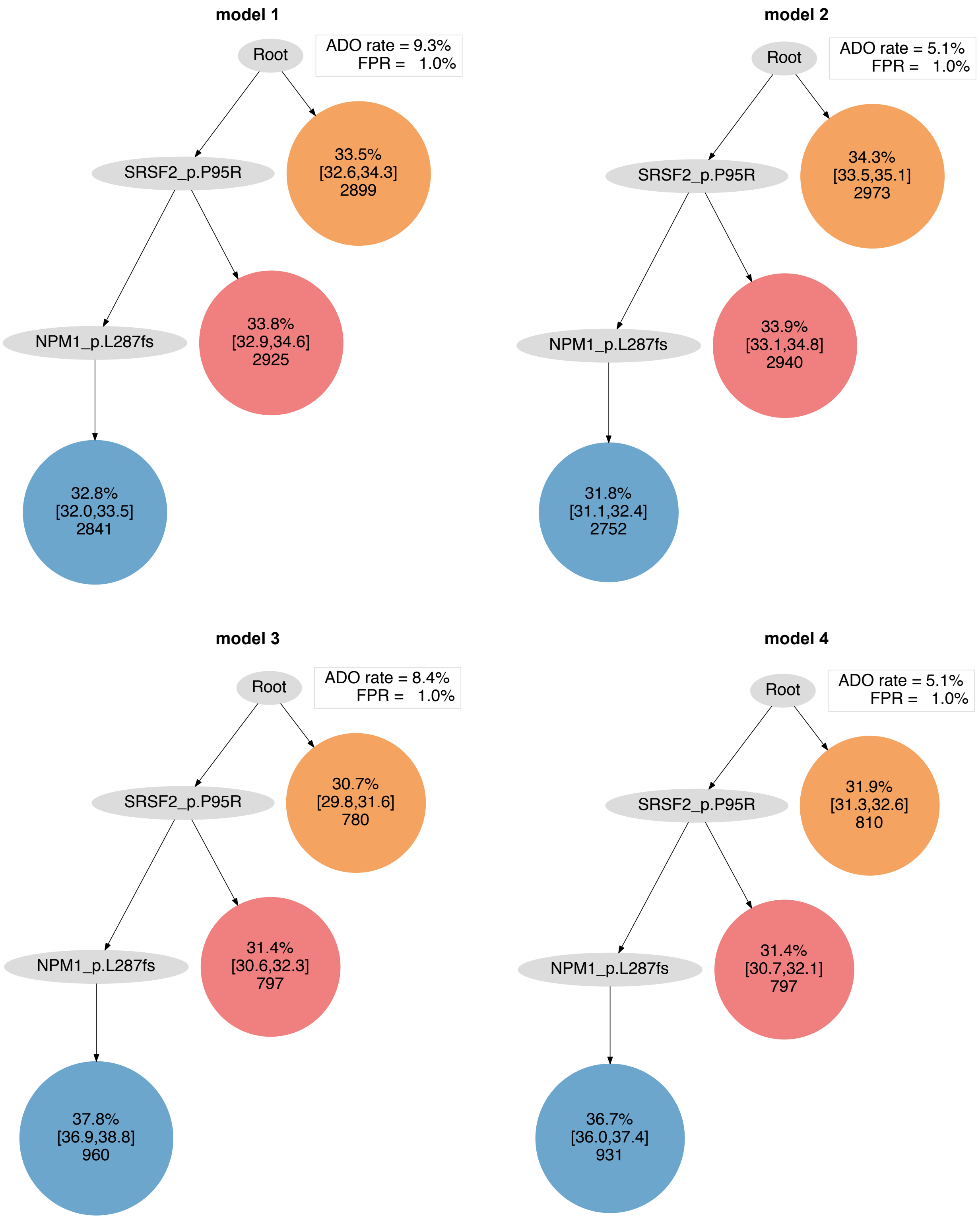
model 1
Root
ADO rate = 9.3%
FPR = 1.0%
SRSF2_p.P95R
33.5%
[32.6,34.3]
2899
NPM1_p.L287fs
33.8%
[32.9,34.6]
2925
32.8%
[32.0,33.5]
2841
model 2
Root
ADO rate = 5.1%
FPR = 1.0%
SRSF2_p.P95R
34.3%
[33.5,35.1]
2973
NPM1_p.L287fs
33.9%
[33.1,34.8]
2940
31.8%
[31.1,32.4]
2752
model 3
Root
ADO rate = 8.4%
FPR = 1.0%
SRSF2_p.P95R
30.7%
[29.8,31.6]
780
NPM1_p.L287fs
31.4%
[30.6,32.3]
797
37.8%
[36.9,38.8]
960
model 4
Root
ADO rate = 5.1%
FPR = 1.0%
SRSF2_p.P95R
31.9%
[31.3,32.6]
810
NPM1_p.L287fs
31.4%
[30.7,32.1]
797
36.7%
[36.0,37.4]
931

### AML-58-001

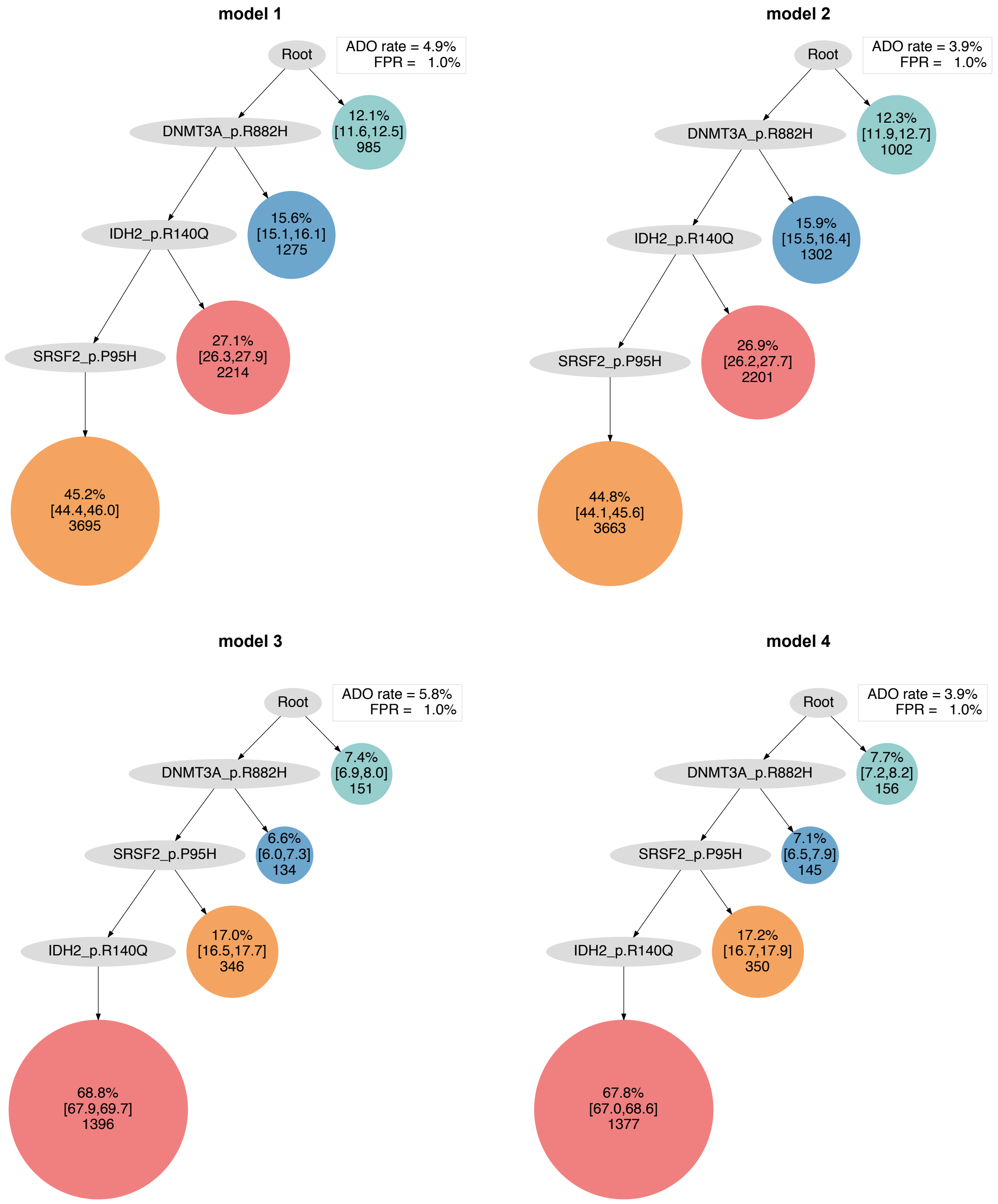
model 1
Root
ADO rate = 4.9%
FPR = 1.0%
DNMT3A_p.R882H
12.1% [11.6,12.5] 985
IDH2_p.R140Q
15.6% [15.1,16.1] 1275
SRSF2_p.P95H
27.1% [26.3,27.9] 2214
45.2% [44.4,46.0] 3695
model 2
Root
ADO rate = 3.9%
FPR = 1.0%
DNMT3A_p.R882H
12.3% [11.9,12.7] 1002
IDH2_p.R140Q
15.9% [15.5,16.4] 1302
SRSF2_p.P95H
26.9% [26.2,27.7] 2201
44.8% [44.1,45.6] 3663
model 3
Root
ADO rate = 5.8%
FPR = 1.0%
DNMT3A_p.R882H
7.4% [6.9,8.0] 151
SRSF2_p.P95H
6.6% [6.0,7.3] 134
IDH2_p.R140Q
17.0% [16.5,17.7] 346
68.8% [67.9,69.7] 1396
model 4
Root
ADO rate = 3.9%
FPR = 1.0%
DNMT3A_p.R882H
7.7% [7.2,8.2] 156
SRSF2_p.P95H
7.1% [6.5,7.9] 145
IDH2_p.R140Q
17.2% [16.7,17.9] 350
67.8% [67.0,68.6] 1377

#### AML-59-001

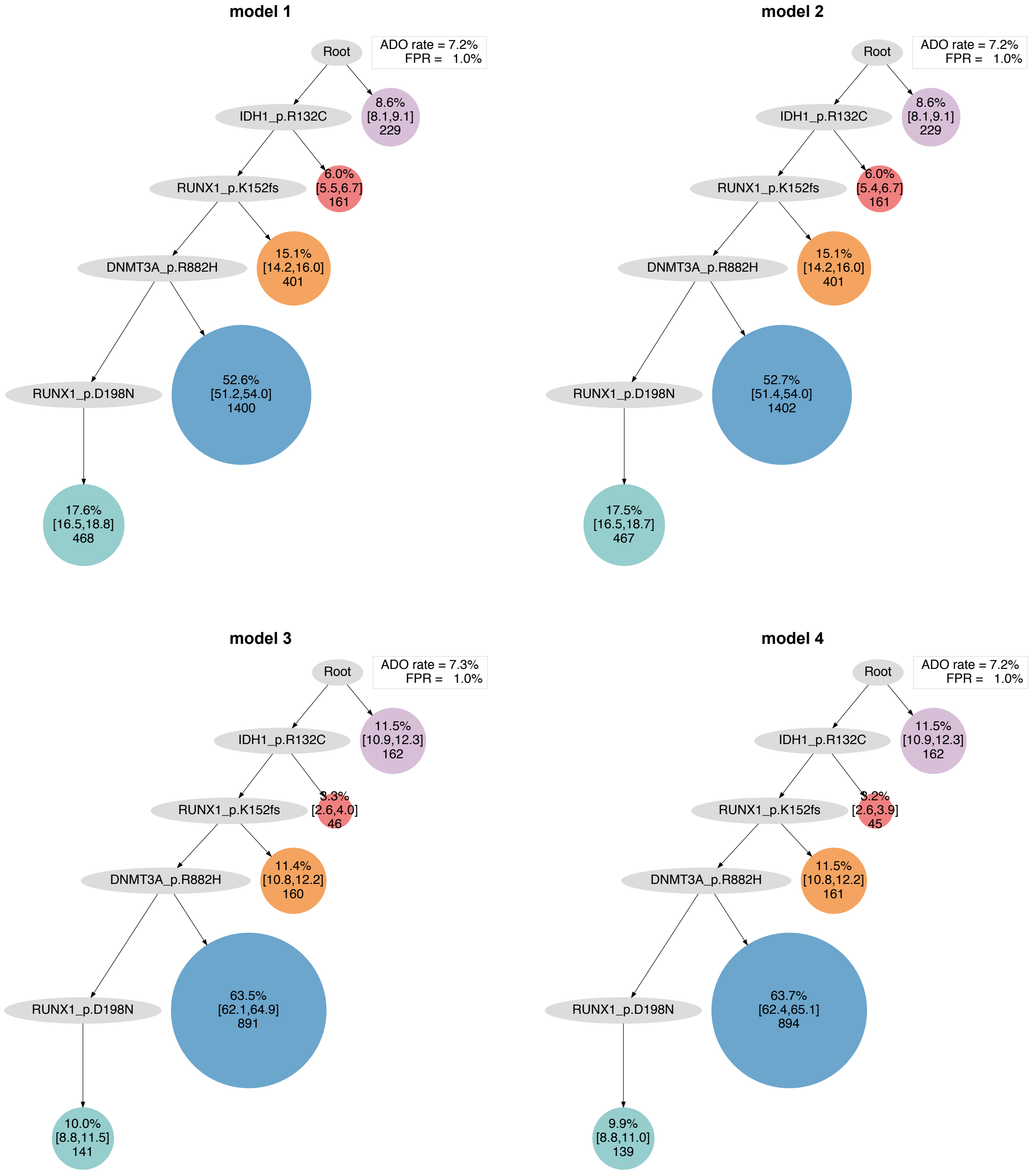
model 1
Root
ADO rate = 7.2%
FPR = 1.0%
IDH1_p.R132C
8.6%
[8.1,9.1]
229
RUNX1_p.K152fs
6.0%
[5.5,6.7]
161
DNMT3A_p.R882H
15.1%
[14.2,16.0]
401
RUNX1_p.D198N
52.6%
[51.2,54.0]
1400
17.6%
[16.5,18.8]
468
model 2
Root
ADO rate = 7.2%
FPR = 1.0%
IDH1_p.R132C
8.6%
[8.1,9.1]
229
RUNX1_p.K152fs
6.0%
[5.4,6.7]
161
DNMT3A_p.R882H
15.1%
[14.2,16.0]
401
RUNX1_p.D198N
52.7%
[51.4,54.0]
1402
17.5%
[16.5,18.7]
467
model 3
Root
ADO rate = 7.3%
FPR = 1.0%
IDH1_p.R132C
11.5%
[10.9,12.3]
162
RUNX1_p.K152fs
3.3%
[2.6,4.0]
46
DNMT3A_p.R882H
11.4%
[10.8,12.2]
160
RUNX1_p.D198N
63.5%
[62.1,64.9]
891
10.0%
[8.8,11.5]
141
model 4
Root
ADO rate = 7.2%
FPR = 1.0%
IDH1_p.R132C
11.5%
[10.9,12.3]
162
RUNX1_p.K152fs
3.2%
[2.6,3.9]
45
DNMT3A_p.R882H
11.5%
[10.8,12.2]
161
RUNX1_p.D198N
63.7%
[62.4,65.1]
894
9.9%
[8.8,11.0]
139

### AML-60-001

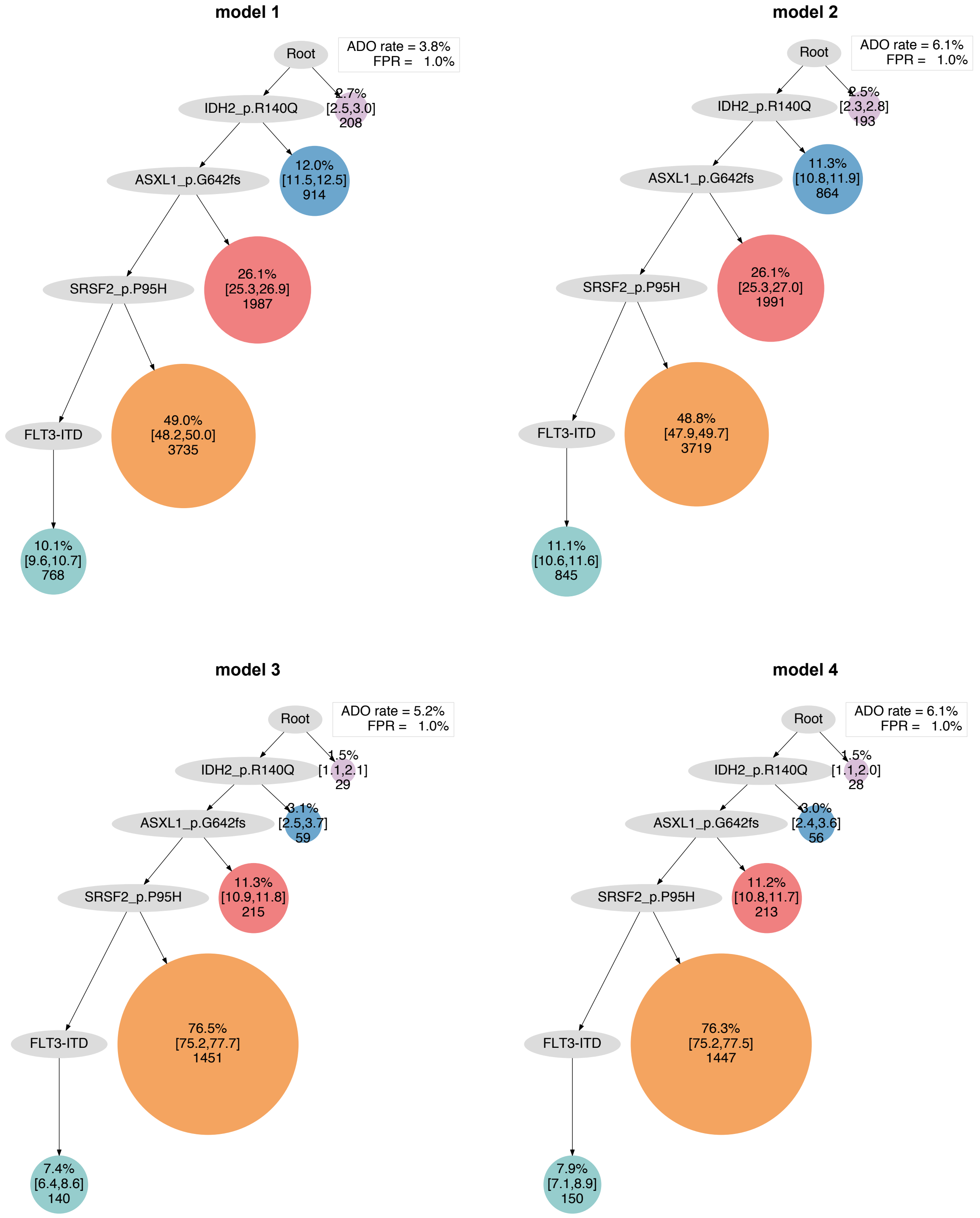

model 1
Root
ADO rate = 3.8%
FPR = 1.0%
IDH2_p.R140Q
2.7%
[2.5,3.0]
208
ASXL1_p.G642fs
12.0%
[11.5,12.5]
914
SRSF2_p.P95H
26.1%
[25.3,26.9]
1987
FLT3-ITD
49.0%
[48.2,50.0]
3735
10.1%
[9.6,10.7]
768
model 2
Root
ADO rate = 6.1%
FPR = 1.0%
IDH2_p.R140Q
2.5%
[2.3,2.8]
193
ASXL1_p.G642fs
11.3%
[10.8,11.9]
864
SRSF2_p.P95H
26.1%
[25.3,27.0]
1991
FLT3-ITD
48.8%
[47.9,49.7]
3719
11.1%
[10.6,11.6]
845
model 3
Root
ADO rate = 5.2%
FPR = 1.0%
IDH2_p.R140Q
1.5%
[1.1,2.1]
29
ASXL1_p.G642fs
3.1%
[2.5,3.7]
59
SRSF2_p.P95H
11.3%
[10.9,11.8]
215
FLT3-ITD
76.5%
[75.2,77.7]
1451
7.4%
[6.4,8.6]
140
model 4
Root
ADO rate = 6.1%
FPR = 1.0%
IDH2_p.R140Q
1.5%
[1.1,2.0]
28
ASXL1_p.G642fs
3.0%
[2.4,3.6]
56
SRSF2_p.P95H
11.2%
[10.8,11.7]
213
FLT3-ITD
76.3%
[75.2,77.5]
1447
7.9%
[7.1,8.9]
150

### AML-61-001

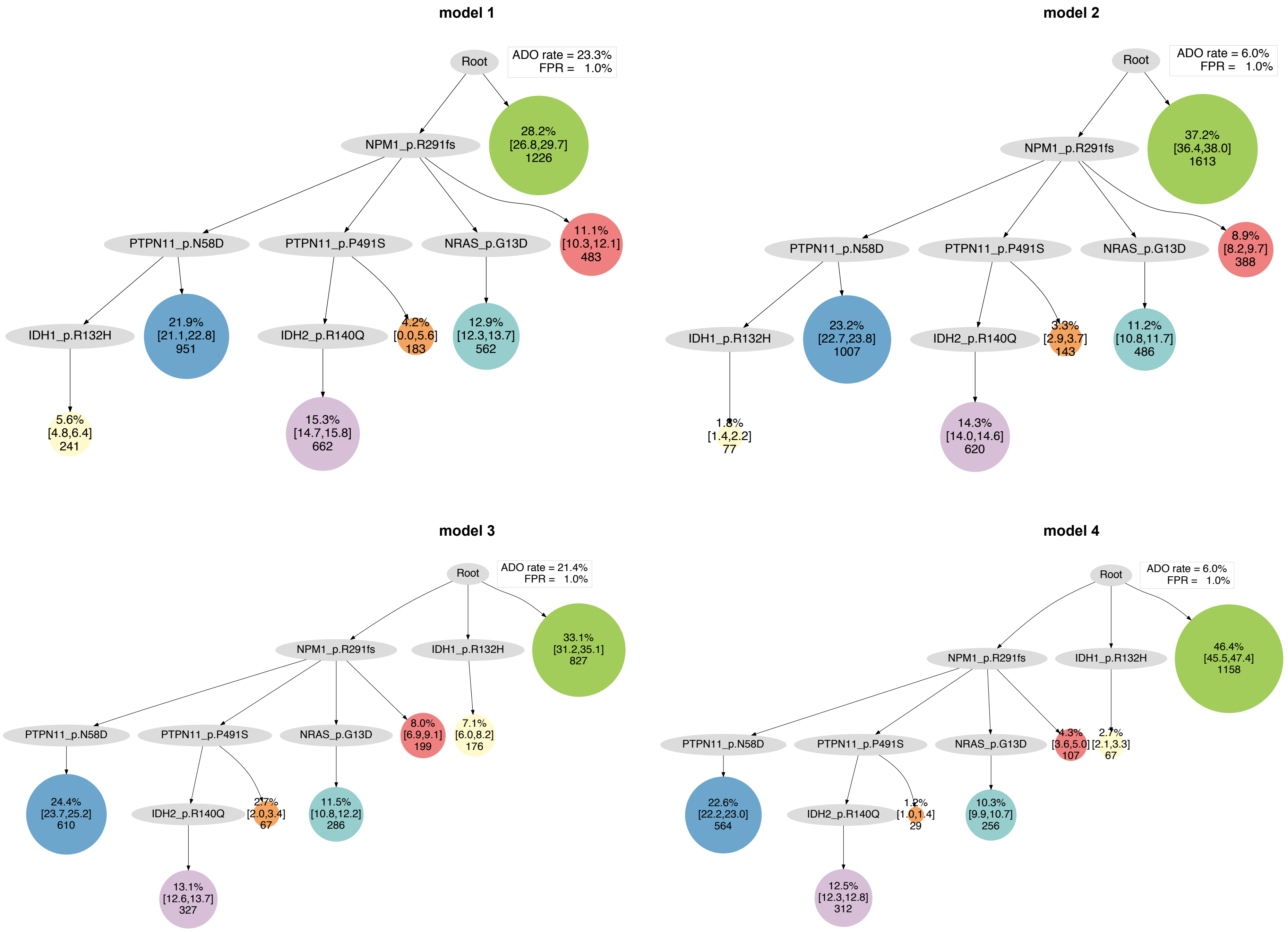

model 1
Root
ADO rate = 23.3%
FPR = 1.0%
NPM1_p.R291fs
28.2% [26.8,29.7] 1226
PTPN11_p.N58D
PTPN11_p.P491S
NRAS_p.G13D
11.1% [10.3,12.1] 483
IDH1_p.R132H
21.9% [21.1,22.8] 951
IDH2_p.R140Q
4.2% [0.0,5.6] 183
12.9% [12.3,13.7] 562
5.6% [4.8,6.4] 241
15.3% [14.7,15.8] 662
model 2
Root
ADO rate = 6.0%
FPR = 1.0%
NPM1_p.R291fs
37.2% [36.4,38.0] 1613
PTPN11_p.N58D
PTPN11_p.P491S
NRAS_p.G13D
8.9% [8.2,9.7] 388
IDH1_p.R132H
23.2% [22.7,23.8] 1007
IDH2_p.R140Q
3.3% [2.9,3.7] 143
11.2% [10.8,11.7] 486
1.8% [1.4,2.2] 77
14.3% [14.0,14.6] 620
model 3
Root
ADO rate = 21.4%
FPR = 1.0%
NPM1_p.R291fs
IDH1_p.R132H
33.1% [31.2,35.1] 827
PTPN11_p.N58D
PTPN11_p.P491S
NRAS_p.G13D
8.0% [6.9,9.1] 199
7.1% [6.0,8.2] 176
24.4% [23.7,25.2] 610
IDH2_p.R140Q
2.7% [2.0,3.4] 67
11.5% [10.8,12.2] 286
13.1% [12.6,13.7] 327
model 4
Root
ADO rate = 6.0%
FPR = 1.0%
NPM1_p.R291fs
IDH1_p.R132H
46.4% [45.5,47.4] 1158
PTPN11_p.N58D
PTPN11_p.P491S
NRAS_p.G13D
4.3% [3.6,5.0] 107
2.7% [2.1,3.3] 67
22.6% [22.2,23.0] 564
IDH2_p.R140Q
1.2% [1.0,1.4] 29
10.3% [9.9,10.7] 256
12.5% [12.3,12.8] 312

#### AML-62-001

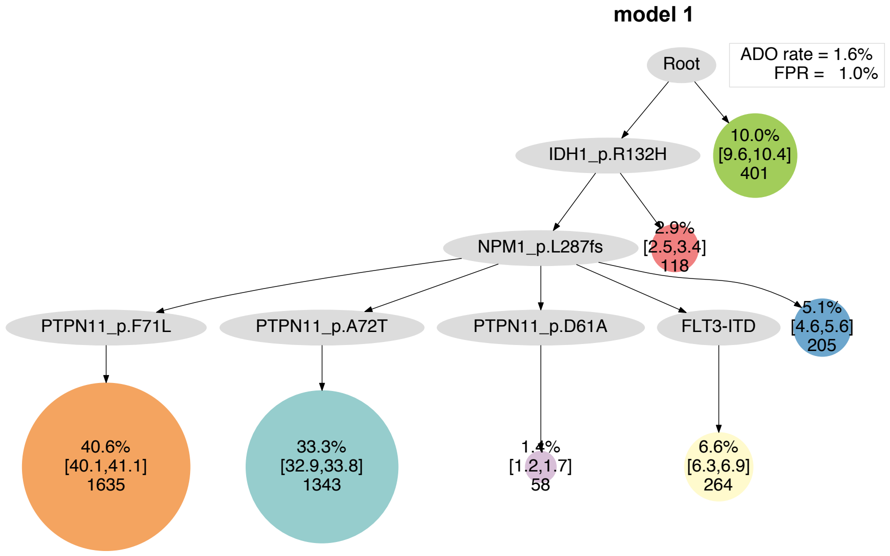
model 1
ADO rate = 1.6%
FPR = 1.0%
Root
IDH1_p.R132H
10.0%
[9.6,10.4]
401
NPM1_p.L287fs
2.9%
[2.5,3.4]
118
PTPN11_p.F71L
PTPN11_p.A72T
PTPN11_p.D61A
FLT3-ITD
5.1%
[4.6,5.6]
205
40.6%
[40.1,41.1]
1635
33.3%
[32.9,33.8]
1343
1.4%
[1.2,1.7]
58
6.6%
[6.3,6.9]
264

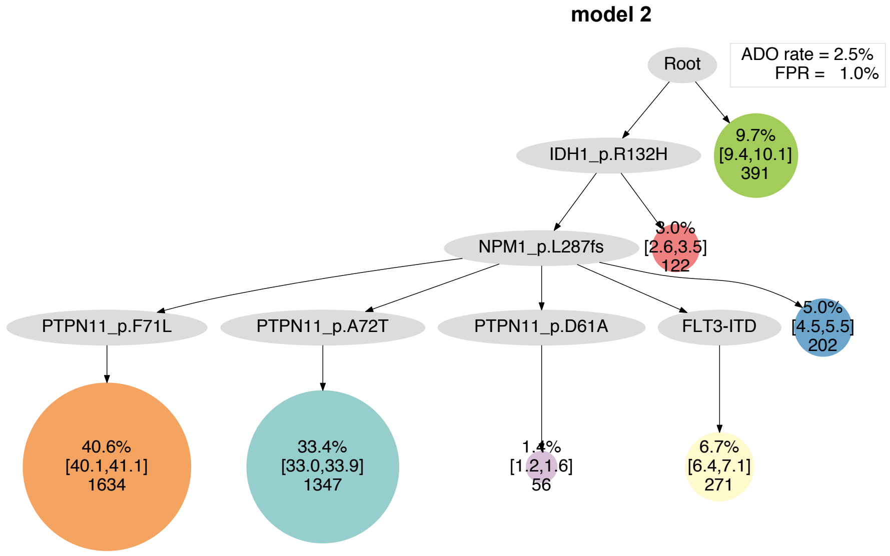
model 2
ADO rate = 2.5%
FPR = 1.0%
Root
IDH1_p.R132H
9.7%
[9.4,10.1]
391
NPM1_p.L287fs
3.0%
[2.6,3.5]
122
PTPN11_p.F71L
PTPN11_p.A72T
PTPN11_p.D61A
FLT3-ITD
5.0%
[4.5,5.5]
202
40.6%
[40.1,41.1]
1634
33.4%
[33.0,33.9]
1347
1.4%
[1.2,1.6]
56
6.7%
[6.4,7.1]
271

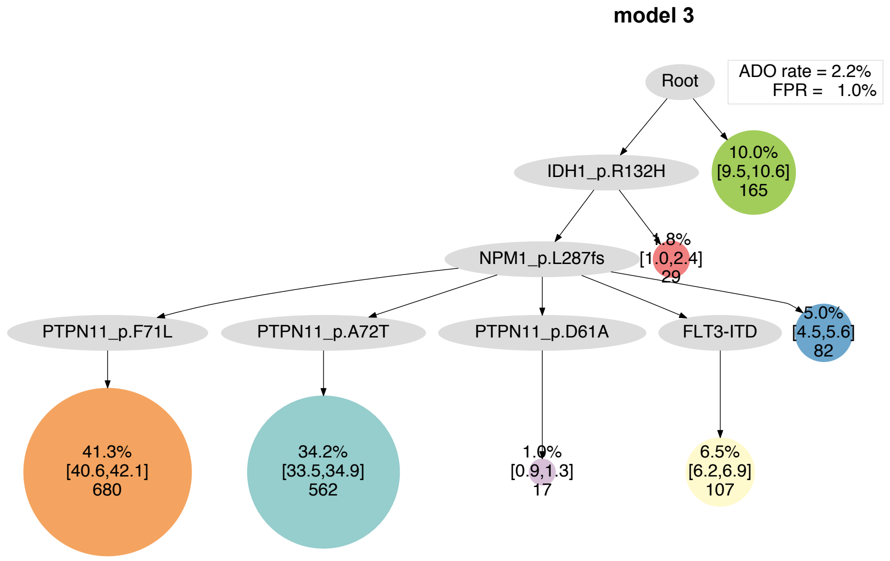
model 3
ADO rate = 2.2%
FPR = 1.0%
Root
IDH1_p.R132H
10.0%
[9.5,10.6]
165
NPM1_p.L287fs
1.8%
[1.0,2.4]
29
PTPN11_p.F71L
PTPN11_p.A72T
PTPN11_p.D61A
FLT3-ITD
5.0%
[4.5,5.6]
82
41.3%
[40.6,42.1]
680
34.2%
[33.5,34.9]
562
1.0%
[0.9,1.3]
17
6.5%
[6.2,6.9]
107

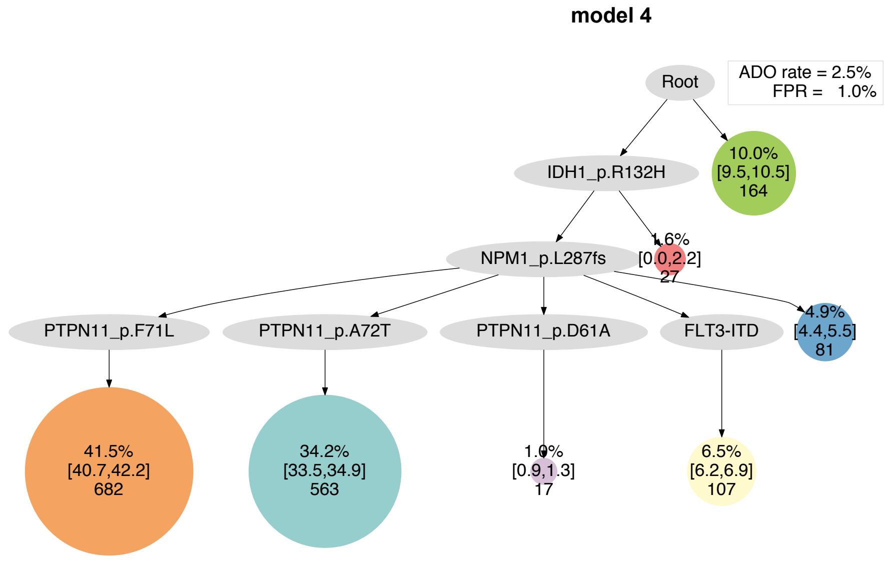
model 4
ADO rate = 2.5%
FPR = 1.0%
Root
IDH1_p.R132H
10.0%
[9.5,10.5]
164
NPM1_p.L287fs
1.6%
[0.0,2.2]
27
PTPN11_p.F71L
PTPN11_p.A72T
PTPN11_p.D61A
FLT3-ITD
4.9%
[4.4,5.5]
81
41.5%
[40.7,42.2]
682
34.2%
[33.5,34.9]
563
1.0%
[0.9,1.3]
17
6.5%
[6.2,6.9]
107

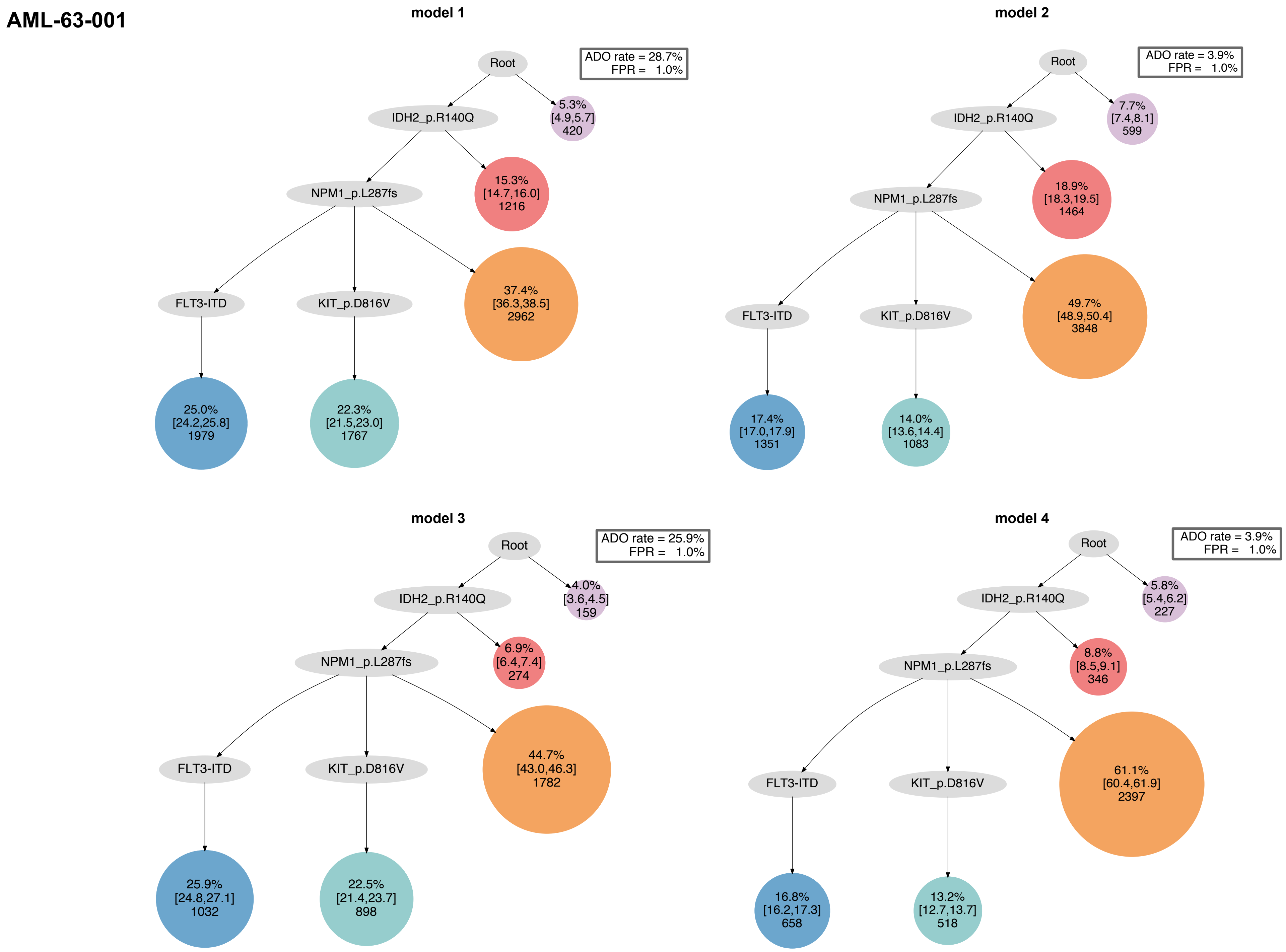
AML-63-001
model 1
Root
ADO rate = 28.7%
FPR = 1.0%
IDH2_p.R140Q
5.3%
[4.9,5.7]
420
NPM1_p.L287fs
15.3%
[14.7,16.0]
1216
FLT3-ITD
KIT_p.D816V
37.4%
[36.3,38.5]
2962
25.0%
[24.2,25.8]
1979
22.3%
[21.5,23.0]
1767
model 2
Root
ADO rate = 3.9%
FPR = 1.0%
IDH2_p.R140Q
7.7%
[7.4,8.1]
599
NPM1_p.L287fs
18.9%
[18.3,19.5]
1464
FLT3-ITD
KIT_p.D816V
49.7%
[48.9,50.4]
3848
17.4%
[17.0,17.9]
1351
14.0%
[13.6,14.4]
1083
model 3
Root
ADO rate = 25.9%
FPR = 1.0%
IDH2_p.R140Q
4.0%
[3.6,4.5]
159
NPM1_p.L287fs
6.9%
[6.4,7.4]
274
FLT3-ITD
KIT_p.D816V
44.7%
[43.0,46.3]
1782
25.9%
[24.8,27.1]
1032
22.5%
[21.4,23.7]
898
model 4
Root
ADO rate = 3.9%
FPR = 1.0%
IDH2_p.R140Q
5.8%
[5.4,6.2]
227
NPM1_p.L287fs
8.8%
[8.5,9.1]
346
FLT3-ITD
KIT_p.D816V
61.1%
[60.4,61.9]
2397
16.8%
[16.2,17.3]
658
13.2%
[12.7,13.7]
518

#### AML-64-001

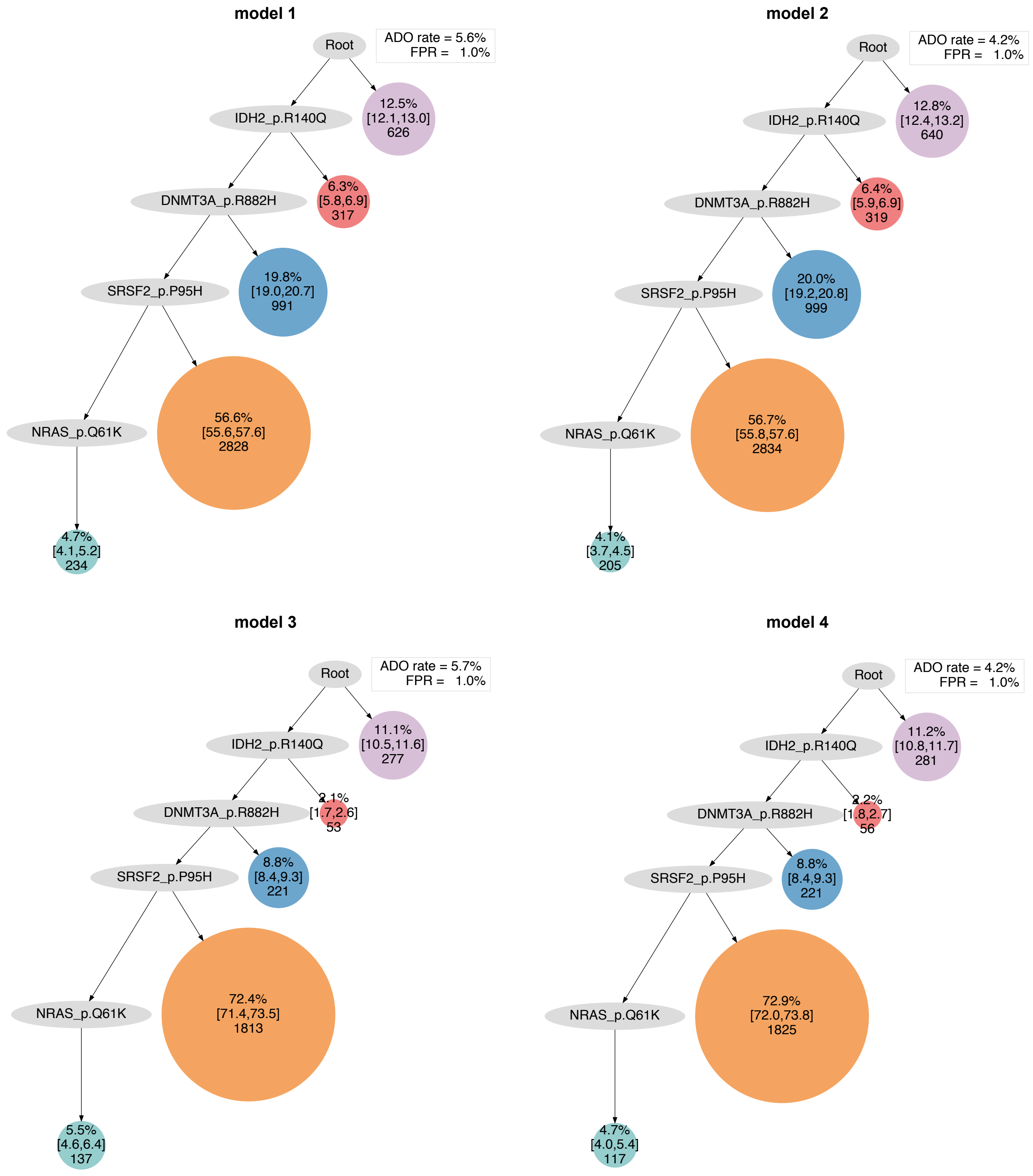
model 1
Root
ADO rate = 5.6%
FPR = 1.0%
IDH2_p.R140Q
12.5%
[12.1,13.0]
626
DNMT3A_p.R882H
6.3%
[5.8,6.9]
317
SRSF2_p.P95H
19.8%
[19.0,20.7]
991
NRAS_p.Q61K
56.6%
[55.6,57.6]
2828
4.7%
[4.1,5.2]
234
model 2
Root
ADO rate = 4.2%
FPR = 1.0%
IDH2_p.R140Q
12.8%
[12.4,13.2]
640
DNMT3A_p.R882H
6.4%
[5.9,6.9]
319
SRSF2_p.P95H
20.0%
[19.2,20.8]
999
NRAS_p.Q61K
56.7%
[55.8,57.6]
2834
4.1%
[3.7,4.5]
205
model 3
Root
ADO rate = 5.7%
FPR = 1.0%
IDH2_p.R140Q
11.1%
[10.5,11.6]
277
DNMT3A_p.R882H
2.1%
[1.7,2.6]
53
SRSF2_p.P95H
8.8%
[8.4,9.3]
221
NRAS_p.Q61K
72.4%
[71.4,73.5]
1813
5.5%
[4.6,6.4]
137
model 4
Root
ADO rate = 4.2%
FPR = 1.0%
IDH2_p.R140Q
11.2%
[10.8,11.7]
281
DNMT3A_p.R882H
2.2%
[1.8,2.7]
56
SRSF2_p.P95H
8.8%
[8.4,9.3]
221
NRAS_p.Q61K
72.9%
[72.0,73.8]
1825
4.7%
[4.0,5.4]
117

### AML-65-001

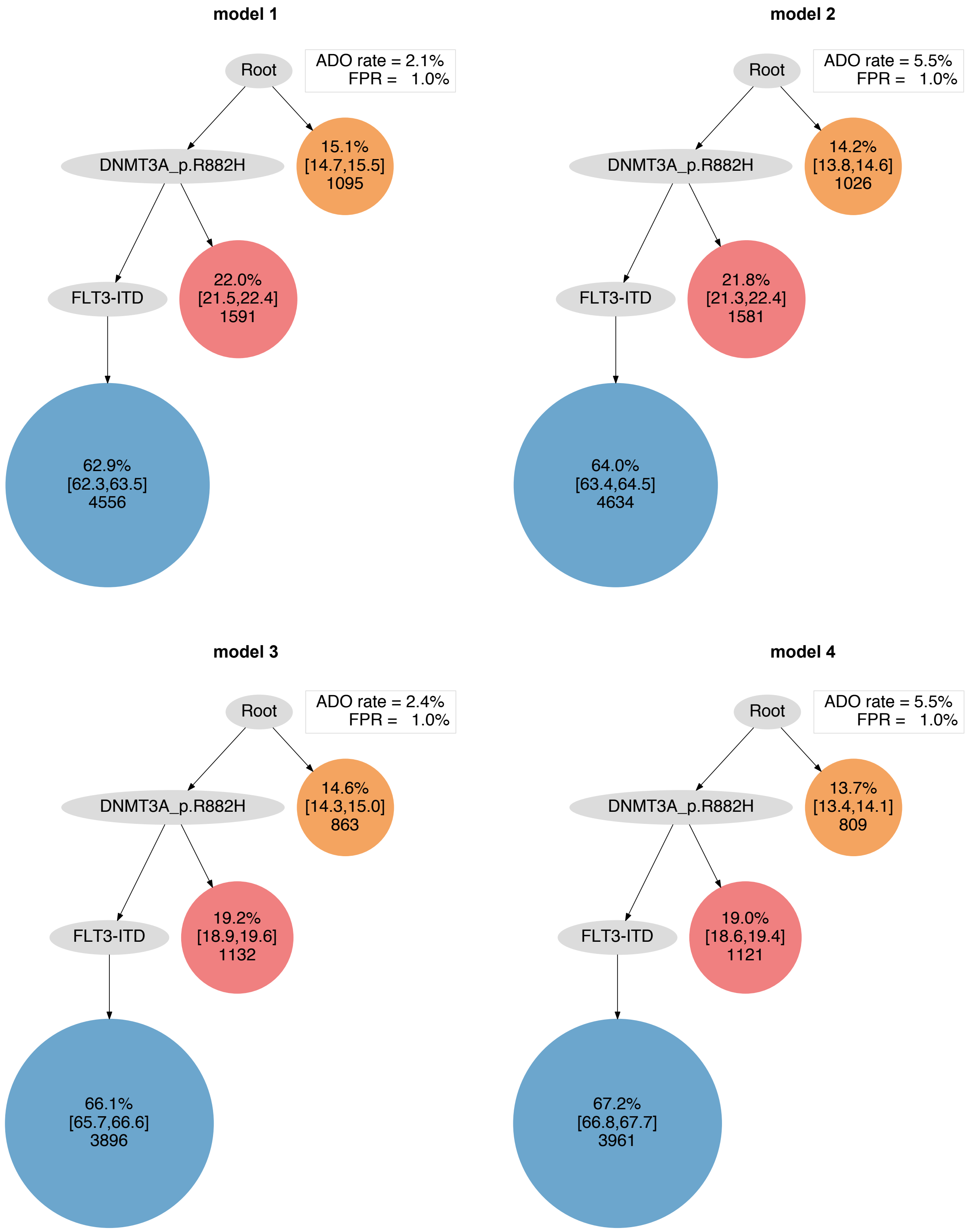

model 1
Root
ADO rate = 2.1%
FPR = 1.0%
DNMT3A_p.R882H
15.1%
[14.7,15.5]
1095
FLT3-ITD
22.0%
[21.5,22.4]
1591
62.9%
[62.3,63.5]
4556
model 2
Root
ADO rate = 5.5%
FPR = 1.0%
DNMT3A_p.R882H
14.2%
[13.8,14.6]
1026
FLT3-ITD
21.8%
[21.3,22.4]
1581
64.0%
[63.4,64.5]
4634
model 3
Root
ADO rate = 2.4%
FPR = 1.0%
DNMT3A_p.R882H
14.6%
[14.3,15.0]
863
FLT3-ITD
19.2%
[18.9,19.6]
1132
66.1%
[65.7,66.6]
3896
model 4
Root
ADO rate = 5.5%
FPR = 1.0%
DNMT3A_p.R882H
13.7%
[13.4,14.1]
809
FLT3-ITD
19.0%
[18.6,19.4]
1121
67.2%
[66.8,67.7]
3961

**AML-66-001**

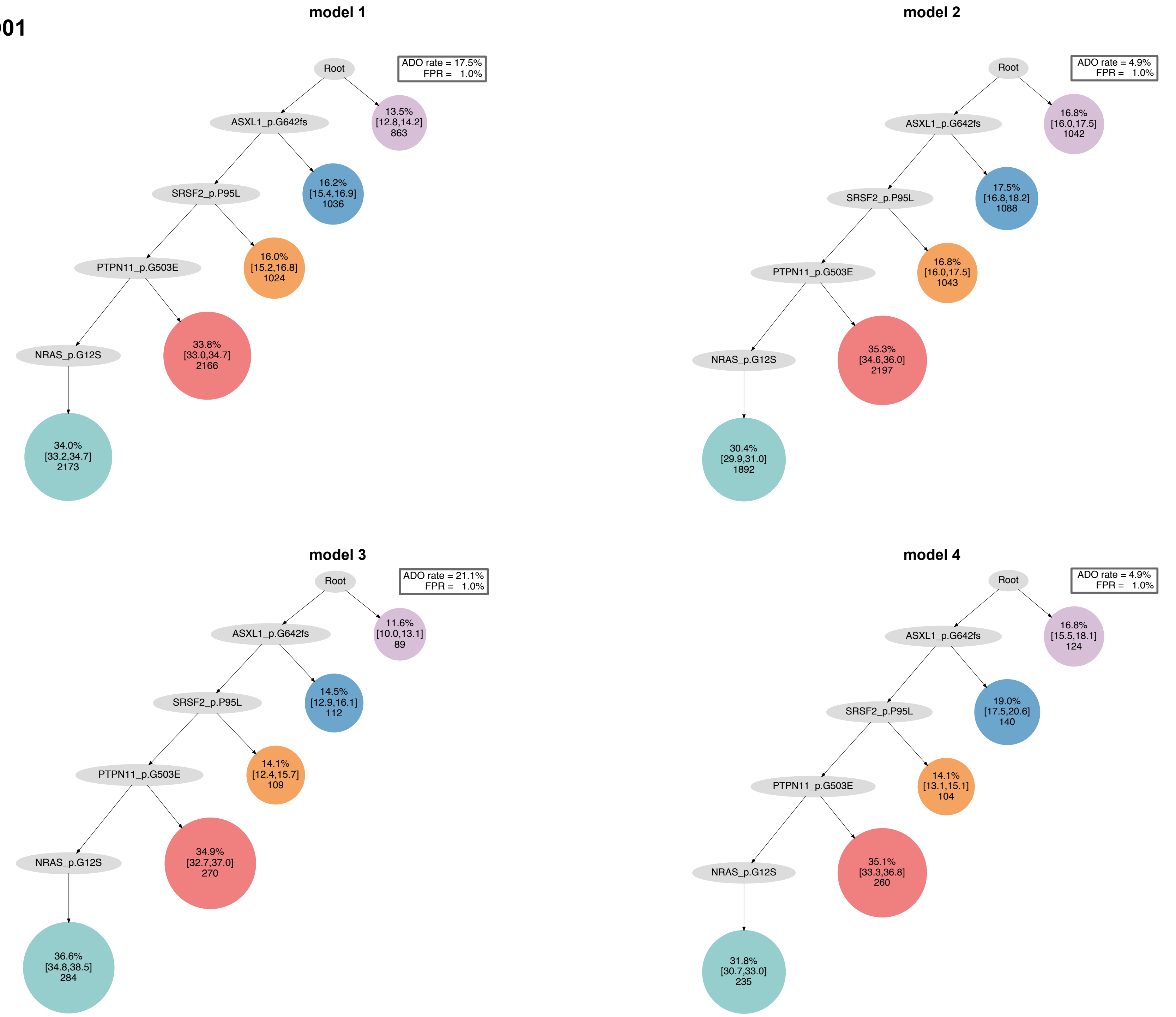
model 1
Root
ADO rate = 17.5%
FPR = 1.0%
ASXL1_p.G642fs
13.5% [12.8,14.2] 863
SRSF2_p.P95L
16.2% [15.4,16.9] 1036
PTPN11_p.G503E
16.0% [15.2,16.8] 1024
NRAS_p.G12S
33.8% [33.0,34.7] 2166
34.0% [33.2,34.7] 2173
model 2
Root
ADO rate = 4.9%
FPR = 1.0%
ASXL1_p.G642fs
16.8% [16.0,17.5] 1042
SRSF2_p.P95L
17.5% [16.8,18.2] 1088
PTPN11_p.G503E
16.8% [16.0,17.5] 1043
NRAS_p.G12S
35.3% [34.6,36.0] 2197
30.4% [29.9,31.0] 1892
model 3
Root
ADO rate = 21.1%
FPR = 1.0%
ASXL1_p.G642fs
11.6% [10.0,13.1] 89
SRSF2_p.P95L
14.5% [12.9,16.1] 112
PTPN11_p.G503E
14.1% [12.4,15.7] 109
NRAS_p.G12S
34.9% [32.7,37.0] 270
36.6% [34.8,38.5] 284
model 4
Root
ADO rate = 4.9%
FPR = 1.0%
ASXL1_p.G642fs
16.8% [15.5,18.1] 124
SRSF2_p.P95L
19.0% [17.5,20.6] 140
PTPN11_p.G503E
14.1% [13.1,15.1] 104
NRAS_p.G12S
35.1% [33.3,36.8] 260
31.8% [30.7,33.0] 235

### AML-67-001

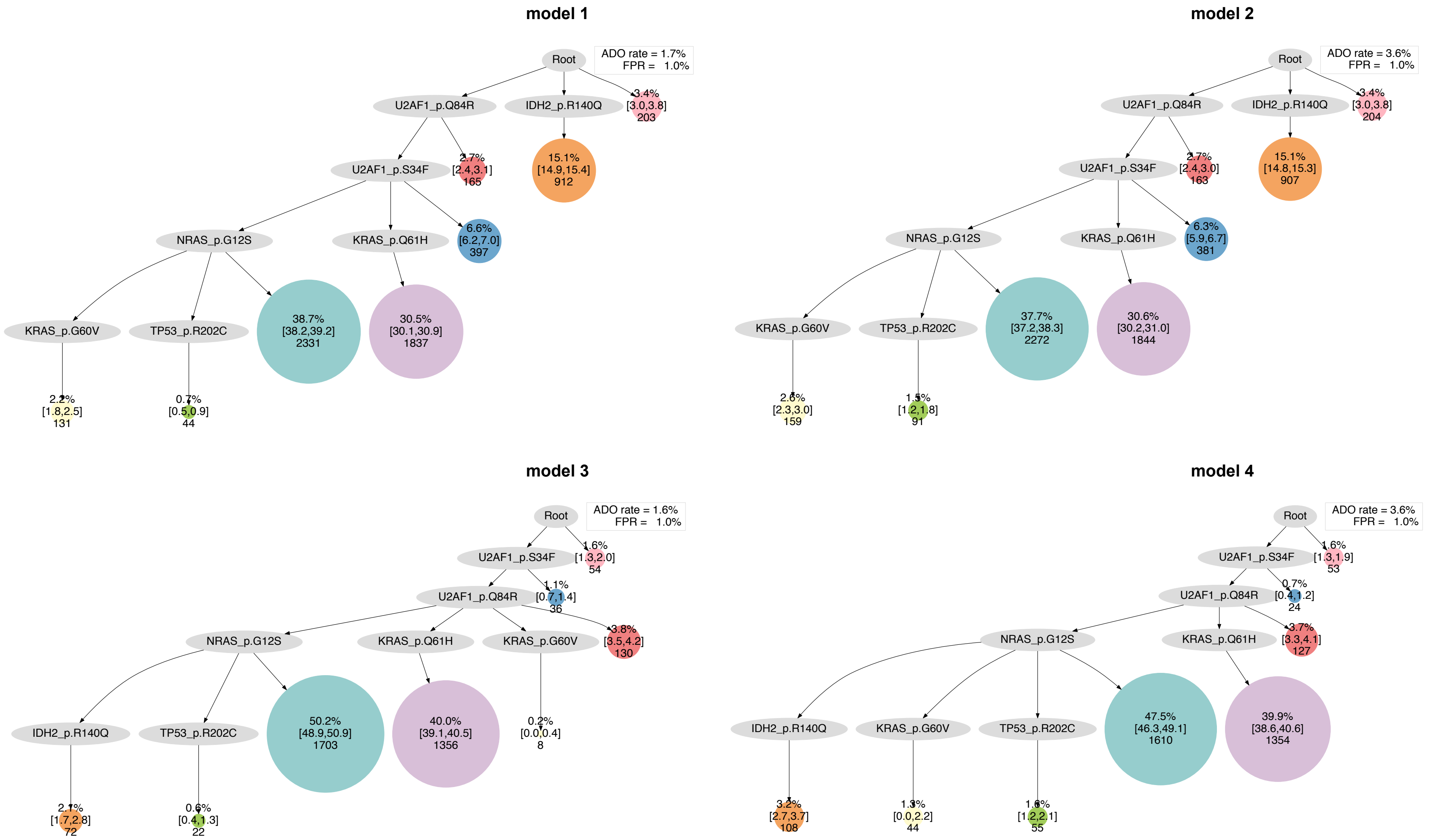
model 1
Root
ADO rate = 1.7%
FPR = 1.0%
U2AF1_p.Q84R
IDH2_p.R140Q
3.4%
[3.0,3.8]
203
U2AF1_p.S34F
2.7%
[2.4,3.1]
165
15.1%
[14.9,15.4]
912
NRAS_p.G12S
KRAS_p.Q61H
6.6%
[6.2,7.0]
397
KRAS_p.G60V
TP53_p.R202C
38.7%
[38.2,39.2]
2331
30.5%
[30.1,30.9]
1837
2.2%
[1.8,2.5]
131
0.7%
[0.5,0.9]
44
model 2
Root
ADO rate = 3.6%
FPR = 1.0%
U2AF1_p.Q84R
IDH2_p.R140Q
3.4%
[3.0,3.8]
204
U2AF1_p.S34F
2.7%
[2.4,3.0]
163
15.1%
[14.8,15.3]
907
NRAS_p.G12S
KRAS_p.Q61H
6.3%
[5.9,6.7]
381
KRAS_p.G60V
TP53_p.R202C
37.7%
[37.2,38.3]
2272
30.6%
[30.2,31.0]
1844
2.6%
[2.3,3.0]
159
1.5%
[1.2,1.8]
91
model 3
Root
ADO rate = 1.6%
FPR = 1.0%
U2AF1_p.S34F
1.6%
[1.3,2.0]
54
U2AF1_p.Q84R
1.1%
[0.7,1.4]
36
NRAS_p.G12S
KRAS_p.Q61H
KRAS_p.G60V
3.8%
[3.5,4.2]
130
IDH2_p.R140Q
TP53_p.R202C
50.2%
[48.9,50.9]
1703
40.0%
[39.1,40.5]
1356
0.2%
[0.0,0.4]
8
2.1%
[1.7,2.8]
72
0.6%
[0.4,1.3]
22
model 4
Root
ADO rate = 3.6%
FPR = 1.0%
U2AF1_p.S34F
1.6%
[1.3,1.9]
53
U2AF1_p.Q84R
0.7%
[0.4,1.2]
24
NRAS_p.G12S
KRAS_p.Q61H
3.7%
[3.3,4.1]
127
IDH2_p.R140Q
KRAS_p.G60V
TP53_p.R202C
47.5%
[46.3,49.1]
1610
39.9%
[38.6,40.6]
1354
3.2%
[2.7,3.7]
108
1.3%
[0.0,2.2]
44
1.6%
[1.2,2.1]
55

AML-68-001

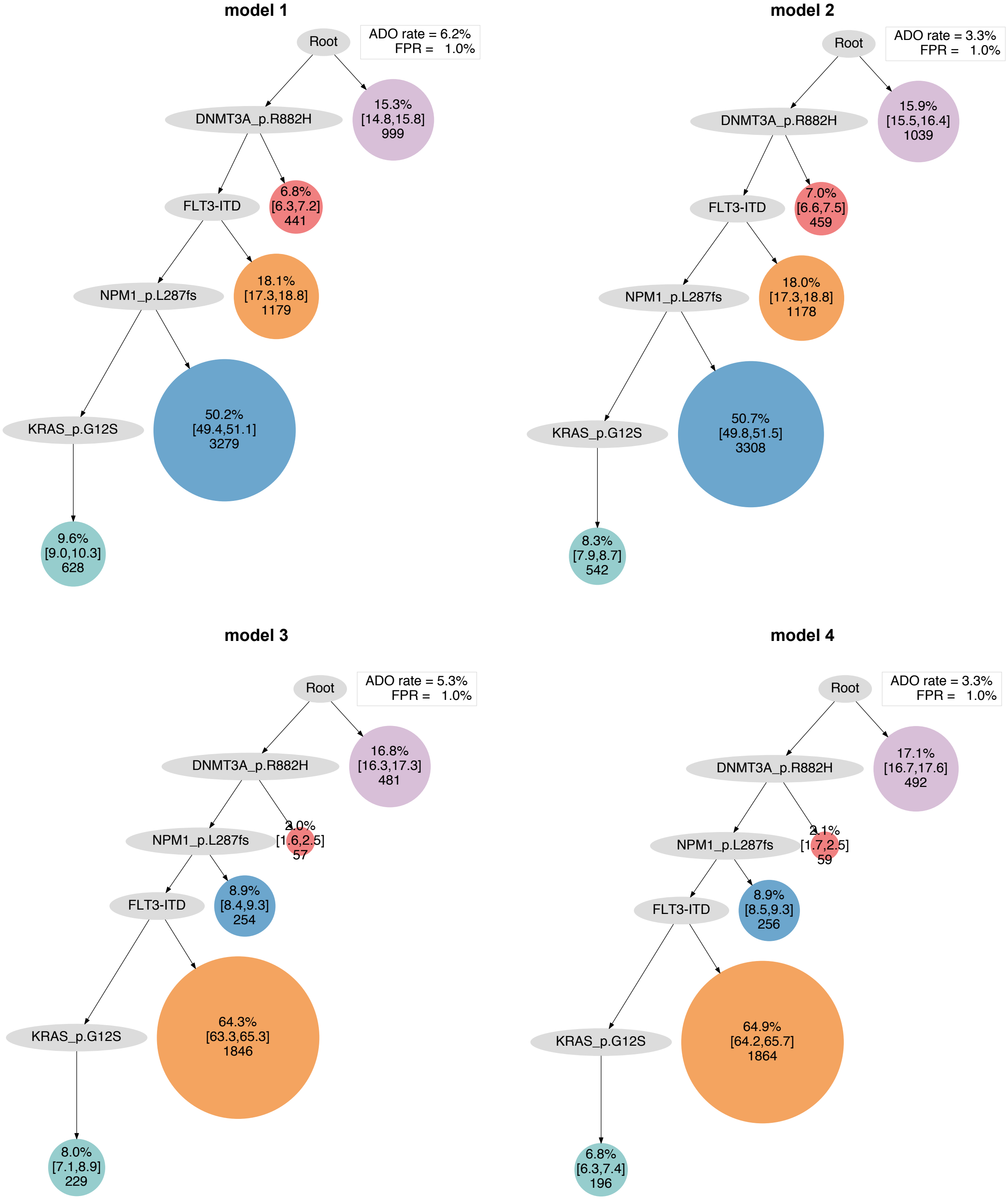
model 1
Root
ADO rate = 6.2%
FPR = 1.0%
DNMT3A_p.R882H
15.3%
[14.8,15.8]
999
FLT3-ITD
6.8%
[6.3,7.2]
441
NPM1_p.L287fs
18.1%
[17.3,18.8]
1179
KRAS_p.G12S
50.2%
[49.4,51.1]
3279
9.6%
[9.0,10.3]
628
model 2
Root
ADO rate = 3.3%
FPR = 1.0%
DNMT3A_p.R882H
15.9%
[15.5,16.4]
1039
FLT3-ITD
7.0%
[6.6,7.5]
459
NPM1_p.L287fs
18.0%
[17.3,18.8]
1178
KRAS_p.G12S
50.7%
[49.8,51.5]
3308
8.3%
[7.9,8.7]
542
model 3
Root
ADO rate = 5.3%
FPR = 1.0%
DNMT3A_p.R882H
16.8%
[16.3,17.3]
481
NPM1_p.L287fs
2.0%
[1.6,2.5]
57
FLT3-ITD
8.9%
[8.4,9.3]
254
KRAS_p.G12S
64.3%
[63.3,65.3]
1846
8.0%
[7.1,8.9]
229
model 4
Root
ADO rate = 3.3%
FPR = 1.0%
DNMT3A_p.R882H
17.1%
[16.7,17.6]
492
NPM1_p.L287fs
2.1%
[1.7,2.5]
59
FLT3-ITD
8.9%
[8.5,9.3]
256
KRAS_p.G12S
64.9%
[64.2,65.7]
1864
6.8%
[6.3,7.4]
196

**AML-69-001**

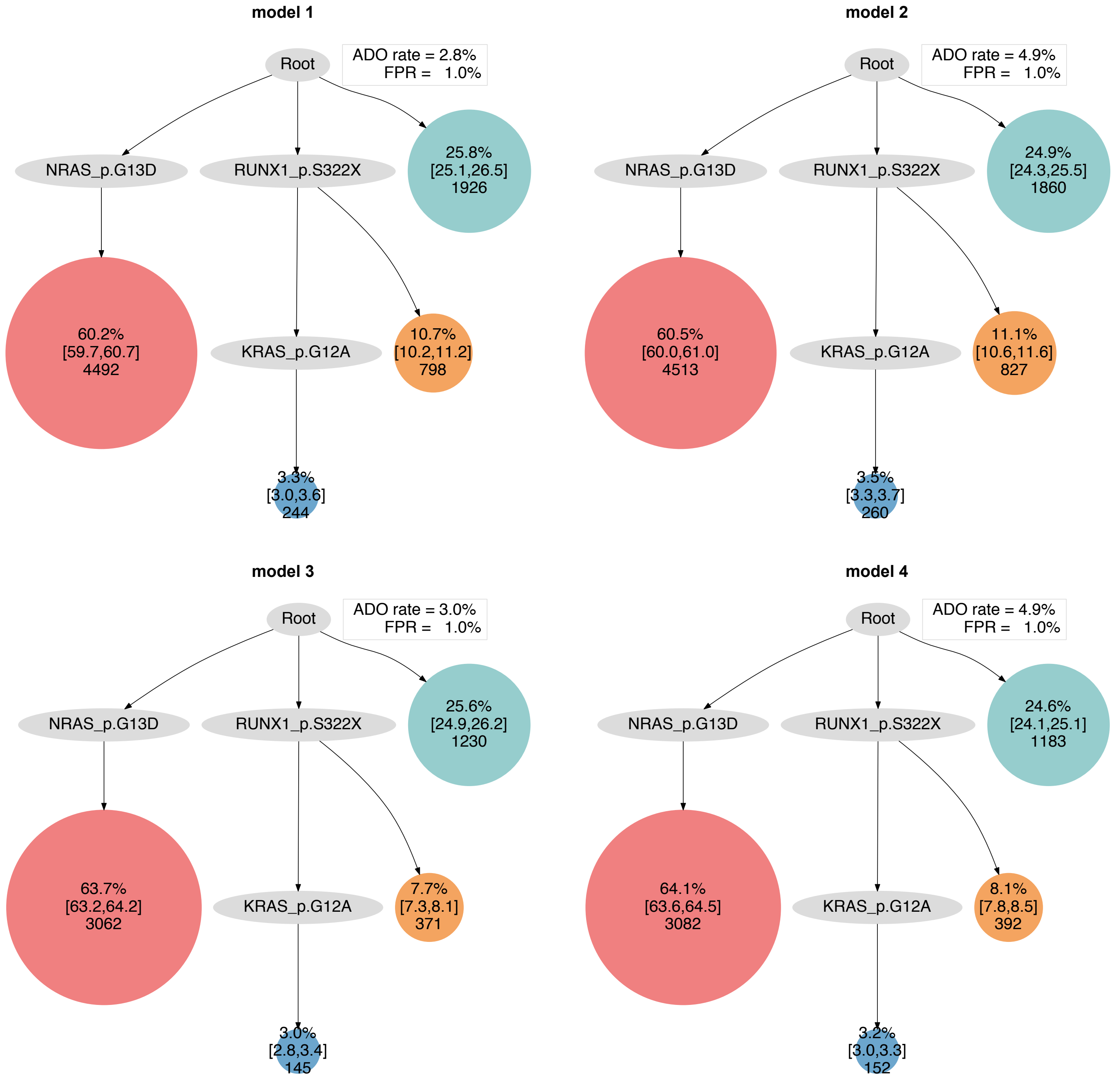
model 1
Root
ADO rate = 2.8%
FPR = 1.0%
NRAS_p.G13D
RUNX1_p.S322X
25.8%
[25.1,26.5]
1926
60.2%
[59.7,60.7]
4492
KRAS_p.G12A
10.7%
[10.2,11.2]
798
3.3%
[3.0,3.6]
244
model 2
Root
ADO rate = 4.9%
FPR = 1.0%
NRAS_p.G13D
RUNX1_p.S322X
24.9%
[24.3,25.5]
1860
60.5%
[60.0,61.0]
4513
KRAS_p.G12A
11.1%
[10.6,11.6]
827
3.5%
[3.3,3.7]
260
model 3
Root
ADO rate = 3.0%
FPR = 1.0%
NRAS_p.G13D
RUNX1_p.S322X
25.6%
[24.9,26.2]
1230
63.7%
[63.2,64.2]
3062
KRAS_p.G12A
7.7%
[7.3,8.1]
371
3.0%
[2.8,3.4]
145
model 4
Root
ADO rate = 4.9%
FPR = 1.0%
NRAS_p.G13D
RUNX1_p.S322X
24.6%
[24.1,25.1]
1183
64.1%
[63.6,64.5]
3082
KRAS_p.G12A
8.1%
[7.8,8.5]
392
3.2%
[3.0,3.3]
152

#### AML-70-001

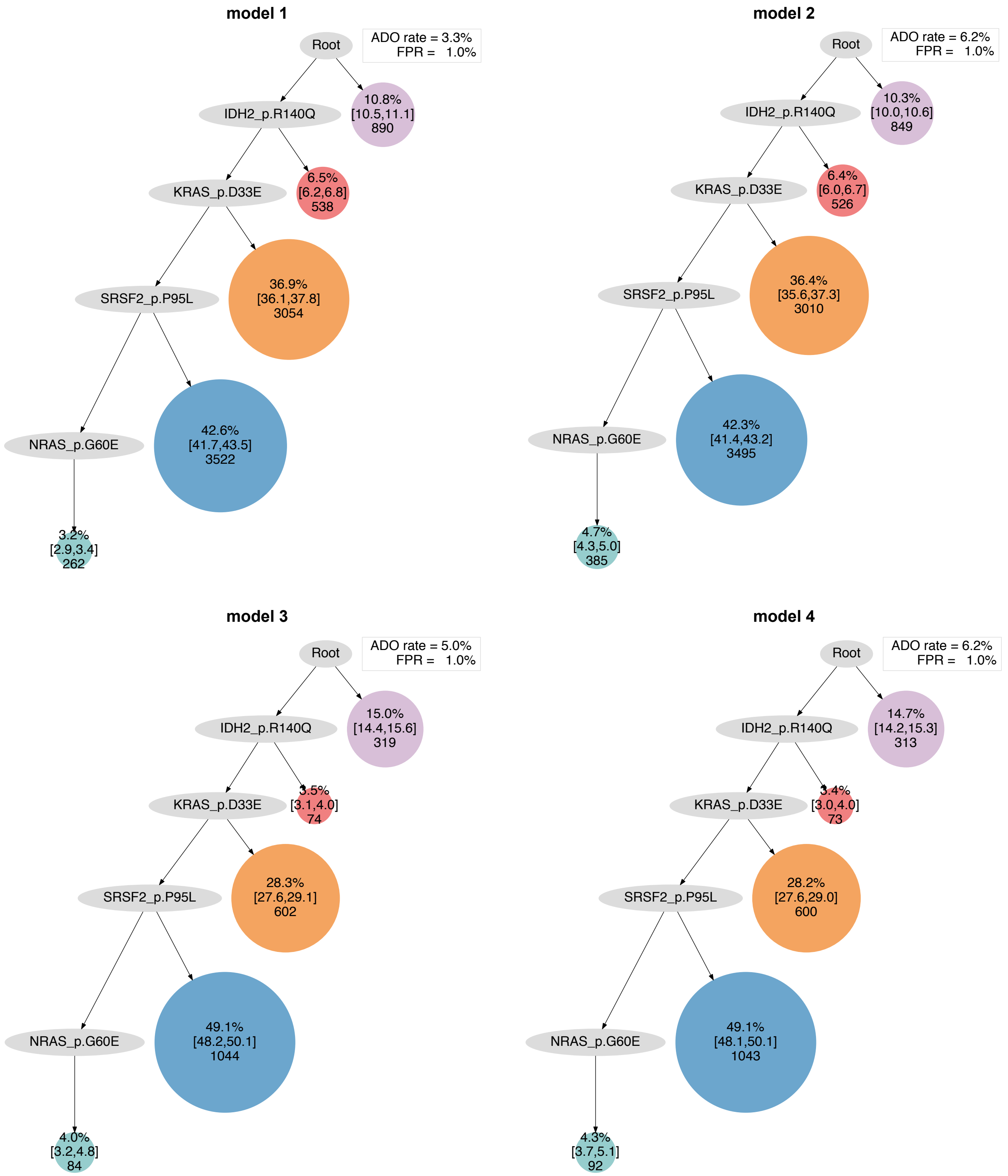

model 1
Root
ADO rate = 3.3%
FPR = 1.0%
IDH2_p.R140Q
10.8% [10.5,11.1] 890
KRAS_p.D33E
6.5% [6.2,6.8] 538
SRSF2_p.P95L
36.9% [36.1,37.8] 3054
NRAS_p.G60E
42.6% [41.7,43.5] 3522
3.2% [2.9,3.4] 262
model 2
Root
ADO rate = 6.2%
FPR = 1.0%
IDH2_p.R140Q
10.3% [10.0,10.6] 849
KRAS_p.D33E
6.4% [6.0,6.7] 526
SRSF2_p.P95L
36.4% [35.6,37.3] 3010
NRAS_p.G60E
42.3% [41.4,43.2] 3495
4.7% [4.3,5.0] 385
model 3
Root
ADO rate = 5.0%
FPR = 1.0%
IDH2_p.R140Q
15.0% [14.4,15.6] 319
KRAS_p.D33E
3.5% [3.1,4.0] 74
SRSF2_p.P95L
28.3% [27.6,29.1] 602
NRAS_p.G60E
49.1% [48.2,50.1] 1044
4.0% [3.2,4.8] 84
model 4
Root
ADO rate = 6.2%
FPR = 1.0%
IDH2_p.R140Q
14.7% [14.2,15.3] 313
KRAS_p.D33E
3.4% [3.0,4.0] 73
SRSF2_p.P95L
28.2% [27.6,29.0] 600
NRAS_p.G60E
49.1% [48.1,50.1] 1043
4.3% [3.7,5.1] 92

**AML-71-001**

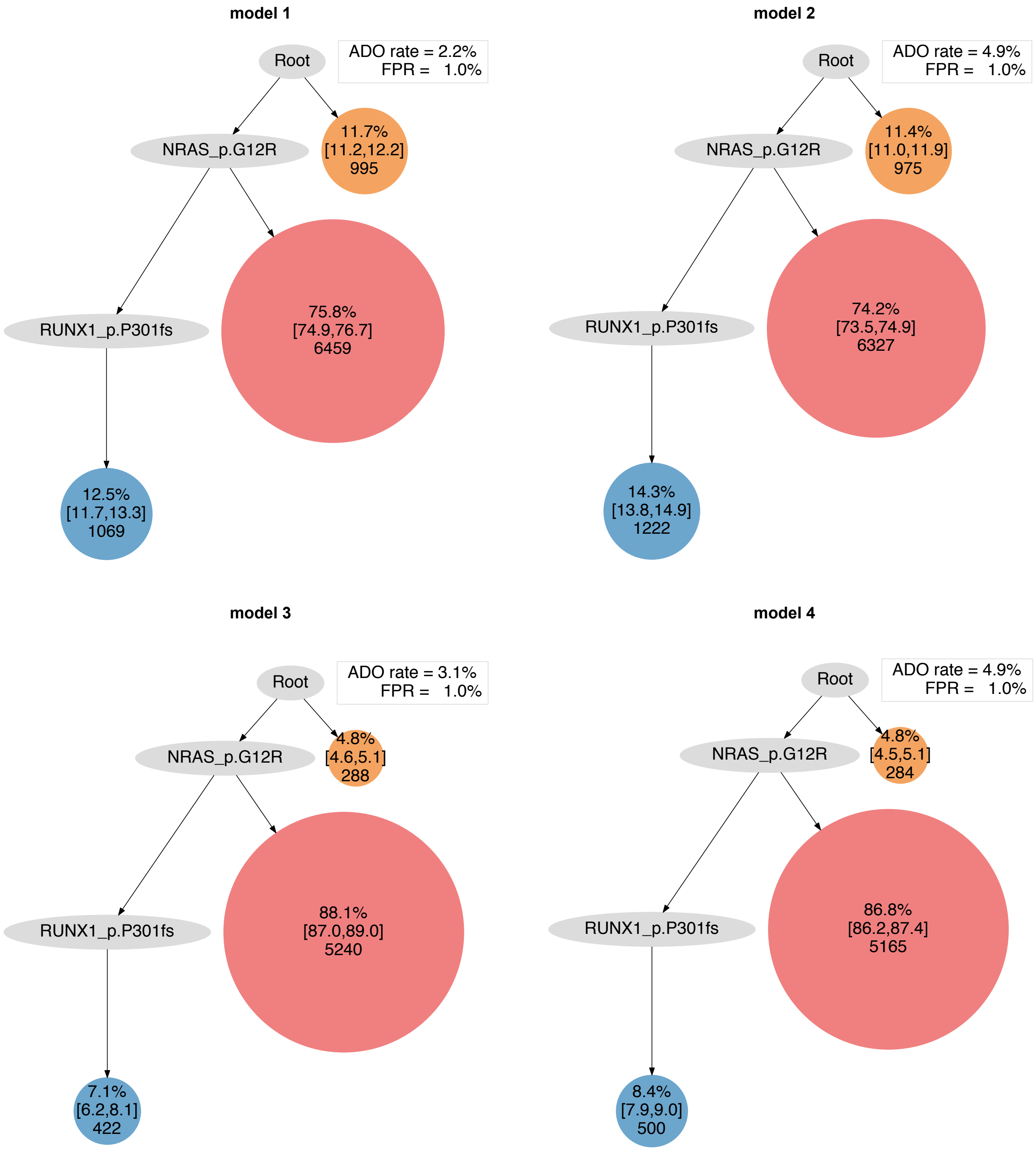
model 1
Root
ADO rate = 2.2%
FPR = 1.0%
NRAS_p.G12R
11.7%
[11.2,12.2]
995
RUNX1_p.P301fs
75.8%
[74.9,76.7]
6459
12.5%
[11.7,13.3]
1069
model 2
Root
ADO rate = 4.9%
FPR = 1.0%
NRAS_p.G12R
11.4%
[11.0,11.9]
975
RUNX1_p.P301fs
74.2%
[73.5,74.9]
6327
14.3%
[13.8,14.9]
1222
model 3
Root
ADO rate = 3.1%
FPR = 1.0%
NRAS_p.G12R
4.8%
[4.6,5.1]
288
RUNX1_p.P301fs
88.1%
[87.0,89.0]
5240
7.1%
[6.2,8.1]
422
model 4
Root
ADO rate = 4.9%
FPR = 1.0%
NRAS_p.G12R
4.8%
[4.5,5.1]
284
RUNX1_p.P301fs
86.8%
[86.2,87.4]
5165
8.4%
[7.9,9.0]
500

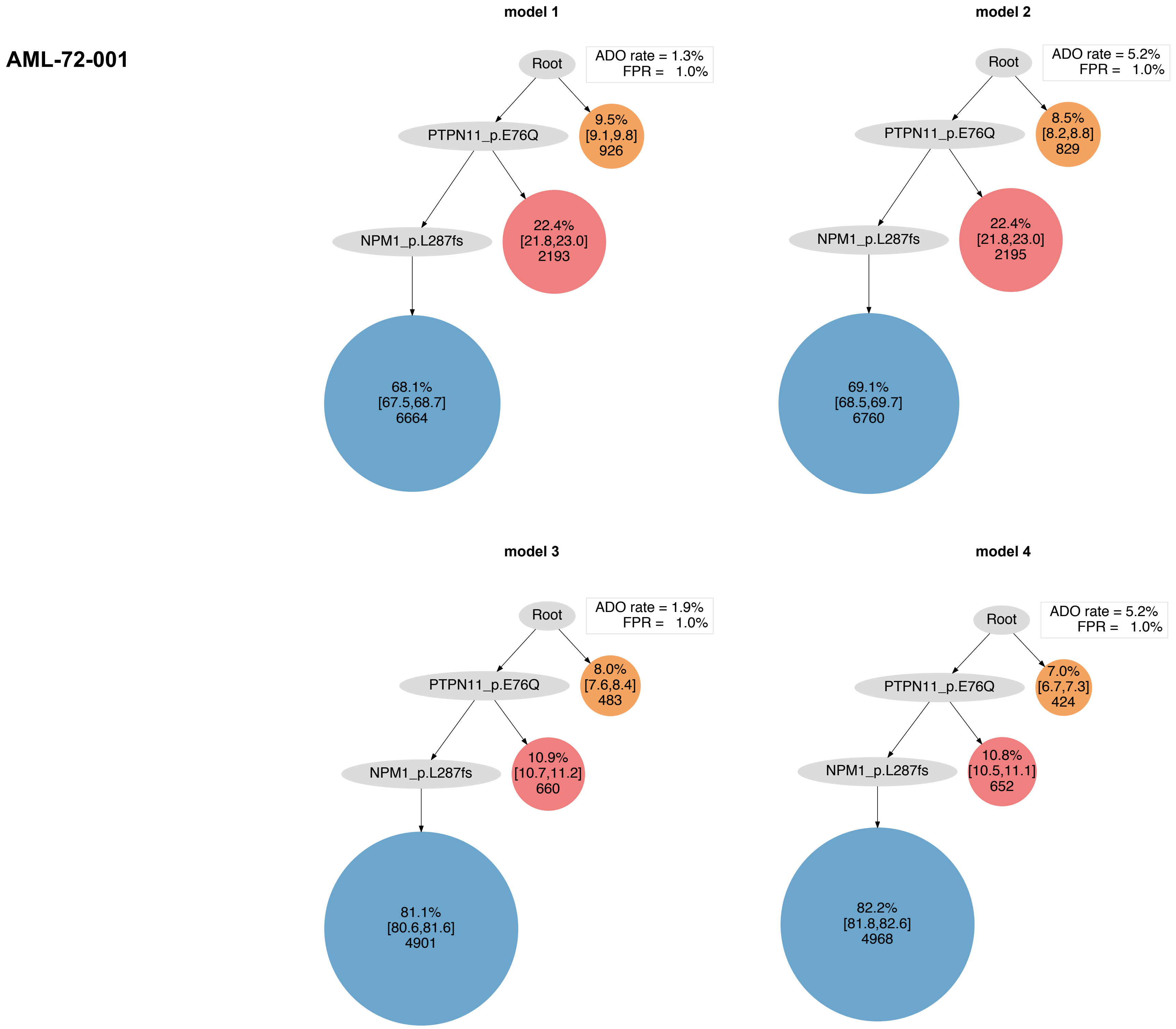
AML-72-001
model 1
Root
ADO rate = 1.3%
FPR = 1.0%
PTPN11_p.E76Q
9.5%
[9.1,9.8]
926
NPM1_p.L287fs
22.4%
[21.8,23.0]
2193
68.1%
[67.5,68.7]
6664
model 2
Root
ADO rate = 5.2%
FPR = 1.0%
PTPN11_p.E76Q
8.5%
[8.2,8.8]
829
NPM1_p.L287fs
22.4%
[21.8,23.0]
2195
69.1%
[68.5,69.7]
6760
model 3
Root
ADO rate = 1.9%
FPR = 1.0%
PTPN11_p.E76Q
8.0%
[7.6,8.4]
483
NPM1_p.L287fs
10.9%
[10.7,11.2]
660
81.1%
[80.6,81.6]
4901
model 4
Root
ADO rate = 5.2%
FPR = 1.0%
PTPN11_p.E76Q
7.0%
[6.7,7.3]
424
NPM1_p.L287fs
10.8%
[10.5,11.1]
652
82.2%
[81.8,82.6]
4968

#### AML-73-001

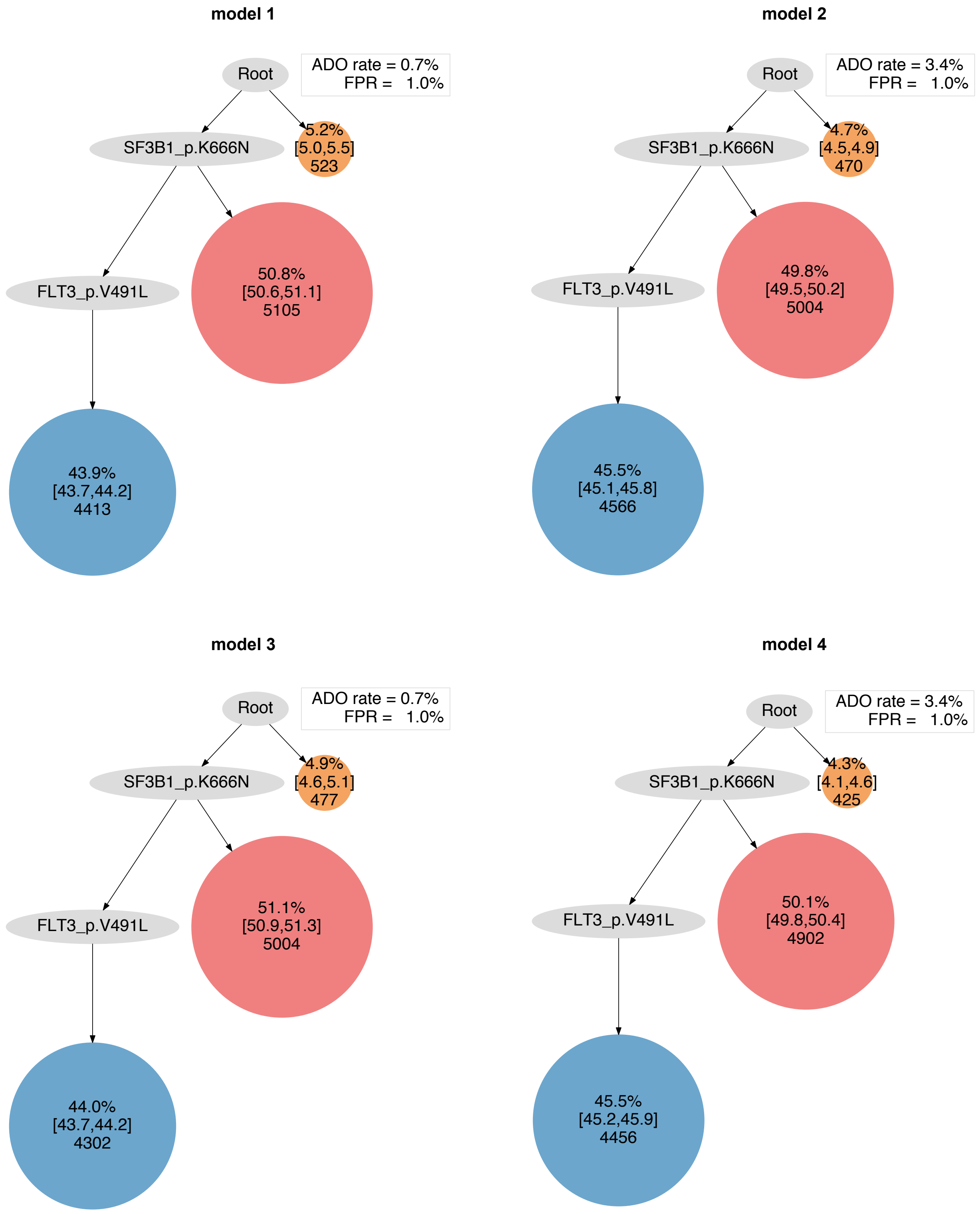
model 1
Root
ADO rate = 0.7%
FPR = 1.0%
SF3B1_p.K666N
5.2%
[5.0,5.5]
523
FLT3_p.V491L
50.8%
[50.6,51.1]
5105
43.9%
[43.7,44.2]
4413
model 2
Root
ADO rate = 3.4%
FPR = 1.0%
SF3B1_p.K666N
4.7%
[4.5,4.9]
470
FLT3_p.V491L
49.8%
[49.5,50.2]
5004
45.5%
[45.1,45.8]
4566
model 3
Root
ADO rate = 0.7%
FPR = 1.0%
SF3B1_p.K666N
4.9%
[4.6,5.1]
477
FLT3_p.V491L
51.1%
[50.9,51.3]
5004
44.0%
[43.7,44.2]
4302
model 4
Root
ADO rate = 3.4%
FPR = 1.0%
SF3B1_p.K666N
4.3%
[4.1,4.6]
425
FLT3_p.V491L
50.1%
[49.8,50.4]
4902
45.5%
[45.2,45.9]
4456

### AML-74-001

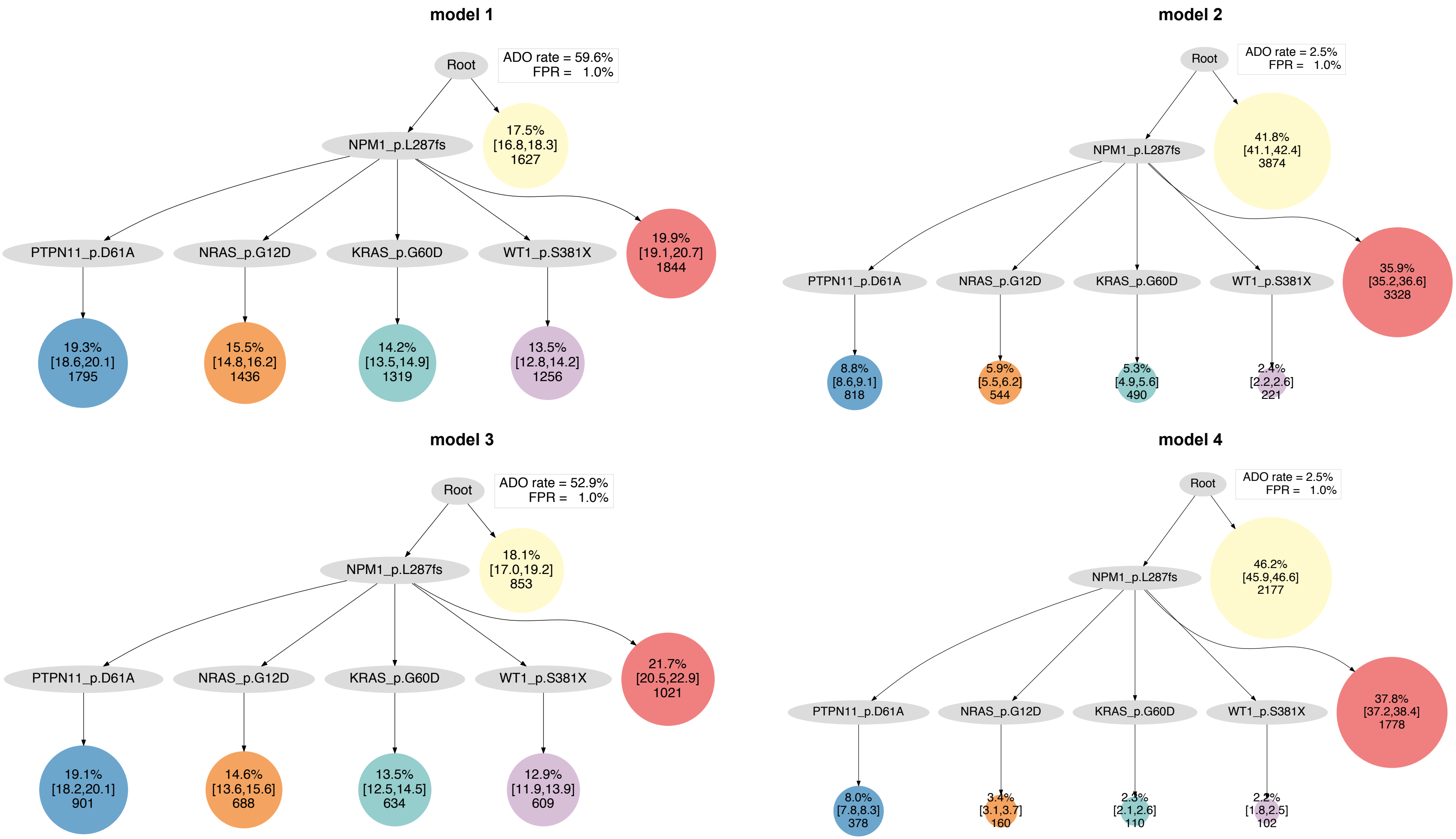
model 1
Root
ADO rate = 59.6%
FPR = 1.0%
NPM1_p.L287fs
17.5%
[16.8,18.3]
1627
PTPN11_p.D61A
NRAS_p.G12D
KRAS_p.G60D
WT1_p.S381X
19.9%
[19.1,20.7]
1844
19.3%
[18.6,20.1]
1795
15.5%
[14.8,16.2]
1436
14.2%
[13.5,14.9]
1319
13.5%
[12.8,14.2]
1256
model 2
Root
ADO rate = 2.5%
FPR = 1.0%
NPM1_p.L287fs
41.8%
[41.1,42.4]
3874
PTPN11_p.D61A
NRAS_p.G12D
KRAS_p.G60D
WT1_p.S381X
35.9%
[35.2,36.6]
3328
8.8%
[8.6,9.1]
818
5.9%
[5.5,6.2]
544
5.3%
[4.9,5.6]
490
2.4%
[2.2,2.6]
221
model 3
Root
ADO rate = 52.9%
FPR = 1.0%
NPM1_p.L287fs
18.1%
[17.0,19.2]
853
PTPN11_p.D61A
NRAS_p.G12D
KRAS_p.G60D
WT1_p.S381X
21.7%
[20.5,22.9]
1021
19.1%
[18.2,20.1]
901
14.6%
[13.6,15.6]
688
13.5%
[12.5,14.5]
634
12.9%
[11.9,13.9]
609
model 4
Root
ADO rate = 2.5%
FPR = 1.0%
NPM1_p.L287fs
46.2%
[45.9,46.6]
2177
PTPN11_p.D61A
NRAS_p.G12D
KRAS_p.G60D
WT1_p.S381X
37.8%
[37.2,38.4]
1778
8.0%
[7.8,8.3]
378
3.4%
[3.1,3.7]
160
2.3%
[2.1,2.6]
110
2.2%
[1.8,2.5]
102

### AML-75-001

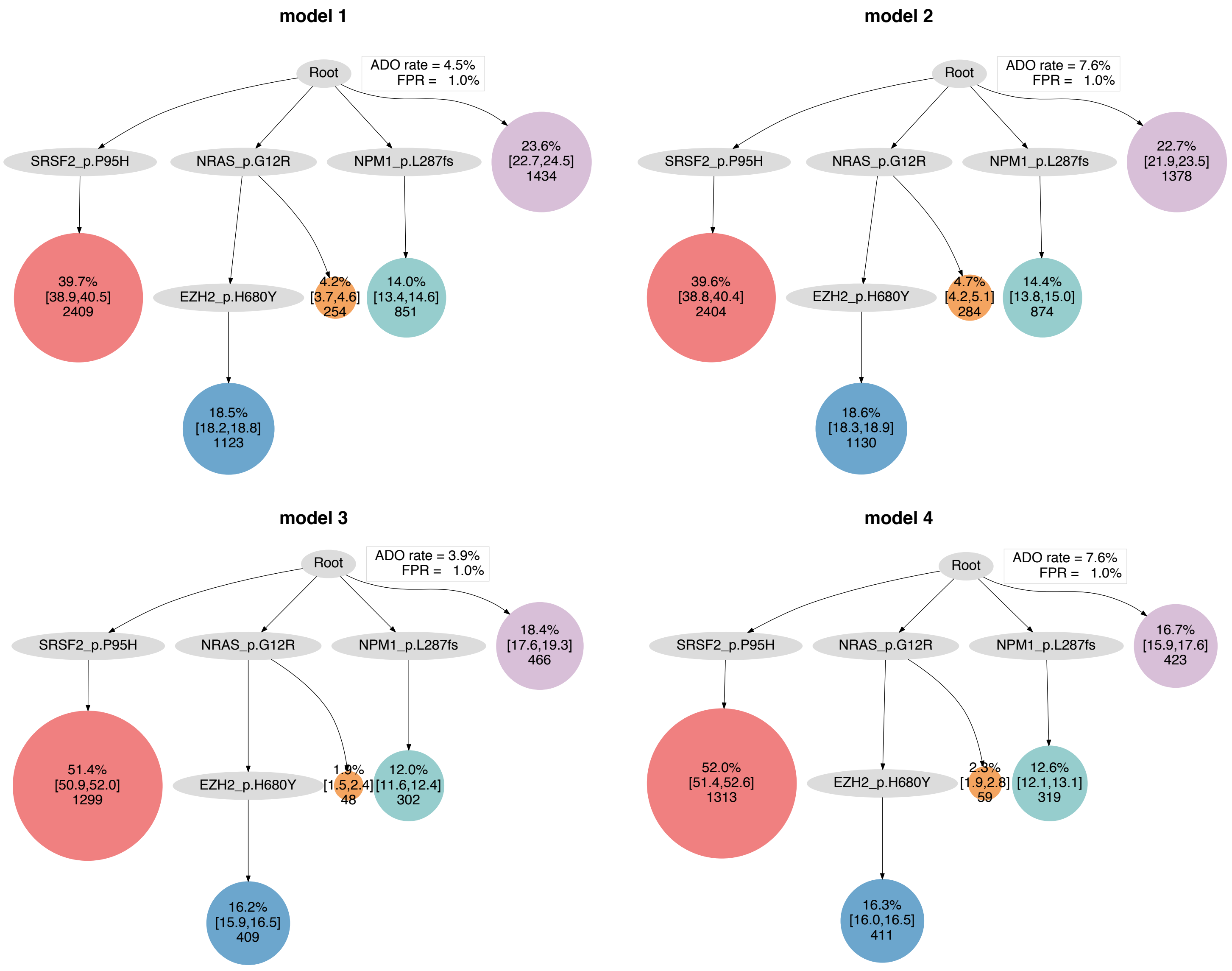
model 1
Root
ADO rate = 4.5%
FPR = 1.0%
SRSF2_p.P95H
NRAS_p.G12R
NPM1_p.L287fs
23.6% [22.7,24.5] 1434
39.7% [38.9,40.5] 2409
EZH2_p.H680Y
4.2% [3.7,4.6] 254
14.0% [13.4,14.6] 851
18.5% [18.2,18.8] 1123
model 2
Root
ADO rate = 7.6%
FPR = 1.0%
SRSF2_p.P95H
NRAS_p.G12R
NPM1_p.L287fs
22.7% [21.9,23.5] 1378
39.6% [38.8,40.4] 2404
EZH2_p.H680Y
4.7% [4.2,5.1] 284
14.4% [13.8,15.0] 874
18.6% [18.3,18.9] 1130
model 3
Root
ADO rate = 3.9%
FPR = 1.0%
SRSF2_p.P95H
NRAS_p.G12R
NPM1_p.L287fs
18.4% [17.6,19.3] 466
51.4% [50.9,52.0] 1299
EZH2_p.H680Y
1.9% [1.5,2.4] 48
12.0% [11.6,12.4] 302
16.2% [15.9,16.5] 409
model 4
Root
ADO rate = 7.6%
FPR = 1.0%
SRSF2_p.P95H
NRAS_p.G12R
NPM1_p.L287fs
16.7% [15.9,17.6] 423
52.0% [51.4,52.6] 1313
EZH2_p.H680Y
2.3% [1.9,2.8] 59
12.6% [12.1,13.1] 319
16.3% [16.0,16.5] 411

### AML-76-001

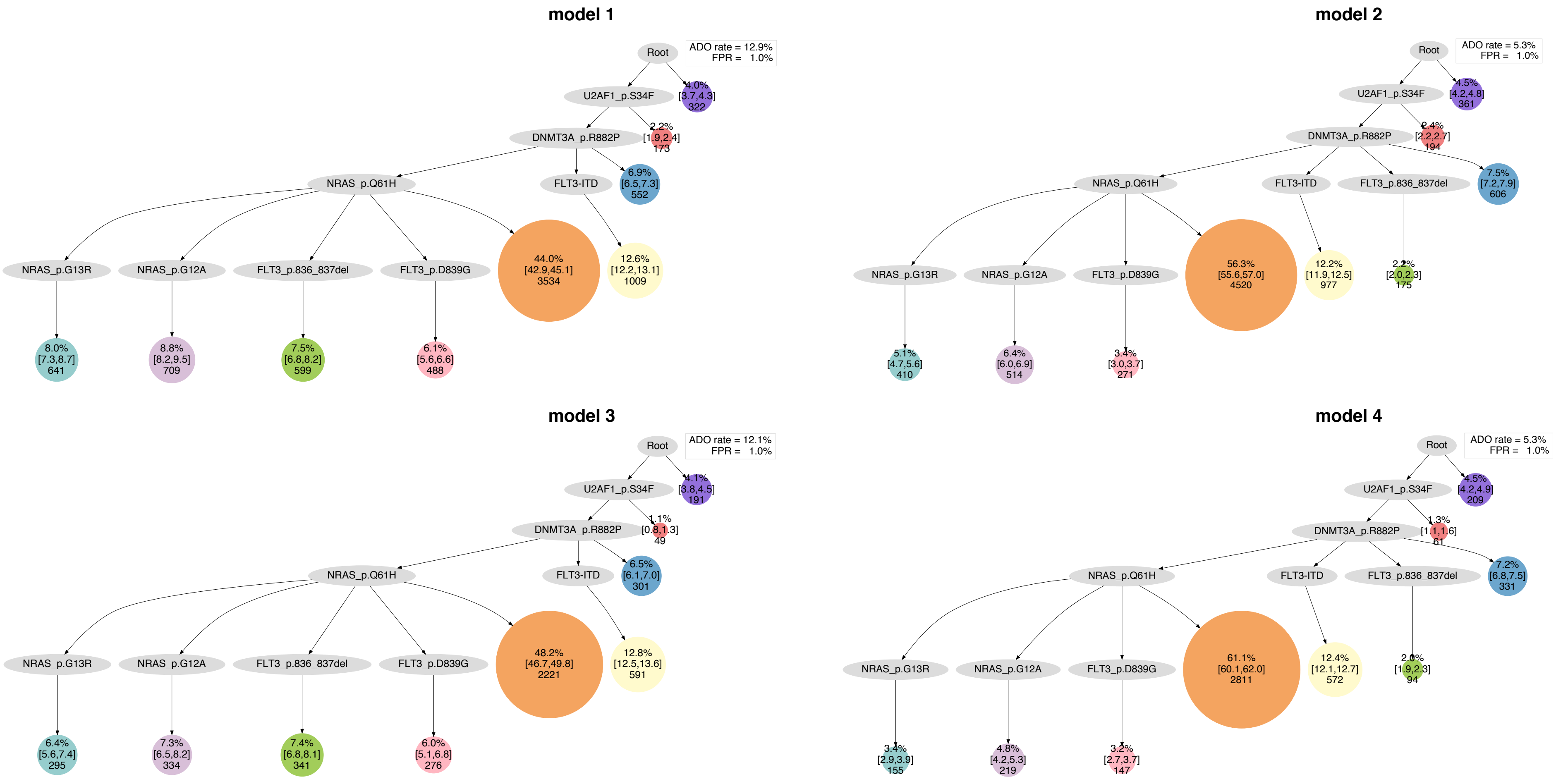
model 1
Root
ADO rate = 12.9%
FPR = 1.0%
U2AF1_p.S34F
4.0% [3.7,4.3] 322
DNMT3A_p.R882P
2.2% [1.9,2.4] 173
NRAS_p.Q61H
FLT3-ITD
6.9% [6.5,7.3] 552
NRAS_p.G13R
NRAS_p.G12A
FLT3_p.836_837del
FLT3_p.D839G
44.0% [42.9,45.1] 3534
12.6% [12.2,13.1] 1009
8.0% [7.3,8.7] 641
8.8% [8.2,9.5] 709
7.5% [6.8,8.2] 599
6.1% [5.6,6.6] 488
model 2
Root
ADO rate = 5.3%
FPR = 1.0%
U2AF1_p.S34F
4.5% [4.2,4.8] 361
DNMT3A_p.R882P
2.4% [2.2,2.7] 194
NRAS_p.Q61H
FLT3-ITD
FLT3_p.836_837del
7.5% [7.2,7.9] 606
NRAS_p.G13R
NRAS_p.G12A
FLT3_p.D839G
56.3% [55.6,57.0] 4520
12.2% [11.9,12.5] 977
2.2% [2.0,2.3] 175
5.1% [4.7,5.6] 410
6.4% [6.0,6.9] 514
3.4% [3.0,3.7] 271
model 3
Root
ADO rate = 12.1%
FPR = 1.0%
U2AF1_p.S34F
4.1% [3.8,4.5] 191
DNMT3A_p.R882P
1.1% [0.8,1.3] 49
NRAS_p.Q61H
FLT3-ITD
6.5% [6.1,7.0] 301
NRAS_p.G13R
NRAS_p.G12A
FLT3_p.836_837del
FLT3_p.D839G
48.2% [46.7,49.8] 2221
12.8% [12.5,13.6] 591
6.4% [5.6,7.4] 295
7.3% [6.5,8.2] 334
7.4% [6.8,8.1] 341
6.0% [5.1,6.8] 276
model 4
Root
ADO rate = 5.3%
FPR = 1.0%
U2AF1_p.S34F
4.5% [4.2,4.9] 209
DNMT3A_p.R882P
1.3% [1.1,1.6] 61
NRAS_p.Q61H
FLT3-ITD
FLT3_p.836_837del
7.2% [6.8,7.5] 331
NRAS_p.G13R
NRAS_p.G12A
FLT3_p.D839G
61.1% [60.1,62.0] 2811
12.4% [12.1,12.7] 572
2.0% [1.9,2.3] 94
3.4% [2.9,3.9] 155
4.8% [4.2,5.3] 219
3.2% [2.7,3.7] 147

### AML-77-001

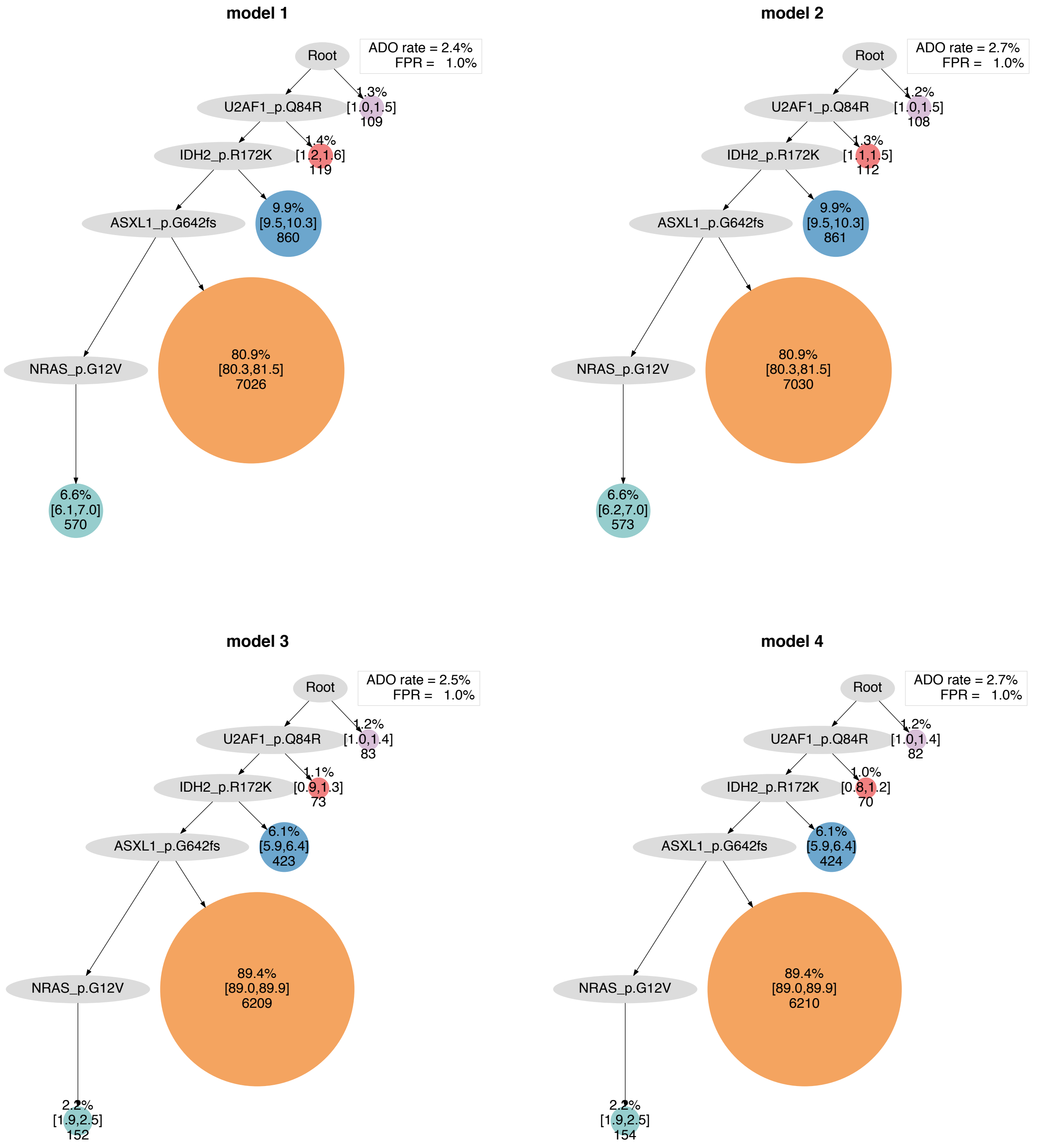

model 1
Root
ADO rate = 2.4%
FPR = 1.0%
U2AF1_p.Q84R
1.3%
[1.0,1.5]
109
IDH2_p.R172K
1.4%
[1.2,1.6]
119
ASXL1_p.G642fs
9.9%
[9.5,10.3]
860
NRAS_p.G12V
80.9%
[80.3,81.5]
7026
6.6%
[6.1,7.0]
570
model 2
Root
ADO rate = 2.7%
FPR = 1.0%
U2AF1_p.Q84R
1.2%
[1.0,1.5]
108
IDH2_p.R172K
1.3%
[1.1,1.5]
112
ASXL1_p.G642fs
9.9%
[9.5,10.3]
861
NRAS_p.G12V
80.9%
[80.3,81.5]
7030
6.6%
[6.2,7.0]
573
model 3
Root
ADO rate = 2.5%
FPR = 1.0%
U2AF1_p.Q84R
1.2%
[1.0,1.4]
83
IDH2_p.R172K
1.1%
[0.9,1.3]
73
ASXL1_p.G642fs
6.1%
[5.9,6.4]
423
NRAS_p.G12V
89.4%
[89.0,89.9]
6209
2.2%
[1.9,2.5]
152
model 4
Root
ADO rate = 2.7%
FPR = 1.0%
U2AF1_p.Q84R
1.2%
[1.0,1.4]
82
IDH2_p.R172K
1.0%
[0.8,1.2]
70
ASXL1_p.G642fs
6.1%
[5.9,6.4]
424
NRAS_p.G12V
89.4%
[89.0,89.9]
6210
2.2%
[1.9,2.5]
154

**Extended Data Fig. 12. Estimation of clonal evolution from single-cell genotype data using the SCITE algorithm.** Phylogenetic trees visualizing the distinct patterns of clonal evolution are shown. Four phylogenetic models are shown for each patient based on the different combinations of parameters: model 1) use all cells including missing genotype information with 1% false positive rate (FPR) and SCITE inferred false negative rate (FNR)/allele dropout (ADO) rate, model 2) use all cells including missing genotype information with 1% FPR and platform provided FNR, model 3) use only cells with full genotype information with 1% FPR and SCITE inferred FNR, and model 4) use only cells with full genotype information with 1% FPR and platform provided FNR.

**Extended Data Fig. 13**

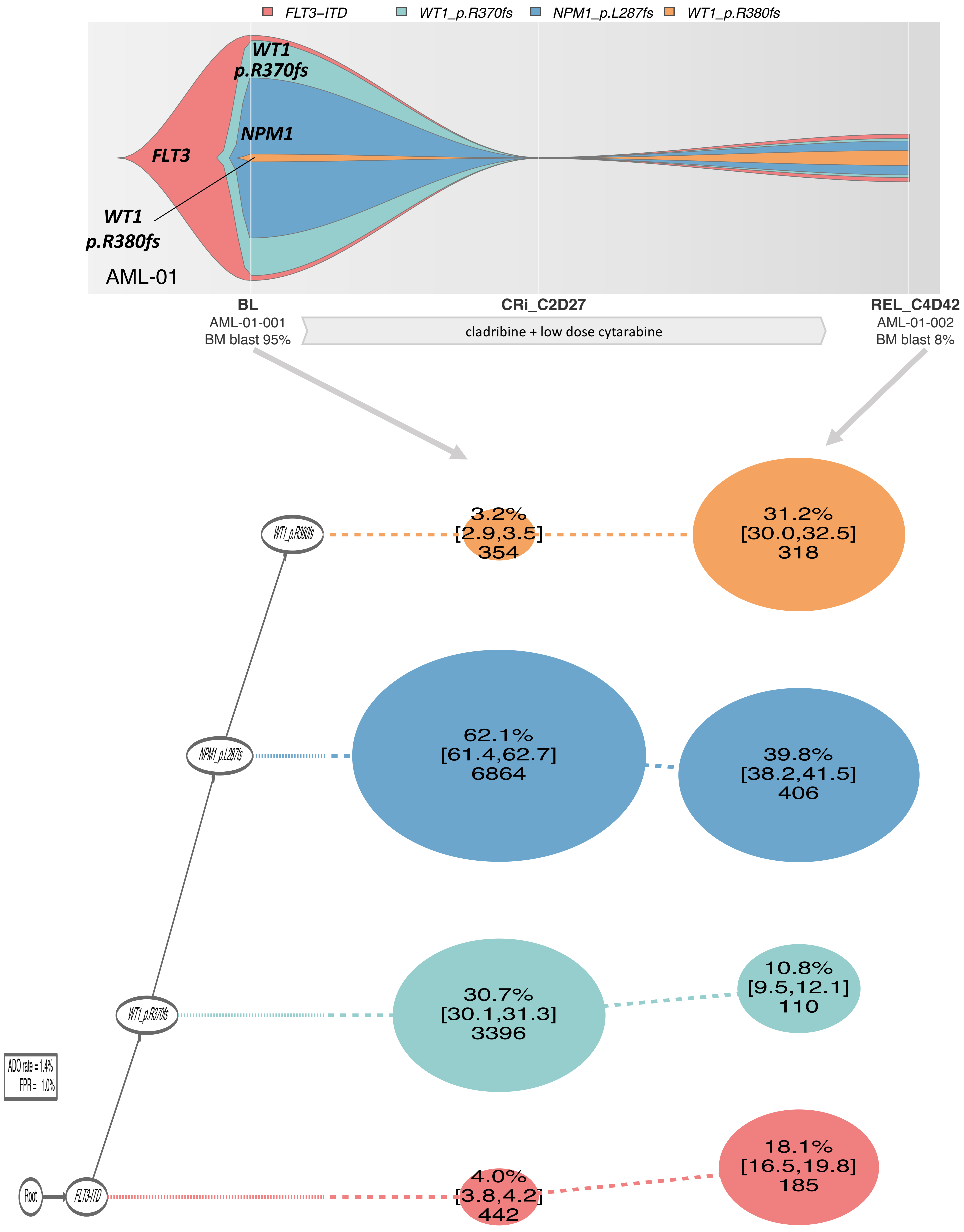

**Extended Data Fig. 13. Patterns of clonal evolution at single-cell resolution.** Fish plots and phylogeny trees show the inferred patterns of clonal evolution based on the mutation data obtained from longitudinal single-cell DNA sequencing.

**AML-01**

A 66-year-old man with a history of radiation treatment for prostate cancer was diagnosed with *FLT3*-positive AML (AML-01-001). He was treated with cladribine plus low-dose cytarabine alternating with decitabine and achieved complete remission with incomplete platelet recovery after 2 cycles. He received additional 2 cycles of consolidation therapy, but his disease relapsed and was positive for *FLT3*-ITD (AML-01-002). He was then treated with azacitidine and sorafenib. Although he achieved complete remission with incomplete platelet recovery after 1 cycle, he developed *Mucor* cellulitis. He died of multiorgan failure approximately 9 months after the diagnosis of therapy-related AML. The clonal architecture was similar in the pretreatment and relapse samples.

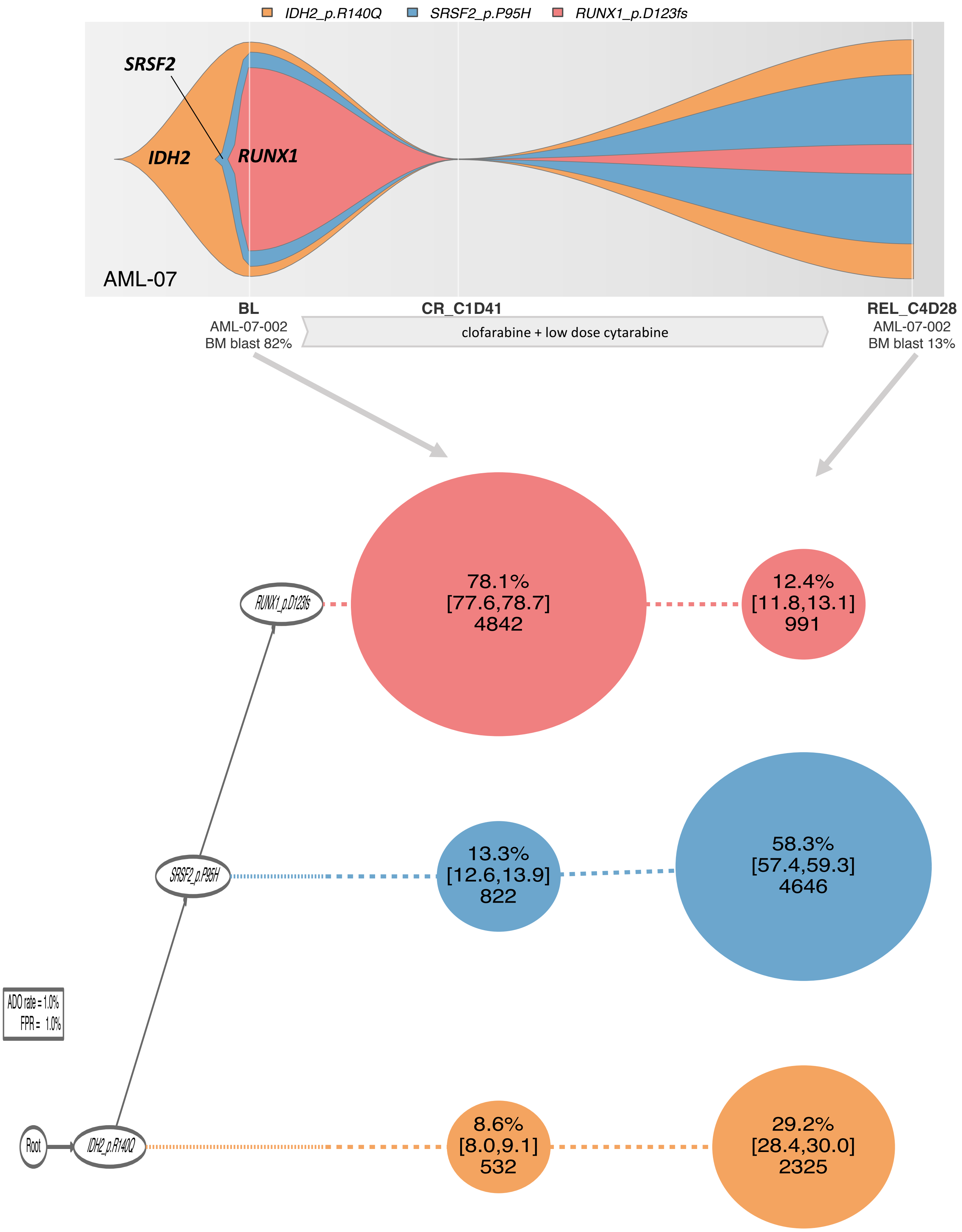
IDH2_p.R140Q
SRSF2_p.P95H
RUNX1_p.D123fs
SRSF2
IDH2
RUNX1
AML-07
BL
AML-07-002
BM blast 82%
CR_C1D41
clofarabine + low dose cytarabine
REL_C4D28
AML-07-002
BM blast 13%
RUNX1_p.D123fs
78.1%
[77.6,78.7]
4842
12.4%
[11.8,13.1]
991
SRSF2_p.P95H
13.3%
[12.6,13.9]
822
58.3%
[57.4,59.3]
4646
ADO rate = 1.0%
FPR = 1.0%
Root
IDH2_p.R140Q
8.6%
[8.0,9.1]
532
29.2%
[28.4,30.0]
2325

**AML-07**

A 75-year old man was diagnosed with pure erythroid leukemia (AML-M6B) (AML-07-001). He received induction chemotherapy with clofarabine plus low-dose cytarabine and achieved complete remission after 1 cycle. He was found to have a relapse after receiving 3 cycles of consolidation therapy (AML-07-002). Five cycles of salvage chemotherapy consisting of fludarabine and cytarabine failed to induce remission. He was then started on clofarabine, azacitidine, and low-dose cytarabine and achieved remission but relapsed approximately 2 years later. The patient was given 5 cycles of azacitidine with no response. He was switched to clofarabine followed by cytarabine and initially responded well, but his disease later progressed. He died of unknown causes approximately 6 years after the diagnosis of AML. A subclone with *RUNX1* mutations, which was estimated to have occurred at the latest evolutionary stage, substantially shrank at relapse, whereas the 2 founder mutations (*IDH2* and *SRSF2*) were persistent, showing the differential chemosensitivity of mutations.

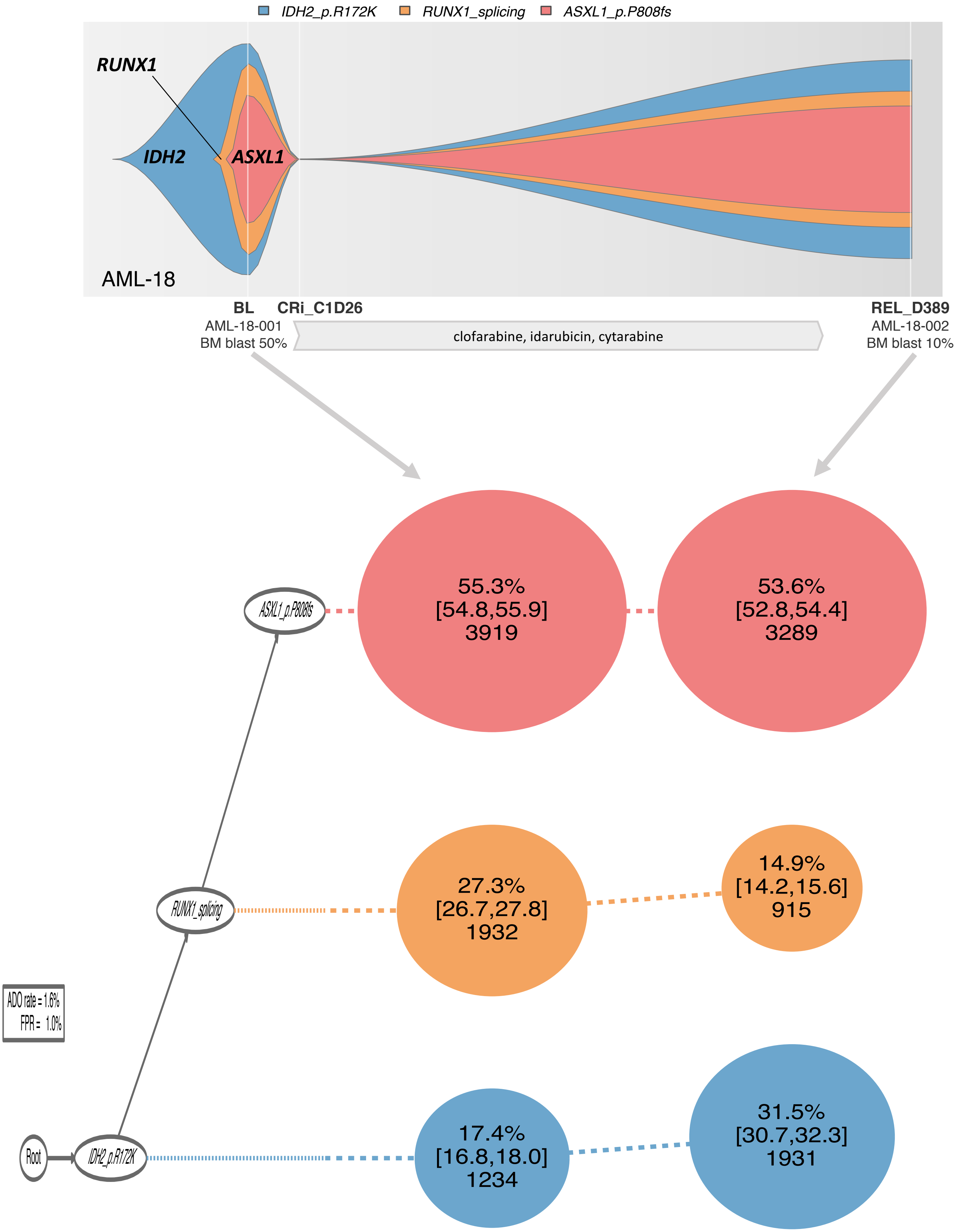

IDH2_p.R172K
RUNX1_splicing
ASXL1_p.P808fs
RUNX1
IDH2
ASXL1
AML-18
BL
AML-18-001
BM blast 50%
CRi_C1D26
clofarabine, idarubicin, cytarabine
REL_D389
AML-18-002
BM blast 10%
55.3%
[54.8,55.9]
3919
53.6%
[52.8,54.4]
3289
27.3%
[26.7,27.8]
1932
14.9%
[14.2,15.6]
915
17.4%
[16.8,18.0]
1234
31.5%
[30.7,32.3]
1931
ASXL1_p.P808fs
RUNX1_splicing
IDH2_p.R172K
Root
ADO rate = 1.6%
FPR = 1.0%

**AML-18**

A 30-year-old-man was diagnosed with AML with maturation (AML-M2) (AML-18-001). He was treated with induction chemotherapy with clofarabine, idarubicin, and cytarabine (CIA). He achieved remission and completed 6 cycles of consolidation therapy, but had a relapse after approximately 1 year of remission (AML-18-002). The relapsed disease was refractory to multiple regimens, including vosaroxin/placebo plus cytarabine, guadecitabine, and CIA. He underwent an allogeneic stem cell transplant from a matched unrelated donor, and achieved complete remission. He received azacitidine maintenance therapy and a donor leukocyte infusion, but had a relapse approximately 6 months after stem cell transplantation. His disease was refractory to multiple regimens, including evofosfamide, buparlisib, decitabine, AZD-1208, erlotinib, PRI724, decitabine plus cytarabine, IGN523, fludarabine plus cytarabine, enasidenib, uprosertib plus trametinib, and APTO253. He developed CNS leukemia and myeloid sarcoma and died approximately 4 years after the diagnosis of AML. The clonal architecture was similar in his pretreatment and relapse samples.

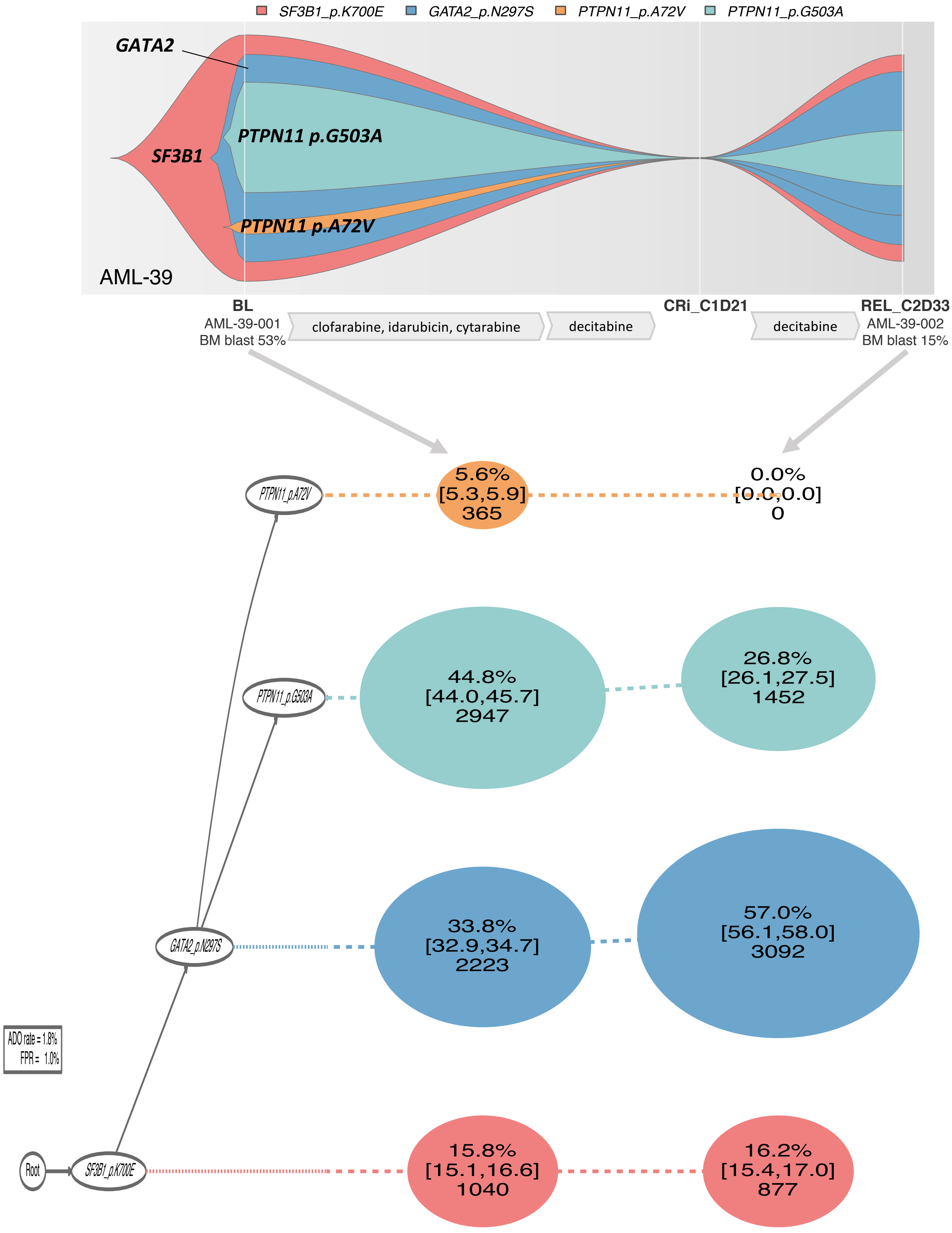

SF3B1_p.K700E
GATA2_p.N297S
PTPN11_p.A72V
PTPN11_p.G503A
GATA2
SF3B1
PTPN11 p.G503A
PTPN11 p.A72V
AML-39
BL
AML-39-001
BM blast 53%
clofarabine, idarubicin, cytarabine
decitabine
CRi_C1D21
decitabine
REL_C2D33
AML-39-002
BM blast 15%
PTPN11_p.A72V
5.6%
[5.3,5.9]
365
0.0%
[0.0,0.0]
0
PTPN11_p.G503A
44.8%
[44.0,45.7]
2947
26.8%
[26.1,27.5]
1452
GATA2_p.N297S
33.8%
[32.9,34.7]
2223
57.0%
[56.1,58.0]
3092
ADO rate = 1.8%
FPR = 1.0%
Root
SF3B1_p.K700E
15.8%
[15.1,16.6]
1040
16.2%
[15.4,17.0]
877

**AML-39**

A 55-year-old man with a history of myelodysplastic syndrome experienced a progression into secondary AML (AML-39-001). His disease was refractory to induction chemotherapy with clofarabine, idarubicin, and cytarabine, and he was switched to decitabine. He achieved complete remission with incomplete platelet and neutrophil recovery, but relapsed after the second cycle (AML-39-002). He was treated with cyclophosphamide, etoposide, carboplatin, and cytarabine, but died approximately 6 months after the diagnosis of AML. The 2 *PTPN11* mutations comprised independent branching subclones that shared *SF3B1* and *GATA2* mutations. The clone with the *PTPN11* p.A72V mutation was cleared at relapse, whereas the *PTPN11* p.G503A mutation persisted, illustrating the selection of *PTPN11* p.G503A-mutated clone over *PTPN11* p.A72V-mutated clone under the selective pressure of treatment.

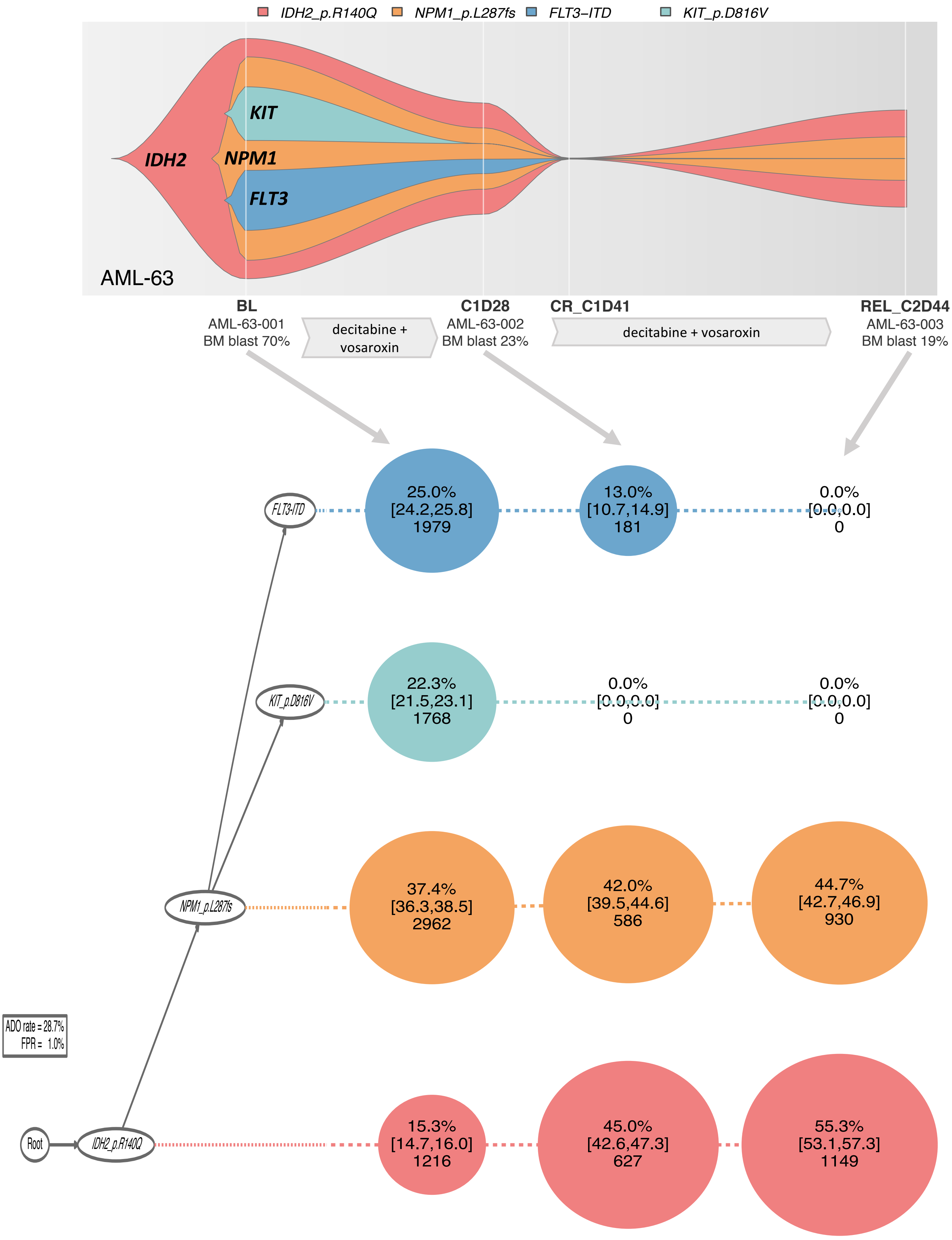

IDH2_p.R140Q
NPM1_p.L287fs
FLT3–ITD
KIT_p.D816V
KIT
IDH2
NPM1
FLT3
AML-63
BL
AML-63-001
BM blast 70%
decitabine + vosaroxin
C1D28
AML-63-002
BM blast 23%
CR_C1D41
decitabine + vosaroxin
REL_C2D44
AML-63-003
BM blast 19%
FLT3-ITD
25.0% [24.2,25.8] 1979
13.0% [10.7,14.9] 181
0.0% [0.0,0.0] 0
KIT_p.D816V
22.3% [21.5,23.1] 1768
0.0% [0.0,0.0] 0
0.0% [0.0,0.0] 0
NPM1_p.L287fs
37.4% [36.3,38.5] 2962
42.0% [39.5,44.6] 586
44.7% [42.7,46.9] 930
ADO rate = 28.7%
FPR = 1.0%
Root
IDH2_p.R140Q
15.3% [14.7,16.0] 1216
45.0% [42.6,47.3] 627
55.3% [53.1,57.3] 1149

**AML-63**

A 65-year-old woman was diagnosed with AML with *FLT3*-ITD (AML-63-001). She was treated with induction therapy consisting of decitabine and vosaroxin (AML-63-002). She achieved complete remission after 1 cycle. After an additional cycle of decitabine and vosaroxin, she was found to have relapsed (AML-63-003). She continued to receive decitabine and vosaroxine, and achieved complete remission with incomplete neutrophil recovery. After a total of 6 courses of decitabine and vosaroxin therapy, she underwent an allogeneic stem cell transplant from a matched related donor. She remains in remission 3 years after the transplant. The clone with *IDH2* and *NPM1* mutations acquired *KIT* or *FLT3* mutations in parallel. Decitabine and vosaroxin therapy suppressed the *KIT*- and *FLT3*- mutated clones, whereas *IDH2* and *NPM1* mutations survived the therapy, illustrating the differential sensitivity of mutations to the therapy with hypomethylating agent.

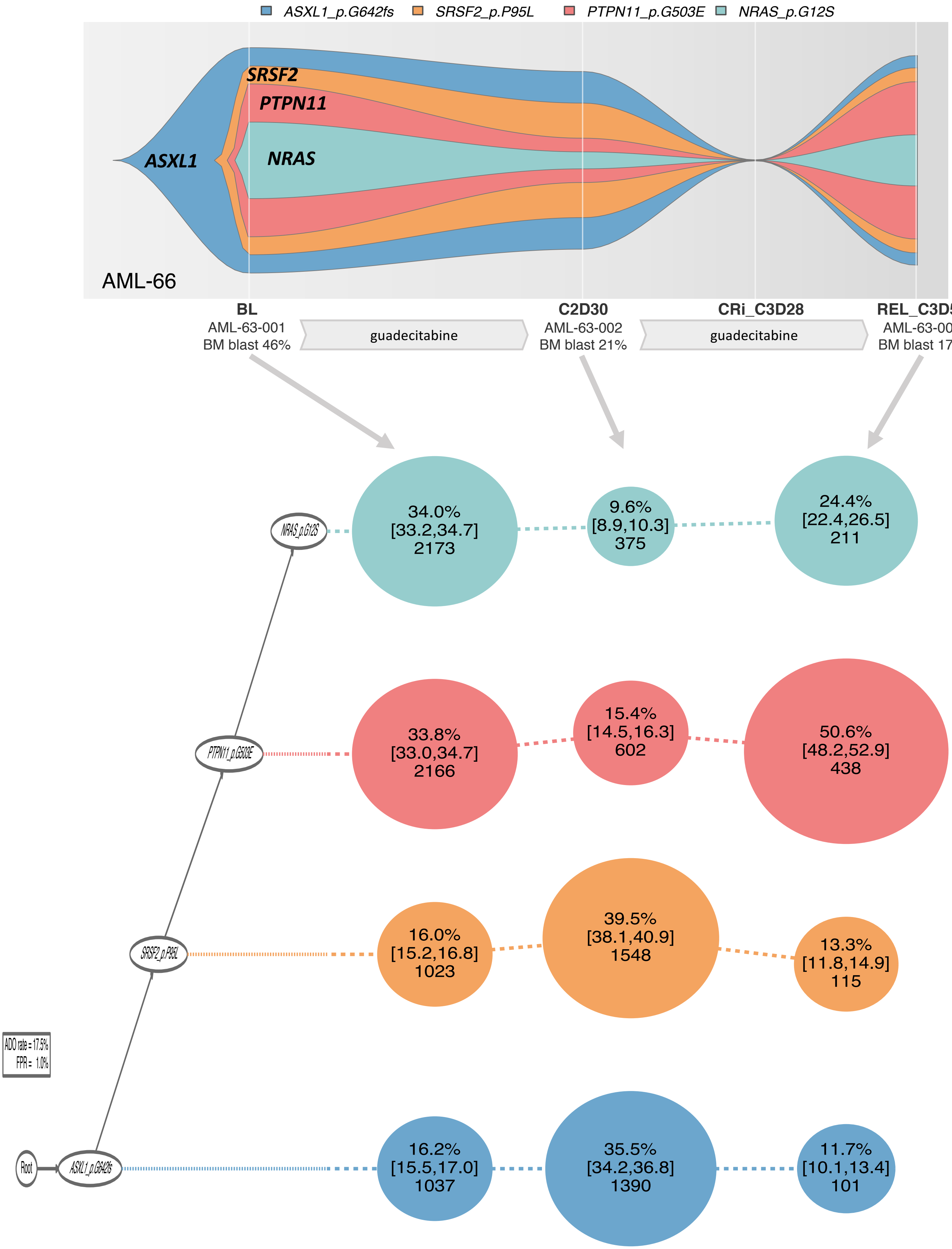

ASXL1_p.G642fs
SRSF2_p.P95L
PTPN11_p.G503E
NRAS_p.G12S
SRSF2
PTPN11
ASXL1
NRAS
AML-66
BL
AML-63-001
BM blast 46%
guadecitabine
C2D30
AML-63-002
BM blast 21%
guadecitabine
CRi_C3D28
REL_C3D55
AML-63-003
BM blast 17%
34.0%
[33.2,34.7]
2173
9.6%
[8.9,10.3]
375
24.4%
[22.4,26.5]
211
33.8%
[33.0,34.7]
2166
15.4%
[14.5,16.3]
602
50.6%
[48.2,52.9]
438
16.0%
[15.2,16.8]
1023
39.5%
[38.1,40.9]
1548
13.3%
[11.8,14.9]
115
16.2%
[15.5,17.0]
1037
35.5%
[34.2,36.8]
1390
11.7%
[10.1,13.4]
101
NRAS_p.G12S
PTPN11_p.G503E
SRSF2_p.P95L
ASXL1_p.G642fs
Root
ADO rate = 17.5%
FPR = 1.0%

**AML-66**

A 70-year-old-man was diagnosed with acute myelomonocytic leukemia (AML-M4) (AML-63-001). He was treated with guadecitabine (AML-63-002) and achieved complete remission with incomplete platelet and neutrophil recovery after 3 cycles, but relapsed after approximately 1 month (AML-63-003). The patient's disease was refractory to salvage therapies, including azacitidine plus nivolumab and venetoclax plus idasanutlin, and he died approximately 1 year after the diagnosis of AML. The clonal architecture was unchanged from baseline to post-treatment with guadecitabine.

**AML-09 (supplementary case description for Fig. 4a)**

A 74 year-old-man who had undergone radiation therapy for prostate cancer was diagnosed with therapy-related AML (AML-09-001). He was found to have *FLT3*-ITD and kinase domain (D835) mutations and was started on induction therapy consisting of azacitidine and sorafenib. He achieved complete remission, but his disease recurred after 9 cycles (AML-09-002). The patient was started on sorafenib and was lost to follow-up. The *FLT3*-ITD mutation that was found in the majority of cells at diagnosis and small independent branched subclones with *FLT3* p.D835E or *KRAS* mutations were cleared after azacitidine plus sorafenib therapy, whereas a small subclone with an *FLT3* p.D835Y mutation significantly expanded and acquired additional mutation in *WT1* at the time of relapse. The evolution pattern of the different *FLT3* variants was in accordance with the differential sensitivity of various *FLT3* mutations to sorafenib.

**AML-21 (supplementary case description for Fig. 4b)**

A 56-year-old woman was diagnosed with acute myelomonocytic leukemia (AML-M4) (AML-21-001). She was treated with induction chemotherapy with clofarabine, idarubicin, and cytarabine, followed by 4 cycles of consolidation therapy. After approximately 7 months of remission, the patient was found to have a relapse (AML-21-002). She was started on vosaroxin/placebo plus cytarabine, followed by fludarabine plus cytarabine, and died approximately after 1 year after the diagnosis of AML. The *NRAS*-mutated clone that was presented as the dominant clone at diagnosis substantially shrank at relapse, while the 2 mutations (*WT1* p.A382fs and *NPM1*) that are estimated to have been acquired prior to the *NRAS* mutation persisted, suggesting the better chemosensitivity of the late-occurring *NRAS* mutation. *FLT3*-ITD and a secondary *WT1* mutation (p.S381fs) were undetectable at diagnosis and acquired at relapse.

**AML-38 (supplementary case description for Fig. 4c)**

A 58-year-old-man with refractory AML, who had been treated with azacitidine and decitabine at an outside institution, was referred to MD Anderson Cancer Center. The patient was found to have *FLT3*-ITD and was started on cytarabine and quizartinib. He achieved complete remission with incomplete platelet and neutrophil recovery after 1 cycle. The patient was found to have recurrent disease prior to the start of the second cycle (AML-38-001), which had been delayed because the patient experienced prolonged myelosuppression. He received a total of 4 cycles of cytarabine and quizartinib therapy (AML-38-002, AML-38-003), but his disease was refractory. He received 2 cycles of chemotherapy consisting of fludarabine, cytarabine, and sorafenib, which markedly reduced the bone marrow blast percentage, from 53% to 7%. He then underwent an allogeneic stem cell transplant from a matched unrelated donor. The patient died of pneumonia approximately 1 year after transplant, even though his AML remained in remission. The founder clone with an *NPM1* mutation independently acquired *IDH1* and *IDH2* mutations. The *NPM1/IDH1*-mutated clone acquired *PTPN11* p.D61H and *KRAS* mutations in a parallel manner, whereas the *NPM1/IDH2*-mutated clone acquired *FLT3*-ITD, *PTPN11* p.A72G, and 2 *NRAS* mutations. After cytarabine and quizartinib therapy, *FLT3*-ITD-mutated clone became undetectable, whereas the remaining clones persisted or expanded. The pattern of mutation phylogeny suggests that there is a preferential order of mutation acquisition (*NPM1/IDH/RTK-RAS* pathway alteration).

**AML-04 (supplementary case description for Fig. 4d)**

A 76-year-old man who had undergone brachytherapy and external-beam radiotherapy for prostate cancer presented with secondary AML arising from essential thrombocythemia, which had been refractory to decitabine and ruxolitinib (AML-04-001). He was found to have *FLT3*-ITD mutation and was treated with azacitidine plus quizartinib. His bone marrow blast percentage markedly decreased, from 40% to 11%, after 2 cycles, but no further response to the treatment was observed after 7 cycles (AML-04-002). The patient was then treated with 3 cycles of crenolanib, but the disease remained refractory (AML-04-003). The patient began azacitidine plus sorafenib therapy but died of unknown causes after approximately 5 months. The *FLT3*-ITD–mutated clone that was the main clone at the first time point substantially shrank after treatment with azacitidine plus quizartinib, whereas the *NRAS-IDH1* p.R132C-mutated clones and *IDH1* p.R132S-mutated clone emerged and expanded. The *FLT3*-ITD mutated clones further shrank after crenolanib therapy, whereas the remaining clones persisted or expanded. Two founder mutations (*SF3B1* and *SRSF2*) survived both courses of therapy.

#### Extended Data Fig. 14

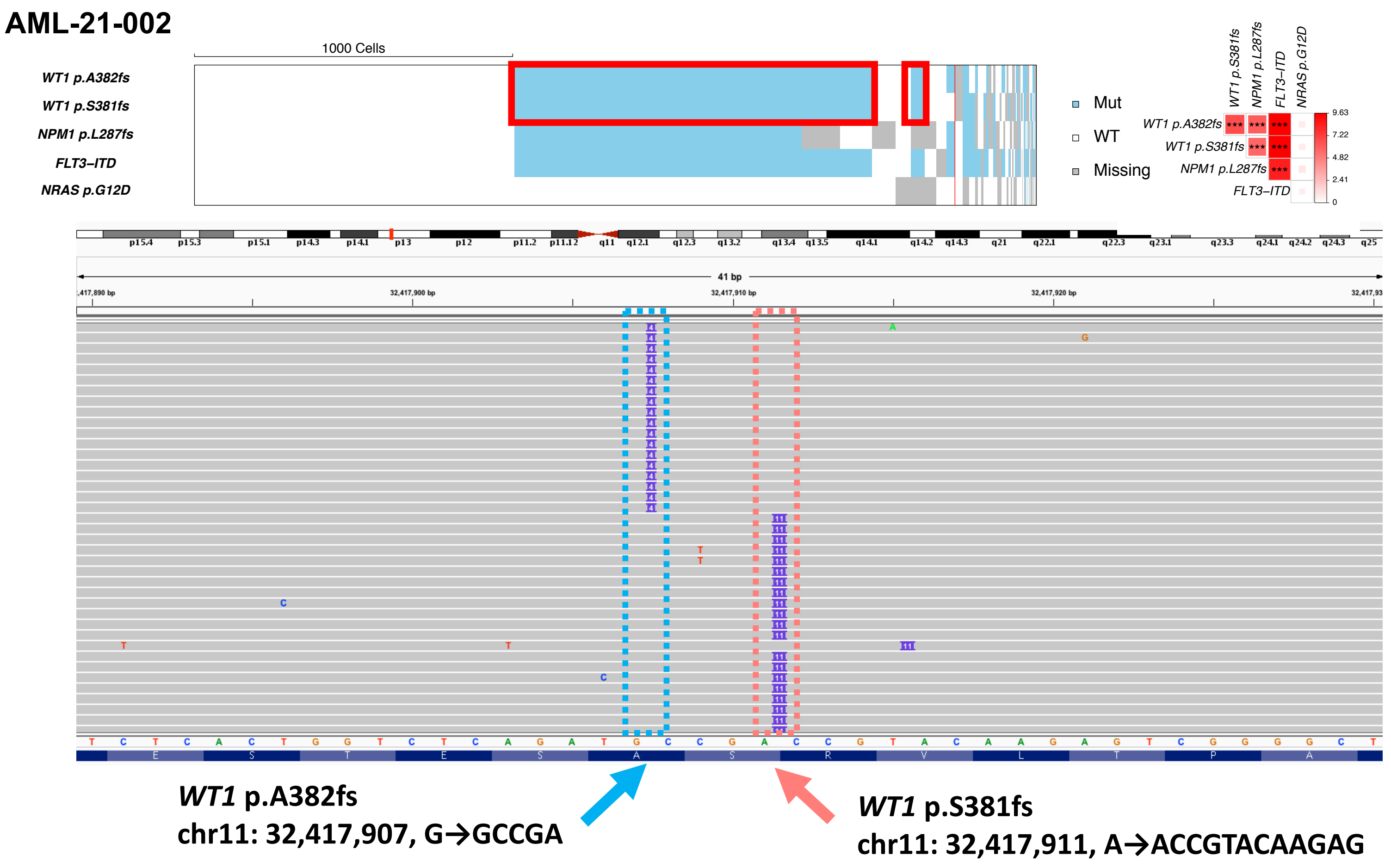


**Extended Data Fig. 14. Allelic-level exclusivity of the two driver mutations in *WT1* that were co-occurring within the same cells.** Individual heat map and Integrative Genomics Viewer (IGV) track showing the cellular-level co-occurrence and allelic-level exclusivity of the 2 *WT1* mutations that were detected in AML-21-002. The cellular populations where the 2 *WT1* mutations were co-occurring are highlighted with red rectangles in the heat map. The region shown in IGV track is covered by one amplicon (*WT1_3*), and each sequencing read (grey) represents each allele. The sequencing reads with *WT1* p.A382fs mutations did not harbor *WT1* p.S381fs mutations, and vice versa, indicating that these mutations did not co-occur on the same alleles, and the two mutations presented as biallelic mutations.
