## Supplemental Table for "Clonal Evolution of Acute Myeloid Leukemia Revealed by High-Throughput Single-Cell Genomics"

**Contents**

**Supplemental Methods**
**Supplementary Table 1**
**Supplementary Table 2**
**Supplementary Table 3**
**Supplementary Table 4**
**Supplementary Table 5**
**Supplementary Table 6**
**Extended Data Figure 1**
**Extended Data Figure 2**
**Extended Data Figure 3**
**Extended Data Figure 4**
**Extended Data Figure 5**
**Extended Data Figure 6**
**Extended Data Figure 7**
**Extended Data Figure 8**
**Extended Data Figure 9**
**Extended Data Figure 10**
**Extended Data Figure 11**
**Extended Data Figure 12**
**Extended Data Figure 13**
**Extended Data Figure 14**

**Supplemental References**

### Supplemental Methods

#### Variant calling and genotyping using single-cell DNA sequencing data

Fastq files generated by an Illumina MiSeq sequencer were processed using the Tapestri Analysis Pipeline. Adapter sequences were trimmed from the sequenced reads using Cutadapt [1,2]. Reads that were too short were discarded, and the Tapestri Barcode structures were extracted from the reads. The reads were then mapped to human genome version hg19 using the BWA-MEM algorithm [3 4]. The extracted barcodes on the mapped reads were error-corrected against a whitelist of known barcodes using a hamming distance approach. Reads lacking an insert sequence between gene-specific primers or mapped to off-target loci were discarded. The barcodes were identified as cells using a 2-step process. First, amplification patterns of all amplicons were reduced to principal components to separate barcodes with high read-load (putative cells) from barcodes with low read-load (noise). Then a coverage parameter of at least 10 reads for at least 60% of the amplicons was used for each barcode to discard putative cells with low data completeness from cells with high data completeness.

The cells were genotyped with the Genome Analysis Toolkit [5] using a joint calling approach that followed GATK Best Practices recommendations [6,7]. Each cell was haplotyped in reference confidence mode to enable per-base pair (bp) confidence estimates for a site's being strictly homozygous (reference). The per-bp resolution was maintained while merging the genomic-VCFs (gVCFs) for all cells using GATK's CombineGVCFs tool. Finally, joint genotyping was performed for all cells using GATK's GenotypeGVCFs tool. Loci found to be nonvariant were maintained in the final output. Genotyping parameters were optimized for high sensitivity: a maximum of 2 alternate alleles were reported for each site, the minimum base quality for variant calling was set at 10, and the heterozygosity value was set at 0.001.

Internal tandem duplications in the *FLT3* gene were identified using a custom genotyping method. The designed panel had 2 amplicons targeting exon 14 and exon 15 in *FLT3*. We looked for insertions in the *FLT3* amplicons and mapped them to the locus. If the read did not go through the insertion, our mapper soft-clipped it at the end. For each cell, we scanned for these soft-clips and insertions; all insertions and clippings were considered as possible insertions. If the total number of reads was greater than a cutoff (10), and the number and the ratio of non-reference reads were greater than a cutoff (4 and 0.1 respectively), the cell was considered to have a non-reference allele. If the ratio of non-reference reads was greater than a cutoff (0.9), a homozygous event was called; otherwise it was considered a heterozygous event. If the cell had enough total reads but not enough alternate reads, it was considered a homozygous reference. Otherwise, it was reported as "no call." Multiallelic variants were decomposed into biallelic variants and then normalized to ensure that each VCF entry was left-aligned and parsimonious [8]. Blacklisted loci were filtered out, and all loci with a quality value <1000 were tagged for downstream processing. The positions that passed our filtering criteria were called as variants. The genotypes and the cell matrix were converted into an open-source loom format [9], which allows efficient storage, data retrieval, and sharing of large omics data sets.

Allele dropout (ADO) was calculated based on the fraction of cells that were genotyped as not heterozygous at a specific locus that is expected to be heterozygous in the general population. Non-heterozygous genotype is either homozygous reference genotype when the mutant allele failed to amplify, or homozygous genotype of the alternate allele when a reference allele failed to amplify. ADO rate was defined based on the variant allele frequency, which was calculated as (number of cells with non-heterozygous genotype at known heterozygous locus) / (number of total sequenced cells) × 0.5.

We estimated the doublet rate of Tapestri system using a 50:50 mixture data of K562 and Raji cell lines that was provided by Missino Bio, Inc. We investigated 4 loci with known truth genotype that are distinct between these cell lines for cell assignment. We assigned cells that were homozygous reference at the 4 loci to Raji, cells that were homozygous alternate at the 4 loci to K562 and cells that are heterozygous at these loci to doublet. The mean doublet rate based on the 45 Tapestri runs with 50:50 K562 Raji mixture was 5.86% with a standard deviation of 2.43%. This doublet rate correlates with the total number of cells to the bead ratio.

**Supplementary Table 1.** Clinical and demographic characteristics of the study cohort (N = 77).

| Characteristics | Median | IQR |
|---|---|---|
| **WBC** (cells/µL) | 11.1 | 4.3-37.7 |
| **HGB** (g/dL) | 9.1 | 8.4-10.0 |
| **PLT** ($\times 10^3$/µL) | 60 | 31-98 |
| **BM blasts** (%) | 44 | 29-67 |
| **PB blasts** (%) | 18 | 4-49 |
| **LDH** (U/L) | 888 | 613-1603 |
| **Age** (y) | 59 | 48-72 |
| | **No.** | **%** |
| **Diagnosis** | | |
| AML | 75 | 97 |
| Ontogeny | | |
| De novo | 57 | 76 |
| Secondary/therapy-related | 18 | 24 |
| MDS | 2 | 3 |
| **Prior treatment** | | |
| Untreated | 64 | 83 |
| Treated | 13 | 17 |
| **Karyotype** | | |
| Normal karyotype | 68 | 88 |
| Others | 9 | 12 |
| **Treatment** | | |
| IA-based chemotherapy | 43 | 56 |
| AraC-based chemotherapy | 12 | 16 |
| HMA | 16 | 21 |
| Others | 6 | 8 |
| **Sex** | | |
| Female | 30 | 39 |
| Male | 47 | 61 |

Abbreviations: IQR, interquartile range; WBC, white blood cells; HGB, hemoglobin; PLT, platelets; BM, bone marrow; PB, peripheral blood; LDH, lactate dehydrogenase; AML, acute myeloid leukemia; FAB, French-American-British; MDS, myelodysplastic syndrome; IA, idarubicin and cytarabine; AraC, cytarabine; HMA, hypomethylating agents

**Supplementary Table 2.** Estimated limit of detection of the single-cell DNA sequencing platform based on the dilution assay using cell line.

The single-cell DNA sequencing data for RAJI and K562 mixture was downloaded from http://tapestriportal.com. RAJI and K562 were mixed with a ratio of 50:50, 99:1, 99.5:0.5, and 99.9:0.1. Three loci with known truth genotype that are distinct between these cell lines were used to determine the sensitivity of the single-cell DNA sequencing platform. The first table shows the known zygosity for the 3 loci for each cell line. In the second table, RAJI and K562 indicate RAJI- and K562-type genotypes as shown in the first table. The number and percentage of cells showing RAJI- and K562-type genotypes are shown for each mixed sample.

Het, heterozygous; WT, wildtype; Homo, homozygous.

| Variant | RAJI | K562 |
|---|---|---|
| TP53: chr17:7577581A>G | Het | WT |
| TP53: chr17:757815T>C | Het | Homo |
| TP53: chr17:7578211C>T | Het | WT |

| Sample | RAJI | K562 |
|---|---|---|
| RAJI 50% + K562 50% | 2404 (42.7%) | 3358 (58.3%) |
| RAJI 99% + K562 1% | 6502 (99.1%) | 59 (0.9%) |
| RAJI 99.5% + K562 0.5% | 3570 (99.4%) | 21 (0.6%) |
| RAJI 99.9% + K562 0.1% | 4519 (99.9%) | 3 (0.1%) |

**Supplementary Table 3.** Somatic mutations detected by single-cell DNA sequencing

| sample ID | gene | chr | start | ref allele | alt allele | amino acid change | function | exonic function | mutated % | scDNA-seq VAF | WT (No. of cells) | Het (No. of cells) | Homo (No. of cells) | Missing (No. of cells) | validated | validation method | bulk VAF |
|---|---|---|---|---|---|---|---|---|---|---|---|---|---|---|---|---|---|
| AML-01-001 | *FLT3* | 13 | 28608259 | X | XAACTCTAAATTTTCTCTTGGAAACTCCCATTTGAGATCATATTCATATTCTCTGAAAT | p.Y599_D600delinsX | exonic | stopgain | 0.9352 | 0.7949 | 239 | 8444 | 2190 | 498 | yes | bulk-NGS | 0.1186 |
| AML-01-001 | *NPM1* | 5 | 170837544 | T | TCTGC | p.L287fs | exonic | frameshift insertion | 0.4249 | 0.4608 | 1285 | 4226 | 606 | 5254 | yes | bulk-NGS | 0.1828 |
| AML-01-001 | *WT1* | 11 | 32417913 | C | CGTACAAGA | p.R380fs | exonic | frameshift insertion | 0.0034 | 0.0015 | 10779 | 39 | 0 | 553 | yes | bulk-NGS in paired sample | 0 |
| AML-01-001 | *WT1* | 11 | 32417943 | C | CG | p.R370fs | exonic | frameshift insertion | 0.8836 | 0.4865 | 568 | 9799 | 248 | 756 | yes | bulk-NGS | 0.236 |
| AML-02-001 | *DNMT3A* | 2 | 25457242 | C | T | p.R882H | exonic | nonsynonymous SNV | 0.7603 | 0.4639 | 468 | 5861 | 169 | 1433 | yes | bulk-NGS | 0.423885 |
| AML-02-001 | *FLT3* | 13 | 28592641 | T | A | p.D835V | exonic | nonsynonymous SNV | 0.8274 | 0.4271 | 1186 | 6335 | 227 | 183 | yes | bulk-NGS | 0.406964 |
| AML-02-001 | *JAK2* | 9 | 5073770 | G | T | p.V617F | exonic | nonsynonymous SNV | 0.02 | 0.0412 | 3165 | 40 | 119 | 4607 | yes | bulk-NGS | 0.075206 |
| AML-02-001 | *RUNX1* | 21 | 36252939 | C | CT | p.S141fs | exonic | frameshift insertion | 0.5369 | 0.4347 | 1052 | 3971 | 287 | 2621 | yes | bulk-NGS | 0.3471 |
| AML-02-001 | *SF3B1* | 2 | 198267359 | C | G | p.K666N | exonic | nonsynonymous SNV | 0.8516 | 0.4416 | 1038 | 6517 | 237 | 139 | yes | bulk-NGS | 0.440258 |
| AML-03-001 | *FLT3* | 13 | 28608271 | X | XTCCCATTTGAGATCATATTCATATTCTCTGAAATCAACGTAGAAGT | p.R595_E596delinsRLLRX | exonic | stopgain | 0.8723 | 0.857 | 632 | 2642 | 5182 | 495 | yes | bulk-NGS | 0 |
| AML-03-001 | *NPM1* | 5 | 170837543 | C | CTCTG | p.L287fs | exonic | frameshift insertion | 0.5709 | 0.476 | 860 | 4730 | 380 | 2981 | yes | bulk-NGS | 0.2048 |
| AML-04-001 | *FLT3* | 13 | 28592640 | A | T | p.D835E | exonic | nonsynonymous SNV | 0.043 | 0.0215 | 5610 | 245 | 8 | 15 | yes | bulk-NGS | 0.023128 |
| AML-04-001 | *FLT3* | 13 | 28608286 | X | XAAGTACTCATTATCTGAGGAGCCGGTCACCTGTACCATC | p.F590delinsLMVQVTGSSDNEYF | exonic | nonframeshift insertion | 0.7383 | 0.6385 | 653 | 3881 | 1283 | 61 | yes | bulk-NGS | 0 |
| AML-04-001 | *IDH1* | 2 | 209113113 | G | A | p.R132C | exonic | nonsynonymous SNV | 0.001 | 0.0089 | 5808 | 6 | 0 | 64 | yes | bulk-NGS in paired sample | 0 |
| AML-04-001 | *NRAS* | 1 | 115258747 | C | T | p.G12D | exonic | nonsynonymous SNV | 0.0012 | 0.005 | 4746 | 7 | 0 | 1125 | yes | bulk-NGS in paired sample | 0 |
| AML-04-001 | *PTPN11* | 12 | 112888197 | T | A | p.F71L | exonic | nonsynonymous SNV | 0.1157 | 0.0606 | 5146 | 668 | 12 | 52 | yes | bulk-NGS | 0.034158 |
| AML-04-001 | *SF3B1* | 2 | 198267359 | C | G | p.K666N | exonic | nonsynonymous SNV | 0.9268 | 0.489 | 298 | 5226 | 222 | 132 | yes | bulk-NGS | 0.459384 |
| AML-04-001 | *SRSF2* | 17 | 74732959 | G | T | p.P95H | exonic | nonsynonymous SNV | 0.5645 | 0.5473 | 580 | 2539 | 779 | 1980 | yes | bulk-NGS | 0.450549 |

| | | | | | | | | | | | | | | | | | |
|---|---|---|---|---|---|---|---|---|---|---|---|---|---|---|---|---|---|
| AML-04-001 | *WT1* | 11 | 32417912 | C | CCT | p.R380fs | exonic | frameshift insertion | 0.0432 | 0.0192 | 5609 | 248 | 6 | 15 | yes | clinical sequencing | 0 |
| AML-04-001 | *WT1* | 11 | 32417914 | G | A | p.R380W | exonic | nonsynonymous SNV | 0.0446 | 0.0206 | 5601 | 256 | 6 | 15 | yes | bulk-NGS | 0.003958 |
| AML-05-001 | *FLT3* | 13 | 28592641 | T | G | p.D835A | exonic | nonsynonymous SNV | 0.3747 | 0.1811 | 6007 | 3757 | 62 | 365 | yes | bulk-NGS | 0.138702 |
| AML-05-001 | *FLT3* | 13 | 28592642 | C | A | p.D835Y | exonic | nonsynonymous SNV | 0.0193 | 0.0093 | 9820 | 191 | 6 | 174 | yes | bulk-NGS | 0.006711 |
| AML-05-001 | *FLT3* | 13 | 28602340 | G | T | p.N676K | exonic | nonsynonymous SNV | 0.0137 | 0.0058 | 9955 | 137 | 3 | 96 | yes | bulk-NGS | 0.005297 |
| AML-05-001 | *FLT3* | 13 | 28608262 | T | TTCATATTCTCTGAAG | p.E598delinsDFREYE | exonic | nonframeshift insertion | 0.015 | 0.009 | 7950 | 152 | 1 | 2088 | yes | bulk-NGS | 0.0136 |
| AML-05-001 | *NRAS* | 1 | 115258744 | C | T | p.G13D | exonic | nonsynonymous SNV | 0.3376 | 0.2416 | 3751 | 3369 | 71 | 3000 | yes | bulk-NGS | 0.20064 |
| AML-06-001 | *FLT3* | 13 | 28608262 | X | XACAAACTCTAAATTTTCTCTTGGAAACTCCCATTTGAGATCATATTCATATTCTC | p.V592_D593delinsVX | exonic | stopgain | 0.0153 | 0.0115 | 8152 | 457 | 8 | 2781 | yes | bulk-NGS | 0.0293 |
| AML-06-001 | *NRAS* | 1 | 115256528 | T | G | p.Q61H | exonic | nonsynonymous SNV | 0.1204 | 0.0547 | 9805 | 1341 | 31 | 221 | yes | bulk-NGS | 0.034483 |
| AML-06-001 | *NRAS* | 1 | 115258744 | C | T | p.G13D | exonic | nonsynonymous SNV | 0.0028 | 0.002 | 9134 | 32 | 0 | 2232 | yes | ddPCR (positive) | 0 |
| AML-06-001 | *NRAS* | 1 | 115258745 | C | G | p.G13R | exonic | nonsynonymous SNV | 0.0052 | 0.0031 | 9109 | 59 | 0 | 2230 | yes | ddPCR (positive) | 0 |
| AML-06-001 | *NRAS* | 1 | 115258747 | C | T | p.G12D | exonic | nonsynonymous SNV | 0.3737 | 0.2473 | 4165 | 4210 | 49 | 2974 | yes | bulk-NGS | 0.219721 |
| AML-07-001 | *IDH2* | 15 | 90631934 | C | T | p.R140Q | exonic | nonsynonymous SNV | 0.8974 | 0.466 | 487 | 5952 | 77 | 202 | yes | bulk-NGS | 0.459245 |
| AML-07-001 | *RUNX1* | 21 | 36252994 | T | TCC | p.D123fs | exonic | frameshift insertion | 0.7063 | 0.7813 | 1152 | 199 | 4546 | 821 | yes | bulk-NGS | 0.1774 |
| AML-07-001 | *SRSF2* | 17 | 74732959 | G | T | p.P95H | exonic | nonsynonymous SNV | 0.4896 | 0.556 | 334 | 2744 | 545 | 3095 | yes | bulk-NGS | 0.54321 |
| AML-08-001 | *DNMT3A* | 2 | 25457242 | C | T | p.R882H | exonic | nonsynonymous SNV | 0.7221 | 0.3826 | 706 | 3246 | 130 | 593 | yes | bulk-NGS | 0.467669 |
| AML-08-001 | *IDH2* | 15 | 90631934 | C | T | p.R140Q | exonic | nonsynonymous SNV | 0.8607 | 0.4381 | 524 | 3912 | 112 | 127 | yes | bulk-NGS | 0.462639 |
| AML-08-001 | *SRSF2* | 17 | 74732959 | G | A | p.P95L | exonic | nonsynonymous SNV | 0.271 | 0.2593 | 936 | 1149 | 118 | 2472 | yes | bulk-NGS | 0.453297 |
| AML-09-001 | *FLT3* | 13 | 28592640 | A | T | p.D835E | exonic | nonsynonymous SNV | 0.071 | 0.0314 | 7622 | 572 | 21 | 139 | yes | clinical sequencing | not tested |
| AML-09-001 | *FLT3* | 13 | 28592642 | C | A | p.D835Y | exonic | nonsynonymous SNV | 0.0085 | 0.0041 | 8187 | 66 | 5 | 96 | yes | clinical sequencing | not tested |
| AML-09-001 | *FLT3* | 13 | 28608312 | X | XTCCGA | p.F590_Y591delinsLVLLRX | exonic | frameshift insertion | 0.4381 | 0.3129 | 2706 | 4890 | 162 | 596 | yes | qPCR | not tested |
| AML-09-001 | *KRAS* | 12 | 25398281 | C | T | p.G13D | exonic | nonsynonymous SNV | 0.0393 | 0.0197 | 7788 | 304 | 24 | 238 | yes | clinical sequencing | not tested |

| | | | | | | | | | | | | | | | | | |
|---|---|---|---|---|---|---|---|---|---|---|---|---|---|---|---|---|---|
| AML-09-001 | *NPM1* | 5 | 170837543 | C | CTCTG | p.L287fs | exonic | frameshift insertion | 0.4682 | 0.4519 | 852 | 3650 | 261 | 3591 | yes | clinical sequencing | not tested |
| AML-10-001 | *FLT3* | 13 | 28608274 | X | XGGAAACTCCCATTTGAGATCATATTCATATTCTCTGAAATCAA | p.F594_R595delinsFX | exonic | stopgain | 0.1871 | 0.1325 | 6257 | 1592 | 44 | 836 | yes | bulk-NGS | 0 |
| AML-10-001 | *GATA2* | 3 | 128202813 | T | C | p.T303A | exonic | nonsynonymous SNV | 0.496 | 0.4122 | 1698 | 3808 | 522 | 2701 | yes | bulk-NGS | 0.319797 |
| AML-10-001 | *SRSF2* | 17 | 74732959 | G | A | p.P95L | exonic | nonsynonymous SNV | 0.0459 | 0.2236 | 474 | 336 | 65 | 7854 | yes | bulk-NGS | 0.356021 |
| AML-10-001 | *WT1* | 11 | 32414262 | C | G | p.R430P | exonic | nonsynonymous SNV | 0.513 | 0.4525 | 535 | 4384 | 94 | 3716 | yes | bulk-NGS | 0.358228 |
| AML-11-001 | *ASXL1* | 20 | 31022441 | A | AG | p.G642fs | exonic | frameshift insertion | 0.2378 | 0.2582 | 693 | 1252 | 2 | 3327 | yes | bulk-NGS | 0.1387 |
| AML-11-001 | *SRSF2* | 17 | 74732960 | G | C | p.P95A | exonic | nonsynonymous SNV | 0.2108 | 0.5819 | 251 | 759 | 353 | 3911 | yes | bulk-NGS | 0.488778 |
| AML-12-001 | *FLT3* | 13 | 28592634 | CATG | C | p.836_837del | exonic | nonframeshift deletion | 0.0651 | 0.0275 | 6765 | 467 | 8 | 60 | yes | bulk-NGS | 0.0903 |
| AML-12-001 | *NPM1* | 5 | 170837545 | C | CTGCT | p.L287fs | exonic | frameshift insertion | 0.617 | 0.514 | 646 | 4330 | 174 | 2150 | yes | bulk-NGS | 0.2 |
| AML-12-001 | *PTPN11* | 12 | 112888211 | A | T | p.E76V | exonic | nonsynonymous SNV | 0.1349 | 0.064 | 6168 | 968 | 17 | 147 | yes | bulk-NGS | 0.0425 |
| AML-13-001 | *IDH2* | 15 | 90631934 | C | T | p.R140Q | exonic | nonsynonymous SNV | 0.4609 | 0.2341 | 4585 | 4130 | 177 | 453 | yes | bulk-NGS | 0.221805 |
| AML-13-001 | *NPM1* | 5 | 170837543 | C | CTCTG | p.L287fs | exonic | frameshift insertion | 0.2037 | 0.1703 | 4139 | 1648 | 256 | 3302 | yes | ddPCR (positive) | 0 |
| AML-13-001 | *TP53* | 17 | 7578415 | AC | A | p.V172fs | exonic | frameshift deletion | 0.0138 | 0.0062 | 8714 | 127 | 2 | 502 | yes | ddPCR (positive) | 0 |
| AML-13-001 | *WT1* | 11 | 32413565 | C | G | p.R462P | exonic | nonsynonymous SNV | 0.2901 | 0.1865 | 4307 | 2684 | 27 | 2327 | yes | bulk-NGS | 0.11236 |
| AML-14-001 | *KRAS* | 12 | 25398281 | C | T | p.G13D | exonic | nonsynonymous SNV | 0.0357 | 0.0183 | 6706 | 245 | 7 | 95 | yes | bulk-NGS | 0.018203 |
| AML-14-001 | *KRAS* | 12 | 25398284 | C | A | p.G12V | exonic | nonsynonymous SNV | 0.1199 | 0.0592 | 6078 | 796 | 50 | 129 | yes | bulk-NGS | 0.026226 |
| AML-14-001 | *KRAS* | 12 | 25398284 | C | T | p.G12D | exonic | nonsynonymous SNV | 0.0095 | 0.005 | 6857 | 64 | 3 | 129 | yes | ddPCR (positive) | 0 |
| AML-14-001 | *NPM1* | 5 | 170837543 | C | CTCTG | p.L287fs | exonic | frameshift insertion | 0.6468 | 0.461 | 736 | 4360 | 202 | 1755 | yes | bulk-NGS | 0.1644 |
| AML-14-001 | *NRAS* | 1 | 115258745 | C | G | p.G13R | exonic | nonsynonymous SNV | 0.1411 | 0.0714 | 5319 | 968 | 27 | 739 | yes | bulk-NGS | 0.05235 |
| AML-14-001 | *NRAS* | 1 | 115258747 | C | T | p.G12D | exonic | nonsynonymous SNV | 0.0306 | 0.0165 | 6145 | 208 | 8 | 692 | yes | bulk-NGS | 0.013845 |
| AML-14-001 | *PTPN11* | 12 | 112888198 | G | A | p.A72T | exonic | nonsynonymous SNV | 0.2015 | 0.0966 | 5527 | 1388 | 33 | 105 | yes | bulk-NGS | 0.056229 |
| AML-15-001 | *IDH1* | 2 | 209113113 | G | C | p.R132G | exonic | nonsynonymous SNV | 0.868 | 0.4564 | 524 | 4973 | 123 | 251 | yes | bulk-NGS | 0.427012 |
| AML-15-001 | *NPM1* | 5 | 170837543 | C | CTCTG | p.L287fs | exonic | frameshift insertion | 0.4866 | 0.4684 | 521 | 2687 | 170 | 2493 | yes | bulk-NGS | 0.216 |

| | | | | | | | | | | | | | | | | | |
|---|---|---|---|---|---|---|---|---|---|---|---|---|---|---|---|---|---|
| AML-16-001 | *DNMT3A* | 2 | 25457242 | C | T | p.R882H | exonic | nonsynonymous SNV | 0.295 | 0.168 | 5040 | 2812 | 98 | 1914 | yes | bulk-NGS | 0.180952 |
| AML-16-001 | *IDH1* | 2 | 209113112 | C | T | p.R132H | exonic | nonsynonymous SNV | 0.4111 | 0.1702 | 5599 | 3956 | 99 | 210 | yes | bulk-NGS | 0.18543 |
| AML-16-001 | *IDH2* | 15 | 90631934 | C | T | p.R140Q | exonic | nonsynonymous SNV | 0.0057 | 0.0022 | 9426 | 54 | 2 | 382 | yes | ddPCR (positive) | 0 |
| AML-17-001 | *DNMT3A* | 2 | 25457283 | A | T | p.F868L | exonic | nonsynonymous SNV | 0.7286 | 0.4711 | 571 | 4564 | 262 | 1227 | yes | bulk-NGS | 0.348485 |
| AML-17-001 | *NPM1* | 5 | 170837543 | C | CTCTG | p.L287fs | exonic | frameshift insertion | 0.5436 | 0.4825 | 641 | 3265 | 336 | 2382 | yes | bulk-NGS | 0.2927 |
| AML-17-001 | *NRAS* | 1 | 115258747 | C | T | p.G12D | exonic | nonsynonymous SNV | 0.7097 | 0.4659 | 489 | 4576 | 125 | 1434 | yes | bulk-NGS | 0.337638 |
| AML-18-001 | *ASXL1* | 20 | 31022936 | TC | T | p.P808fs | exonic | frameshift deletion | 0.4866 | 0.2644 | 3190 | 3664 | 124 | 807 | yes | bulk-NGS | 0.1989 |
| AML-18-001 | *IDH2* | 15 | 90631838 | C | T | p.R172K | exonic | nonsynonymous SNV | 0.7747 | 0.5024 | 22 | 5867 | 164 | 1732 | yes | bulk-NGS | 0.319149 |
| AML-18-001 | *RUNX1* | 21 | 36252852 | A | ACCT | splicing | splicing | . | 0.5941 | 0.3918 | 1523 | 4408 | 217 | 1637 | yes | bulk-NGS | 0.2841 |
| AML-19-001 | *FLT3* | 13 | 28608265 | X | XCCCCTTGGAAACTCCCATTTGAGATCATATTCATATTCTCTGAAATCAA | p.D593_F594delinsELLRX | exonic | stopgain | 0.8006 | 0.804 | 575 | 1420 | 2815 | 456 | yes | bulk-NGS | 0 |
| AML-19-001 | *RUNX1* | 21 | 36206720 | CTG | C | p.Q264fs | exonic | frameshift deletion | 0.6673 | 0.4107 | 1020 | 3389 | 125 | 732 | yes | bulk-NGS | 0.1127 |
| AML-19-001 | *WT1* | 11 | 32417909 | CG | C | p.S381fs | exonic | frameshift deletion | 0.2801 | 0.1347 | 3350 | 1470 | 5 | 441 | yes | bulk-NGS | 0.1193 |
| AML-20-001 | *DNMT3A* | 2 | 25457243 | G | A | p.R882C | exonic | nonsynonymous SNV | 0.6134 | 0.389 | 1516 | 6048 | 117 | 2369 | yes | bulk-NGS | 0.476056 |
| AML-20-001 | *NPM1* | 5 | 170837543 | C | CTCTG | p.L287fs | exonic | frameshift insertion | 0.3078 | 0.2525 | 2967 | 2932 | 161 | 3990 | yes | bulk-NGS | 0.1556 |
| AML-20-001 | *NRAS* | 1 | 115258744 | C | A | p.G13V | exonic | nonsynonymous SNV | 0.3298 | 0.2441 | 3530 | 3260 | 54 | 3206 | yes | bulk-NGS | 0.331135 |
| AML-21-001 | *NPM1* | 5 | 170837543 | C | CTCTG | p.L287fs | exonic | frameshift insertion | 0.5089 | 0.417 | 1093 | 3844 | 411 | 3013 | yes | bulk-NGS | 0.2931 |
| AML-21-001 | *NRAS* | 1 | 115258747 | C | T | p.G12D | exonic | nonsynonymous SNV | 0.5923 | 0.4597 | 624 | 4841 | 111 | 2785 | yes | bulk-NGS | 0.370618 |
| AML-21-001 | *WT1* | 11 | 32417907 | G | GCCGA | p.A382fs | exonic | frameshift insertion | 0.9235 | 0.4713 | 484 | 7567 | 154 | 156 | yes | bulk-NGS | 0.3192 |
| AML-22-001 | *KRAS* | 12 | 25398284 | C | A | p.G12V | exonic | nonsynonymous SNV | 0.3843 | 0.1886 | 1640 | 1064 | 60 | 161 | yes | bulk-NGS | 0.132517 |
| AML-22-001 | *KRAS* | 12 | 25398285 | C | T | p.G12S | exonic | nonsynonymous SNV | 0.026 | 0.0207 | 2722 | 73 | 3 | 127 | yes | bulk-NGS | 0.010067 |
| AML-22-001 | *NRAS* | 1 | 115258744 | C | T | p.G13D | exonic | nonsynonymous SNV | 0.1262 | 0.0635 | 2254 | 363 | 6 | 302 | yes | bulk-NGS | 0.061202 |
| AML-22-001 | *NRAS* | 1 | 115258745 | C | A | p.G13C | exonic | nonsynonymous SNV | 0.0239 | 0.0179 | 2569 | 68 | 2 | 286 | yes | bulk-NGS | 0.016484 |
| AML-22-001 | *NRAS* | 1 | 115258747 | C | A | p.G12V | exonic | nonsynonymous SNV | 0.0062 | 0.006 | 2609 | 17 | 1 | 298 | yes | ddPCR (positive) | 0 |

| | | | | | | | | | | | | | | | | | |
|---|---|---|---|---|---|---|---|---|---|---|---|---|---|---|---|---|---|
| AML-22-001 | *NRAS* | 1 | 115258747 | C | T | p.G12D | exonic | nonsynonymous SNV | 0.0615 | 0.0329 | 2447 | 172 | 8 | 298 | yes | bulk-NGS | 0.036545 |
| AML-22-001 | *NRAS* | 1 | 115258748 | C | A | p.G12C | exonic | nonsynonymous SNV | 0.0988 | 0.0559 | 2323 | 281 | 8 | 313 | yes | bulk-NGS | 0.050847 |
| AML-22-001 | *NRAS* | 1 | 115258748 | C | T | p.G12S | exonic | nonsynonymous SNV | 0.0537 | 0.0312 | 2455 | 152 | 5 | 313 | yes | ddPCR (positive) | 0 |
| AML-23-001 | *NRAS* | 1 | 115258745 | C | G | p.G13R | exonic | nonsynonymous SNV | 0.2775 | 0.1463 | 1222 | 528 | 45 | 270 | yes | bulk-NGS | 0.15572 |
| AML-23-001 | *WT1* | 11 | 32417916 | A | ACAAGAGTC | p.V379fs | exonic | frameshift insertion | 0.6378 | 0.3072 | 683 | 1265 | 52 | 65 | yes | bulk-NGS | 0.2582 |
| AML-24-001 | *DNMT3A* | 2 | 25457242 | C | T | p.R882H | exonic | nonsynonymous SNV | 0.7778 | 0.4411 | 561 | 4944 | 141 | 892 | yes | bulk-NGS | 0.403846 |
| AML-24-001 | *FLT3* | 13 | 28602329 | G | A | p.A680V | exonic | nonsynonymous SNV | 0.0043 | 0.0015 | 6450 | 28 | 0 | 60 | yes | ddPCR (positive) | 0 |
| AML-24-001 | *FLT3* | 13 | 28608301 | X | XCTGAAATCAACGTAGA | p.T582delinsKYA | exonic | stopgain | 0.2213 | 0.1771 | 2507 | 3646 | 100 | 285 | yes | qPCR | 0 |
| AML-24-001 | *U2AF1* | 21 | 44524456 | G | T | p.S34Y | exonic | nonsynonymous SNV | 0.8796 | 0.4547 | 587 | 5602 | 149 | 200 | yes | bulk-NGS | 0.405333 |
| AML-24-001 | *WT1* | 11 | 32413592 | C | T | p.C453Y | exonic | nonsynonymous SNV | 0.2583 | 0.1603 | 3549 | 1663 | 26 | 1300 | yes | bulk-NGS | 0.081028 |
| AML-25-001 | *ASXL1* | 20 | 31022441 | A | AG | p.G642fs | exonic | frameshift insertion | 0.6734 | 0.243 | 1327 | 4263 | 24 | 752 | yes | bulk-NGS | 0.1438 |
| AML-25-001 | *EZH2* | 7 | 148506462 | G | A | p.R670C | exonic | nonsynonymous SNV | 0.7537 | 0.3818 | 1448 | 4625 | 173 | 120 | yes | bulk-NGS | 0.459459 |
| AML-25-001 | *FLT3* | 13 | 28608270 | C | CTCTGAAGGGG | p.E596fs | exonic | frameshift insertion | 0.017 | 0.0085 | 6223 | 108 | 0 | 35 | yes | qPCR | 0 |
| AML-25-001 | *RUNX1* | 21 | 36164872 | G | A | p.Q335X | exonic | stopgain | 0.2336 | 0.7353 | 518 | 47 | 1440 | 4361 | yes | bulk-NGS | 0.688119 |
| AML-26-001 | *DNMT3A* | 2 | 25457242 | C | T | p.R882H | exonic | nonsynonymous SNV | 0.5265 | 0.2864 | 3322 | 4961 | 94 | 1224 | yes | bulk-NGS | 0.311346 |
| AML-26-001 | *FLT3* | 13 | 28592641 | T | G | p.D835A | exonic | nonsynonymous SNV | 0.1035 | 0.0498 | 8490 | 971 | 23 | 117 | yes | bulk-NGS | 0.056522 |
| AML-26-001 | *FLT3* | 13 | 28592642 | C | A | p.D835Y | exonic | nonsynonymous SNV | 0.0134 | 0.0062 | 9407 | 126 | 3 | 65 | yes | bulk-NGS | 0.006522 |
| AML-26-001 | *KRAS* | 12 | 25380276 | T | A | p.Q61L | exonic | nonsynonymous SNV | 0.0534 | 0.0281 | 8452 | 495 | 18 | 636 | yes | bulk-NGS | 0.031183 |
| AML-26-001 | *NPM1* | 5 | 170837543 | C | CTCTG | p.L287fs | exonic | frameshift insertion | 0.1768 | 0.1138 | 5586 | 1628 | 69 | 2318 | yes | bulk-NGS | 0.0924 |
| AML-26-001 | *NRAS* | 1 | 115258744 | C | T | p.G13D | exonic | nonsynonymous SNV | 0.0269 | 0.0143 | 7982 | 254 | 4 | 1361 | yes | bulk-NGS | 0.010787 |
| AML-27-001 | *DNMT3A* | 2 | 25457243 | G | A | p.R882C | exonic | nonsynonymous SNV | 0.6681 | 0.4421 | 743 | 2533 | 197 | 613 | yes | bulk-NGS | 0.497664 |
| AML-27-001 | *FLT3* | 13 | 28608273 | X | XTGGAAACTCCCATTTGAGATCATATTCATATTCTCTGAAATCAA | p.R595fs | exonic | frameshift insertion | 0.8245 | 0.4324 | 656 | 3261 | 108 | 61 | yes | bulk-NGS | 0.1224 |
| AML-27-001 | *NPM1* | 5 | 170837543 | C | CTCTG | p.L287fs | exonic | frameshift insertion | 0.502 | 0.4435 | 571 | 1908 | 143 | 1464 | yes | bulk-NGS | 0.1754 |
| AML-27-001 | *TP53* | 17 | 7578275 | G | A | p.Q192X | exonic | stopgain | 0.7359 | 0.369 | 994 | 2889 | 118 | 85 | yes | bulk-NGS | 0.352853 |

| | | | | | | | | | | | | | | | | | |
|---|---|---|---|---|---|---|---|---|---|---|---|---|---|---|---|---|---|
| AML-28-001 | *DNMT3A* | 2 | 25457243 | G | A | p.R882C | exonic | nonsynonymous SNV | 0.7177 | 0.4516 | 698 | 7143 | 170 | 2178 | yes | bulk-NGS | 0.439024 |
| AML-28-001 | *FLT3* | 13 | 28608282 | C | CTGCAAAGACAAATGGTGAGTACGTGCATTTTAAAGATTTTCCAATG | p.V592fs | exonic | frameshift insertion | 0.1021 | 0.0366 | 7794 | 1040 | 0 | 1355 | yes | bulk-NGS | 0.0569 |
| AML-28-001 | *IDH1* | 2 | 209113112 | C | T | p.R132H | exonic | nonsynonymous SNV | 0.7791 | 0.39 | 1918 | 7762 | 176 | 333 | yes | bulk-NGS | 0.444444 |
| AML-28-001 | *IDH2* | 15 | 90631934 | C | T | p.R140Q | exonic | nonsynonymous SNV | 0.1207 | 0.0569 | 8356 | 1182 | 48 | 603 | yes | bulk-NGS | 0.061224 |
| AML-28-001 | *NPM1* | 5 | 170837543 | C | CTCTG | p.L287fs | exonic | frameshift insertion | 0.4446 | 0.4639 | 1034 | 4086 | 444 | 4625 | yes | clinical sequencing | 0 |
| AML-29-001 | *ASXL1* | 20 | 31022440 | G | GA | p.G642fs | exonic | frameshift insertion | 0.7923 | 0.4624 | 477 | 5983 | 193 | 1142 | yes | bulk-NGS | 0.1299 |
| AML-29-001 | *IDH2* | 15 | 90631934 | C | T | p.R140Q | exonic | nonsynonymous SNV | 0.0233 | 0.0135 | 7569 | 176 | 6 | 44 | yes | bulk-NGS | 0.020548 |
| AML-29-001 | *RUNX1* | 21 | 36231782 | C | T | p.R201Q | exonic | nonsynonymous SNV | 0.2409 | 0.4756 | 340 | 1708 | 170 | 5577 | yes | bulk-NGS | 0.272727 |
| AML-29-001 | *SRSF2* | 17 | 74732959 | G | T | p.P95H | exonic | nonsynonymous SNV | 0.4076 | 0.5642 | 538 | 2379 | 798 | 4080 | yes | bulk-NGS | 0.361538 |
| AML-30-001 | *DNMT3A* | 2 | 25457242 | C | T | p.R882H | exonic | nonsynonymous SNV | 0.3494 | 0.1745 | 3744 | 2298 | 32 | 594 | yes | bulk-NGS | 0.227554 |
| AML-30-001 | *IDH2* | 15 | 90631934 | C | T | p.R140Q | exonic | nonsynonymous SNV | 0.0396 | 0.0182 | 6329 | 255 | 9 | 75 | yes | bulk-NGS | 0.021992 |
| AML-30-001 | *NPM1* | 5 | 170837543 | C | CTCTG | p.L287fs | exonic | frameshift insertion | 0.0301 | 0.0181 | 5239 | 193 | 8 | 1228 | yes | bulk-NGS | 0.0146 |
| AML-30-001 | *NRAS* | 1 | 115258744 | C | T | p.G13D | exonic | nonsynonymous SNV | 0.0354 | 0.0186 | 5738 | 233 | 3 | 694 | yes | bulk-NGS | 0.019459 |
| AML-31-001 | *FLT3* | 13 | 28608286 | X | XTCATATTCTCTGAAATCAACGTAGAAGTACT | p.F590_Y591delinsLVLLRX | exonic | stopgain | 0.0372 | 0.0262 | 8702 | 375 | 5 | 1141 | yes | qPCR | 0 |
| AML-31-001 | *NPM1* | 5 | 170837543 | C | CTCTG | p.L287fs | exonic | frameshift insertion | 0.2621 | 0.2468 | 2950 | 2576 | 103 | 4594 | yes | clinical sequencing | 0 |
| AML-31-001 | *NRAS* | 1 | 115258744 | C | A | p.G13V | exonic | nonsynonymous SNV | 0.1195 | 0.0772 | 6374 | 1201 | 21 | 2627 | yes | bulk-NGS | 0.083333 |
| AML-31-001 | *PTPN11* | 12 | 112888139 | C | A | p.T52N | exonic | nonsynonymous SNV | 0.0229 | 0.0099 | 9604 | 229 | 5 | 385 | yes | ddPCR (positive) | 0 |
| AML-31-001 | *WT1* | 11 | 32417907 | G | GCCGA | p.A382fs | exonic | frameshift insertion | 0.0664 | 0.0317 | 9226 | 674 | 5 | 318 | yes | ddPCR (positive) | 0 |
| AML-32-001 | *DNMT3A* | 2 | 25457242 | C | T | p.R882H | exonic | nonsynonymous SNV | 0.6849 | 0.4248 | 902 | 5551 | 93 | 1695 | yes | bulk-NGS | 0.376623 |
| AML-32-001 | *FLT3* | 13 | 28592642 | C | A | p.D835Y | exonic | nonsynonymous SNV | 0.6763 | 0.3344 | 2500 | 5422 | 151 | 168 | yes | bulk-NGS | 0.246032 |
| AML-32-001 | *FLT3* | 13 | 28608283 | X | XGCATATTCATATTCTCTGAAATCAACGTAGAAGT | p.V581fs | exonic | stopgain | 0.0546 | 0.0395 | 6345 | 878 | 17 | 1001 | yes | qPCR | 0 |
| AML-32-001 | *RUNX1* | 21 | 36252866 | G | C | p.R166G | exonic | nonsynonymous SNV | 0.6178 | 0.8748 | 676 | 125 | 4966 | 2474 | yes | bulk-NGS | 0.732143 |

| | | | | | | | | | | | | | | | | | |
|---|---|---|---|---|---|---|---|---|---|---|---|---|---|---|---|---|---|
| AML-32-001 | *SF3B1* | 2 | 198266834 | T | C | p.K700E | exonic | nonsynonymous SNV | 0.8537 | 0.4406 | 1038 | 6830 | 205 | 168 | yes | bulk-NGS | 0.397059 |
| AML-33-001 | *IDH2* | 15 | 90631934 | C | T | p.R140Q | exonic | nonsynonymous SNV | 0.5454 | 0.2812 | 3256 | 4248 | 181 | 435 | yes | bulk-NGS | 0.306667 |
| AML-33-001 | *RUNX1* | 21 | 36231792 | C | T | p.D198N | exonic | nonsynonymous SNV | 0.1915 | 0.217 | 1990 | 1507 | 48 | 4575 | yes | bulk-NGS | 0.193548 |
| AML-33-001 | *SRSF2* | 17 | 74732959 | G | T | p.P95H | exonic | nonsynonymous SNV | 0.2361 | 0.4082 | 1004 | 1487 | 430 | 5199 | yes | bulk-NGS | 0.35514 |
| AML-34-001 | *KRAS* | 12 | 25398284 | C | T | p.G12D | exonic | nonsynonymous SNV | 0.0629 | 0.0359 | 3859 | 267 | 7 | 226 | yes | bulk-NGS | 0.017083 |
| AML-34-001 | *NRAS* | 1 | 115258744 | C | T | p.G13D | exonic | nonsynonymous SNV | 0.0053 | 0.0067 | 3849 | 22 | 1 | 487 | yes | ddPCR (positive) | 0 |
| AML-34-001 | *NRAS* | 1 | 115258747 | C | T | p.G12D | exonic | nonsynonymous SNV | 0.0014 | 0.0044 | 3863 | 6 | 0 | 490 | yes | ddPCR (positive) | 0 |
| AML-34-001 | *SF3B1* | 2 | 198266834 | T | C | p.K700E | exonic | nonsynonymous SNV | 0.0679 | 0.0365 | 3995 | 286 | 10 | 68 | yes | bulk-NGS | 0.015038 |
| AML-35-001 | *FLT3* | 13 | 28608275 | X | XCGACGCCGCGCCATTTGAGATCATATTCATATTCTCTGAAAT | p.F594delinsYFREYEYDLKWRGVV | exonic | nonframeshift insertion | 0.0024 | 0.0015 | 6546 | 18 | 0 | 1020 | yes | bulk-NGS | 0.0811 |
| AML-35-001 | *NRAS* | 1 | 115258747 | C | T | p.G12D | exonic | nonsynonymous SNV | 0.3891 | 0.2264 | 3554 | 2846 | 105 | 1079 | yes | bulk-NGS | 0.249377 |
| AML-35-001 | *PTPN11* | 12 | 112888199 | C | T | p.A72V | exonic | nonsynonymous SNV | 0.0145 | 0.008 | 7082 | 109 | 1 | 392 | yes | bulk-NGS | 0.010152 |
| AML-36-001 | *FLT3* | 13 | 28592640 | A | T | p.D835E | exonic | nonsynonymous SNV | 0.1333 | 0.0642 | 6812 | 1016 | 71 | 254 | yes | bulk-NGS | 0.058219 |
| AML-36-001 | *NRAS* | 1 | 115258747 | C | T | p.G12D | exonic | nonsynonymous SNV | 0.0115 | 0.0061 | 7257 | 90 | 4 | 802 | yes | bulk-NGS | 0.00655 |
| AML-36-001 | *NRAS* | 1 | 115258748 | C | T | p.G12S | exonic | nonsynonymous SNV | 0.0623 | 0.0328 | 6817 | 495 | 13 | 828 | yes | bulk-NGS | 0.023913 |
| AML-37-001 | *DNMT3A* | 2 | 25457252 | T | C | p.N879D | exonic | nonsynonymous SNV | 0.8184 | 0.4716 | 469 | 4924 | 186 | 665 | yes | bulk-NGS | 0.475 |
| AML-37-001 | *NPM1* | 5 | 170837543 | C | CTCTG | p.L287fs | exonic | frameshift insertion | 0.6156 | 0.4805 | 498 | 3657 | 187 | 1902 | yes | bulk-NGS | 0.25 |
| AML-37-001 | *NRAS* | 1 | 115258747 | C | T | p.G12D | exonic | nonsynonymous SNV | 0.8061 | 0.4755 | 474 | 4887 | 146 | 737 | yes | bulk-NGS | 0.419183 |
| AML-38-001 | *FLT3* | 13 | 28592642 | C | G | p.D835H | exonic | nonsynonymous SNV | 0.0224 | 0.0112 | 7054 | 156 | 6 | 19 | yes | bulk-NGS | 0.006472 |
| AML-38-001 | *FLT3* | 13 | 28608261 | X | XTGAGATCATATTCATATTCTCTGAAATCAACGTAGAAGTACTCATTATCTGAGGAGCCGGTCACCTGTACCAT | p.M578delinsKQFRYESQLQMVQVTGSSDNEYFYVDFREYEM | exonic | frameshift insertion | 0.2202 | 0.1963 | 5147 | 1593 | 133 | 362 | yes | bulk-NGS | 0.133 |
| AML-38-001 | *IDH1* | 2 | 209113112 | C | T | p.R132H | exonic | nonsynonymous SNV | 0.5084 | 0.2356 | 3432 | 3603 | 75 | 125 | yes | bulk-NGS | 0.221122 |
| AML-38-001 | *IDH2* | 15 | 90631934 | C | T | p.R140Q | exonic | nonsynonymous SNV | 0.4137 | 0.1823 | 4103 | 2811 | 182 | 139 | yes | bulk-NGS | 0.183515 |
| AML-38-001 | *KRAS* | 12 | 25398284 | C | G | p.G12A | exonic | nonsynonymous SNV | 0.3797 | 0.2023 | 4348 | 2618 | 129 | 140 | yes | bulk-NGS | 0.180068 |

| | | | | | | | | | | | | | | | | | |
|---|---|---|---|---|---|---|---|---|---|---|---|---|---|---|---|---|---|
| AML-38-001 | *KRAS* | 12 | 25398284 | C | T | p.G12D | exonic | nonsynonymous SNV | 0.0023 | 0.0043 | 7078 | 16 | 1 | 140 | yes | ddPCR in paired sample | 0 |
| AML-38-001 | *NPM1* | 5 | 170837544 | T | TCTGC | p.L287fs | exonic | frameshift insertion | 0.6012 | 0.43 | 764 | 4119 | 231 | 2121 | yes | bulk-NGS | 0.1962 |
| AML-38-001 | *NRAS* | 1 | 115258745 | C | G | p.G13R | exonic | nonsynonymous SNV | 0.1035 | 0.0565 | 5560 | 728 | 21 | 926 | yes | bulk-NGS | 0.032644 |
| AML-38-001 | *NRAS* | 1 | 115258747 | C | G | p.G12A | exonic | nonsynonymous SNV | 0.0437 | 0.0267 | 6042 | 309 | 7 | 877 | yes | bulk-NGS | 0.017372 |
| AML-38-001 | *PTPN11* | 12 | 112888165 | G | C | p.D61H | exonic | nonsynonymous SNV | 0.098 | 0.0494 | 6474 | 688 | 21 | 52 | yes | bulk-NGS | 0.032301 |
| AML-38-001 | *PTPN11* | 12 | 112888199 | C | G | p.A72G | exonic | nonsynonymous SNV | 0.0135 | 0.009 | 7088 | 90 | 8 | 49 | yes | bulk-NGS | 0.010101 |
| AML-38-001 | *PTPN11* | 12 | 112926888 | G | C | p.G503A | exonic | nonsynonymous SNV | 0.0015 | 0.0023 | 6750 | 11 | 0 | 474 | yes | bulk-NGS in paired sample | 0 |
| AML-39-001 | *GATA2* | 3 | 128202830 | T | C | p.N297S | exonic | nonsynonymous SNV | 0.4595 | 0.9488 | 100 | 35 | 3060 | 3540 | yes | bulk-NGS | 0.831169 |
| AML-39-001 | *PTPN11* | 12 | 112888199 | C | T | p.A72V | exonic | nonsynonymous SNV | 0.0457 | 0.0198 | 6300 | 302 | 6 | 127 | yes | bulk-NGS | 0.030303 |
| AML-39-001 | *PTPN11* | 12 | 112888211 | A | G | p.E76G | exonic | nonsynonymous SNV | 0.0199 | 0.0092 | 6477 | 131 | 3 | 124 | no | | 0 |
| AML-39-001 | *PTPN11* | 12 | 112926885 | C | T | p.S502L | exonic | nonsynonymous SNV | 0.0105 | 0.0104 | 3514 | 69 | 2 | 3150 | no | | 0 |
| AML-39-001 | *PTPN11* | 12 | 112926888 | G | C | p.G503A | exonic | nonsynonymous SNV | 0.2598 | 0.271 | 1551 | 1740 | 10 | 3434 | yes | bulk-NGS | 0.315476 |
| AML-39-001 | *SF3B1* | 2 | 198266834 | T | C | p.K700E | exonic | nonsynonymous SNV | 0.9661 | 0.4971 | 167 | 6353 | 154 | 61 | yes | bulk-NGS | 0.468619 |
| AML-40-001 | *FLT3* | 13 | 28602340 | G | C | p.N676K | exonic | nonsynonymous SNV | 0.0636 | 0.0301 | 5661 | 370 | 24 | 139 | yes | bulk-NGS | 0.026989 |
| AML-40-001 | *FLT3* | 13 | 28602340 | G | T | p.N676K | exonic | nonsynonymous SNV | 0.0076 | 0.0034 | 6008 | 42 | 5 | 139 | yes | ddPCR (positive) | 0 |
| AML-40-001 | *FLT3* | 13 | 28608313 | X | XTCTC | p.V581fs | exonic | frameshift insertion | 0.006 | 0.0042 | 5436 | 34 | 3 | 721 | yes | qPCR | 0 |
| AML-40-001 | *IDH2* | 15 | 90631934 | C | T | p.R140Q | exonic | nonsynonymous SNV | 0.8074 | 0.4647 | 785 | 4560 | 441 | 408 | yes | bulk-NGS | 0.419558 |
| AML-40-001 | *NPM1* | 5 | 170837543 | C | CTCTG | p.L287fs | exonic | frameshift insertion | 0.444 | 0.4683 | 736 | 2344 | 406 | 2708 | yes | bulk-NGS | 0.2584 |
| AML-40-001 | *NRAS* | 1 | 115258744 | C | T | p.G13D | exonic | nonsynonymous SNV | 0.0063 | 0.0039 | 5225 | 38 | 1 | 930 | yes | ddPCR (positive) | 0 |
| AML-40-001 | *NRAS* | 1 | 115258747 | C | T | p.G12D | exonic | nonsynonymous SNV | 0.289 | 0.1781 | 3343 | 1705 | 85 | 1061 | yes | bulk-NGS | 0.138848 |
| AML-41-001 | *IDH2* | 15 | 90631934 | C | T | p.R140Q | exonic | nonsynonymous SNV | 0.817 | 0.4526 | 660 | 3827 | 325 | 270 | yes | bulk-NGS | 0.422512 |
| AML-41-001 | *KRAS* | 12 | 25398284 | C | G | p.G12A | exonic | nonsynonymous SNV | 0.0201 | 0.019 | 4584 | 95 | 7 | 396 | yes | ddPCR (positive) | 0 |
| AML-41-001 | *NRAS* | 1 | 115258745 | C | G | p.G13R | exonic | nonsynonymous SNV | 0.1281 | 0.0778 | 3580 | 628 | 23 | 851 | yes | bulk-NGS | 0.047312 |

| | | | | | | | | | | | | | | | | | |
|---|---|---|---|---|---|---|---|---|---|---|---|---|---|---|---|---|---|
| AML-41-001 | *NRAS* | 1 | 115258747 | C | G | p.G12A | exonic | nonsynonymous SNV | 0.0031 | 0.0055 | 4254 | 16 | 0 | 812 | yes | ddPCR (positive) | 0 |
| AML-41-001 | *PTPN11* | 12 | 112888165 | G | C | p.D61H | exonic | nonsynonymous SNV | 0.0183 | 0.0136 | 4743 | 88 | 5 | 246 | yes | bulk-NGS | 0.010825 |
| AML-41-001 | *PTPN11* | 12 | 112888198 | G | A | p.A72T | exonic | nonsynonymous SNV | 0.0187 | 0.0173 | 4732 | 84 | 11 | 255 | yes | bulk-NGS | 0.004938 |
| AML-41-001 | *SRSF2* | 17 | 74732959 | G | T | p.P95H | exonic | nonsynonymous SNV | 0.5085 | 0.5189 | 785 | 1818 | 766 | 1713 | yes | bulk-NGS | 0.448148 |
| AML-41-001 | *TP53* | 17 | 7577577 | T | C | p.N235S | exonic | nonsynonymous SNV | 0.8933 | 0.4826 | 363 | 4245 | 295 | 179 | yes | bulk-NGS | 0.506329 |
| AML-41-001 | *WT1* | 11 | 32417947 | G | A | p.R369X | exonic | stopgain | 0.9044 | 0.6498 | 415 | 2625 | 1971 | 71 | yes | bulk-NGS | 0.625407 |
| AML-42-001 | *ASXL1* | 20 | 31022441 | A | AG | p.G642fs | exonic | frameshift insertion | 0.5244 | 0.2781 | 323 | 1178 | 3 | 748 | yes | bulk-NGS | 0.1525 |
| AML-42-001 | *EZH2* | 7 | 148506443 | C | T | p.R676H | exonic | nonsynonymous SNV | 0.8668 | 0.4577 | 235 | 1880 | 72 | 65 | yes | bulk-NGS | 0.447917 |
| AML-42-001 | *NRAS* | 1 | 115258744 | C | A | p.G13V | exonic | nonsynonymous SNV | 0.0044 | 0.0064 | 1887 | 9 | 1 | 355 | yes | ddPCR (positive) | 0 |
| AML-42-001 | *NRAS* | 1 | 115258744 | C | T | p.G13D | exonic | nonsynonymous SNV | 0.7402 | 0.435 | 230 | 1629 | 38 | 355 | yes | bulk-NGS | 0.38512 |
| AML-42-001 | *NRAS* | 1 | 115258747 | C | T | p.G12D | exonic | nonsynonymous SNV | 0.0071 | 0.0072 | 2002 | 16 | 0 | 234 | yes | bulk-NGS | 0.012862 |
| AML-42-001 | *RUNX1* | 21 | 36252937 | G | GCCGGGAA | p.A142fs | exonic | frameshift insertion | 0.0315 | 0.0194 | 2002 | 69 | 2 | 179 | yes | bulk-NGS | 0.1706 |
| AML-43-001 | *FLT3* | 13 | 28608310 | X | XAAATCAA | p.T582_G583delinsTX | exonic | stopgain | 0.8731 | 0.5067 | 906 | 10024 | 228 | 548 | yes | bulk-NGS | 0 |
| AML-43-001 | *WT1* | 11 | 32417802 | C | G | splicing | splicing | . | 0.891 | 0.4643 | 674 | 10263 | 167 | 602 | yes | bulk-NGS | 0.406103 |
| AML-43-001 | *WT1* | 11 | 32417909 | C | CGACCGTACAA | p.S381fs | exonic | frameshift insertion | 0.8287 | 0.4351 | 1338 | 9566 | 135 | 667 | yes | bulk-NGS | 0.2898 |
| AML-44-001 | *FLT3* | 13 | 28608243 | A | ACTCCCATTTGAGATCATATTCATATTCTCTGAAATCAAGGGGACCAAACTCTAAATTTTCTCTTGGAAG | p.F605delinsSPRENLEFGPLDFREYEYDLKWEF | exonic | nonframeshift insertion | 0.0012 | 0.0054 | 6702 | 23 | 0 | 34 | yes | qPCR | 0 |
| AML-44-001 | *RUNX1* | 21 | 36252867 | A | ACC | p.G165fs | exonic | frameshift insertion | 0.5084 | 0.3097 | 2272 | 3038 | 398 | 1051 | yes | bulk-NGS | 0.3046 |
| AML-44-001 | *WT1* | 11 | 32413565 | C | A | p.R462L | exonic | nonsynonymous SNV | 0.4945 | 0.2826 | 2720 | 3183 | 159 | 697 | yes | bulk-NGS | 0.288037 |
| AML-45-001 | *ASXL1* | 20 | 31022916 | G | T | p.E801X | exonic | stopgain | 0.8129 | 0.459 | 694 | 6050 | 222 | 750 | yes | bulk-NGS | 0.495082 |
| AML-45-001 | *KRAS* | 12 | 25398285 | C | G | p.G12R | exonic | nonsynonymous SNV | 0.5941 | 0.3105 | 2842 | 4405 | 179 | 290 | yes | bulk-NGS | 0.211892 |
| AML-45-001 | *NRAS* | 1 | 115258747 | C | G | p.G12A | exonic | nonsynonymous SNV | 0.0035 | 0.0019 | 6499 | 27 | 0 | 1190 | yes | ddPCR (positive) | 0 |
| AML-45-001 | *NRAS* | 1 | 115258747 | C | T | p.G12D | exonic | nonsynonymous SNV | 0.1103 | 0.0619 | 5675 | 833 | 18 | 1190 | yes | bulk-NGS | 0.055615 |
| AML-46-001 | *NPM1* | 5 | 170837543 | C | CTCTG | p.L287fs | exonic | frameshift insertion | 0.4944 | 0.467 | 597 | 3772 | 168 | 3433 | yes | bulk-NGS | 0.2099 |

| | | | | | | | | | | | | | | | | | |
|---|---|---|---|---|---|---|---|---|---|---|---|---|---|---|---|---|---|
| AML-46-001 | *PTPN11* | 12 | 112888198 | G | A | p.A72T | exonic | nonsynonymous SNV | 0.8765 | 0.4676 | 515 | 6875 | 111 | 469 | yes | bulk-NGS | 0.432075 |
| AML-47-001 | *FLT3* | 13 | 28592642 | C | A | p.D835Y | exonic | nonsynonymous SNV | 0.7993 | 0.419 | 1076 | 4955 | 238 | 228 | yes | bulk-NGS | 0.426941 |
| AML-47-001 | *NPM1* | 5 | 170837543 | C | CTCTG | p.L287fs | exonic | frameshift insertion | 0.5045 | 0.4664 | 620 | 3080 | 198 | 2599 | yes | bulk-NGS | 0.2987 |
| AML-47-001 | *WT1* | 11 | 32417910 | G | T | p.S381X | exonic | stopgain | 0.8816 | 0.4545 | 646 | 5579 | 149 | 123 | yes | bulk-NGS | 0.472112 |
| AML-48-001 | *IDH2* | 15 | 90631838 | C | T | p.R172K | exonic | nonsynonymous SNV | 0.2739 | 0.1449 | 537 | 722 | 33 | 1464 | yes | bulk-NGS | 0.110454 |
| AML-48-001 | *KRAS* | 12 | 25398279 | C | T | p.V14I | exonic | nonsynonymous SNV | 0.0878 | 0.0497 | 2420 | 229 | 13 | 94 | yes | bulk-NGS | 0.045147 |
| AML-49-001 | *FLT3* | 13 | 28608262 | X | XTCATATTCTCTGAAATCAACG | p.E598delinsDVDFREYE | exonic | nonframeshift insertion | 0.7681 | 0.8682 | 743 | 2386 | 3589 | 1061 | yes | bulk-NGS | 0.2491 |
| AML-49-001 | *NPM1* | 5 | 170837543 | C | CTCTG | p.L287fs | exonic | frameshift insertion | 0.715 | 0.462 | 595 | 5285 | 277 | 1622 | yes | bulk-NGS | 0.2599 |
| AML-49-001 | *NRAS* | 1 | 115258747 | C | T | p.G12D | exonic | nonsynonymous SNV | 0.6955 | 0.4895 | 322 | 5296 | 114 | 2047 | yes | bulk-NGS | 0.41189 |
| AML-50-001 | *FLT3* | 13 | 28608341 | T | C | p.Y572C | exonic | nonsynonymous SNV | 0.7136 | 0.4489 | 887 | 6615 | 98 | 1807 | yes | bulk-NGS | 0.41863 |
| AML-50-001 | *NPM1* | 5 | 170837543 | C | CTCTG | p.L287fs | exonic | frameshift insertion | 0.5619 | 0.4534 | 1141 | 4843 | 443 | 2980 | yes | bulk-NGS | 0.1689 |
| AML-50-001 | *RUNX1* | 21 | 36252940 | G | A | p.S141L | exonic | nonsynonymous SNV | 0.0167 | 0.0082 | 7642 | 157 | 0 | 1608 | no | ddPCR (negative) | 0 |
| AML-51-001 | *DNMT3A* | 2 | 25457243 | G | A | p.R882C | exonic | nonsynonymous SNV | 0.8547 | 0.464 | 749 | 6704 | 321 | 445 | yes | bulk-NGS | 0.464835 |
| AML-51-001 | *FLT3* | 13 | 28592642 | C | A | p.D835Y | exonic | nonsynonymous SNV | 0.2772 | 0.1345 | 5863 | 2208 | 70 | 78 | yes | bulk-NGS | 0.095135 |
| AML-51-001 | *FLT3* | 13 | 28608243 | X | XCCCATTTGAGATCATATTCATATTCTCTGAAATCAACGTAGAAGTACTCATTATT | p.F612delinsYFYVDFREYEYV | exonic | frameshift insertion | 0.1177 | 0.2997 | 6029 | 150 | 935 | 1105 | yes | bulk-NGS | 0.1268 |
| AML-51-001 | *NPM1* | 5 | 170837543 | C | CTCTG | p.L287fs | exonic | frameshift insertion | 0.6596 | 0.4077 | 847 | 5174 | 247 | 1951 | yes | bulk-NGS | 0.2262 |
| AML-51-001 | *NRAS* | 1 | 115258747 | C | A | p.G12V | exonic | nonsynonymous SNV | 0.019 | 0.0134 | 7104 | 150 | 6 | 959 | yes | bulk-NGS | 0.007447 |
| AML-51-001 | *NRAS* | 1 | 115258747 | C | T | p.G12D | exonic | nonsynonymous SNV | 0.0051 | 0.0065 | 7218 | 40 | 2 | 959 | yes | ddPCR (positive) | 0 |
| AML-52-001 | *FLT3* | 13 | 28608296 | X | XCATATTCATATTCTCTGAAAT | p.N587delinsNFREYEYD | exonic | nonframeshift insertion | 0.0814 | 0.0679 | 5477 | 513 | 25 | 595 | yes | qPCR | 0 |
| AML-52-001 | *IDH2* | 15 | 90631934 | C | T | p.R140Q | exonic | nonsynonymous SNV | 0.6165 | 0.3172 | 1964 | 3801 | 274 | 571 | yes | bulk-NGS | 0.258373 |
| AML-52-001 | *NPM1* | 5 | 170837543 | C | CTCTG | p.L287fs | exonic | frameshift insertion | 0.3495 | 0.3174 | 1582 | 2077 | 233 | 2718 | yes | bulk-NGS | 0.1143 |

| | | | | | | | | | | | | | | | | | |
|---|---|---|---|---|---|---|---|---|---|---|---|---|---|---|---|---|---|
| AML-53-001 | *FLT3* | 13 | 28608251 | X | XCATATTCTCTGAAATCAACGTAGAAGTACTCATTATCTGAGGAGCCGGTCACCTGTACCCCCC | p.M578_V579delinsIMKASYRWYRX | exonic | nonframeshift insertion | 0.0482 | 0.0531 | 7313 | 482 | 33 | 185 | yes | bulk-NGS | 0.0481 |
| AML-53-001 | *NRAS* | 1 | 115258745 | C | G | p.G13R | exonic | nonsynonymous SNV | 0.0116 | 0.0101 | 7008 | 83 | 10 | 912 | yes | ddPCR (positive) | 0 |
| AML-53-001 | *NRAS* | 1 | 115258747 | C | T | p.G12D | exonic | nonsynonymous SNV | 0.1033 | 0.059 | 6233 | 774 | 54 | 952 | yes | bulk-NGS | 0.111356 |
| AML-54-001 | *ASXL1* | 20 | 31022441 | A | AG | p.G642fs | exonic | frameshift insertion | 0.279 | 0.2585 | 1032 | 2178 | 3 | 4603 | yes | bulk-NGS | 0.2216 |
| AML-54-001 | *FLT3* | 13 | 28608224 | X | XAGAAGTACTCATTATCTGAGGAGCCGGTCCCCTGTA | p.E611delinsVQGTGSSDNEYFX | exonic | stopgain | 0.7593 | 0.5857 | 387 | 6628 | 286 | 515 | yes | bulk-NGS | 0 |
| AML-54-001 | *SRSF2* | 17 | 74732959 | G | C | p.P95R | exonic | nonsynonymous SNV | 0.2033 | 0.4677 | 546 | 1250 | 339 | 5681 | yes | bulk-NGS | 0.422642 |
| AML-55-001 | *ASXL1* | 20 | 31022402 | TCACCACTGCCATAGAGAGGCGGC | T | p.H630fs | exonic | frameshift deletion | 0.2519 | 0.0946 | 2281 | 796 | 14 | 124 | yes | bulk-NGS | 0.0621 |
| AML-55-001 | *DNMT3A* | 2 | 25457243 | G | A | p.R882C | exonic | nonsynonymous SNV | 0.2233 | 0.1048 | 2186 | 707 | 11 | 311 | yes | bulk-NGS | 0.020979 |
| AML-55-001 | *IDH1* | 2 | 209113113 | G | A | p.R132C | exonic | nonsynonymous SNV | 0.275 | 0.1213 | 2314 | 873 | 11 | 17 | yes | ddPCR (positive) | 0 |
| AML-55-001 | *IDH2* | 15 | 90631934 | C | T | p.R140Q | exonic | nonsynonymous SNV | 0.0019 | 0.0005 | 3179 | 6 | 0 | 30 | no | | 0 |
| AML-55-001 | *RUNX1* | 21 | 36252865 | C | T | p.R166Q | exonic | nonsynonymous SNV | 0.246 | 0.1601 | 2041 | 777 | 14 | 383 | yes | bulk-NGS | 0.055814 |
| AML-55-001 | *SRSF2* | 17 | 74732959 | G | T | p.P95H | exonic | nonsynonymous SNV | 0.1779 | 0.2047 | 1209 | 486 | 86 | 1434 | yes | bulk-NGS | 0.04902 |
| AML-56-001 | *IDH2* | 15 | 90631934 | C | T | p.R140Q | exonic | nonsynonymous SNV | 0.8952 | 0.4917 | 141 | 6001 | 90 | 572 | yes | bulk-NGS | 0.401544 |
| AML-57-001 | *KRAS* | 12 | 25398281 | C | T | p.G13D | exonic | nonsynonymous SNV | 0.0104 | 0.0051 | 8363 | 86 | 4 | 213 | no | | 0 |
| AML-57-001 | *NPM1* | 5 | 170837543 | C | CTCTG | p.L287fs | exonic | frameshift insertion | 0.1971 | 0.1499 | 4030 | 1646 | 62 | 2928 | yes | bulk-NGS | 0.0921 |
| AML-57-001 | *SRSF2* | 17 | 74732959 | G | C | p.P95R | exonic | nonsynonymous SNV | 0.2347 | 0.3584 | 1136 | 1687 | 347 | 5496 | yes | bulk-NGS | 0.326923 |
| AML-58-001 | *DNMT3A* | 2 | 25457242 | C | T | p.R882H | exonic | nonsynonymous SNV | 0.6151 | 0.4221 | 797 | 4912 | 113 | 2348 | yes | bulk-NGS | 0.465868 |
| AML-58-001 | *IDH2* | 15 | 90631934 | C | T | p.R140Q | exonic | nonsynonymous SNV | 0.6602 | 0.3281 | 2421 | 5293 | 101 | 355 | yes | bulk-NGS | 0.276243 |
| AML-58-001 | *SRSF2* | 17 | 74732959 | G | T | p.P95H | exonic | nonsynonymous SNV | 0.2275 | 0.4743 | 541 | 1377 | 482 | 5770 | yes | bulk-NGS | 0.448343 |
| AML-59-001 | *DNMT3A* | 2 | 25457242 | C | T | p.R882H | exonic | nonsynonymous SNV | 0.6153 | 0.294 | 488 | 1549 | 89 | 536 | yes | bulk-NGS | 0.420635 |
| AML-59-001 | *IDH1* | 2 | 209113113 | G | A | p.R132C | exonic | nonsynonymous SNV | 0.885 | 0.4484 | 278 | 2281 | 75 | 28 | yes | bulk-NGS | 0.258065 |

| | | | | | | | | | | | | | | | | | |
|---|---|---|---|---|---|---|---|---|---|---|---|---|---|---|---|---|---|
| AML-59-001 | *RUNX1* | 21 | 36231792 | C | T | p.D198N | exonic | nonsynonymous SNV | 0.0496 | 0.0351 | 1765 | 130 | 2 | 765 | yes | bulk-NGS | 0.039216 |
| AML-59-001 | *RUNX1* | 21 | 36252906 | C | CT | p.K152fs | exonic | frameshift insertion | 0.6642 | 0.3736 | 584 | 1645 | 123 | 310 | yes | bulk-NGS | 0.2295 |
| AML-60-001 | *ASXL1* | 20 | 31022441 | A | AG | p.G642fs | exonic | frameshift insertion | 0.6545 | 0.3388 | 648 | 4929 | 55 | 1983 | yes | bulk-NGS | 0.1736 |
| AML-60-001 | *FLT3* | 13 | 28608244 | X | XTCTAAATTTTCTCTTGGAAACTCCCATTTGAGATCATATTCATATTCTCTGAACCTT | p.E604delinsERFREYEYDLKWEFPRENLE | exonic | nonframeshift insertion | 0.0232 | 0.0318 | 6027 | 97 | 176 | 1315 | yes | qPCR | 0 |
| AML-60-001 | *IDH2* | 15 | 90631934 | C | T | p.R140Q | exonic | nonsynonymous SNV | 0.9368 | 0.4938 | 362 | 6798 | 336 | 119 | yes | bulk-NGS | 0.423423 |
| AML-60-001 | *SRSF2* | 17 | 74732959 | G | T | p.P95H | exonic | nonsynonymous SNV | 0.3135 | 0.5576 | 397 | 1797 | 590 | 4831 | yes | bulk-NGS | 0.444444 |
| AML-61-001 | *IDH1* | 2 | 209113112 | C | T | p.R132H | exonic | nonsynonymous SNV | 0.0037 | 0.0076 | 4272 | 16 | 0 | 52 | yes | ddPCR (positive) | 0 |
| AML-61-001 | *IDH2* | 15 | 90631934 | C | T | p.R140Q | exonic | nonsynonymous SNV | 0.1445 | 0.0603 | 3592 | 604 | 23 | 121 | yes | bulk-NGS | 0.036866 |
| AML-61-001 | *NPM1* | 5 | 170837555 | A | AT | p.R291fs | exonic | frameshift insertion | 0.2622 | 0.1701 | 1558 | 1080 | 59 | 1643 | yes | clinical sequencing | 0 |
| AML-61-001 | *NRAS* | 1 | 115258744 | C | T | p.G13D | exonic | nonsynonymous SNV | 0.0952 | 0.051 | 3376 | 391 | 22 | 551 | yes | bulk-NGS | 0.026371 |
| AML-61-001 | *PTPN11* | 12 | 112888156 | A | G | p.N58D | exonic | nonsynonymous SNV | 0.2507 | 0.1212 | 3123 | 1036 | 52 | 129 | yes | bulk-NGS | 0.075594 |
| AML-61-001 | *PTPN11* | 12 | 112926851 | C | T | p.P491S | exonic | nonsynonymous SNV | 0.1346 | 0.0738 | 3382 | 561 | 23 | 374 | yes | bulk-NGS | 0.043431 |
| AML-62-001 | *FLT3* | 13 | 28608311 | X | XCTCCGA | p.T582delinsIGA | exonic | nonframeshift insertion | 0.0618 | 0.0444 | 3112 | 238 | 11 | 666 | yes | qPCR | 0 |
| AML-62-001 | *IDH1* | 2 | 209113112 | C | T | p.R132H | exonic | nonsynonymous SNV | 0.8644 | 0.4449 | 414 | 3422 | 59 | 132 | yes | bulk-NGS | 0.468966 |
| AML-62-001 | *NPM1* | 5 | 170837543 | C | CTCTG | p.L287fs | exonic | frameshift insertion | 0.4467 | 0.4438 | 324 | 1758 | 41 | 1904 | yes | bulk-NGS | 0.2195 |
| AML-62-001 | *PTPN11* | 12 | 112888166 | A | C | p.D61A | exonic | nonsynonymous SNV | 0.0079 | 0.0036 | 3838 | 32 | 0 | 157 | yes | ddPCR (positive) | 0 |
| AML-62-001 | *PTPN11* | 12 | 112888195 | T | C | p.F71L | exonic | nonsynonymous SNV | 0.4246 | 0.2068 | 2119 | 1682 | 28 | 198 | yes | bulk-NGS | 0.226381 |
| AML-62-001 | *PTPN11* | 12 | 112888198 | G | A | p.A72T | exonic | nonsynonymous SNV | 0.3496 | 0.1713 | 2416 | 1383 | 25 | 203 | yes | bulk-NGS | 0.166963 |
| AML-63-001 | *FLT3* | 13 | 28608303 | X | XAGTTTCTCTTGGAA | p.S585fs | exonic | frameshift insertion | 0.101 | 0.097 | 6484 | 914 | 44 | 905 | yes | bulk-NGS | 0 |
| AML-63-001 | *IDH2* | 15 | 90631934 | C | T | p.R140Q | exonic | nonsynonymous SNV | 0.8404 | 0.4693 | 663 | 6709 | 306 | 669 | yes | bulk-NGS | 0.46755 |
| AML-63-001 | *KIT* | 4 | 55599321 | A | T | p.D816V | exonic | nonsynonymous SNV | 0.1059 | 0.0508 | 7120 | 827 | 57 | 343 | yes | bulk-NGS | 0.060325 |
| AML-63-001 | *NPM1* | 5 | 170837543 | C | CTCTG | p.L287fs | exonic | frameshift insertion | 0.5135 | 0.4534 | 734 | 3921 | 365 | 3327 | yes | bulk-NGS | 0.1458 |
| AML-64-001 | *DNMT3A* | 2 | 25457242 | C | T | p.R882H | exonic | nonsynonymous SNV | 0.6713 | 0.3639 | 778 | 3255 | 101 | 865 | yes | bulk-NGS | 0.392265 |

| | | | | | | | | | | | | | | | | | |
|---|---|---|---|---|---|---|---|---|---|---|---|---|---|---|---|---|---|
| AML-64-001 | *IDH1* | 2 | 209113113 | G | A | p.R132C | exonic | nonsynonymous SNV | 0.0014 | 0.0004 | 4970 | 7 | 0 | 22 | no | ddPCR (negative) | 0 |
| AML-64-001 | *IDH2* | 15 | 90631934 | C | T | p.R140Q | exonic | nonsynonymous SNV | 0.8056 | 0.4043 | 827 | 3910 | 117 | 145 | yes | bulk-NGS | 0.464078 |
| AML-64-001 | *NRAS* | 1 | 115256530 | G | T | p.Q61K | exonic | nonsynonymous SNV | 0.0174 | 0.0069 | 4893 | 80 | 7 | 19 | yes | bulk-NGS | 0.00753 |
| AML-64-001 | *SRSF2* | 17 | 74732959 | G | T | p.P95H | exonic | nonsynonymous SNV | 0.4579 | 0.4834 | 549 | 1843 | 446 | 2161 | yes | bulk-NGS | 0.407407 |
| AML-65-001 | *DNMT3A* | 2 | 25457242 | C | T | p.R882H | exonic | nonsynonymous SNV | 0.7823 | 0.426 | 1117 | 5358 | 308 | 460 | yes | bulk-NGS | 0.454545 |
| AML-65-001 | *FLT3* | 13 | 28608274 | X | XAAATCAACGTAGAAGTACTCATTA | p.F594delinsFNEYFYVDF | exonic | nonframeshift insertion | 0.5873 | 0.7227 | 1981 | 1599 | 2663 | 1000 | yes | bulk-NGS | 0 |
| AML-66-001 | *ASXL1* | 20 | 31022441 | A | AG | p.G642fs | exonic | frameshift insertion | 0.5189 | 0.2737 | 1024 | 3750 | 20 | 2471 | yes | bulk-NGS in paired sample | not tested |
| AML-66-001 | *KRAS* | 12 | 25398284 | C | T | p.G12D | exonic | nonsynonymous SNV | 0.0162 | 0.0121 | 7027 | 111 | 7 | 120 | no | | not tested |
| AML-66-001 | *NRAS* | 1 | 115256529 | T | G | p.Q61P | exonic | nonsynonymous SNV | 0.0134 | 0.0174 | 7050 | 94 | 3 | 118 | no | | not tested |
| AML-66-001 | *NRAS* | 1 | 115258744 | C | A | p.G13V | exonic | nonsynonymous SNV | 0.0139 | 0.011 | 6260 | 98 | 3 | 904 | no | | not tested |
| AML-66-001 | *NRAS* | 1 | 115258747 | C | T | p.G12D | exonic | nonsynonymous SNV | 0.0187 | 0.0136 | 6221 | 133 | 3 | 908 | no | | not tested |
| AML-66-001 | *NRAS* | 1 | 115258748 | C | T | p.G12S | exonic | nonsynonymous SNV | 0.2208 | 0.129 | 4646 | 1562 | 42 | 1015 | yes | bulk-NGS in paired sample | not tested |
| AML-66-001 | *PTPN11* | 12 | 112926888 | G | A | p.G503E | exonic | nonsynonymous SNV | 0.5217 | 0.3048 | 2515 | 3662 | 128 | 960 | yes | bulk-NGS in paired sample | not tested |
| AML-66-001 | *SRSF2* | 17 | 74732959 | G | A | p.P95L | exonic | nonsynonymous SNV | 0.0728 | 0.2165 | 766 | 454 | 75 | 5970 | yes | bulk-NGS in paired sample | not tested |
| AML-67-001 | *ASXL1* | 20 | 31023108 | G | A | p.E865K | exonic | nonsynonymous SNV | 0.0116 | 0.0085 | 5475 | 70 | 0 | 479 | no | ddPCR (negative) | 0 |
| AML-67-001 | *IDH1* | 2 | 209113113 | G | A | p.R132C | exonic | nonsynonymous SNV | 0.0012 | 0.0004 | 5989 | 7 | 0 | 28 | no | ddPCR (negative) | 0 |
| AML-67-001 | *IDH2* | 15 | 90631934 | C | T | p.R140Q | exonic | nonsynonymous SNV | 0.1703 | 0.0713 | 4879 | 1005 | 21 | 119 | yes | bulk-NGS | 0.085526 |
| AML-67-001 | *KRAS* | 12 | 25380275 | T | A | p.Q61H | exonic | nonsynonymous SNV | 0.3025 | 0.1611 | 3676 | 1758 | 64 | 526 | yes | bulk-NGS | 0.219024 |
| AML-67-001 | *KRAS* | 12 | 25380279 | C | A | p.G60V | exonic | nonsynonymous SNV | 0.0023 | 0.0012 | 5579 | 13 | 1 | 431 | yes | ddPCR (positive) | 0 |
| AML-67-001 | *NRAS* | 1 | 115258748 | C | T | p.G12S | exonic | nonsynonymous SNV | 0.3958 | 0.207 | 3051 | 2351 | 33 | 589 | yes | bulk-NGS | 0.172414 |
| AML-67-001 | *RUNX1* | 21 | 36171613 | A | ACC | p.S318fs | exonic | frameshift insertion | 0.2729 | 0.1503 | 3899 | 1438 | 206 | 481 | no | ddPCR (negative) | 0 |
| AML-67-001 | *RUNX1* | 21 | 36171614 | A | AAGG | p.L317delinsLL | exonic | nonframeshift insertion | 0.2726 | 0.1495 | 3929 | 1434 | 208 | 453 | no | ddPCR (negative) | 0 |

| | | | | | | | | | | | | | | | | | |
|---|---|---|---|---|---|---|---|---|---|---|---|---|---|---|---|---|---|
| AML-67-001 | *TP53* | 17 | 7578245 | G | A | p.R202C | exonic | nonsynonymous SNV | 0.0015 | 0.0007 | 5931 | 9 | 0 | 84 | yes | clinical sequencing | 0 |
| AML-67-001 | *U2AF1* | 21 | 44514777 | T | C | p.Q84R | exonic | nonsynonymous SNV | 0.6716 | 0.4593 | 123 | 3983 | 63 | 1855 | yes | bulk-NGS | 0.406114 |
| AML-67-001 | *U2AF1* | 21 | 44524456 | G | A | p.S34F | exonic | nonsynonymous SNV | 0.7656 | 0.3833 | 1247 | 4547 | 65 | 165 | yes | bulk-NGS | 0.358531 |
| AML-68-001 | *DNMT3A* | 2 | 25457242 | C | T | p.R882H | exonic | nonsynonymous SNV | 0.599 | 0.3811 | 1018 | 3825 | 85 | 1600 | yes | bulk-NGS | 0.396146 |
| AML-68-001 | *FLT3* | 13 | 28608275 | X | XGATCATATTCATATTCTCTGAAATCAACGTAGAAGTACTCAT | p.G583_S584delinsGRIX | exonic | nonframeshift insertion | 0.6351 | 0.3723 | 1471 | 4511 | 40 | 506 | yes | bulk-NGS | 0.1905 |
| AML-68-001 | *KRAS* | 12 | 25398285 | C | T | p.G12S | exonic | nonsynonymous SNV | 0.0487 | 0.03 | 5709 | 301 | 17 | 501 | yes | bulk-NGS | 0.021823 |
| AML-68-001 | *NPM1* | 5 | 170837543 | C | CTCTG | p.L287fs | exonic | frameshift insertion | 0.4205 | 0.4077 | 813 | 2633 | 112 | 2970 | yes | bulk-NGS | 0.2362 |
| AML-69-001 | *KRAS* | 12 | 25398284 | C | G | p.G12A | exonic | nonsynonymous SNV | 0.0293 | 0.0147 | 7030 | 203 | 16 | 213 | yes | bulk-NGS | 0.007968 |
| AML-69-001 | *NRAS* | 1 | 115258744 | C | T | p.G13D | exonic | nonsynonymous SNV | 0.5702 | 0.3224 | 2357 | 4115 | 140 | 850 | yes | bulk-NGS | 0.305353 |
| AML-69-001 | *RUNX1* | 21 | 36171600 | G | C | p.S322X | exonic | stopgain | 0.0741 | 0.0574 | 4927 | 377 | 176 | 1982 | yes | bulk-NGS | 0.065306 |
| AML-69-001 | *TP53* | 17 | 7578290 | C | T | splicing | splicing | . | 0.0039 | 0.0026 | 7216 | 29 | 0 | 217 | no | ddPCR (negative) | 0 |
| AML-70-001 | *IDH2* | 15 | 90631934 | C | T | p.R140Q | exonic | nonsynonymous SNV | 0.8382 | 0.4402 | 1033 | 6660 | 270 | 305 | yes | bulk-NGS | 0.5 |
| AML-70-001 | *KRAS* | 12 | 25398220 | A | T | p.D33E | exonic | nonsynonymous SNV | 0.7916 | 0.4336 | 1269 | 6070 | 475 | 454 | yes | bulk-NGS | 0.3125 |
| AML-70-001 | *NRAS* | 1 | 115256532 | C | T | p.G60E | exonic | nonsynonymous SNV | 0.0195 | 0.0089 | 8011 | 152 | 9 | 96 | yes | clinical sequencing | 0 |
| AML-70-001 | *SRSF2* | 17 | 74732959 | G | A | p.P95L | exonic | nonsynonymous SNV | 0.1451 | 0.2368 | 1053 | 1079 | 121 | 6015 | yes | bulk-NGS | 0.454369 |
| AML-71-001 | *NRAS* | 1 | 115258748 | C | G | p.G12R | exonic | nonsynonymous SNV | 0.8012 | 0.491 | 309 | 6703 | 127 | 1386 | yes | bulk-NGS | 0.452731 |
| AML-71-001 | *RUNX1* | 21 | 36171664 | G | GCGGA | p.P301fs | exonic | frameshift insertion | 0.0304 | 0.0206 | 6779 | 202 | 57 | 1487 | yes | bulk-NGS | 0.0672 |
| AML-72-001 | *NPM1* | 5 | 170837543 | C | CTCTG | p.L287fs | exonic | frameshift insertion | 0.5444 | 0.4558 | 1033 | 4993 | 334 | 3425 | yes | bulk-NGS | 0.2938 |
| AML-72-001 | *PTPN11* | 12 | 112888210 | G | C | p.E76Q | exonic | nonsynonymous SNV | 0.8528 | 0.4653 | 880 | 8032 | 313 | 560 | yes | bulk-NGS | 0.437186 |
| AML-73-001 | *FLT3* | 13 | 28609758 | C | A | p.V491L | exonic | nonsynonymous SNV | 0.4392 | 0.2089 | 5496 | 4338 | 72 | 136 | yes | bulk-NGS | 0.204962 |
| AML-73-001 | *KRAS* | 12 | 25380276 | T | A | p.Q61L | exonic | nonsynonymous SNV | 0.007 | 0.0063 | 8860 | 70 | 0 | 1112 | no | | 0 |
| AML-73-001 | *KRAS* | 12 | 25380276 | T | C | p.Q61R | exonic | nonsynonymous SNV | 0.0041 | 0.0034 | 8860 | 41 | 0 | 1141 | no | | 0 |
| AML-73-001 | *SF3B1* | 2 | 198267359 | C | G | p.K666N | exonic | nonsynonymous SNV | 0.944 | 0.4927 | 437 | 9166 | 314 | 125 | yes | bulk-NGS | 0.505938 |
| AML-74-001 | *KRAS* | 12 | 25380279 | C | T | p.G60D | exonic | nonsynonymous SNV | 0.0141 | 0.008 | 8030 | 129 | 2 | 1118 | yes | bulk-NGS | 0.009544 |

| | | | | | | | | | | | | | | | | | |
|---|---|---|---|---|---|---|---|---|---|---|---|---|---|---|---|---|---|
| AML-74-001 | *NPM1* | 5 | 170837543 | C | CTCTG | p.L287fs | exonic | frameshift insertion | 0.2974 | 0.2593 | 2466 | 2681 | 79 | 4053 | yes | bulk-NGS | 0.1659 |
| AML-74-001 | *NRAS* | 1 | 115258747 | C | T | p.G12D | exonic | nonsynonymous SNV | 0.0258 | 0.0137 | 7970 | 234 | 5 | 1070 | yes | bulk-NGS | 0.014909 |
| AML-74-001 | *PTPN11* | 12 | 112888166 | A | C | p.D61A | exonic | nonsynonymous SNV | 0.075 | 0.0359 | 8317 | 680 | 16 | 266 | yes | bulk-NGS | 0.021954 |
| AML-74-001 | *WT1* | 11 | 32417910 | G | T | p.S381X | exonic | stopgain | 0.0136 | 0.0058 | 9030 | 123 | 3 | 123 | yes | clinical sequencing | 0 |
| AML-75-001 | *EZH2* | 7 | 148506432 | G | A | p.H680Y | exonic | nonsynonymous SNV | 0.1798 | 0.0824 | 4586 | 1047 | 45 | 395 | yes | bulk-NGS | 0.1152 |
| AML-75-001 | *NPM1* | 5 | 170837543 | C | CTCTG | p.L287fs | exonic | frameshift insertion | 0.0845 | 0.0548 | 4081 | 494 | 19 | 1479 | yes | bulk-NGS | 0.0276 |
| AML-75-001 | *NRAS* | 1 | 115258748 | C | G | p.G12R | exonic | nonsynonymous SNV | 0.1841 | 0.0911 | 4561 | 1077 | 41 | 394 | yes | bulk-NGS | 0.081051 |
| AML-75-001 | *SRSF2* | 17 | 74732959 | G | T | p.P95H | exonic | nonsynonymous SNV | 0.2621 | 0.3264 | 1317 | 1269 | 323 | 3164 | yes | bulk-NGS | 0.226829 |
| AML-76-001 | *DNMT3A* | 2 | 25457242 | C | G | p.R882P | exonic | nonsynonymous SNV | 0.6892 | 0.4534 | 567 | 5365 | 171 | 1930 | yes | bulk-NGS | 0.477361 |
| AML-76-001 | *FLT3* | 13 | 28592629 | T | C | p.D839G | exonic | nonsynonymous SNV | 0.0055 | 0.0027 | 7887 | 43 | 1 | 102 | yes | ddPCR (positive) | 0 |
| AML-76-001 | *FLT3* | 13 | 28592634 | CATG | C | p.836_837del | exonic | nonframeshift deletion | 0.0193 | 0.008 | 7795 | 153 | 2 | 83 | yes | bulk-NGS | 0.0413 |
| AML-76-001 | *FLT3* | 13 | 28608309 | X | XTCTGAAAT | p.G583fs | exonic | frameshift insertion | 0.1145 | 0.0841 | 5843 | 889 | 31 | 1270 | yes | qPCR | 0 |
| AML-76-001 | *NRAS* | 1 | 115256528 | T | A | p.Q61H | exonic | nonsynonymous SNV | 0.7005 | 0.3621 | 2233 | 5385 | 242 | 173 | yes | bulk-NGS | 0.347403 |
| AML-76-001 | *NRAS* | 1 | 115258745 | C | G | p.G13R | exonic | nonsynonymous SNV | 0.0065 | 0.0033 | 7218 | 52 | 0 | 763 | yes | ddPCR (positive) | 0 |
| AML-76-001 | *NRAS* | 1 | 115258747 | C | G | p.G12A | exonic | nonsynonymous SNV | 0.0168 | 0.008 | 7127 | 132 | 3 | 771 | yes | bulk-NGS | 0.008649 |
| AML-76-001 | *U2AF1* | 21 | 44524456 | G | A | p.S34F | exonic | nonsynonymous SNV | 0.8486 | 0.4738 | 554 | 6576 | 241 | 662 | yes | bulk-NGS | 0.455726 |
| AML-77-001 | *ASXL1* | 20 | 31022441 | A | AG | p.G642fs | exonic | frameshift insertion | 0.8508 | 0.3462 | 475 | 7337 | 54 | 821 | yes | bulk-NGS | 0.3333 |
| AML-77-001 | *IDH2* | 15 | 90631838 | C | T | p.R172K | exonic | nonsynonymous SNV | 0.9512 | 0.5055 | 351 | 7951 | 312 | 73 | yes | bulk-NGS | 0.537915 |
| AML-77-001 | *NRAS* | 1 | 115258747 | C | A | p.G12V | exonic | nonsynonymous SNV | 0.0031 | 0.004 | 7698 | 27 | 0 | 962 | yes | ddPCR (positive) | 0 |
| AML-77-001 | *U2AF1* | 21 | 44514777 | T | C | p.Q84R | exonic | nonsynonymous SNV | 0.9687 | 0.4939 | 216 | 8239 | 176 | 56 | yes | bulk-NGS | 0.451372 |

**Abbreviations:** chr, chromosome; ref allele, reference allele; alt allele, alternate allele; scDNA-seq VAF, variant allele frequency (VAF) based on single-cell DNA sequencing; WT, wild type; Het, heterozygous; Homo, homozygous; Missing, missing genotype; bulk-NGS, next-generation sequencing using the bulk sample; bulk VAF, VAF based on bulk-NGS.

**Supplementary Table 4.** List of 50 amplicons covered by the single-cell DNA sequencing panel.

| Amplicon | Chr | Primer Start (based on hg19) | Insert Start (based on hg19) | Insert End (based on hg19) | Primer End (based on hg19) | Amplicon Length (without primer) [bp] | Amplicon Length (with primer) [bp] | Primer length [bp] |
|---|---|---|---|---|---|---|---|---|
| *ASXL1_1* | 20 | 31022348 | 31022369 | 31022586 | 31022608 | 217 | 260 | 43 |
| *ASXL1_2_a1* | 20 | 31022880 | 31022899 | 31023110 | 31023130 | 211 | 250 | 39 |
| *DNMT3A_10* | 2 | 25457114 | 25457135 | 25457351 | 25457372 | 216 | 258 | 42 |
| *EZH2_1* | 7 | 148504627 | 148504654 | 148504874 | 148504901 | 220 | 274 | 54 |
| *EZH2_2* | 7 | 148506303 | 148506332 | 148506547 | 148506577 | 215 | 274 | 59 |
| *FLT3_1* | 13 | 28592473 | 28592494 | 28592723 | 28592747 | 229 | 274 | 45 |
| *FLT3_2_a3* | 13 | 28608168 | 28608191 | 28608368 | 28608392 | 177 | 224 | 47 |
| *FLT3_3* | 13 | 28602155 | 28602179 | 28602404 | 28602429 | 225 | 274 | 49 |
| *FLT3_4* | 13 | 28609521 | 28609547 | 28609769 | 28609795 | 222 | 274 | 52 |
| *FLT3_5_4* | 13 | 28607997 | 28608018 | 28608153 | 28608176 | 135 | 179 | 44 |
| *GATA2_1* | 3 | 128202704 | 128202723 | 128202891 | 128202911 | 168 | 207 | 39 |
| *IDH1_1* | 2 | 209112875 | 209112898 | 209113123 | 209113149 | 225 | 274 | 49 |
| *IDH2_1_4* | 15 | 90631738 | 90631759 | 90631985 | 90632009 | 226 | 271 | 45 |
| *JAK2_1* | 9 | 5073541 | 5073563 | 5073785 | 5073815 | 222 | 274 | 52 |
| *KIT_1* | 4 | 55599204 | 55599231 | 55599448 | 55599478 | 217 | 274 | 57 |
| *KIT_2* | 4 | 55589585 | 55589607 | 55589829 | 55589859 | 222 | 274 | 52 |
| *KRAS_1* | 12 | 25398161 | 25398183 | 25398405 | 25398435 | 222 | 274 | 52 |
| *KRAS_2* | 12 | 25380238 | 25380259 | 25380466 | 25380490 | 207 | 252 | 45 |
| *NPM1_1_2* | 5 | 170837385 | 170837412 | 170837636 | 170837659 | 224 | 274 | 50 |
| *NRAS_1* | 1 | 115256296 | 115256324 | 115256546 | 115256570 | 222 | 274 | 52 |
| *NRAS_2* | 1 | 115258525 | 115258553 | 115258776 | 115258799 | 223 | 274 | 51 |
| *PTPN11_1_1* | 12 | 112926827 | 112926848 | 112927043 | 112927063 | 195 | 236 | 41 |
| *PTPN11_2* | 12 | 112888095 | 112888116 | 112888327 | 112888351 | 211 | 256 | 45 |
| *RUNX1_2* | 21 | 36171458 | 36171482 | 36171710 | 36171732 | 228 | 274 | 46 |
| *RUNX1_3* | 21 | 36206684 | 36206705 | 36206884 | 36206907 | 179 | 223 | 44 |
| *RUNX1_4* | 21 | 36231583 | 36231604 | 36231834 | 36231857 | 230 | 274 | 44 |
| *RUNX1_5* | 21 | 36164768 | 36164785 | 36164997 | 36165018 | 212 | 250 | 38 |
| *RUNX1_7* | 21 | 36252789 | 36252811 | 36253007 | 36253030 | 196 | 241 | 45 |
| *SF3B1_1* | 2 | 198266733 | 198266761 | 198266977 | 198267007 | 216 | 274 | 58 |
| *SF3B1_2* | 2 | 198267134 | 198267156 | 198267384 | 198267406 | 228 | 272 | 44 |
| *SRSF2_2_2* | 17 | 74732865 | 74732882 | 74733050 | 74733069 | 168 | 204 | 36 |
| *TP53_1* | 17 | 7578062 | 7578086 | 7578296 | 7578319 | 210 | 257 | 47 |
| *TP53_2* | 17 | 7577376 | 7577397 | 7577615 | 7577637 | 218 | 261 | 43 |
| *TP53_3* | 17 | 7578363 | 7578384 | 7578605 | 7578627 | 221 | 264 | 43 |
| *TP53_4* | 17 | 7576930 | 7576953 | 7577180 | 7577204 | 227 | 274 | 47 |
| *U2AF1_1* | 21 | 44514679 | 44514700 | 44514922 | 44514947 | 222 | 268 | 46 |
| *U2AF1_2* | 21 | 44524258 | 44524279 | 44524505 | 44524532 | 226 | 274 | 48 |
| *WT1_1_a2* | 11 | 32414174 | 32414193 | 32414411 | 32414432 | 218 | 258 | 40 |
| *WT1_2* | 11 | 32413389 | 32413415 | 32413641 | 32413663 | 226 | 274 | 48 |

| | | | | | | | | |
|---|---|---|---|---|---|---|---|---|
| *WT1_3* | 11 | 32417744 | 32417765 | 32417992 | 32418018 | 227 | 274 | 47 |
| chr10_106721610 | 10 | 106721487 | 106721508 | 106721711 | 106721736 | 203 | 249 | 46 |
| chr10_5554293 | 10 | 5554171 | 5554192 | 5554401 | 5554419 | 209 | 248 | 39 |
| chr10_77210191 | 10 | 77210064 | 77210083 | 77210294 | 77210313 | 211 | 249 | 38 |
| chr14_56969005 | 14 | 56968884 | 56968905 | 56969106 | 56969129 | 201 | 245 | 44 |
| chr16_55770629 | 16 | 55770512 | 55770529 | 55770735 | 55770757 | 206 | 245 | 39 |
| chr16_8569820 | 16 | 8569695 | 8569720 | 8569926 | 8569944 | 206 | 249 | 43 |
| chr18_9750662 | 18 | 9750543 | 9750561 | 9750767 | 9750791 | 206 | 248 | 42 |
| chr6_17076840 | 6 | 17076720 | 17076739 | 17076941 | 17076969 | 202 | 249 | 47 |
| chr6_40116264 | 6 | 40116143 | 40116164 | 40116366 | 40116388 | 202 | 245 | 43 |
| chr6_62094287 | 6 | 62094166 | 62094187 | 62094388 | 62094411 | 201 | 245 | 44 |

**Abbreviations:** Chr, chromosome; bp, base pair.

**Supplementary Table 5.** List of 295 genes targeted by bulk next-generation sequencing.

| **Gene name** | | | | | | | | | |
|---|---|---|---|---|---|---|---|---|---|
| *ABCC9* | *CALR* | *CUL5* | *FANCD2* | *HIST1H2BF* | *LEF1* | *NBN* | *PLA2G2D* | *SF3B1* | *TINF2 (TIN2)* |
| *ABL1* | *CARD11* | *CUX1* | *FANCE* | *HIST1H3D* | *LRP1B* | *NCOR1* | *PLCG2* | *SFRS1* | *TLR2* |
| *ACTG1* | *CBL* | *CYLD* | *FANCG* | *HIST1H4D* | *LTB* | *NCOR2* | *POT1* | *SFRS7* | *TLR9* |
| *AKT1* | *CBLB* | *DAXX* | *FANCI* | *HNRNPK* | *LUC7L2* | *NF1* | *POU2AF1* | *SGK1* | *TNFAIP3* |
| *ANKRD11* | *CCND1* | *DCLRE1C* | *FANCL* | *HRAS* | *LYN* | *NFE2* | *PRDM1* | *SH2B3* | *TNFRSF14* |
| *ARID1A* | *CCND3* | *DDX3X* | *FAS* | *ICOS* | *MALT1* | *NFKB1* | *PRKCB* | *SHH* | *TNKS* |
| *ARID1B* | *CD200* | *DIS3* | *FAT1* | *ID3* | *MAP2K1* | *NFKB2* | *PTEN* | *SMAD2* | *TOX* |
| *ARID2* | *CD274* | *DKC1* | *FAT3* | *IDH1* | *MAPK1* | *NFKBIA* | *PTPN1* | *SMC1A* | *TP53* |
| *ARID5B* | *CD58* | *DLC1* | *FBXW7* | *IDH2* | *MAX* | *NFKBIE* | *PTPN11* | *SMC3* | *TRAF3* |
| *ARPP21* | *CD79A* | *DNM2* | *FGFR3* | *IKBKA* | *MDM2* | *NOTCH1* | *RAD21* | *SMC5* | *TRAF6* |
| *ASXL1* | *CD79B* | *DNMT1* | *FLI1* | *IKZF1* | *MED12* | *NOTCH2* | *RAD51C* | *SNX7* | *TYK2* |
| *ATF7IP* | *CDK4* | *DNMT3A* | *FLT3* | *IKZF2* | *MEF2B* | *NPM1* | *RAG1* | *SOCS1* | *TYK3* |
| *ATM* | *CDKN2A* | *DNMT3B* | *FNDC3A* | *IKZF3* | *MEF2C* | *NR3C2* | *RAG2* | *SOX5* | *U2AF1* |
| *ATRX* | *CDKN2B* | *EBF1* | *FOXP1* | *IL7R* | *MGA* | *NRAS* | *RASA2* | *SP140* | *U2AF2* |
| *B2M* | *CDKN2C* | *ECT2L* | *FYN* | *IRAK1* | *miR125a* | *NSD2* | *RB1* | *SPEN* | *UBR5* |
| *BCL10* | *CEBPA* | *EED* | *G6PC3* | *IRAK4* | *miR-142* | *NT5C2* | *REL* | *SPIB* | *USP29* |
| *BCL2* | *CEBPE* | *EGR1* | *GAB2* | *IRF1* | *mIR155* | *PAG1* | *RELA* | *SRSF2* | *VPREB1* |
| *BCL6* | *CHD2* | *EGR2* | *GATA1* | *IRF4* | *mIR15a* | *PALB2* | *RELB* | *STAG1* | *WHSC1* |
| *BCL7A* | *CHK2* | *ELANE* | *GATA2* | *IRF7* | *mIR16-1* | *PAX5* | *RELN* | *STAG2* | *WHSC1L1* |
| *BCOR* | *CIITA* | *EP300* | *GATA3* | *ITPKB* | *MIR17HG* | *PDCD1* | *RHOA* | *STAT1* | *WT1* |
| *BCR* | *CNOT3* | *EPHA7* | *GCET2* | *JAK1* | *mIR21* | *PDCD1LG2* | *RIPK1* | *STAT3* | *XPO1* |
| *BIRC3* | *CREBBP* | *EPOR* | *GFI1B* | *JAK2* | *mir34b* | *PDGFRB* | *ROBO1* | *SUZ12* | *ZAP70* |
| *BLK* | *CRLF2* | *ERG* | *GNA13* | *JAK3* | *mir34c* | *PEG3* | *ROR1* | *SYK* | *ZMYM2* |
| *BMI1* | *CSF2RA* | *ETV6* | *GNAS* | *JARID2* | *MLL* | *PHF6* | *RPL10* | *TBL1XR1* | *ZMYM3* |
| *BRAF* | *CSF3R* | *EZH2* | *GNB1* | *KDM4C* | *MLL2* | *PHIP* | *RPL5* | *TCF3* | *ZRSR2* |
| *BRIP1* | *CTBP1* | *FAM46C* | *GPRC5A* | *KDM6A* | *MLL3* | *PIGA* | *RUNX1* | *TERC* | |
| *BTG1* | *CTBP2* | *FAM5C* | *HAX1* | *KIT* | *MPL* | *PIK3CA* | *RUNX2* | *TERT* | |
| *BTK* | *CTCF* | *FANCA* | *HIST1H1E* | *KLHL6* | *MS4A1* | *PIK3CB* | *SAMHD1* | *TET1* | |
| *BTLA* | *CTLA4* | *FANCB* | *HIST1H2AD* | *KRAS* | *MYB* | *PIK3CG* | *SETBP1* | *TET2* | |
| *C22orf194* | *CTNNA1* | *FANCC* | *HIST1H2BE* | *LAMB4* | *MYD88* | *PIK3R1* | *SETD2* | *TGDS* | |

**Supplementary Table 6.** List of probes/primers used for ddPCR assay.

| primer/probe ID | sequence and chromosome coordinates of the assay's amplicon | Variant |
|---|---|---|
| dHsaMDS708288136 | hg19\|chr20:31023047-31023169:+ CCCCAGTTCCACACCTGAATCCTCACCGACTGATTGCCTGCAG AACAGAGCATTTGATGAC[G/A]AATTAGGGCTTGGTGGCTCAT GCCCTCCTATGAGGGAAAGTGATACTAGACAAGAAAACTT | *ASXL1*:exon13:c.G2593A:p.E865K |
| dHsaMDS299498199 | hg19\|chr13:28602279-28602401:- AGGCACTCATGTCAGAACTCAAGATGATGACCCAGCTGGGAA GCCACGAGAATATTGTGAA[C/A]CTGCTGGGGGCGTGCACAC TGTCAGGTAACCCACTTCCACGAAAATCACCTCATCAAAAAG | *FLT3*:exon16:c.C2028A:p.N676K |
| dHsaMDS887479234 | hg19\|chr13:28602268-28602390:- TCAGAACTCAAGATGATGACCCAGCTGGGAAGCCACGAGAAT ATTGTGAACCTGCTGGGGG[C/T]GTGCACACTGTCAGGTAACC CACTTCCACGAAAATCACCTCATCAAAAAGACTGTAGCTTG | *FLT3*:exon16:c.C2039T:p.A680V |
| dHsaMDS600877565 | hg19\|chr13:28592568-28592690:- GTCACCCACGGGAAAGTGGTGAAGATATGTGACTTTGGATTG GCTCGAGATATCATGAGTG[A/G]TTCCAACTATGTTGTCAGGG GCAATGTGAGGCTGCTATTTCCTACTTATTTTTATACGGCT | *FLT3*:exon20:c.A2516G:p.D839G |
| dHsaMDV2010053 | hg19\|chr2:209113052-209113174:- CATTATCTGCAAAAATATCCCCCGGCTTGTGAGTGGATGGGTA AAACCTATCATCATAGGT[C/T]GTCATGCTTATGGGGATCAAG TAAGTCATGTTGGCAATAATGTGATTTTGCATGTTTTTTT | *IDH1*:exon4:c.C394T:p.R132C |
| dHsaMDV2010055 | hg19\|chr2:209113051-209113173:- ATTATCTGCAAAAATATCCCCCGGCTTGTGAGTGGATGGGTAA AACCTATCATCATAGGTC[G/A]TCATGCTTATGGGGATCAAGT AAGTCATGTTGGCAATAATGTGATTTTGCATGTTTTTTTT | *IDH1*:exon4:c.G395A:p.R132H |
| dHsaMDV2010057 | hg19\|chr15:90631873-90631995:- ATCTCTGTCCTCACAGAGTTCAAGCTGAAGAAGATGTGGAAA AGTCCCAATGGAACTATCC[G/A]GAACATCCTGGGGGGGACT GTCTTCCGGGAGCCCATCATCTGCAAAAACATCCCACGCCTA | *IDH2*:exon4:c.G419A:p.R140Q |
| dHsaMDV2510596 | hg19\|chr12:25398223-25398345:- TTATTTTTATTATAAGGCCTGCTGAAAATGACTGAATATAAACT TGTGGTAGTTGGAGCTG[G/A]TGGCGTAGGCAAGAGTGCCTT GACGATACAGCTAATTCAGAATCATTTTGTGGACGAATAT | *KRAS*:exon2:c.G35A:p.G12D |

| | | |
|---|---|---|
| dHsaMDV2510586 | hg19\|chr12:25398223-25398345:-<br>TTATTTTTATTATAAGGCCTGCTGAAAATGACTGAATATAAACT<br>TGTGGTAGTTGGAGCTG[G/C]TGGCGTAGGCAAGAGTGCCTT<br>GACGATACAGCTAATTCAGAATCATTTTGTGGACGAATAT | *KRAS*:exon2:c.G35C:p.G12A |
| dHsaMDS589521946 | hg19\|chr12:25380218-25380340:-<br>TACAGGAAGCAAGTAGTAATTGATGGAGAAACCTGTCTCTTG<br>GATATTCTCGACACAGCAG[G/T]TCAAGAGGAGTACAGTGCA<br>ATGAGGGACCAGTACATGAGGACTGGGGAGGGCTTTCTTTGT | *KRAS*:exon3:c.G179T:p.G60V |
| dHsaMDS890511303 | hg19\|chr5:170837482-170837604:+<br>TATGAAGTGTTGTGGTTCCTTAACCACATTTCTTTTTTTTTTTTT<br>CCAGGCTATTCAAGAT[C/CTCTG]TCTGGCAGTGGAGGAAGTC<br>TCTTTAAGAAAATAGTTTAAACAATTTGTTAAAAAATTTTCC | *NPM1*:exon11:c.859_860insTCTG:<br>p.L287fs |
| dHsaMDV2010093 | hg19\|chr1:115258687-115258809:-<br>GTTTCCAACAGGTTCTTGCTGGTGTGAAATGACTGAGTACAAA<br>CTGGTGGTGGTTGGAGCA[G/A]GTGGTGTTGGGAAAAGCGC<br>ACTGACAATCCAGCTAATCCAGAACCACTTTGTAGATGAATA | *NRAS*:exon2:c.G34A:p.G12S |
| dHsaMDV2010095 | hg19\|chr1:115258686-115258808:-<br>TTTCCAACAGGTTCTTGCTGGTGTGAAATGACTGAGTACAAAC<br>TGGTGGTGGTTGGAGCAG[G/A]TGGTGTTGGGAAAAGCGCA<br>CTGACAATCCAGCTAATCCAGAACCACTTTGTAGATGAATAT | *NRAS*:exon2:c.G35A:p.G12D |
| dHsaMDS721630771 | hg19\|chr1:115258686-115258808:-<br>TTTCCAACAGGTTCTTGCTGGTGTGAAATGACTGAGTACAAAC<br>TGGTGGTGGTTGGAGCAG[G/C]TGGTGTTGGGAAAAGCGCA<br>CTGACAATCCAGCTAATCCAGAACCACTTTGTAGATGAATAT | *NRAS*:exon2:c.G35C:p.G12A |
| dHsaMDV2510528 | hg19\|chr1:115258686-115258808:-<br>TTTCCAACAGGTTCTTGCTGGTGTGAAATGACTGAGTACAAAC<br>TGGTGGTGGTTGGAGCAG[G/T]TGGTGTTGGGAAAAGCGCAC<br>TGACAATCCAGCTAATCCAGAACCACTTTGTAGATGAATAT | *NRAS*:exon2:c.G35T:p.G12V |
| dHsaMDV2510534 | hg19\|chr1:115258684-115258806:-<br>TCCAACAGGTTCTTGCTGGTGTGAAATGACTGAGTACAAACTG<br>GTGGTGGTTGGAGCAGGT[G/C]GTGTTGGGAAAAGCGCACT<br>GACAATCCAGCTAATCCAGAACCACTTTGTAGATGAATATGA | *NRAS*:exon2:c.G37C:p.G13R |
| dHsaMDV2510526 | hg19\|chr1:115258683-115258805:-<br>CCAACAGGTTCTTGCTGGTGTGAAATGACTGAGTACAAACTG<br>GTGGTGGTTGGAGCAGGTG[G/A]TGTTGGGAAAAGCGCACT<br>GACAATCCAGCTAATCCAGAACCACTTTGTAGATGAATATGAT | *NRAS*:exon2:c.G38A:p.G13D |

| | | |
|---|---|---|
| dHsaMDV2510524 | hg19\|chr1:115258683-115258805:- CCAACAGGTTCTTGCTGGTGTGAAATGACTGAGTACAAACTG GTGGTGGTTGGAGCAGGTG[G/T]TGTTGGGAAAAGCGCACTG ACAATCCAGCTAATCCAGAACCACTTTGTAGATGAATATGAT | *NRAS*:exon2:c.G38T:p.G13V |
| dHsaMDS381711539 | hg19\|chr12:112888105-112888227:+ CCAATGGACTATTTTAGAAGAAATGGAGCTGTCACCCACATCA AGATTCAGAACACTGGTG[A/C]TTACTATGACCTGTATGGAGG GGAGAAATTTGCCACTTTGGCTGAGTTGGTCCAGTATTAC | *PTPN11*:exon3:c.A182C:p.D61A |
| dHsaMDS585736928 | hg19\|chr12:112888078-112888200:+ TCTTTATTTGTCCCCTTGCCTCCCTTTCCAATGGACTATTTTAGA AGAAATGGAGCTGTCA[C/A]CCACATCAAGATTCAGAACACTG GTGATTACTATGACCTGTATGGAGGGGAGAAATTTGCC | *PTPN11*:exon3:c.C155A:p.T52N |
| dHsaMDS797586326 | hg19\|chr21:36252879-36253001:- CTAGGGGATGTTCCAGATGGCACTCTGGTCACTGTGATGGCT GGCAATGATGAAAACTACT[C/T]GGCTGAGCTGAGAAATGCT ACCGCAGCCATGAAGAACCAGGTTGCAAGATTTAATGACCTC | *RUNX1*:exon5:c.C422T:p.S141L |
| dHsaMDS704617272 | hg19\|chr21:36171553-36171675:+ TCCACCCCAGCTCAGCTGCAAAGAATGTGTTTTCAAGTGGCTT ACTTGAGAGTCGACTGGA[A/AAGG]AGTTCTGCAGAGAGGGT TGTCATGCCGCTGGCACGTCCAGGTGAAATGGGCGTTGCTGG GT | *RUNX1*:exon8:c.950_951insCCT: p.L317delinsLL |
| dHsaMDS837236043 | hg19\|chr21:36171552-36171674:+ TTCCACCCCAGCTCAGCTGCAAAGAATGTGTTTTCAAGTGGCT TACTTGAGAGTCGACTGG[A/ACC]AAGTTCTGCAGAGAGGGT TGTCATGCCGCTGGCACGTCCAGGTGAAATGGGCGTTGCTGG G | *RUNX1*:exon8:c.951_952insGG:p.S318fs |
| dHsaMDS911558603 | hg19\|chr17:7578355-7578477:+ AGCCCCAGCTGCTCACCATCGCTATCTGAGCAGCGCTCATGGT GGGGGCAGCGCCTCACAA[C/-]CTCCGTCATGTGCTGTGACTG CTTGTAGATGGCCATGGCGCGGACGCGGGTGCCGGGCGGG | *TP53*:exon5:c.514delG:p.V172fs |
| dHsaMDS234249306 | hg19\|chr17:7578229-7578351:+ TCCAAATACTCCACACGCAAATTTCCTTCCACTCGGATAAGATG CTGAGGAGGGGCCAGAC[C/T]TAAGAGCAATCAGTGAGGAAT CAGAGGCCTGGGGACCCTGGGCAACCAGCCCTGTCGTCTC | *TP53*:splicing (chr17: 7,578,290 C>T) |
| dHsaMDS590501846 | hg19\|chr11:32417846-32417968:+ ATATCTCTTATTGCAGCCTGGGTAAGCACACATGAAGGGGCG TTTCTCACTGGTCTCAGAT[G/GCCGA]CCGACCGTACAAGAGT CGGGGCTACTCCAGGCACACGTCGCACATCCTGCAGGCAGAG AGT | *WT1*:exon7:c.1144_1145insTCGG: p.A382fs |
